# Zebrafish larvae as a model system for systematic characterization of drugs and genes in dyslipidemia and atherosclerosis

**DOI:** 10.1101/502674

**Authors:** Manoj K Bandaru, Anastasia Emmanouilidou, Petter Ranefall, Benedikt von der Heyde, Eugenia Mazzaferro, Tiffany Klingström, Mauro Masiero, Olga Dethlefsen, Johan Ledin, Anders Larsson, Hannah L Brooke, Carolina Wählby, Erik Ingelsson, Marcel den Hoed

**Affiliations:** The Beijer Laboratory and Department of Immunology, Genetics and Pathology, Uppsala University, Uppsala, Sweden; Science for Life Laboratory, Uppsala University, Uppsala, Sweden; Science for Life Laboratory - BioImage Informatics Facility, Uppsala, Sweden; Department of Information Technology, Division of Visual Information and Interaction, Uppsala University, Uppsala, Sweden; Science for Life Laboratory - Genome Engineering Zebrafish National Facility, Uppsala University, Uppsala, Sweden; Science for Life Laboratory - National Bioinformatics Infrastructure Sweden, Stockholm University, Stockholm, Sweden; Department of Organismal Biology, Evolutionary and Developmental Biology, Uppsala University, Uppsala, Sweden; Department of Medical Sciences, Biochemical structure and function, Uppsala University, Uppsala, Sweden; Department of Public Health and Caring Sciences, Uppsala University, Uppsala, Sweden; Department of Medicine, Division of Cardiovascular Medicine, Stanford University School of Medicine, Stanford, CA 94305, USA; Department of Medical Sciences, Molecular Epidemiology, Uppsala University, Uppsala, Sweden

**Author notes:** Address for correspondence: Marcel den Hoed, The Beijer Laboratory, Department of Immunology, Genetics and Pathology and SciLifeLab, BMC, Husargatan 3, Box 815, 75 108 Uppsala, Sweden.

## Abstract

**Background:** Hundreds of loci have been robustly associated with circulating lipids, atherosclerosis and coronary artery disease; but for most loci the causal genes and mechanisms remain uncharacterized.

**Methods:** We developed a semi-automated experimental pipeline for systematic, quantitative, large-scale characterization of mechanisms, drugs and genes associated with dyslipidemia and atherosclerosis in a zebrafish model system. We validated our pipeline using a dietary (n>2000), drug treatment (n>1000), and genetic intervention (n=384), and used it to characterize three candidate genes in a GWAS-identified pleiotropic locus on chr 19p13.11 (n>500).

**Results:** Our results show that five days of overfeeding and cholesterol supplementation had independent pro-atherogenic effects, which could be diminished by concomitant treatment with atorvastatin and ezetimibe. CRISPR-Cas9-induced mutations in orthologues of proof-of-concept genes resulted in higher LDL cholesterol levels (*apoea*), and more early stage atherosclerosis (*apobb.1*). Finally, our pipeline helped identify putative causal genes for circulating lipids and early-stage atherosclerosis (*LPAR2* and *GATAD2A*).

**Conclusions:** In summary, our pipeline facilitates systematic, *in vivo* characterization of drugs and candidate genes to increase our understanding of disease etiology, and can likely help identify novel targets for therapeutic intervention.

## Introduction

Coronary artery disease (CAD) is the main cause of death worldwide, and results from the progression of atherosclerosis in the coronary arteries^1^. Since 2006, increasingly large genome-wide association studies (GWAS) have identified hundreds of genetic loci that are robustly associated with circulating lipid levels and CAD susceptibility^2–4^. Some of these loci harbor genes with well-known roles in cholesterol metabolism – e.g. *APOE*^5^, *APOB*^6^ and *LDLR*^7^ – and some encode the targets of lipid-lowering drugs, i.e. *HMGCR* (statins)^8^, *NPC1L1* (ezetimibe)^9^ and *PCSK9* (evolocumab)^10,11^. It thus seems plausible that identifying and characterizing causal genes in the remaining loci would further increase our understanding of cholesterol metabolism, atherosclerosis and CAD pathophysiology, and yield new targets for prevention and treatment of CAD^11,12^.

Murine model systems have traditionally been used to characterize genes that play a role in familial hypercholesterolemia and atherosclerosis^13,14^. However, such screens are too time consuming and costly to facilitate systematic screens across hundreds of candidate genes. In addition, mice differ in cholesterol ester and triglyceride transport between lipoprotein particles due to lack of *CETP*^15^, and only develop atherosclerosis in an *LDLR* or *APOE* knockout background. Alternative, *in vivo* model systems that are suitable for high-throughput characterization of disease-related traits are thus desirable. In this context, zebrafish (*Danio rerio*) provide a promising opportunity^16,17^.

Proof-of-principle experiments on using zebrafish larvae as a model system for early-stage atherosclerosis have so far provided promising results^18–23^, but were based on observations in fewer than 25 larvae per condition. This was at least in part because mounting larvae in low melting agarose for imaging is time-consuming. In addition, whole-body cholesterol and triglyceride levels were usually quantified in samples of 20-100 pooled larvae^24,25^. While suitable and efficient for dietary and drug treatment interventions, pooling larvae for phenotypic characterization is not optimal in CRISPR-based genetic interventions, where – depending on the approach - sequencing of individual larvae may be required. Hence, confirmation of initial findings, an improved resolution of quantitative readouts, and a higher throughput are desirable before zebrafish larvae can be used as a model system for large-scale characterization of candidate genes for dyslipidemia, atherosclerosis and CAD. To this end, we performed a large-scale dietary intervention, a treatment intervention with lipid lowering drugs, and a multiplexed, CRISPR-Cas9-based genetic intervention for proof-of-concept genes, followed by the characterization of three candidate genes in a GWAS-identified, pleiotropic locus on chr 19p13.11^26^.

## Results

### Overfeeding and cholesterol supplementation have independent pro-atherogenic effects

To quantify and distinguish between the atherogenic potential of overfeeding and dietary cholesterol, >2000 larvae from three transgenic backgrounds (Fig. 1, Supplementary Tables 1 and 2) were fed on one of six diets, from the age of 5 days post-fertilization (dpf) until 9 dpf (**Methods**).

**Figure 1.**
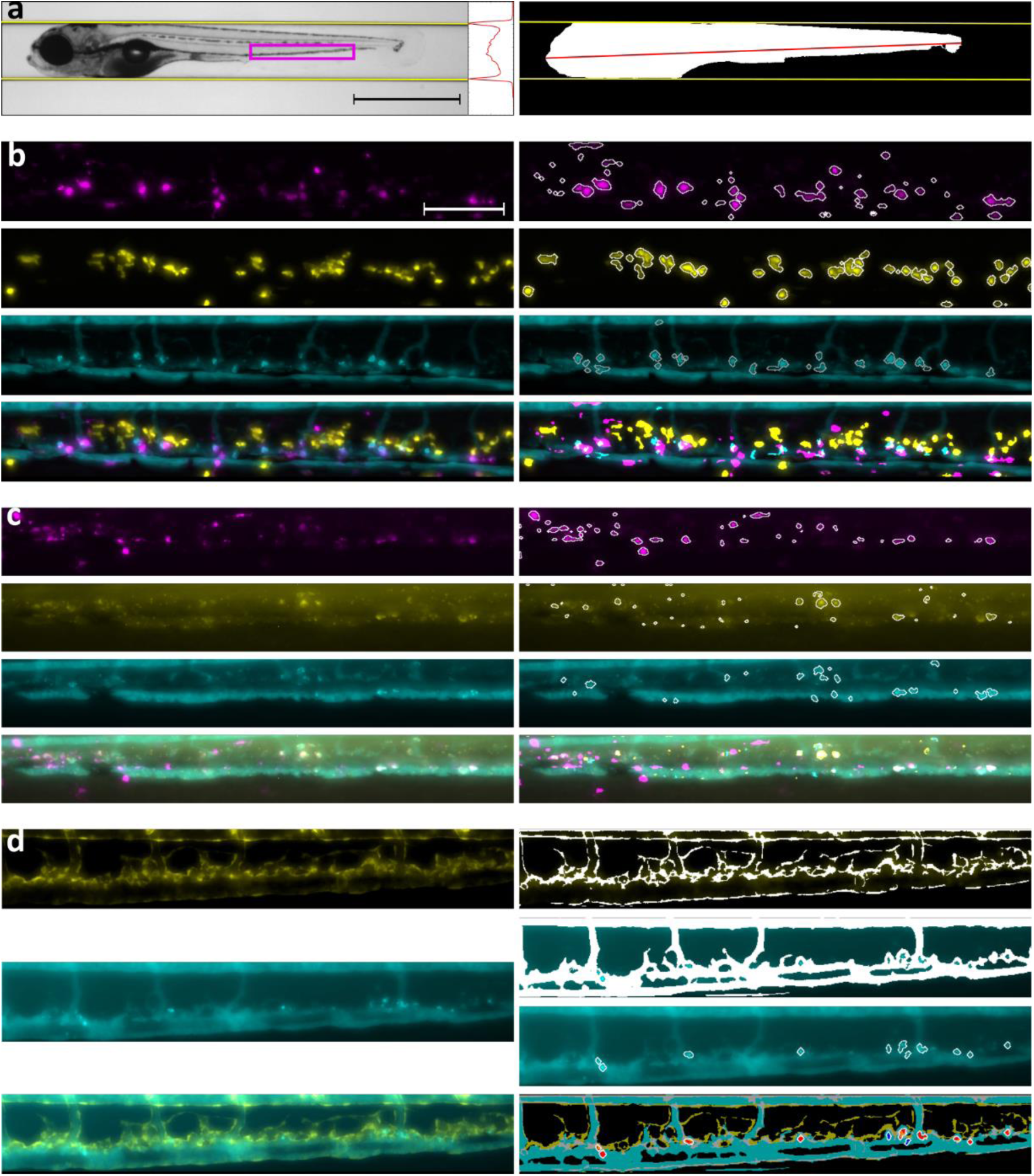
Raw data (left) and objective, semi-automated quantification (right) of body size and early-stage atherosclerosis in 10-day-old zebrafish larvae. a) Left: A bright field image of a zebrafish larva in lateral orientation with projection of all intensity values to the y-axis. The two distinct minima in the projection represent the walls of the capillary, outlined in yellow (scale bar = 1 mm). The region of the tail that was imaged to quantify vascular atherogenic traits is highlighted in magenta. Right: a binary mask of the same larva, with lateral surface area in white, and body length in red. b) A Tg(mpeg1-mCherry; mpo-EGFP) transgenic larva with fluorescently labelled macrophages (top, magenta) and neutrophils (2^nd^ from top, yellow). Circulating lipids and vascular lipid deposits were stained with a dye (3^rd^ from the top, cyan). The overlay (bottom) shows co-localization of all traits (scale bar = 100µm). c) A Tg(mpeg1-mCherry; hsp70:IK17-EGFP) transgenic larva with fluorescently labelled macrophages (top, magenta) and oxidized LDL (2^nd^ from top, yellow) with stained lipids (3^rd^ from top, cyan). The overlay shows co-localization of all traits (bottom). d) A Tg(flk-EGFP) transgenic larva with fluorescently labelled endothelial cells showing endothelial surface area (top, yellow); stained lipids (2^nd^ from top, cyan) from which both circulating lipids (right, 2^nd^ from top) and vascular lipid deposition (right, 3^rd^ from top) were quantified; and an overlay that enabled distinguishing between lipid deposition inside (in red) and outside the endothelium (bottom right, blue).

Five days of overfeeding on average resulted in longer larvae, with a larger body surface area and volume normalized for length (Fig. 2a-i, Supplementary Fig. 1, Supplementary Table 3). Overfeeding induced a triglyceride-driven increase in total cholesterol levels, without materially affecting LDLc, HDLc or glucose levels (Fig. 2a-iii, Supplementary Fig. 2, Supplementary Table 4). Overfeeding also resulted in more lipid deposition (Fig. 2a-ii, Supplementary Fig. 3, Supplementary Table 5). We further ensured that the observed lipid deposition was indeed located inside the vascular endothelium using larvae with fluorescently labelled endothelial cells (*Tg:flk-EGFP*) (Fig. 1d, Supplementary Table 1). Of the 361 *Tg:flk-EGFP* positive larvae that showed at least some vascular lipid deposition, 355 (98.3%) had all lipid deposits co-localize with circulating lipids and/or vascular endothelial cells, implying that almost all deposits were at least partly located inside the endothelial cell layer. Deposits in the remaining six larvae were false positives, illustrating the high sensitivity (72%) and specificity (93%) of our image quantification pipeline for detection of vascular lipid deposition. Overfeeding also resulted in more vascular accumulation of oxLDL; more co-localization of oxLDL with macrophages; and more vascular co-localization of lipids with neutrophils. We also observed some evidence for a positive effect of overfeeding on vascular infiltration by neutrophils and on endothelial thickness (Fig. 2a-ii, Supplementary Fig. 3, Supplementary Table 5).

**Figure 2.**
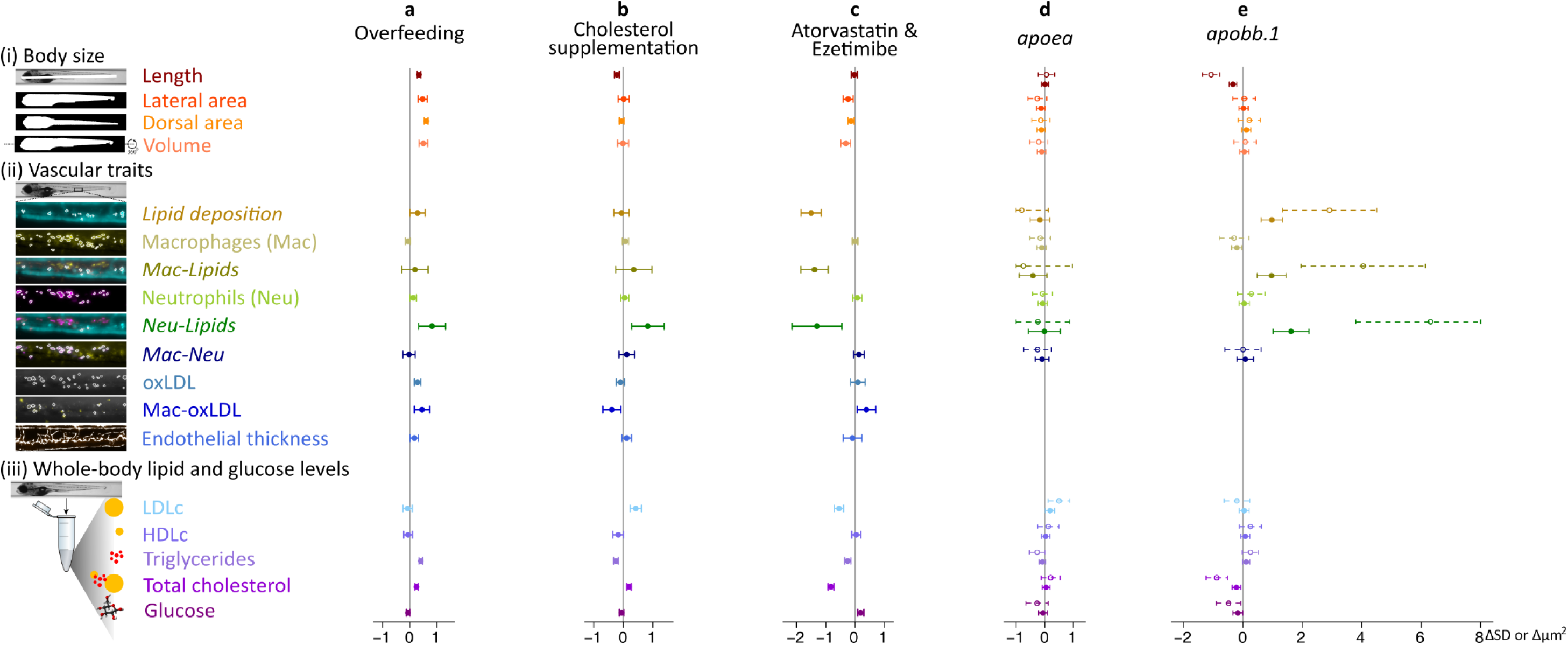
The effect of overfeeding and cholesterol supplementation (n>2000); treatment with atorvastatin and ezetimibe (n>1000); and mutations in apoea and apobb.1 (n=384) on body size (i), vascular atherogenic traits (ii) and whole-body lipid and glucose levels (iii). Across a-e, dorsal and lateral body surface area and body volume were normalized for body length before the analysis; whole-body lipid and glucose levels were normalized for protein levels; and endothelial thickness was normalized for surface area of the circulation. For normally distributed traits, associations were examined using hierarchical linear models on inverse-normally transformed outcomes. For these traits effect sizes and 95% confidence intervals are expressed in standard deviation units (SD). The remaining vascular atherogenic traits (shown in italics) showed a negative binomial distribution and data were analyzed accordingly. For these traits, effect sizes and 95% confidence intervals are expressed in µm^2^. In d and e, open circles and the dotted lines represent the effect of two functionally knocked-out alleles vs. two unmodified alleles, and full circles and filled lines represent the additive per mutated allele effect. Associations were adjusted for time of day; use of diethyl ether (for overfeeding and cholesterol supplementation); cholesterol supplementation (for overfeeding); the amount fed (for cholesterol supplementation); body length and dorsal body surface area (for vascular outcomes); batch; and transgenic background.

Five days of dietary cholesterol supplementation resulted in shorter larvae, without affecting body surface area or volume normalized for length (Fig. 2b-i, Supplementary Fig. 1, Supplementary Table 3), and without influencing food intake (Supplementary Fig. 4). Cholesterol supplementation induced an LDLc-driven increase in total cholesterol levels, while lowering HDLc (trend) and triglyceride levels (Fig. 2b-iii, Supplementary Fig. 2, Supplementary Table 4). Cholesterol supplementation did not influence vascular accumulation of lipids and oxLDL, but tended to result in more co-localization of lipids with neutrophils, and in less co-localization of oxLDL with macrophages (Fig. 2b-ii, Supplementary Fig. 3, Supplementary Table 5).

As described in earlier studies^18,19,25^, we supplemented regular dry food with extra cholesterol using diethyl ether, which may itself affect endogenous cholesterol levels^27^. Our results show that diethyl ether per se indeed resulted in: 1) higher triglyceride (trend) and total cholesterol levels (Supplementary Table 4); 2) less vascular co-localization of lipids with neutrophils; 3) a lower endothelial thickness (Supplementary Table 5); and 5) a lower food intake (Supplementary Fig. 4). Hence, absence of a control group fed on diethyl ether-treated food without cholesterol supplementation would have resulted in biased estimates for the effect of cholesterol supplementation.

Neither overfeeding, nor cholesterol supplementation or diethyl ether supplementation was associated with suboptimal image quality or image quantification. Hence, exclusion of larvae based on these criteria likely did not influence the results of the dietary intervention (Supplementary Table 6).

### Combined treatment with atorvastatin and ezetimibe has an atheroprotective effect

To examine whether the commonly prescribed LDLc lowering drugs atorvastatin and ezetimibe exert similar effect in zebrafish, we overfed >1,000 larvae on a cholesterol supplemented diet with or without concomitant atorvastatin and ezetimibe treatment, from 5 dpf until 9 dpf. Compared with untreated larvae, five days of combined treatment with atorvastatin and ezetimibe resulted in leaner larvae (Fig. 2c-i, Supplementary Fig. 1, Supplementary Table 7), without affecting food intake (Supplementary Fig. 4). On average, atorvastatin and ezetimibe treatment also resulted in lower whole-body LDLc, triglyceride and total cholesterol levels, and in higher glucose levels (Fig. 2c-iii, Supplementary Fig. 2, Supplementary Table 8). In larvae with data on LDLc (n=564), atorvastatin and ezetimibe’s effect on glucose levels was independent of triglyceride (beta: 0.13 SD; 95% CI: 0.04 to 0.22 SD), but not LDLc levels (0.05, −0.05 to 0.15 SD).

Treatment with atorvastatin and ezetimibe resulted in less vascular lipid deposition and less co-localization of lipids with macrophages and with neutrophils. On the other hand, treated larvae on average had more vascular co-localization of oxLDL with macrophages (Fig. 2c-ii, Supplementary Fig. 3, Supplementary Table 9).

Larvae treated with atorvastatin and ezetimibe were more likely to move during imaging (due to their leaner bodies) and had lower odds of many false positive oxLDL deposits. Exclusion of larvae with such suboptimal imaging or quantification data is unlikely to have influenced the results (Supplementary Table 10).

### Mutations in zebrafish orthologues of APOE and APOB exert pro-atherogenic effects

Next, orthologues of genes with an established role in dyslipidemia and atherosclerosis - *APOE*, *APOB* and *LDLR* - were targeted together using a multiplexed CRISPR-Cas9 approach. These three genes together have seven orthologues in zebrafish (*apoea*, *apoeb*, *apoba*, *apobb.1*, *apobb.2*, *ldlra* and *ldlrb*) (Supplementary Tables 11 and 12). Across the seven CRISPR-Cas9-targeted orthologues, we observed a median of 15 unique amplicons per targeted site in the 384 sequenced F1 larvae (Supplementary Table 13). Compared with the reference genome, the 384 sequenced F1 larvae together contained 55 frameshift variants, nine variants that introduced a premature stop codon, 34 missense variants, 13 in frame deletions, two in frame insertions, four synonymous variants, and 18 upstream variants within ±30 bp of the targeted sites (Supplementary Table 14). The mutant allele frequency was typically high across the seven targeted sites (i.e. median 0.883, Supplementary Table 15).

Most larvae carried two functionally knocked alleles in *apoba* and *apobb.2* – i.e. frame shift mutations and/or variants introducing a premature stop codon in both alleles – as well as bi-allelic mutations immediately upstream of *ldlrb* that were predicted to modify *ldlrb* gene expression (Supplementary Tables 14 and 15). Since there were no wildtype larvae for *apoba*, *apobb.2* and *ldlrb*, we could not examine the role of these three orthologues. For the four remaining orthologues (*apoea, apoeb*, *apobb.1* and *ldlra*), a genetic burden score comprising the sum of the number of mutated alleles across the four genes, weighted by their predicted effect on protein function was normally distributed. The score was associated with higher HDLc levels, more vascular lipid deposition, and more vascular co-localization of lipids with neutrophils (Supplementary Fig. 5, Supplementary Tables 16-18).

When examining the influence of mutations in each gene separately, we observed that larvae carrying two functionally knocked alleles in *apoea* had higher whole-body LDLc and lower triglyceride (trend) levels, without an effect on body size or early-stage atherosclerosis (Fig. 2d-ii, Supplementary Tables 20 and 21). Larvae with two functionally knocked *apoeb* alleles showed at most a trend for more vascular accumulation of lipids and co-localization of macrophages with neutrophils, without affecting whole-body lipoprotein or glucose levels (Supplementary Fig. 6b, Supplementary Tables 20 and 21). On average, larvae with two functionally knocked *apobb.1* alleles were shorter than larvae with two unmodified alleles (Fig. 2e-i, Supplementary Table 19), and had higher triglyceride (trend) and lower total cholesterol and glucose levels (Fig. 2e-iii, Supplementary Table 20). They were also characterized by more vascular accumulation of lipids, and by more vascular co-localization of lipids with macrophages and with neutrophils, independently of whole-body lipoprotein or glucose levels (Fig. 2e-ii, Supplementary Table 21). While less than half of the larvae free from mutated *apobb.1* alleles showed any vascular co-localization of lipids with macrophages (44%) and neutrophils (15%), more than two out of three larvae carrying two functionally knocked *apobb.1* alleles showed such vascular co-localization. Finally, larvae with two functionally knocked *ldlra* alleles were similar on all accounts to larvae free from CRISPR-induced mutations in the gene, except for having less vascular co-localization of macrophages with neutrophils (Supplementary Fig. 6d, Supplementary Tables 19-21).

Across the four zebrafish orthologues (*apoea*, *apoeb*, *apobb.1* and *ldlra*), results were similar when data were analyzed using an additive model in which the number of mutated alleles was weighted by their predicted effect on protein function (Supplementary Tables 22-24). Additional analyses showed that suboptimal image quality or image quantification in a small subset of larvae is unlikely to have influenced the results (Supplementary Table 25). Since *APOE*, *APOB* and *LDLR* interact to process triglyceride-rich LDLc in humans, we next examined two-way gene x gene interactions for *apoea*, *apobb.1* and *ldlra*; i.e. the genes that showed the most promising results individually. Focusing on interactions that were observed under an additive model and that were confirmed when comparing larvae with two vs. zero functionally knocked alleles only shows cautious evidence of a positive interaction between mutations in *apoea* and *ldlra* for vascular accumulation of lipids and co-localization of lipids with macrophages (Supplementary Tables 26-28).

### Vascular atherogenic traits are associated with whole-body triglyceride levels

Data from the dietary, drug treatment and genetic interventions combined showed that LDLc, HDLc and triglyceride levels together explained 47% of the variance in directly assessed total cholesterol levels (n=1,867). Interestingly, the Friedewald equation (i.e. LDLc + HDLc + triglycerides/5)^28^ did not perform much worse, explaining 43% of the variance in total cholesterol levels (Supplementary Fig. 7). We next explored the mutually adjusted association of vascular atherogenic traits with whole-body LDLc, HDLc, triglyceride and glucose levels in data from the dietary, drug treatment and proof of concept genetic intervention studies combined. Vascular accumulation of lipids, co-localization of lipids with macrophages and neutrophils, and co-localization of oxLDL with macrophages were all positively associated with whole-body triglyceride levels, independently of LDLc, HDLc and glucose levels. Furthermore, vascular accumulation of oxLDL and co-localization of lipids with neutrophils showed some evidence of a positive association with whole-body HDLc levels, in line with the absence of effects on primary clinical endpoint events observed in large clinical trials that therapeutically elevated HDLc levels and reduced triglyceride and/or LDLc levels^29–32^. Interestingly, vascular co-localization of lipids with neutrophils showed independent positive associations with LDLc, HDLc and triglyceride levels. Finally, we observed negative associations of whole-body glucose levels with vascular accumulation of lipids, co-localization of lipids with macrophages, and co-localization of macrophages with oxLDL, suggesting that hyperglycemia per se is perhaps not responsible for the elevated risk of CAD in diabetes patients, at least not by increasing early stage atherosclerosis (Supplementary Fig. 8, Supplementary Table 29).

### Identifying putative causal genes for circulating lipids and early-stage atherosclerosis

Based on the positive association of vascular atherogenic traits with triglyceride levels in our combined analysis, in combination with the known causal effect of high triglyceride levels on CAD incidence^33^, we used DEPICT^34^ to prioritize candidate genes in 23 triglyceride-associated loci^26^. In one of these loci, represented by the intronic rs10401969 in *SUGP1* on chr 19p13.11, DEPICT prioritized *LPAR2, GMIP, GATAD2A* and *TM6SF2*. The four prioritized genes together have six orthologues in zebrafish (Supplementary Table 30), which we targeted simultaneously (Supplementary Table 31).

Across the six CRISPR-Cas9-targeted orthologues (*lpar2a*, *lpar2b*, *gmip*, *gatad2ab*, *tm6sf2* and *zgc:85843*), we observed a median of 2.5 unique amplicons per targeted site (Supplementary Table 32). Compared with the reference genome, the 547 sequenced F1 larvae together contained four frameshift variants, three missense variants, and four in-frame deletions that were located within ±30 bp of the CRISPR targeted sites (Supplementary Table 33). In spite of having pre-tested the CRISPR gRNAs for efficiency, all F1 larvae carried two unmodified alleles for the zebrafish orthologues of *TM6SF2* (*tm6sf2* and *zgc:85843*) and *GMIP* (*gmip*), and mutant allele frequencies were low for *gatad2ab* (0.029), *lpar2a* (0.010), and *lpar2b* (0.005) (Supplementary Table 34).

In spite of the low statistical power to find associations, we observed evidence for lower LDLc, triglyceride and total cholesterol levels in the 11 larvae with a mutated *lpar2a* allele, when compared with larvae with two unmodified alleles. Counterintuitively, these larvae also showed some evidence for having more vascular co-localization of lipids with macrophages and with neutrophils. (Fig. 3a, Supplementary Tables 35-37). In addition to the effects observed for *lpar2a*, the six larvae with a mutated *lpar2b* allele were longer and tended to have lower HDLc levels compared with larvae free from CRISPR-induced *lpar2b* mutations (Fig. 3b, Supplementary Tables 35-37). Finally, the 32 larvae with a mutated *gatad2ab* allele were larger and tended to have lower HDL and higher triglyceride levels when compared with larvae with two unmodified alleles (Fig. 3c, Supplementary Table 36). Exclusion of larvae with suboptimal image quality or image quantification is unlikely to have influenced the results (Supplementary Table 38).

**Figure 3.**
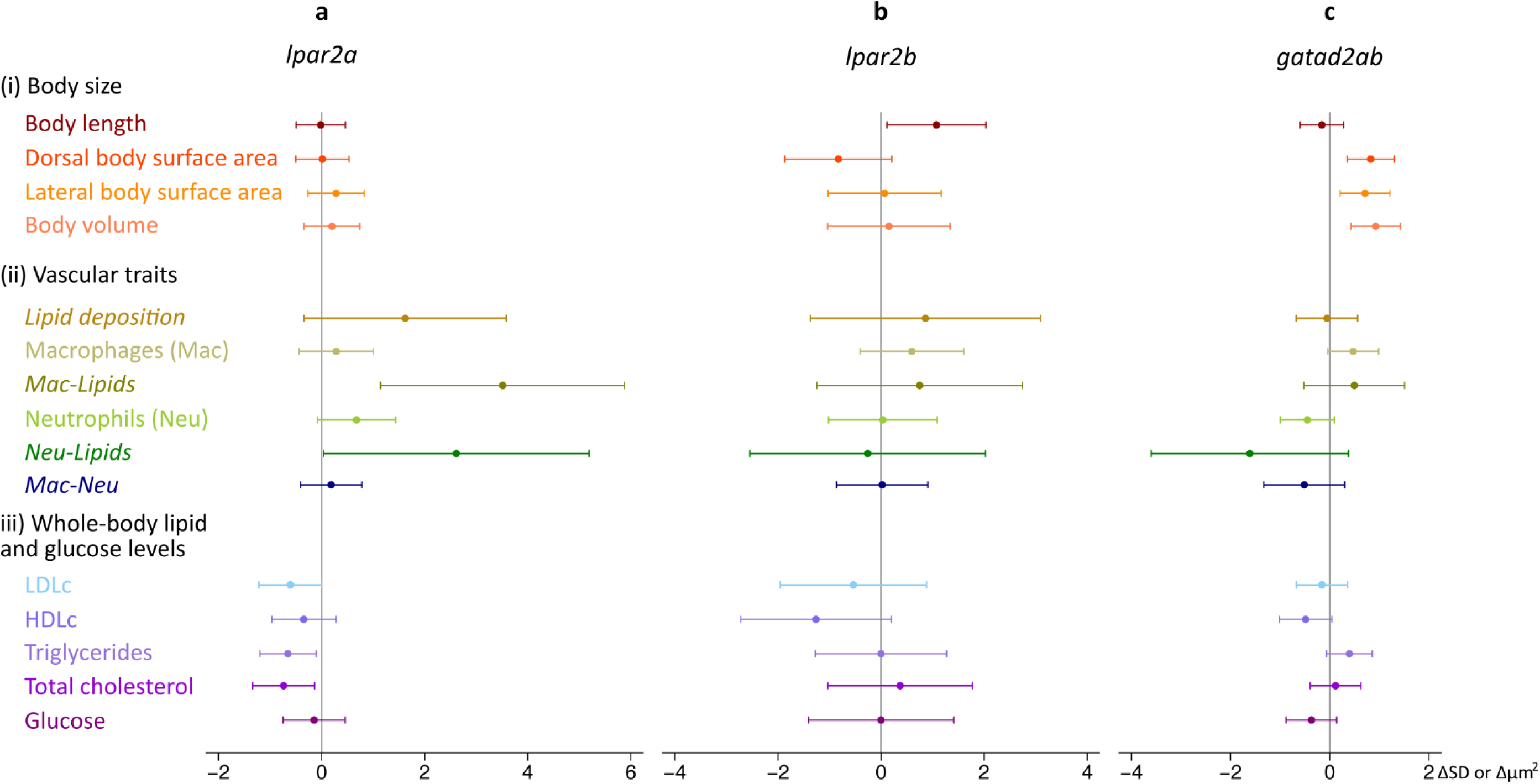
The mutually adjusted effect of mutations in zebrafish orthologues of LPAR2 and GATAD2A (n=547) on body size (i), vascular atherogenic traits (ii) and whole-body lipid and glucose levels (iii) using an additive model. Dorsal and lateral body surface area and body volume were normalized for body length; and whole-body lipid and glucose levels were normalized for protein levels before the analysis. For normally distributed traits, associations were examined using hierarchical linear models on inverse-normally transformed outcomes. For these traits, effect sizes and 95% confidence intervals are expressed in standard deviation units (SD). Some vascular atherogenic traits showed a negative binomial distribution and associations were analyzed accordingly. For these traits (shown in italics), effect sizes and 95% confidence intervals are expressed in µm^2^. Associations were adjusted for time of day; body length and dorsal body surface area (for vascular outcomes); and batch.

## Discussion

We developed and validated a largely image-based experimental pipeline in zebrafish larvae that is suitable to systematically characterize candidate genes and drugs for a role in dyslipidemia and early-stage atherosclerosis and inflammation. Our dietary intervention showed that five days of overfeeding is sufficient to induce early-stage atherosclerosis and vascular inflammation in zebrafish larvae, without the need to use an *APOE* or *LDLR* knockout background as is customary in mouse models. Our drug treatment intervention showed that the pro-atherogenic effects of overfeeding and cholesterol supplementation can be diminished by concomitant treatment with atorvastatin and ezetimibe. A proof-of-concept genetic screen showed that CRISPR-Cas9-induced mutations in zebrafish orthologues of *APOE* and *APOB* trigger a pro-dyslipidemia, pro-atherogenic and pro-inflammatory phenotype that is in line with the known role of these genes. Finally, we illustrated the merit of our pipeline by attributing a role in cholesterol metabolism and atherosclerosis to *LPAR2* and *GATAD2A*; two genes in a pleiotropic locus on chr 19p13.11^26^.

In an adequately powered dietary intervention, we showed that overfeeding and cholesterol supplementation have independent pro-inflammatory and pro-atherogenic effects in zebrafish larvae. Both induced higher whole-body total cholesterol levels, albeit via different mechanisms. While overfeeding resulted in higher triglyceride levels, cholesterol supplementation induced higher LDLc levels. At the vascular level, overfeeding resulted in an overall more pro-atherogenic profile; cholesterol supplementation only resulted in more vascular co-localization of lipids with neutrophils. These results suggest that primary accumulation of lipids in the vessel wall was likely mostly driven by elevated triglyceride levels. In line with this, data from the dietary, drug treatment and genetic interventions combined showed a positive association for triglyceride levels – but not LDLc – with vascular lipid deposition. Furthermore, mutations in *apobb.1* resulted in higher whole-body triglyceride levels as well as in more vascular accumulation of lipids – albeit independently of triglyceride levels – while mutations in *apoea* resulted in higher whole-body LDLc levels but did not influence vascular lipid deposition.

Treating larvae with atorvastatin and ezetimibe resulted in lower whole-body LDLc and total cholesterol levels; and in less vascular co-localization of lipids with macrophages; yet – paradoxically – in more vascular co-localization of oxLDL with macrophages. A directionally consistent (i.e. lowering) effect was observed for dietary cholesterol supplementation on vascular co-localization of oxLDL with macrophages. Moreover, cholesterol supplementation or drug treatment did not affect accumulation of oxLDL or macrophages per se. Taken together, these observations suggest that ezetimibe’s exogenous cholesterol lowering effect may be responsible for improved: 1) recruitment of macrophages to oxLDL; 2) engulfing of oxLDL by macrophages; 3) survival of macrophages that successfully engulfed oxLDL; and/or 4) clearing of neutral lipid deposits by macrophages. The opposite rationale applies to elevated exogenous cholesterol levels following dietary cholesterol supplementation.

In line with results from clinical trials^35,36^ and genetic association studies^37,38^ in humans, zebrafish larvae treated with atorvastatin and ezetimibe were characterized by higher glucose levels, on average. Additional analyses indicated that the drugs’ effect on glucose levels is likely mediated by LDLc. Hence, main effects and mediation analyses based on whole-body cholesterol and glucose levels in zebrafish larvae should be sufficiently sensitive to provide valuable new insights.

Of the three *APOB* orthologues in zebrafish (*apoba*, *apobb.1* and *apobb.2*), *apobb.1,* accounts for 95% of apob protein^21^. Apobb.1 is mainly expressed in intestine and liver^21^, where it facilitates the assembly and secretion of triglyceride-rich chylomicrons and VLDL, respectively^39^. Unlike mice^40^, but similar to humans^41^, zebrafish larvae with two loss of function mutations in *apobb.1* alongside similar mutations in *apoba* and *apobb.2* were viable, at least until 10dpf. Viability during early development despite absence of functional apob protein suggests that Apo B is more essential for processing fetal nutrition in mice than in humans and zebrafish. However, homozygous mutations described in humans^42^ and observed here in zebrafish result in a truncated protein, while those described in mice inhibit protein synthesis completely^40^. Hence, alternatively, complete absence of Apo B protein may induce a more severe phenotype than presence of a truncated protein. The CRISPR-induced mutations in *apobb.1* may result in a truncated apobb.1 protein that cannot act as a ligand for lipoprotein lipase; the enzyme that catalyzes the conversion of triglyceride-rich-chylomicrons and VLDL to cholesterol-rich-IDL and LDL. This would result in higher triglyceride and lower total cholesterol levels, as we observed in this study. In line with results in humans^43^, we also observed a more severe pro-atherogenic and pro-inflammatory profile in *apobb.1* mutant zebrafish larvae. These findings together suggest that *apobb.1*^-/-^ zebrafish are likely a promising model to examine candidate genes and drugs for a role in dyslipidemia and atherosclerosis. The lower glucose levels in *apobb.1* mutants are in line with results in humans^44,45^, and may result from impaired insulin clearance in the presence of hypobetalipoproteinemia^46^.

Mutations in *ldlra* were not associated with dyslipidemia, early-stage atherosclerosis or vascular inflammation in our study. This contrast with established results in humans and mouse models likely reflects the presence of a second – albeit downregulated - *LDLR* orthologue in zebrafish (*ldlrb*); the possibility of *cetp*-mediated reverse cholesterol transport to remove excess cholesterol from the body in zebrafish; or the early stage of development at which we performed our screen. Two studies previously implicated *ldlra* in dyslipidemia and early-stage atherosclerosis in zebrafish larvae^25,47^. While differences in age, food intake, microbial environment, enzymatic assays, normalization for protein content^48^, genetic manipulation^49^ and adjustment for co-variables across studies may have influenced the results, the difference in sample size between studies is most noteworthy. O’Hare et al. compared combined LDLc and VLDLc levels using repeated measures on a sample of 100 pooled morpholino-injected and a sample of 100 pooled control-injected larvae^25^, while Liu et al. compared triglyceride and total cholesterol levels in four wildtype larvae and four larvae that were homozygous for mutations in *ldlra*^47^. In contrast, we compared cholesterol levels in 181 and 120 individual larvae with two and zero functionally knocked alleles, respectively, and included data from 381 larvae in our additive analyses. While mutations in *ldlra* and *apoea* alone did not trigger early-stage atherosclerosis, mutations in these genes showed a positive interaction for vascular accumulation of lipids and co-localization of lipids with macrophages. It appears that absence of both *ldlra* and *apoea* cannot be compensated in zebrafish larvae.

Like humans, zebrafish are genetically heterogeneous, and we observed a normal or negative binomial distribution for the examined outcomes, with substantial variance by transgenic background and batch in the dietary, drug treatment and genetic interventions. These findings stress the importance of including data from a large number of larvae to acquire meaningful results. We show it is now feasible to objectively quantify relevant traits in a large number of larvae with relative ease. Hence, adequately powered, *in vivo*, systematic gene and drug screens are now possible for these complex outcomes.

Characterizing zebrafish orthologues of candidate genes in a pleiotropic locus on chr 19p13.11 highlighted a role for *LPAR2* in cholesterol metabolism and early-stage atherosclerosis, and for *GATAD2A* in cholesterol metabolism. *LPAR2* belongs to family I of the G-protein coupled receptors and functions to mobilize calcium in response to lysophosphatidic acid (LPA), while *GATAD2A* encodes a transcriptional repressor. Unfortunately, all larvae in our screen were wildtype for the two CRISPR-targeted orthologues of *TM6SF2*. Knockdown and knockout of *Tm6sf2* in mice was previously shown to result in lower circulating triglyceride, LDLc, HDLc and total cholesterol levels; as well as in higher hepatic triglyceride and cholesteryl esters; more and larger neutral lipid droplets in the liver; a higher risk of hepatic steatosis; and less atherosclerosis^50–52^. Like *Tm6sf2* deficient mice, *lpar2a* mutant zebrafish larvae had lower triglyceride, LDLc and total cholesterol levels. However, in contrast with *Tm6sf2* mutant mice, *lpar2a* mutant zebrafish larvae had more early-stage atherosclerosis, possibly driven by higher lysophosphatidic acid levels. Lysophosphatidic acid has been shown to increase NFκB, IL-8 and MCP-1 secretion from endothelial cells, which attract neutrophils and macrophages^53,54^; and induces barrier dysfunction and elevated monocyte adhesion to the minimally modified LDL within the vascular intima^55^. The bi-directional effects of mutations in putative causal genes in this pleiotropic locus on cardiovascular risk factors likely explains why the C allele of the regulatory rs10401969 is only associated with a trend towards a lower risk of CAD in humans (OR=0.95, P=2.8•10^-3^, n=268,744)^2^.

In conclusion, zebrafish larvae can be used as a time and cost-efficient model system for systematic image- and CRISPR-Cas9-based genetic interventions, as illustrated by the identification of putative causal genes for cholesterol metabolism (*LPAR2* and *GATAD2A*) and for early-stage atherosclerosis and inflammation (*LPAR2*). Our approach represents an opportunity to reduce the hundreds of candidate genes in GWAS-identified loci to a more feasible number for: 1) further in-depth characterization using animal models; 2) more targeted whole-genome or whole-exome sequencing and genotype-based recall efforts; 3) target-driven drug discovery.

## Online Methods

### 1 Transgenic backgrounds and atherogenic traits

We used three combinations of fluorescent transgenes (backgrounds) with a lipid-staining dye^56^ (see below) to visualize and quantify (see ‘*Image quantification’*) molecular processes that are known to play a role in early-stage atherosclerosis (**Table 1**). Firstly, zebrafish carrying transgenes to fluorescently label macrophages (*Tg:mpeg1-mCherry*^57^) and neutrophils (*Tg:mpo-EGFP*^58^) were crossed to yield a stable line in which we can visualize and quantify vascular accumulation and co-localization of lipids^56^, macrophages^57^ and neutrophils^58^ (Fig. 1b). Secondly, we in-crossed zebrafish that express a fluorescently labelled antibody (IK17) against oxLDL (*Tg:hsp70:IK17-EGFP*)^20^ to allow visualization and quantification of vascular accumulation of lipids^56^ and oxLDL^20^, and we crossed *Tg:hsp70:IK17-EGFP*^20^ carriers with *Tg:mpeg1-mCherry* carriers to yield a stable line in which we can visualize and quantify vascular accumulation and co-localization of lipids^56^, oxLDL^20^ and macrophages^57^ (Fig. 1b-c). Thirdly, carriers of the flk:EGFP transgene (*Tg:flk-EGFP*^59^) allowed us to quantify vascular accumulation of lipids, confirm or refute whether vascular lipid deposits are located inside the endothelial cell layer, and quantify the endothelial thickness (Fig. 1d). In all backgrounds, circulating lipids and vascular lipid deposits were visualized using a dye that preferentially partitions in lipid droplets and that has a blue-shifted, highly enhanced emission in lipophilic environments (monodansylpentane cadaverase [MDH], Abgent, Nordic Biosite, Täby, Sweden)^56^.

After imaging, we used enzymatic assays to assess whole-body LDLc, HDLc, triglyceride, total cholesterol and glucose levels. DNA was isolated from the remaining tissue for paired-end sequencing of CRISPR-Cas9 targeted sites in the genetic interventions (see ‘*genetic intervention*’).

### 2 Husbandry

All experiments described below were performed in zebrafish larvae. Adult transgenic fish and CRISPR founders were raised and kept solely for breeding purposes. Adult fish were fed twice daily on rotifers and dry food (Sparos, Olhão, Portugal), and were maintained in circulating and filtered water (Aquaneering Inc., San Diego, CA), in accordance with Swedish regulations. To generate the required offspring, transgenic adult fish were in-crossed, and fertilized eggs were raised in an incubator at 28.5°C until 5 days post-fertilization (dpf). At 3dpf, embryos were optically screened for fluorescence in 96-well plates (EVOS FL Auto, Thermo Fisher Scientific, MA, USA), and embryos carrying the fluorescent transgene(s) were retained and placed back in the incubator. From 5 to 10dpf, zebrafish larvae were kept in 1L tanks filled with 300mL of water at a density of 30 larvae/tank. Larvae were fed twice daily until 9dpf. At 7dpf, waste products and debris were removed from the water, followed by replenishing of the water level to 300ml. *Tg:hsp70:IK17-EGFP* larvae were subject to a 37°C heat shock for 1 hour at 9dpf, to induce expression of the transgene for screening (see ‘Experimental procedure, imaging’).

### 3 Dietary intervention

To identify the atherogenic potential of overfeeding and dietary cholesterol supplementation, larvae from all backgrounds were fed on one of six diets from 5 to 9dpf before being screened at 10dpf. Diets consisted of a normal (∼5mg/feeding/tank) or larger amount (∼15mg/feeding/tank) of: 1) standard dry food (Golden Pearls, 50-100 μm particles, Alcester, UK); 2) standard dry food supplemented with 4% (wt/wt) extra cholesterol (≥99%, Sigma-Aldrich, Stockholm, Sweden) using a 1:1 volume ratio of diethyl ether to food (>99%, Fisher Scientific, Stockholm, Sweden)^18^; or 3) standard dry food treated with the same amount of diethyl ether without extra cholesterol. The latter condition was added to distinguish between effects of dietary cholesterol supplementation and/or treatment of the food with diethyl ether per se. To ensure the standard and cholesterol-supplemented diets were provided in energy balance, we assessed the energy density of both diets using blinded bomb calorimetry measurements on four samples per diet (C200 calorimeter, IKA-Werke GmbH & Co. Kg., Staufen, Germany). Since the energy density was on average slightly higher for the cholesterol-supplemented diet than for the regular dry food (i.e. 22.40 vs. 21.70 kJ/g), we fed larvae on slightly more regular dry food with and without treatment with diethyl ether (5.2mg and 15.5mg/feeding/tank for normal and overfeeding) than cholesterol-supplemented diet (5mg and 15mg/feeding/tank).

At 10dpf, larvae were subject to optical screening of atherogenic traits (see ‘*Imaging*’), followed by assessment of whole-body lipid and glucose levels (see ‘*Lipid, glucose and protein quantification*’). To reach a sample size of ∼100 larvae per background per dietary condition, we repeated the experimental procedure 5-9 times per background (total 27 times). To avoid batch effects, all dietary conditions were included on each occasion, and we adjusted for batch in the statistical analysis. To avoid bias by the time of imaging, the six dietary conditions were imaged in a randomized manner across imaging days, and time of imaging was recorded for each larva and adjusted for in the statistical analysis.

### 4 Treatment with atorvastatin and ezetimibe

Combined treatment with atorvastatin and ezetimibe is a widely-used strategy to lower LDLc – as well as other key atherogenic parameters – in patients with hypercholesterolemia^60–62^. Results from small-scale studies in samples of 20 to 100 pooled larvae suggest that treating larvae fed on a cholesterol-supplemented diet with statins and/or ezetimibe may prevent the elevated whole-body LDLc and/or total cholesterol levels that are otherwise observed^24,25^. In addition, evidence suggests that treatment with atorvastatin and ezetimibe may reduce vascular lipid deposition^25,63^.

To examine the anti-atherogenic potential of combined treatment with atorvastatin and ezetimibe, we overfed larvae of three backgrounds on a cholesterol-supplemented diet – as described earlier – in the presence or absence of 6μg atorvastatin and 80μg ezetimibe per 1g of dry food^63^ from 5 to 9dpf. At 10dpf, larvae were optically screened for atherogenic traits (see ‘*Imaging*’) and used for enzymatic assessment of whole-body lipid and glucose levels (see ‘*Lipid, glucose and protein quantification*’). To reach a sample size of 100 to 200 larvae per background per condition (treated vs. untreated), we repeated the experimental procedure 3 to 4 times across the three backgrounds (total 10 times). To avoid batch effects, treated and untreated larvae were included on each occasion. To avoid bias by the time of imaging, both conditions were alternated during imaging and time of imaging was recorded for each larva.

### 5 Food intake

To examine if supplementation of food with extra cholesterol and/or atorvastatin and ezetimibe affect food intake, we examined food intake in 204 additional larvae that were overfed on: 1) standard dry food; 2) standard dry food supplemented with 4% extra cholesterol using diethyl ether; 3) standard dry food treated with diethyl ether without cholesterol supplementation; or 4) standard dry food supplemented with 4% extra cholesterol using diethyl ether and further enrichment with atorvastatin and ezetimibe. Larvae were overfed on one of the four diets from 5 to 7dpf as described earlier. Before the morning feeding of 8dpf, larvae were transferred to fresh water before feeding them on diet that had (additionally) been supplemented with a fluorescent tracer. The fluorescently labelled diet was prepared as described previously^64^. Briefly, 75μl of yellow-green 2.0 μm polysterene microspheres (FluoSpheres carboxylate-modified microspheres, Invitrogen, Carlsbad, CA, USA), supplied as a 2% solution, were mixed with 50 mg of food and 25μl of deionized water for each of the four diets. The mixture was left to dry overnight in the dark and crashed into fine powder the next day. We subsequently acquired Z-stacks of the gastrointestinal tract (30 images, 1.5μm apart) between 20 mins and 5 hours after the morning feeding (Supplementary Fig. 4). Two rounds of imaging were performed to reach the final sample size. With the exception of atorvastatin and ezetimibe (2^nd^ round only), all conditions were included in both rounds to avoid batch effects. To avoid bias by the time of imaging, conditions were alternated during imaging – consistently imaging two consecutive larvae per condition – and time of imaging was recorded for each larva to allow for statistical adjustment.

### 6 Genetic interventions

A rich body of literature supports the role of *APOE*, *APOB*, and *LDLR* in familial hypercholesterolemia and CAD^5–7^. To examine if zebrafish can be used for high-throughput screening of candidate genes for dyslipidemia, atherosclerosis and CAD, we performed a multiplexed, CRISPR-Cas9-based genetic intervention for these genes using the protocol described by Varshney et al.^65^.

The zebrafish can have multiple orthologues of any human gene, thanks to a duplication of the genome early in the evolution of teleost fish. Hence, we firstly identified two zebrafish orthologues for *APOE*, three for *APOB* and two for *LDLR* using Ensembl, a synteny search in Genomicus^66^, and a literature search^21^ (Supplementary Table 10). We then designed CRISPR guide RNAs (gRNAs) to target these zebrafish orthologues – aiming for an early exon – using chopchop^67^ and CRISPRscan^68^, and tested their efficiency by micro-injecting gRNAs and Cas9 into the cell at the single-cell stage in multiplex. Eight larvae per multiplex were sacrificed at 3dpf and used for fragment length PCR analysis, to establish the efficiency of the gRNAs. For each orthologue, we selected a gRNA that showed moderate to high mutagenic efficiency, i.e. additional peaks in the fragment length spectrum in at least four of the eight larvae, while also retaining the wildtype peak (Supplementary Table 11). Pilot experiments in our lab indicated this approach can be anticipated to yield an adequate number of homozygous mutants for each orthologue in the F1 generation.

Identifying a gRNA with moderate to high mutagenic efficiency on average required six attempts for the seven orthologues (range 2 to 10) (Supplementary Table 11). The seven selected gRNAs for orthologues of *APOE*, *APOB*, and *LDLR* – one per orthologue – were subsequently co-injected in the cell of fertilized eggs from *Tg*(*mpeg1-mCherry*; *mpo-EGFP*) parents at the single-cell stage. Founder mutants were optically screened for the presence of the *Tg:mpeg1-mCherry* and *Tg:mpo-EGFP* transgenes at 4dpf, and carriers were raised to maturity. Founder mutants were then in-crossed, and offspring (F1) were overfed on a cholesterol-supplemented diet from 5 to 9dpf, followed by optical screening for atherogenic traits at 10dpf (see ‘*Imaging*’) and enzymatic assessment of whole-body LDLc, HDLc, triglyceride, total cholesterol, and glucose levels (see ‘*Lipid, glucose and protein quantification*’). DNA was then extracted and larvae were paired-end sequenced for the CRISPR-targeted sites (see ‘*DNA extraction and paired-end sequencing*’). To reach a sample size of 384 larvae per multiplex and background, we repeated the experimental procedure eight times. This sample size allows automated downstream sample preparation for paired-end sequencing in multiplex (see ‘*DNA extraction, sample preparation and paired-end sequencing*’).

The procedure described for the proof-of-concept genetic intervention was repeated for four DEPICT^34^-identified candidate genes in triglyceride-associated loci^26^, i.e. *LPAR2*, *TM6SF2*, *GATAD2A* and *GMIP*. Together, these four genes have six orthologues in zebrafish (Supplementary Table 29), and identifying moderate or highly active gRNAs on average required two attempts (range 2 to 4) (Supplementary Table 30). Phenotypically characterizing 384 larvae for the multiplexed mutant line with six targeted candidate genes required repeating the experiment four times.

### 7 Experimental procedure

#### 7.1 Imaging

On the morning of 10dpf – before the usual morning feeding – all tanks were blinded for dietary or drug treatment condition (not applicable to the genetic interventions) to ensure unbiased imaging, annotation, and quality control of images (see ‘*Image quantification*’). Immediately before imaging each tank, 15 to 20 larvae were simultaneously soaked in 25μM MDH in PBS for 30 mins, to enable visualization of circulating lipids and vascular lipid deposition. After soaking in MDH, the larvae were placed in a petri dish and anesthetized using tricaine (0.04 mg/ml). Larvae were subsequently aspirated one-by-one using a Vertebrate Automated Screening Technology (VAST) BioImager (Union Biometrica Inc, Geel, Belgium)^69,70^, which was mounted on the stage of a Leica DM6000B LED automated upright fluorescence microscope (MicroMedic AB, Stockholm, Sweden). The VAST BioImager automatically loads and positions larvae into a borosilicate capillary, where they are detected by the system’s camera. Whole-body images (n=12) were acquired during one full rotation, followed by automated rotation to the lateral orientation, as pre-specified using template images. The VAST BioImager subsequently positioned the larva so the caudal vein and dorsal aorta immediately caudal of the rectum were located within the field of view of the microscope (i.e. ∼2.9 mm from the tip of the nose), and triggered the microscope to start imaging. The researcher then manually focused on the center of the vasculature in z using the MDH channel, followed by the automated acquisition of 17 optical sections above, and 17 below the focal point – one every 1.5μm – using a Leica dip-in objective with 20X magnification (Leica OBJ HCX APO L 20X/0.50 W). This procedure was automatically repeated for each of the up-to-three channels per larva – i.e. to visualize the MDH dye as well as the *EGFP-* and *mCherry*-labelled transgenes – using the Leica 405, L5, and TXR filters, respectively. For each channel, the fluorescence signal was recorded using a Leica DFC365 FX CCD camera. Upon completion of optical sectioning in all channels, the larva was dispensed into a 96-well plate for further processing, and the next larva was loaded for imaging. This procedure takes up to 2 mins per larva.

#### 7.2.1 Quantification of morphological features in zebrafish larvae

Whole-body images of larvae acquired by the bright field camera of the VAST BioImager were used to quantify body length, dorsal and lateral body surface area, and body volume. To distinguish the larva from the capillary in which it was positioned, the capillary was assumed to be horizontal and the position of the capillary was obtained by projecting all pixel intensity values to the y-axis. That is, for each y-level, the sum of all pixels on that level in x was computed (Fig. 1a). The edges of the capillary appeared as minima of the projection, and the position of the inner walls were defined as the inner slopes of those minima with the steepest angle, i.e. the highest absolute derivative. Once the region inside the capillary was defined, larvae were segmented from the image background using grey-level thresholding based on optimized precision with regards to a given size interval^71^. This method efficiently tries all possible threshold levels and selects the threshold level that maximizes the per-threshold-precision (true positive / (true positive + false positive), where true positive is defined as the number of pixels in objects within the size interval and false positive is defined as the number of pixels in objects outside the size interval. This pre-processing was performed in ImageJ. Holes within the binary mask were filled automatically using CellProfiler^72^, and the largest connected component was extracted as the final segmentation. The dorsal and lateral surface area of the larva was computed as the number of pixels in this final mask, and the body length was estimated as the largest distance between two points on the larva outline touching a bounding box in the dorsal orientation (Fig. 1a).

Fluorescence signals from MDH (lipids), *mCherry* (macrophages) and *EGFP* (neutrophils, oxLDL, endothelial cells) were quantified using custom-written scripts in ImageJ, CellProfiler and ilastik^73^. Firstly, the maximal projection of each fluorescent channel was computed across all optical sections in z using ImageJ, to yield a single image containing signal (and noise) from multiple focal depths. Next, CellProfiler was used to quantify the surface area for each fluorescence signal across the z-stack. Images were first cropped in y using the MDH signal to only include the region from the center of the dorsal aorta to immediately caudal of the caudal vein. The fluorescence signal was then separated from background and noise using an ilastik-based, lenient pixel classifier that takes fluorescence intensity into account. Further segmentation was performed using a CellProfilerAnalyst-based object classifier in which criteria based on area, shape, texture and intensity were summarized in ten rules (see ‘*code availability*’). Subsequent feature extraction in which the surface area and shape of each identified object were quantified provided the total number of objects and the surface area covered by those objects. In addition, we created a mask to quantify two-way co-localization of MDH-stained lipid deposits or oxLDL with macrophages and neutrophils. Similarly, the proportion of lipid deposits that was located inside the vascular endothelium was calculated by creating a mask for the lipid deposits on top of the segmented vascular endothelial cells. Lipid deposits that overlapped with the endothelial cell layer and/or circulating lipids were considered to reside inside the endothelium; a requirement for atherogenic lipid deposits.

Food intake was quantified in the acquired images by first computing maximal projection of the acquired z-stacks using ImageJ^74^, to yield a single image containing signal (and noise) from multiple focal depths. Next, grey-level thresholding was applied in CellProfiler to quantify the total surface area of the fluorescence signal.

All procedures have been incorporated in pipelines that can be run in an automated manner on at least 2000 images simultaneously.

#### 7.2.2 Sensitivity and specificity for vascular lipid deposition

Accurate identification and quantification of vascular lipid deposits in an automated manner is challenging, since the MDH dye stains both vascular lipid deposits and circulating lipids. To ensure adequate detection of vascular lipid deposits, we calculated the sensitivity and specificity of the image quantification pipeline. To this end, researchers MKB and MdH manually annotated vascular lipid deposits in 30 randomly selected images from the *Tg:mpeg1-mCherry*; *mpo-EGFP* background across the six dietary conditions (blinded) in 3D using the acquired z-stacks. These 6×30 images had not been used to train the pixel and object classifiers. MKB and MdH subsequently discussed the results of the manual annotation process, and resolved discrepancies in judgment where needed. The results of the manual annotation process (gold standard) were then compared with the projections generated by the image quantification pipeline, in which the lipid deposits identified by both pixel and object classifier had been highlighted. Doing so allowed us to quantify the number of true positive (TP), false positive (FP) and false negative (FN) lipid deposits. The average sensitivity and specificity of the image quantification pipeline across the six dietary conditions were 72% and 93%, respectively (calculated using Stata’s ‘diagti’).

#### 7.3 Lipid, glucose and protein profiling

After imaging was completed, the anesthetized larvae were euthanized by exposure to tricaine (MS-222, Sigma, Sweden) and ice. All excess liquid was removed from the well, and one 1.4mm zirconium bead (Diagnostics, NJ, USA) and 75μl ice-cold PBS 1X were added to each well. The tissue was subsequently homogenized for 2 mins at 1000 rpm (1600 MiniG-Automated homogenizer, Gammadata Instruments, Uppsala, Sweden) and centrifuged at 3500 rpm for 5 mins at 4°C (13,000 rpm when using tubes). After centrifugation, 12.5μl of supernatant was removed and added to a new 96-well plate for protein quantification, together with 12.5μl of ice-cold PBS per well. The remaining supernatant (∼60μl/well) was transferred to Eppendorf tubes, together with 160μl of ice-cold PBS 1X (to a total volume of 220μl/well), and stored at −80°C for profiling of LDLc, HDLc, triglyceride, total cholesterol and glucose levels. Samples were subsequently stored at −80°C prior to analysis.

Protein content was assessed using the Pierce bicinchoninic acid (BCA) Protein Assay Kit (Thermo Fisher Scientific, Waltham, MA, USA) and a Varioscan LUX Microplate Reader (Thermo Fisher Scientific, Waltham, MA USA). LDLc, HDLc, triglyceride, total cholesterol and glucose levels were quantified using a fully automated Mindray TM BS-380 analyzer (Mindray Medical International, Shenzhen, China) using direct LDLc (1E31), HDLc (3K33), triglyceride (7D74), cholesterol (7D62), and glucose (3L82) reagents from Abott Laboratories (Abott Park, IL, USA). All analyses were blinded to dietary or treatment condition or genotype, respectively.

### 7.4.1 DNA extraction, sample preparation and paired-end sequencing

For larvae that were part of the genetic intervention, the pellet that remained after lipid, glucose and protein profiling was used to extract DNA. To this end, 50μl of lysis buffer containing proteinase K (diluted 1:100) was added to each well, followed by incubation at 55°C for 2h, and incubation at 95°C for 10 mins to heat-inactivate the proteinase K. Samples were then centrifuged at 3500 rpm for 2 mins and the supernatant was transferred to a new 96-well plate. A two-step PCR reaction subsequently incorporated Illumina Nextera XT v2 indices into the PCR products (Illumina Inc, San Diego, CA) using a Hamilton Nimbus 96 liquid handling system (Hamilton Robotics AB, Kista, Sweden), followed by paired-end sequencing (2×250 bp) on a MiSeq (Illumina Inc, San Diego, CA) at the National Genomics Infrastructure (NGI) Sweden. This procedure allows us to combine samples from up to eight 384-well plates – i.e. 8×384 larvae with 8×8 different target sites – in a single sequencing lane while retaining >100X coverage, on average. The combination of sequence and indices allows post-sequencing linking of reads to individual larvae (see ‘post-sequencing data analysis’).

#### 7.4.2 Post-sequencing data analysis

The MiSeq generated two de-multiplexed, paired-end .fastq files per larva (2×250 bp). A custom-written Perl script was used to split the reads into separate .fastq files for each CRISPR-Cas9 targeted site, and remove the insert sequences from the .fastq files. The paired-end sequences were processed at the same time, extracting the sequence in between the two primers if both primers were present in the read, or the sequence downstream of the first primer if only the first primer was observed (to prevent excluding longer reads *a priori*). No mismatches in the primer(s) were allowed to ensure optimal data quality. We subsequently used the fast and accurate Illumina Paired-End read mergeR (PEAR)^75^ to merge the trimmed paired-end reads; FastX version 0.0.14^76^ to remove reads containing bases with a quality score below 20 (-q 20, -p 100); and Spliced Transcripts Alignment to a Reference (STAR) version 2.4.1c^77^ to map the reads to the reference genome (Danio_rerio.GRCz11.dna.toplevel.fa as downloaded from Ensembl). SAMtools version 0.1.19^78^ was used to convert files from SAM to BAM format and sort and index BAM files, as well as to generate a summary of the coverage of mapped reads on a reference sequence at a single bp resolution (using the ‘mpileup’ utility).

A custom-written variant calling algorithm in R (DIVaH - Danio rerio Identification of Variants by Haplotype) was used to identify the two most prominent reads per larva and target site that passed quality control. That is, reads with a length difference compared with the reference sequence of less than 170 bp, and with an alignment report string (Concise Idiosyncratic Gapped Alignment Report [CIGAR]) shorter than 50 characters (Supplementary Tables 12 and 31). Variants located within 30 bp of the CRISPR target site were subsequently functionally annotated using Ensembl’s variant effect predictor (VEP) (Supplementary Tables 13 and 32). At each larva, target site and allele, the variant with the highest predicted likelihood of functionally affecting protein function was retained. Allele-specific scores (no annotation=0; modifier=0.2; low=0.33; moderate=0.66; high=1) were then calculated, and summing across the two alleles yielded an orthologue-specific dosage score for each targeted site (i.e. orthologue) in each larva.

In the proof-of-concept genetic screen, a median of 18 unique CRISPR-Cas9 induced mutations with a predicted detrimental effect on protein function were observed across the seven targeted orthologues in offspring of founder mutants (interquartile range 16 to 24.5 mutations, Supplementary Table 13). Of the 138 unique mutations identified across the seven target sites, 54 were frameshift deletions (40.0%), nine introduced a premature stop codon, and 34 were missense variants (24.6%). VEP predicted that of the 138 unique mutations, 18 were modifiers (13.0%), while four, 51 and 65 were assigned a low, moderate or high likelihood of affecting protein function (2.9%, 37.0% and 47.1%, respectively, Supplementary Table 13). The mutant allele frequency across the seven targeted zebrafish orthologues was typically high in the F1 generation (median 0.88, interquartile range 0.52 to 0.98, Supplementary Table 14).

In the discovery screen for candidate genes in the triglyceride, LDLc, total cholesterol and type-2 diabetes-associated locus on chr 19p13.11, three, one and seven unique CRISPR-Cas9 induced mutations with a predicted detrimental effect on protein function were observed in *lpar2a*, *lpar2b* and *gatad2ab* in offspring of founder mutants (Supplementary Table 32). All larvae were wildtype for the zebrafish orthologues of *TM6SF2* and *GMIP*, in spite of having pre-tested the CRISPR gRNAs for efficiency. Of the 11 unique mutations identified across *lpar2a*, *lpar2b* and *gatad2ab*, four were frameshift deletions (36.4%), four were inframe deletions (36.4%), and three were missense variants (27.3%, Supplementary Table 32). In addition to all F1 larvae being wildtype for three of the six targeted orthologues, the mutant allele frequency was very low across *lpar2a*, *lpar2b* and *gatad2ab*, with only 11, 6 and 18 of the 376 successfully sequenced larvae carrying one mutated allele. None of the larvae carried a mutated allele in more than one gene (Supplementary Table 33).

### 8 Quality control

After image quantification and before the statistical analysis, all quantified images were manually screened to ensure adequate quantification had occurred. Larvae for which the automated quantification pipeline had failed for a trait were annotated and excluded from the analysis for that trait, as well as for any traits that rely on adequate quantification of that trait. Annotations include: weak staining of the circulation by MDH (possibly reflecting low levels of circulating lipids), resulting in incorrect segmentation of the region of interest; inadequate segmentation of the region of interest for other reasons, like movement during imaging; more than 20% of true negative objects being detected as objects (i.e. many false positives); less than 20% of true objects being detected (i.e. many false negatives); and circulating neutrophils being present, resulting in the same neutrophil being quantified multiple times (Supplementary Table 1). Annotations that resulted in the exclusion of at least ten larvae were examined in more detail, to examine if the underlying reason for exclusion may have influenced the results (see below). Based on the large proportion of affected larvae and the absence of influence on the results, larvae with many false positive or many false negative oxLDL deposits were included in the analysis.

In the dietary and drug treatment interventions, all continuous outcomes and exposures outside the mean ± 5×SD (standard deviation) range were set to missing before the association analysis, to prevent outliers – be it biological or methodological – from driving the results. This step was omitted in the genetic interventions because larvae carrying two mutated alleles for causal genes were *a priori* anticipated to show extreme phenotypes. In addition, total cholesterol levels were set to missing if triglyceride levels were missing and vice versa. This resulted in the exclusion of images from a median of 2.5 larvae across all outcomes in the dietary intervention (inter quartile range 2.5 to 7), and one larva in the drug treatment intervention (interquartile range 0 to 2.75). Next, residuals were calculated to normalize: 1) vascular endothelial surface area for the surface area of circulating lipids, yielding a variable that reflects endothelial thickness; 2) LDLc, HDLc, triglyceride, total cholesterol and glucose levels for protein content of the sample; and 3) dorsal and lateral body surface area as well as body volume for body length. All analyses with these variables as outcomes or exposures were performed using normalized values. Finally, all continuous outcomes showing an approximately normal distribution were inverse-normally transformed to a mean of 0 and standard deviation (SD) of 1, to ensure all residuals in the association analyses were normally distributed. This transformation implies that all effect sizes (β), standard errors (SE) and 95% confidence intervals (95% CI) for these outcomes can be interpreted as z-scores, allowing a comparison of effect sizes across outcomes, conditions and experiments. Image-based vascular atherogenic outcomes that showed a negative binomial distribution – with or without inflation of zeros –were not inverse-normally transformed.

### 9 Statistical analysis

In the main analysis, we examined the effect of: 1) overfeeding and cholesterol supplementation (in the dietary intervention); 2) treatment with atorvastatin and ezetimibe (in the drug treatment intervention); and 3) mutations in proof-of-concept and candidate genes (in the genetic interventions) on body size; early-stage vascular atherogenic traits; and whole-body LDLc, HDLc, triglyceride, total cholesterol and glucose levels. This was accomplished using hierarchical linear models (xtmixed in Stata) or negative binomial regression (nbreg). Hierarchical linear models on inverse-normally transformed outcomes provide effect sizes and standard errors for the fixed factors, while providing the standard deviation of the outcome across random factors, for which the intercept – i.e. the value of non-exposed larvae – is allowed to vary. Body size, whole-body lipid and glucose levels, and some image-based atherogenic traits were analyzed this way, i.e. typically vascular infiltration by macrophages or neutrophils. However, most image-based vascular atherogenic traits showed negative binomial distributions. For such traits, the effects of dietary, drug treatment and genetic factors were examined using negative binomial regression.

All models were adjusted for: a) the use of diethyl ether (in the dietary intervention); and b) time of day at which the image was acquired (in all experiments) as fixed factors (xtmixed) or as regular co-variables (nbreg). Models were additionally adjusted for transgenic background and batch as random factors or as regular co-variables. For image-based vascular atherogenic traits, associations were examined with and without adjusting for body length and dorsal body surface area, by adding them as fixed factors or co-variables to the models. To ensure unbiased estimates, we only included data from larvae with information on body length and dorsal body surface area in the adjusted and unadjusted analyses, to ensure that effect estimates were based on data from the same larvae. This step was omitted in the genetic interventions to maximize the sample size of the analysis that was not adjusted for body size. For image-based atherogenic traits, we also examined if LDLc, HDLc, triglyceride, and/or glucose levels mediated the main effect of dietary, drug treatment and genetic factors, by adding them as additional fixed factors or co-variables to the size-adjusted model. The sample size was typically somewhat lower for the latter analyses due to missing data. Directed acyclic graphs (DAGs) indicated that based on the anticipated causal paths, this analysis plan should not have resulted in biased effect estimates. For image-based atherogenic traits, results from models that were additionally adjusted for body size were considered the main results, and are referred to throughout the results and figures.

A small subset of larvae with suboptimal image or image quantification quality were excluded from the analyses (Supplementary Table 2). To examine if exclusion of these larvae may have influenced the results, we examined if the odds of being affected was associated with the main exposures, i.e. diet, drug treatment or mutations. These analyses were performed using logistic regression models for annotations that affected at least ten larvae.

For all analyses, effect sizes, standard errors (robust standard errors for nbreg) and 95% confidence intervals are reported for the exposed compared with the unexposed group. Odds ratios (OR) and 95% confidence intervals are provided for analyses of image-based exclusions. All data management and statistical analyses were performed using Stata/MP 14.0 for Mac.

### 10 Ethical approval

All procedures were performed in line with Swedish regulations, and all experiments have been approved by Uppsala Djurförsöksetiska nämnd, Uppsala, Sweden (Permit numbers C142/13 and C14/16).

### 11 Code availability

All custom-written image analysis scripts; all post-sequencing QC and alignment scripts; the custom-written variant calling algorithm in R (i.e. DIVaH - Danio rerio Identification of Variants by Haplotype); and all Stata scripts used for statistical analysis are available from the corresponding author upon request.

### 12 Data availability

The data that support the findings of the current study are available from the corresponding author upon reasonable request.

## Acknowledgments

We are very grateful to Stephen A Renshaw from the University of Sheffield, Graham Lieschke from Monash University, Yury Miller from UCSD, and Dimitris Beis from the Biomedical Research Foundation Academy of Science for kindly providing us with carriers of the *Tg:mpo-EGFP*, *Tg:mpeg1-mCherry*, *Tg:hsp70:IK17-EGFP* and *Tg:flk-EGFP* transgenes, respectively. Input from Shawn Burgess and Gaurav Varshney on mutagenesis using CRISPR-Cas9 is also much appreciated. João Campos Costa’s efforts when setting up the lab are gratefully acknowledged, and we would also like to thank Francis Smet from Union Biometrica Inc and Thommie Karlsson from Leica for their help with installing the imaging set-up. Pilou Janssens’ work on bomb calorimetry of diets, and Lingjie Tao and Lisa Conrad’s help with sample preparation and pre-screening of larvae are also much appreciated. The computations were performed on resources provided by SNIC through Uppsala Multidisciplinary Center for Advanced Computational Science (UPPMAX) under Project SNIC b2015283. The authors would like to acknowledge support from Science for Life Laboratory, the National Genomics Infrastructure (NGI) and UPPMAX for aiding in massive parallel sequencing and computational infrastructure. Support from the National Bioinformatics Infrastructure Sweden (NBIS) is also gratefully acknowledged, and constructive discussions with the Genome Engineering Zebrafish (GEZ) facility are appreciated. Data on coronary artery disease / myocardial infarction have been contributed by the CARDIoGRAMplusC4D and UK Biobank CardioMetabolic Consortium CHD working group who used the UK Biobank Resource (application number 9922). Data have been downloaded from www.CARDIOGRAMPLUSC4D.ORG. MdH is a fellow of the Swedish Heart-Lung Foundation (20170872) and a Kjell and Märta Beijer Foundation researcher. This research was supported by project grants from the Swedish Heart-Lung Foundation (20140543, 20170678, 20180706), the Swedish Research Council (2015-03657), the Knut och Alice Wallenberg Foundation (2013.0126), the European Research Council (ERC-StG-335395) and NIH (R01DK106236, R01DK107786, U01DK105554).

## Author contributions

MKB, EI and MdH conceived the study; EI and MdH ascertained funding and provided material support; MKB, AE, BvdH, EM, MMM, TK and MdH performed the experiments; MKB, PR, CW and MdH generated the image quantification pipelines and performed the image-based analysis; MKB, AE, BvdH, TK and MdH optimized the CRISPR-Cas9 multiplex pipeline; EM, OD and MDH generated the post NGS QC and variant calling pipeline; AL assessed whole-body lipid and glucose levels; HLB and MdH performed the statistical analysis; MKB and MdH wrote the manuscript; all authors provided critical feedback to the manuscript.

## SUPPLEMENTARY FIGURES

**Supplementary Figure 1.**
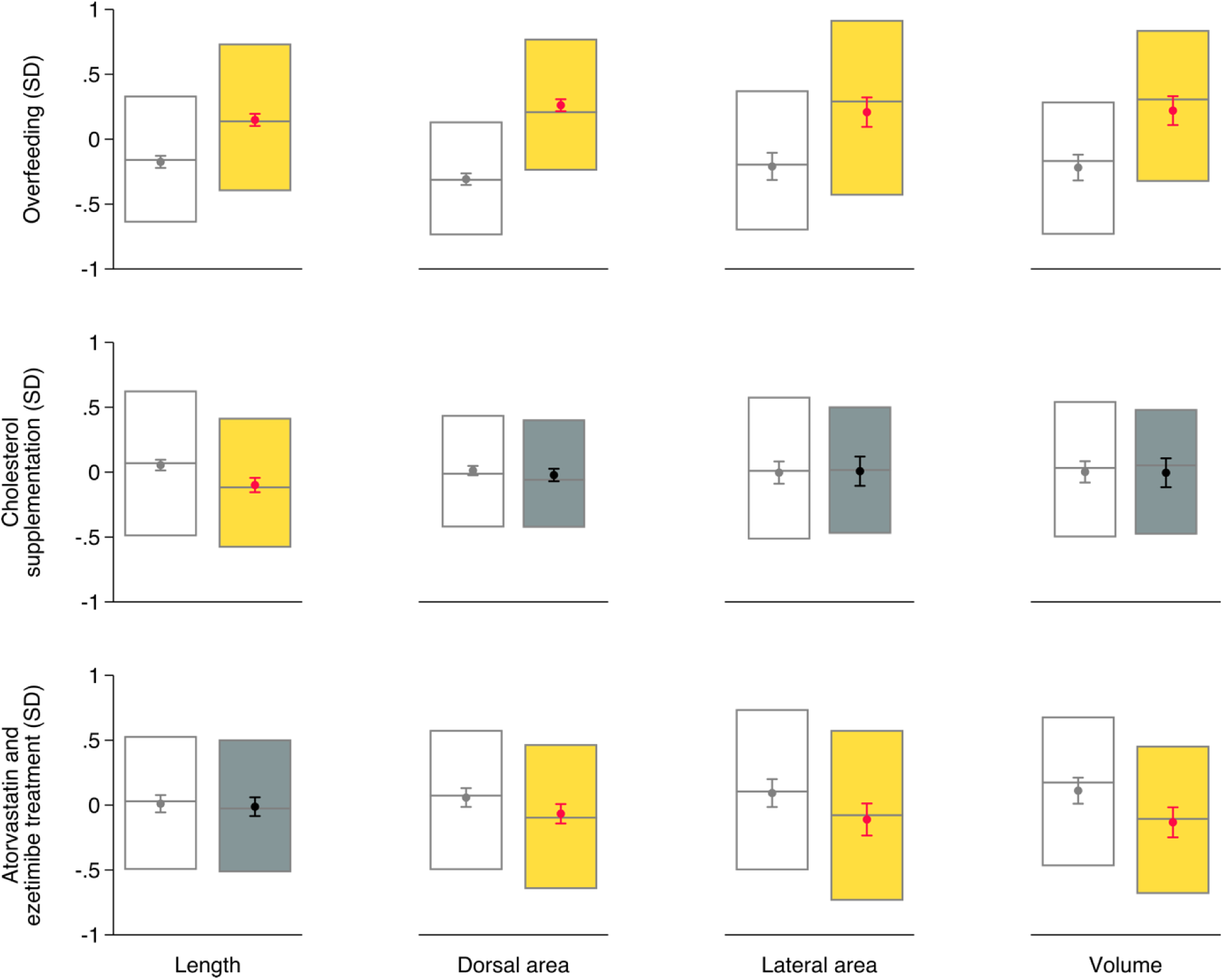
The effect of overfeeding (top), cholesterol supplementation (middle) and treatment with atorvastatin and ezetimibe (bottom) on body size. Dots and whiskers show mean and 95% confidence interval (CI); boxes show median and inter quantile range. Analyses were performed using residuals acquired using hierarchical linear models on inverse-normally transformed outcomes, adjusted for the use of diethyl ether (for overfeeding and cholesterol supplementation), cholesterol supplementation (for overfeeding), the amount fed (for cholesterol supplementation), and time of day as fixed factors. Larvae were nested in batches and transgenic backgrounds (random factors). White boxes with grey mean and 95% CI (left) show results for unexposed larvae; grey boxes with black mean and 95% CI (right) show results for exposed larvae that are not different from unexposed ones; yellow boxes with red mean and 95% CI (right) show results for exposed larvae that are different from unexposed ones at P<0.05.

**Supplementary Figure 2.**
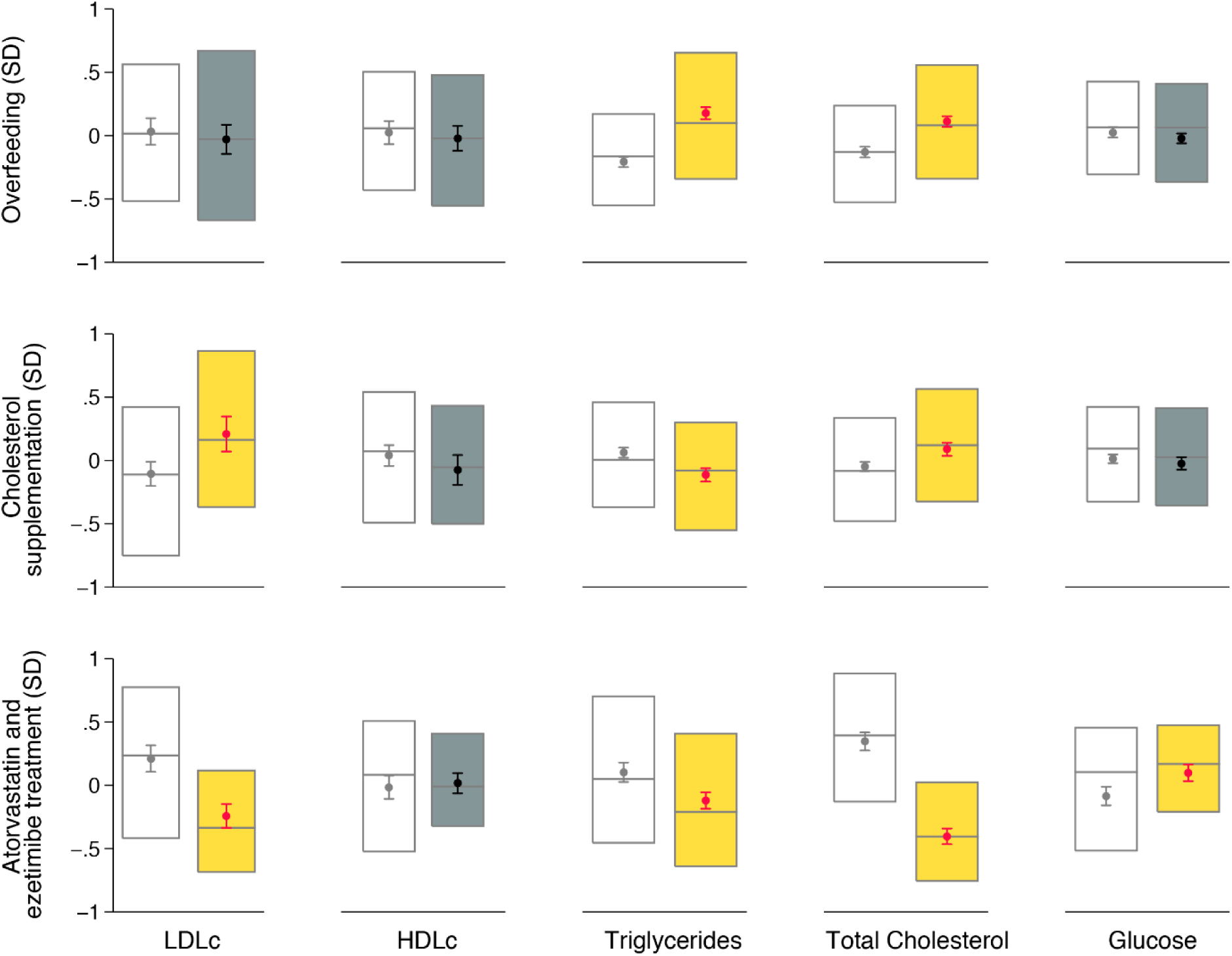
The effect of overfeeding (top), cholesterol supplementation (middle) and treatment with atorvastatin and ezetimibe (bottom) on whole-body lipid and glucose levels. Dots and whiskers show mean and 95% confidence interval (CI); boxes show median and inter quantile range. Analyses were performed using residuals acquired using hierarchical linear models on inverse-normally transformed outcomes, adjusting for the use of diethyl ether (for overfeeding and cholesterol supplementation), cholesterol supplementation (for overfeeding), the amount fed (for cholesterol supplementation) and time of day as fixed factors. Larvae were nested in batches and transgenic backgrounds (random factors). White boxes with grey mean and 95% CI (left) show results for unexposed larvae; grey boxes with black mean and 95% CI (right) show results for exposed larvae that are not different from unexposed ones; yellow boxes with red mean and 95% CI (right) show results for exposed larvae that are different from unexposed ones at P<0.05.

**Supplementary Figure 3.**
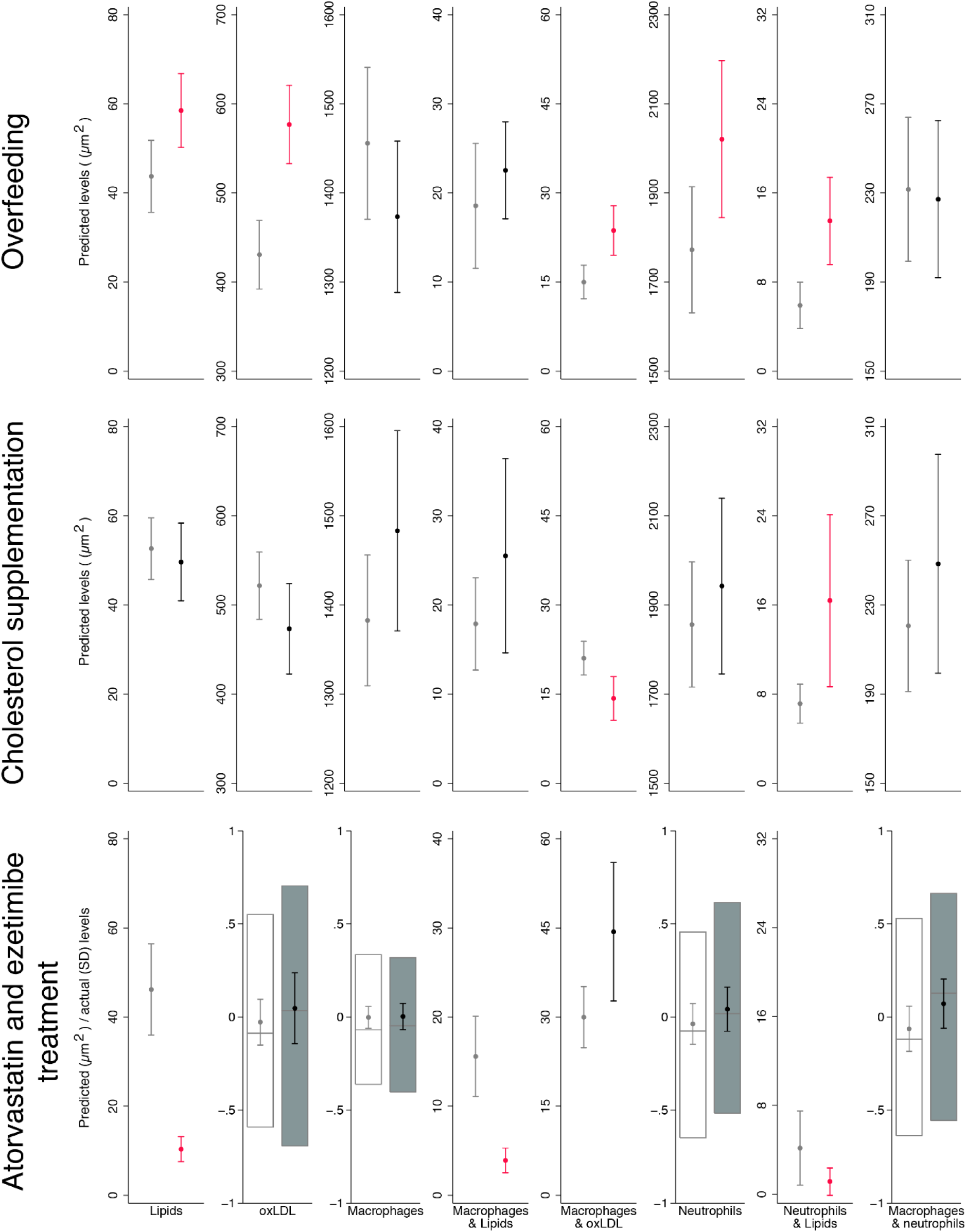
The effect of overfeeding (top), cholesterol supplementation (middle) and treatment with atorvastatin and ezetimibe (bottom) on vascular atherogenic traits. Outcomes showing effect estimate and 95% CI for predicted values have been analyzed using negative binomial regression, with adjustment for the same co-variables. Outcomes showing only effect estimate and 95% CI for predicted values have been analyzed using regular negative binomial regression, adjusting for the use of diethyl ether (for overfeeding and cholesterol supplementation), cholesterol supplementation (for overfeeding), the amount fed (for cholesterol supplementation), body length, dorsal body surface area, and time of day. Outcomes showing mean and 95% confidence interval (CI) as well as boxes for median and inter quartile range have been analyzed using hierarchical linear models on residuals after adjusting inverse-normally transformed outcomes for body length, dorsal body surface area, and time of day as fixed factors. Larvae were nested in batches and transgenic backgrounds (random factors). White boxes and/or light grey mean and 95% CI (left) show results for unexposed larvae; grey boxes and/or black mean and 95% CI (right) show results for exposed larvae that are not different from unexposed ones; red mean and 95% CI (right) show results for exposed larvae that are different from unexposed ones at P<0.05.

**Supplementary Figure 4.**
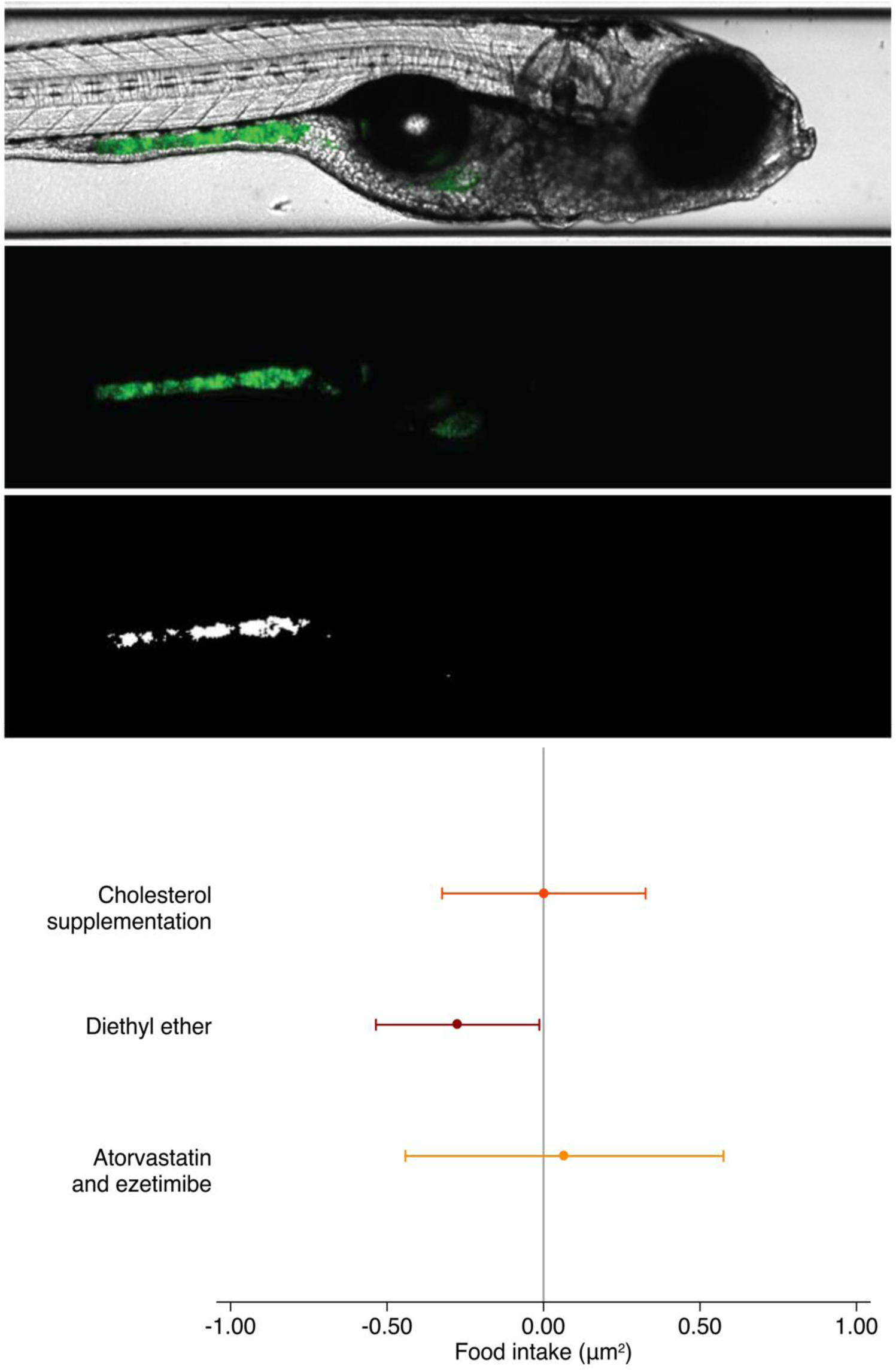
Food intake as a function of dietary or drug treatment intervention. Mixing fluorescently labelled tracers in with standard dry food, standard dry food enriched with 4% extra cholesterol using diethyl ether, standard dry food treated with diethyl ether, and standard dry food enriched with 4% extra cholesterol using diethyl ether and further enriched with atorvastatin and ezetimibe allowed image-based quantification of food intake - i.e. surface area of fluorescence in the gastrointestinal tract - in eight-day-old zebrafish larvae (top). Bottom: mutually adjusted effect of cholesterol supplementation, treatment of the diet with diethyl ether, and enrichment with atorvastatin and ezetimibe on food intake, assessed using dummy variables and negative binomial regression, additionally adjusted for time since feeding and batch (n=204). Dots and whiskers show effect size and 95% confidence interval.

**Supplementary Figure 5.**
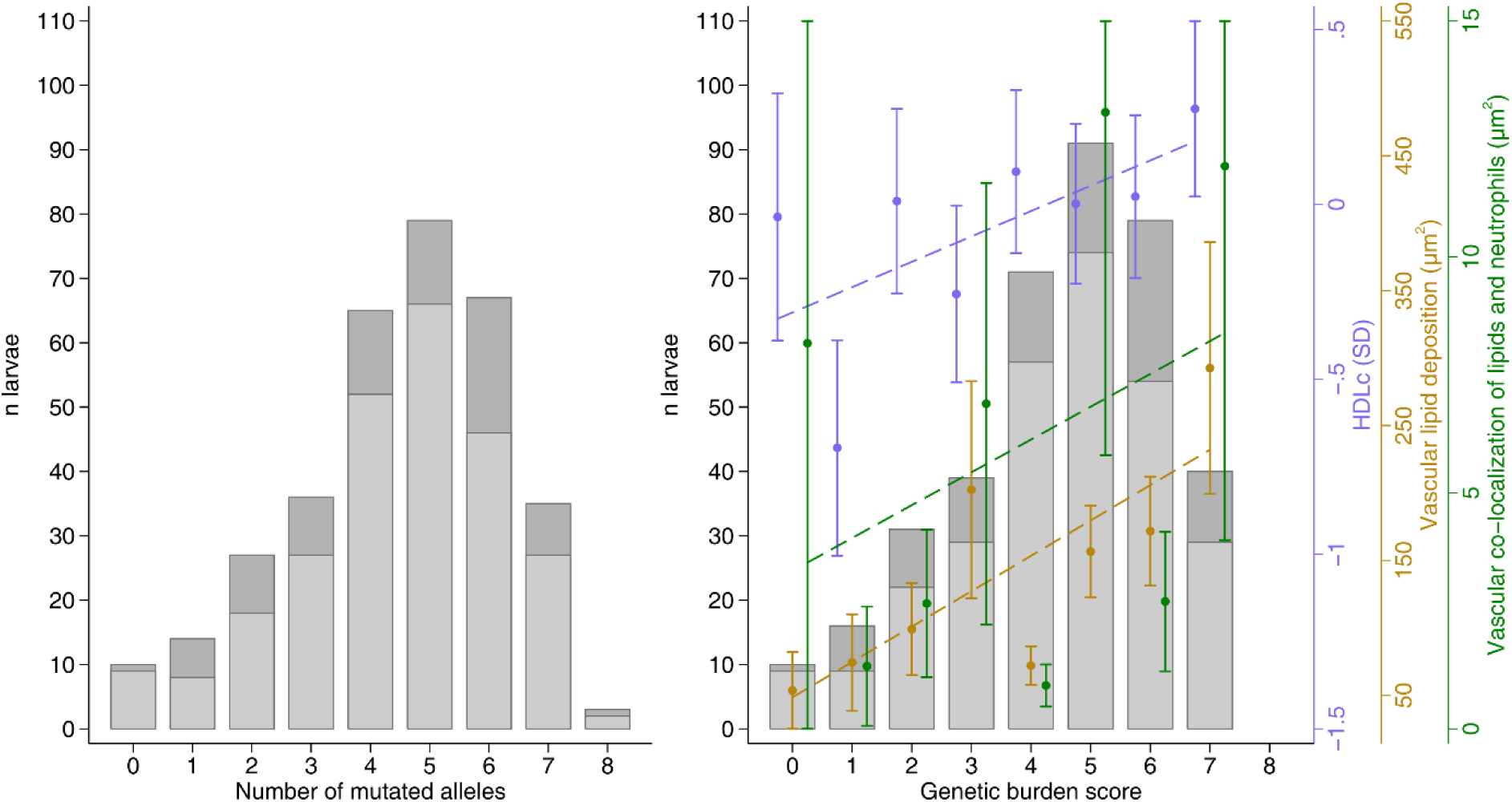
Histogram of the number of mutated alleles and genetic burden score across apoea, apoeb, apobb.1 and ldlra and association of whole-body HDL cholesterol levels, vascular lipid deposition and vascular co-localization of lipids and neutrophils with the genetic burden score. Left: histogram of the number of mutated alleles across apoea, apoeb, apobb.1 and ldlra. Larvae with two mutated alleles in apoba, apobb.2 and ldlrb are shown in light grey (bottom); larvae with at least one unaffected allele in these three genes are shown in dark grey (top). Right: as before, but with each affected allele weighed by the probability that it affects protein function, based on annotation using Ensembl’s variant effect predictor (VEP) (i.e. a genetic burden score). This figure also shows the association between atherogenic traits and the genetic burden score for significantly associated traits, adjusted for the number of mutated alleles in apoba, apobb.2 and ldlrb, i.e: 1) HDLc (n=381, in purple), assessed using a hierarchical linear model after inverse-normal transformation of LDLc, adjusted for time of day (fixed factors) and with larvae nested in batches; 2) vascular lipid deposition (n=272, in yellow); and 3) vascular co-localization of lipids and neutrophils (n=271, in green), using negative binomial regression, adjusted for body length, dorsal body surface area, time of day and batch. Dots and whiskers show mean and standard error of the mean, acquired using the margins command.

**Supplementary Figure 6.**
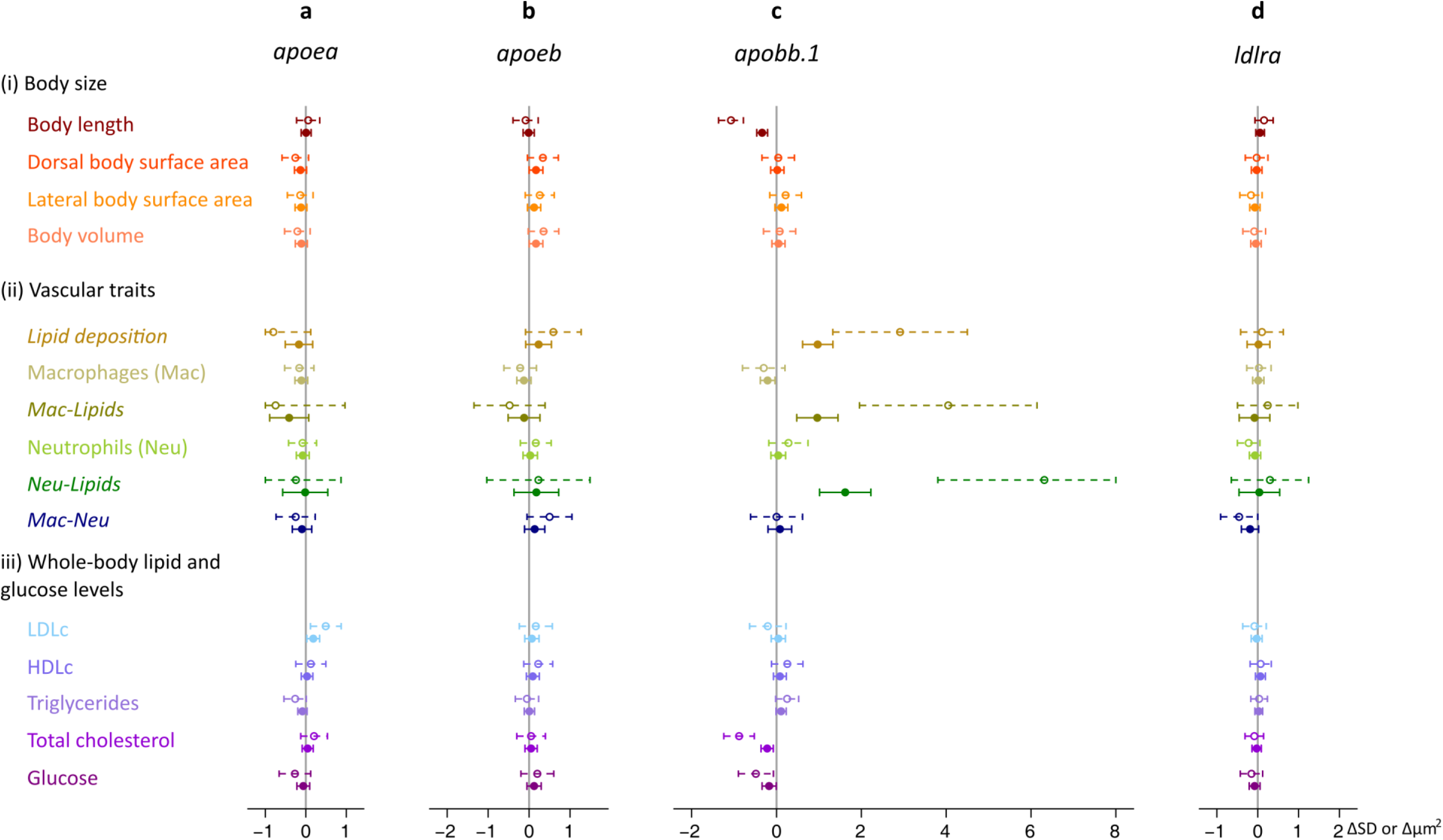
The effect of mutations in apoea, apoeb, apobb.1 and ldlra on body size (i), vascular atherogenic traits (ii) and whole-body lipid and glucose levels (iii). Dorsal and lateral body surface area and body volume were normalized for body length before the analysis; and whole-body lipid and glucose levels were normalized for protein levels. For normally distributed traits (shown in regular font), associations were examined using hierarchical linear models on inverse-normally transformed values. For these traits effect sizes and 95% confidence intervals are expressed in standard deviation units (SD). The remaining vascular atherogenic traits (shown in italic) were analyzed using negative binomial regression analyses. For these traits, effect sizes and 95% confidence intervals are expressed in µm^2^. Dotted lines represent the effect of two functionally knocked out alleles compared with zero mutated alleles. Regular lines show the additive per-allele effect. Associations were adjusted for time of day; batch; body length and dorsal body surface area (for vascular outcomes); and the number of mutated alleles in the other genes.

**Supplementary Figure 7.**
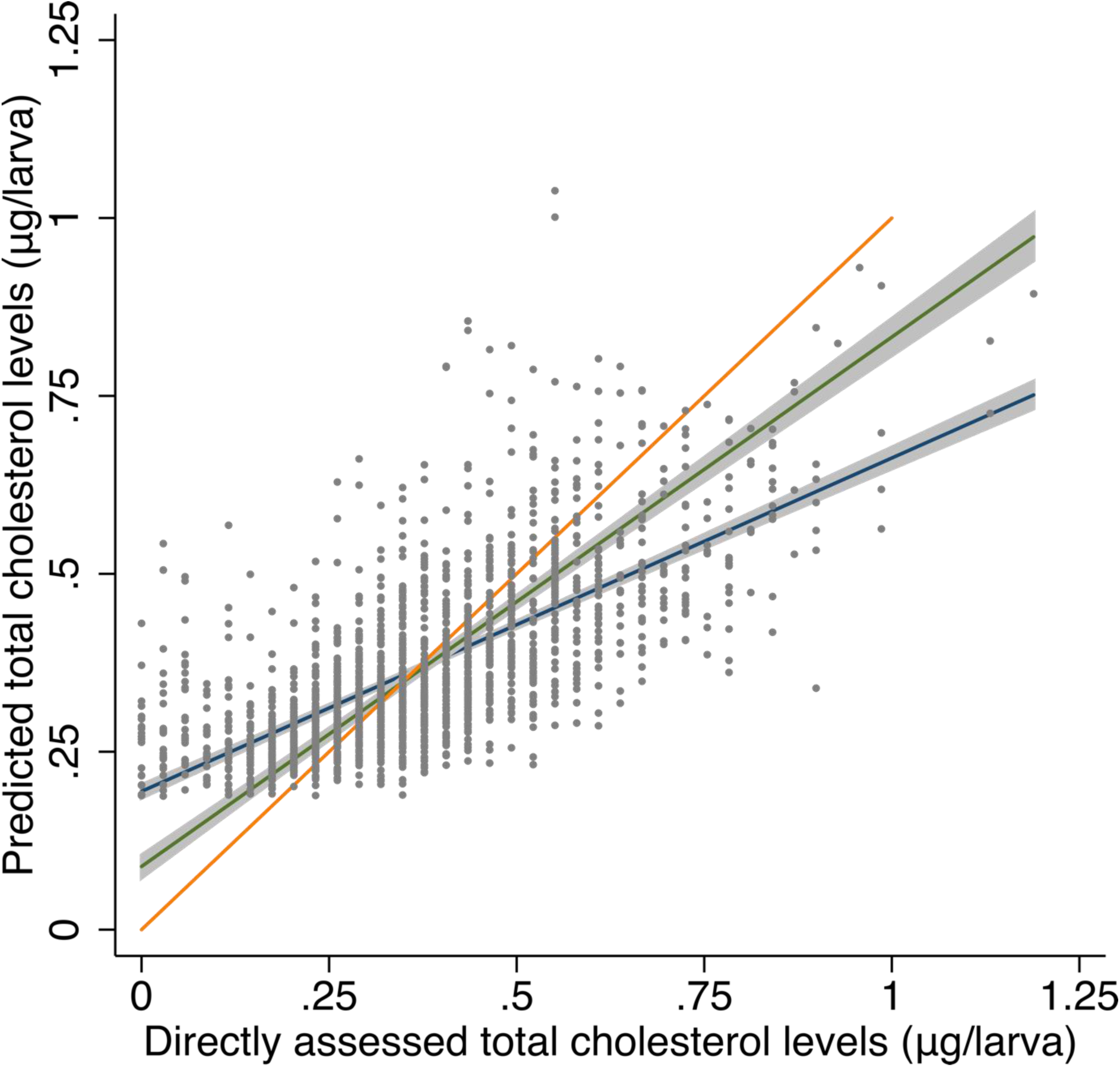
The association of predicted total cholesterol levels using regression of directly assessed LDLc, HDLc and triglyceride levels with directly assessed total cholesterol levels. In blue and grey are the regression line and 95% confidence interval (CI) (r^2^=0.468). In green and grey are the regression line and 95% CI for the association of total cholesterol levels calculated using the formula that is typically applied in humans (i.e. LDLc + HDLc + triglycerides/5) with directly assessed total cholesterol levels (r^2^=0.430). In orange is a line with a slope of 1 (n=1,867 larvae).

**Supplementary Figure 8.**
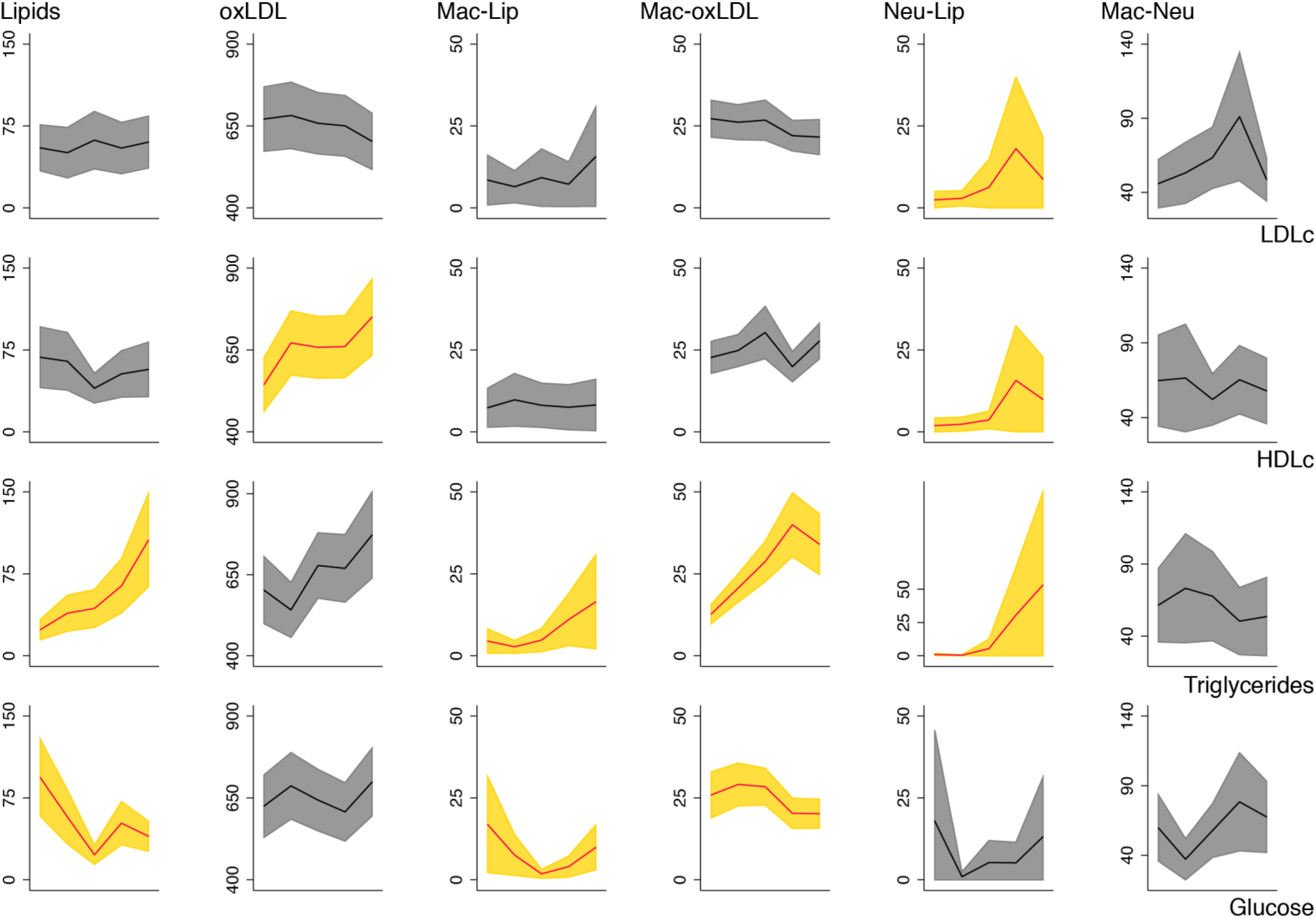
The association of vascular atherogenic traits with whole-body lipid and glucose levels in data from the dietary, drug treatment and genetic intervention for proof-of-concept genes combined. For each vascular atherogenic outcome (i.e. vascular lipid deposition [Lip] and accumulation of oxidized LDL [oxLDL]; and vascular co-localization of lipids and oxLDL with macrophages [Mac] and neutrophils [Neu]), mutually adjusted associations with protein-normalized levels of LDL cholesterol (LDLc), HDL cholesterol (HDLc), triglyceride and glucose levels were examined using negative binomial regression. Besides for the other main exposures, associations were adjusted for body length and dorsal body surface area, transgenic background and batch. Graphs show margins plots – highlighting mean and 95% confidence intervals – for the vascular atherogenic outcomes, expressed in µm^2^ (y-axes) with exposures grouped by quintile (x-axes). Significant associations are shown in yellow.

## SUPPLEMENTARY TABLES

**Supplementary Table 1.**
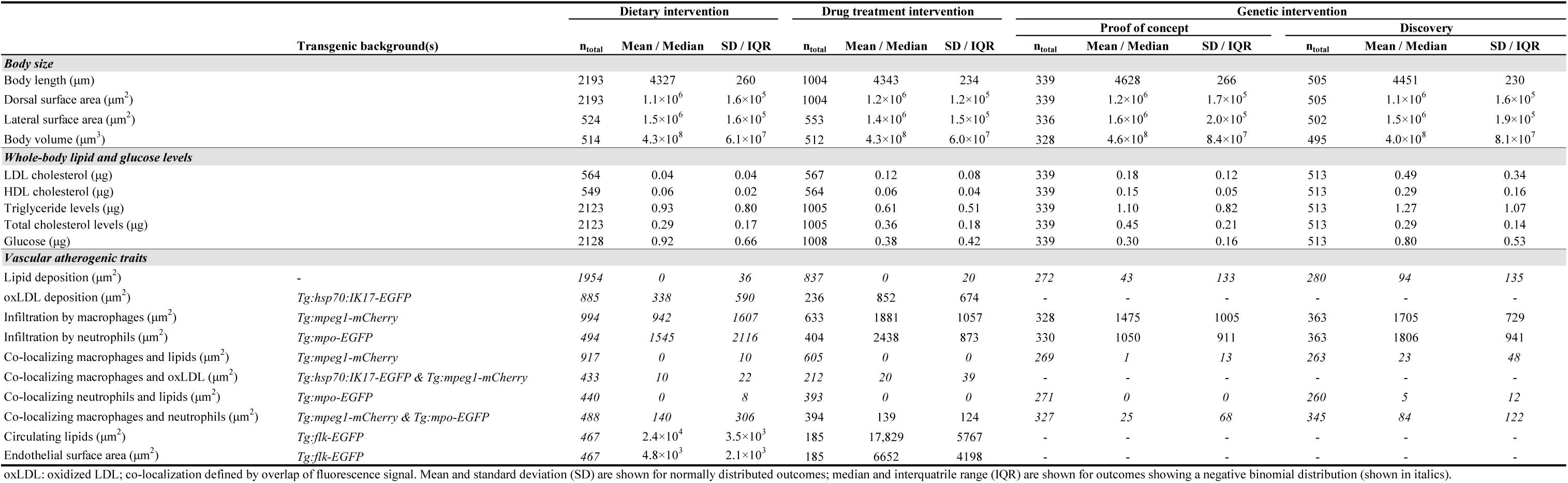
Descriptive information for larvae at 10 days post-fertilization in the dietary, drug treatment and genetic interventions

**Supplementary Table 2.**
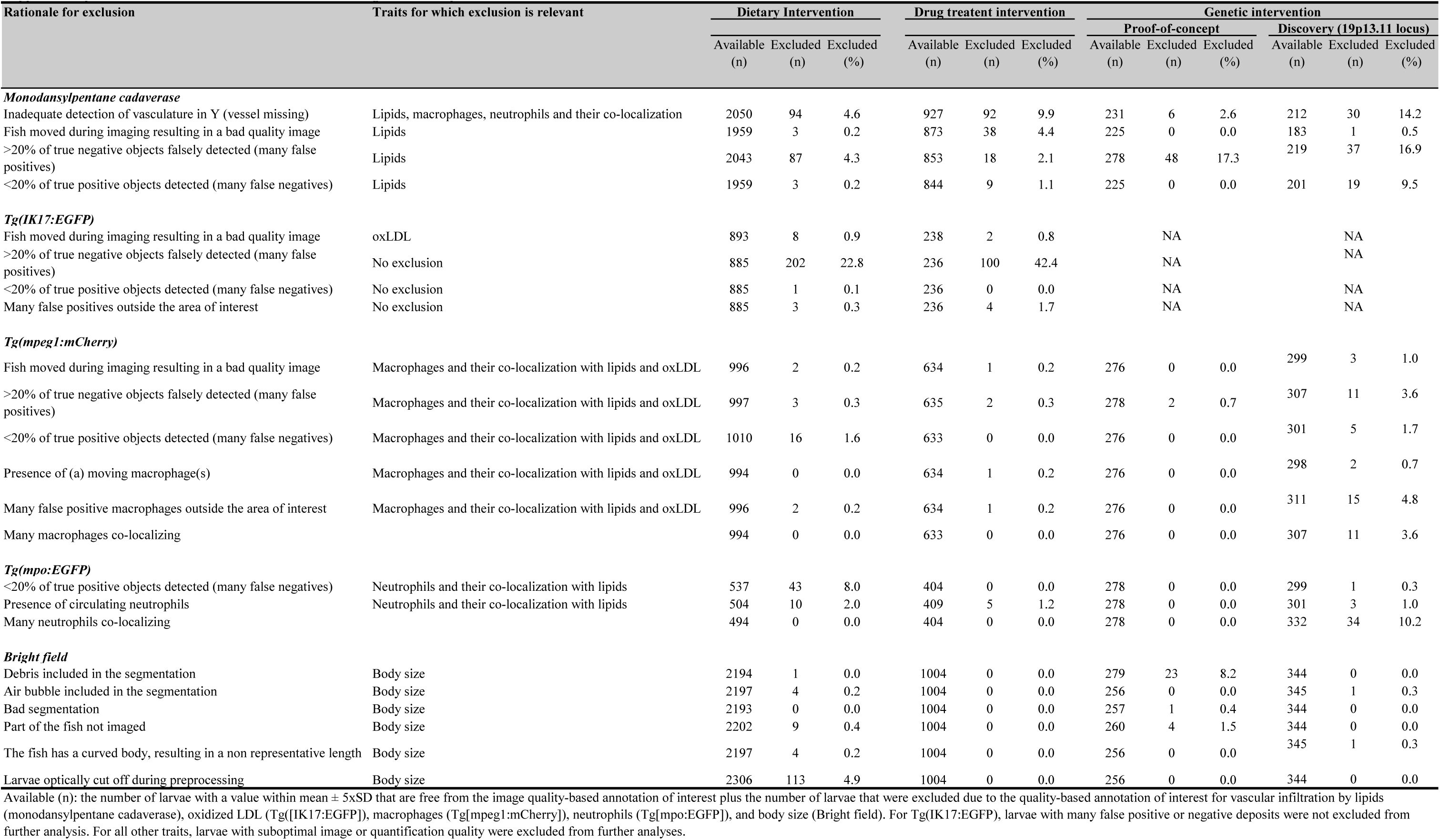
Annotation-based exclusions in the image-based analyses

**Supplementary Table 3.**
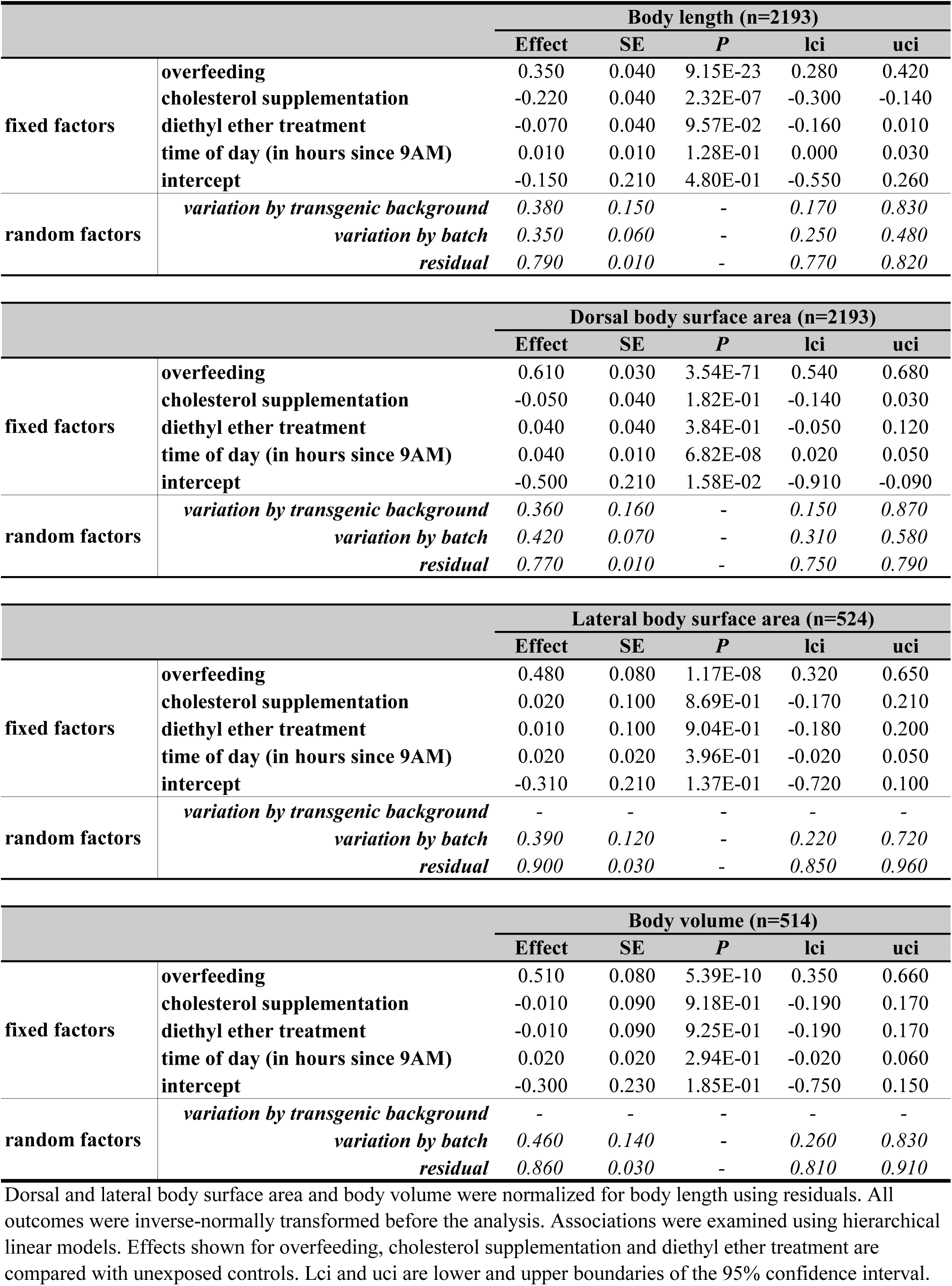
The effect of overfeeding and cholesterol supplementation on body size

**Supplementary Table 4.**
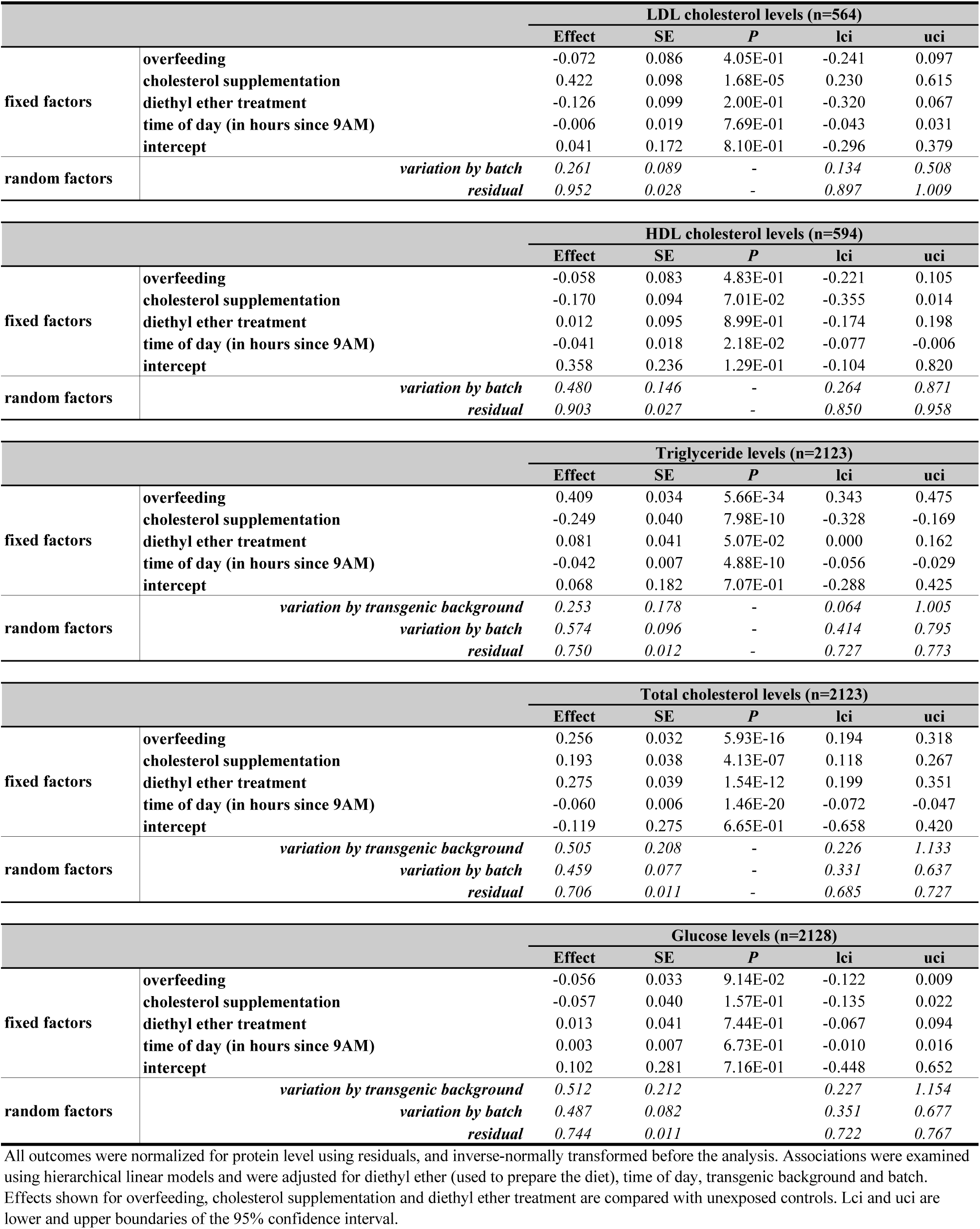
The effect of overfeeding and cholesterol supplementation on whole-body lipid and glucose levels

**Supplementary Table 5.**
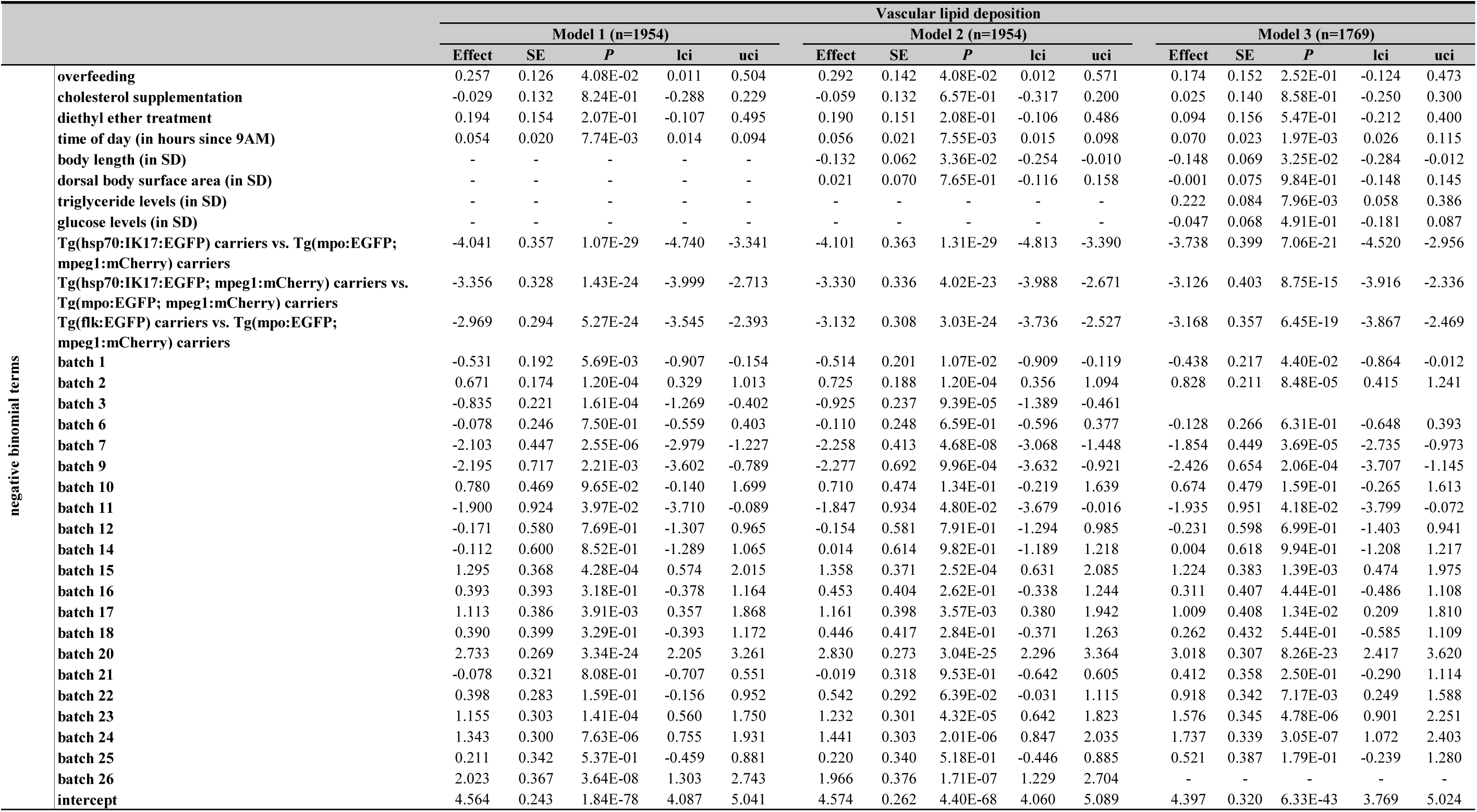
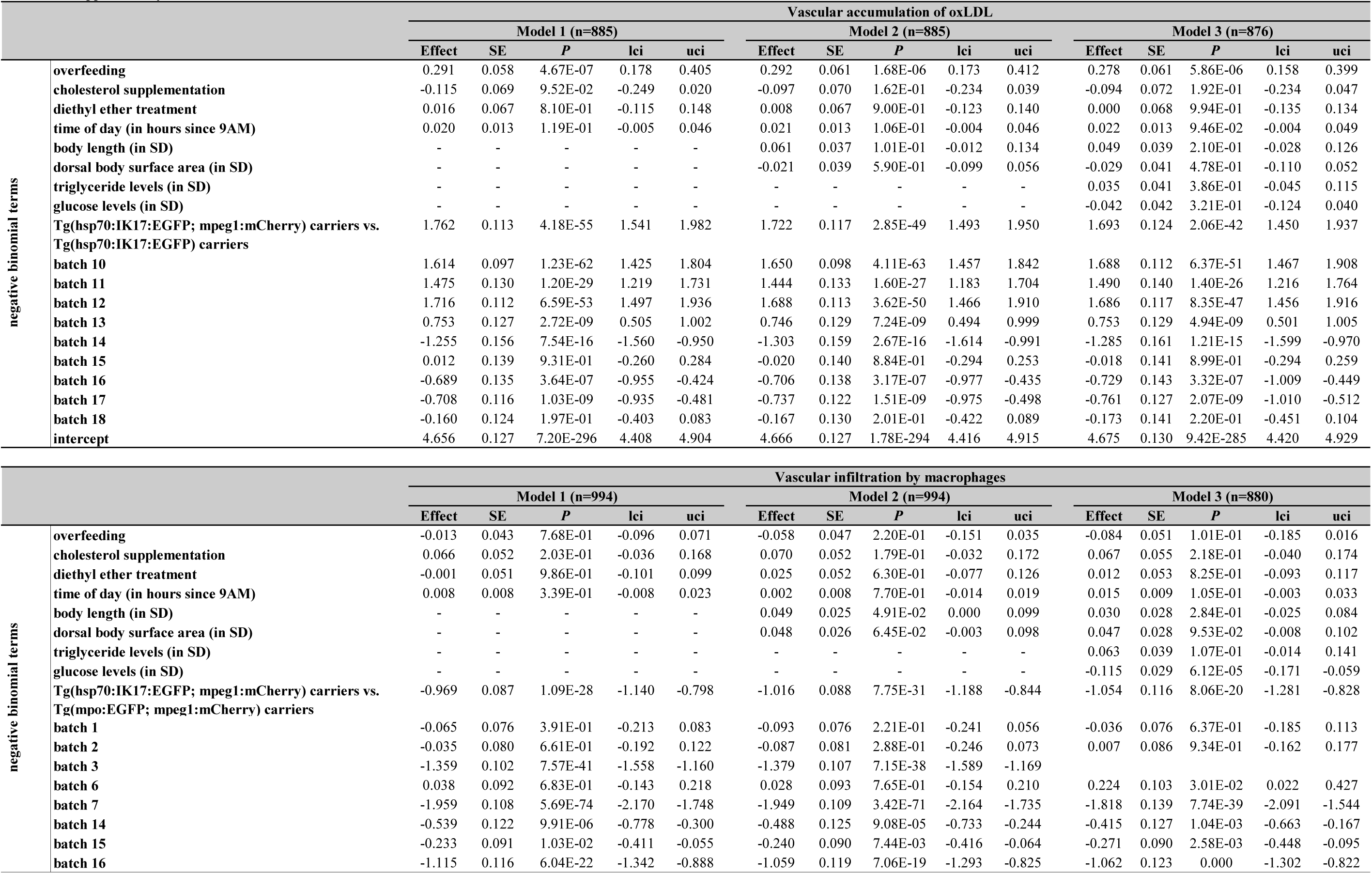
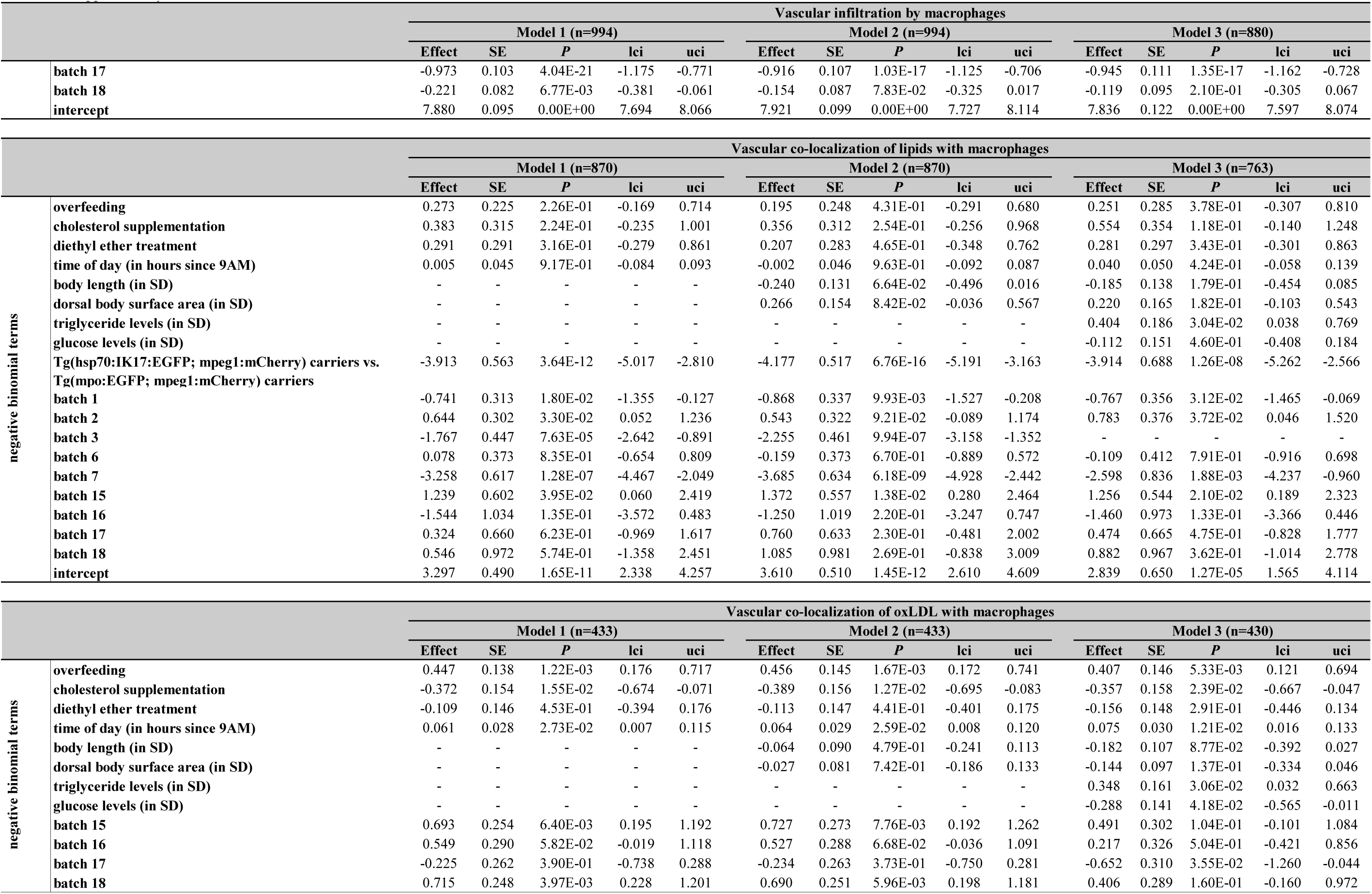
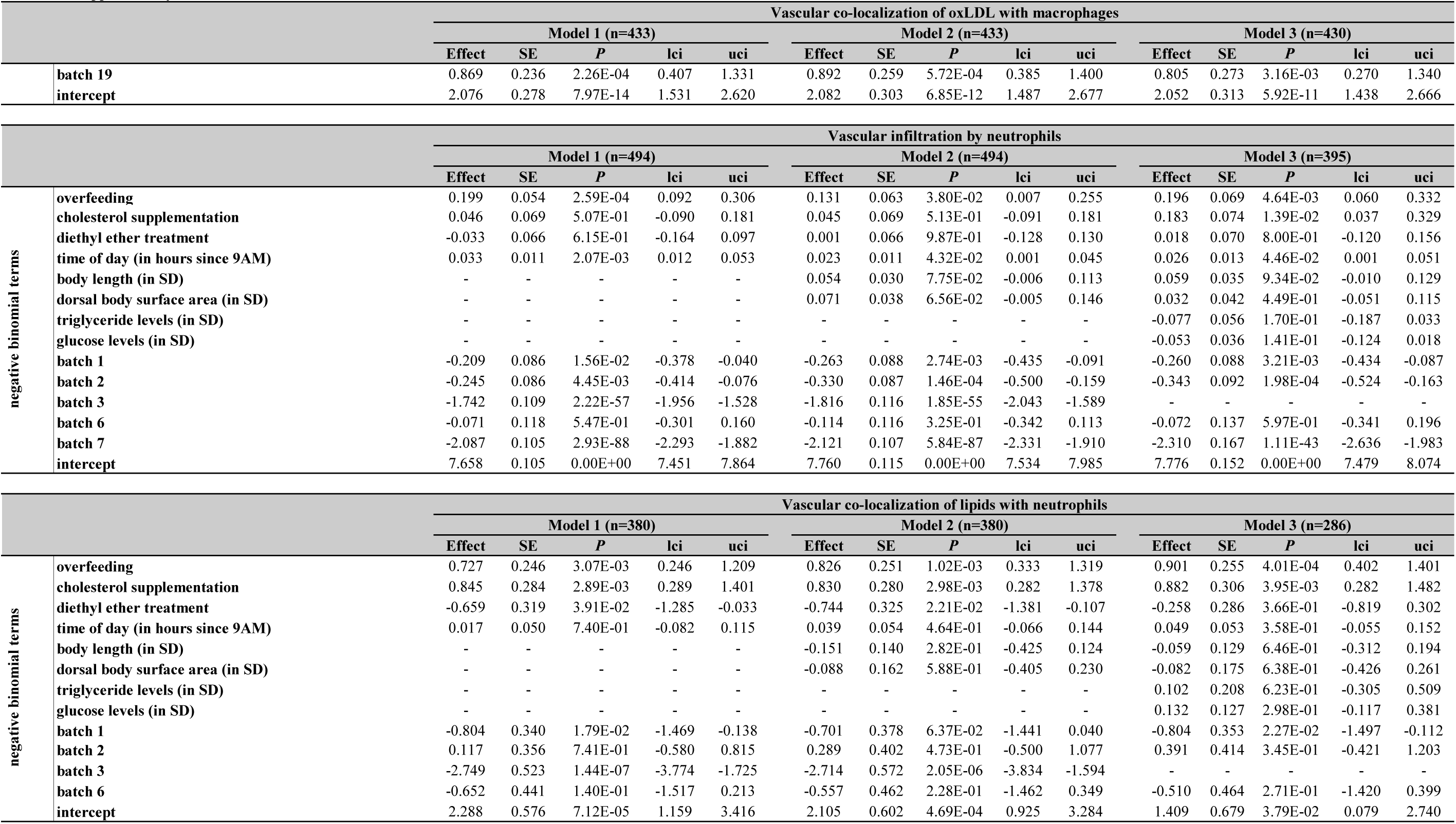
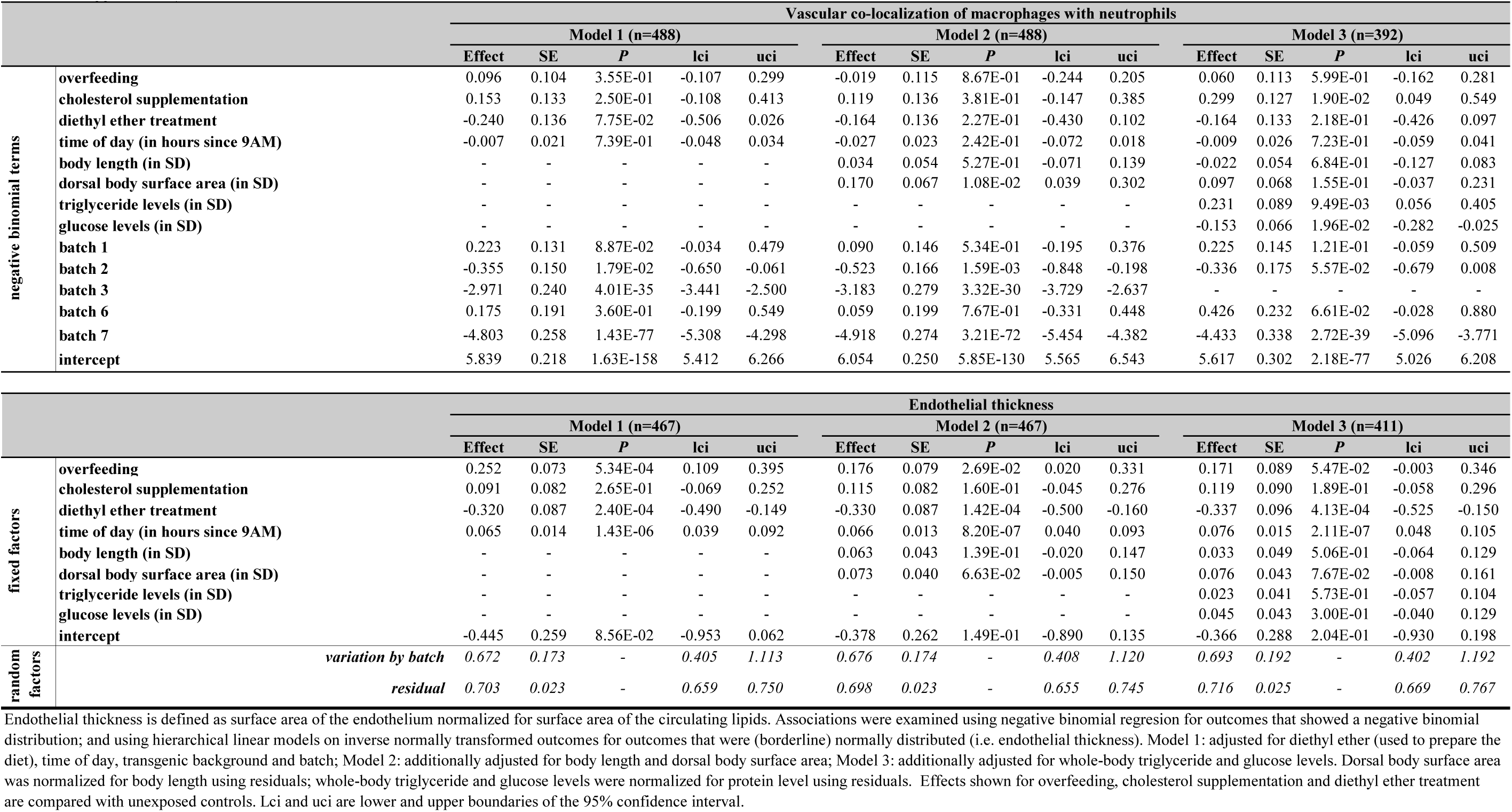
The effect of overfeeding and cholesterol supplementation on image-based vascular atherogenic traits

**Supplementary Table 6.**
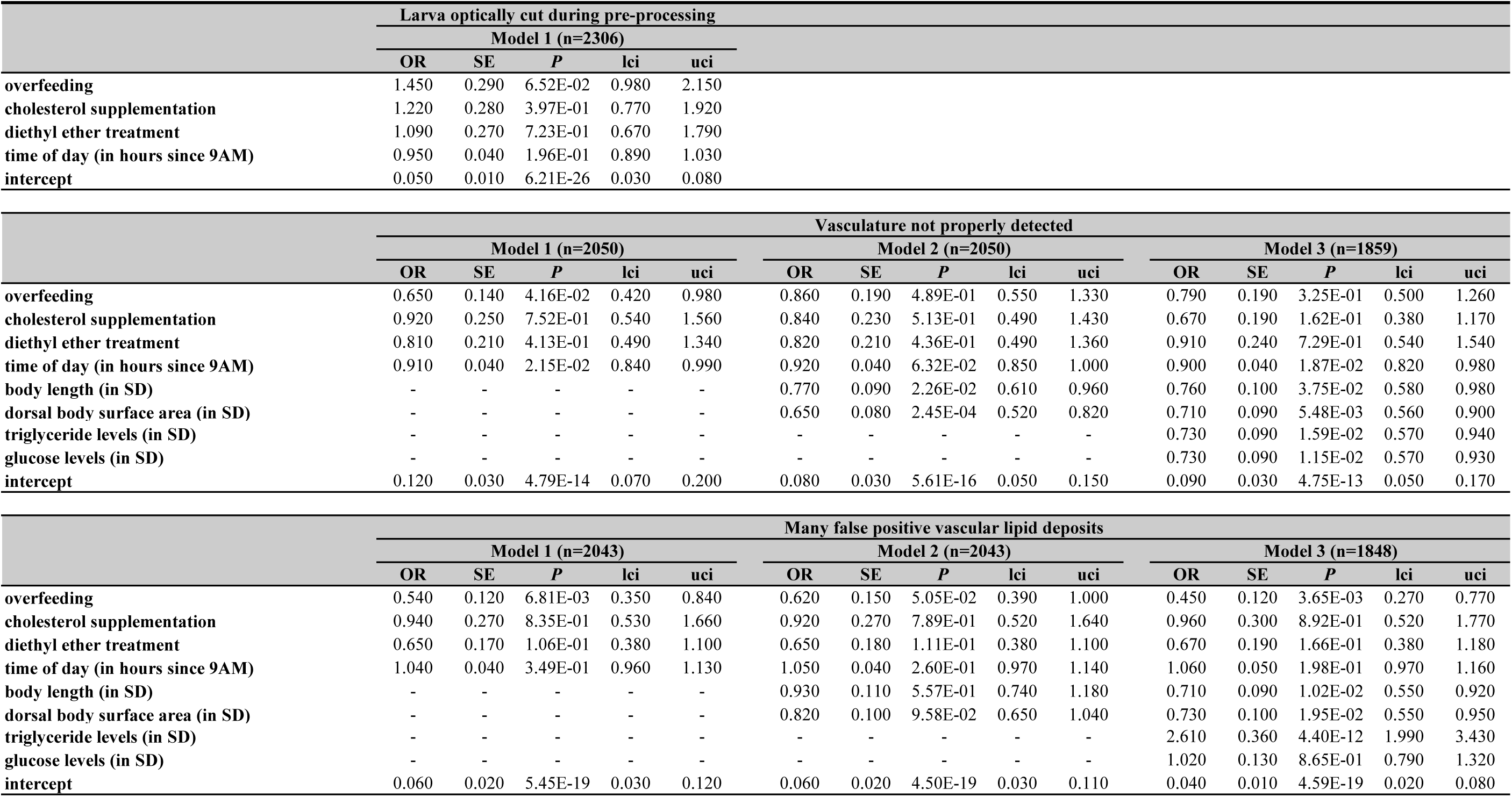
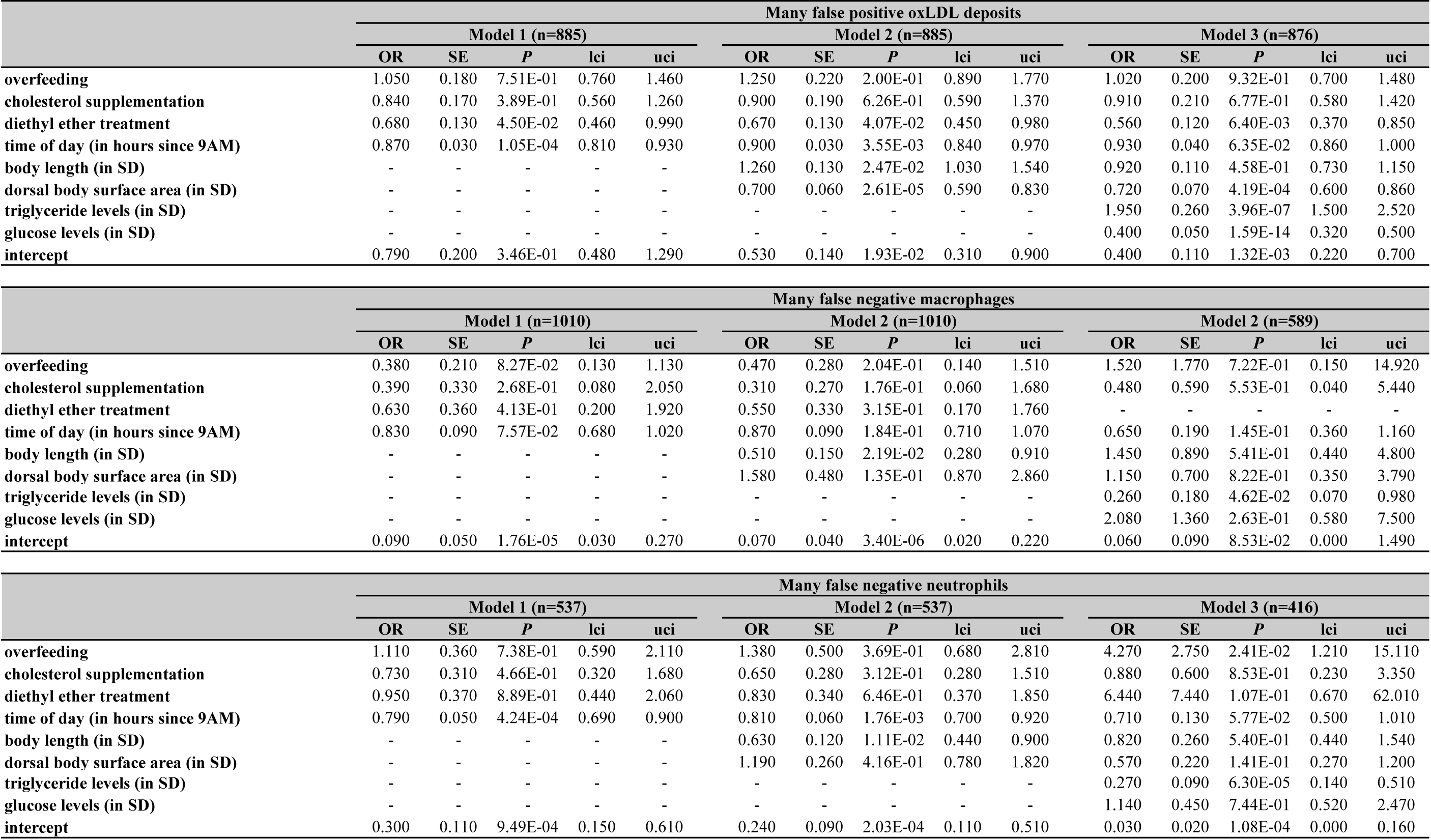
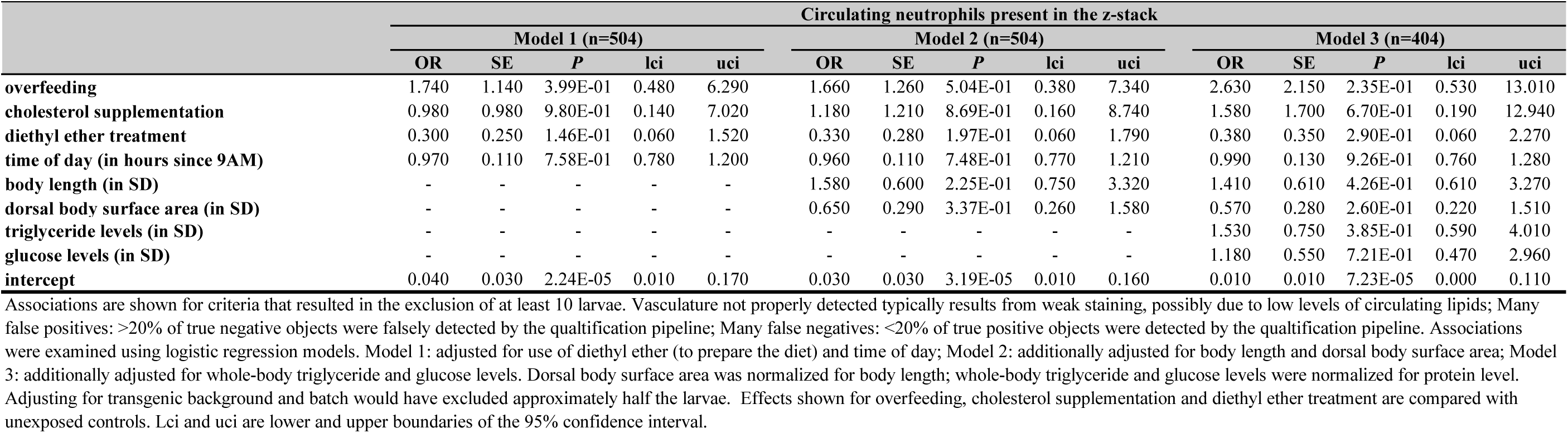
The effect of overfeeding and cholesterol supplementation on suboptimal image or quantification quality

**Supplementary Table 7.**
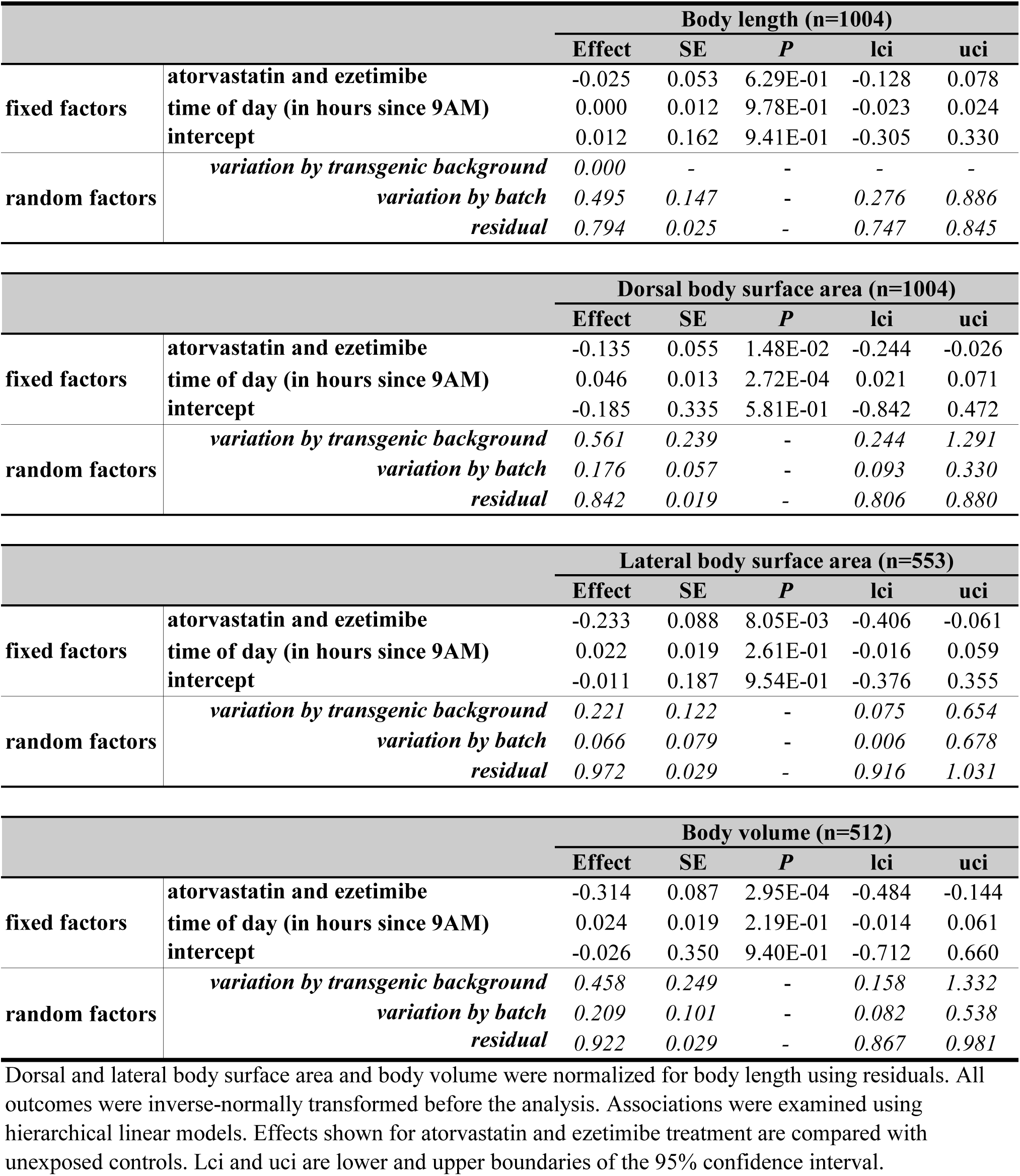
The effect of treatment with atorvastatin and ezetimibe on body size

**Supplementary Table 8.**
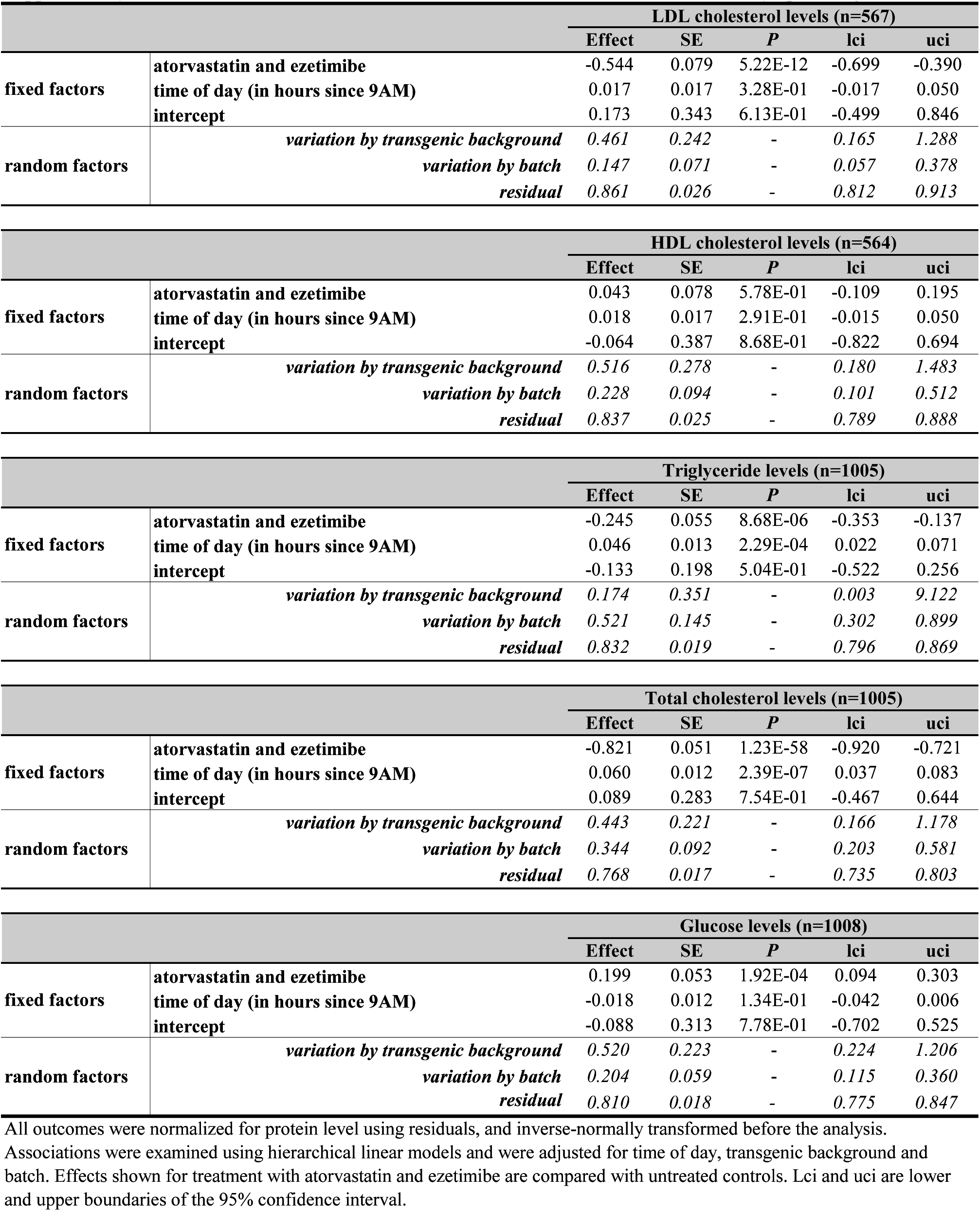
The effect of treatment with atorvastatin and ezetimibe on whole-body lipid and glucose levels

**Supplementary Table 9.**
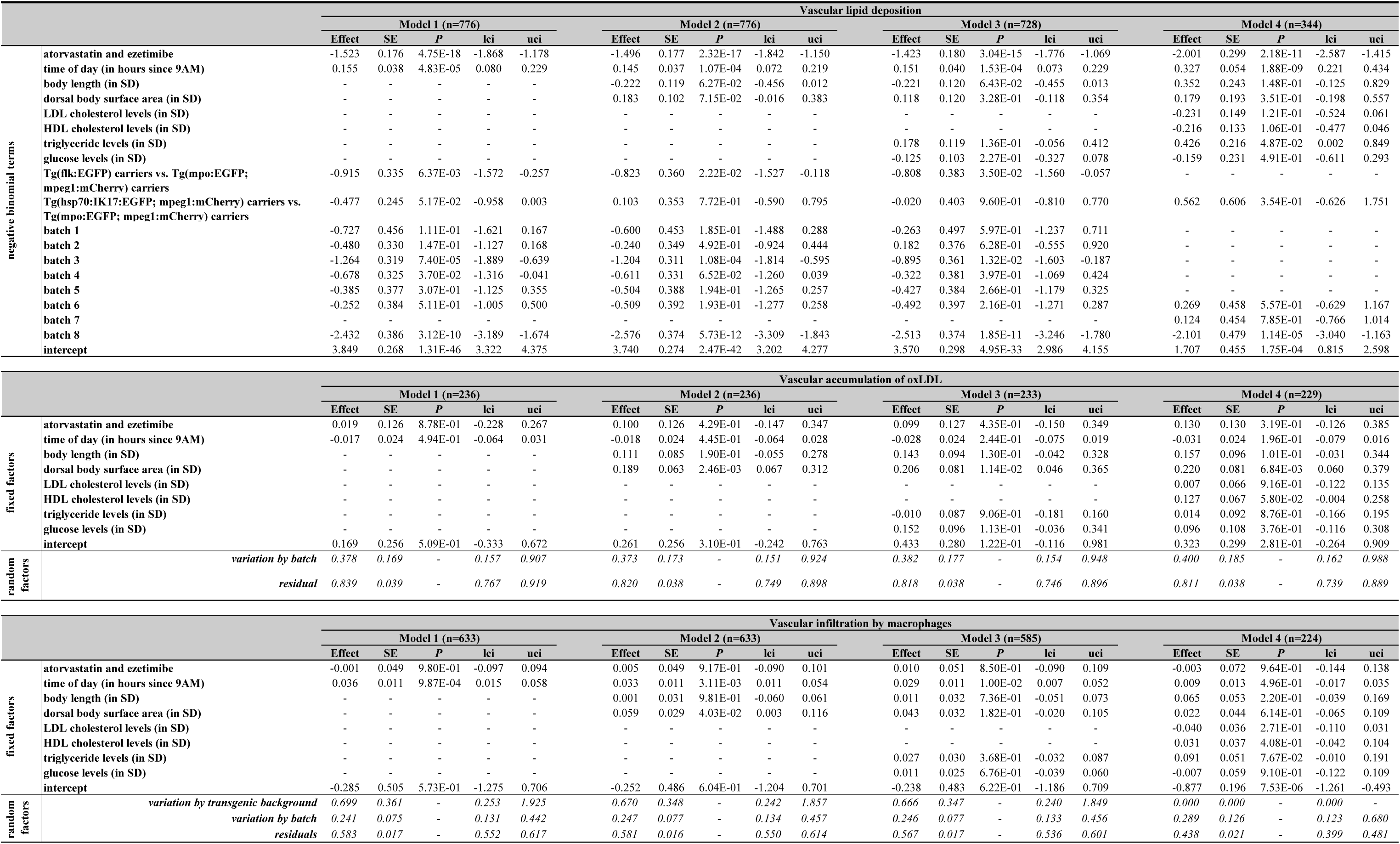
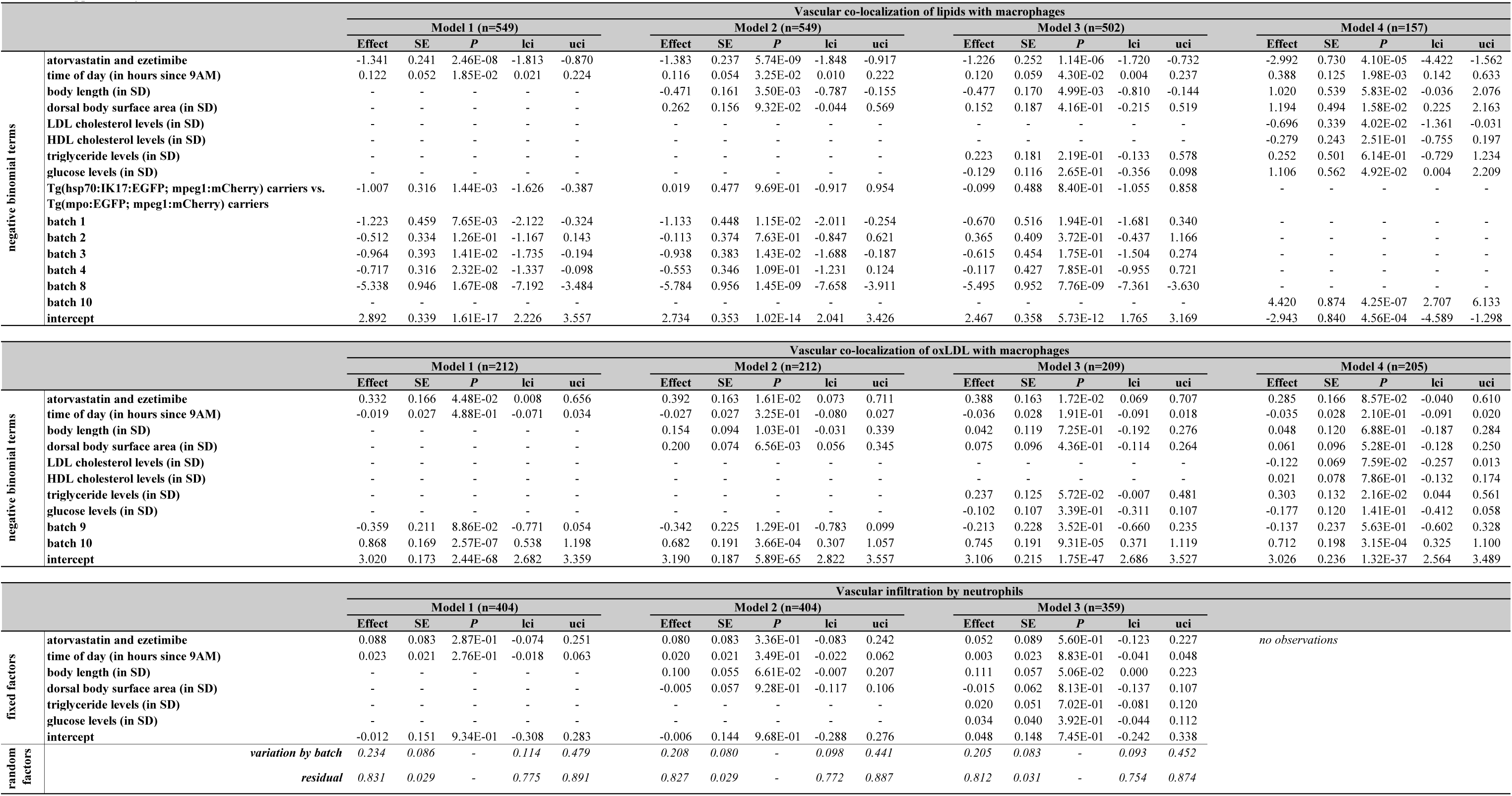
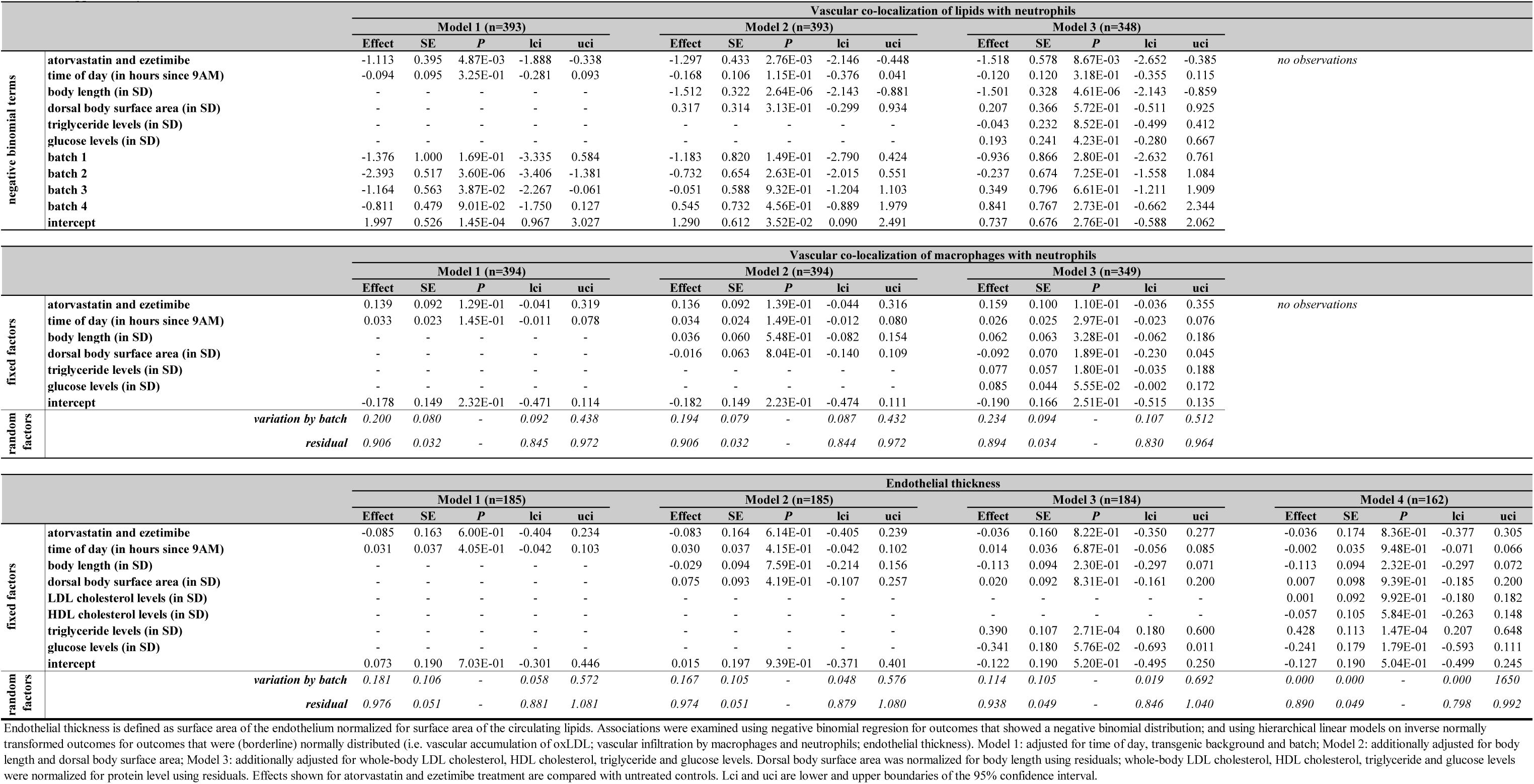
The effect of atorvastatin and ezetimibe on image-based vascular atherogenic traits

**Supplementary Table 10.**
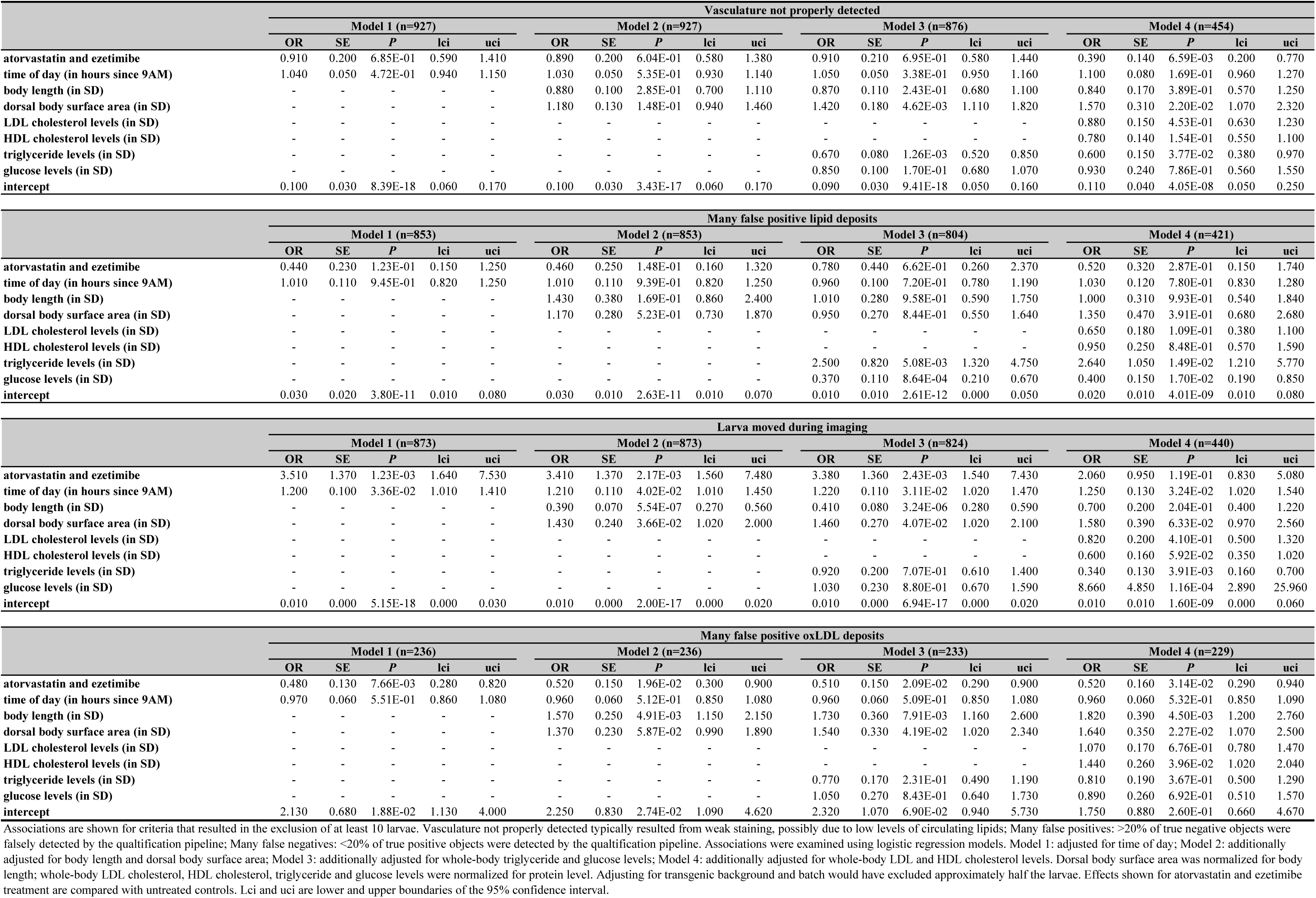
The effect of treatment with atorvastatin and ezetimibe on suboptimal image or image quantification quality

**Supplementary Table 11.**
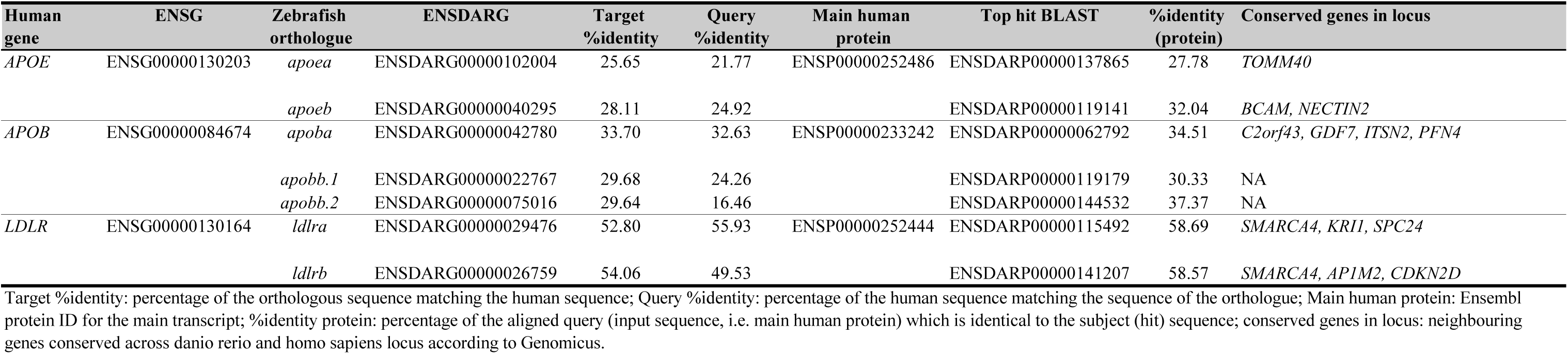
Orthologues of proof-of-concept genes for dyslipidemia, atherosclerosis and coronary artery disease

**Supplementary Table 12.**
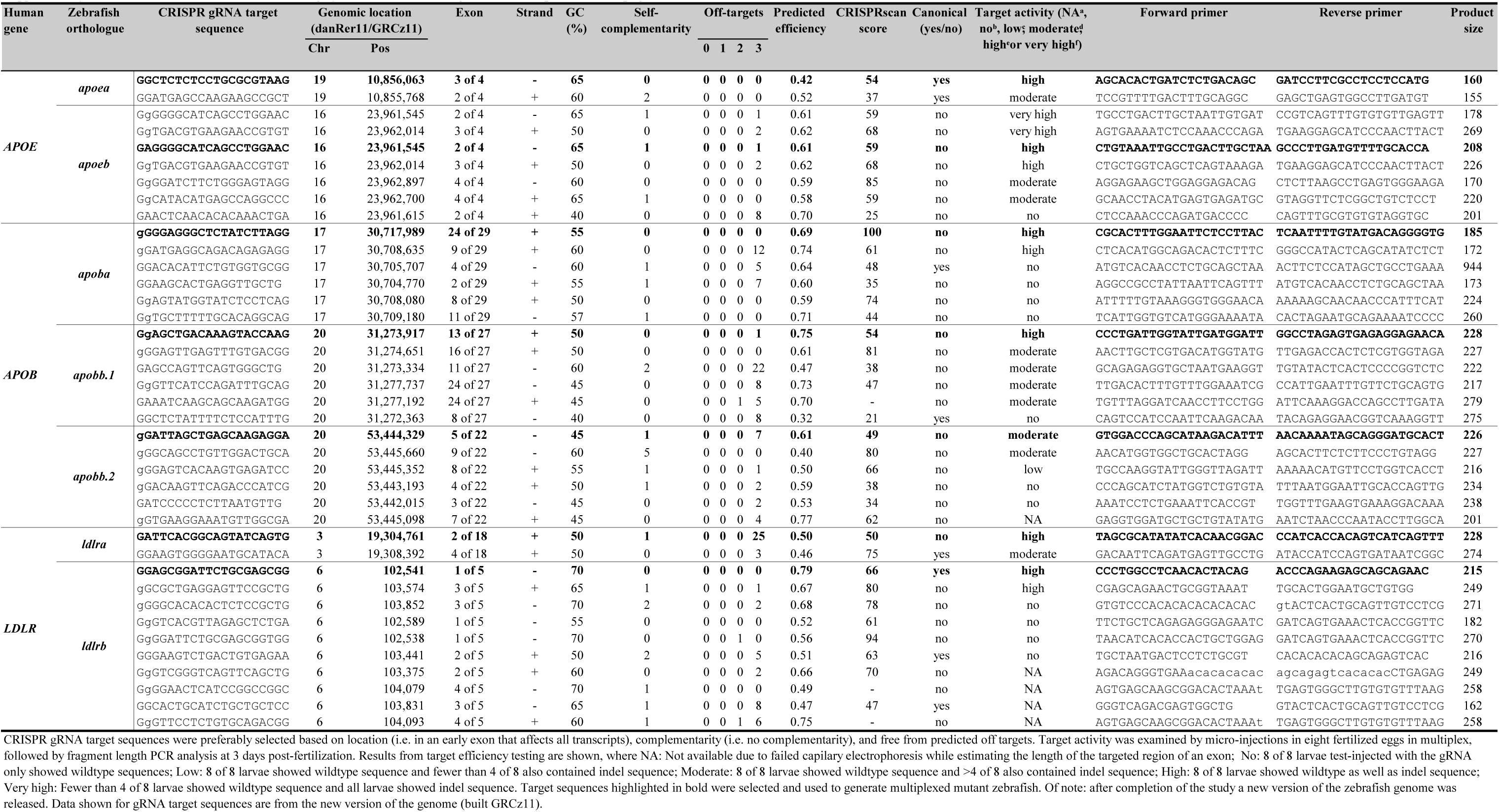
Identification of moderate-to-highly active CRISPR-Cas9 guide RNAs for proof-of-concept genes

**Supplementary Table 13.**
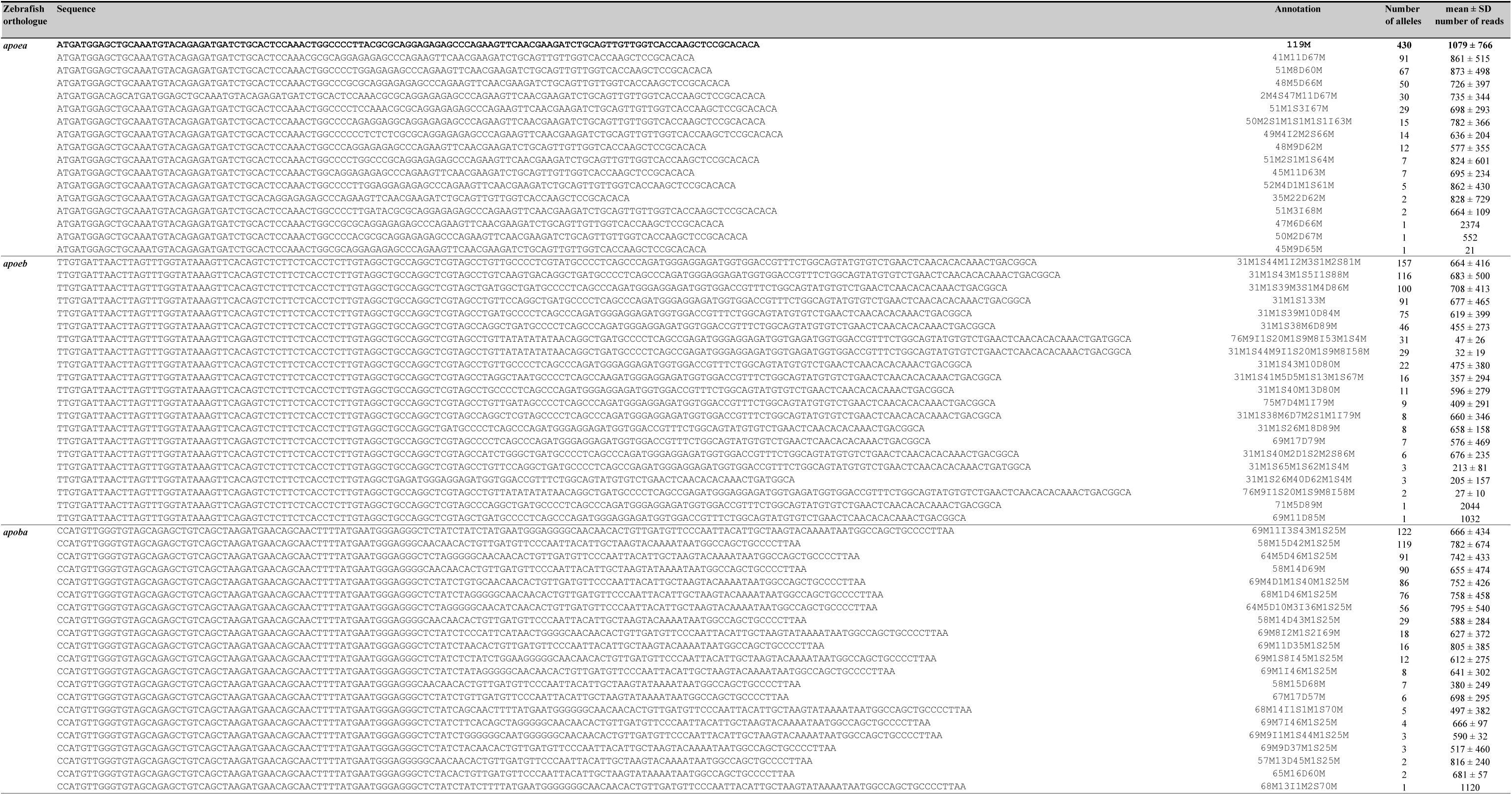
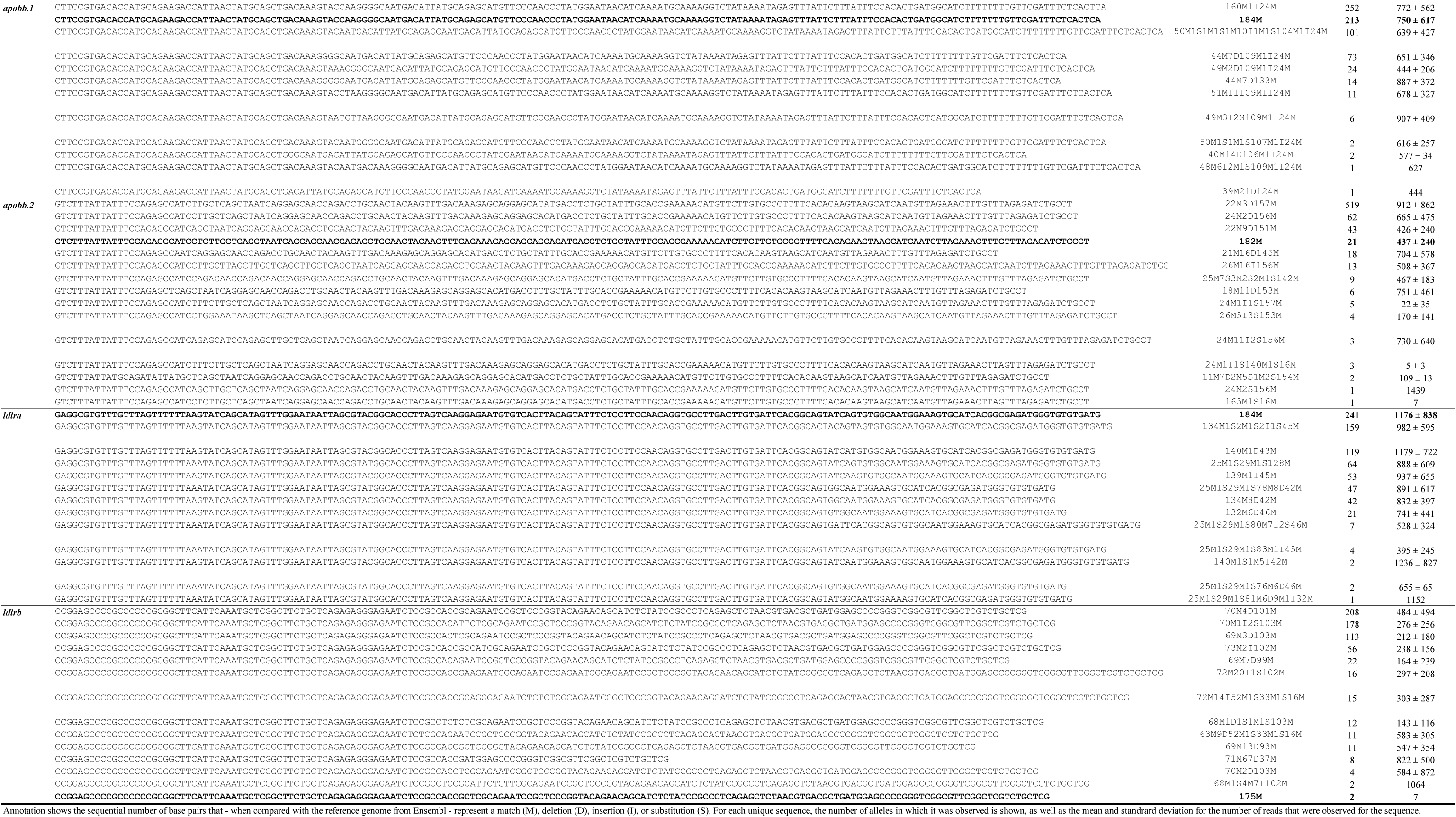
Unique CRISPR-Cas9-induced mutations for orthologues of proof-of-concept genes

**Supplementary Table 14.**
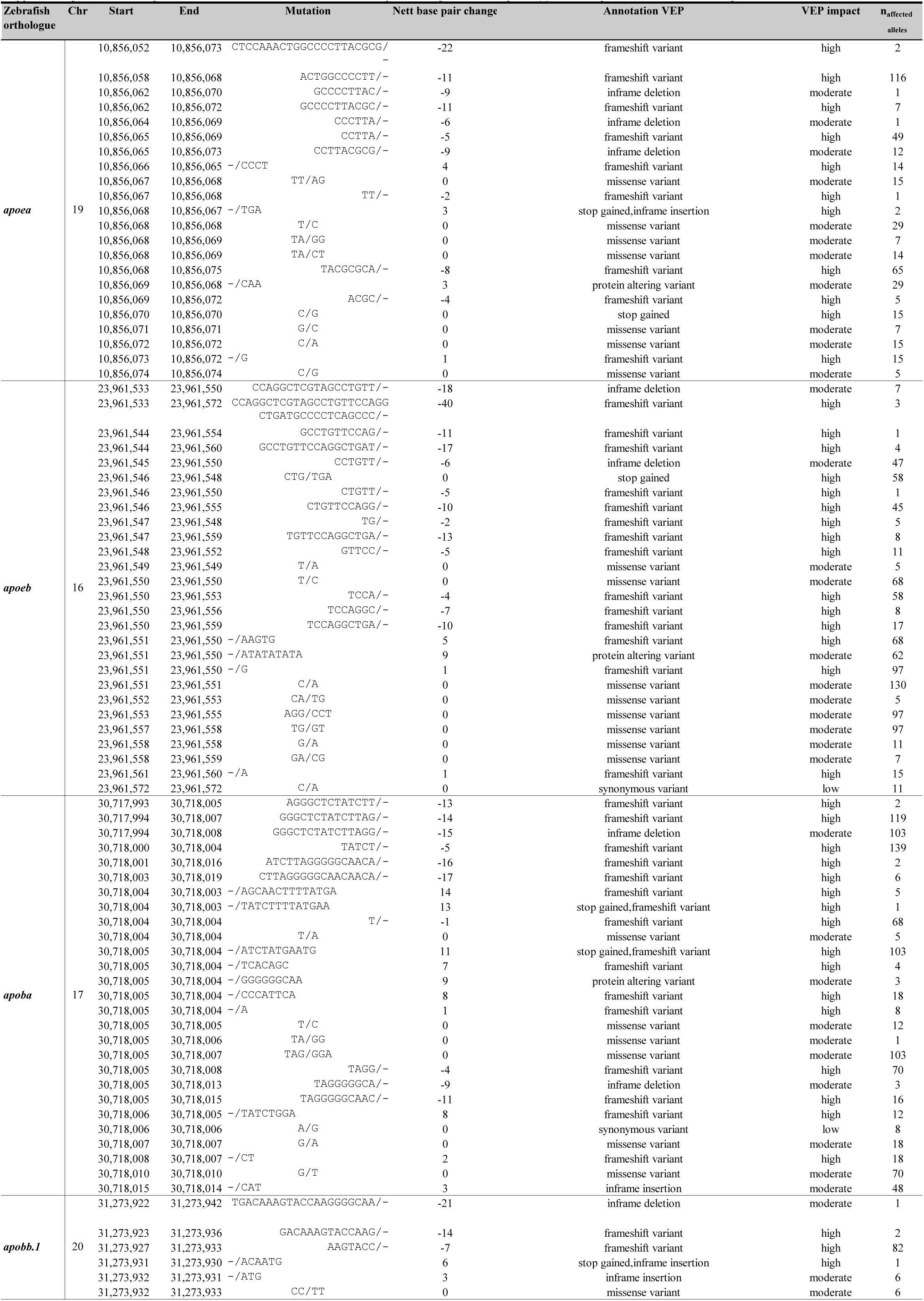
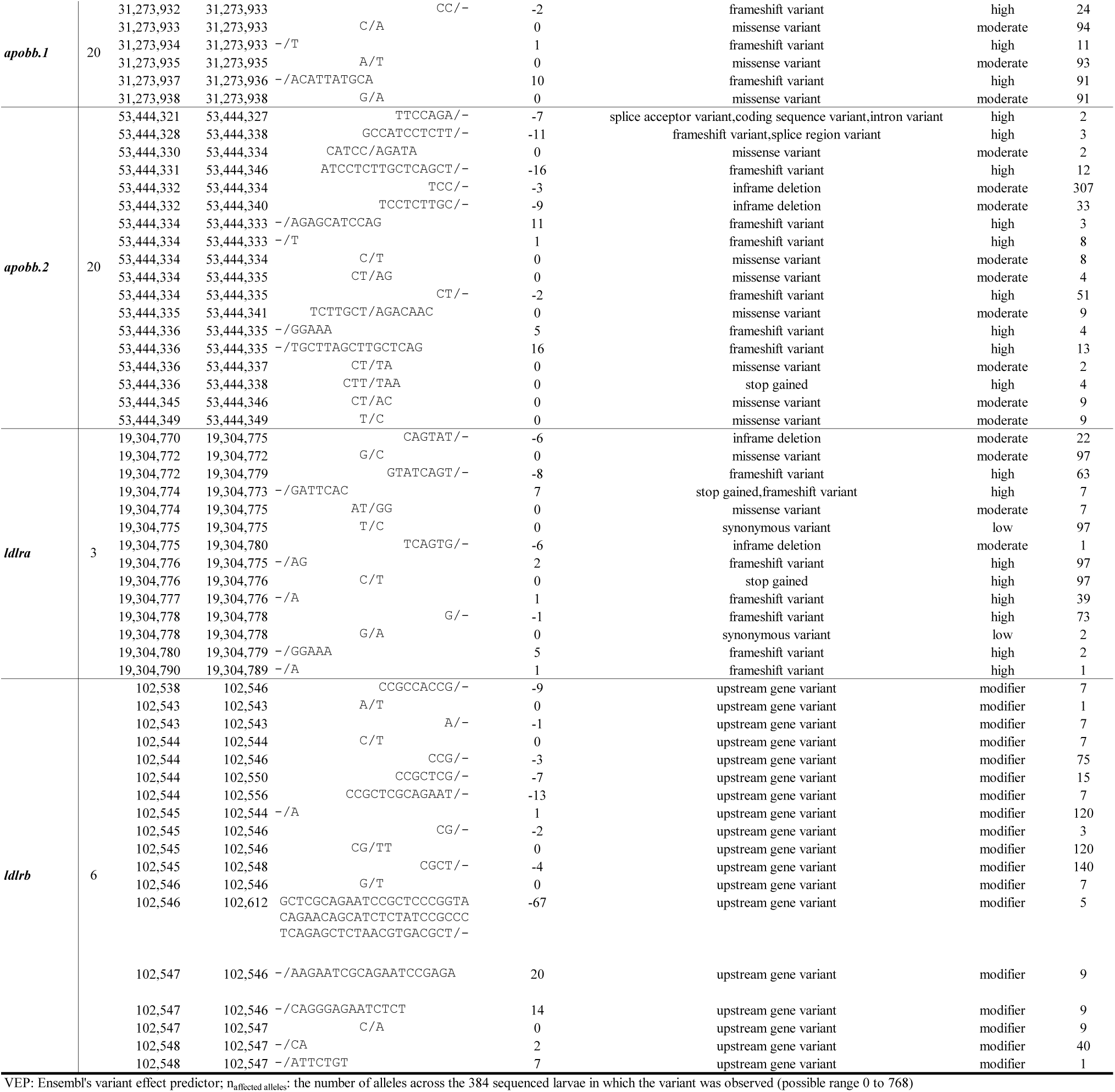
Unique CRISPR-Cas9-induced variants in the most prominently observed sequence(s) and their predicted functional consequences

**Supplementary Table 15.**
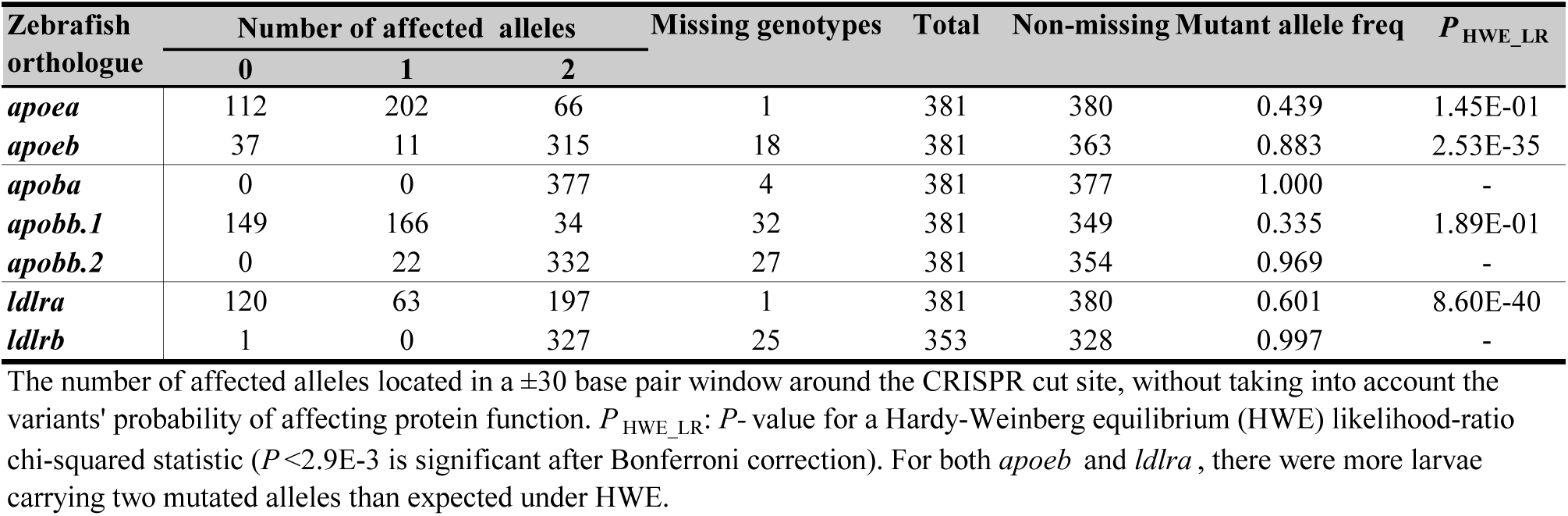
Sequencing results expressed in number of mutated alleles for proof-of-concept genes

**Supplementary Table 16.**
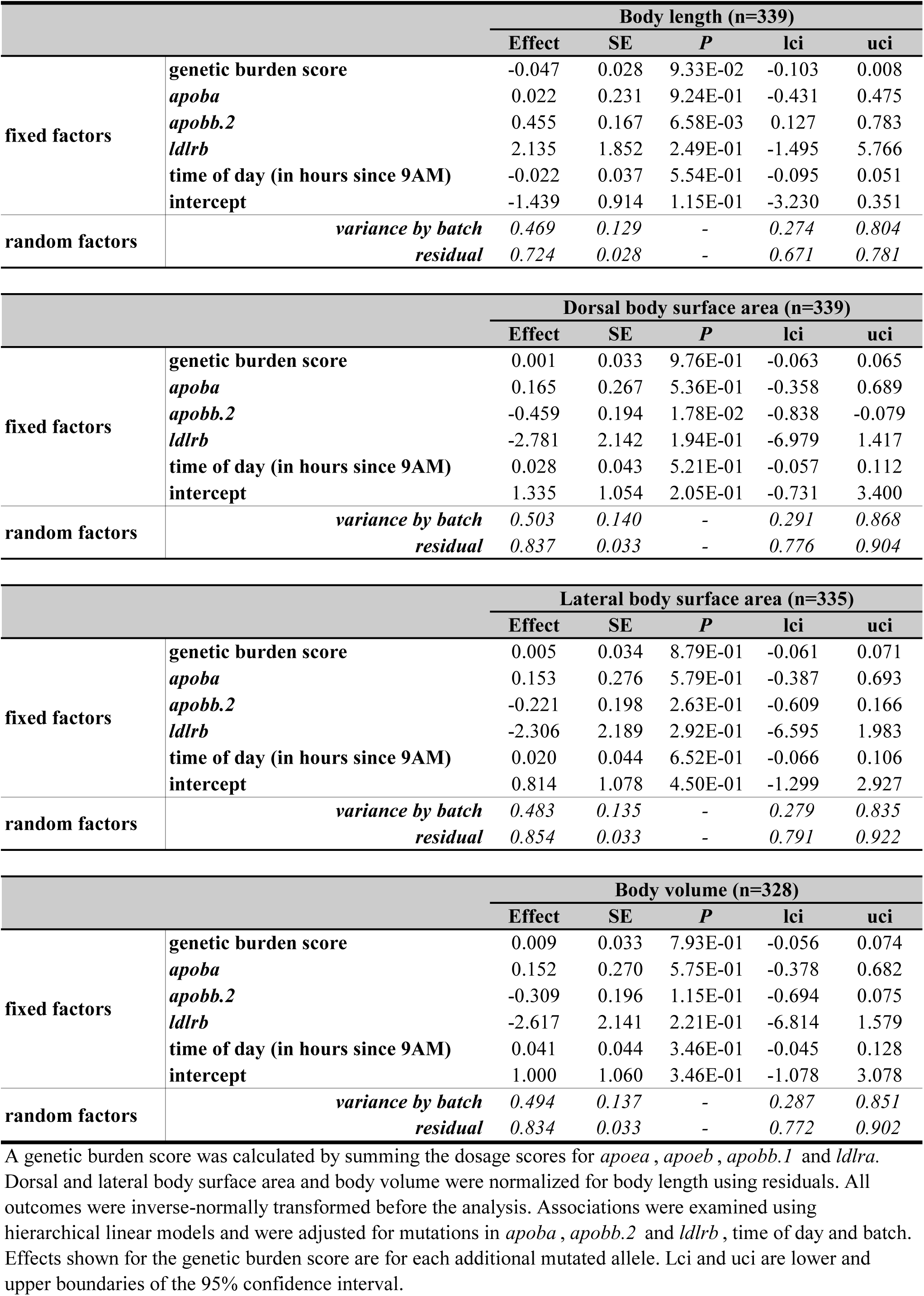
The effect of a genetic burden score on body size

**Supplementary Table 17.**
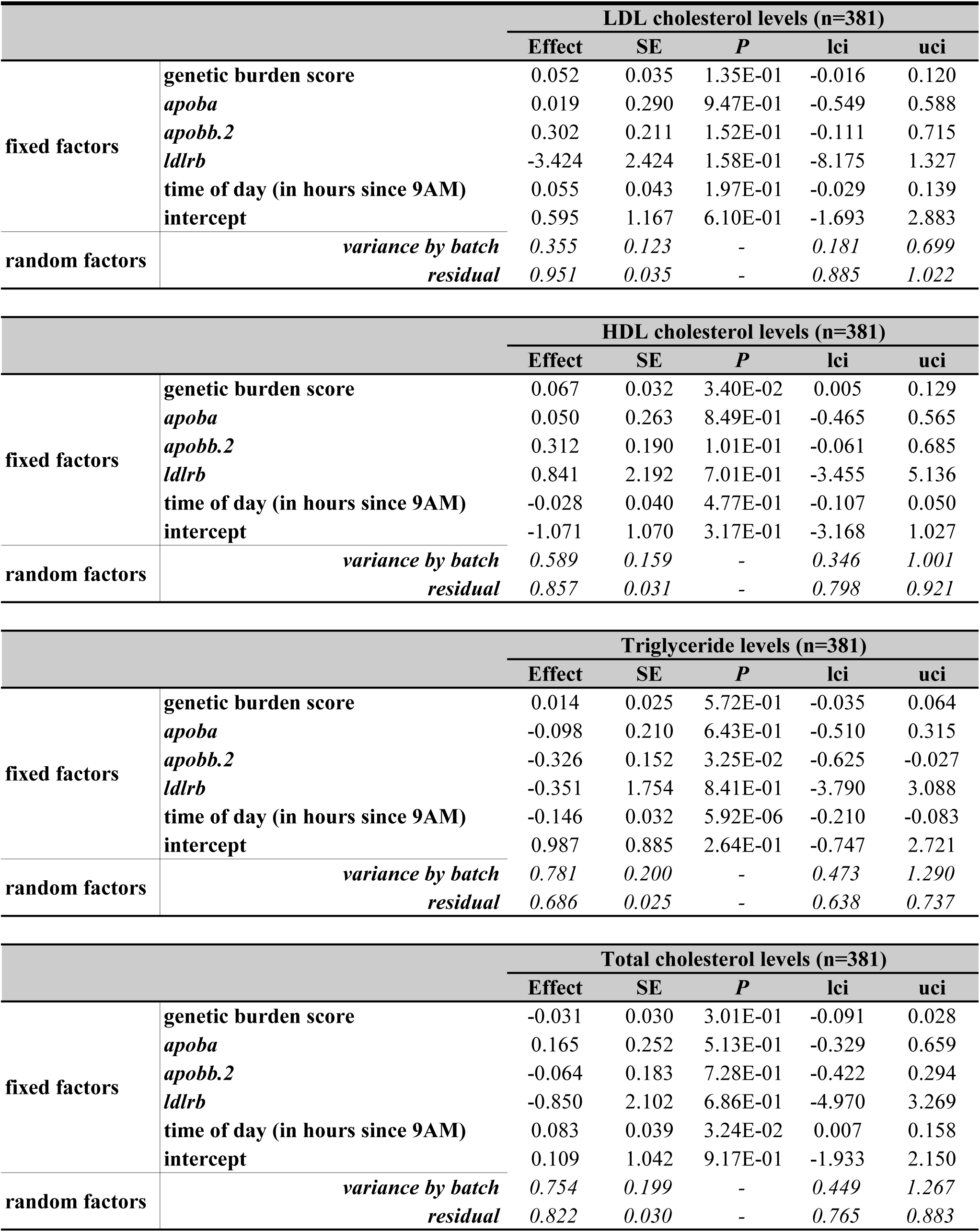
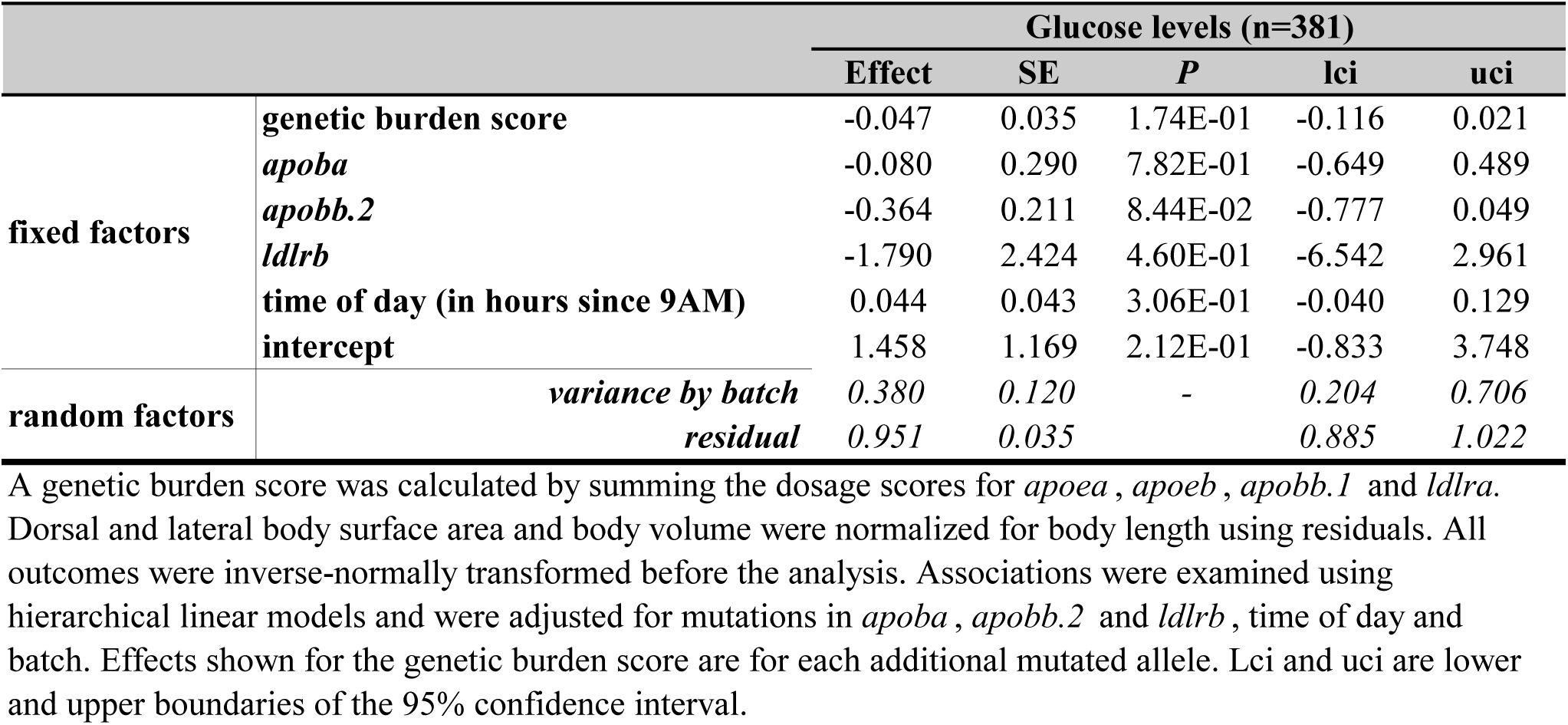
The effect of a genetic burden score on whole-body lipid and glucose levels

**Supplementary Table 18.**
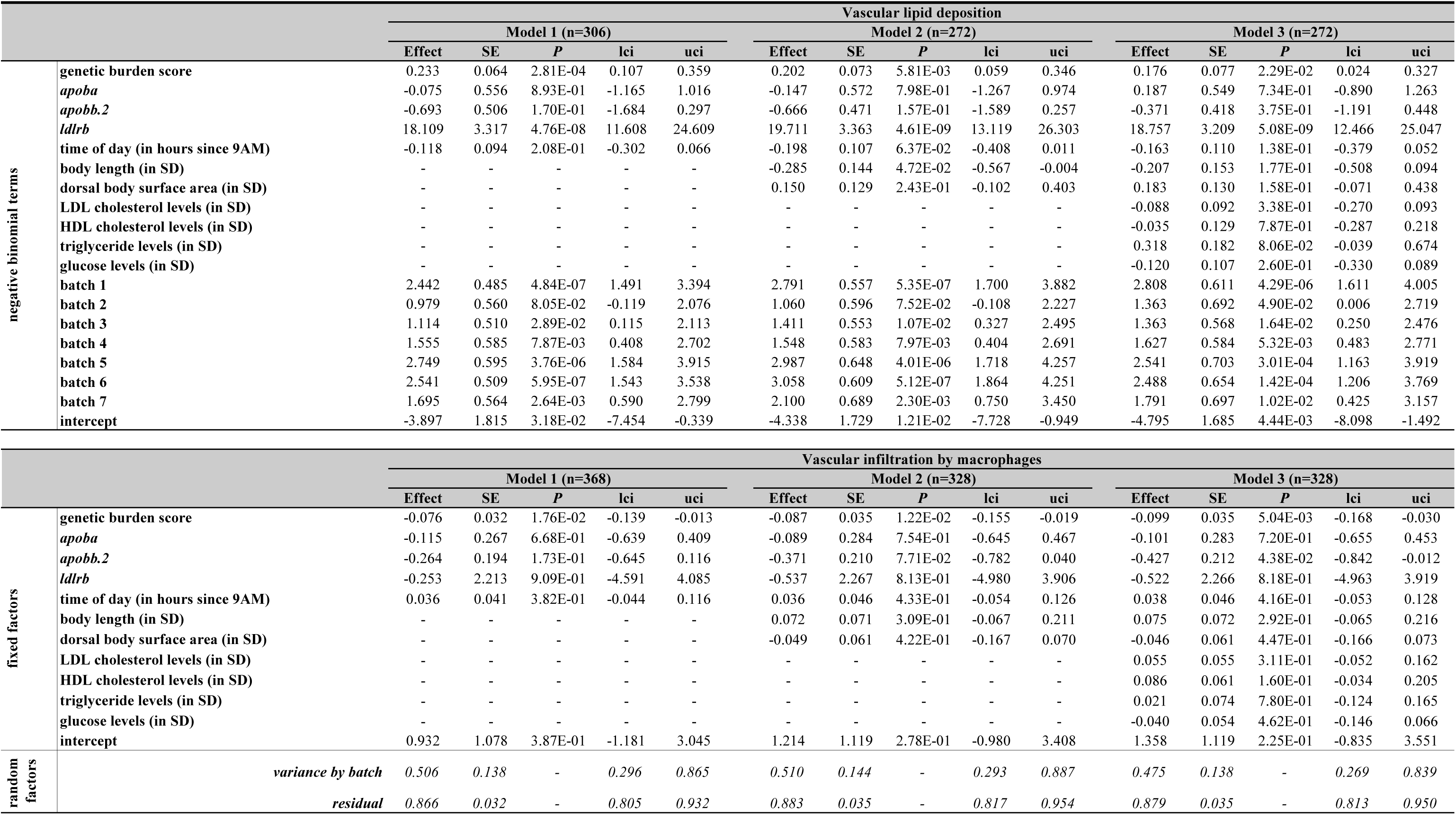
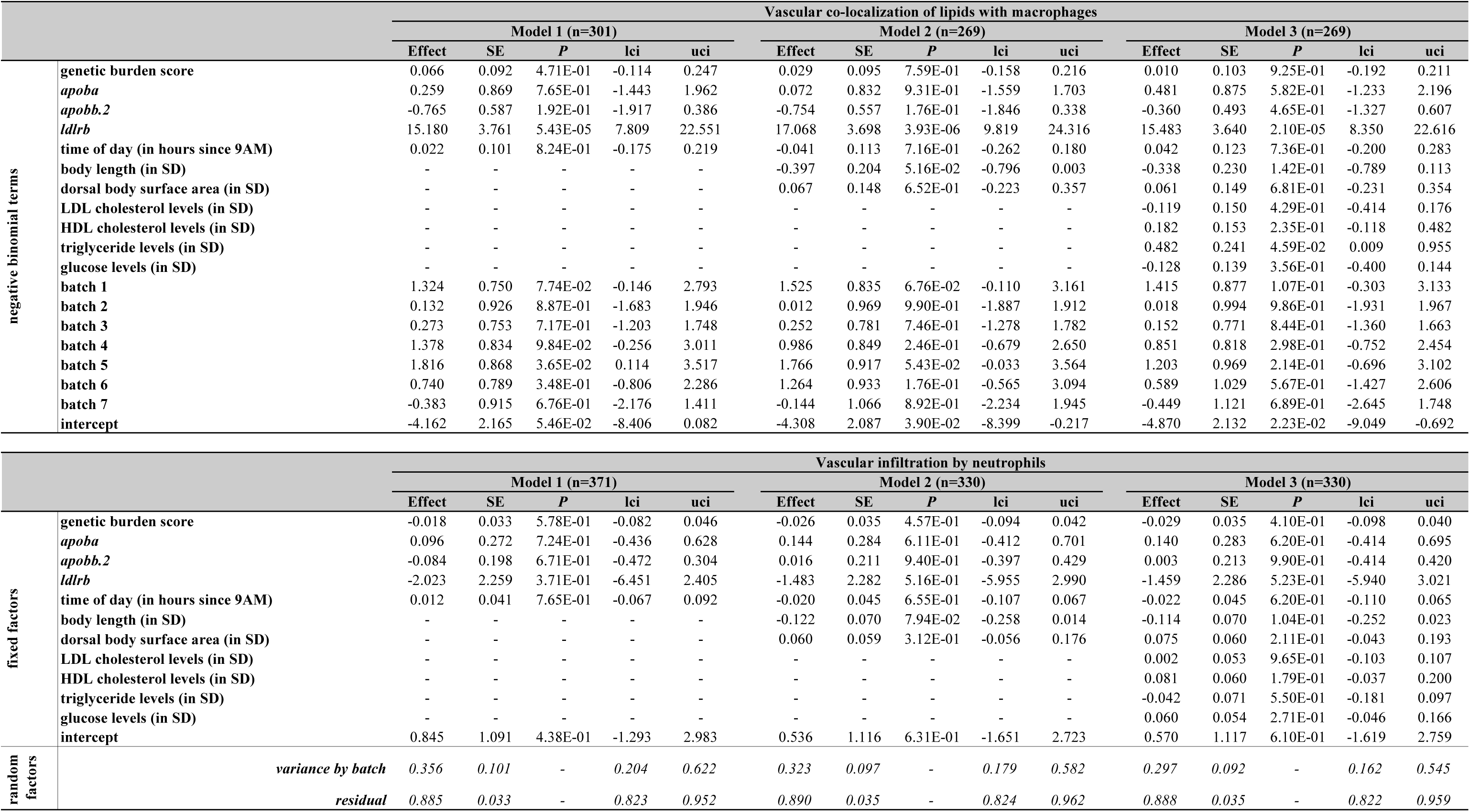
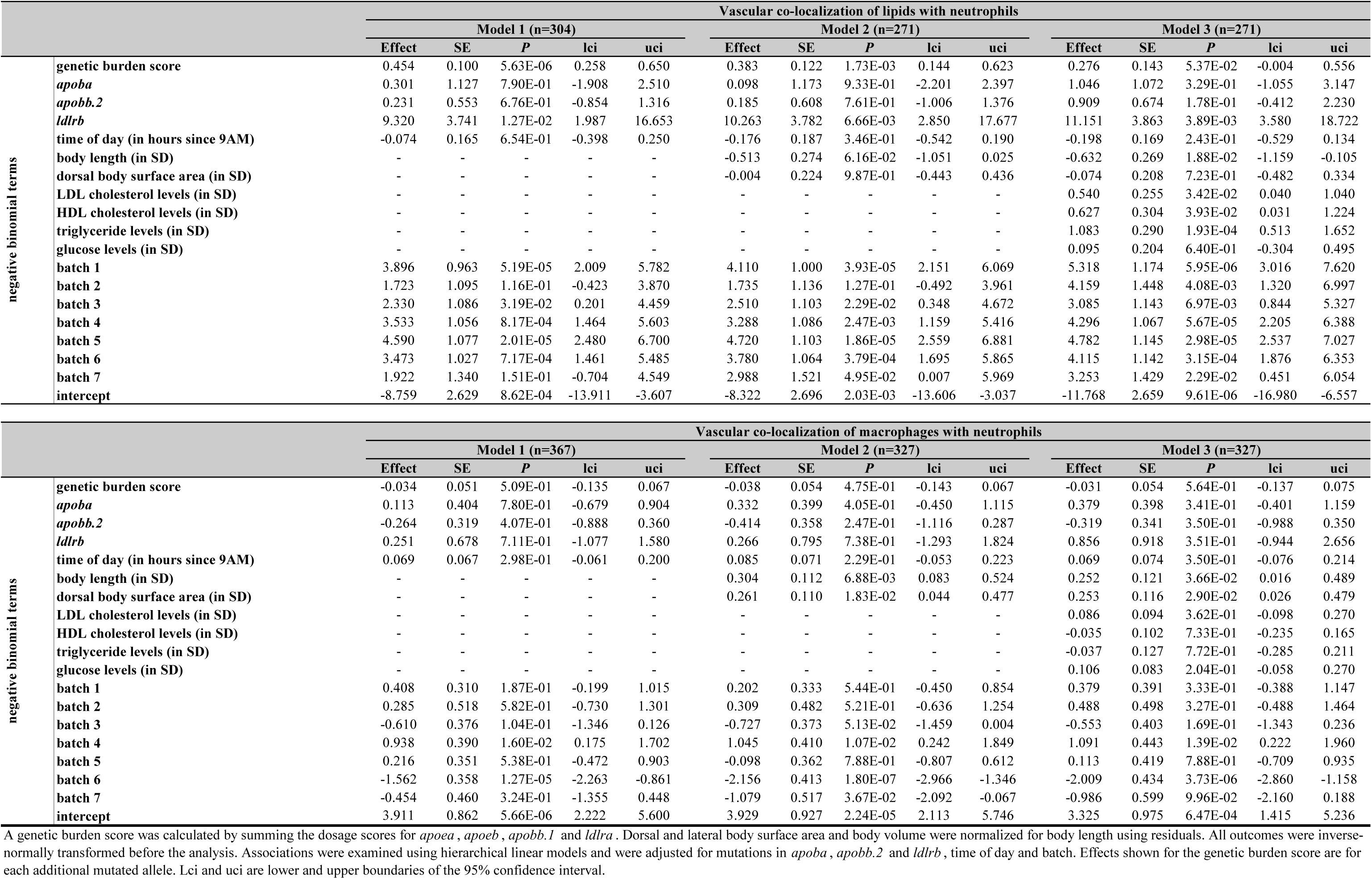
The effect of a genetic burden score on vascular atherogenic traits

**Supplementary Table 19.**
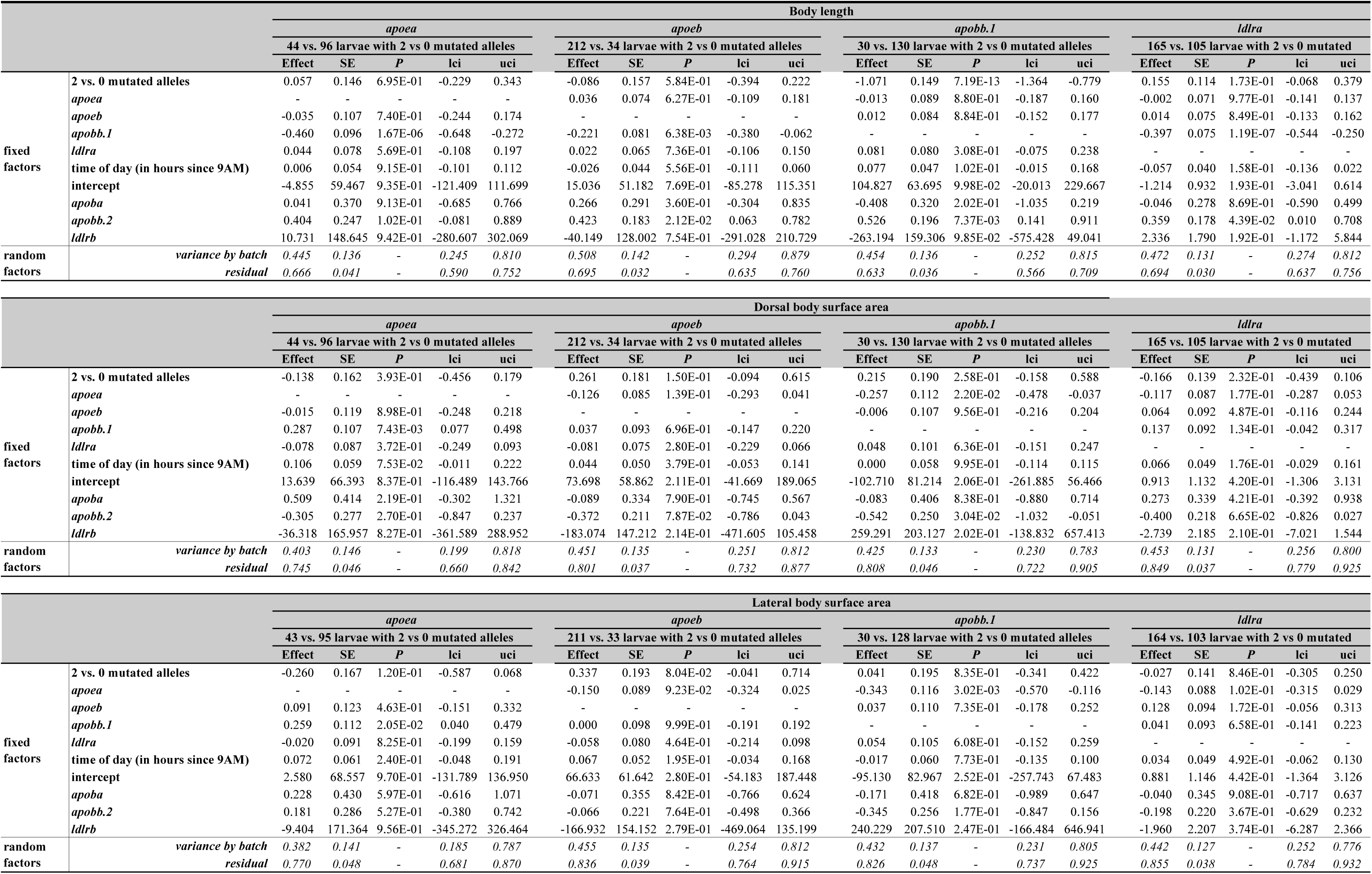
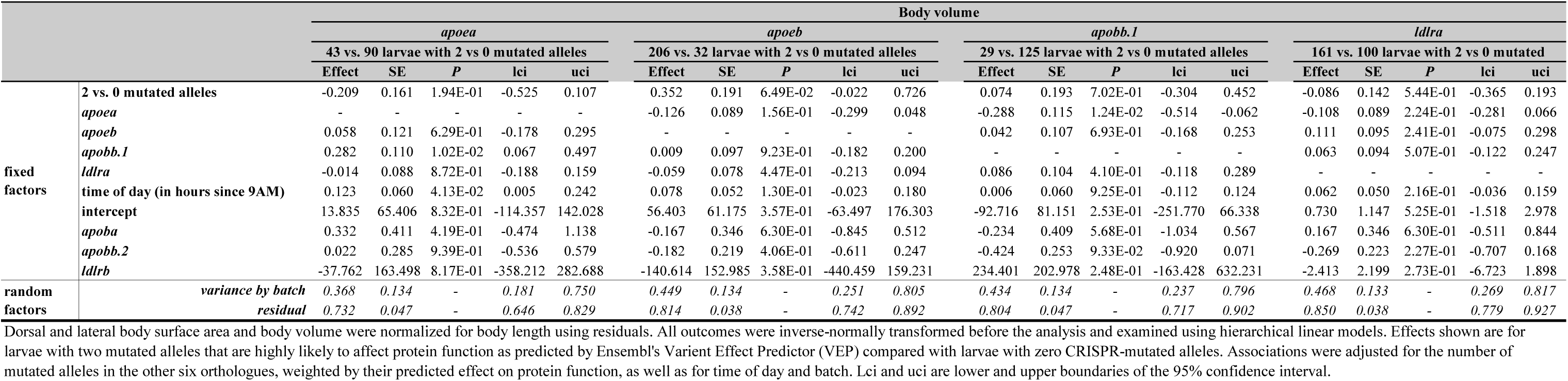
The effect of two vs. zero mutated alleles in apoea, apoeb, apobb.1 or ldlra on body size

**Supplementary Table 20.**
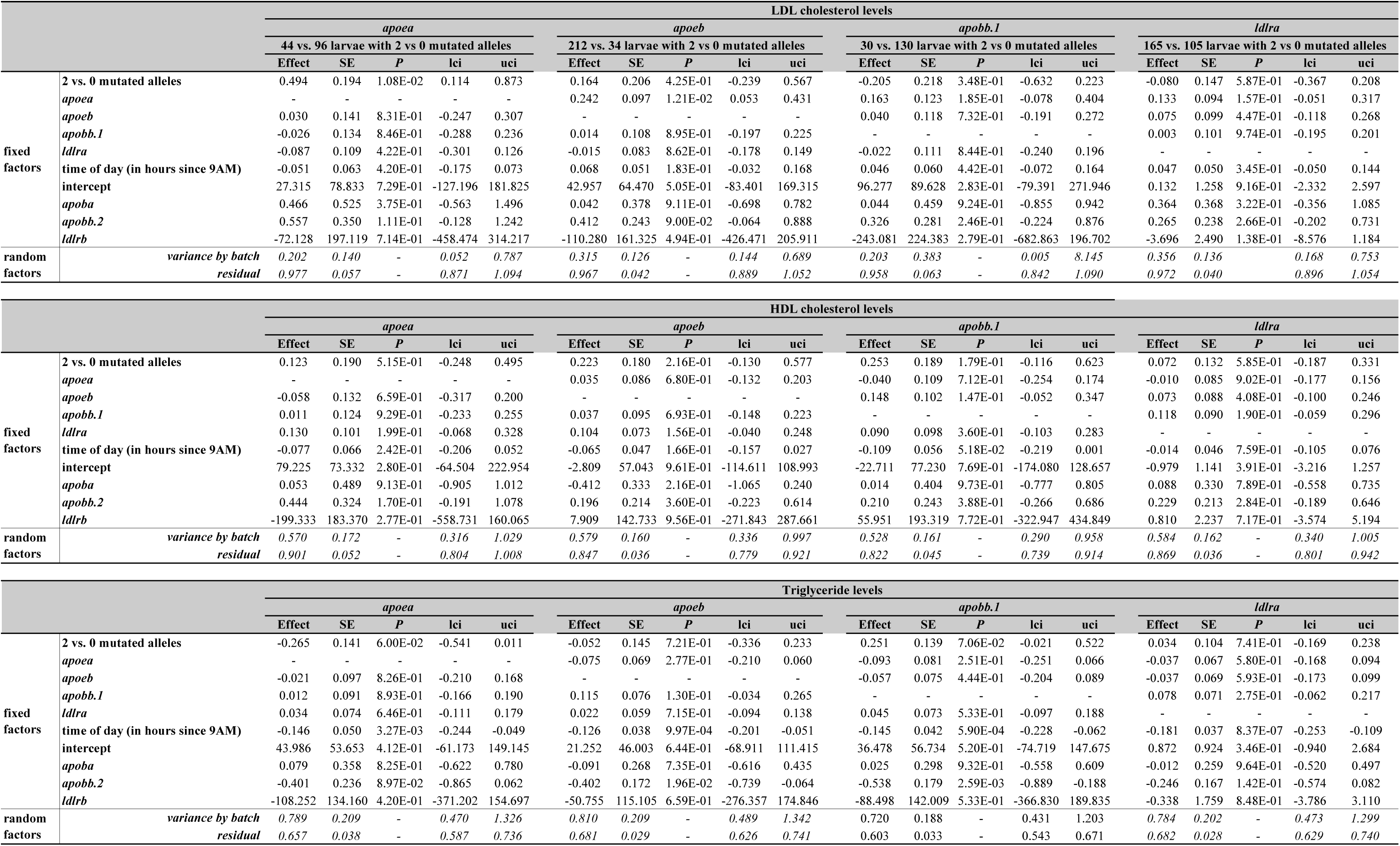
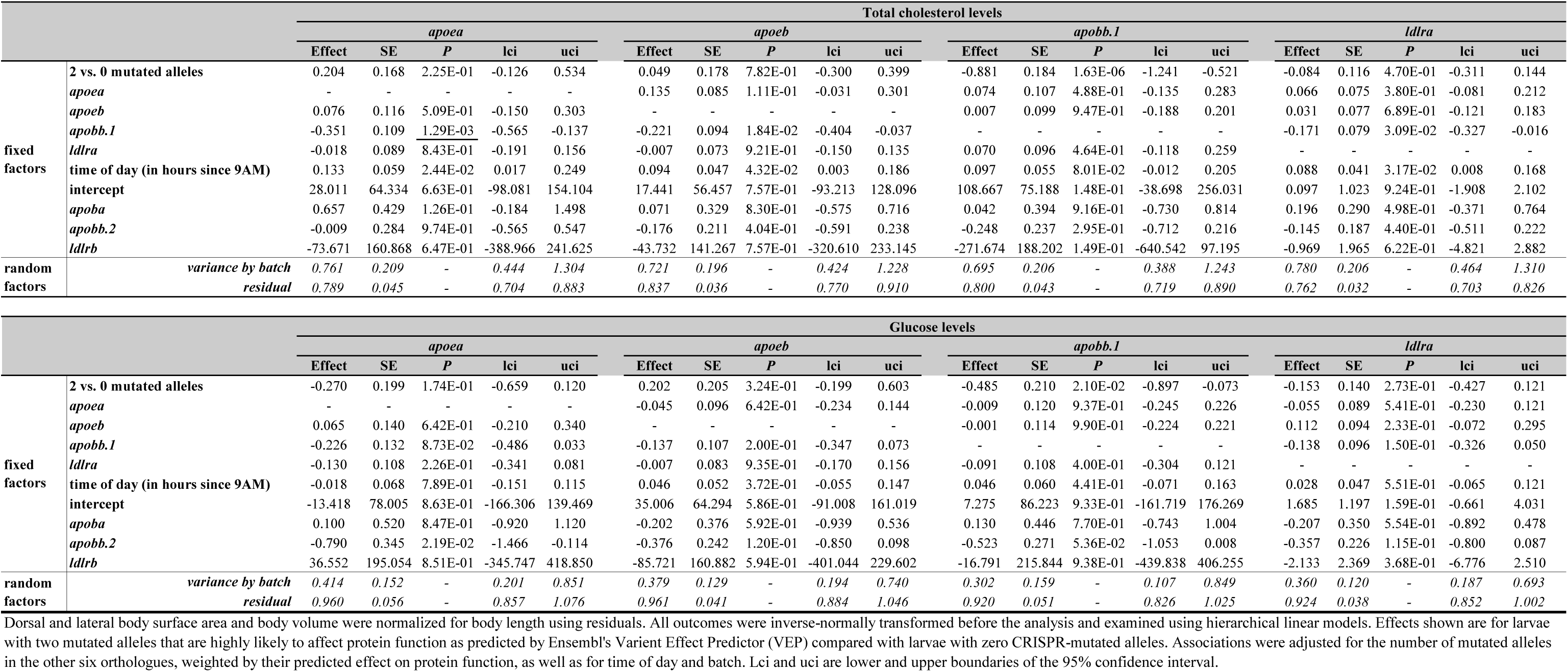
The effect of two vs. zero mutated alleles in *apoea*, *apoeb*, *apobb.1* or *ldlra* on whole-body lipid and glucose levels

**Supplementary Table 21.**
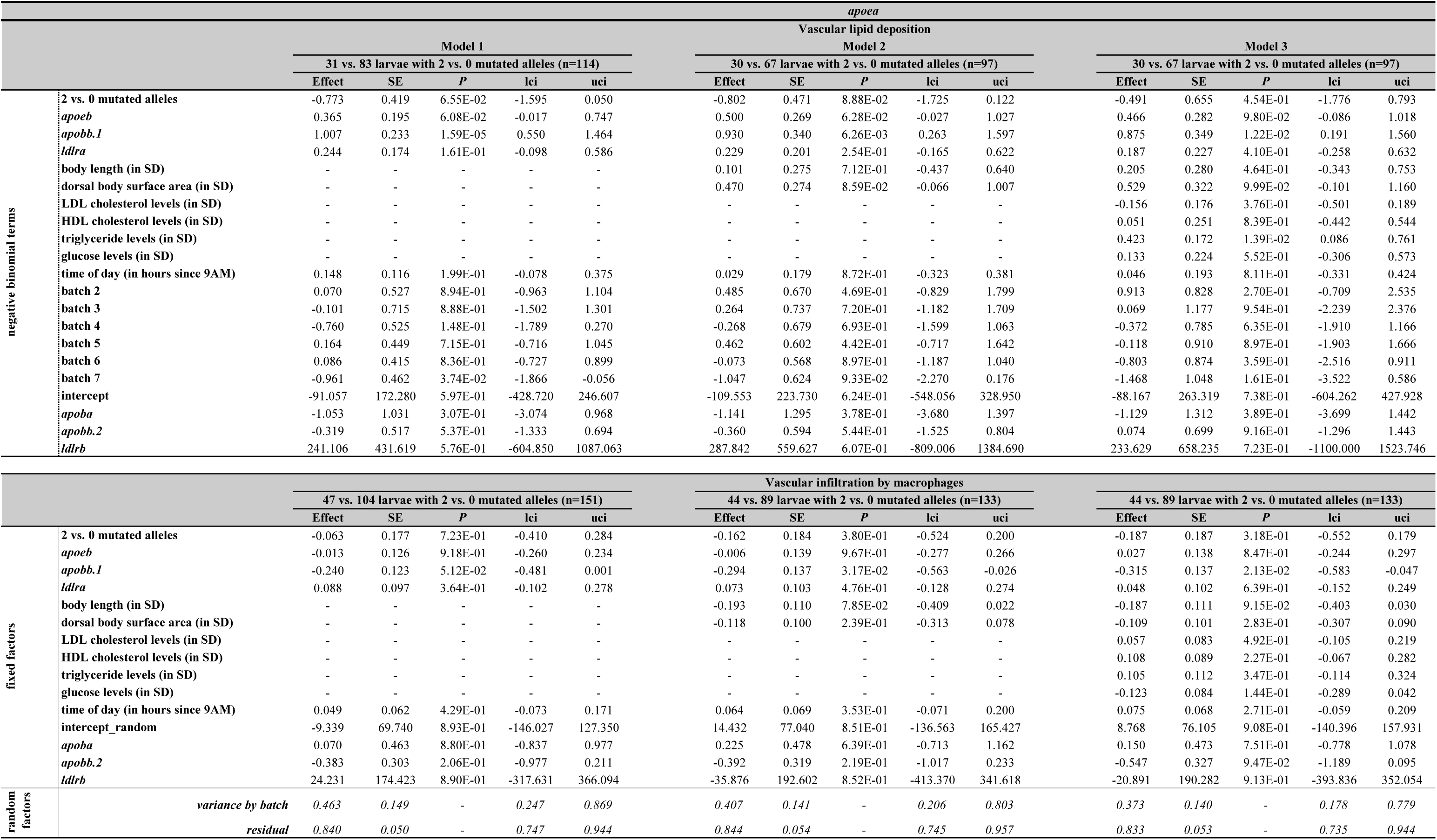
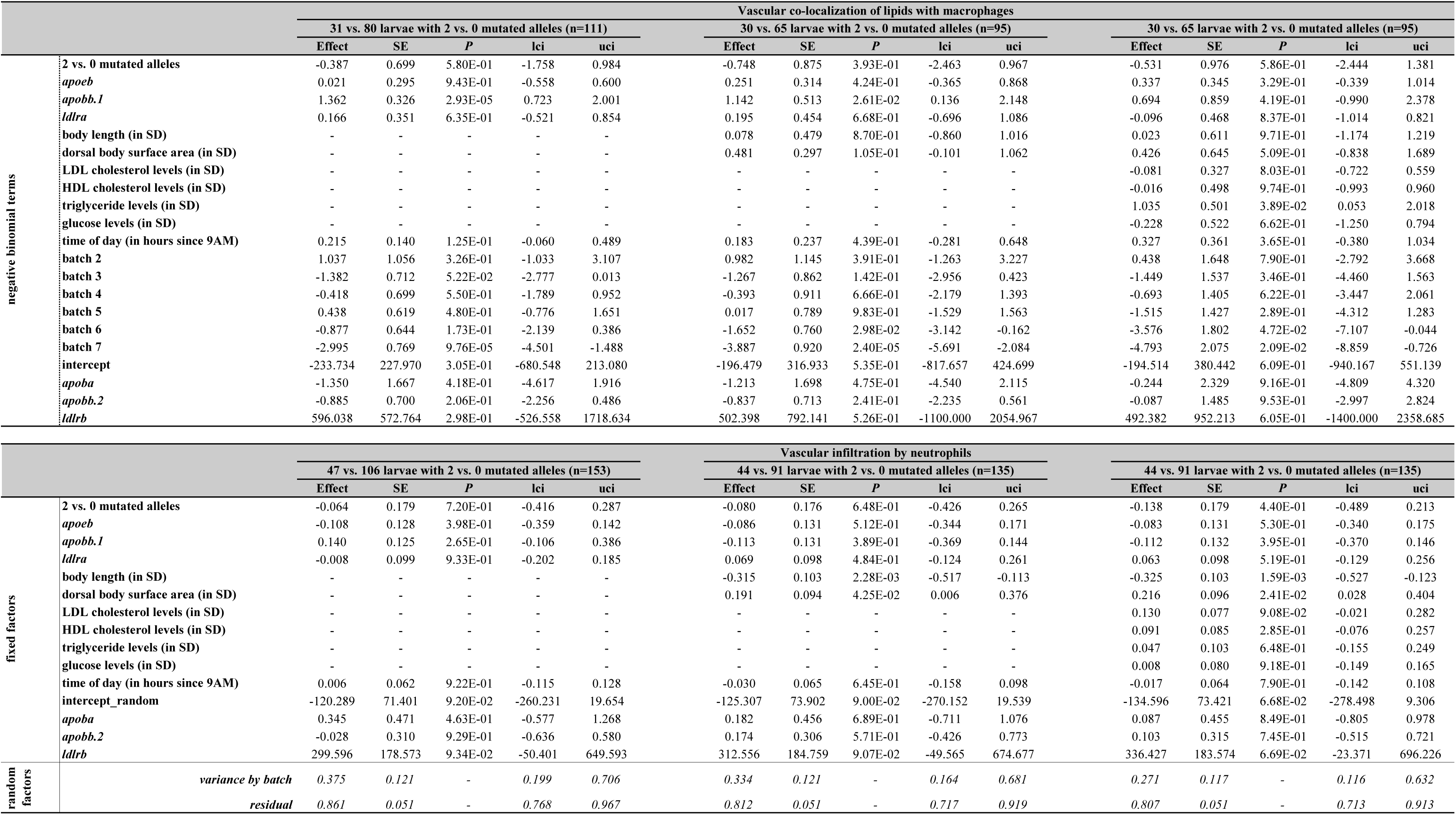
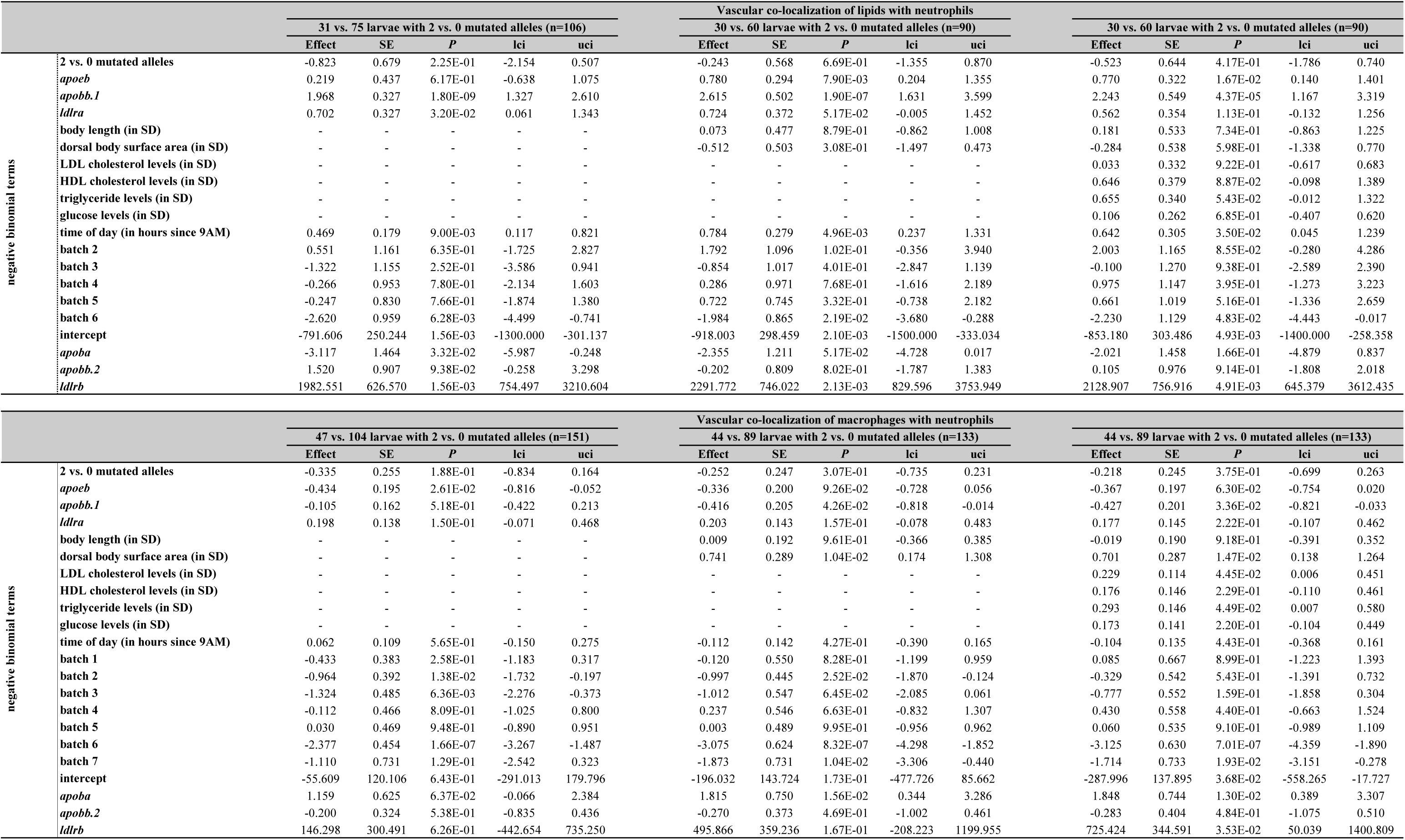
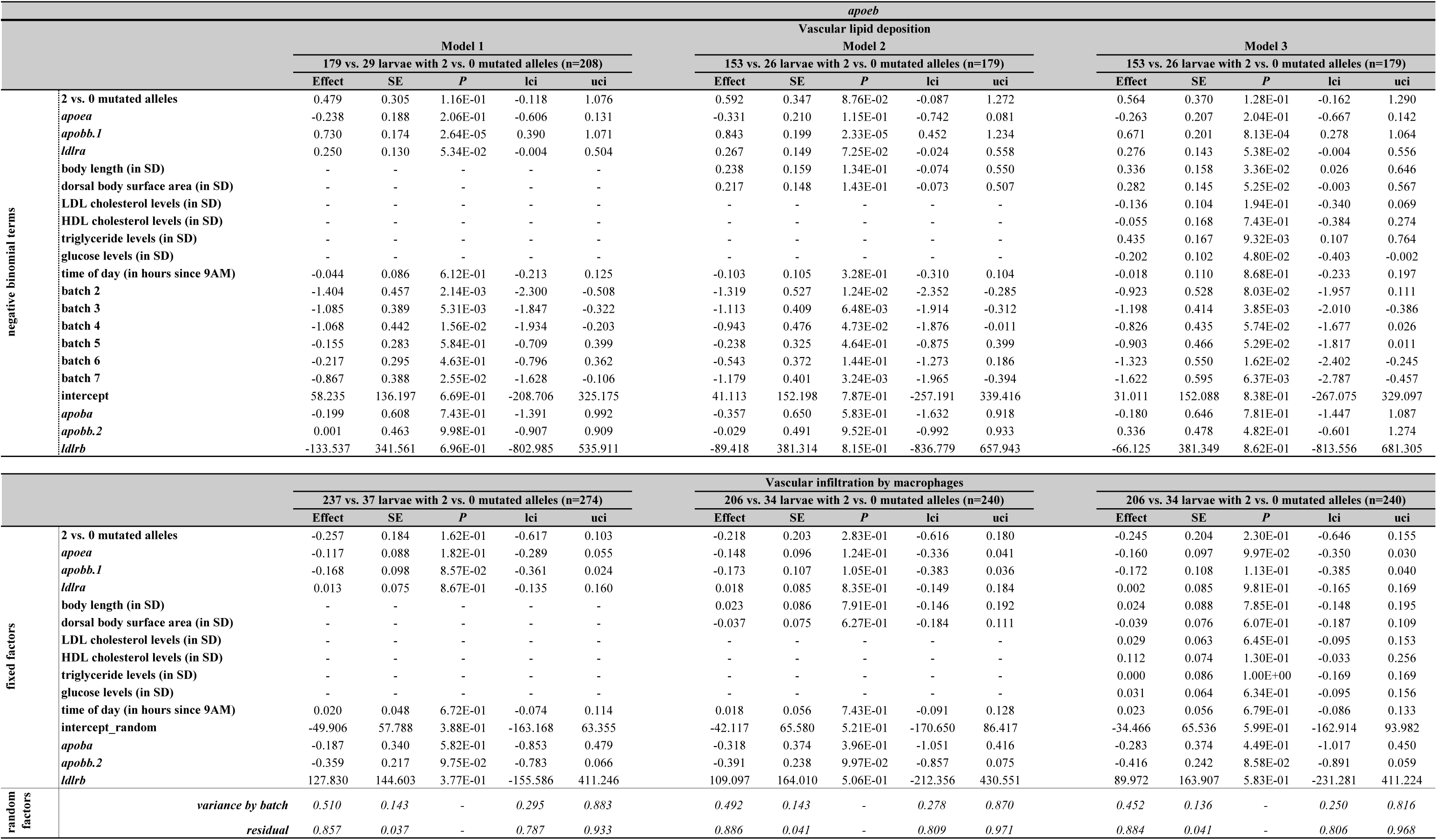
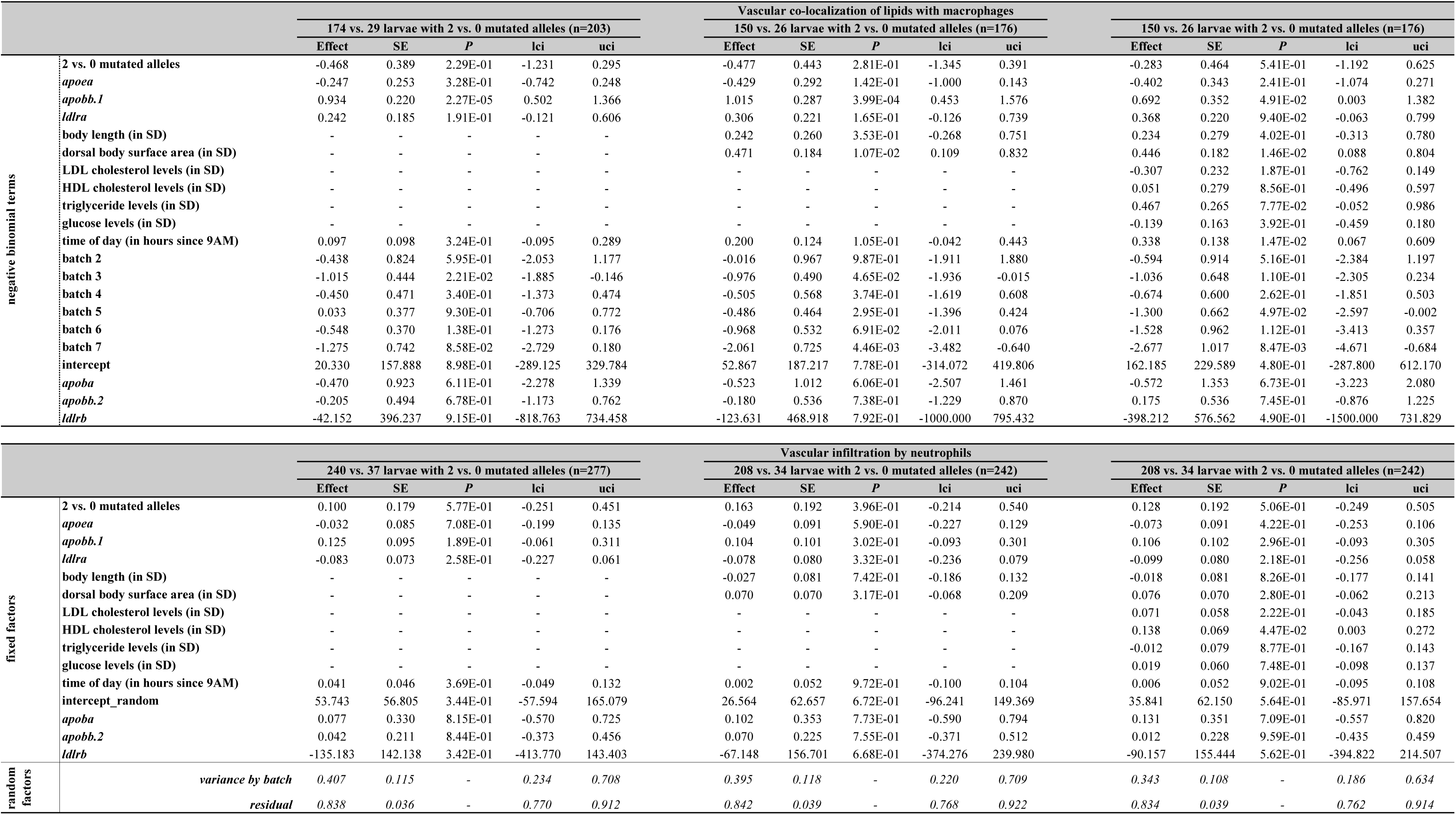
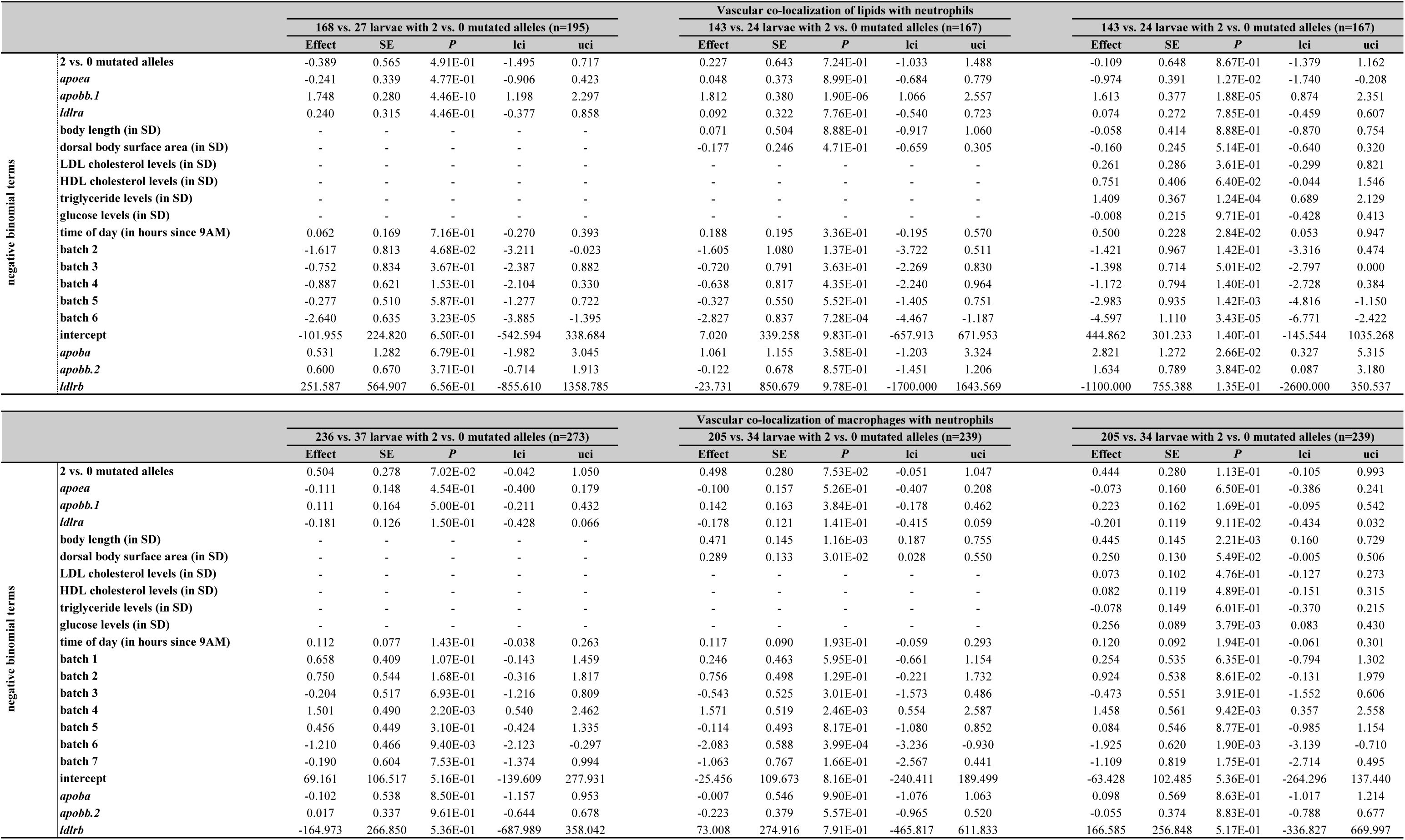
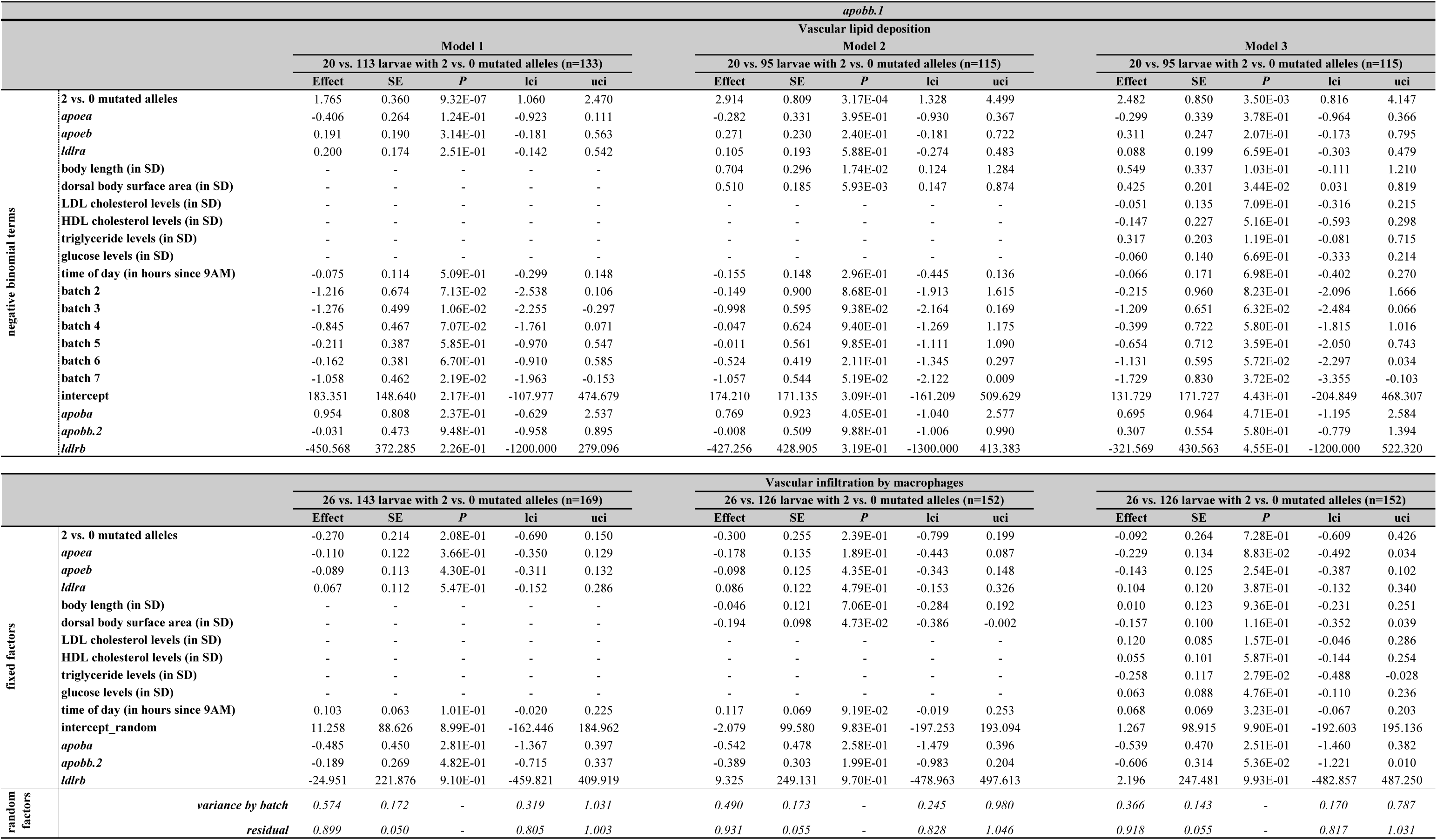
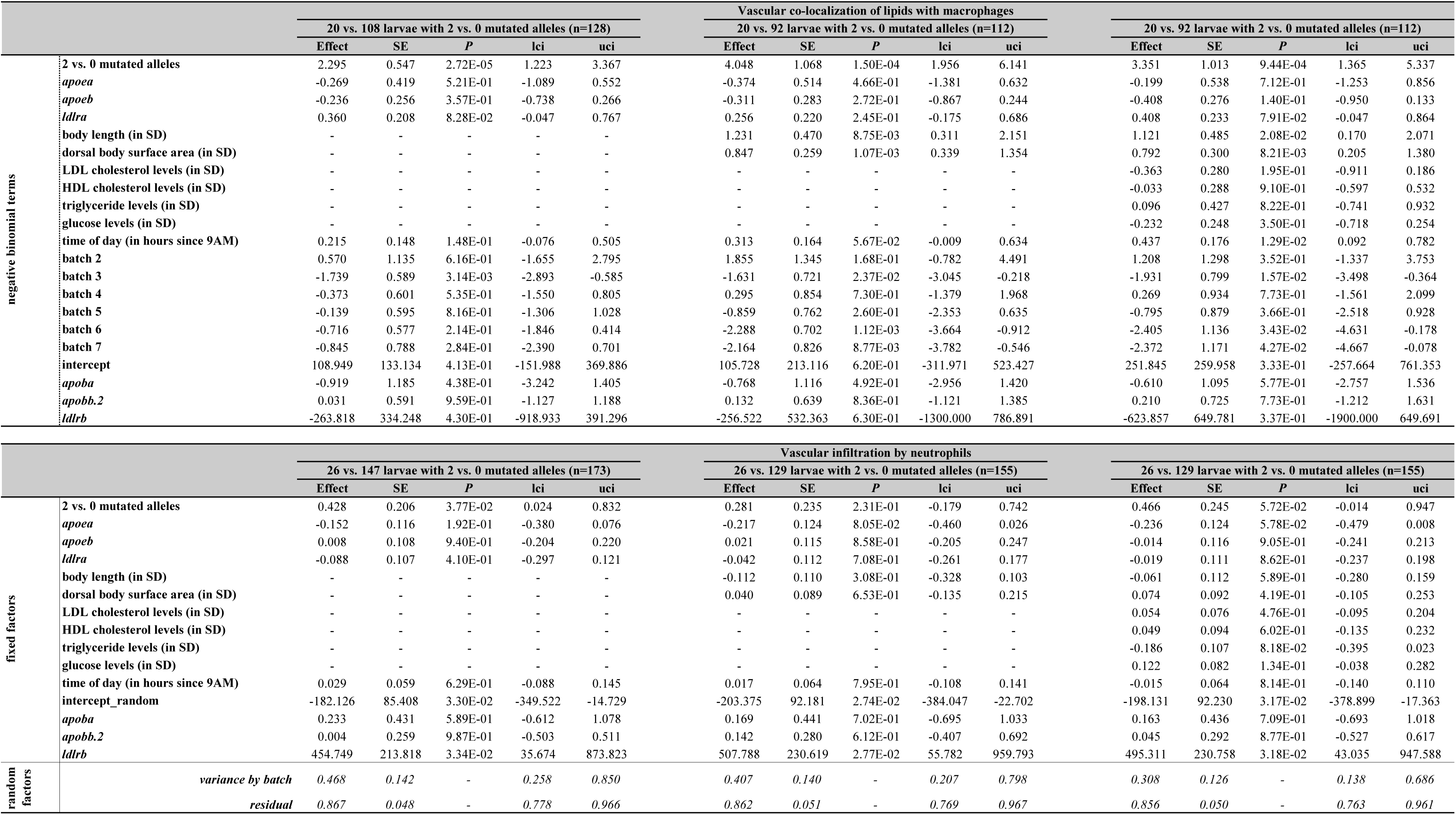
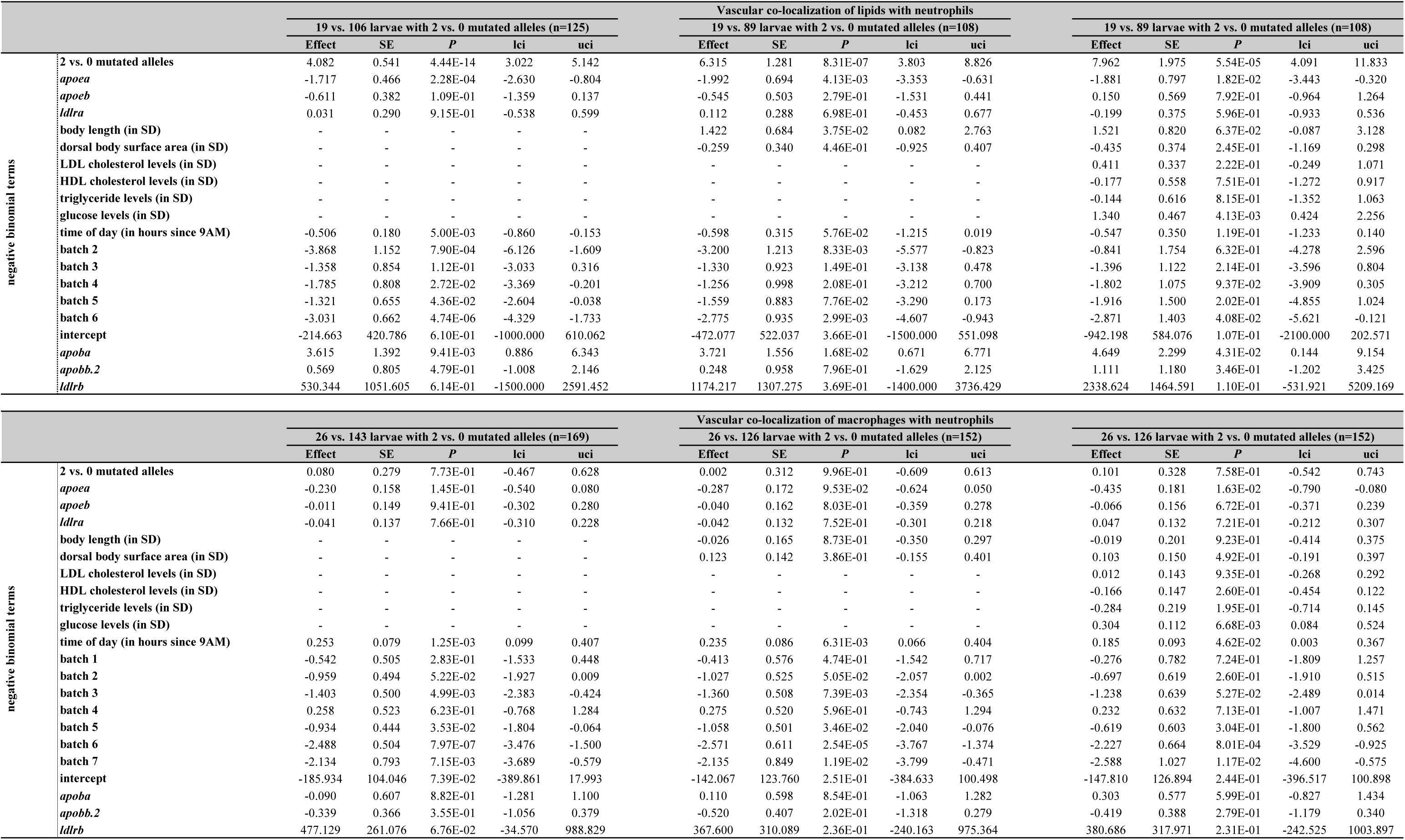
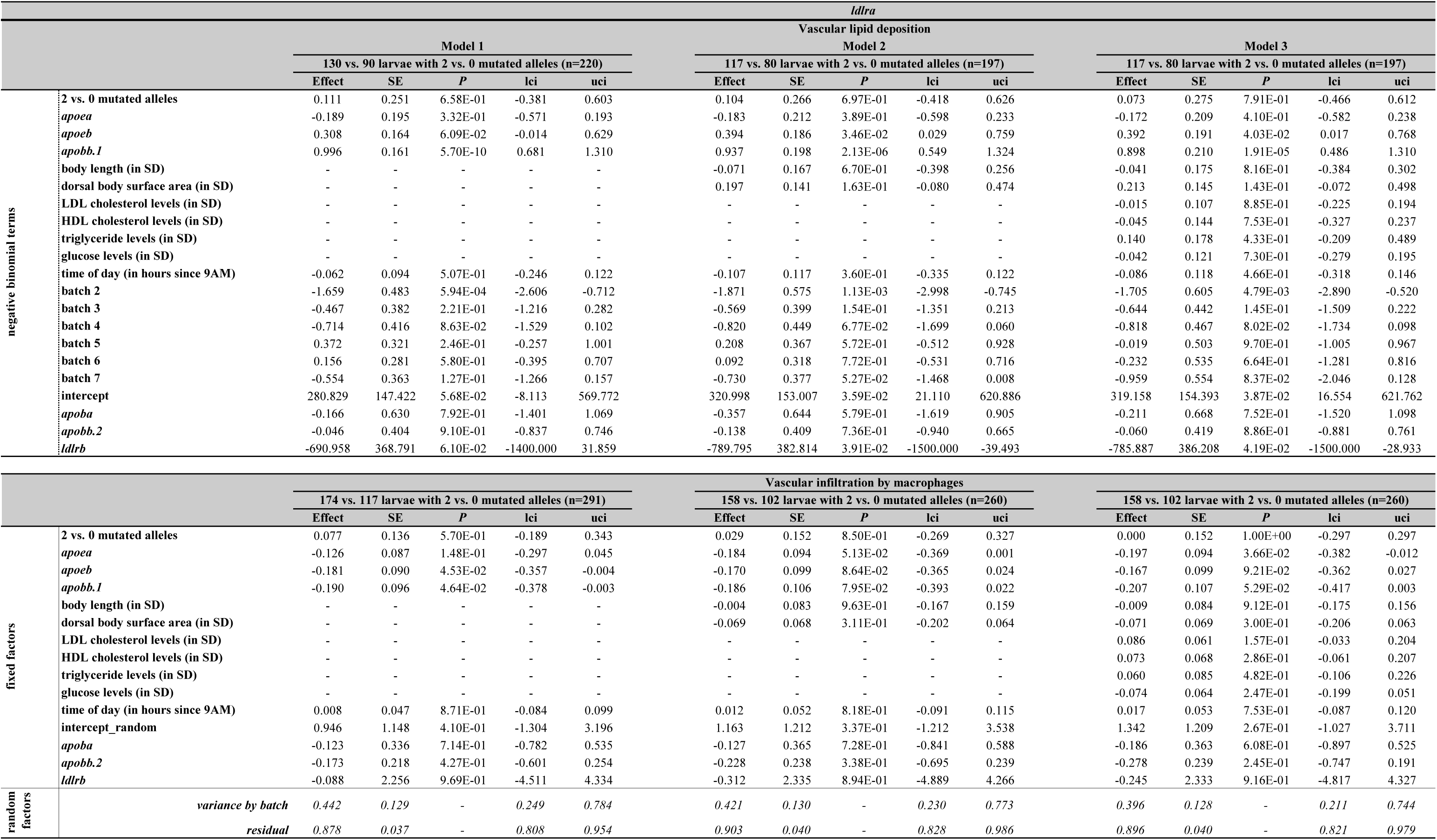
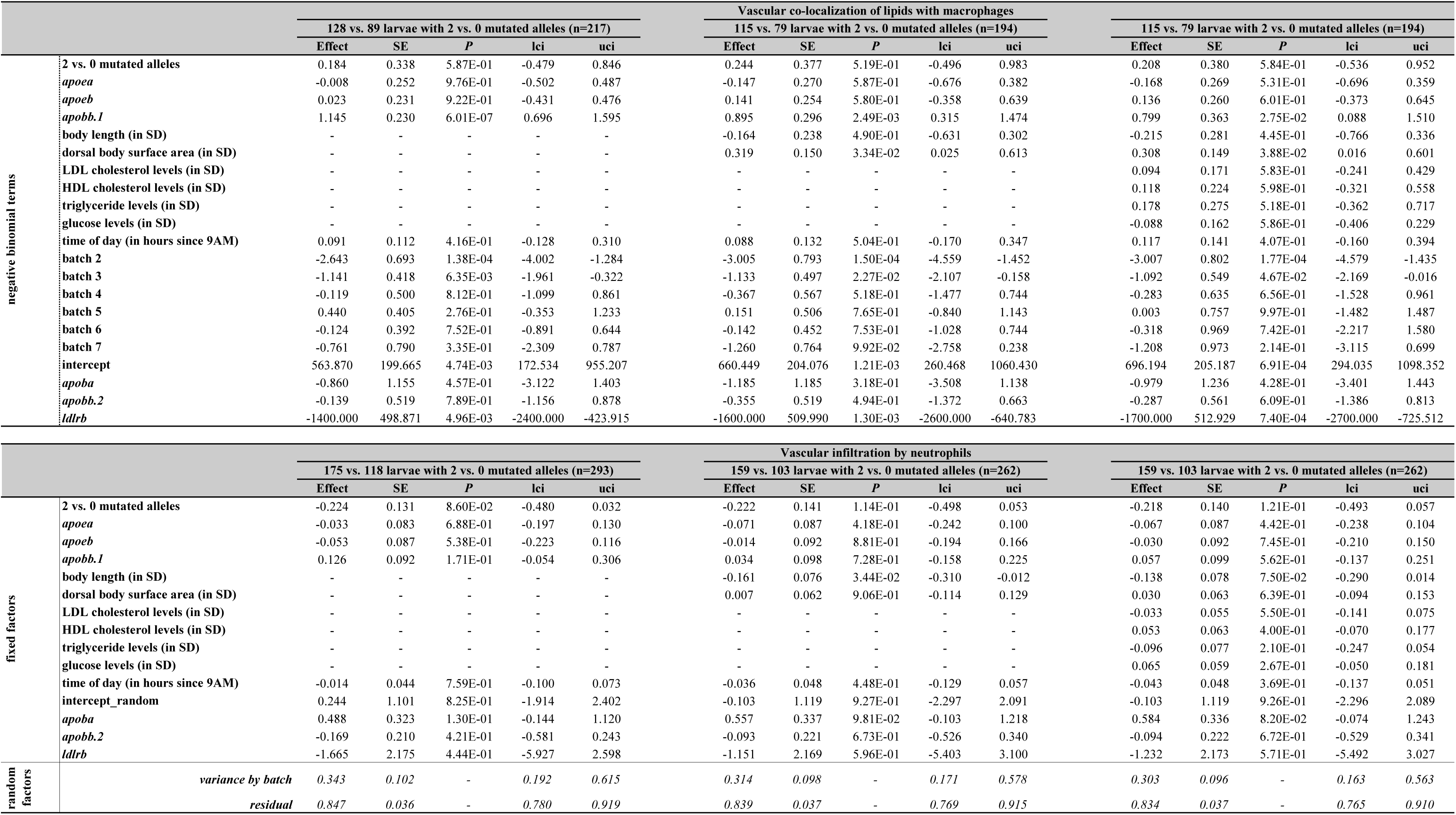
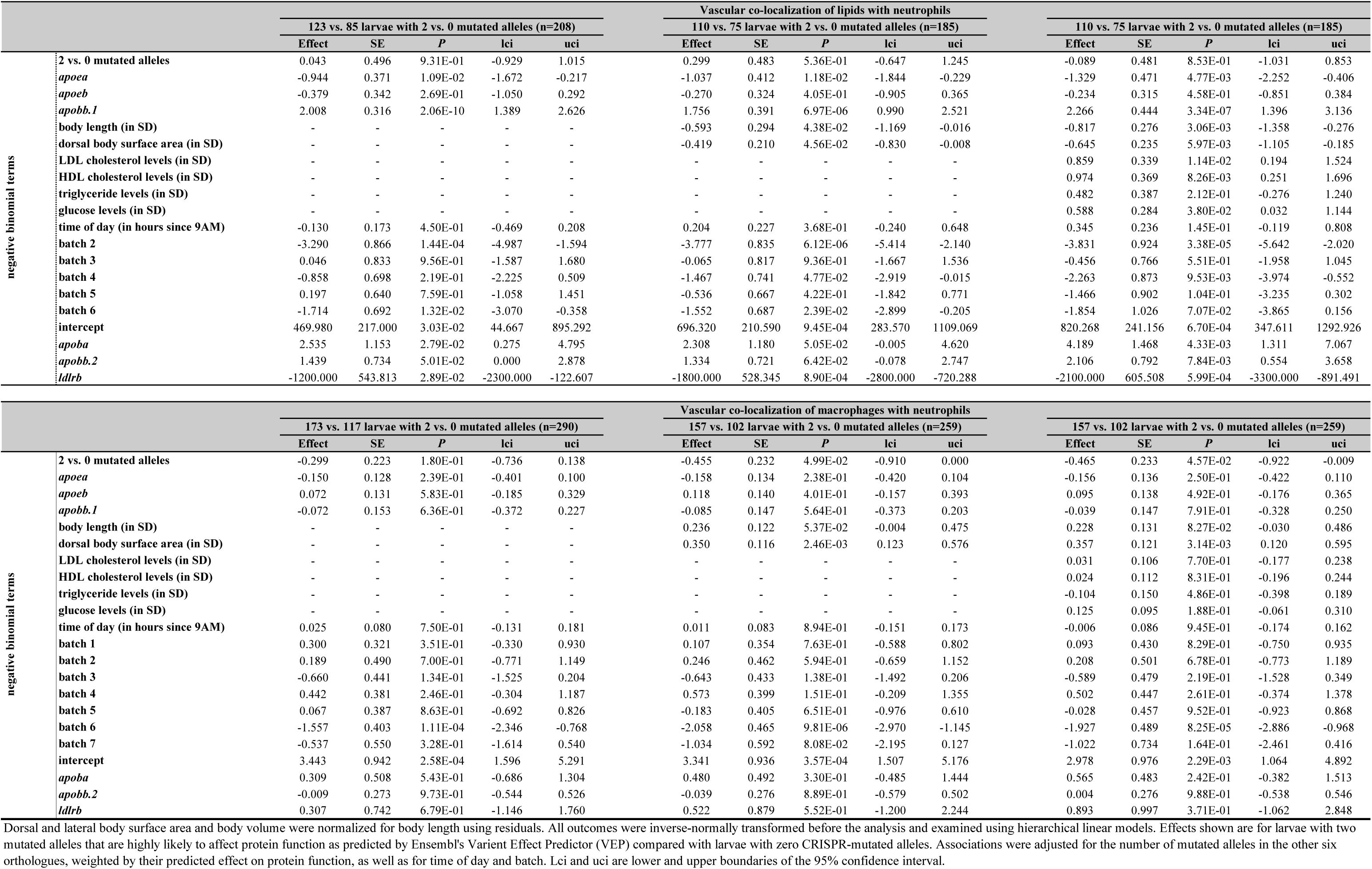
The effect of two vs. zero mutated alleles in *apoea*, *apoeb*, *apobb.1* or *ldlra* on vascular atherogenic traits

**Supplementary Table 22.**
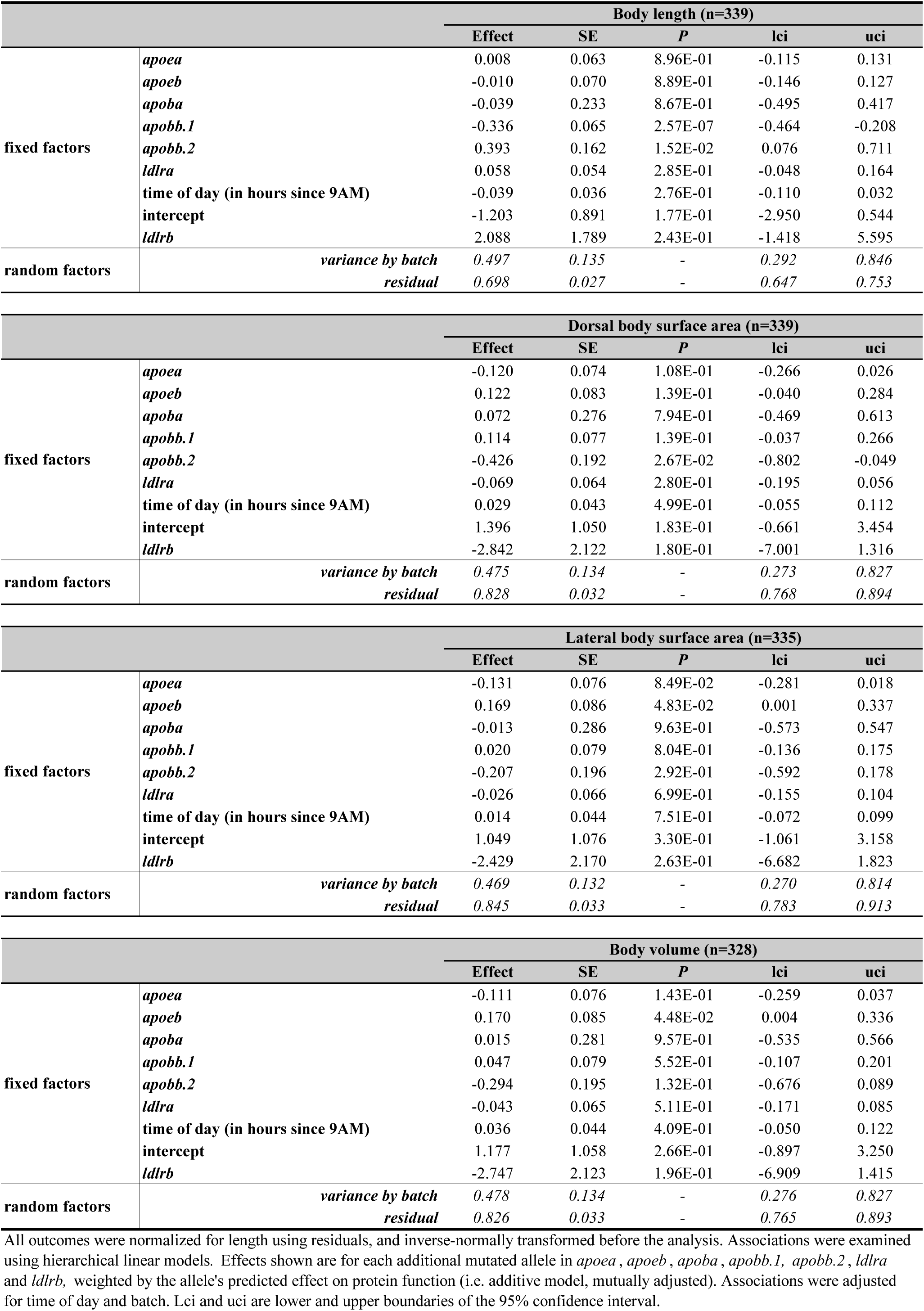
The additive effect of mutated alleles in *apoea*, *apoeb*, *apobb.1* and *ldlra* on body size

**Supplementary Table 23.**
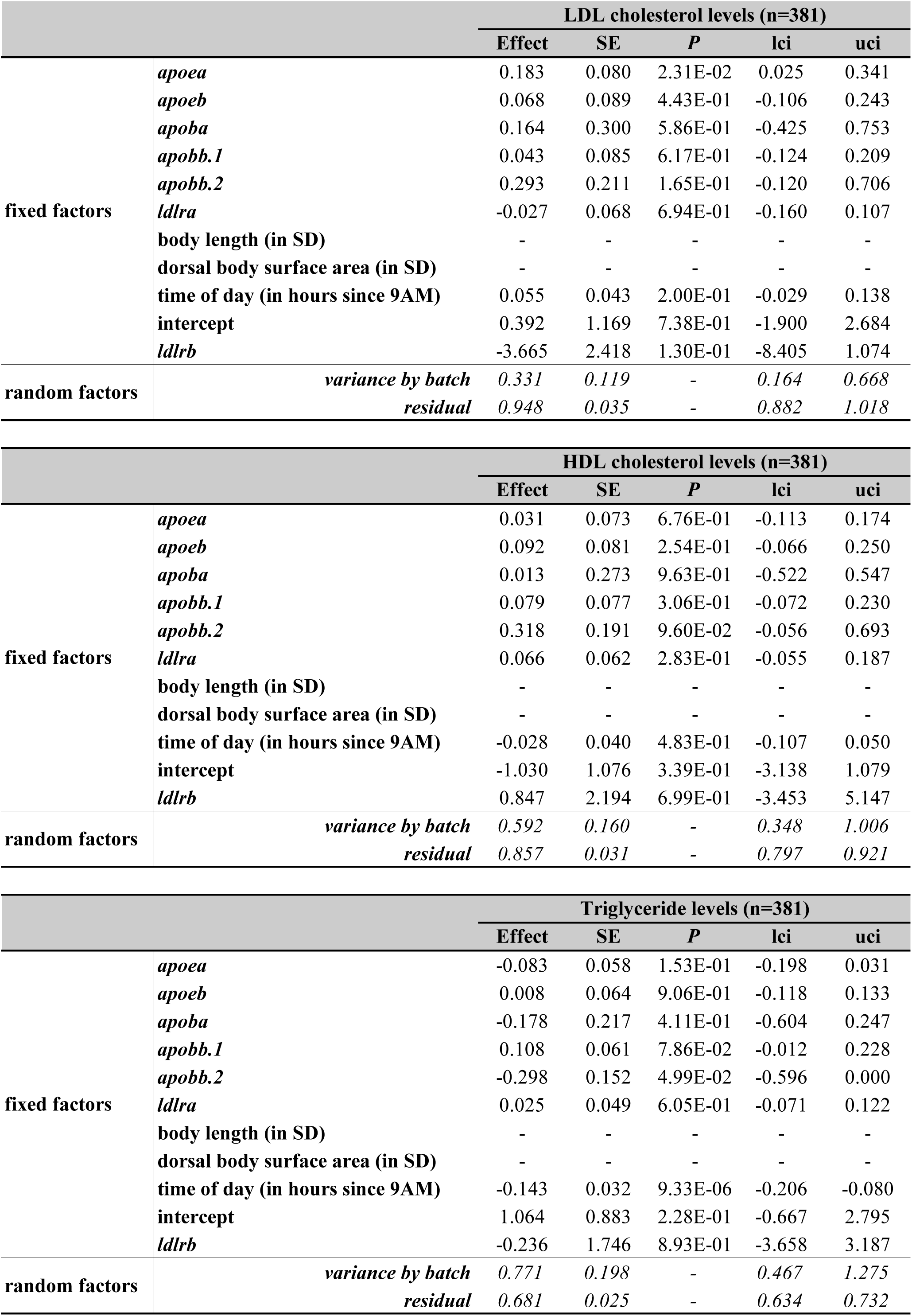
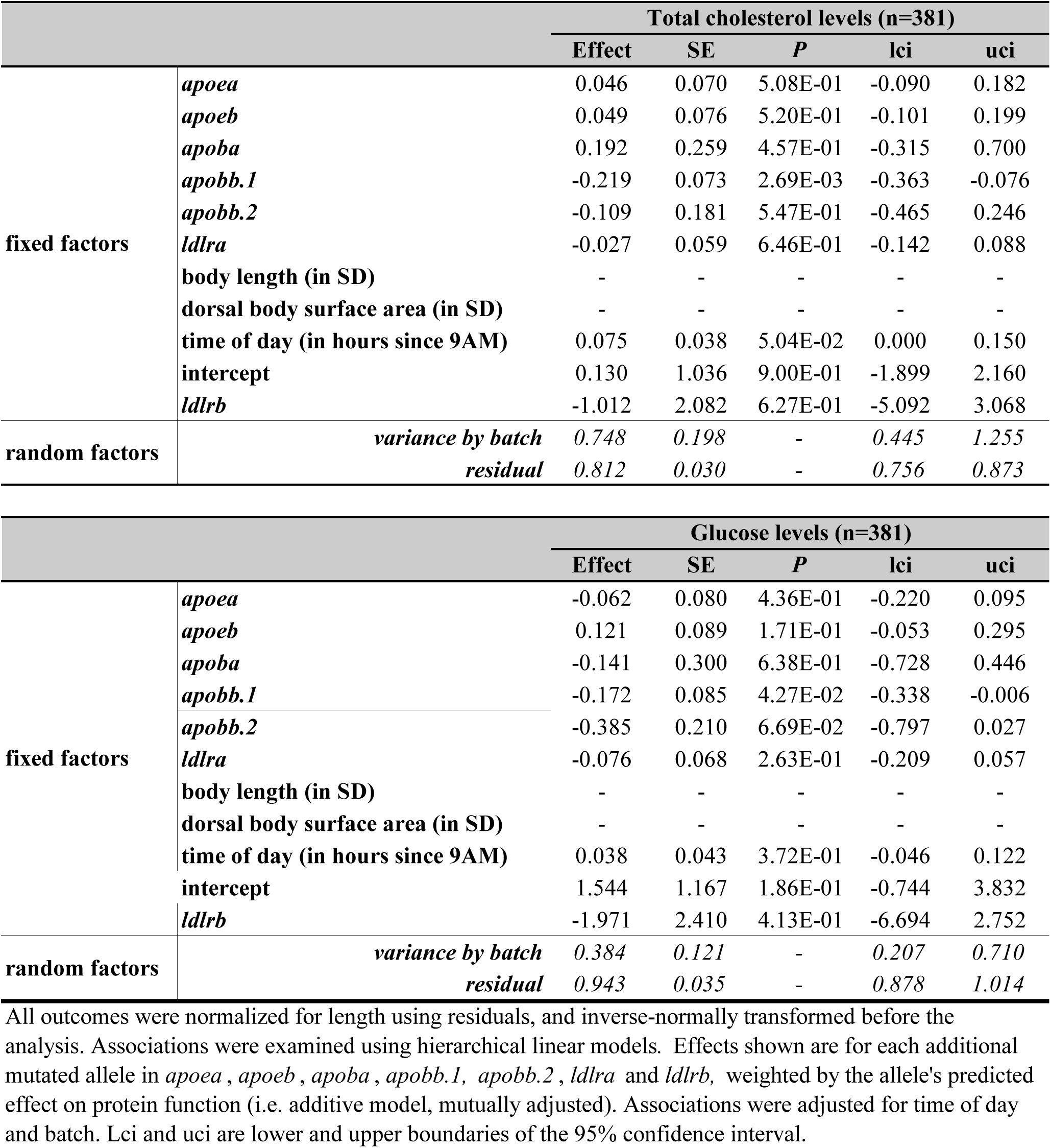
The additive effect of mutated alleles in *apoea*, *apoeb*, *apobb.1* and *ldlra* on

**Supplementary Table 24.**
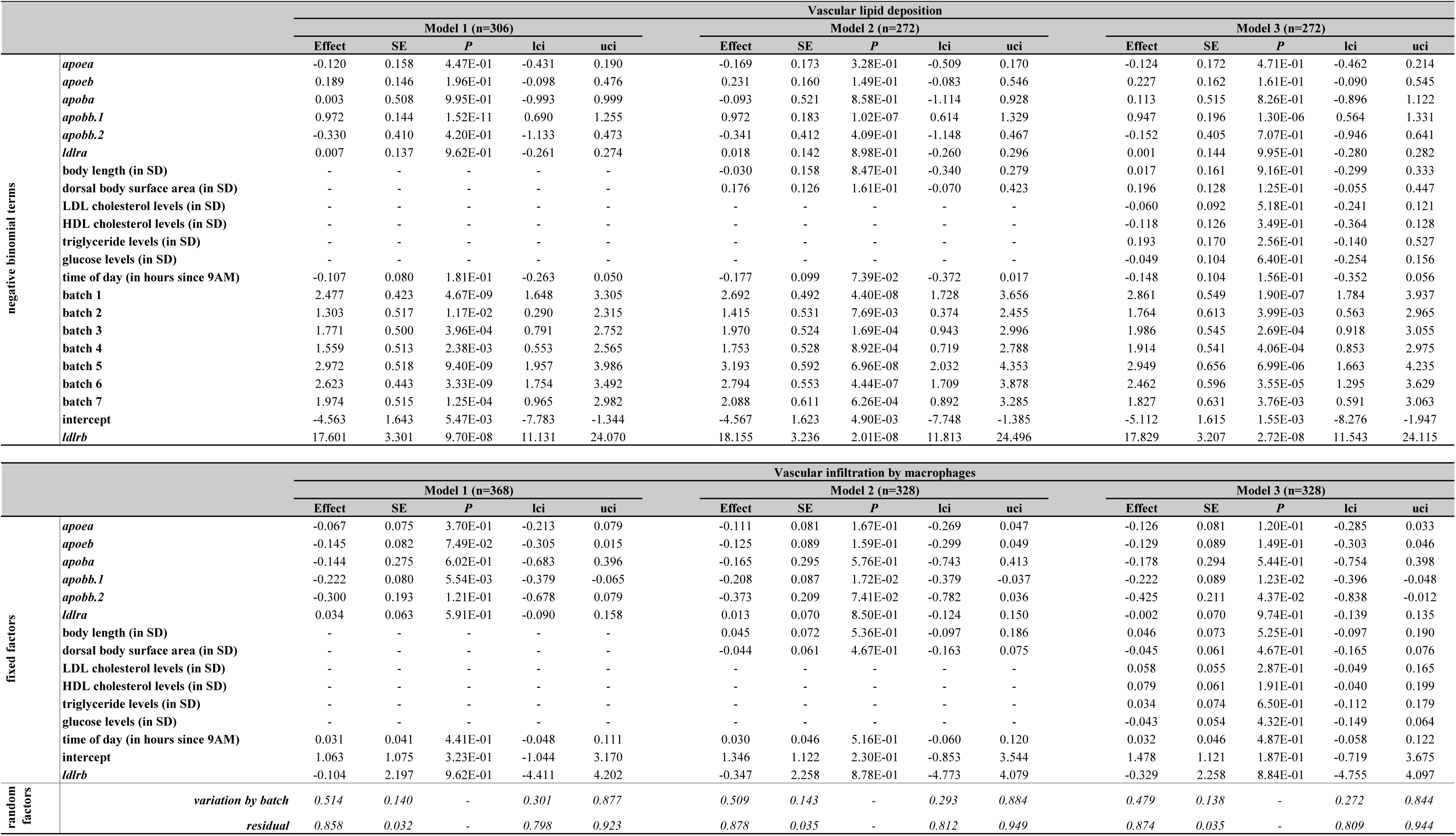
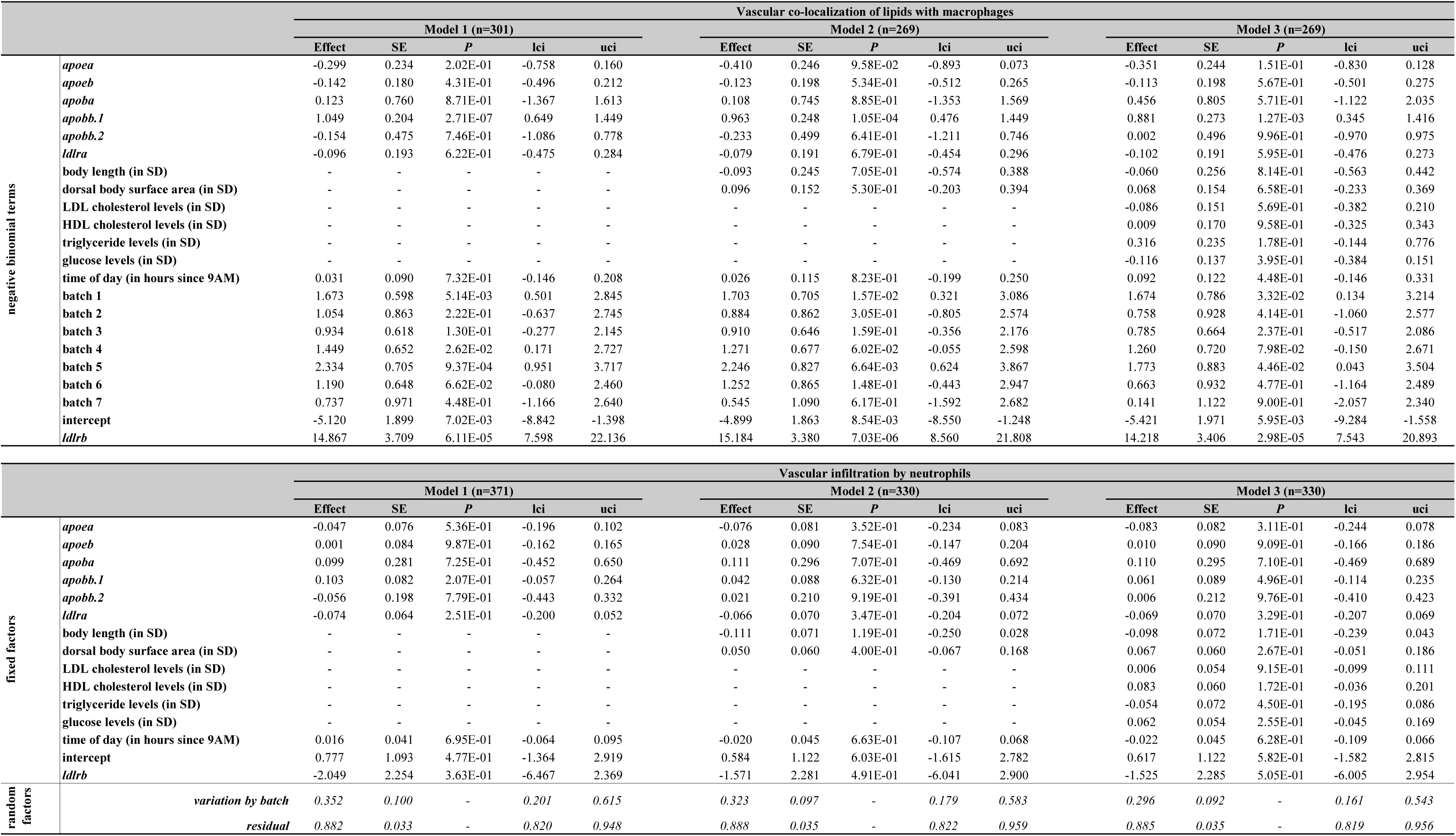
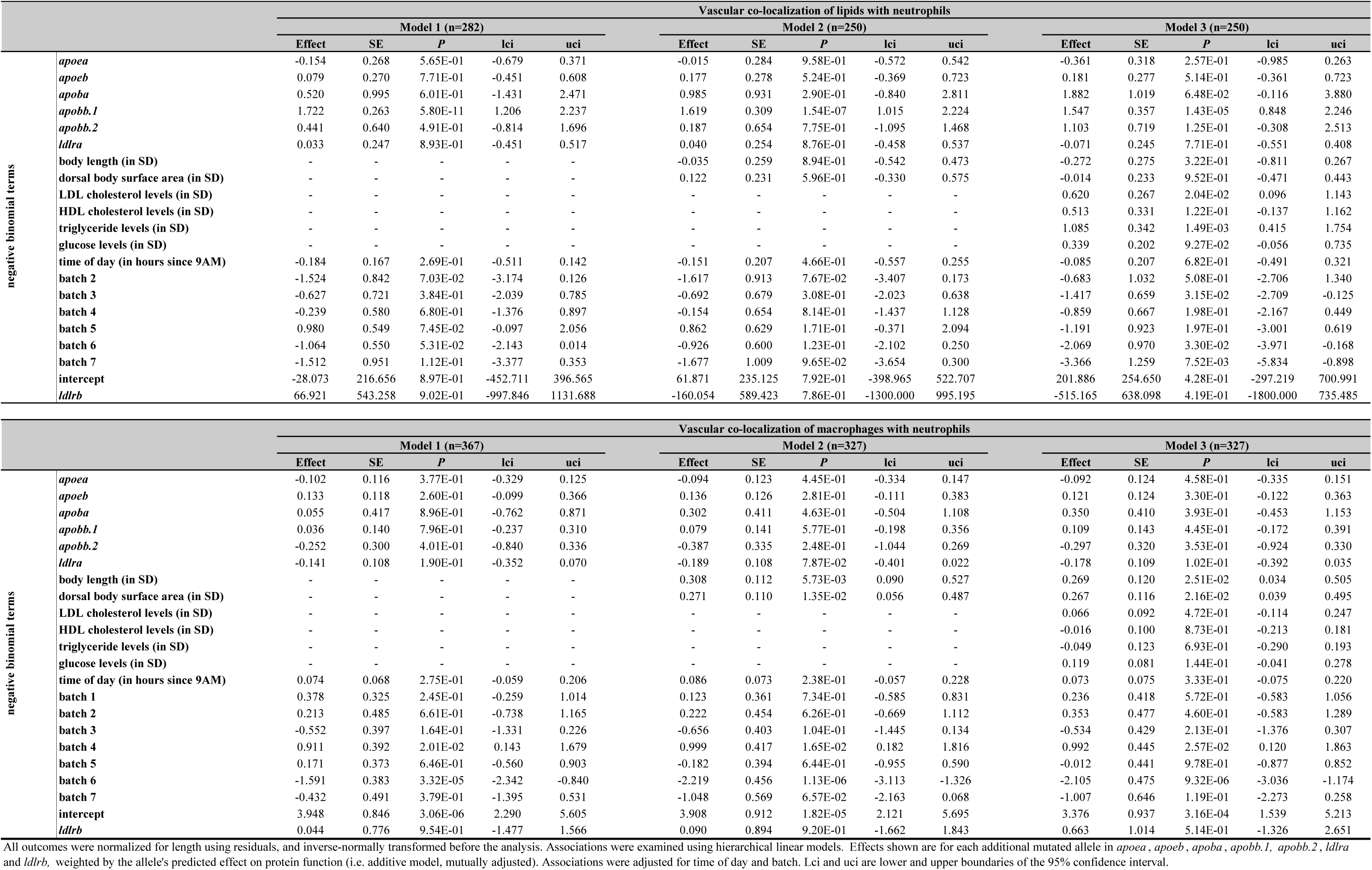
The additive effect of mutated alleles in *apoea*, *apoeb*, *apobb.1* and *ldlra* on image-based vascular atherogenic traits

**Supplementary Table 25.**
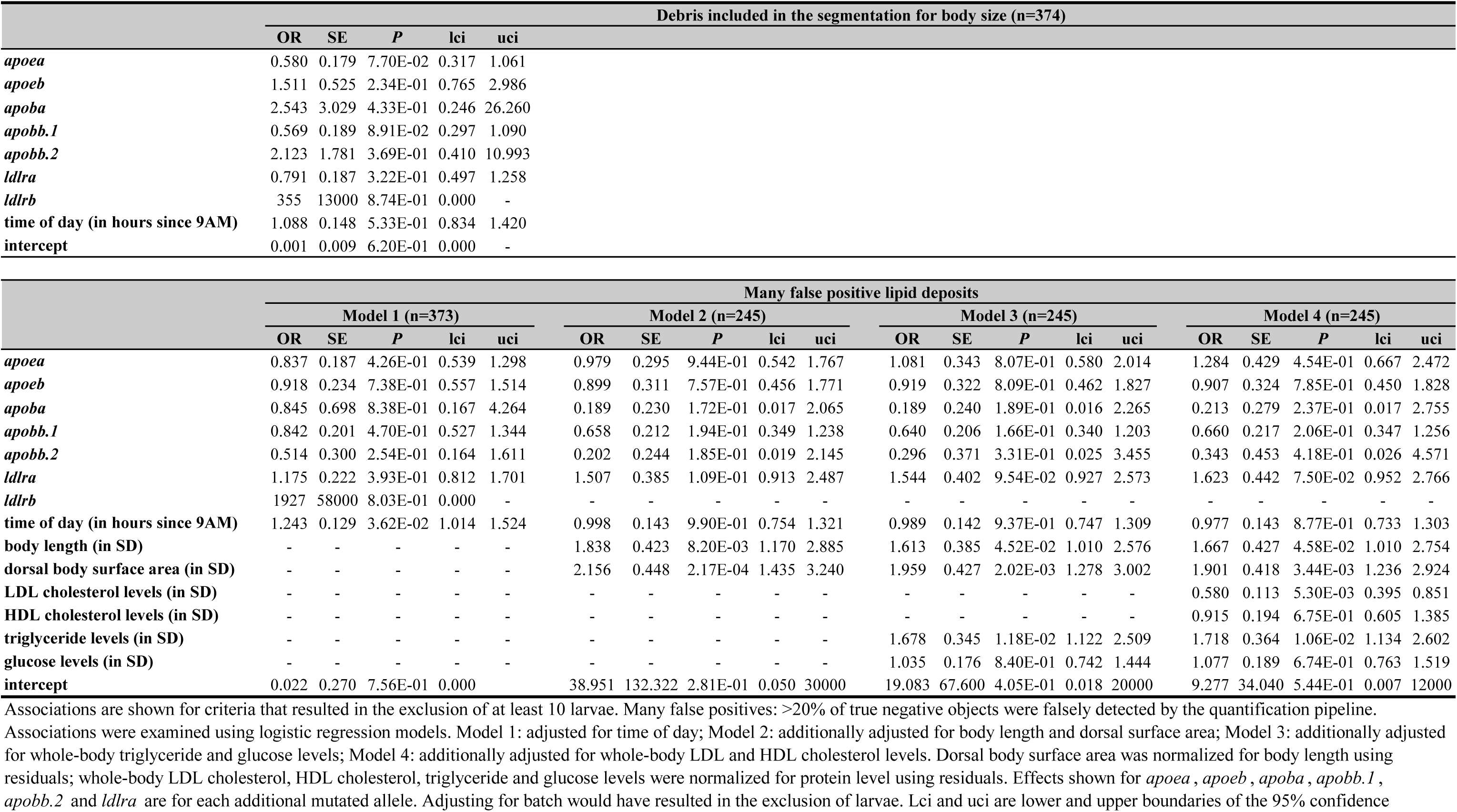
The additive effect of mutated alleles in *apoea*, *apoeb*, *apoba*, *apobb.1, apobb.2, ldlra* and *ldlrb* on image and image quantification quality

**Supplementary Table 26.**
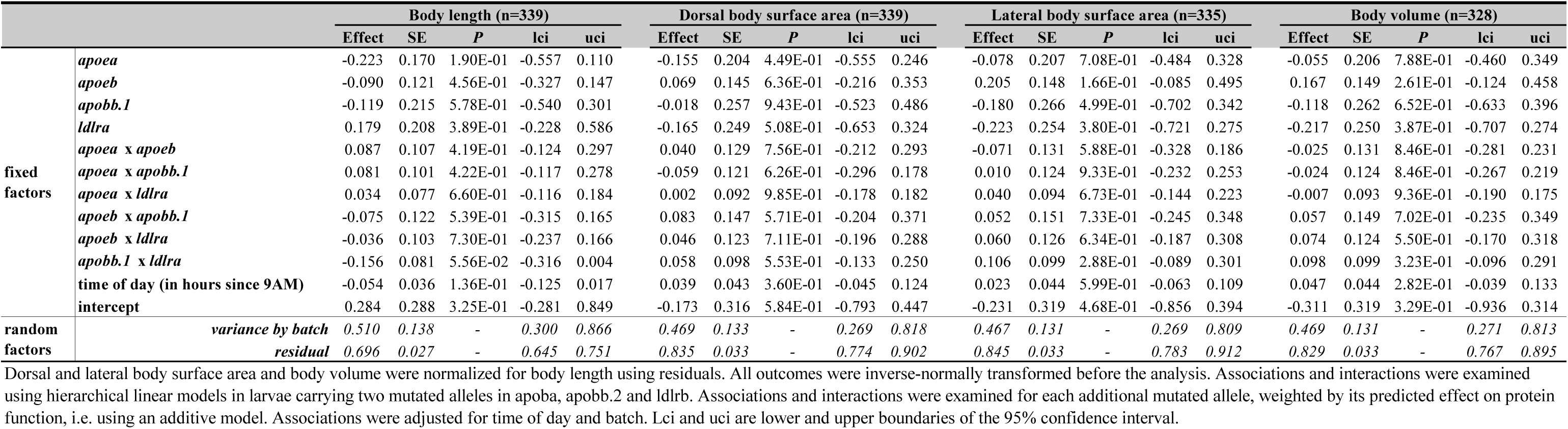
The effect of gene x gene interactions on body size

**Supplementary Table 27.**
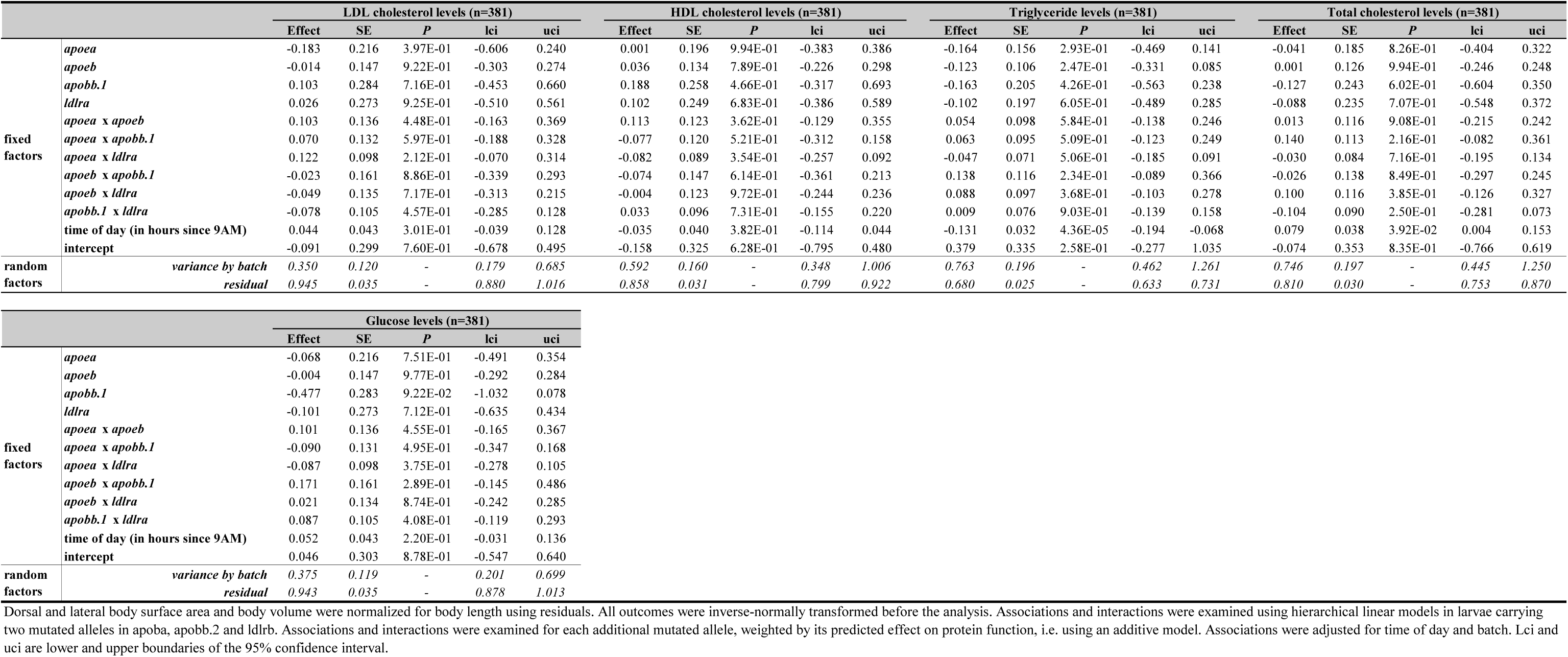
The effect of gene x gene interactions on whole-body lipid and glucose levels

**Supplementary Table 28.**
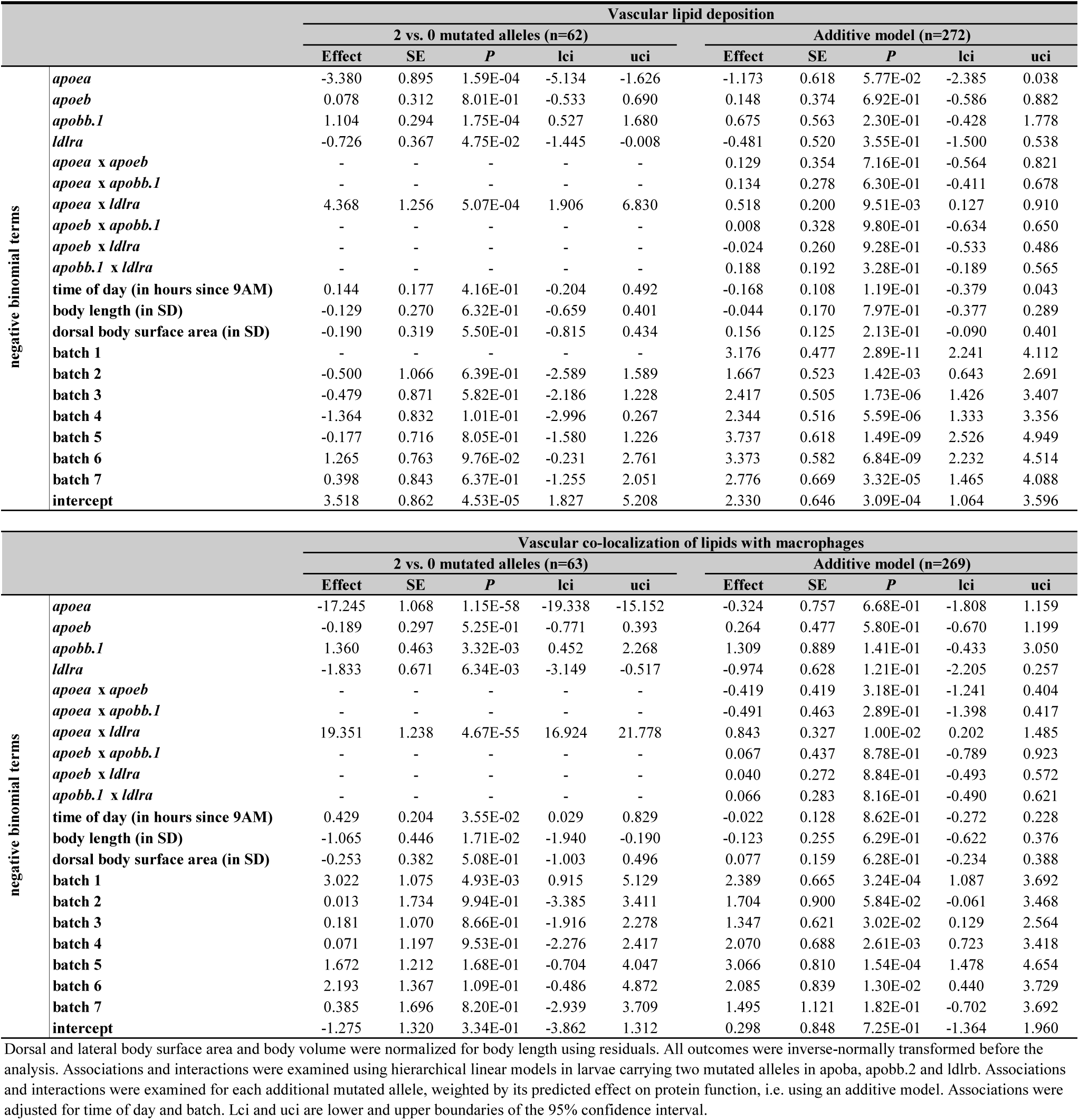
The effect of gene x gene interactions on image-based vascular atherogenic traits

**Supplementary Table 29.**
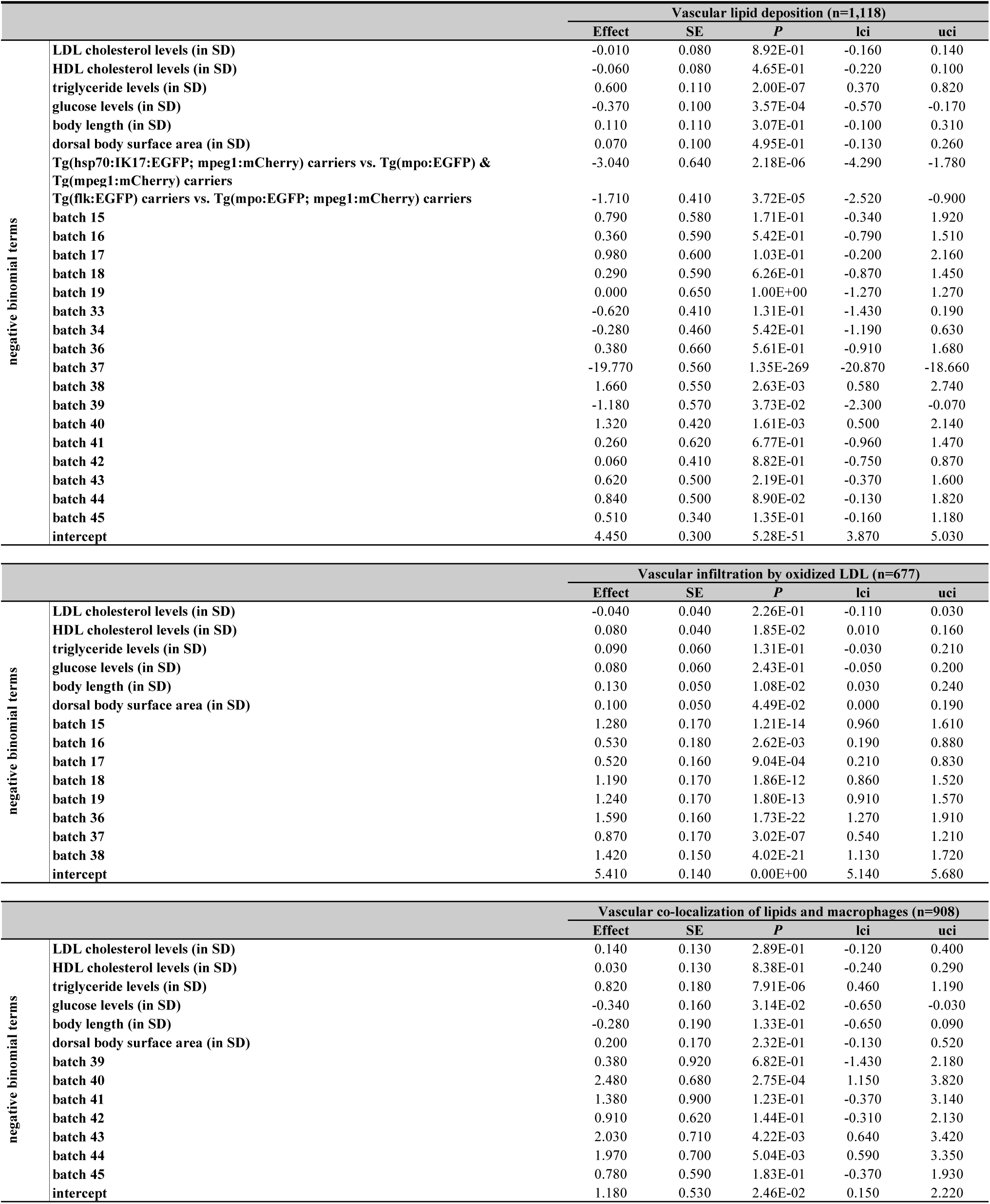
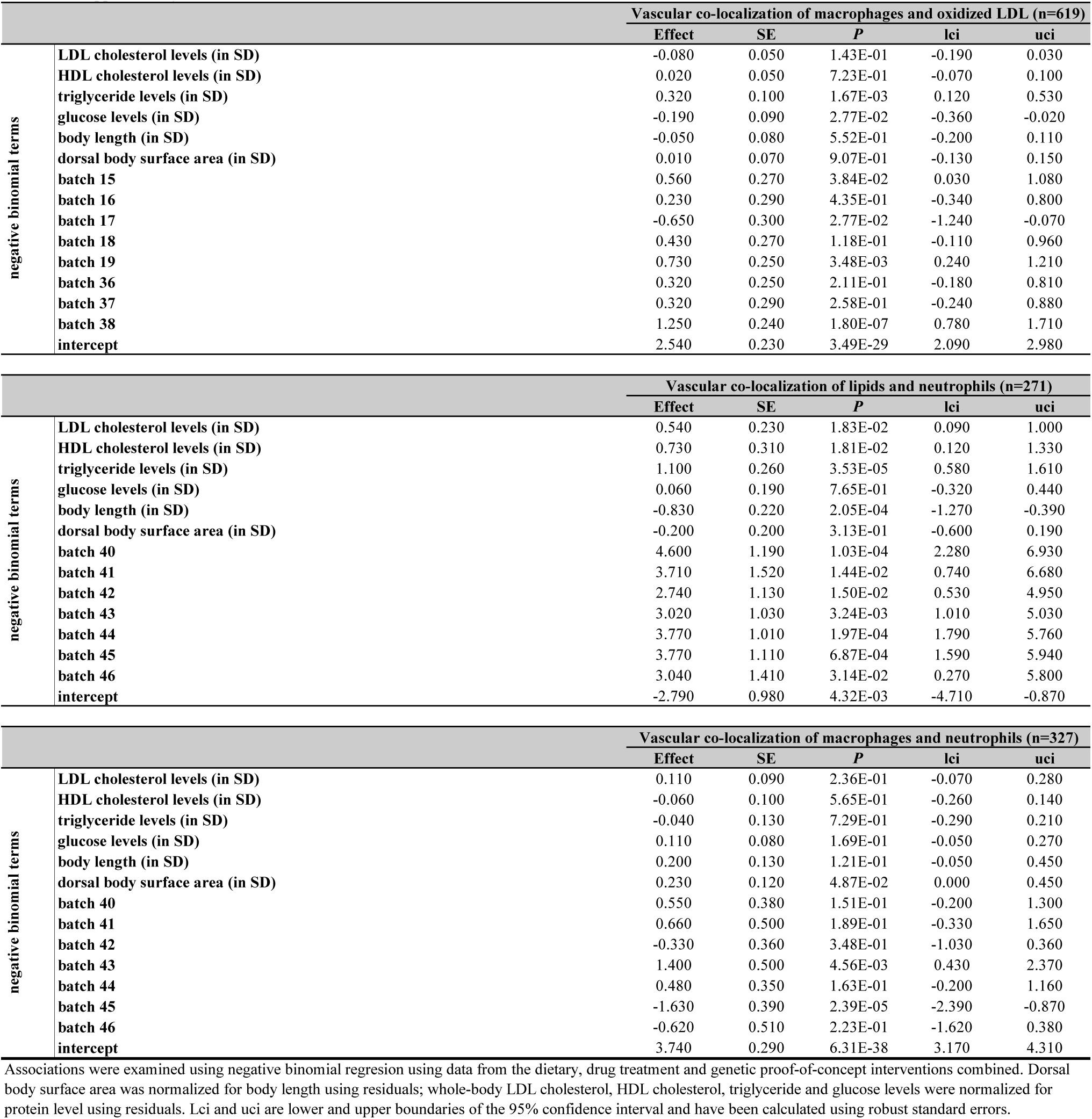
The association of image-based vascular atherogenic traits with whole-body lipid and glucose levels

**Supplementary Table 30.**
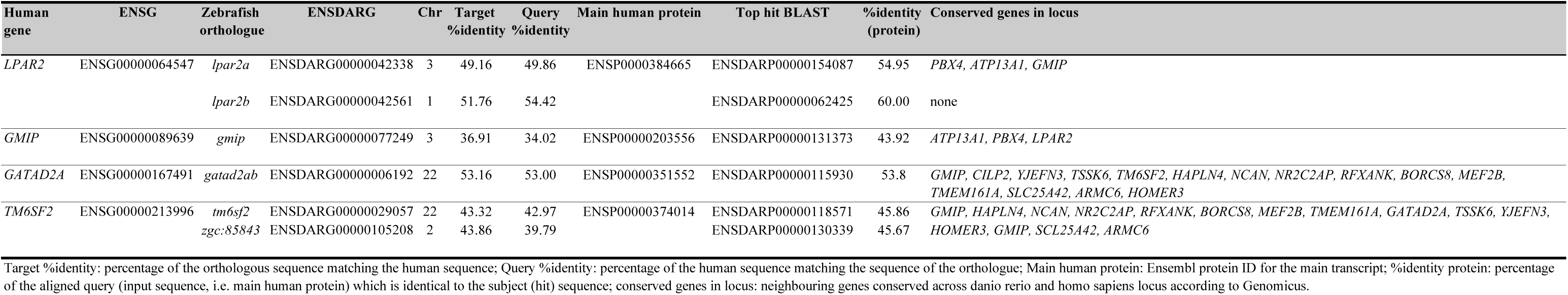
Orthologues of candidate genes in the triglyceride, LDLc and total cholesterol-associated locus on chr 19p13.11

| Human gene | ENSG | Zebrafish orthologue | ENSDARG | Chr | Target %identity | Query %identity | Main human protein | Top hit BLAST | %identity (protein) | Conserved genes in locus |
| --- | --- | --- | --- | --- | --- | --- | --- | --- | --- | --- |
| *LPAR2* | ENSG000000064547 | *lpar2a* | ENSDARG000000042338 | 3 | 49.16 | 49.86 | ENSP00000384665 | ENSDARP00000154087 | 54.95 | *PBX4, ATP13A1, GMIP* |
|  |  | *lpar2b* | ENSDARG000000042561 | 1 | 51.76 | 54.42 |  | ENSDARP00000062425 | 60.00 | none |
| *GMIP* | ENSG000000089639 | *gmip* | ENSDARG000000077249 | 3 | 36.91 | 34.02 | ENSP00000203556 | ENSDARP00000131373 | 43.92 | *ATP13A1, PBX4, LPAR2* |
| *GATAD2A* | ENSG00000167491 | *gata2ab* | ENSDARG000000006192 | 22 | 53.16 | 53.00 | ENSP00000351552 | ENSDARP00000115930 | 53.8 | *GMIP, CILP2, YIEFN3, TSSK6, TM6SF2, HAPLN4, NCAN, NR2C2AP, RFXANK, BORCS8, MEF2B, TMEM161A, SLC25A42, ARMC6, HOMER3* |
| *TM6SF2* | ENSG00000213996 | *tm6sf2* | ENSDARG00000029057 | 22 | 43.32 | 42.97 | ENSP00000374014 | ENSDARP00000118571 | 45.86 | *GMIP, HAPLN4, NCAN, NR2C2AP, RFXANK, BORCS8, MEF2B, TMEM161A, GATAD2A, TSSK6, YIEFN3,* |
|  |  | *zgc:85843* | ENSDARG00000105208 | 2 | 43.86 | 39.79 |  | ENSDARP00000130339 | 45.67 | *HOMER3, GMIP, SCL25A42, ARMC6* |

Target %identity: percentage of the orthologous sequence matching the human sequence; Query %identity: percentage of the human sequence matching the sequence of the orthologue; Main human protein: Ensembl protein ID for the main transcript; %identity protein: percentage of the aligned query (input sequence, i.e. main human protein) which is identical to the subject (hit) sequence; conserved genes in locus: neighbouring genes conserved across danio rerio and homo sapiens locus according to Genomicus.

**Supplementary Table 31.**
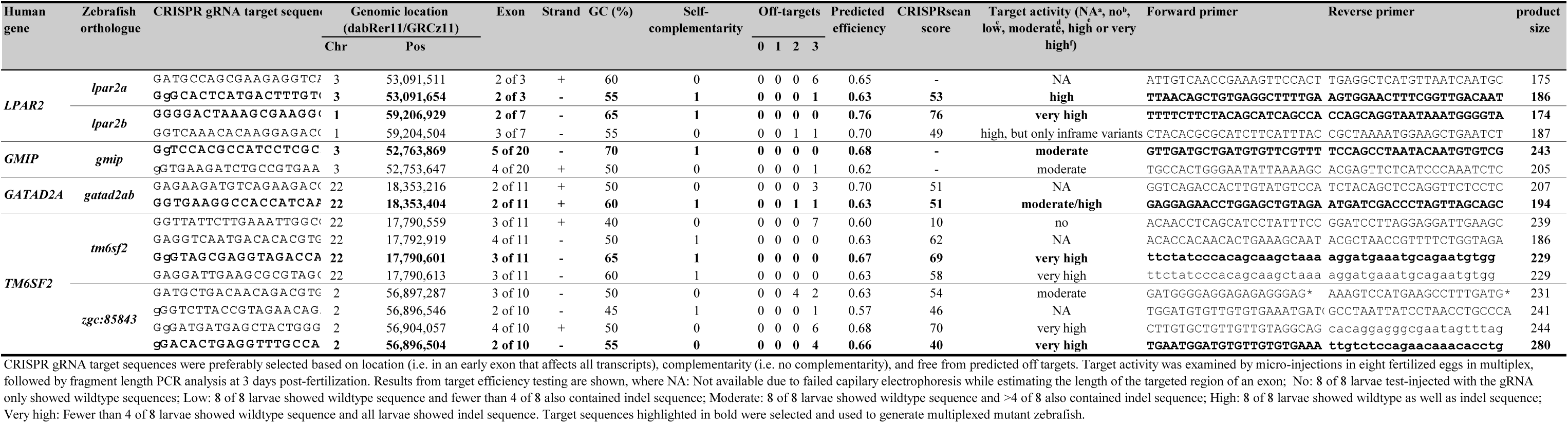
Identification of moderate-to-highly active CRISPR-Cas9 guide RNAs for 19p13.11 candidate genes

**Supplementary Table 32.**
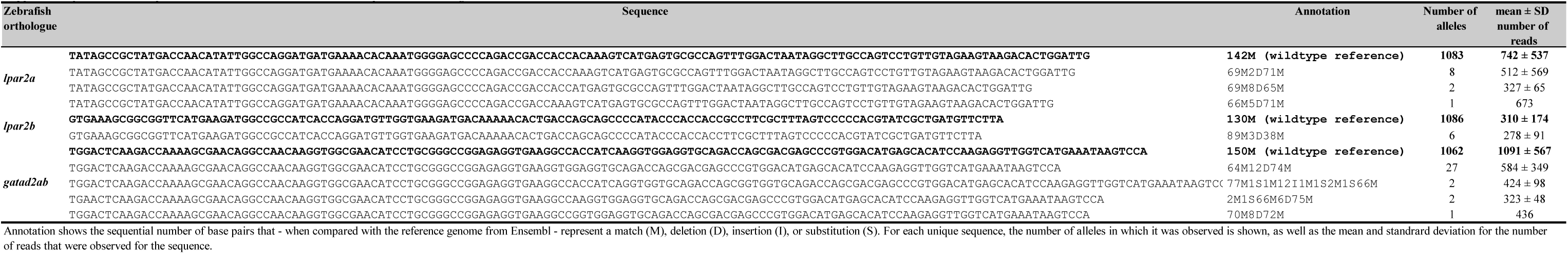
Unique CRISPR-Cas9-induced mutations for 19p13.11 candidate genes

**Supplementary Table 33.**
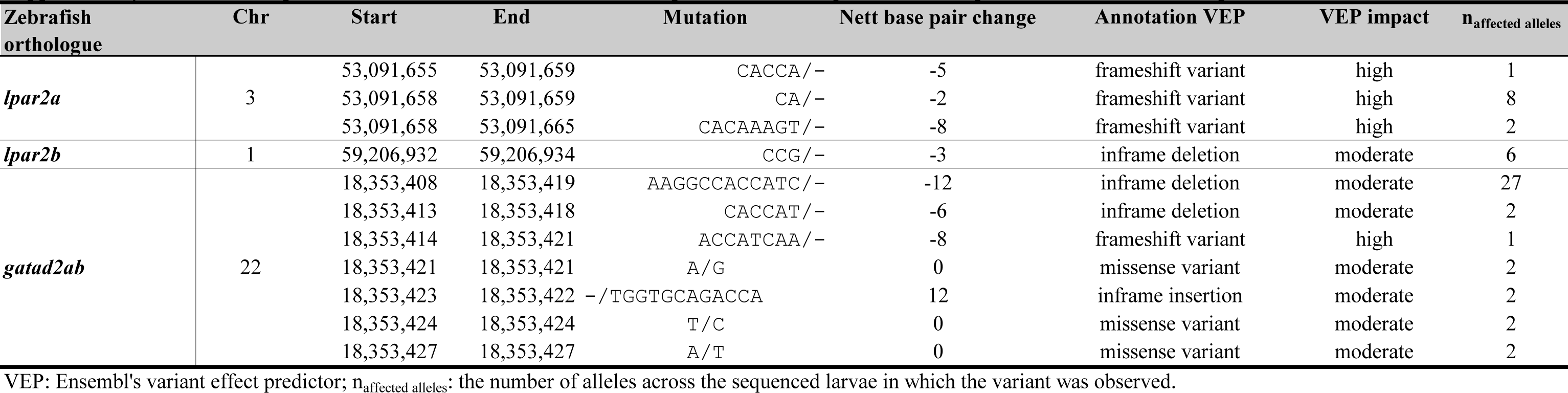
Unique CRISPR-Cas9-induced mutations in 19p13.11 candidate genes and their predicted functional consequences

**Supplementary Table 34.**
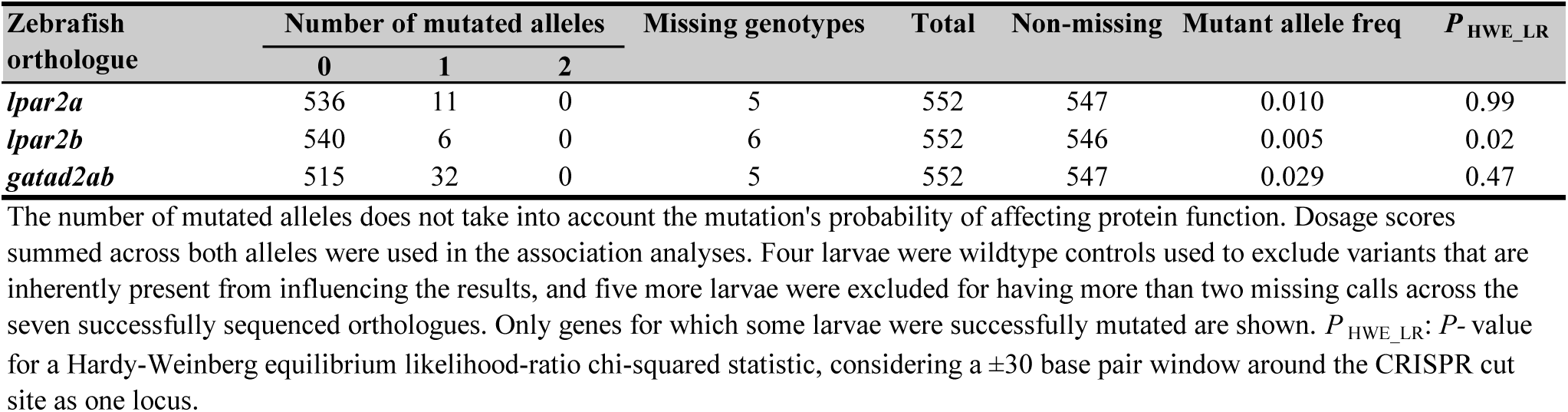
Sequencing results expressed in number of mutated alleles for 19p13.11 candidate genes

**Supplementary Table 35.**
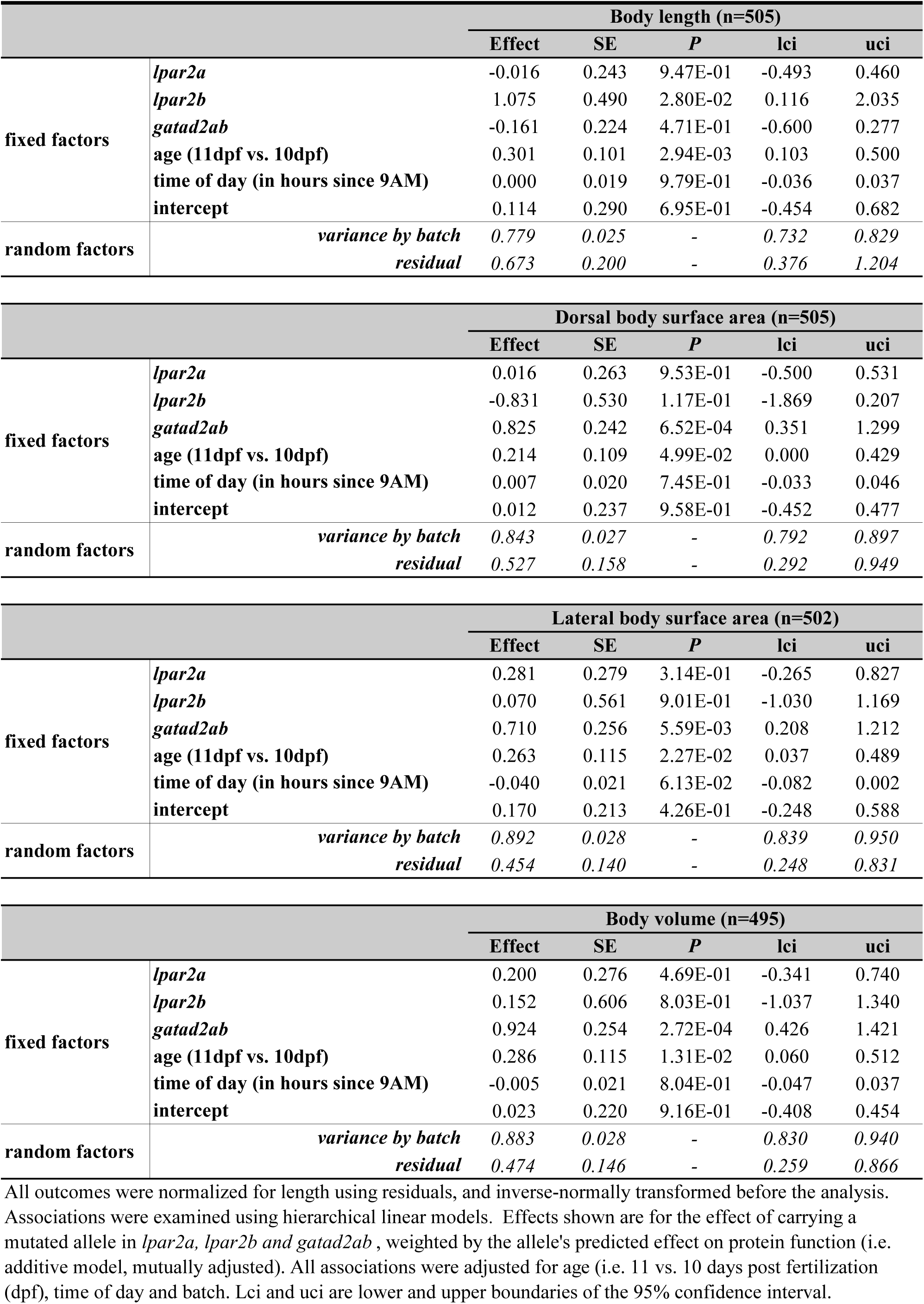
The effect of a mutated allele in *lpar2a*, *lpar2b* and *gatad2ab* on body size

**Supplementary Table 36.**
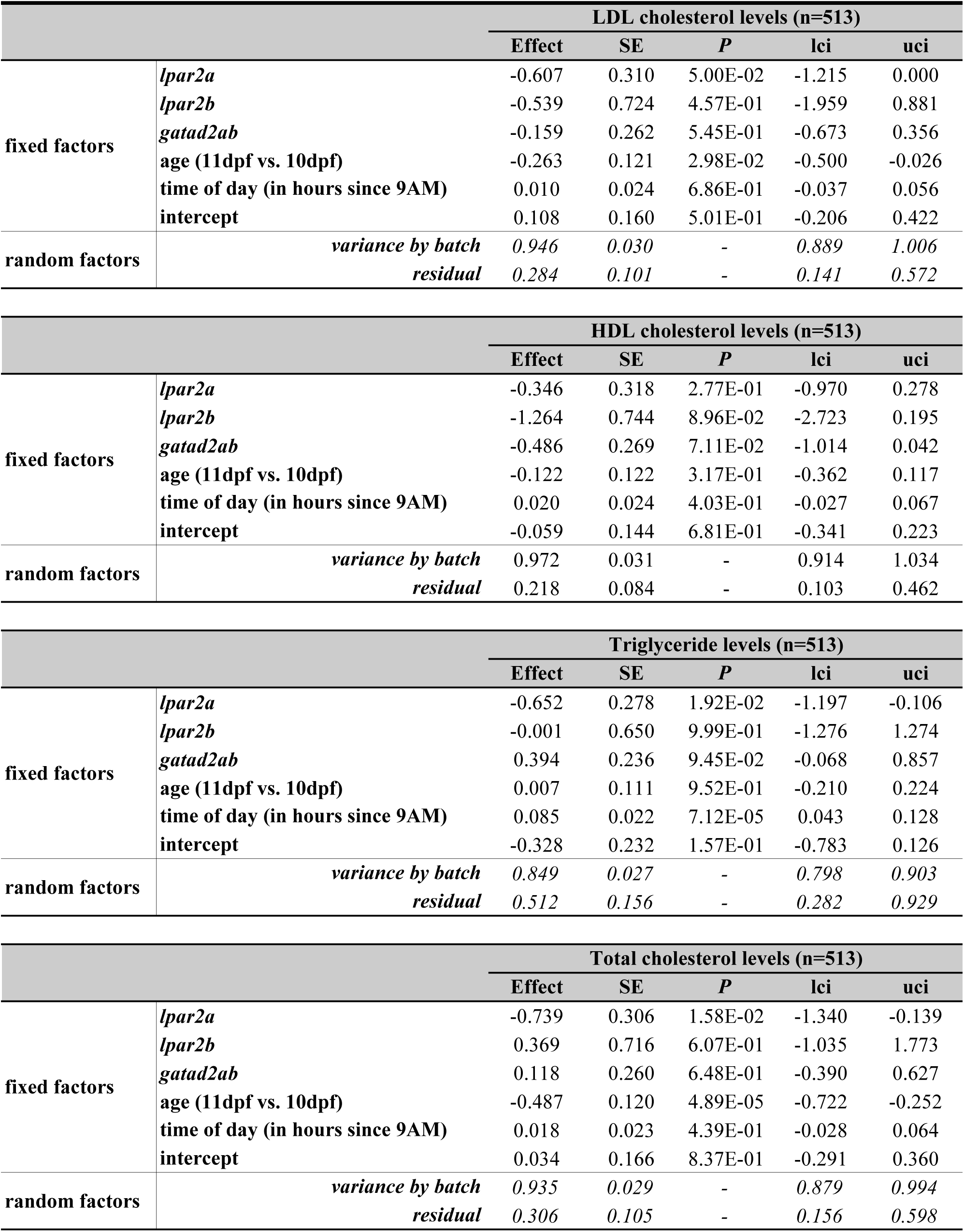
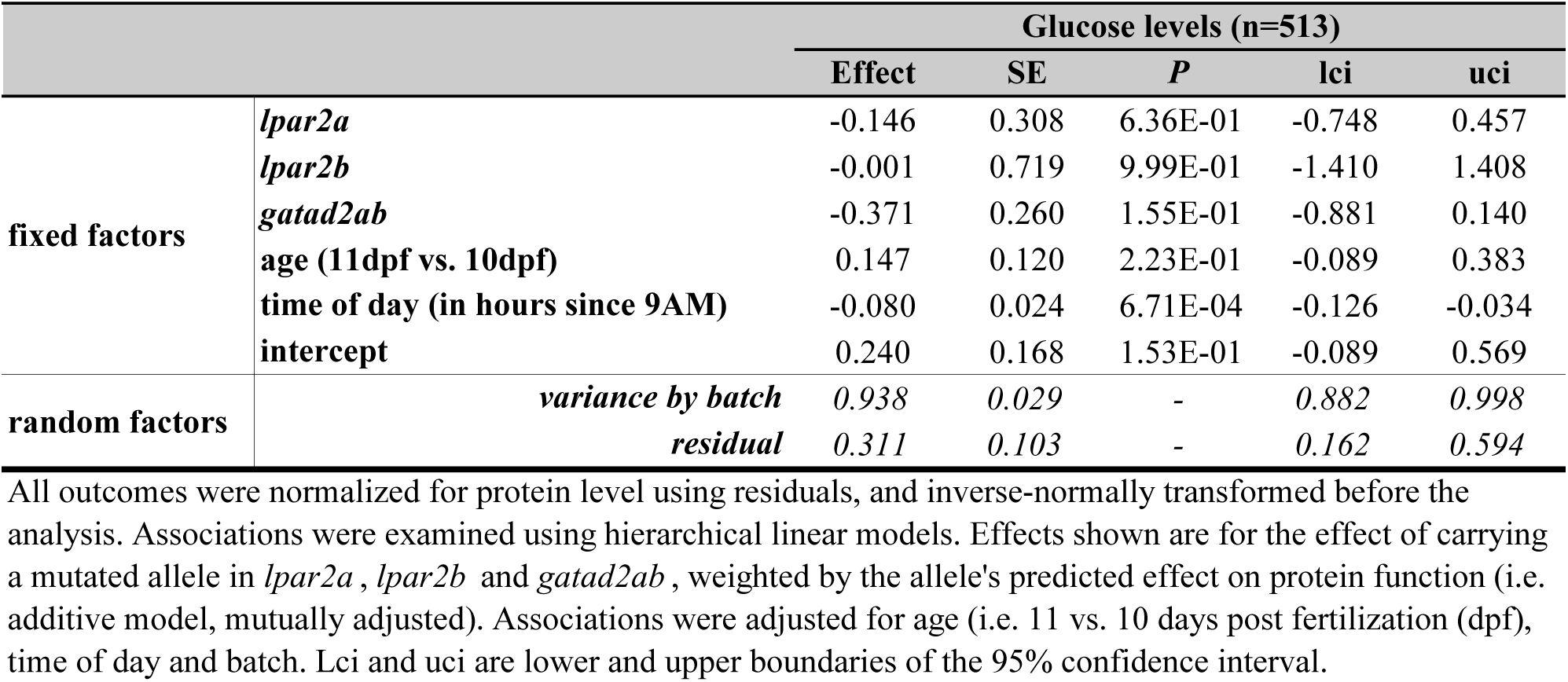
The effect of a mutated allele in *lpar2a, lpar2b* and *gatad2ab* on whole-body lipid

**Supplementary Table 37.**
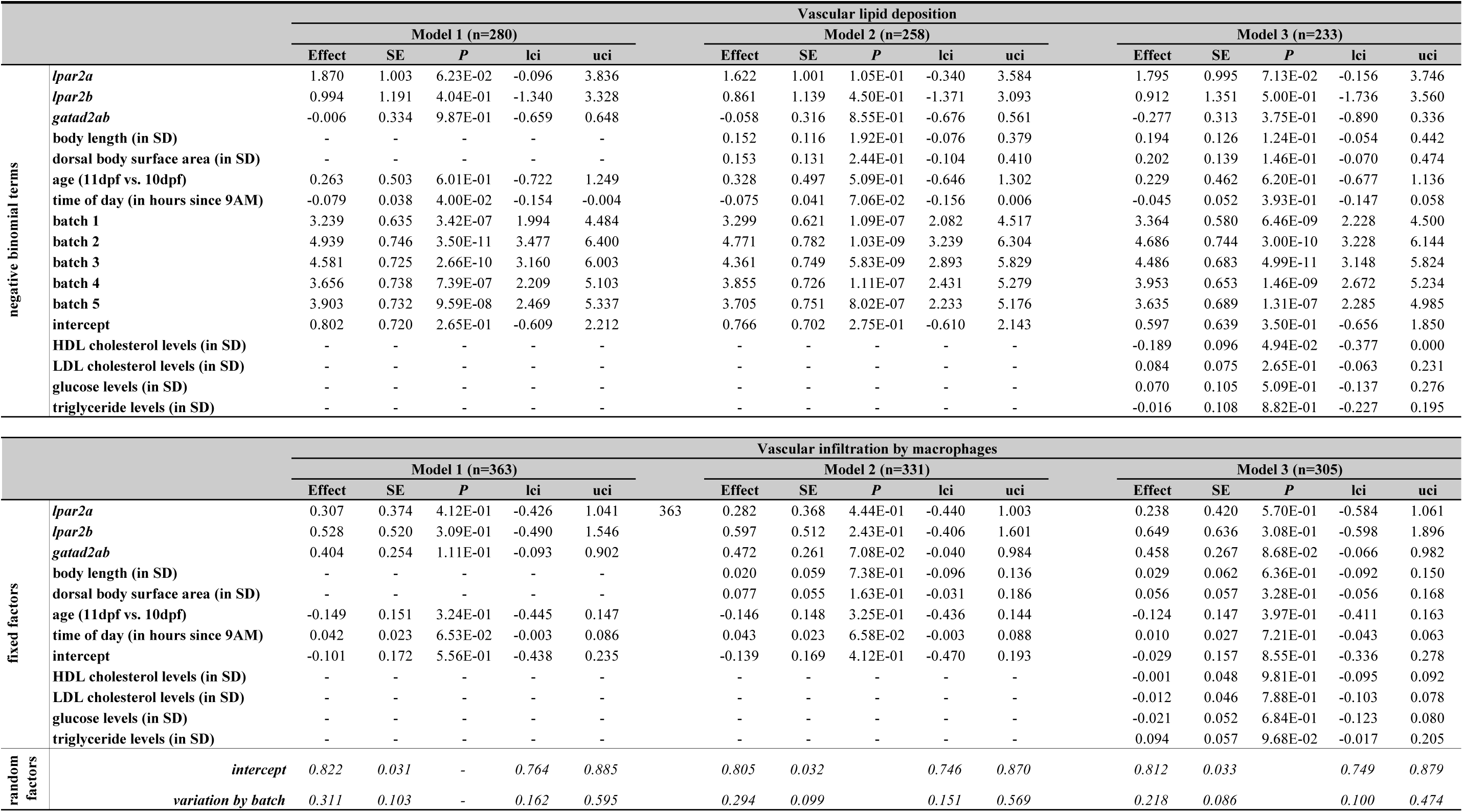
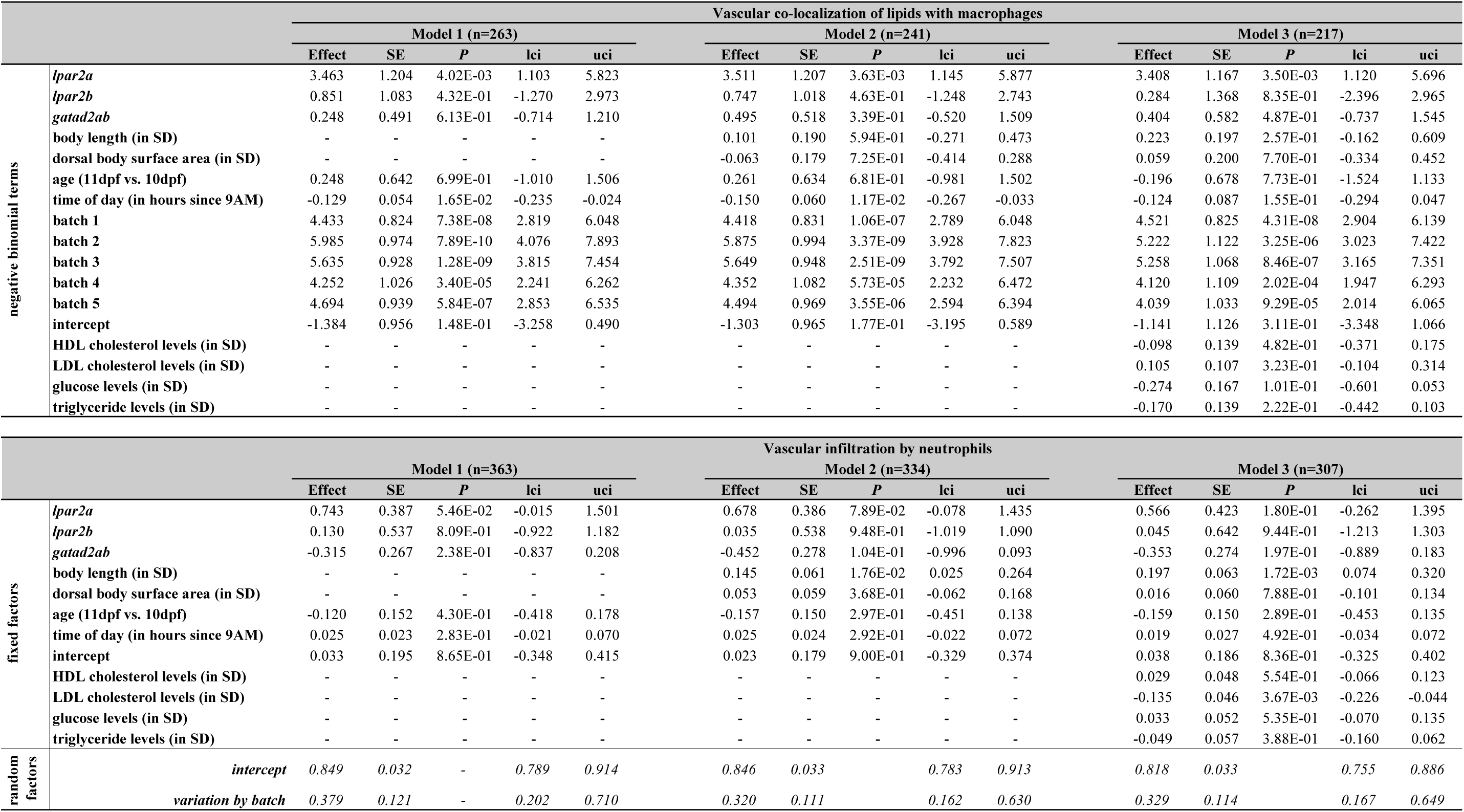
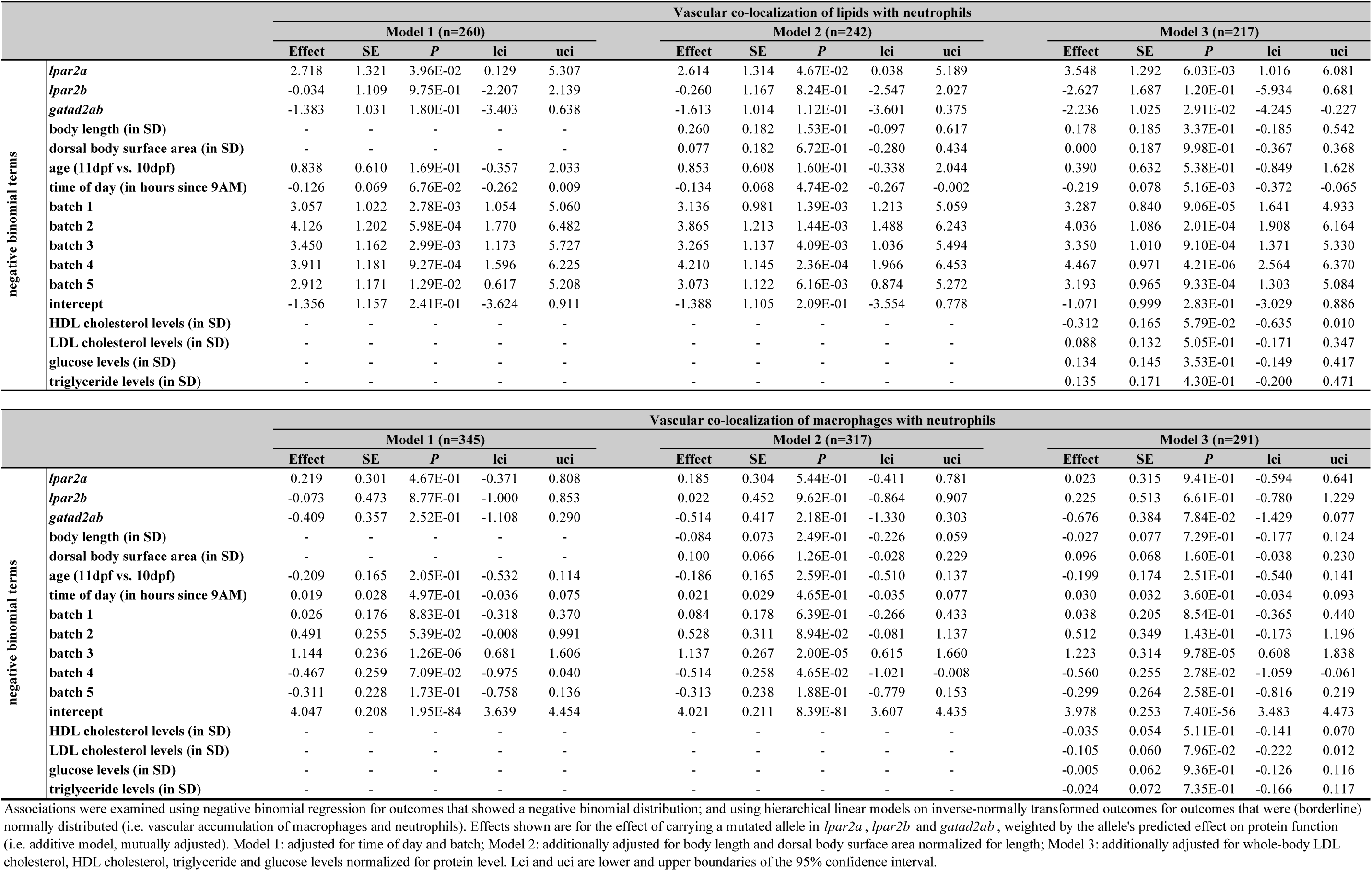
The effect of a mutated allele in *lpar2a, lpar2b* and *gatad2ab* on vascular atherogenic traits

**Supplementary Table 38.**
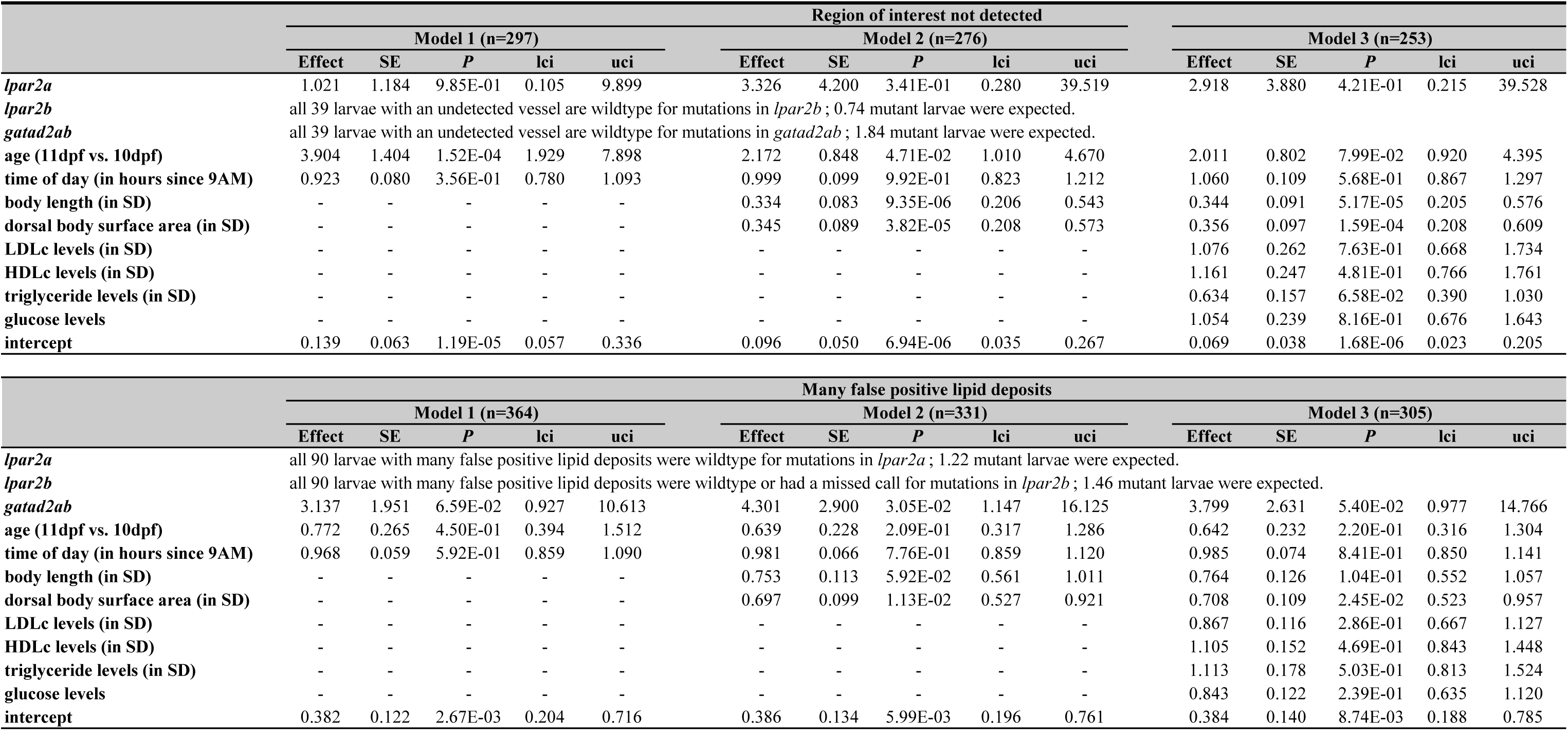
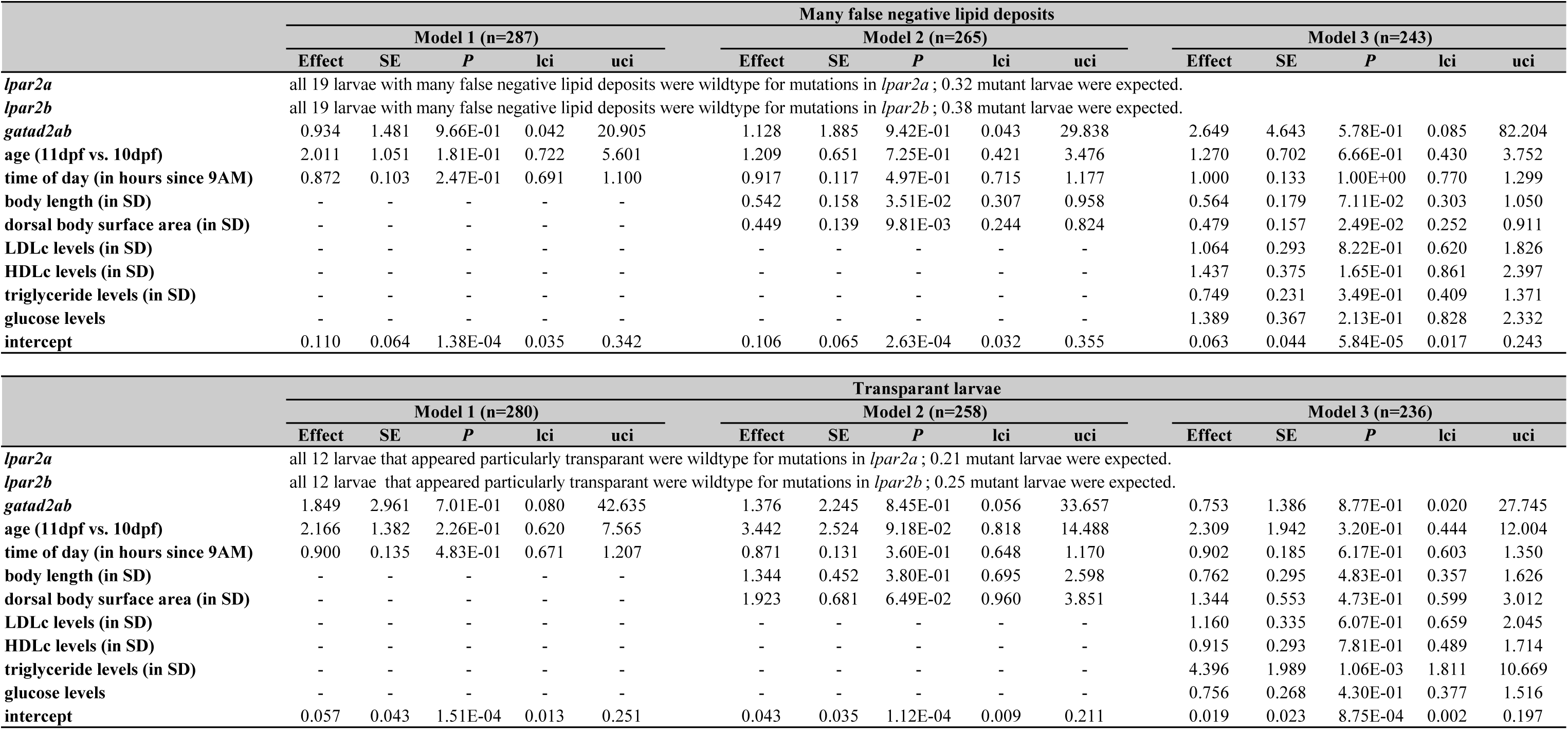
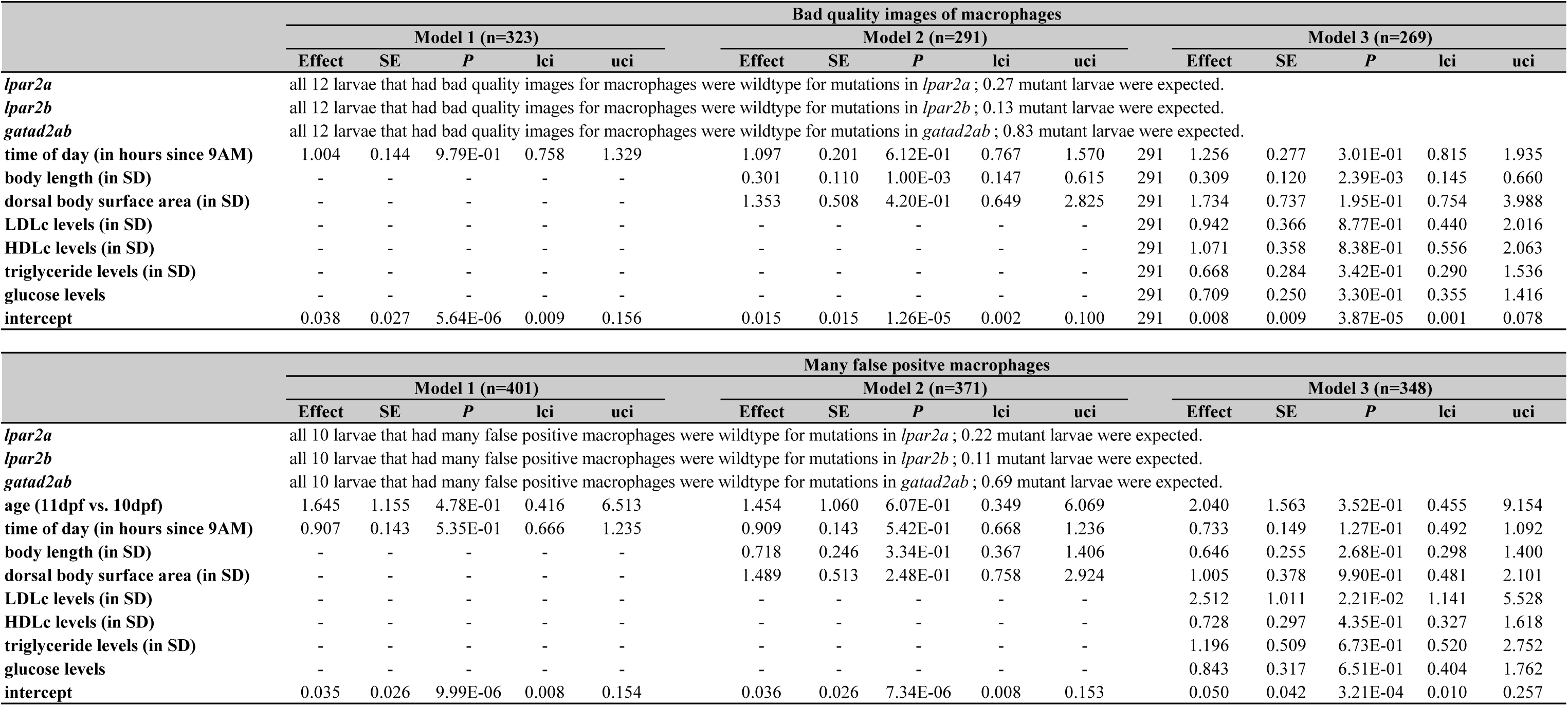
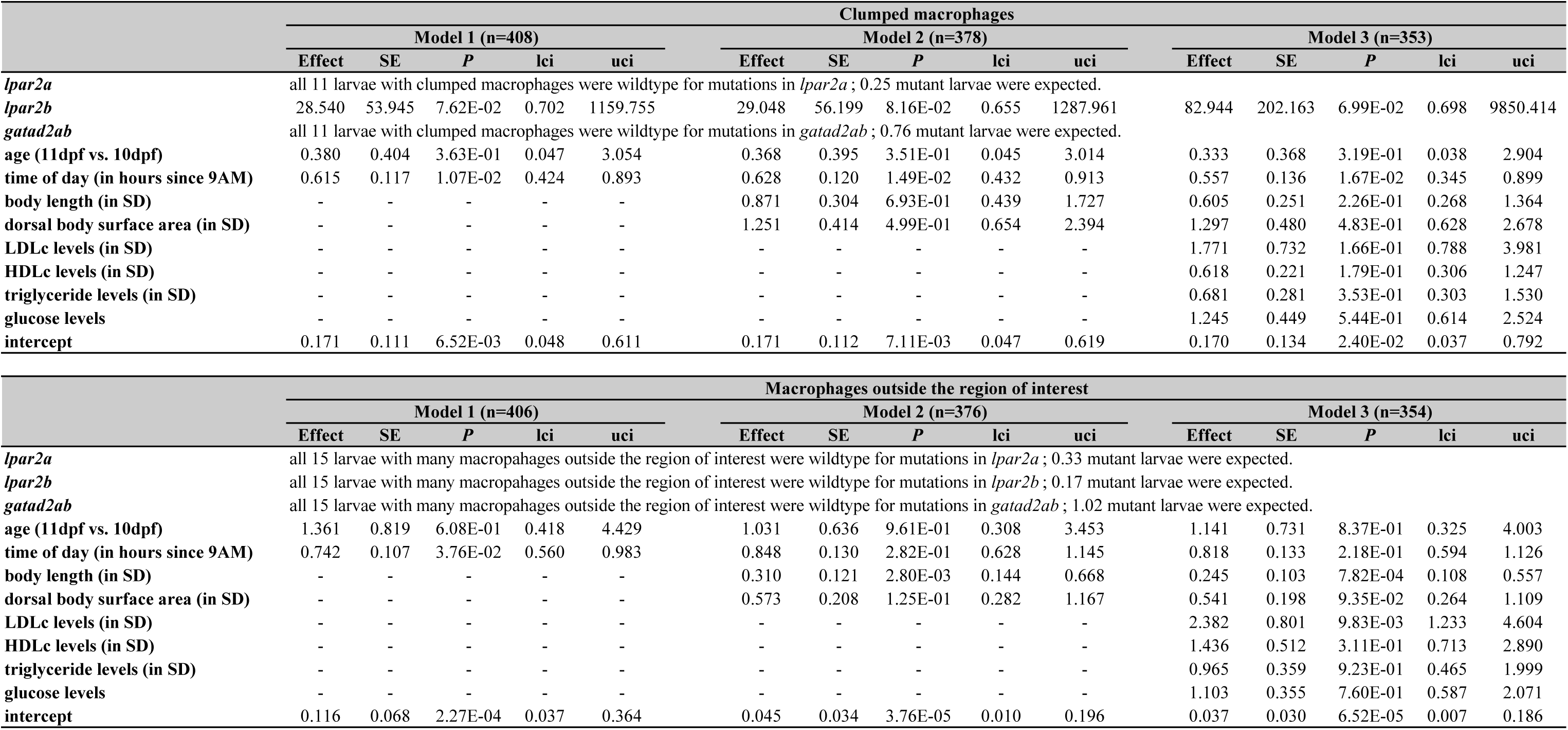
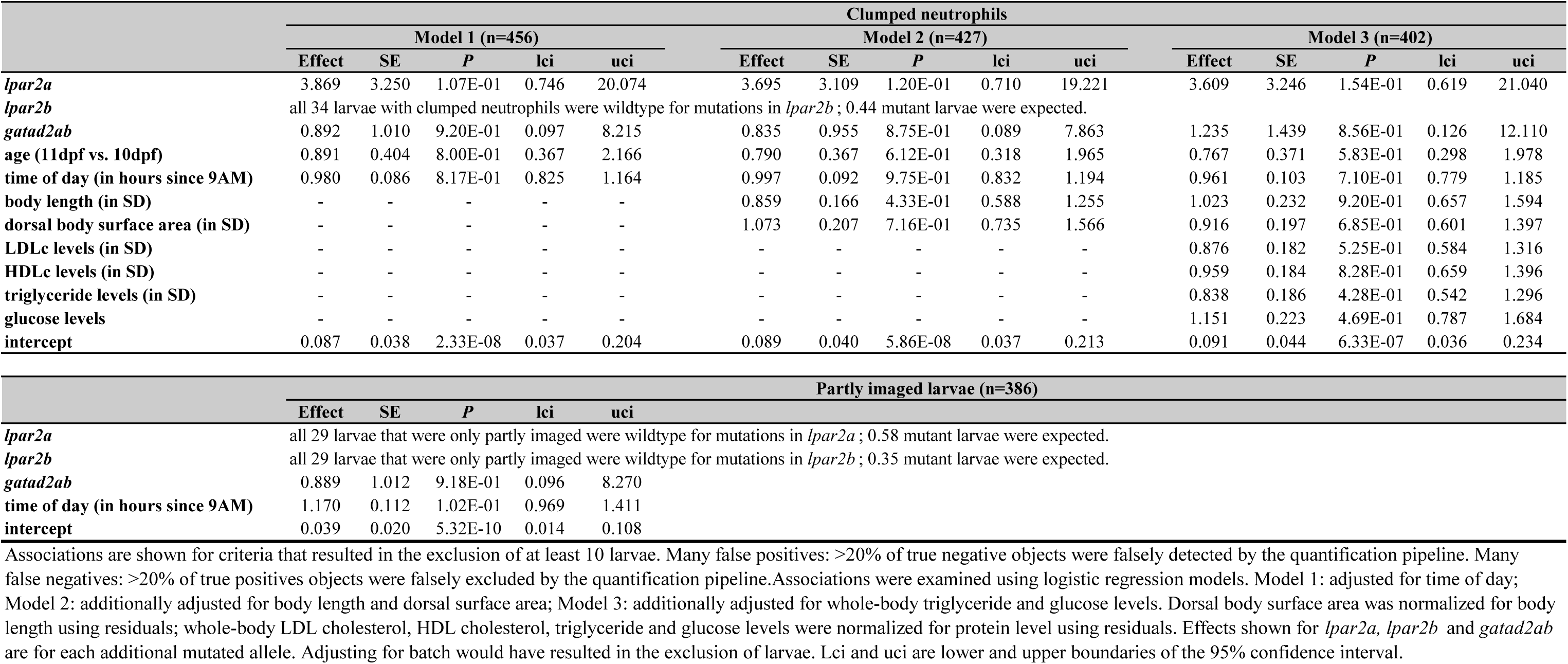
The effect of a mutated allele in *lpar2a, lpar2b* and *gatad2ab* on image and image quantification quality

