## Supplementary Figure 1 for "Zebrafish larvae as a model system for systematic characterization of drugs and genes in dyslipidemia and atherosclerosis"

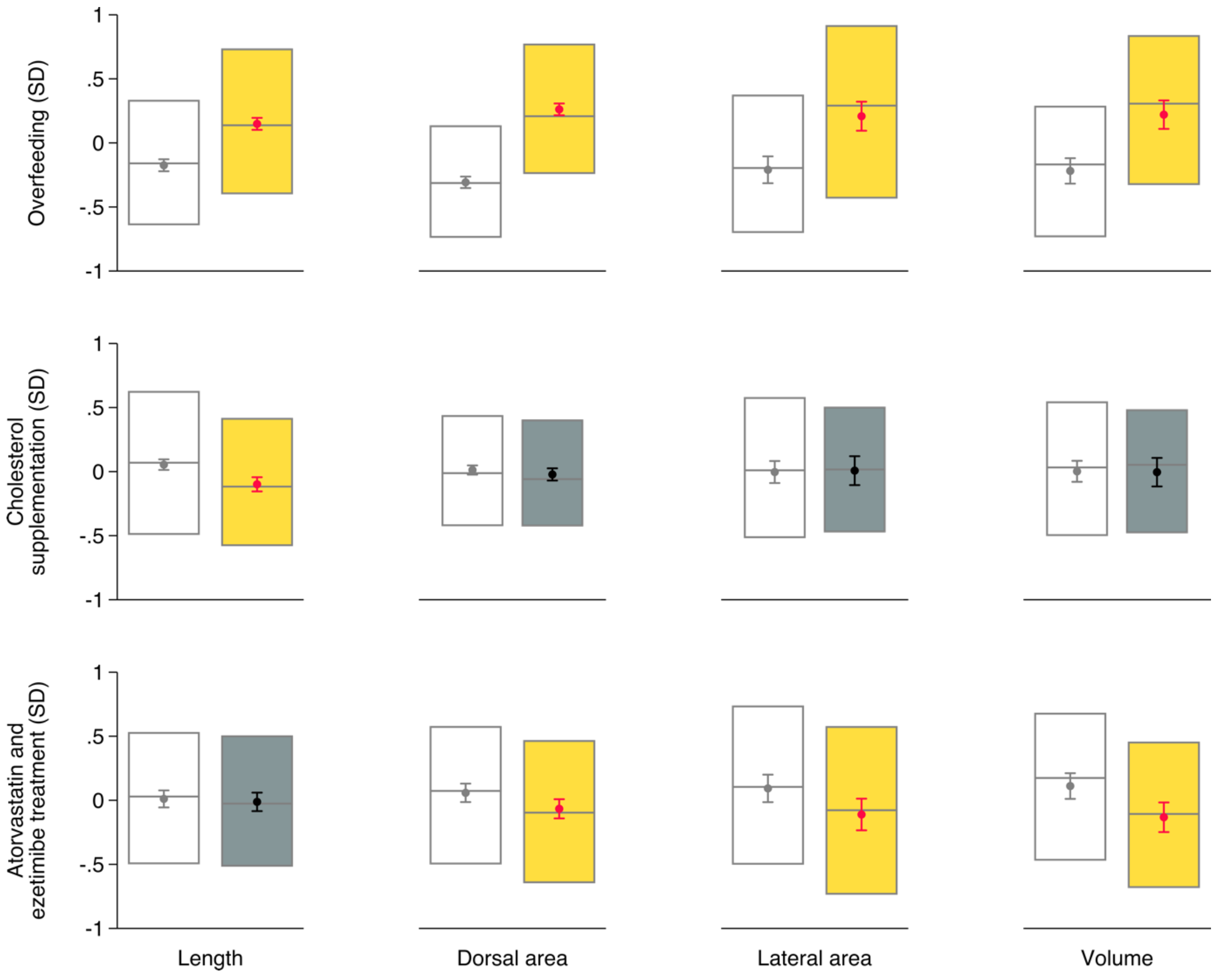


Supplementary Figure 1. The effect of overfeeding (top), cholesterol supplementation (middle) and treatment with atorvastatin and ezetimibe (bottom) on body size. Dots and whiskers show mean and 95% confidence interval (CI); boxes show median and inter quantile range. Analyses were performed using residuals acquired using hierarchical linear models on inverse-normally transformed outcomes, adjusted for the use of diethyl ether (for overfeeding and cholesterol supplementation), cholesterol supplementation (for overfeeding), the amount fed (for cholesterol supplementation), and time of day as fixed factors. Larvae were nested in batches and transgenic backgrounds (random factors). White boxes with grey mean and 95% CI (left) show results for unexposed larvae; grey boxes with black mean and 95% CI (right) show results for exposed larvae that are not different from unexposed ones; yellow boxes with red mean and 95% CI (right) show results for exposed larvae that are different from unexposed ones at $P<0.05$.
