## Supplementary Figure 3 for "Zebrafish larvae as a model system for systematic characterization of drugs and genes in dyslipidemia and atherosclerosis"

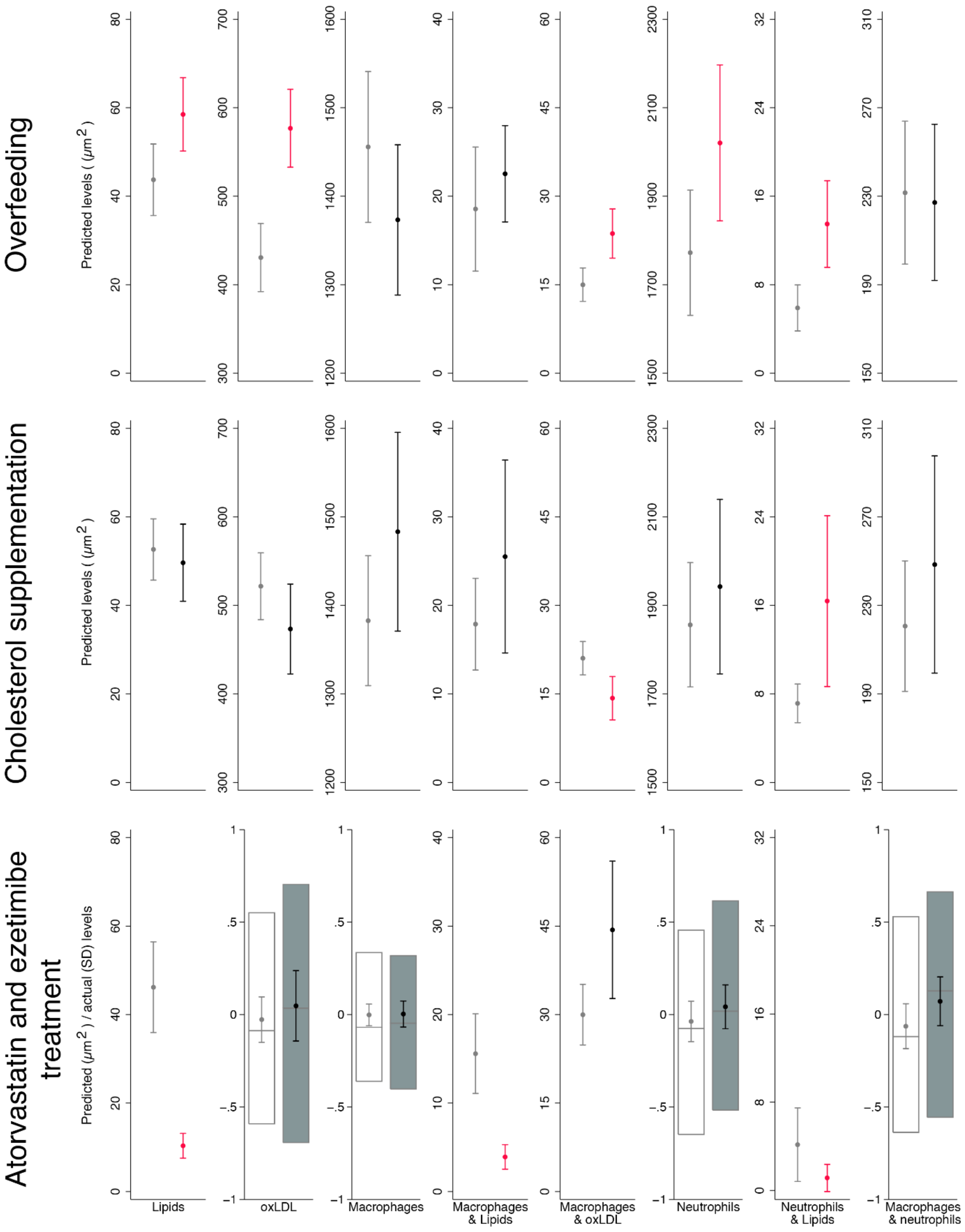


Supplementary Figure 3. The effect of overfeeding (top), cholesterol supplementation (middle) and treatment with atorvastatin and ezetimibe (bottom) on vascular atherogenic traits. Outcomes showing effect estimate and 95% CI for predicted values have been analyzed using negative binomial regression, with adjustment for the same co-variables. Outcomes showing only effect estimate and 95% CI for predicted values have been analyzed using regular negative binomial regression, adjusting for the use of diethyl ether (for overfeeding and cholesterol supplementation), cholesterol supplementation (for overfeeding), the amount fed (for cholesterol supplementation), body length, dorsal body surface area, and time of day. Outcomes showing mean and 95% confidence interval (CI) as well as boxes for median and inter quartile range have been analyzed using hierarchical linear models on residuals after adjusting inverse-normally transformed outcomes for body length, dorsal body surface area, and time of day as fixed factors. Larvae were nested in batches and transgenic backgrounds (random factors). White boxes and/or light grey mean and 95% CI (left) show results for unexposed larvae; grey boxes and/or black mean and 95% CI (right) show results for exposed larvae that are not different from unexposed ones; red mean and 95% CI (right) show results for exposed larvae that are different from unexposed ones at $P<0.05$.
