## Supplementary Figure 4 for "Zebrafish larvae as a model system for systematic characterization of drugs and genes in dyslipidemia and atherosclerosis"

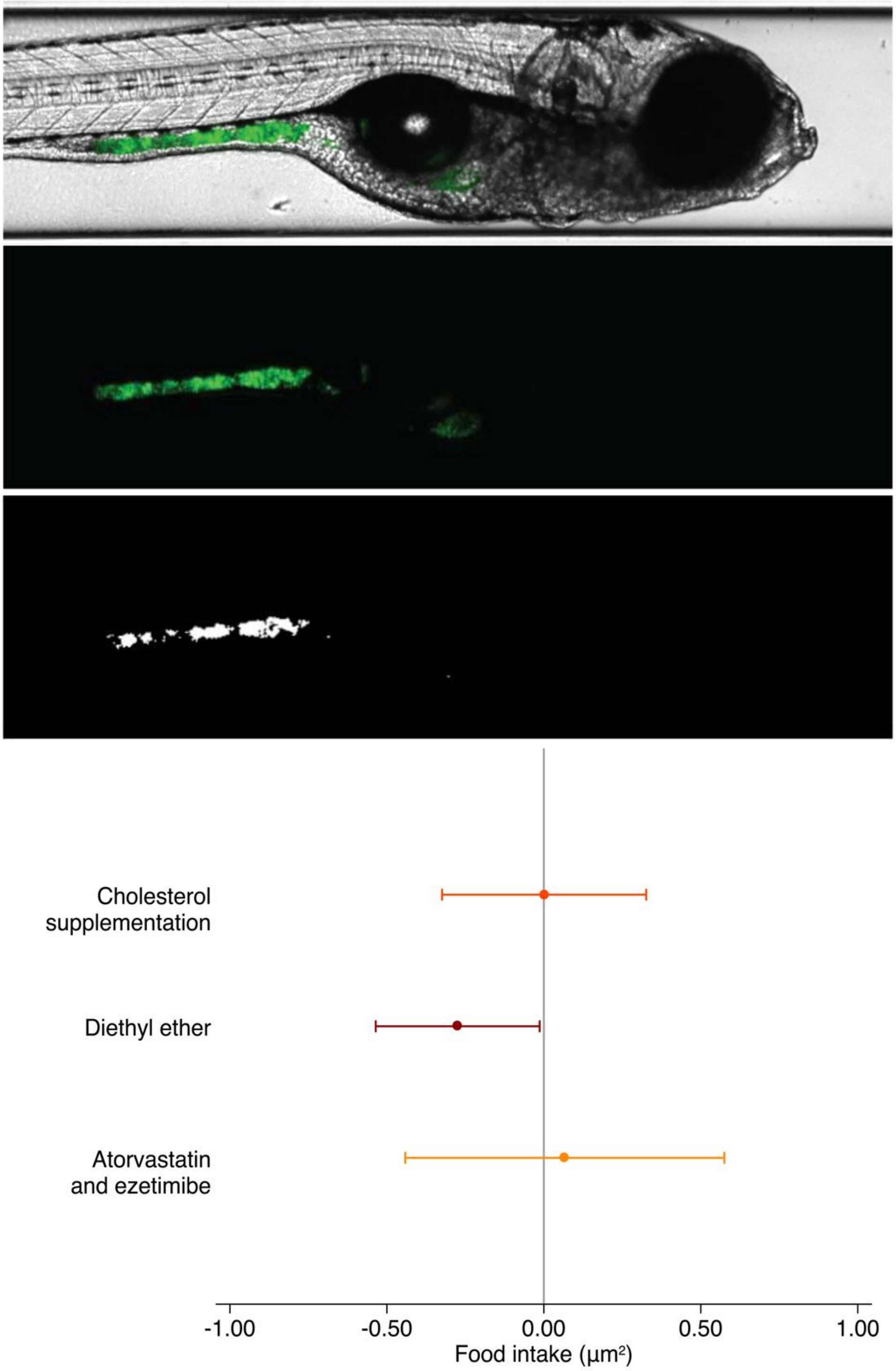


Supplementary Figure 4. Food intake as a function of dietary or drug treatment intervention. Mixing fluorescently labelled tracers in with standard dry food, standard dry food enriched with 4% extra cholesterol using diethyl ether, standard dry food treated with diethyl ether, and standard dry food enriched with 4% extra cholesterol using diethyl ether and further enriched with atorvastatin and ezetimibe allowed image-based quantification of food intake - i.e. surface area of fluorescence in the gastrointestinal tract - in eight-day-old zebrafish larvae (top). Bottom: mutually adjusted effect of cholesterol supplementation, treatment of the diet with diethyl ether, and enrichment with atorvastatin and ezetimibe on food intake, assessed using dummy variables and negative binomial regression, additionally adjusted for time since feeding and batch (n=204). Dots and whiskers show effect size and 95% confidence interval.
