## Supplementary Figure 5 for "Zebrafish larvae as a model system for systematic characterization of drugs and genes in dyslipidemia and atherosclerosis"

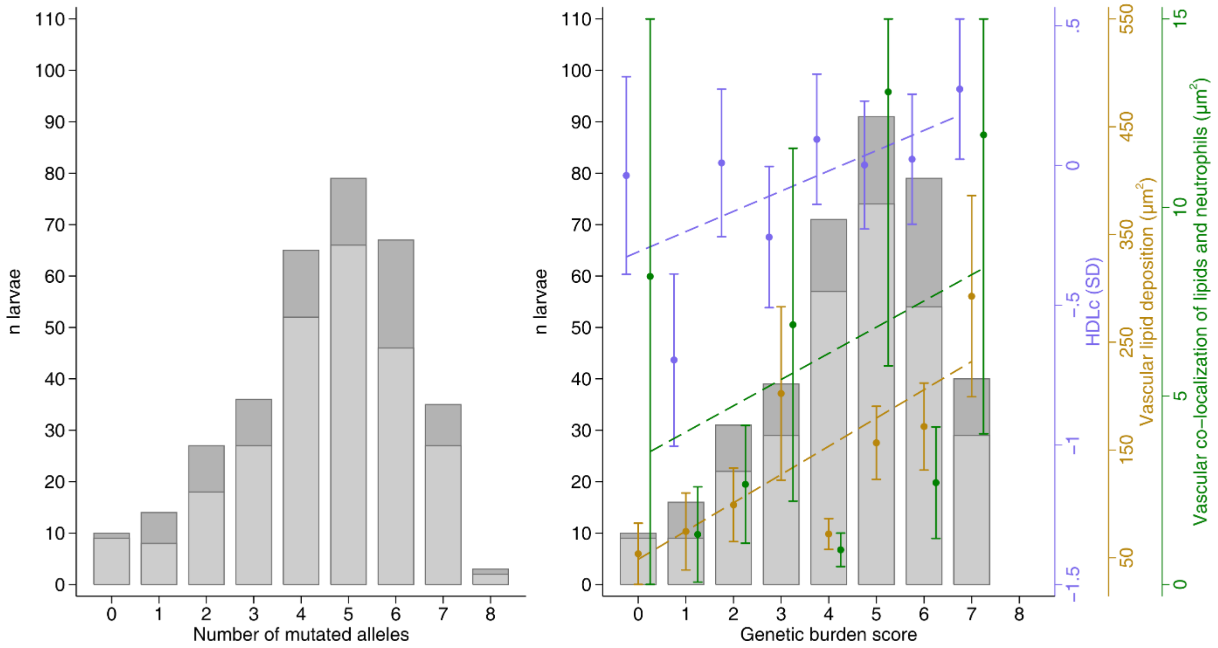


Supplementary Figure 5. Histogram of the number of mutated alleles and genetic burden score across apoea, apoeb, apobb.1 and ldlra and association of whole-body HDL cholesterol levels, vascular lipid deposition and vascular co-localization of lipids and neutrophils with the genetic burden score. Left: histogram of the number of mutated alleles across apoea, apoeb, apobb.1 and ldlra. Larvae with two mutated alleles in apoba, apobb.2 and ldlrb are shown in light grey (bottom); larvae with at least one unaffected allele in these three genes are shown in dark grey (top). Right: as before, but with each affected allele weighed by the probability that it affects protein function, based on annotation using Ensembl's variant effect predictor (VEP) (i.e. a genetic burden score). This figure also shows the association between atherogenic traits and the genetic burden score for significantly associated traits, adjusted for the number of mutated alleles in apoba, apobb.2 and ldlrb, i.e: 1) HDLc (n=381, in purple), assessed using a hierarchical linear model after inverse-normal transformation of LDLc, adjusted for time of day (fixed factors) and with larvae nested in batches; 2) vascular lipid deposition (n=272, in yellow); and 3) vascular co-localization of lipids and neutrophils (n=271, in green), using negative binomial regression, adjusted for body length, dorsal body surface area, time of day and batch. Dots and whiskers show mean and standard error of the mean, acquired using the margins command.
