## Supplementary Figure 6 for "Zebrafish larvae as a model system for systematic characterization of drugs and genes in dyslipidemia and atherosclerosis"

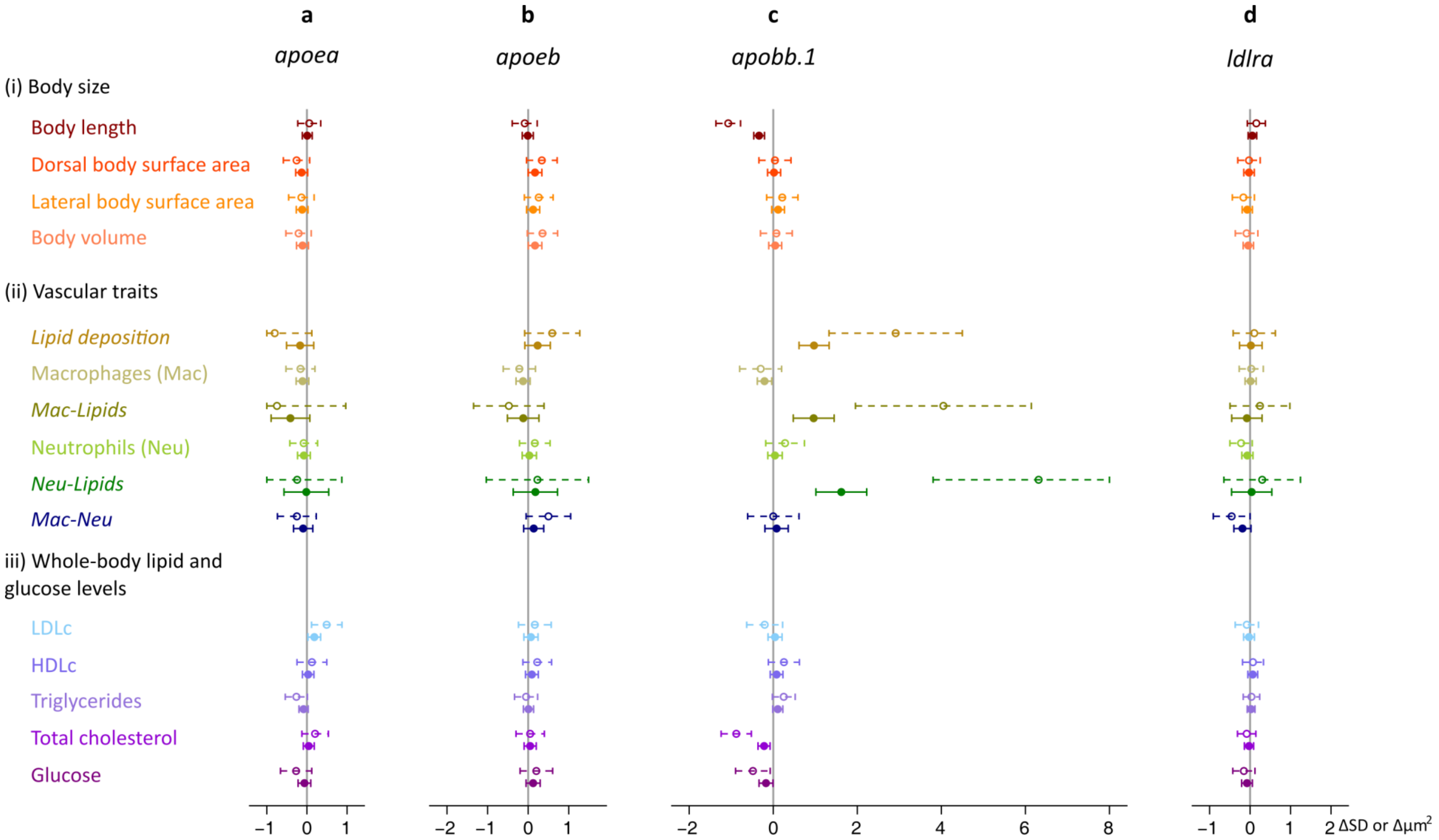


Supplementary Figure 6. The effect of mutations in apoea, apoeb, apobb.1 and ldlra on body size (i), vascular atherogenic traits (ii) and whole-body lipid and glucose levels (iii). Dorsal and lateral body surface area and body volume were normalized for body length before the analysis; and whole-body lipid and glucose levels were normalized for protein levels. For normally distributed traits (shown in regular font), associations were examined using hierarchical linear models on inverse-normally transformed values. For these traits effect sizes and 95% confidence intervals are expressed in standard deviation units (SD). The remaining vascular atherogenic traits (shown in italic) were analyzed using negative binomial regression analyses. For these traits, effect sizes and 95% confidence intervals are expressed in $\mu m^2$. Dotted lines represent the effect of two functionally knocked out alleles compared with zero mutated alleles. Regular lines show the additive per-allele effect. Associations were adjusted for time of day; batch; body length and dorsal body surface area (for vascular outcomes); and the number of mutated alleles in the other genes.
