## Supplementary Figure 7 for "Zebrafish larvae as a model system for systematic characterization of drugs and genes in dyslipidemia and atherosclerosis"

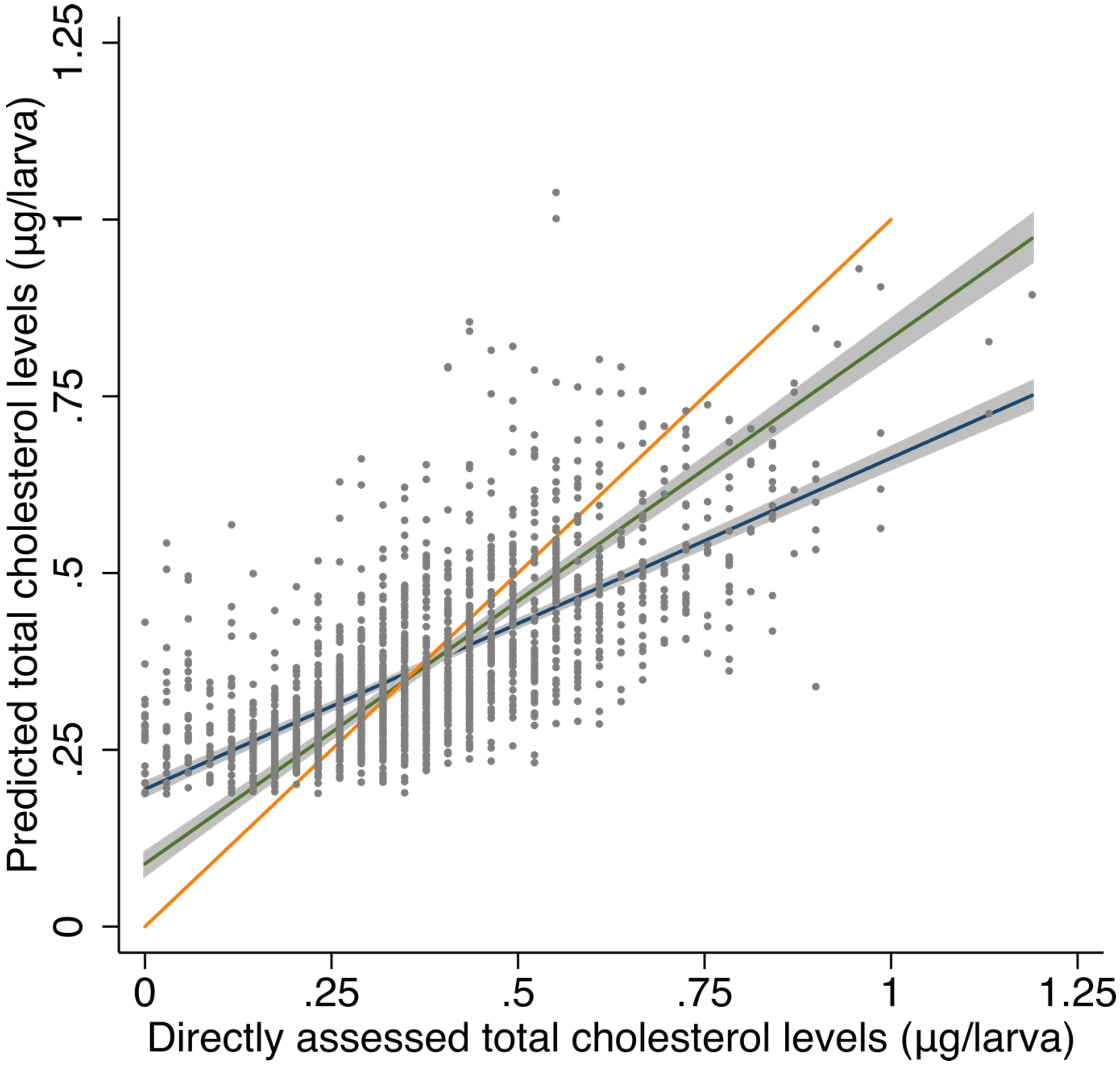


Supplementary Figure 7. The association of predicted total cholesterol levels using regression of directly assessed LDLc, HDLc and triglyceride levels with directly assessed total cholesterol levels. In blue and grey are the regression line and 95% confidence interval (CI) ($r^2$=0.468). In green and grey are the regression line and 95% CI for the association of total cholesterol levels calculated using the formula that is typically applied in humans (i.e. LDLc + HDLc + triglycerides/5) with directly assessed total cholesterol levels ($r^2$=0.430). In orange is a line with a slope of 1 (n=1,867 larvae).
