## Supplementary Figure 8 for "Zebrafish larvae as a model system for systematic characterization of drugs and genes in dyslipidemia and atherosclerosis"

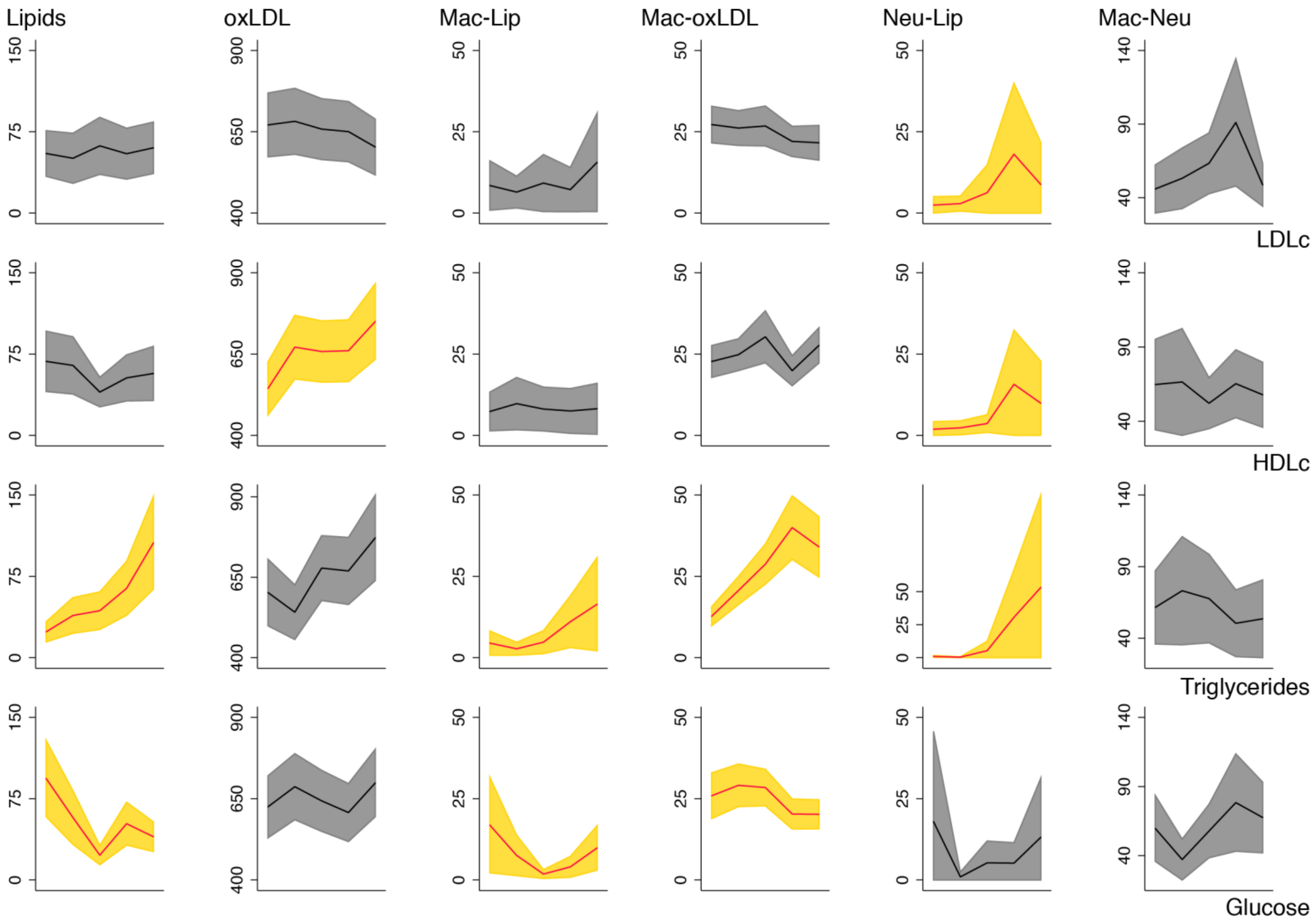


Supplementary Figure 8. The association of vascular atherogenic traits with whole-body lipid and glucose levels in data from the dietary, drug treatment and genetic intervention for proof-of-concept genes combined. For each vascular atherogenic outcome (i.e. vascular lipid deposition [Lip] and accumulation of oxidized LDL [oxLDL]; and vascular co-localization of lipids and oxLDL with macrophages [Mac] and neutrophils [Neu]), mutually adjusted associations with protein-normalized levels of LDL cholesterol (LDLc), HDL cholesterol (HDLc), triglyceride and glucose levels were examined using negative binomial regression. Besides for the other main exposures, associations were adjusted for body length and dorsal body surface area, transgenic background and batch. Graphs show margins plots – highlighting mean and 95% confidence intervals – for the vascular atherogenic outcomes, expressed in $\mu m^2$ (y-axes) with exposures grouped by quintile (x-axes). Significant associations are shown in yellow.
