## Supplementary Tables 1-38 for "Zebrafish larvae as a model system for systematic characterization of drugs and genes in dyslipidemia and atherosclerosis"

**Supplementary Table 1 - Descriptive information for larvae at 10 days post-fertilization in the dietary, drug treatment and genetic interventions**

| | | Dietary intervention | | | Drug treatment intervention | | | Genetic intervention | | | | | |
|---|---|---|---|---|---|---|---|---|---|---|---|---|---|
| | | | | | | | | Proof of concept | | | Discovery | | |
| | **Transgenic background(s)** | $n_{total}$ | **Mean / Median** | **SD / IQR** | $n_{total}$ | **Mean / Median** | **SD / IQR** | $n_{total}$ | **Mean / Median** | **SD / IQR** | $n_{total}$ | **Mean / Median** | **SD / IQR** |
| ***Body size*** | | | | | | | | | | | | | |
| Body length (μm) | | 2193 | 4327 | 260 | 1004 | 4343 | 234 | 339 | 4628 | 266 | 505 | 4451 | 230 |
| Dorsal surface area (μm$^2$) | | 2193 | $1.1\times10^6$ | $1.6\times10^5$ | 1004 | $1.2\times10^6$ | $1.2\times10^5$ | 339 | $1.2\times10^6$ | $1.7\times10^5$ | 505 | $1.1\times10^6$ | $1.6\times10^5$ |
| Lateral surface area (μm$^2$) | | 524 | $1.5\times10^6$ | $1.6\times10^5$ | 553 | $1.4\times10^6$ | $1.5\times10^5$ | 336 | $1.6\times10^6$ | $2.0\times10^5$ | 502 | $1.5\times10^6$ | $1.9\times10^5$ |
| Body volume (μm$^3$) | | 514 | $4.3\times10^8$ | $6.1\times10^7$ | 512 | $4.3\times10^8$ | $6.0\times10^7$ | 328 | $4.6\times10^8$ | $8.4\times10^7$ | 495 | $4.0\times10^8$ | $8.1\times10^7$ |
| ***Whole-body lipid and glucose levels*** | | | | | | | | | | | | | |
| LDL cholesterol (μg) | | 564 | 0.04 | 0.04 | 567 | 0.12 | 0.08 | 339 | 0.18 | 0.12 | 513 | 0.49 | 0.34 |
| HDL cholesterol (μg) | | 549 | 0.06 | 0.02 | 564 | 0.06 | 0.04 | 339 | 0.15 | 0.05 | 513 | 0.29 | 0.16 |
| Triglyceride levels (μg) | | 2123 | 0.93 | 0.80 | 1005 | 0.61 | 0.51 | 339 | 1.10 | 0.82 | 513 | 1.27 | 1.07 |
| Total cholesterol levels (μg) | | 2123 | 0.29 | 0.17 | 1005 | 0.36 | 0.18 | 339 | 0.45 | 0.21 | 513 | 0.29 | 0.14 |
| Glucose (μg) | | 2128 | 0.92 | 0.66 | 1008 | 0.38 | 0.42 | 339 | 0.30 | 0.16 | 513 | 0.80 | 0.53 |
| ***Vascular atherogenic traits*** | | | | | | | | | | | | | |
| Lipid deposition (μm$^2$) | - | *1954* | *0* | *36* | *837* | *0* | *20* | *272* | *43* | *133* | *280* | *94* | *135* |
| oxLDL deposition (μm$^2$) | *Tg:hsp70:IK17-EGFP* | *885* | *338* | *590* | 236 | 852 | 674 | - | - | - | - | - | - |
| Infiltration by macrophages (μm$^2$) | *Tg:mpeg1-mCherry* | *994* | *942* | *1607* | 633 | 1881 | 1057 | 328 | 1475 | 1005 | 363 | 1705 | 729 |
| Infiltration by neutrophils (μm$^2$) | *Tg:mpo-EGFP* | *494* | *1545* | *2116* | 404 | 2438 | 873 | 330 | 1050 | 911 | 363 | 1806 | 941 |
| Co-localizing macrophages and lipids (μm$^2$) | *Tg:mpeg1-mCherry* | *917* | *0* | *10* | *605* | *0* | *0* | *269* | *1* | *13* | *263* | *23* | *48* |
| Co-localizing macrophages and oxLDL (μm$^2$) | *Tg:hsp70:IK17-EGFP & Tg:mpeg1-mCherry* | *433* | *10* | *22* | *212* | *20* | *39* | - | - | - | - | - | - |
| Co-localizing neutrophils and lipids (μm$^2$) | *Tg:mpo-EGFP* | *440* | *0* | *8* | *393* | *0* | *0* | *271* | *0* | *0* | *260* | *5* | *12* |
| Co-localizing macrophages and neutrophils (μm$^2$) | *Tg:mpeg1-mCherry & Tg:mpo-EGFP* | *488* | *140* | *306* | 394 | 139 | 124 | *327* | *25* | *68* | *345* | *84* | *122* |
| Circulating lipids (μm$^2$) | *Tg:flk-EGFP* | *467* | $2.4\times10^4$ | $3.5\times10^3$ | 185 | 17,829 | 5767 | - | - | - | - | - | - |
| Endothelial surface area (μm$^2$) | *Tg:flk-EGFP* | *467* | $4.8\times10^3$ | $2.1\times10^3$ | 185 | 6652 | 4198 | - | - | - | - | - | - |

oxLDL: oxidized LDL; co-localization defined by overlap of fluorescence signal. Mean and standard deviation (SD) are shown for normally distributed outcomes; median and interquatrile range (IQR) are shown for outcomes showing a negative binomial distribution (shown in italics).

**Supplementary Table 2 - Annotation-based exclusions in the image-based analyses**

| Rationale for exclusion | Traits for which exclusion is relevant | Dietary Intervention | | | Drug treatent intervention | | | Genetic intervention | | | | | |
|---|---|---|---|---|---|---|---|---|---|---|---|---|---|
| | | | | | | | | Proof-of-concept | | | Discovery (19p13.11 locus) | | |
| | | Available (n) | Excluded (n) | Excluded (%) | Available (n) | Excluded (n) | Excluded (%) | Available (n) | Excluded (n) | Excluded (%) | Available (n) | Excluded (n) | Excluded (%) |
| ***Monodansylpentane cadaverase*** | | | | | | | | | | | | | |
| Inadequate detection of vasculature in Y (vessel missing) | Lipids, macrophages, neutrophils and their co-localization | 2050 | 94 | 4.6 | 927 | 92 | 9.9 | 231 | 6 | 2.6 | 212 | 30 | 14.2 |
| Fish moved during imaging resulting in a bad quality image | Lipids | 1959 | 3 | 0.2 | 873 | 38 | 4.4 | 225 | 0 | 0.0 | 183 | 1 | 0.5 |
| >20% of true negative objects falsely detected (many false positives) | Lipids | 2043 | 87 | 4.3 | 853 | 18 | 2.1 | 278 | 48 | 17.3 | 219 | 37 | 16.9 |
| <20% of true positive objects detected (many false negatives) | Lipids | 1959 | 3 | 0.2 | 844 | 9 | 1.1 | 225 | 0 | 0.0 | 201 | 19 | 9.5 |
| ***Tg(IK17:EGFP)*** | | | | | | | | | | | | | |
| Fish moved during imaging resulting in a bad quality image | oxLDL | 893 | 8 | 0.9 | 238 | 2 | 0.8 | | NA | | | NA | |
| >20% of true negative objects falsely detected (many false positives) | No exclusion | 885 | 202 | 22.8 | 236 | 100 | 42.4 | | NA | | | NA | |
| <20% of true positive objects detected (many false negatives) | No exclusion | 885 | 1 | 0.1 | 236 | 0 | 0.0 | | NA | | | NA | |
| Many false positives outside the area of interest | No exclusion | 885 | 3 | 0.3 | 236 | 4 | 1.7 | | NA | | | NA | |
| ***Tg(mpeg1:mCherry)*** | | | | | | | | | | | | | |
| Fish moved during imaging resulting in a bad quality image | Macrophages and their co-localization with lipids and oxLDL | 996 | 2 | 0.2 | 634 | 1 | 0.2 | 276 | 0 | 0.0 | 299 | 3 | 1.0 |
| >20% of true negative objects falsely detected (many false positives) | Macrophages and their co-localization with lipids and oxLDL | 997 | 3 | 0.3 | 635 | 2 | 0.3 | 278 | 2 | 0.7 | 307 | 11 | 3.6 |
| <20% of true positive objects detected (many false negatives) | Macrophages and their co-localization with lipids and oxLDL | 1010 | 16 | 1.6 | 633 | 0 | 0.0 | 276 | 0 | 0.0 | 301 | 5 | 1.7 |
| Presence of (a) moving macrophage(s) | Macrophages and their co-localization with lipids and oxLDL | 994 | 0 | 0.0 | 634 | 1 | 0.2 | 276 | 0 | 0.0 | 298 | 2 | 0.7 |
| Many false positive macrophages outside the area of interest | Macrophages and their co-localization with lipids and oxLDL | 996 | 2 | 0.2 | 634 | 1 | 0.2 | 276 | 0 | 0.0 | 311 | 15 | 4.8 |
| Many macrophages co-localizing | | 994 | 0 | 0.0 | 633 | 0 | 0.0 | 276 | 0 | 0.0 | 307 | 11 | 3.6 |
| ***Tg(mpo:EGFP)*** | | | | | | | | | | | | | |
| <20% of true positive objects detected (many false negatives) | Neutrophils and their co-localization with lipids | 537 | 43 | 8.0 | 404 | 0 | 0.0 | 278 | 0 | 0.0 | 299 | 1 | 0.3 |
| Presence of circulating neutrophils | Neutrophils and their co-localization with lipids | 504 | 10 | 2.0 | 409 | 5 | 1.2 | 278 | 0 | 0.0 | 301 | 3 | 1.0 |
| Many neutrophils co-localizing | | 494 | 0 | 0.0 | 404 | 0 | 0.0 | 278 | 0 | 0.0 | 332 | 34 | 10.2 |
| ***Bright field*** | | | | | | | | | | | | | |
| Debris included in the segmentation | Body size | 2194 | 1 | 0.0 | 1004 | 0 | 0.0 | 279 | 23 | 8.2 | 344 | 0 | 0.0 |
| Air bubble included in the segmentation | Body size | 2197 | 4 | 0.2 | 1004 | 0 | 0.0 | 256 | 0 | 0.0 | 345 | 1 | 0.3 |
| Bad segmentation | Body size | 2193 | 0 | 0.0 | 1004 | 0 | 0.0 | 257 | 1 | 0.4 | 344 | 0 | 0.0 |
| Part of the fish not imaged | Body size | 2202 | 9 | 0.4 | 1004 | 0 | 0.0 | 260 | 4 | 1.5 | 344 | 0 | 0.0 |
| The fish has a curved body, resulting in a non representative length | Body size | 2197 | 4 | 0.2 | 1004 | 0 | 0.0 | 256 | 0 | 0.0 | 345 | 1 | 0.3 |
| Larvae optically cut off during preprocessing | Body size | 2306 | 113 | 4.9 | 1004 | 0 | 0.0 | 256 | 0 | 0.0 | 344 | 0 | 0.0 |

Available (n): the number of larvae with a value within mean ± 5xSD that are free from the image quality-based annotation of interest plus the number of larvae that were excluded due to the quality-based annotation of interest for vascular infiltration by lipids (monodansylpentane cadaverase), oxidized LDL (Tg([IK17:EGFP]), macrophages (Tg[mpeg1:mCherry]), neutrophils (Tg[mpo:EGFP]), and body size (Bright field). For Tg(IK17:EGFP), larvae with many false positive or negative deposits were not excluded from further analysis. For all other traits, larvae with suboptimal image or quantification quality were excluded from further analyses.

**Supplementary Table 3 - The effect of overfeeding and cholesterol supplementation on body size**

| | | Body length (n=2193) | | | | |
|---|---|---|---|---|---|---|
| | | **Effect** | **SE** | ***P*** | **lci** | **uci** |
| **fixed factors** | **overfeeding** | 0.350 | 0.040 | 9.15E-23 | 0.280 | 0.420 |
| | **cholesterol supplementation** | -0.220 | 0.040 | 2.32E-07 | -0.300 | -0.140 |
| | **diethyl ether treatment** | -0.070 | 0.040 | 9.57E-02 | -0.160 | 0.010 |
| | **time of day (in hours since 9AM)** | 0.010 | 0.010 | 1.28E-01 | 0.000 | 0.030 |
| | **intercept** | -0.150 | 0.210 | 4.80E-01 | -0.550 | 0.260 |
| **random factors** | ***variation by transgenic background*** | *0.380* | *0.150* | - | *0.170* | *0.830* |
| | ***variation by batch*** | *0.350* | *0.060* | - | *0.250* | *0.480* |
| | ***residual*** | *0.790* | *0.010* | - | *0.770* | *0.820* |

| | | Dorsal body surface area (n=2193) | | | | |
|---|---|---|---|---|---|---|
| | | **Effect** | **SE** | ***P*** | **lci** | **uci** |
| **fixed factors** | **overfeeding** | 0.610 | 0.030 | 3.54E-71 | 0.540 | 0.680 |
| | **cholesterol supplementation** | -0.050 | 0.040 | 1.82E-01 | -0.140 | 0.030 |
| | **diethyl ether treatment** | 0.040 | 0.040 | 3.84E-01 | -0.050 | 0.120 |
| | **time of day (in hours since 9AM)** | 0.040 | 0.010 | 6.82E-08 | 0.020 | 0.050 |
| | **intercept** | -0.500 | 0.210 | 1.58E-02 | -0.910 | -0.090 |
| **random factors** | ***variation by transgenic background*** | *0.360* | *0.160* | - | *0.150* | *0.870* |
| | ***variation by batch*** | *0.420* | *0.070* | - | *0.310* | *0.580* |
| | ***residual*** | *0.770* | *0.010* | - | *0.750* | *0.790* |

| | | Lateral body surface area (n=524) | | | | |
|---|---|---|---|---|---|---|
| | | **Effect** | **SE** | ***P*** | **lci** | **uci** |
| **fixed factors** | **overfeeding** | 0.480 | 0.080 | 1.17E-08 | 0.320 | 0.650 |
| | **cholesterol supplementation** | 0.020 | 0.100 | 8.69E-01 | -0.170 | 0.210 |
| | **diethyl ether treatment** | 0.010 | 0.100 | 9.04E-01 | -0.180 | 0.200 |
| | **time of day (in hours since 9AM)** | 0.020 | 0.020 | 3.96E-01 | -0.020 | 0.050 |
| | **intercept** | -0.310 | 0.210 | 1.37E-01 | -0.720 | 0.100 |
| **random factors** | ***variation by transgenic background*** | - | - | - | - | - |
| | ***variation by batch*** | *0.390* | *0.120* | - | *0.220* | *0.720* |
| | ***residual*** | *0.900* | *0.030* | - | *0.850* | *0.960* |

| | | Body volume (n=514) | | | | |
|---|---|---|---|---|---|---|
| | | **Effect** | **SE** | ***P*** | **lci** | **uci** |
| **fixed factors** | **overfeeding** | 0.510 | 0.080 | 5.39E-10 | 0.350 | 0.660 |
| | **cholesterol supplementation** | -0.010 | 0.090 | 9.18E-01 | -0.190 | 0.170 |
| | **diethyl ether treatment** | -0.010 | 0.090 | 9.25E-01 | -0.190 | 0.170 |
| | **time of day (in hours since 9AM)** | 0.020 | 0.020 | 2.94E-01 | -0.020 | 0.060 |
| | **intercept** | -0.300 | 0.230 | 1.85E-01 | -0.750 | 0.150 |
| **random factors** | ***variation by transgenic background*** | - | - | - | - | - |
| | ***variation by batch*** | *0.460* | *0.140* | - | *0.260* | *0.830* |
| | ***residual*** | *0.860* | *0.030* | - | *0.810* | *0.910* |

Dorsal and lateral body surface area and body volume were normalized for body length using residuals. All outcomes were inverse-normally transformed before the analysis. Associations were examined using hierarchical linear models. Effects shown for overfeeding, cholesterol supplementation and diethyl ether treatment are compared with unexposed controls. Lci and uci are lower and upper boundaries of the 95% confidence interval.

**Supplementary Table 4 - The effect of overfeeding and cholesterol supplementation on whole-body lipid and glucose levels**

| | | LDL cholesterol levels (n=564) | | | | |
|---|---|---|---|---|---|---|
| | | **Effect** | **SE** | ***P*** | **lci** | **uci** |
| **fixed factors** | **overfeeding** | -0.072 | 0.086 | 4.05E-01 | -0.241 | 0.097 |
| | **cholesterol supplementation** | 0.422 | 0.098 | 1.68E-05 | 0.230 | 0.615 |
| | **diethyl ether treatment** | -0.126 | 0.099 | 2.00E-01 | -0.320 | 0.067 |
| | **time of day (in hours since 9AM)** | -0.006 | 0.019 | 7.69E-01 | -0.043 | 0.031 |
| | **intercept** | 0.041 | 0.172 | 8.10E-01 | -0.296 | 0.379 |
| **random factors** | ***variation by batch*** | *0.261* | *0.089* | - | *0.134* | *0.508* |
| | ***residual*** | *0.952* | *0.028* | - | *0.897* | *1.009* |

| | | HDL cholesterol levels (n=594) | | | | |
|---|---|---|---|---|---|---|
| | | **Effect** | **SE** | ***P*** | **lci** | **uci** |
| **fixed factors** | **overfeeding** | -0.058 | 0.083 | 4.83E-01 | -0.221 | 0.105 |
| | **cholesterol supplementation** | -0.170 | 0.094 | 7.01E-02 | -0.355 | 0.014 |
| | **diethyl ether treatment** | 0.012 | 0.095 | 8.99E-01 | -0.174 | 0.198 |
| | **time of day (in hours since 9AM)** | -0.041 | 0.018 | 2.18E-02 | -0.077 | -0.006 |
| | **intercept** | 0.358 | 0.236 | 1.29E-01 | -0.104 | 0.820 |
| **random factors** | ***variation by batch*** | *0.480* | *0.146* | - | *0.264* | *0.871* |
| | ***residual*** | *0.903* | *0.027* | - | *0.850* | *0.958* |

| | | Triglyceride levels (n=2123) | | | | |
|---|---|---|---|---|---|---|
| | | **Effect** | **SE** | ***P*** | **lci** | **uci** |
| **fixed factors** | **overfeeding** | 0.409 | 0.034 | 5.66E-34 | 0.343 | 0.475 |
| | **cholesterol supplementation** | -0.249 | 0.040 | 7.98E-10 | -0.328 | -0.169 |
| | **diethyl ether treatment** | 0.081 | 0.041 | 5.07E-02 | 0.000 | 0.162 |
| | **time of day (in hours since 9AM)** | -0.042 | 0.007 | 4.88E-10 | -0.056 | -0.029 |
| | **intercept** | 0.068 | 0.182 | 7.07E-01 | -0.288 | 0.425 |
| **random factors** | ***variation by transgenic background*** | *0.253* | *0.178* | - | *0.064* | *1.005* |
| | ***variation by batch*** | *0.574* | *0.096* | - | *0.414* | *0.795* |
| | ***residual*** | *0.750* | *0.012* | - | *0.727* | *0.773* |

| | | Total cholesterol levels (n=2123) | | | | |
|---|---|---|---|---|---|---|
| | | **Effect** | **SE** | ***P*** | **lci** | **uci** |
| **fixed factors** | **overfeeding** | 0.256 | 0.032 | 5.93E-16 | 0.194 | 0.318 |
| | **cholesterol supplementation** | 0.193 | 0.038 | 4.13E-07 | 0.118 | 0.267 |
| | **diethyl ether treatment** | 0.275 | 0.039 | 1.54E-12 | 0.199 | 0.351 |
| | **time of day (in hours since 9AM)** | -0.060 | 0.006 | 1.46E-20 | -0.072 | -0.047 |
| | **intercept** | -0.119 | 0.275 | 6.65E-01 | -0.658 | 0.420 |
| **random factors** | ***variation by transgenic background*** | *0.505* | *0.208* | - | *0.226* | *1.133* |
| | ***variation by batch*** | *0.459* | *0.077* | - | *0.331* | *0.637* |
| | ***residual*** | *0.706* | *0.011* | - | *0.685* | *0.727* |

| | | Glucose levels (n=2128) | | | | |
|---|---|---|---|---|---|---|
| | | **Effect** | **SE** | ***P*** | **lci** | **uci** |
| **fixed factors** | **overfeeding** | -0.056 | 0.033 | 9.14E-02 | -0.122 | 0.009 |
| | **cholesterol supplementation** | -0.057 | 0.040 | 1.57E-01 | -0.135 | 0.022 |
| | **diethyl ether treatment** | 0.013 | 0.041 | 7.44E-01 | -0.067 | 0.094 |
| | **time of day (in hours since 9AM)** | 0.003 | 0.007 | 6.73E-01 | -0.010 | 0.016 |
| | **intercept** | 0.102 | 0.281 | 7.16E-01 | -0.448 | 0.652 |
| **random factors** | ***variation by transgenic background*** | *0.512* | *0.212* | | *0.227* | *1.154* |
| | ***variation by batch*** | *0.487* | *0.082* | | *0.351* | *0.677* |
| | ***residual*** | *0.744* | *0.011* | | *0.722* | *0.767* |

All outcomes were normalized for protein level using residuals, and inverse-normally transformed before the analysis. Associations were examined using hierarchical linear models and were adjusted for diethyl ether (used to prepare the diet), time of day, transgenic background and batch. Effects shown for overfeeding, cholesterol supplementation and diethyl ether treatment are compared with unexposed controls. Lci and uci are lower and upper boundaries of the 95% confidence interval.

**Supplementary Table 5 - The effect of overfeeding and cholesterol supplementation on image-based vascular atherogenic traits**

| | | Vascular lipid deposition | | | | | | | | | | | | | | |
|---|---|---|---|---|---|---|---|---|---|---|---|---|---|---|---|---|
| | | Model 1 (n=1954) | | | | | Model 2 (n=1954) | | | | | Model 3 (n=1769) | | | | |
| | | **Effect** | **SE** | ***P*** | **lci** | **uci** | **Effect** | **SE** | ***P*** | **lci** | **uci** | **Effect** | **SE** | ***P*** | **lci** | **uci** |
| **negative binomial terms** | **overfeeding** | 0.257 | 0.126 | 4.08E-02 | 0.011 | 0.504 | 0.292 | 0.142 | 4.08E-02 | 0.012 | 0.571 | 0.174 | 0.152 | 2.52E-01 | -0.124 | 0.473 |
| | **cholesterol supplementation** | -0.029 | 0.132 | 8.24E-01 | -0.288 | 0.229 | -0.059 | 0.132 | 6.57E-01 | -0.317 | 0.200 | 0.025 | 0.140 | 8.58E-01 | -0.250 | 0.300 |
| | **diethyl ether treatment** | 0.194 | 0.154 | 2.07E-01 | -0.107 | 0.495 | 0.190 | 0.151 | 2.08E-01 | -0.106 | 0.486 | 0.094 | 0.156 | 5.47E-01 | -0.212 | 0.400 |
| | **time of day (in hours since 9AM)** | 0.054 | 0.020 | 7.74E-03 | 0.014 | 0.094 | 0.056 | 0.021 | 7.55E-03 | 0.015 | 0.098 | 0.070 | 0.023 | 1.97E-03 | 0.026 | 0.115 |
| | **body length (in SD)** | - | - | - | - | - | -0.132 | 0.062 | 3.36E-02 | -0.254 | -0.010 | -0.148 | 0.069 | 3.25E-02 | -0.284 | -0.012 |
| | **dorsal body surface area (in SD)** | - | - | - | - | - | 0.021 | 0.070 | 7.65E-01 | -0.116 | 0.158 | -0.001 | 0.075 | 9.84E-01 | -0.148 | 0.145 |
| | **triglyceride levels (in SD)** | - | - | - | - | - | - | - | - | - | - | 0.222 | 0.084 | 7.96E-03 | 0.058 | 0.386 |
| | **glucose levels (in SD)** | - | - | - | - | - | - | - | - | - | - | -0.047 | 0.068 | 4.91E-01 | -0.181 | 0.087 |
| | **Tg(hsp70:IK17:EGFP) carriers vs. Tg(mpo:EGFP; mpeg1:mCherry) carriers** | -4.041 | 0.357 | 1.07E-29 | -4.740 | -3.341 | -4.101 | 0.363 | 1.31E-29 | -4.813 | -3.390 | -3.738 | 0.399 | 7.06E-21 | -4.520 | -2.956 |
| | **Tg(hsp70:IK17:EGFP; mpeg1:mCherry) carriers vs. Tg(mpo:EGFP; mpeg1:mCherry) carriers** | -3.356 | 0.328 | 1.43E-24 | -3.999 | -2.713 | -3.330 | 0.336 | 4.02E-23 | -3.988 | -2.671 | -3.126 | 0.403 | 8.75E-15 | -3.916 | -2.336 |
| | **Tg(flk:EGFP) carriers vs. Tg(mpo:EGFP; mpeg1:mCherry) carriers** | -2.969 | 0.294 | 5.27E-24 | -3.545 | -2.393 | -3.132 | 0.308 | 3.03E-24 | -3.736 | -2.527 | -3.168 | 0.357 | 6.45E-19 | -3.867 | -2.469 |
| | **batch 1** | -0.531 | 0.192 | 5.69E-03 | -0.907 | -0.154 | -0.514 | 0.201 | 1.07E-02 | -0.909 | -0.119 | -0.438 | 0.217 | 4.40E-02 | -0.864 | -0.012 |
| | **batch 2** | 0.671 | 0.174 | 1.20E-04 | 0.329 | 1.013 | 0.725 | 0.188 | 1.20E-04 | 0.356 | 1.094 | 0.828 | 0.211 | 8.48E-05 | 0.415 | 1.241 |
| | **batch 3** | -0.835 | 0.221 | 1.61E-04 | -1.269 | -0.402 | -0.925 | 0.237 | 9.39E-05 | -1.389 | -0.461 | | | | | |
| | **batch 6** | -0.078 | 0.246 | 7.50E-01 | -0.559 | 0.403 | -0.110 | 0.248 | 6.59E-01 | -0.596 | 0.377 | -0.128 | 0.266 | 6.31E-01 | -0.648 | 0.393 |
| | **batch 7** | -2.103 | 0.447 | 2.55E-06 | -2.979 | -1.227 | -2.258 | 0.413 | 4.68E-08 | -3.068 | -1.448 | -1.854 | 0.449 | 3.69E-05 | -2.735 | -0.973 |
| | **batch 9** | -2.195 | 0.717 | 2.21E-03 | -3.602 | -0.789 | -2.277 | 0.692 | 9.96E-04 | -3.632 | -0.921 | -2.426 | 0.654 | 2.06E-04 | -3.707 | -1.145 |
| | **batch 10** | 0.780 | 0.469 | 9.65E-02 | -0.140 | 1.699 | 0.710 | 0.474 | 1.34E-01 | -0.219 | 1.639 | 0.674 | 0.479 | 1.59E-01 | -0.265 | 1.613 |
| | **batch 11** | -1.900 | 0.924 | 3.97E-02 | -3.710 | -0.089 | -1.847 | 0.934 | 4.80E-02 | -3.679 | -0.016 | -1.935 | 0.951 | 4.18E-02 | -3.799 | -0.072 |
| | **batch 12** | -0.171 | 0.580 | 7.69E-01 | -1.307 | 0.965 | -0.154 | 0.581 | 7.91E-01 | -1.294 | 0.985 | -0.231 | 0.598 | 6.99E-01 | -1.403 | 0.941 |
| | **batch 14** | -0.112 | 0.600 | 8.52E-01 | -1.289 | 1.065 | 0.014 | 0.614 | 9.82E-01 | -1.189 | 1.218 | 0.004 | 0.618 | 9.94E-01 | -1.208 | 1.217 |
| | **batch 15** | 1.295 | 0.368 | 4.28E-04 | 0.574 | 2.015 | 1.358 | 0.371 | 2.52E-04 | 0.631 | 2.085 | 1.224 | 0.383 | 1.39E-03 | 0.474 | 1.975 |
| | **batch 16** | 0.393 | 0.393 | 3.18E-01 | -0.378 | 1.164 | 0.453 | 0.404 | 2.62E-01 | -0.338 | 1.244 | 0.311 | 0.407 | 4.44E-01 | -0.486 | 1.108 |
| | **batch 17** | 1.113 | 0.386 | 3.91E-03 | 0.357 | 1.868 | 1.161 | 0.398 | 3.57E-03 | 0.380 | 1.942 | 1.009 | 0.408 | 1.34E-02 | 0.209 | 1.810 |
| | **batch 18** | 0.390 | 0.399 | 3.29E-01 | -0.393 | 1.172 | 0.446 | 0.417 | 2.84E-01 | -0.371 | 1.263 | 0.262 | 0.432 | 5.44E-01 | -0.585 | 1.109 |
| | **batch 20** | 2.733 | 0.269 | 3.34E-24 | 2.205 | 3.261 | 2.830 | 0.273 | 3.04E-25 | 2.296 | 3.364 | 3.018 | 0.307 | 8.26E-23 | 2.417 | 3.620 |
| | **batch 21** | -0.078 | 0.321 | 8.08E-01 | -0.707 | 0.551 | -0.019 | 0.318 | 9.53E-01 | -0.642 | 0.605 | 0.412 | 0.358 | 2.50E-01 | -0.290 | 1.114 |
| | **batch 22** | 0.398 | 0.283 | 1.59E-01 | -0.156 | 0.952 | 0.542 | 0.292 | 6.39E-02 | -0.031 | 1.115 | 0.918 | 0.342 | 7.17E-03 | 0.249 | 1.588 |
| | **batch 23** | 1.155 | 0.303 | 1.41E-04 | 0.560 | 1.750 | 1.232 | 0.301 | 4.32E-05 | 0.642 | 1.823 | 1.576 | 0.345 | 4.78E-06 | 0.901 | 2.251 |
| | **batch 24** | 1.343 | 0.300 | 7.63E-06 | 0.755 | 1.931 | 1.441 | 0.303 | 2.01E-06 | 0.847 | 2.035 | 1.737 | 0.339 | 3.05E-07 | 1.072 | 2.403 |
| | **batch 25** | 0.211 | 0.342 | 5.37E-01 | -0.459 | 0.881 | 0.220 | 0.340 | 5.18E-01 | -0.446 | 0.885 | 0.521 | 0.387 | 1.79E-01 | -0.239 | 1.280 |
| | **batch 26** | 2.023 | 0.367 | 3.64E-08 | 1.303 | 2.743 | 1.966 | 0.376 | 1.71E-07 | 1.229 | 2.704 | - | - | - | - | - |
| | **intercept** | 4.564 | 0.243 | 1.84E-78 | 4.087 | 5.041 | 4.574 | 0.262 | 4.40E-68 | 4.060 | 5.089 | 4.397 | 0.320 | 6.33E-43 | 3.769 | 5.024 |

*continued* **Supplementary Table 5**

| | | Vascular accumulation of oxLDL | | | | | | | | | | | | | | |
|---|---|---|---|---|---|---|---|---|---|---|---|---|---|---|---|---|
| | | Model 1 (n=885) | | | | | Model 2 (n=885) | | | | | Model 3 (n=876) | | | | |
| | | **Effect** | **SE** | ***P*** | **lci** | **uci** | **Effect** | **SE** | ***P*** | **lci** | **uci** | **Effect** | **SE** | ***P*** | **lci** | **uci** |
| **negative binomial terms** | **overfeeding** | 0.291 | 0.058 | 4.67E-07 | 0.178 | 0.405 | 0.292 | 0.061 | 1.68E-06 | 0.173 | 0.412 | 0.278 | 0.061 | 5.86E-06 | 0.158 | 0.399 |
| | **cholesterol supplementation** | -0.115 | 0.069 | 9.52E-02 | -0.249 | 0.020 | -0.097 | 0.070 | 1.62E-01 | -0.234 | 0.039 | -0.094 | 0.072 | 1.92E-01 | -0.234 | 0.047 |
| | **diethyl ether treatment** | 0.016 | 0.067 | 8.10E-01 | -0.115 | 0.148 | 0.008 | 0.067 | 9.00E-01 | -0.123 | 0.140 | 0.000 | 0.068 | 9.94E-01 | -0.135 | 0.134 |
| | **time of day (in hours since 9AM)** | 0.020 | 0.013 | 1.19E-01 | -0.005 | 0.046 | 0.021 | 0.013 | 1.06E-01 | -0.004 | 0.046 | 0.022 | 0.013 | 9.46E-02 | -0.004 | 0.049 |
| | **body length (in SD)** | - | - | - | - | - | 0.061 | 0.037 | 1.01E-01 | -0.012 | 0.134 | 0.049 | 0.039 | 2.10E-01 | -0.028 | 0.126 |
| | **dorsal body surface area (in SD)** | - | - | - | - | - | -0.021 | 0.039 | 5.90E-01 | -0.099 | 0.056 | -0.029 | 0.041 | 4.78E-01 | -0.110 | 0.052 |
| | **triglyceride levels (in SD)** | - | - | - | - | - | - | - | - | - | - | 0.035 | 0.041 | 3.86E-01 | -0.045 | 0.115 |
| | **glucose levels (in SD)** | - | - | - | - | - | - | - | - | - | - | -0.042 | 0.042 | 3.21E-01 | -0.124 | 0.040 |
| | **Tg(hsp70:IK17:EGFP; mpeg1:mCherry) carriers vs. Tg(hsp70:IK17:EGFP) carriers** | 1.762 | 0.113 | 4.18E-55 | 1.541 | 1.982 | 1.722 | 0.117 | 2.85E-49 | 1.493 | 1.950 | 1.693 | 0.124 | 2.06E-42 | 1.450 | 1.937 |
| | **batch 10** | 1.614 | 0.097 | 1.23E-62 | 1.425 | 1.804 | 1.650 | 0.098 | 4.11E-63 | 1.457 | 1.842 | 1.688 | 0.112 | 6.37E-51 | 1.467 | 1.908 |
| | **batch 11** | 1.475 | 0.130 | 1.20E-29 | 1.219 | 1.731 | 1.444 | 0.133 | 1.60E-27 | 1.183 | 1.704 | 1.490 | 0.140 | 1.40E-26 | 1.216 | 1.764 |
| | **batch 12** | 1.716 | 0.112 | 6.59E-53 | 1.497 | 1.936 | 1.688 | 0.113 | 3.62E-50 | 1.466 | 1.910 | 1.686 | 0.117 | 8.35E-47 | 1.456 | 1.916 |
| | **batch 13** | 0.753 | 0.127 | 2.72E-09 | 0.505 | 1.002 | 0.746 | 0.129 | 7.24E-09 | 0.494 | 0.999 | 0.753 | 0.129 | 4.94E-09 | 0.501 | 1.005 |
| | **batch 14** | -1.255 | 0.156 | 7.54E-16 | -1.560 | -0.950 | -1.303 | 0.159 | 2.67E-16 | -1.614 | -0.991 | -1.285 | 0.161 | 1.21E-15 | -1.599 | -0.970 |
| | **batch 15** | 0.012 | 0.139 | 9.31E-01 | -0.260 | 0.284 | -0.020 | 0.140 | 8.84E-01 | -0.294 | 0.253 | -0.018 | 0.141 | 8.99E-01 | -0.294 | 0.259 |
| | **batch 16** | -0.689 | 0.135 | 3.64E-07 | -0.955 | -0.424 | -0.706 | 0.138 | 3.17E-07 | -0.977 | -0.435 | -0.729 | 0.143 | 3.32E-07 | -1.009 | -0.449 |
| | **batch 17** | -0.708 | 0.116 | 1.03E-09 | -0.935 | -0.481 | -0.737 | 0.122 | 1.51E-09 | -0.975 | -0.498 | -0.761 | 0.127 | 2.07E-09 | -1.010 | -0.512 |
| | **batch 18** | -0.160 | 0.124 | 1.97E-01 | -0.403 | 0.083 | -0.167 | 0.130 | 2.01E-01 | -0.422 | 0.089 | -0.173 | 0.141 | 2.20E-01 | -0.451 | 0.104 |
| | **intercept** | 4.656 | 0.127 | 7.20E-296 | 4.408 | 4.904 | 4.666 | 0.127 | 1.78E-294 | 4.416 | 4.915 | 4.675 | 0.130 | 9.42E-285 | 4.420 | 4.929 |

| | | Vascular infiltration by macrophages | | | | | | | | | | | | | | |
|---|---|---|---|---|---|---|---|---|---|---|---|---|---|---|---|---|
| | | Model 1 (n=994) | | | | | Model 2 (n=994) | | | | | Model 3 (n=880) | | | | |
| | | **Effect** | **SE** | ***P*** | **lci** | **uci** | **Effect** | **SE** | ***P*** | **lci** | **uci** | **Effect** | **SE** | ***P*** | **lci** | **uci** |
| **negative binomial terms** | **overfeeding** | -0.013 | 0.043 | 7.68E-01 | -0.096 | 0.071 | -0.058 | 0.047 | 2.20E-01 | -0.151 | 0.035 | -0.084 | 0.051 | 1.01E-01 | -0.185 | 0.016 |
| | **cholesterol supplementation** | 0.066 | 0.052 | 2.03E-01 | -0.036 | 0.168 | 0.070 | 0.052 | 1.79E-01 | -0.032 | 0.172 | 0.067 | 0.055 | 2.18E-01 | -0.040 | 0.174 |
| | **diethyl ether treatment** | -0.001 | 0.051 | 9.86E-01 | -0.101 | 0.099 | 0.025 | 0.052 | 6.30E-01 | -0.077 | 0.126 | 0.012 | 0.053 | 8.25E-01 | -0.093 | 0.117 |
| | **time of day (in hours since 9AM)** | 0.008 | 0.008 | 3.39E-01 | -0.008 | 0.023 | 0.002 | 0.008 | 7.70E-01 | -0.014 | 0.019 | 0.015 | 0.009 | 1.05E-01 | -0.003 | 0.033 |
| | **body length (in SD)** | - | - | - | - | - | 0.049 | 0.025 | 4.91E-02 | 0.000 | 0.099 | 0.030 | 0.028 | 2.84E-01 | -0.025 | 0.084 |
| | **dorsal body surface area (in SD)** | - | - | - | - | - | 0.048 | 0.026 | 6.45E-02 | -0.003 | 0.098 | 0.047 | 0.028 | 9.53E-02 | -0.008 | 0.102 |
| | **triglyceride levels (in SD)** | - | - | - | - | - | - | - | - | - | - | 0.063 | 0.039 | 1.07E-01 | -0.014 | 0.141 |
| | **glucose levels (in SD)** | - | - | - | - | - | - | - | - | - | - | -0.115 | 0.029 | 6.12E-05 | -0.171 | -0.059 |
| | **Tg(hsp70:IK17:EGFP; mpeg1:mCherry) carriers vs. Tg(mpo:EGFP; mpeg1:mCherry) carriers** | -0.969 | 0.087 | 1.09E-28 | -1.140 | -0.798 | -1.016 | 0.088 | 7.75E-31 | -1.188 | -0.844 | -1.054 | 0.116 | 8.06E-20 | -1.281 | -0.828 |
| | **batch 1** | -0.065 | 0.076 | 3.91E-01 | -0.213 | 0.083 | -0.093 | 0.076 | 2.21E-01 | -0.241 | 0.056 | -0.036 | 0.076 | 6.37E-01 | -0.185 | 0.113 |
| | **batch 2** | -0.035 | 0.080 | 6.61E-01 | -0.192 | 0.122 | -0.087 | 0.081 | 2.88E-01 | -0.246 | 0.073 | 0.007 | 0.086 | 9.34E-01 | -0.162 | 0.177 |
| | **batch 3** | -1.359 | 0.102 | 7.57E-41 | -1.558 | -1.160 | -1.379 | 0.107 | 7.15E-38 | -1.589 | -1.169 | | | | | |
| | **batch 6** | 0.038 | 0.092 | 6.83E-01 | -0.143 | 0.218 | 0.028 | 0.093 | 7.65E-01 | -0.154 | 0.210 | 0.224 | 0.103 | 3.01E-02 | 0.022 | 0.427 |
| | **batch 7** | -1.959 | 0.108 | 5.69E-74 | -2.170 | -1.748 | -1.949 | 0.109 | 3.42E-71 | -2.164 | -1.735 | -1.818 | 0.139 | 7.74E-39 | -2.091 | -1.544 |
| | **batch 14** | -0.539 | 0.122 | 9.91E-06 | -0.778 | -0.300 | -0.488 | 0.125 | 9.08E-05 | -0.733 | -0.244 | -0.415 | 0.127 | 1.04E-03 | -0.663 | -0.167 |
| | **batch 15** | -0.233 | 0.091 | 1.03E-02 | -0.411 | -0.055 | -0.240 | 0.090 | 7.44E-03 | -0.416 | -0.064 | -0.271 | 0.090 | 2.58E-03 | -0.448 | -0.095 |
| | **batch 16** | -1.115 | 0.116 | 6.04E-22 | -1.342 | -0.888 | -1.059 | 0.119 | 7.06E-19 | -1.293 | -0.825 | -1.062 | 0.123 | 0.000 | -1.302 | -0.822 |

***continued*** **Supplementary Table 5**

| | | Vascular infiltration by macrophages | | | | | | | | | | | | | | |
|---|---|---|---|---|---|---|---|---|---|---|---|---|---|---|---|---|
| | | Model 1 (n=994) | | | | | Model 2 (n=994) | | | | | Model 3 (n=880) | | | | |
| | | Effect | SE | *P* | lci | uci | Effect | SE | *P* | lci | uci | Effect | SE | *P* | lci | uci |
| | **batch 17** | -0.973 | 0.103 | 4.04E-21 | -1.175 | -0.771 | -0.916 | 0.107 | 1.03E-17 | -1.125 | -0.706 | -0.945 | 0.111 | 1.35E-17 | -1.162 | -0.728 |
| | **batch 18** | -0.221 | 0.082 | 6.77E-03 | -0.381 | -0.061 | -0.154 | 0.087 | 7.83E-02 | -0.325 | 0.017 | -0.119 | 0.095 | 2.10E-01 | -0.305 | 0.067 |
| | **intercept** | 7.880 | 0.095 | 0.00E+00 | 7.694 | 8.066 | 7.921 | 0.099 | 0.00E+00 | 7.727 | 8.114 | 7.836 | 0.122 | 0.00E+00 | 7.597 | 8.074 |

| | | Vascular co-localization of lipids with macrophages | | | | | | | | | | | | | | |
|---|---|---|---|---|---|---|---|---|---|---|---|---|---|---|---|---|
| | | Model 1 (n=870) | | | | | Model 2 (n=870) | | | | | Model 3 (n=763) | | | | |
| | | Effect | SE | *P* | lci | uci | Effect | SE | *P* | lci | uci | Effect | SE | *P* | lci | uci |
| **negative binomial terms** | **overfeeding** | 0.273 | 0.225 | 2.26E-01 | -0.169 | 0.714 | 0.195 | 0.248 | 4.31E-01 | -0.291 | 0.680 | 0.251 | 0.285 | 3.78E-01 | -0.307 | 0.810 |
| | **cholesterol supplementation** | 0.383 | 0.315 | 2.24E-01 | -0.235 | 1.001 | 0.356 | 0.312 | 2.54E-01 | -0.256 | 0.968 | 0.554 | 0.354 | 1.18E-01 | -0.140 | 1.248 |
| | **diethyl ether treatment** | 0.291 | 0.291 | 3.16E-01 | -0.279 | 0.861 | 0.207 | 0.283 | 4.65E-01 | -0.348 | 0.762 | 0.281 | 0.297 | 3.43E-01 | -0.301 | 0.863 |
| | **time of day (in hours since 9AM)** | 0.005 | 0.045 | 9.17E-01 | -0.084 | 0.093 | -0.002 | 0.046 | 9.63E-01 | -0.092 | 0.087 | 0.040 | 0.050 | 4.24E-01 | -0.058 | 0.139 |
| | **body length (in SD)** | - | - | - | - | - | -0.240 | 0.131 | 6.64E-02 | -0.496 | 0.016 | -0.185 | 0.138 | 1.79E-01 | -0.454 | 0.085 |
| | **dorsal body surface area (in SD)** | - | - | - | - | - | 0.266 | 0.154 | 8.42E-02 | -0.036 | 0.567 | 0.220 | 0.165 | 1.82E-01 | -0.103 | 0.543 |
| | **triglyceride levels (in SD)** | - | - | - | - | - | - | - | - | - | - | 0.404 | 0.186 | 3.04E-02 | 0.038 | 0.769 |
| | **glucose levels (in SD)** | - | - | - | - | - | - | - | - | - | - | -0.112 | 0.151 | 4.60E-01 | -0.408 | 0.184 |
| | **Tg(hsp70:IK17:EGFP; mpeg1:mCherry) carriers vs. Tg(mpo:EGFP; mpeg1:mCherry) carriers** | -3.913 | 0.563 | 3.64E-12 | -5.017 | -2.810 | -4.177 | 0.517 | 6.76E-16 | -5.191 | -3.163 | -3.914 | 0.688 | 1.26E-08 | -5.262 | -2.566 |
| | **batch 1** | -0.741 | 0.313 | 1.80E-02 | -1.355 | -0.127 | -0.868 | 0.337 | 9.93E-03 | -1.527 | -0.208 | -0.767 | 0.356 | 3.12E-02 | -1.465 | -0.069 |
| | **batch 2** | 0.644 | 0.302 | 3.30E-02 | 0.052 | 1.236 | 0.543 | 0.322 | 9.21E-02 | -0.089 | 1.174 | 0.783 | 0.376 | 3.72E-02 | 0.046 | 1.520 |
| | **batch 3** | -1.767 | 0.447 | 7.63E-05 | -2.642 | -0.891 | -2.255 | 0.461 | 9.94E-07 | -3.158 | -1.352 | - | - | - | - | - |
| | **batch 6** | 0.078 | 0.373 | 8.35E-01 | -0.654 | 0.809 | -0.159 | 0.373 | 6.70E-01 | -0.889 | 0.572 | -0.109 | 0.412 | 7.91E-01 | -0.916 | 0.698 |
| | **batch 7** | -3.258 | 0.617 | 1.28E-07 | -4.467 | -2.049 | -3.685 | 0.634 | 6.18E-09 | -4.928 | -2.442 | -2.598 | 0.836 | 1.88E-03 | -4.237 | -0.960 |
| | **batch 15** | 1.239 | 0.602 | 3.95E-02 | 0.060 | 2.419 | 1.372 | 0.557 | 1.38E-02 | 0.280 | 2.464 | 1.256 | 0.544 | 2.10E-02 | 0.189 | 2.323 |
| | **batch 16** | -1.544 | 1.034 | 1.35E-01 | -3.572 | 0.483 | -1.250 | 1.019 | 2.20E-01 | -3.247 | 0.747 | -1.460 | 0.973 | 1.33E-01 | -3.366 | 0.446 |
| | **batch 17** | 0.324 | 0.660 | 6.23E-01 | -0.969 | 1.617 | 0.760 | 0.633 | 2.30E-01 | -0.481 | 2.002 | 0.474 | 0.665 | 4.75E-01 | -0.828 | 1.777 |
| | **batch 18** | 0.546 | 0.972 | 5.74E-01 | -1.358 | 2.451 | 1.085 | 0.981 | 2.69E-01 | -0.838 | 3.009 | 0.882 | 0.967 | 3.62E-01 | -1.014 | 2.778 |
| | **intercept** | 3.297 | 0.490 | 1.65E-11 | 2.338 | 4.257 | 3.610 | 0.510 | 1.45E-12 | 2.610 | 4.609 | 2.839 | 0.650 | 1.27E-05 | 1.565 | 4.114 |

| | | Vascular co-localization of oxLDL with macrophages | | | | | | | | | | | | | | |
|---|---|---|---|---|---|---|---|---|---|---|---|---|---|---|---|---|
| | | Model 1 (n=433) | | | | | Model 2 (n=433) | | | | | Model 3 (n=430) | | | | |
| | | Effect | SE | *P* | lci | uci | Effect | SE | *P* | lci | uci | Effect | SE | *P* | lci | uci |
| **negative binomial terms** | **overfeeding** | 0.447 | 0.138 | 1.22E-03 | 0.176 | 0.717 | 0.456 | 0.145 | 1.67E-03 | 0.172 | 0.741 | 0.407 | 0.146 | 5.33E-03 | 0.121 | 0.694 |
| | **cholesterol supplementation** | -0.372 | 0.154 | 1.55E-02 | -0.674 | -0.071 | -0.389 | 0.156 | 1.27E-02 | -0.695 | -0.083 | -0.357 | 0.158 | 2.39E-02 | -0.667 | -0.047 |
| | **diethyl ether treatment** | -0.109 | 0.146 | 4.53E-01 | -0.394 | 0.176 | -0.113 | 0.147 | 4.41E-01 | -0.401 | 0.175 | -0.156 | 0.148 | 2.91E-01 | -0.446 | 0.134 |
| | **time of day (in hours since 9AM)** | 0.061 | 0.028 | 2.73E-02 | 0.007 | 0.115 | 0.064 | 0.029 | 2.59E-02 | 0.008 | 0.120 | 0.075 | 0.030 | 1.21E-02 | 0.016 | 0.133 |
| | **body length (in SD)** | - | - | - | - | - | -0.064 | 0.090 | 4.79E-01 | -0.241 | 0.113 | -0.182 | 0.107 | 8.77E-02 | -0.392 | 0.027 |
| | **dorsal body surface area (in SD)** | - | - | - | - | - | -0.027 | 0.081 | 7.42E-01 | -0.186 | 0.133 | -0.144 | 0.097 | 1.37E-01 | -0.334 | 0.046 |
| | **triglyceride levels (in SD)** | - | - | - | - | - | - | - | - | - | - | 0.348 | 0.161 | 3.06E-02 | 0.032 | 0.663 |
| | **glucose levels (in SD)** | - | - | - | - | - | - | - | - | - | - | -0.288 | 0.141 | 4.18E-02 | -0.565 | -0.011 |
| | **batch 15** | 0.693 | 0.254 | 6.40E-03 | 0.195 | 1.192 | 0.727 | 0.273 | 7.76E-03 | 0.192 | 1.262 | 0.491 | 0.302 | 1.04E-01 | -0.101 | 1.084 |
| | **batch 16** | 0.549 | 0.290 | 5.82E-02 | -0.019 | 1.118 | 0.527 | 0.288 | 6.68E-02 | -0.036 | 1.091 | 0.217 | 0.326 | 5.04E-01 | -0.421 | 0.856 |
| | **batch 17** | -0.225 | 0.262 | 3.90E-01 | -0.738 | 0.288 | -0.234 | 0.263 | 3.73E-01 | -0.750 | 0.281 | -0.652 | 0.310 | 3.55E-02 | -1.260 | -0.044 |
| | **batch 18** | 0.715 | 0.248 | 3.97E-03 | 0.228 | 1.201 | 0.690 | 0.251 | 5.96E-03 | 0.198 | 1.181 | 0.406 | 0.289 | 1.60E-01 | -0.160 | 0.972 |

***continued*** **Supplementary Table 5**

| | | Vascular co-localization of oxLDL with macrophages | | | | | | | | | | | | | | |
|---|---|---|---|---|---|---|---|---|---|---|---|---|---|---|---|---|
| | | Model 1 (n=433) | | | | | Model 2 (n=433) | | | | | Model 3 (n=430) | | | | |
| | | Effect | SE | *P* | lci | uci | Effect | SE | *P* | lci | uci | Effect | SE | *P* | lci | uci |
| | batch 19 | 0.869 | 0.236 | 2.26E-04 | 0.407 | 1.331 | 0.892 | 0.259 | 5.72E-04 | 0.385 | 1.400 | 0.805 | 0.273 | 3.16E-03 | 0.270 | 1.340 |
| | intercept | 2.076 | 0.278 | 7.97E-14 | 1.531 | 2.620 | 2.082 | 0.303 | 6.85E-12 | 1.487 | 2.677 | 2.052 | 0.313 | 5.92E-11 | 1.438 | 2.666 |

| | | Vascular infiltration by neutrophils | | | | | | | | | | | | | | |
|---|---|---|---|---|---|---|---|---|---|---|---|---|---|---|---|---|
| | | Model 1 (n=494) | | | | | Model 2 (n=494) | | | | | Model 3 (n=395) | | | | |
| | | Effect | SE | *P* | lci | uci | Effect | SE | *P* | lci | uci | Effect | SE | *P* | lci | uci |
| negative binomial terms | overfeeding | 0.199 | 0.054 | 2.59E-04 | 0.092 | 0.306 | 0.131 | 0.063 | 3.80E-02 | 0.007 | 0.255 | 0.196 | 0.069 | 4.64E-03 | 0.060 | 0.332 |
| | cholesterol supplementation | 0.046 | 0.069 | 5.07E-01 | -0.090 | 0.181 | 0.045 | 0.069 | 5.13E-01 | -0.091 | 0.181 | 0.183 | 0.074 | 1.39E-02 | 0.037 | 0.329 |
| | diethyl ether treatment | -0.033 | 0.066 | 6.15E-01 | -0.164 | 0.097 | 0.001 | 0.066 | 9.87E-01 | -0.128 | 0.130 | 0.018 | 0.070 | 8.00E-01 | -0.120 | 0.156 |
| | time of day (in hours since 9AM) | 0.033 | 0.011 | 2.07E-03 | 0.012 | 0.053 | 0.023 | 0.011 | 4.32E-02 | 0.001 | 0.045 | 0.026 | 0.013 | 4.46E-02 | 0.001 | 0.051 |
| | body length (in SD) | - | - | - | - | - | 0.054 | 0.030 | 7.75E-02 | -0.006 | 0.113 | 0.059 | 0.035 | 9.34E-02 | -0.010 | 0.129 |
| | dorsal body surface area (in SD) | - | - | - | - | - | 0.071 | 0.038 | 6.56E-02 | -0.005 | 0.146 | 0.032 | 0.042 | 4.49E-01 | -0.051 | 0.115 |
| | triglyceride levels (in SD) | - | - | - | - | - | - | - | - | - | - | -0.077 | 0.056 | 1.70E-01 | -0.187 | 0.033 |
| | glucose levels (in SD) | - | - | - | - | - | - | - | - | - | - | -0.053 | 0.036 | 1.41E-01 | -0.124 | 0.018 |
| | batch 1 | -0.209 | 0.086 | 1.56E-02 | -0.378 | -0.040 | -0.263 | 0.088 | 2.74E-03 | -0.435 | -0.091 | -0.260 | 0.088 | 3.21E-03 | -0.434 | -0.087 |
| | batch 2 | -0.245 | 0.086 | 4.45E-03 | -0.414 | -0.076 | -0.330 | 0.087 | 1.46E-04 | -0.500 | -0.159 | -0.343 | 0.092 | 1.98E-04 | -0.524 | -0.163 |
| | batch 3 | -1.742 | 0.109 | 2.22E-57 | -1.956 | -1.528 | -1.816 | 0.116 | 1.85E-55 | -2.043 | -1.589 | - | - | - | - | - |
| | batch 6 | -0.071 | 0.118 | 5.47E-01 | -0.301 | 0.160 | -0.114 | 0.116 | 3.25E-01 | -0.342 | 0.113 | -0.072 | 0.137 | 5.97E-01 | -0.341 | 0.196 |
| | batch 7 | -2.087 | 0.105 | 2.93E-88 | -2.293 | -1.882 | -2.121 | 0.107 | 5.84E-87 | -2.331 | -1.910 | -2.310 | 0.167 | 1.11E-43 | -2.636 | -1.983 |
| | intercept | 7.658 | 0.105 | 0.00E+00 | 7.451 | 7.864 | 7.760 | 0.115 | 0.00E+00 | 7.534 | 7.985 | 7.776 | 0.152 | 0.00E+00 | 7.479 | 8.074 |

| | | Vascular co-localization of lipids with neutrophils | | | | | | | | | | | | | | |
|---|---|---|---|---|---|---|---|---|---|---|---|---|---|---|---|---|
| | | Model 1 (n=380) | | | | | Model 2 (n=380) | | | | | Model 3 (n=286) | | | | |
| | | Effect | SE | *P* | lci | uci | Effect | SE | *P* | lci | uci | Effect | SE | *P* | lci | uci |
| negative binomial terms | overfeeding | 0.727 | 0.246 | 3.07E-03 | 0.246 | 1.209 | 0.826 | 0.251 | 1.02E-03 | 0.333 | 1.319 | 0.901 | 0.255 | 4.01E-04 | 0.402 | 1.401 |
| | cholesterol supplementation | 0.845 | 0.284 | 2.89E-03 | 0.289 | 1.401 | 0.830 | 0.280 | 2.98E-03 | 0.282 | 1.378 | 0.882 | 0.306 | 3.95E-03 | 0.282 | 1.482 |
| | diethyl ether treatment | -0.659 | 0.319 | 3.91E-02 | -1.285 | -0.033 | -0.744 | 0.325 | 2.21E-02 | -1.381 | -0.107 | -0.258 | 0.286 | 3.66E-01 | -0.819 | 0.302 |
| | time of day (in hours since 9AM) | 0.017 | 0.050 | 7.40E-01 | -0.082 | 0.115 | 0.039 | 0.054 | 4.64E-01 | -0.066 | 0.144 | 0.049 | 0.053 | 3.58E-01 | -0.055 | 0.152 |
| | body length (in SD) | - | - | - | - | - | -0.151 | 0.140 | 2.82E-01 | -0.425 | 0.124 | -0.059 | 0.129 | 6.46E-01 | -0.312 | 0.194 |
| | dorsal body surface area (in SD) | - | - | - | - | - | -0.088 | 0.162 | 5.88E-01 | -0.405 | 0.230 | -0.082 | 0.175 | 6.38E-01 | -0.426 | 0.261 |
| | triglyceride levels (in SD) | - | - | - | - | - | - | - | - | - | - | 0.102 | 0.208 | 6.23E-01 | -0.305 | 0.509 |
| | glucose levels (in SD) | - | - | - | - | - | - | - | - | - | - | 0.132 | 0.127 | 2.98E-01 | -0.117 | 0.381 |
| | batch 1 | -0.804 | 0.340 | 1.79E-02 | -1.469 | -0.138 | -0.701 | 0.378 | 6.37E-02 | -1.441 | 0.040 | -0.804 | 0.353 | 2.27E-02 | -1.497 | -0.112 |
| | batch 2 | 0.117 | 0.356 | 7.41E-01 | -0.580 | 0.815 | 0.289 | 0.402 | 4.73E-01 | -0.500 | 1.077 | 0.391 | 0.414 | 3.45E-01 | -0.421 | 1.203 |
| | batch 3 | -2.749 | 0.523 | 1.44E-07 | -3.774 | -1.725 | -2.714 | 0.572 | 2.05E-06 | -3.834 | -1.594 | - | - | - | - | - |
| | batch 6 | -0.652 | 0.441 | 1.40E-01 | -1.517 | 0.213 | -0.557 | 0.462 | 2.28E-01 | -1.462 | 0.349 | -0.510 | 0.464 | 2.71E-01 | -1.420 | 0.399 |
| | intercept | 2.288 | 0.576 | 7.12E-05 | 1.159 | 3.416 | 2.105 | 0.602 | 4.69E-04 | 0.925 | 3.284 | 1.409 | 0.679 | 3.79E-02 | 0.079 | 2.740 |

***continued*** **Supplementary Table 5**

| | | **Vascular co-localization of macrophages with neutrophils** | | | | | | | | | | | | | | |
|---|---|---|---|---|---|---|---|---|---|---|---|---|---|---|---|---|
| | | **Model 1 (n=488)** | | | | | **Model 2 (n=488)** | | | | | **Model 3 (n=392)** | | | | |
| | | **Effect** | **SE** | ***P*** | **lci** | **uci** | **Effect** | **SE** | ***P*** | **lci** | **uci** | **Effect** | **SE** | ***P*** | **lci** | **uci** |
| **negative binomial terms** | **overfeeding** | 0.096 | 0.104 | 3.55E-01 | -0.107 | 0.299 | -0.019 | 0.115 | 8.67E-01 | -0.244 | 0.205 | 0.060 | 0.113 | 5.99E-01 | -0.162 | 0.281 |
| | **cholesterol supplementation** | 0.153 | 0.133 | 2.50E-01 | -0.108 | 0.413 | 0.119 | 0.136 | 3.81E-01 | -0.147 | 0.385 | 0.299 | 0.127 | 1.90E-02 | 0.049 | 0.549 |
| | **diethyl ether treatment** | -0.240 | 0.136 | 7.75E-02 | -0.506 | 0.026 | -0.164 | 0.136 | 2.27E-01 | -0.430 | 0.102 | -0.164 | 0.133 | 2.18E-01 | -0.426 | 0.097 |
| | **time of day (in hours since 9AM)** | -0.007 | 0.021 | 7.39E-01 | -0.048 | 0.034 | -0.027 | 0.023 | 2.42E-01 | -0.072 | 0.018 | -0.009 | 0.026 | 7.23E-01 | -0.059 | 0.041 |
| | **body length (in SD)** | - | - | - | - | - | 0.034 | 0.054 | 5.27E-01 | -0.071 | 0.139 | -0.022 | 0.054 | 6.84E-01 | -0.127 | 0.083 |
| | **dorsal body surface area (in SD)** | - | - | - | - | - | 0.170 | 0.067 | 1.08E-02 | 0.039 | 0.302 | 0.097 | 0.068 | 1.55E-01 | -0.037 | 0.231 |
| | **triglyceride levels (in SD)** | - | - | - | - | - | - | - | - | - | - | 0.231 | 0.089 | 9.49E-03 | 0.056 | 0.405 |
| | **glucose levels (in SD)** | - | - | - | - | - | - | - | - | - | - | -0.153 | 0.066 | 1.96E-02 | -0.282 | -0.025 |
| | **batch 1** | 0.223 | 0.131 | 8.87E-02 | -0.034 | 0.479 | 0.090 | 0.146 | 5.34E-01 | -0.195 | 0.376 | 0.225 | 0.145 | 1.21E-01 | -0.059 | 0.509 |
| | **batch 2** | -0.355 | 0.150 | 1.79E-02 | -0.650 | -0.061 | -0.523 | 0.166 | 1.59E-03 | -0.848 | -0.198 | -0.336 | 0.175 | 5.57E-02 | -0.679 | 0.008 |
| | **batch 3** | -2.971 | 0.240 | 4.01E-35 | -3.441 | -2.500 | -3.183 | 0.279 | 3.32E-30 | -3.729 | -2.637 | - | - | - | - | - |
| | **batch 6** | 0.175 | 0.191 | 3.60E-01 | -0.199 | 0.549 | 0.059 | 0.199 | 7.67E-01 | -0.331 | 0.448 | 0.426 | 0.232 | 6.61E-02 | -0.028 | 0.880 |
| | **batch 7** | -4.803 | 0.258 | 1.43E-77 | -5.308 | -4.298 | -4.918 | 0.274 | 3.21E-72 | -5.454 | -4.382 | -4.433 | 0.338 | 2.72E-39 | -5.096 | -3.771 |
| | **intercept** | 5.839 | 0.218 | 1.63E-158 | 5.412 | 6.266 | 6.054 | 0.250 | 5.85E-130 | 5.565 | 6.543 | 5.617 | 0.302 | 2.18E-77 | 5.026 | 6.208 |

| | | **Endothelial thickness** | | | | | | | | | | | | | | |
|---|---|---|---|---|---|---|---|---|---|---|---|---|---|---|---|---|
| | | **Model 1 (n=467)** | | | | | **Model 2 (n=467)** | | | | | **Model 3 (n=411)** | | | | |
| | | **Effect** | **SE** | ***P*** | **lci** | **uci** | **Effect** | **SE** | ***P*** | **lci** | **uci** | **Effect** | **SE** | ***P*** | **lci** | **uci** |
| **fixed factors** | **overfeeding** | 0.252 | 0.073 | 5.34E-04 | 0.109 | 0.395 | 0.176 | 0.079 | 2.69E-02 | 0.020 | 0.331 | 0.171 | 0.089 | 5.47E-02 | -0.003 | 0.346 |
| | **cholesterol supplementation** | 0.091 | 0.082 | 2.65E-01 | -0.069 | 0.252 | 0.115 | 0.082 | 1.60E-01 | -0.045 | 0.276 | 0.119 | 0.090 | 1.89E-01 | -0.058 | 0.296 |
| | **diethyl ether treatment** | -0.320 | 0.087 | 2.40E-04 | -0.490 | -0.149 | -0.330 | 0.087 | 1.42E-04 | -0.500 | -0.160 | -0.337 | 0.096 | 4.13E-04 | -0.525 | -0.150 |
| | **time of day (in hours since 9AM)** | 0.065 | 0.014 | 1.43E-06 | 0.039 | 0.092 | 0.066 | 0.013 | 8.20E-07 | 0.040 | 0.093 | 0.076 | 0.015 | 2.11E-07 | 0.048 | 0.105 |
| | **body length (in SD)** | - | - | - | - | - | 0.063 | 0.043 | 1.39E-01 | -0.020 | 0.147 | 0.033 | 0.049 | 5.06E-01 | -0.064 | 0.129 |
| | **dorsal body surface area (in SD)** | - | - | - | - | - | 0.073 | 0.040 | 6.63E-02 | -0.005 | 0.150 | 0.076 | 0.043 | 7.67E-02 | -0.008 | 0.161 |
| | **triglyceride levels (in SD)** | - | - | - | - | - | - | - | - | - | - | 0.023 | 0.041 | 5.73E-01 | -0.057 | 0.104 |
| | **glucose levels (in SD)** | - | - | - | - | - | - | - | - | - | - | 0.045 | 0.043 | 3.00E-01 | -0.040 | 0.129 |
| | **intercept** | -0.445 | 0.259 | 8.56E-02 | -0.953 | 0.062 | -0.378 | 0.262 | 1.49E-01 | -0.890 | 0.135 | -0.366 | 0.288 | 2.04E-01 | -0.930 | 0.198 |
| **random factors** | ***variation by batch*** | *0.672* | *0.173* | - | *0.405* | *1.113* | *0.676* | *0.174* | - | *0.408* | *1.120* | *0.693* | *0.192* | - | *0.402* | *1.192* |
| | ***residual*** | *0.703* | *0.023* | - | *0.659* | *0.750* | *0.698* | *0.023* | - | *0.655* | *0.745* | *0.716* | *0.025* | - | *0.669* | *0.767* |

Endothelial thickness is defined as surface area of the endothelium normalized for surface area of the circulating lipids. Associations were examined using negative binomial regresion for outcomes that showed a negative binomial distribution; and using hierarchical linear models on inverse normally transformed outcomes for outcomes that were (borderline) normally distributed (i.e. endothelial thickness). Model 1: adjusted for diethyl ether (used to prepare the diet), time of day, transgenic background and batch; Model 2: additionally adjusted for body length and dorsal body surface area; Model 3: additionally adjusted for whole-body triglyceride and glucose levels. Dorsal body surface area was normalized for body length using residuals; whole-body triglyceride and glucose levels were normalized for protein level using residuals. Effects shown for overfeeding, cholesterol supplementation and diethyl ether treatment are compared with unexposed controls. Lci and uci are lower and upper boundaries of the 95% confidence interval.

**Supplementary Table 6 - The effect of overfeeding and cholesterol supplementation on suboptimal image or quantification quality**

| | Larva optically cut during pre-processing | | | | |
|---|---|---|---|---|---|
| | Model 1 (n=2306) | | | | |
| | OR | SE | *P* | lci | uci |
| **overfeeding** | 1.450 | 0.290 | 6.52E-02 | 0.980 | 2.150 |
| **cholesterol supplementation** | 1.220 | 0.280 | 3.97E-01 | 0.770 | 1.920 |
| **diethyl ether treatment** | 1.090 | 0.270 | 7.23E-01 | 0.670 | 1.790 |
| **time of day (in hours since 9AM)** | 0.950 | 0.040 | 1.96E-01 | 0.890 | 1.030 |
| **intercept** | 0.050 | 0.010 | 6.21E-26 | 0.030 | 0.080 |

| | Vasculature not properly detected | | | | | | | | | | | | | | |
|---|---|---|---|---|---|---|---|---|---|---|---|---|---|---|---|
| | Model 1 (n=2050) | | | | | Model 2 (n=2050) | | | | | Model 3 (n=1859) | | | | |
| | OR | SE | *P* | lci | uci | OR | SE | *P* | lci | uci | OR | SE | *P* | lci | uci |
| **overfeeding** | 0.650 | 0.140 | 4.16E-02 | 0.420 | 0.980 | 0.860 | 0.190 | 4.89E-01 | 0.550 | 1.330 | 0.790 | 0.190 | 3.25E-01 | 0.500 | 1.260 |
| **cholesterol supplementation** | 0.920 | 0.250 | 7.52E-01 | 0.540 | 1.560 | 0.840 | 0.230 | 5.13E-01 | 0.490 | 1.430 | 0.670 | 0.190 | 1.62E-01 | 0.380 | 1.170 |
| **diethyl ether treatment** | 0.810 | 0.210 | 4.13E-01 | 0.490 | 1.340 | 0.820 | 0.210 | 4.36E-01 | 0.490 | 1.360 | 0.910 | 0.240 | 7.29E-01 | 0.540 | 1.540 |
| **time of day (in hours since 9AM)** | 0.910 | 0.040 | 2.15E-02 | 0.840 | 0.990 | 0.920 | 0.040 | 6.32E-02 | 0.850 | 1.000 | 0.900 | 0.040 | 1.87E-02 | 0.820 | 0.980 |
| **body length (in SD)** | - | - | - | - | - | 0.770 | 0.090 | 2.26E-02 | 0.610 | 0.960 | 0.760 | 0.100 | 3.75E-02 | 0.580 | 0.980 |
| **dorsal body surface area (in SD)** | - | - | - | - | - | 0.650 | 0.080 | 2.45E-04 | 0.520 | 0.820 | 0.710 | 0.090 | 5.48E-03 | 0.560 | 0.900 |
| **triglyceride levels (in SD)** | - | - | - | - | - | - | - | - | - | - | 0.730 | 0.090 | 1.59E-02 | 0.570 | 0.940 |
| **glucose levels (in SD)** | - | - | - | - | - | - | - | - | - | - | 0.730 | 0.090 | 1.15E-02 | 0.570 | 0.930 |
| **intercept** | 0.120 | 0.030 | 4.79E-14 | 0.070 | 0.200 | 0.080 | 0.030 | 5.61E-16 | 0.050 | 0.150 | 0.090 | 0.030 | 4.75E-13 | 0.050 | 0.170 |

| | Many false positive vascular lipid deposits | | | | | | | | | | | | | | |
|---|---|---|---|---|---|---|---|---|---|---|---|---|---|---|---|
| | Model 1 (n=2043) | | | | | Model 2 (n=2043) | | | | | Model 3 (n=1848) | | | | |
| | OR | SE | *P* | lci | uci | OR | SE | *P* | lci | uci | OR | SE | *P* | lci | uci |
| **overfeeding** | 0.540 | 0.120 | 6.81E-03 | 0.350 | 0.840 | 0.620 | 0.150 | 5.05E-02 | 0.390 | 1.000 | 0.450 | 0.120 | 3.65E-03 | 0.270 | 0.770 |
| **cholesterol supplementation** | 0.940 | 0.270 | 8.35E-01 | 0.530 | 1.660 | 0.920 | 0.270 | 7.89E-01 | 0.520 | 1.640 | 0.960 | 0.300 | 8.92E-01 | 0.520 | 1.770 |
| **diethyl ether treatment** | 0.650 | 0.170 | 1.06E-01 | 0.380 | 1.100 | 0.650 | 0.180 | 1.11E-01 | 0.380 | 1.100 | 0.670 | 0.190 | 1.66E-01 | 0.380 | 1.180 |
| **time of day (in hours since 9AM)** | 1.040 | 0.040 | 3.49E-01 | 0.960 | 1.130 | 1.050 | 0.040 | 2.60E-01 | 0.970 | 1.140 | 1.060 | 0.050 | 1.98E-01 | 0.970 | 1.160 |
| **body length (in SD)** | - | - | - | - | - | 0.930 | 0.110 | 5.57E-01 | 0.740 | 1.180 | 0.710 | 0.090 | 1.02E-02 | 0.550 | 0.920 |
| **dorsal body surface area (in SD)** | - | - | - | - | - | 0.820 | 0.100 | 9.58E-02 | 0.650 | 1.040 | 0.730 | 0.100 | 1.95E-02 | 0.550 | 0.950 |
| **triglyceride levels (in SD)** | - | - | - | - | - | - | - | - | - | - | 2.610 | 0.360 | 4.40E-12 | 1.990 | 3.430 |
| **glucose levels (in SD)** | - | - | - | - | - | - | - | - | - | - | 1.020 | 0.130 | 8.65E-01 | 0.790 | 1.320 |
| **intercept** | 0.060 | 0.020 | 5.45E-19 | 0.030 | 0.120 | 0.060 | 0.020 | 4.50E-19 | 0.030 | 0.110 | 0.040 | 0.010 | 4.59E-19 | 0.020 | 0.080 |

*continued* **Supplementary Table 6**

| | Many false positive oxLDL deposits | | | | | | | | | | | | | | |
|---|---|---|---|---|---|---|---|---|---|---|---|---|---|---|---|
| | Model 1 (n=885) | | | | | Model 2 (n=885) | | | | | Model 3 (n=876) | | | | |
| | OR | SE | *P* | lci | uci | OR | SE | *P* | lci | uci | OR | SE | *P* | lci | uci |
| **overfeeding** | 1.050 | 0.180 | 7.51E-01 | 0.760 | 1.460 | 1.250 | 0.220 | 2.00E-01 | 0.890 | 1.770 | 1.020 | 0.200 | 9.32E-01 | 0.700 | 1.480 |
| **cholesterol supplementation** | 0.840 | 0.170 | 3.89E-01 | 0.560 | 1.260 | 0.900 | 0.190 | 6.26E-01 | 0.590 | 1.370 | 0.910 | 0.210 | 6.77E-01 | 0.580 | 1.420 |
| **diethyl ether treatment** | 0.680 | 0.130 | 4.50E-02 | 0.460 | 0.990 | 0.670 | 0.130 | 4.07E-02 | 0.450 | 0.980 | 0.560 | 0.120 | 6.40E-03 | 0.370 | 0.850 |
| **time of day (in hours since 9AM)** | 0.870 | 0.030 | 1.05E-04 | 0.810 | 0.930 | 0.900 | 0.030 | 3.55E-03 | 0.840 | 0.970 | 0.930 | 0.040 | 6.35E-02 | 0.860 | 1.000 |
| **body length (in SD)** | - | - | - | - | - | 1.260 | 0.130 | 2.47E-02 | 1.030 | 1.540 | 0.920 | 0.110 | 4.58E-01 | 0.730 | 1.150 |
| **dorsal body surface area (in SD)** | - | - | - | - | - | 0.700 | 0.060 | 2.61E-05 | 0.590 | 0.830 | 0.720 | 0.070 | 4.19E-04 | 0.600 | 0.860 |
| **triglyceride levels (in SD)** | - | - | - | - | - | - | - | - | - | - | 1.950 | 0.260 | 3.96E-07 | 1.500 | 2.520 |
| **glucose levels (in SD)** | - | - | - | - | - | - | - | - | - | - | 0.400 | 0.050 | 1.59E-14 | 0.320 | 0.500 |
| **intercept** | 0.790 | 0.200 | 3.46E-01 | 0.480 | 1.290 | 0.530 | 0.140 | 1.93E-02 | 0.310 | 0.900 | 0.400 | 0.110 | 1.32E-03 | 0.220 | 0.700 |

| | Many false negative macrophages | | | | | | | | | | | | | | |
|---|---|---|---|---|---|---|---|---|---|---|---|---|---|---|---|
| | Model 1 (n=1010) | | | | | Model 2 (n=1010) | | | | | Model 2 (n=589) | | | | |
| | OR | SE | *P* | lci | uci | OR | SE | *P* | lci | uci | OR | SE | *P* | lci | uci |
| **overfeeding** | 0.380 | 0.210 | 8.27E-02 | 0.130 | 1.130 | 0.470 | 0.280 | 2.04E-01 | 0.140 | 1.510 | 1.520 | 1.770 | 7.22E-01 | 0.150 | 14.920 |
| **cholesterol supplementation** | 0.390 | 0.330 | 2.68E-01 | 0.080 | 2.050 | 0.310 | 0.270 | 1.76E-01 | 0.060 | 1.680 | 0.480 | 0.590 | 5.53E-01 | 0.040 | 5.440 |
| **diethyl ether treatment** | 0.630 | 0.360 | 4.13E-01 | 0.200 | 1.920 | 0.550 | 0.330 | 3.15E-01 | 0.170 | 1.760 | - | - | - | - | - |
| **time of day (in hours since 9AM)** | 0.830 | 0.090 | 7.57E-02 | 0.680 | 1.020 | 0.870 | 0.090 | 1.84E-01 | 0.710 | 1.070 | 0.650 | 0.190 | 1.45E-01 | 0.360 | 1.160 |
| **body length (in SD)** | - | - | - | - | - | 0.510 | 0.150 | 2.19E-02 | 0.280 | 0.910 | 1.450 | 0.890 | 5.41E-01 | 0.440 | 4.800 |
| **dorsal body surface area (in SD)** | - | - | - | - | - | 1.580 | 0.480 | 1.35E-01 | 0.870 | 2.860 | 1.150 | 0.700 | 8.22E-01 | 0.350 | 3.790 |
| **triglyceride levels (in SD)** | - | - | - | - | - | - | - | - | - | - | 0.260 | 0.180 | 4.62E-02 | 0.070 | 0.980 |
| **glucose levels (in SD)** | - | - | - | - | - | - | - | - | - | - | 2.080 | 1.360 | 2.63E-01 | 0.580 | 7.500 |
| **intercept** | 0.090 | 0.050 | 1.76E-05 | 0.030 | 0.270 | 0.070 | 0.040 | 3.40E-06 | 0.020 | 0.220 | 0.060 | 0.090 | 8.53E-02 | 0.000 | 1.490 |

| | Many false negative neutrophils | | | | | | | | | | | | | | |
|---|---|---|---|---|---|---|---|---|---|---|---|---|---|---|---|
| | Model 1 (n=537) | | | | | Model 2 (n=537) | | | | | Model 3 (n=416) | | | | |
| | OR | SE | *P* | lci | uci | OR | SE | *P* | lci | uci | OR | SE | *P* | lci | uci |
| **overfeeding** | 1.110 | 0.360 | 7.38E-01 | 0.590 | 2.110 | 1.380 | 0.500 | 3.69E-01 | 0.680 | 2.810 | 4.270 | 2.750 | 2.41E-02 | 1.210 | 15.110 |
| **cholesterol supplementation** | 0.730 | 0.310 | 4.66E-01 | 0.320 | 1.680 | 0.650 | 0.280 | 3.12E-01 | 0.280 | 1.510 | 0.880 | 0.600 | 8.53E-01 | 0.230 | 3.350 |
| **diethyl ether treatment** | 0.950 | 0.370 | 8.89E-01 | 0.440 | 2.060 | 0.830 | 0.340 | 6.46E-01 | 0.370 | 1.850 | 6.440 | 7.440 | 1.07E-01 | 0.670 | 62.010 |
| **time of day (in hours since 9AM)** | 0.790 | 0.050 | 4.24E-04 | 0.690 | 0.900 | 0.810 | 0.060 | 1.76E-03 | 0.700 | 0.920 | 0.710 | 0.130 | 5.77E-02 | 0.500 | 1.010 |
| **body length (in SD)** | - | - | - | - | - | 0.630 | 0.120 | 1.11E-02 | 0.440 | 0.900 | 0.820 | 0.260 | 5.40E-01 | 0.440 | 1.540 |
| **dorsal body surface area (in SD)** | - | - | - | - | - | 1.190 | 0.260 | 4.16E-01 | 0.780 | 1.820 | 0.570 | 0.220 | 1.41E-01 | 0.270 | 1.200 |
| **triglyceride levels (in SD)** | - | - | - | - | - | - | - | - | - | - | 0.270 | 0.090 | 6.30E-05 | 0.140 | 0.510 |
| **glucose levels (in SD)** | - | - | - | - | - | - | - | - | - | - | 1.140 | 0.450 | 7.44E-01 | 0.520 | 2.470 |
| **intercept** | 0.300 | 0.110 | 9.49E-04 | 0.150 | 0.610 | 0.240 | 0.090 | 2.03E-04 | 0.110 | 0.510 | 0.030 | 0.020 | 1.08E-04 | 0.000 | 0.160 |

*continued* **Supplementary Table 6**

| | Circulating neutrophils present in the z-stack | | | | | | | | | | | | | | |
|---|---|---|---|---|---|---|---|---|---|---|---|---|---|---|---|
| | Model 1 (n=504) | | | | | Model 2 (n=504) | | | | | Model 3 (n=404) | | | | |
| | OR | SE | *P* | lci | uci | OR | SE | *P* | lci | uci | OR | SE | *P* | lci | uci |
| **overfeeding** | 1.740 | 1.140 | 3.99E-01 | 0.480 | 6.290 | 1.660 | 1.260 | 5.04E-01 | 0.380 | 7.340 | 2.630 | 2.150 | 2.35E-01 | 0.530 | 13.010 |
| **cholesterol supplementation** | 0.980 | 0.980 | 9.80E-01 | 0.140 | 7.020 | 1.180 | 1.210 | 8.69E-01 | 0.160 | 8.740 | 1.580 | 1.700 | 6.70E-01 | 0.190 | 12.940 |
| **diethyl ether treatment** | 0.300 | 0.250 | 1.46E-01 | 0.060 | 1.520 | 0.330 | 0.280 | 1.97E-01 | 0.060 | 1.790 | 0.380 | 0.350 | 2.90E-01 | 0.060 | 2.270 |
| **time of day (in hours since 9AM)** | 0.970 | 0.110 | 7.58E-01 | 0.780 | 1.200 | 0.960 | 0.110 | 7.48E-01 | 0.770 | 1.210 | 0.990 | 0.130 | 9.26E-01 | 0.760 | 1.280 |
| **body length (in SD)** | - | - | - | - | - | 1.580 | 0.600 | 2.25E-01 | 0.750 | 3.320 | 1.410 | 0.610 | 4.26E-01 | 0.610 | 3.270 |
| **dorsal body surface area (in SD)** | - | - | - | - | - | 0.650 | 0.290 | 3.37E-01 | 0.260 | 1.580 | 0.570 | 0.280 | 2.60E-01 | 0.220 | 1.510 |
| **triglyceride levels (in SD)** | - | - | - | - | - | - | - | - | - | - | 1.530 | 0.750 | 3.85E-01 | 0.590 | 4.010 |
| **glucose levels (in SD)** | - | - | - | - | - | - | - | - | - | - | 1.180 | 0.550 | 7.21E-01 | 0.470 | 2.960 |
| **intercept** | 0.040 | 0.030 | 2.24E-05 | 0.010 | 0.170 | 0.030 | 0.030 | 3.19E-05 | 0.010 | 0.160 | 0.010 | 0.010 | 7.23E-05 | 0.000 | 0.110 |

Associations are shown for criteria that resulted in the exclusion of at least 10 larvae. Vasculature not properly detected typically results from weak staining, possibly due to low levels of circulating lipids; Many false positives: >20% of true negative objects were falsely detected by the qualtification pipeline; Many false negatives: <20% of true positive objects were detected by the qualtification pipeline. Associations were examined using logistic regression models. Model 1: adjusted for use of diethyl ether (to prepare the diet) and time of day; Model 2: additionally adjusted for body length and dorsal body surface area; Model 3: additionally adjusted for whole-body triglyceride and glucose levels. Dorsal body surface area was normalized for body length; whole-body triglyceride and glucose levels were normalized for protein level. Adjusting for transgenic background and batch would have excluded approximately half the larvae. Effects shown for overfeeding, cholesterol supplementation and diethyl ether treatment are compared with unexposed controls. Lci and uci are lower and upper boundaries of the 95% confidence interval.

**Supplementary Table 7 - The effect of treatment with atorvastatin and ezetimibe on body size**

| | | Body length (n=1004) | | | | |
|---|---|---|---|---|---|---|
| | | **Effect** | **SE** | ***P*** | **lci** | **uci** |
| **fixed factors** | **atorvastatin and ezetimibe** | -0.025 | 0.053 | 6.29E-01 | -0.128 | 0.078 |
| | **time of day (in hours since 9AM)** | 0.000 | 0.012 | 9.78E-01 | -0.023 | 0.024 |
| | **intercept** | 0.012 | 0.162 | 9.41E-01 | -0.305 | 0.330 |
| **random factors** | ***variation by transgenic background*** | *0.000* | - | - | - | - |
| | ***variation by batch*** | *0.495* | *0.147* | - | *0.276* | *0.886* |
| | ***residual*** | *0.794* | *0.025* | - | *0.747* | *0.845* |

| | | Dorsal body surface area (n=1004) | | | | |
|---|---|---|---|---|---|---|
| | | **Effect** | **SE** | ***P*** | **lci** | **uci** |
| **fixed factors** | **atorvastatin and ezetimibe** | -0.135 | 0.055 | 1.48E-02 | -0.244 | -0.026 |
| | **time of day (in hours since 9AM)** | 0.046 | 0.013 | 2.72E-04 | 0.021 | 0.071 |
| | **intercept** | -0.185 | 0.335 | 5.81E-01 | -0.842 | 0.472 |
| **random factors** | ***variation by transgenic background*** | *0.561* | *0.239* | - | *0.244* | *1.291* |
| | ***variation by batch*** | *0.176* | *0.057* | - | *0.093* | *0.330* |
| | ***residual*** | *0.842* | *0.019* | - | *0.806* | *0.880* |

| | | Lateral body surface area (n=553) | | | | |
|---|---|---|---|---|---|---|
| | | **Effect** | **SE** | ***P*** | **lci** | **uci** |
| **fixed factors** | **atorvastatin and ezetimibe** | -0.233 | 0.088 | 8.05E-03 | -0.406 | -0.061 |
| | **time of day (in hours since 9AM)** | 0.022 | 0.019 | 2.61E-01 | -0.016 | 0.059 |
| | **intercept** | -0.011 | 0.187 | 9.54E-01 | -0.376 | 0.355 |
| **random factors** | ***variation by transgenic background*** | *0.221* | *0.122* | - | *0.075* | *0.654* |
| | ***variation by batch*** | *0.066* | *0.079* | - | *0.006* | *0.678* |
| | ***residual*** | *0.972* | *0.029* | - | *0.916* | *1.031* |

| | | Body volume (n=512) | | | | |
|---|---|---|---|---|---|---|
| | | **Effect** | **SE** | ***P*** | **lci** | **uci** |
| **fixed factors** | **atorvastatin and ezetimibe** | -0.314 | 0.087 | 2.95E-04 | -0.484 | -0.144 |
| | **time of day (in hours since 9AM)** | 0.024 | 0.019 | 2.19E-01 | -0.014 | 0.061 |
| | **intercept** | -0.026 | 0.350 | 9.40E-01 | -0.712 | 0.660 |
| **random factors** | ***variation by transgenic background*** | *0.458* | *0.249* | - | *0.158* | *1.332* |
| | ***variation by batch*** | *0.209* | *0.101* | - | *0.082* | *0.538* |
| | ***residual*** | *0.922* | *0.029* | - | *0.867* | *0.981* |

Dorsal and lateral body surface area and body volume were normalized for body length using residuals. All outcomes were inverse-normally transformed before the analysis. Associations were examined using hierarchical linear models. Effects shown for atorvastatin and ezetimibe treatment are compared with unexposed controls. Lci and uci are lower and upper boundaries of the 95% confidence interval.

**Supplementary Table 8 - The effect of treatment with atorvastatin and ezetimibe on whole-body lipid and glucose levels**

| | | Effect | SE | *P* | lci | uci |
|---|---|---|---|---|---|---|
| | | **LDL cholesterol levels (n=567)** | | | | |
| | | **Effect** | **SE** | ***P*** | **lci** | **uci** |
| **fixed factors** | **atorvastatin and ezetimibe** | -0.544 | 0.079 | 5.22E-12 | -0.699 | -0.390 |
| | **time of day (in hours since 9AM)** | 0.017 | 0.017 | 3.28E-01 | -0.017 | 0.050 |
| | **intercept** | 0.173 | 0.343 | 6.13E-01 | -0.499 | 0.846 |
| **random factors** | ***variation by transgenic background*** | *0.461* | *0.242* | - | *0.165* | *1.288* |
| | ***variation by batch*** | *0.147* | *0.071* | - | *0.057* | *0.378* |
| | ***residual*** | *0.861* | *0.026* | - | *0.812* | *0.913* |
| | | **HDL cholesterol levels (n=564)** | | | | |
| | | **Effect** | **SE** | ***P*** | **lci** | **uci** |
| **fixed factors** | **atorvastatin and ezetimibe** | 0.043 | 0.078 | 5.78E-01 | -0.109 | 0.195 |
| | **time of day (in hours since 9AM)** | 0.018 | 0.017 | 2.91E-01 | -0.015 | 0.050 |
| | **intercept** | -0.064 | 0.387 | 8.68E-01 | -0.822 | 0.694 |
| **random factors** | ***variation by transgenic background*** | *0.516* | *0.278* | - | *0.180* | *1.483* |
| | ***variation by batch*** | *0.228* | *0.094* | - | *0.101* | *0.512* |
| | ***residual*** | *0.837* | *0.025* | - | *0.789* | *0.888* |
| | | **Triglyceride levels (n=1005)** | | | | |
| | | **Effect** | **SE** | ***P*** | **lci** | **uci** |
| **fixed factors** | **atorvastatin and ezetimibe** | -0.245 | 0.055 | 8.68E-06 | -0.353 | -0.137 |
| | **time of day (in hours since 9AM)** | 0.046 | 0.013 | 2.29E-04 | 0.022 | 0.071 |
| | **intercept** | -0.133 | 0.198 | 5.04E-01 | -0.522 | 0.256 |
| **random factors** | ***variation by transgenic background*** | *0.174* | *0.351* | - | *0.003* | *9.122* |
| | ***variation by batch*** | *0.521* | *0.145* | - | *0.302* | *0.899* |
| | ***residual*** | *0.832* | *0.019* | - | *0.796* | *0.869* |
| | | **Total cholesterol levels (n=1005)** | | | | |
| | | **Effect** | **SE** | ***P*** | **lci** | **uci** |
| **fixed factors** | **atorvastatin and ezetimibe** | -0.821 | 0.051 | 1.23E-58 | -0.920 | -0.721 |
| | **time of day (in hours since 9AM)** | 0.060 | 0.012 | 2.39E-07 | 0.037 | 0.083 |
| | **intercept** | 0.089 | 0.283 | 7.54E-01 | -0.467 | 0.644 |
| **random factors** | ***variation by transgenic background*** | *0.443* | *0.221* | - | *0.166* | *1.178* |
| | ***variation by batch*** | *0.344* | *0.092* | - | *0.203* | *0.581* |
| | ***residual*** | *0.768* | *0.017* | - | *0.735* | *0.803* |
| | | **Glucose levels (n=1008)** | | | | |
| | | **Effect** | **SE** | ***P*** | **lci** | **uci** |
| **fixed factors** | **atorvastatin and ezetimibe** | 0.199 | 0.053 | 1.92E-04 | 0.094 | 0.303 |
| | **time of day (in hours since 9AM)** | -0.018 | 0.012 | 1.34E-01 | -0.042 | 0.006 |
| | **intercept** | -0.088 | 0.313 | 7.78E-01 | -0.702 | 0.525 |
| **random factors** | ***variation by transgenic background*** | *0.520* | *0.223* | - | *0.224* | *1.206* |
| | ***variation by batch*** | *0.204* | *0.059* | - | *0.115* | *0.360* |
| | ***residual*** | *0.810* | *0.018* | - | *0.775* | *0.847* |

All outcomes were normalized for protein level using residuals, and inverse-normally transformed before the analysis. Associations were examined using hierarchical linear models and were adjusted for time of day, transgenic background and batch. Effects shown for treatment with atorvastatin and ezetimibe are compared with untreated controls. Lci and uci are lower and upper boundaries of the 95% confidence interval.

**Supplementary Table 9 - The effect of atorvastatin and ezetimibe on image-based vascular atherogenic traits**

| | | Vascular lipid deposition | | | | | | | | | | | | | | | | | | | |
|---|---|---|---|---|---|---|---|---|---|---|---|---|---|---|---|---|---|---|---|---|---|
| | | Model 1 (n=776) | | | | | Model 2 (n=776) | | | | | Model 3 (n=728) | | | | | Model 4 (n=344) | | | | |
| | | Effect | SE | *P* | lci | uci | Effect | SE | *P* | lci | uci | Effect | SE | *P* | lci | uci | Effect | SE | *P* | lci | uci |
| **negative binomial terms** | **atorvastatin and ezetimibe** | -1.523 | 0.176 | 4.75E-18 | -1.868 | -1.178 | -1.496 | 0.177 | 2.32E-17 | -1.842 | -1.150 | -1.423 | 0.180 | 3.04E-15 | -1.776 | -1.069 | -2.001 | 0.299 | 2.18E-11 | -2.587 | -1.415 |
| | **time of day (in hours since 9AM)** | 0.155 | 0.038 | 4.83E-05 | 0.080 | 0.229 | 0.145 | 0.037 | 1.07E-04 | 0.072 | 0.219 | 0.151 | 0.040 | 1.53E-04 | 0.073 | 0.229 | 0.327 | 0.054 | 1.88E-09 | 0.221 | 0.434 |
| | **body length (in SD)** | - | - | - | - | - | -0.222 | 0.119 | 6.27E-02 | -0.456 | 0.012 | -0.221 | 0.120 | 6.43E-02 | -0.455 | 0.013 | 0.352 | 0.243 | 1.48E-01 | -0.125 | 0.829 |
| | **dorsal body surface area (in SD)** | - | - | - | - | - | 0.183 | 0.102 | 7.15E-02 | -0.016 | 0.383 | 0.118 | 0.120 | 3.28E-01 | -0.118 | 0.354 | 0.179 | 0.193 | 3.51E-01 | -0.198 | 0.557 |
| | **LDL cholesterol levels (in SD)** | - | - | - | - | - | - | - | - | - | - | - | - | - | - | - | -0.231 | 0.149 | 1.21E-01 | -0.524 | 0.061 |
| | **HDL cholesterol levels (in SD)** | - | - | - | - | - | - | - | - | - | - | - | - | - | - | - | -0.216 | 0.133 | 1.06E-01 | -0.477 | 0.046 |
| | **triglyceride levels (in SD)** | - | - | - | - | - | - | - | - | - | - | 0.178 | 0.119 | 1.36E-01 | -0.056 | 0.412 | 0.426 | 0.216 | 4.87E-02 | 0.002 | 0.849 |
| | **glucose levels (in SD)** | - | - | - | - | - | - | - | - | - | - | -0.125 | 0.103 | 2.27E-01 | -0.327 | 0.078 | -0.159 | 0.231 | 4.91E-01 | -0.611 | 0.293 |
| | **Tg(flk:EGFP) carriers vs. Tg(mpo:EGFP; mpeg1:mCherry) carriers** | -0.915 | 0.335 | 6.37E-03 | -1.572 | -0.257 | -0.823 | 0.360 | 2.22E-02 | -1.527 | -0.118 | -0.808 | 0.383 | 3.50E-02 | -1.560 | -0.057 | - | - | - | - | - |
| | **Tg(hsp70:IK17:EGFP; mpeg1:mCherry) carriers vs. Tg(mpo:EGFP; mpeg1:mCherry) carriers** | -0.477 | 0.245 | 5.17E-02 | -0.958 | 0.003 | 0.103 | 0.353 | 7.72E-01 | -0.590 | 0.795 | -0.020 | 0.403 | 9.60E-01 | -0.810 | 0.770 | 0.562 | 0.606 | 3.54E-01 | -0.626 | 1.751 |
| | **batch 1** | -0.727 | 0.456 | 1.11E-01 | -1.621 | 0.167 | -0.600 | 0.453 | 1.85E-01 | -1.488 | 0.288 | -0.263 | 0.497 | 5.97E-01 | -1.237 | 0.711 | - | - | - | - | - |
| | **batch 2** | -0.480 | 0.330 | 1.47E-01 | -1.127 | 0.168 | -0.240 | 0.349 | 4.92E-01 | -0.924 | 0.444 | 0.182 | 0.376 | 6.28E-01 | -0.555 | 0.920 | - | - | - | - | - |
| | **batch 3** | -1.264 | 0.319 | 7.40E-05 | -1.889 | -0.639 | -1.204 | 0.311 | 1.08E-04 | -1.814 | -0.595 | -0.895 | 0.361 | 1.32E-02 | -1.603 | -0.187 | - | - | - | - | - |
| | **batch 4** | -0.678 | 0.325 | 3.70E-02 | -1.316 | -0.041 | -0.611 | 0.331 | 6.52E-02 | -1.260 | 0.039 | -0.322 | 0.381 | 3.97E-01 | -1.069 | 0.424 | - | - | - | - | - |
| | **batch 5** | -0.385 | 0.377 | 3.07E-01 | -1.125 | 0.355 | -0.504 | 0.388 | 1.94E-01 | -1.265 | 0.257 | -0.427 | 0.384 | 2.66E-01 | -1.179 | 0.325 | - | - | - | - | - |
| | **batch 6** | -0.252 | 0.384 | 5.11E-01 | -1.005 | 0.500 | -0.509 | 0.392 | 1.93E-01 | -1.277 | 0.258 | -0.492 | 0.397 | 2.16E-01 | -1.271 | 0.287 | 0.269 | 0.458 | 5.57E-01 | -0.629 | 1.167 |
| | **batch 7** | - | - | - | - | - | - | - | - | - | - | - | - | - | - | - | 0.124 | 0.454 | 7.85E-01 | -0.766 | 1.014 |
| | **batch 8** | -2.432 | 0.386 | 3.12E-10 | -3.189 | -1.674 | -2.576 | 0.374 | 5.73E-12 | -3.309 | -1.843 | -2.513 | 0.374 | 1.85E-11 | -3.246 | -1.780 | -2.101 | 0.479 | 1.14E-05 | -3.040 | -1.163 |
| | **intercept** | 3.849 | 0.268 | 1.31E-46 | 3.322 | 4.375 | 3.740 | 0.274 | 2.47E-42 | 3.202 | 4.277 | 3.570 | 0.298 | 4.95E-33 | 2.986 | 4.155 | 1.707 | 0.455 | 1.75E-04 | 0.815 | 2.598 |

| | | Vascular accumulation of oxLDL | | | | | | | | | | | | | | | | | | | |
|---|---|---|---|---|---|---|---|---|---|---|---|---|---|---|---|---|---|---|---|---|---|
| | | Model 1 (n=236) | | | | | Model 2 (n=236) | | | | | Model 3 (n=233) | | | | | Model 4 (n=229) | | | | |
| | | Effect | SE | *P* | lci | uci | Effect | SE | *P* | lci | uci | Effect | SE | *P* | lci | uci | Effect | SE | *P* | lci | uci |
| **fixed factors** | **atorvastatin and ezetimibe** | 0.019 | 0.126 | 8.78E-01 | -0.228 | 0.267 | 0.100 | 0.126 | 4.29E-01 | -0.147 | 0.347 | 0.099 | 0.127 | 4.35E-01 | -0.150 | 0.349 | 0.130 | 0.130 | 3.19E-01 | -0.126 | 0.385 |
| | **time of day (in hours since 9AM)** | -0.017 | 0.024 | 4.94E-01 | -0.064 | 0.031 | -0.018 | 0.024 | 4.45E-01 | -0.064 | 0.028 | -0.028 | 0.024 | 2.44E-01 | -0.075 | 0.019 | -0.031 | 0.024 | 1.96E-01 | -0.079 | 0.016 |
| | **body length (in SD)** | - | - | - | - | - | 0.111 | 0.085 | 1.90E-01 | -0.055 | 0.278 | 0.143 | 0.094 | 1.30E-01 | -0.042 | 0.328 | 0.157 | 0.096 | 1.01E-01 | -0.031 | 0.344 |
| | **dorsal body surface area (in SD)** | - | - | - | - | - | 0.189 | 0.063 | 2.46E-03 | 0.067 | 0.312 | 0.206 | 0.081 | 1.14E-02 | 0.046 | 0.365 | 0.220 | 0.081 | 6.84E-03 | 0.060 | 0.379 |
| | **LDL cholesterol levels (in SD)** | - | - | - | - | - | - | - | - | - | - | - | - | - | - | - | 0.007 | 0.066 | 9.16E-01 | -0.122 | 0.135 |
| | **HDL cholesterol levels (in SD)** | - | - | - | - | - | - | - | - | - | - | - | - | - | - | - | 0.127 | 0.067 | 5.80E-02 | -0.004 | 0.258 |
| | **triglyceride levels (in SD)** | - | - | - | - | - | - | - | - | - | - | -0.010 | 0.087 | 9.06E-01 | -0.181 | 0.160 | 0.014 | 0.092 | 8.76E-01 | -0.166 | 0.195 |
| | **glucose levels (in SD)** | - | - | - | - | - | - | - | - | - | - | 0.152 | 0.096 | 1.13E-01 | -0.036 | 0.341 | 0.096 | 0.108 | 3.76E-01 | -0.116 | 0.308 |
| | **intercept** | 0.169 | 0.256 | 5.09E-01 | -0.333 | 0.672 | 0.261 | 0.256 | 3.10E-01 | -0.242 | 0.763 | 0.433 | 0.280 | 1.22E-01 | -0.116 | 0.981 | 0.323 | 0.299 | 2.81E-01 | -0.264 | 0.909 |
| **random factors** | ***variation by batch*** | *0.378* | *0.169* | - | *0.157* | *0.907* | *0.373* | *0.173* | - | *0.151* | *0.924* | *0.382* | *0.177* | - | *0.154* | *0.948* | *0.400* | *0.185* | - | *0.162* | *0.988* |
| | ***residual*** | *0.839* | *0.039* | - | *0.767* | *0.919* | *0.820* | *0.038* | - | *0.749* | *0.898* | *0.818* | *0.038* | - | *0.746* | *0.896* | *0.811* | *0.038* | - | *0.739* | *0.889* |

| | | Vascular infiltration by macrophages | | | | | | | | | | | | | | | | | | | |
|---|---|---|---|---|---|---|---|---|---|---|---|---|---|---|---|---|---|---|---|---|---|
| | | Model 1 (n=633) | | | | | Model 2 (n=633) | | | | | Model 3 (n=585) | | | | | Model 4 (n=224) | | | | |
| | | Effect | SE | *P* | lci | uci | Effect | SE | *P* | lci | uci | Effect | SE | *P* | lci | uci | Effect | SE | *P* | lci | uci |
| **fixed factors** | **atorvastatin and ezetimibe** | -0.001 | 0.049 | 9.80E-01 | -0.097 | 0.094 | 0.005 | 0.049 | 9.17E-01 | -0.090 | 0.101 | 0.010 | 0.051 | 8.50E-01 | -0.090 | 0.109 | -0.003 | 0.072 | 9.64E-01 | -0.144 | 0.138 |
| | **time of day (in hours since 9AM)** | 0.036 | 0.011 | 9.87E-04 | 0.015 | 0.058 | 0.033 | 0.011 | 3.11E-03 | 0.011 | 0.054 | 0.029 | 0.011 | 1.00E-02 | 0.007 | 0.052 | 0.009 | 0.013 | 4.96E-01 | -0.017 | 0.035 |
| | **body length (in SD)** | - | - | - | - | - | 0.001 | 0.031 | 9.81E-01 | -0.060 | 0.061 | 0.011 | 0.032 | 7.36E-01 | -0.051 | 0.073 | 0.065 | 0.053 | 2.20E-01 | -0.039 | 0.169 |
| | **dorsal body surface area (in SD)** | - | - | - | - | - | 0.059 | 0.029 | 4.03E-02 | 0.003 | 0.116 | 0.043 | 0.032 | 1.82E-01 | -0.020 | 0.105 | 0.022 | 0.044 | 6.14E-01 | -0.065 | 0.109 |
| | **LDL cholesterol levels (in SD)** | - | - | - | - | - | - | - | - | - | - | - | - | - | - | - | -0.040 | 0.036 | 2.71E-01 | -0.110 | 0.031 |
| | **HDL cholesterol levels (in SD)** | - | - | - | - | - | - | - | - | - | - | - | - | - | - | - | 0.031 | 0.037 | 4.08E-01 | -0.042 | 0.104 |
| | **triglyceride levels (in SD)** | - | - | - | - | - | - | - | - | - | - | 0.027 | 0.030 | 3.68E-01 | -0.032 | 0.087 | 0.091 | 0.051 | 7.67E-02 | -0.010 | 0.191 |
| | **glucose levels (in SD)** | - | - | - | - | - | - | - | - | - | - | 0.011 | 0.025 | 6.76E-01 | -0.039 | 0.060 | -0.007 | 0.059 | 9.10E-01 | -0.122 | 0.109 |
| | **intercept** | -0.285 | 0.505 | 5.73E-01 | -1.275 | 0.706 | -0.252 | 0.486 | 6.04E-01 | -1.204 | 0.701 | -0.238 | 0.483 | 6.22E-01 | -1.186 | 0.709 | -0.877 | 0.196 | 7.53E-06 | -1.261 | -0.493 |
| **random factors** | ***variation by transgenic background*** | *0.699* | *0.361* | - | *0.253* | *1.925* | *0.670* | *0.348* | - | *0.242* | *1.857* | *0.666* | *0.347* | - | *0.240* | *1.849* | *0.000* | *0.000* | - | *0.000* | - |
| | ***variation by batch*** | *0.241* | *0.075* | - | *0.131* | *0.442* | *0.247* | *0.077* | - | *0.134* | *0.457* | *0.246* | *0.077* | - | *0.133* | *0.456* | *0.289* | *0.126* | - | *0.123* | *0.680* |
| | ***residuals*** | *0.583* | *0.017* | - | *0.552* | *0.617* | *0.581* | *0.016* | - | *0.550* | *0.614* | *0.567* | *0.017* | - | *0.536* | *0.601* | *0.438* | *0.021* | - | *0.399* | *0.481* |

***continued*** **Supplementary Table 9**

| | | **Vascular co-localization of lipids with macrophages** | | | | | | | | | | | | | | | | | | | |
|---|---|---|---|---|---|---|---|---|---|---|---|---|---|---|---|---|---|---|---|---|---|
| | | **Model 1 (n=549)** | | | | | **Model 2 (n=549)** | | | | | **Model 3 (n=502)** | | | | | **Model 4 (n=157)** | | | | |
| | | **Effect** | **SE** | ***P*** | **lci** | **uci** | **Effect** | **SE** | ***P*** | **lci** | **uci** | **Effect** | **SE** | ***P*** | **lci** | **uci** | **Effect** | **SE** | ***P*** | **lci** | **uci** |
| **negative binomial terms** | **atorvastatin and ezetimibe** | -1.341 | 0.241 | 2.46E-08 | -1.813 | -0.870 | -1.383 | 0.237 | 5.74E-09 | -1.848 | -0.917 | -1.226 | 0.252 | 1.14E-06 | -1.720 | -0.732 | -2.992 | 0.730 | 4.10E-05 | -4.422 | -1.562 |
| | **time of day (in hours since 9AM)** | 0.122 | 0.052 | 1.85E-02 | 0.021 | 0.224 | 0.116 | 0.054 | 3.25E-02 | 0.010 | 0.222 | 0.120 | 0.059 | 4.30E-02 | 0.004 | 0.237 | 0.388 | 0.125 | 1.98E-03 | 0.142 | 0.633 |
| | **body length (in SD)** | - | - | - | - | - | -0.471 | 0.161 | 3.50E-03 | -0.787 | -0.155 | -0.477 | 0.170 | 4.99E-03 | -0.810 | -0.144 | 1.020 | 0.539 | 5.83E-02 | -0.036 | 2.076 |
| | **dorsal body surface area (in SD)** | - | - | - | - | - | 0.262 | 0.156 | 9.32E-02 | -0.044 | 0.569 | 0.152 | 0.187 | 4.16E-01 | -0.215 | 0.519 | 1.194 | 0.494 | 1.58E-02 | 0.225 | 2.163 |
| | **LDL cholesterol levels (in SD)** | - | - | - | - | - | - | - | - | - | - | - | - | - | - | - | -0.696 | 0.339 | 4.02E-02 | -1.361 | -0.031 |
| | **HDL cholesterol levels (in SD)** | - | - | - | - | - | - | - | - | - | - | - | - | - | - | - | -0.279 | 0.243 | 2.51E-01 | -0.755 | 0.197 |
| | **triglyceride levels (in SD)** | - | - | - | - | - | - | - | - | - | - | 0.223 | 0.181 | 2.19E-01 | -0.133 | 0.578 | 0.252 | 0.501 | 6.14E-01 | -0.729 | 1.234 |
| | **glucose levels (in SD)** | - | - | - | - | - | - | - | - | - | - | -0.129 | 0.116 | 2.65E-01 | -0.356 | 0.098 | 1.106 | 0.562 | 4.92E-02 | 0.004 | 2.209 |
| | **Tg(hsp70:IK17:EGFP; mpeg1:mCherry) carriers vs. Tg(mpo:EGFP; mpeg1:mCherry) carriers** | -1.007 | 0.316 | 1.44E-03 | -1.626 | -0.387 | 0.019 | 0.477 | 9.69E-01 | -0.917 | 0.954 | -0.099 | 0.488 | 8.40E-01 | -1.055 | 0.858 | - | - | - | - | - |
| | **batch 1** | -1.223 | 0.459 | 7.65E-03 | -2.122 | -0.324 | -1.133 | 0.448 | 1.15E-02 | -2.011 | -0.254 | -0.670 | 0.516 | 1.94E-01 | -1.681 | 0.340 | - | - | - | - | - |
| | **batch 2** | -0.512 | 0.334 | 1.26E-01 | -1.167 | 0.143 | -0.113 | 0.374 | 7.63E-01 | -0.847 | 0.621 | 0.365 | 0.409 | 3.72E-01 | -0.437 | 1.166 | - | - | - | - | - |
| | **batch 3** | -0.964 | 0.393 | 1.41E-02 | -1.735 | -0.194 | -0.938 | 0.383 | 1.43E-02 | -1.688 | -0.187 | -0.615 | 0.454 | 1.75E-01 | -1.504 | 0.274 | - | - | - | - | - |
| | **batch 4** | -0.717 | 0.316 | 2.32E-02 | -1.337 | -0.098 | -0.553 | 0.346 | 1.09E-01 | -1.231 | 0.124 | -0.117 | 0.427 | 7.85E-01 | -0.955 | 0.721 | - | - | - | - | - |
| | **batch 8** | -5.338 | 0.946 | 1.67E-08 | -7.192 | -3.484 | -5.784 | 0.956 | 1.45E-09 | -7.658 | -3.911 | -5.495 | 0.952 | 7.76E-09 | -7.361 | -3.630 | - | - | - | - | - |
| | **batch 10** | - | - | - | - | - | - | - | - | - | - | - | - | - | - | - | 4.420 | 0.874 | 4.25E-07 | 2.707 | 6.133 |
| | **intercept** | 2.892 | 0.339 | 1.61E-17 | 2.226 | 3.557 | 2.734 | 0.353 | 1.02E-14 | 2.041 | 3.426 | 2.467 | 0.358 | 5.73E-12 | 1.765 | 3.169 | -2.943 | 0.840 | 4.56E-04 | -4.589 | -1.298 |

| | | **Vascular co-localization of oxLDL with macrophages** | | | | | | | | | | | | | | | | | | | |
|---|---|---|---|---|---|---|---|---|---|---|---|---|---|---|---|---|---|---|---|---|---|
| | | **Model 1 (n=212)** | | | | | **Model 2 (n=212)** | | | | | **Model 3 (n=209)** | | | | | **Model 4 (n=205)** | | | | |
| | | **Effect** | **SE** | ***P*** | **lci** | **uci** | **Effect** | **SE** | ***P*** | **lci** | **uci** | **Effect** | **SE** | ***P*** | **lci** | **uci** | **Effect** | **SE** | ***P*** | **lci** | **uci** |
| **negative binomial terms** | **atorvastatin and ezetimibe** | 0.332 | 0.166 | 4.48E-02 | 0.008 | 0.656 | 0.392 | 0.163 | 1.61E-02 | 0.073 | 0.711 | 0.388 | 0.163 | 1.72E-02 | 0.069 | 0.707 | 0.285 | 0.166 | 8.57E-02 | -0.040 | 0.610 |
| | **time of day (in hours since 9AM)** | -0.019 | 0.027 | 4.88E-01 | -0.071 | 0.034 | -0.027 | 0.027 | 3.25E-01 | -0.080 | 0.027 | -0.036 | 0.028 | 1.91E-01 | -0.091 | 0.018 | -0.035 | 0.028 | 2.10E-01 | -0.091 | 0.020 |
| | **body length (in SD)** | - | - | - | - | - | 0.154 | 0.094 | 1.03E-01 | -0.031 | 0.339 | 0.042 | 0.119 | 7.25E-01 | -0.192 | 0.276 | 0.048 | 0.120 | 6.88E-01 | -0.187 | 0.284 |
| | **dorsal body surface area (in SD)** | - | - | - | - | - | 0.200 | 0.074 | 6.56E-03 | 0.056 | 0.345 | 0.075 | 0.096 | 4.36E-01 | -0.114 | 0.264 | 0.061 | 0.096 | 5.28E-01 | -0.128 | 0.250 |
| | **LDL cholesterol levels (in SD)** | - | - | - | - | - | - | - | - | - | - | - | - | - | - | - | -0.122 | 0.069 | 7.59E-02 | -0.257 | 0.013 |
| | **HDL cholesterol levels (in SD)** | - | - | - | - | - | - | - | - | - | - | - | - | - | - | - | 0.021 | 0.078 | 7.86E-01 | -0.132 | 0.174 |
| | **triglyceride levels (in SD)** | - | - | - | - | - | - | - | - | - | - | 0.237 | 0.125 | 5.72E-02 | -0.007 | 0.481 | 0.303 | 0.132 | 2.16E-02 | 0.044 | 0.561 |
| | **glucose levels (in SD)** | - | - | - | - | - | - | - | - | - | - | -0.102 | 0.107 | 3.39E-01 | -0.311 | 0.107 | -0.177 | 0.120 | 1.41E-01 | -0.412 | 0.058 |
| | **batch 9** | -0.359 | 0.211 | 8.86E-02 | -0.771 | 0.054 | -0.342 | 0.225 | 1.29E-01 | -0.783 | 0.099 | -0.213 | 0.228 | 3.52E-01 | -0.660 | 0.235 | -0.137 | 0.237 | 5.63E-01 | -0.602 | 0.328 |
| | **batch 10** | 0.868 | 0.169 | 2.57E-07 | 0.538 | 1.198 | 0.682 | 0.191 | 3.66E-04 | 0.307 | 1.057 | 0.745 | 0.191 | 9.31E-05 | 0.371 | 1.119 | 0.712 | 0.198 | 3.15E-04 | 0.325 | 1.100 |
| | **intercept** | 3.020 | 0.173 | 2.44E-68 | 2.682 | 3.359 | 3.190 | 0.187 | 5.89E-65 | 2.822 | 3.557 | 3.106 | 0.215 | 1.75E-47 | 2.686 | 3.527 | 3.026 | 0.236 | 1.32E-37 | 2.564 | 3.489 |

| | | **Vascular infiltration by neutrophils** | | | | | | | | | | | | | | | |
|---|---|---|---|---|---|---|---|---|---|---|---|---|---|---|---|---|---|
| | | **Model 1 (n=404)** | | | | | **Model 2 (n=404)** | | | | | **Model 3 (n=359)** | | | | | |
| | | **Effect** | **SE** | ***P*** | **lci** | **uci** | **Effect** | **SE** | ***P*** | **lci** | **uci** | **Effect** | **SE** | ***P*** | **lci** | **uci** | |
| **fixed factors** | **atorvastatin and ezetimibe** | 0.088 | 0.083 | 2.87E-01 | -0.074 | 0.251 | 0.080 | 0.083 | 3.36E-01 | -0.083 | 0.242 | 0.052 | 0.089 | 5.60E-01 | -0.123 | 0.227 | *no observations* |
| | **time of day (in hours since 9AM)** | 0.023 | 0.021 | 2.76E-01 | -0.018 | 0.063 | 0.020 | 0.021 | 3.49E-01 | -0.022 | 0.062 | 0.003 | 0.023 | 8.83E-01 | -0.041 | 0.048 | |
| | **body length (in SD)** | - | - | - | - | - | 0.100 | 0.055 | 6.61E-02 | -0.007 | 0.207 | 0.111 | 0.057 | 5.06E-02 | 0.000 | 0.223 | |
| | **dorsal body surface area (in SD)** | - | - | - | - | - | -0.005 | 0.057 | 9.28E-01 | -0.117 | 0.106 | -0.015 | 0.062 | 8.13E-01 | -0.137 | 0.107 | |
| | **triglyceride levels (in SD)** | - | - | - | - | - | - | - | - | - | - | 0.020 | 0.051 | 7.02E-01 | -0.081 | 0.120 | |
| | **glucose levels (in SD)** | - | - | - | - | - | - | - | - | - | - | 0.034 | 0.040 | 3.92E-01 | -0.044 | 0.112 | |
| | **intercept** | -0.012 | 0.151 | 9.34E-01 | -0.308 | 0.283 | -0.006 | 0.144 | 9.68E-01 | -0.288 | 0.276 | 0.048 | 0.148 | 7.45E-01 | -0.242 | 0.338 | |
| **random factors** | ***variation by batch*** | *0.234* | *0.086* | - | *0.114* | *0.479* | *0.208* | *0.080* | - | *0.098* | *0.441* | *0.205* | *0.083* | - | *0.093* | *0.452* | |
| | ***residual*** | *0.831* | *0.029* | - | *0.775* | *0.891* | *0.827* | *0.029* | - | *0.772* | *0.887* | *0.812* | *0.031* | - | *0.754* | *0.874* | |

***continued*** **Supplementary Table 9**

| | | **Vascular co-localization of lipids with neutrophils** | | | | | | | | | | | | | | | |
|---|---|---|---|---|---|---|---|---|---|---|---|---|---|---|---|---|---|
| | | **Model 1 (n=393)** | | | | | **Model 2 (n=393)** | | | | | **Model 3 (n=348)** | | | | | |
| | | **Effect** | **SE** | ***P*** | **lci** | **uci** | **Effect** | **SE** | ***P*** | **lci** | **uci** | **Effect** | **SE** | ***P*** | **lci** | **uci** | |
| **negative binomial terms** | **atorvastatin and ezetimibe** | -1.113 | 0.395 | 4.87E-03 | -1.888 | -0.338 | -1.297 | 0.433 | 2.76E-03 | -2.146 | -0.448 | -1.518 | 0.578 | 8.67E-03 | -2.652 | -0.385 | *no observations* |
| | **time of day (in hours since 9AM)** | -0.094 | 0.095 | 3.25E-01 | -0.281 | 0.093 | -0.168 | 0.106 | 1.15E-01 | -0.376 | 0.041 | -0.120 | 0.120 | 3.18E-01 | -0.355 | 0.115 | |
| | **body length (in SD)** | - | - | - | - | - | -1.512 | 0.322 | 2.64E-06 | -2.143 | -0.881 | -1.501 | 0.328 | 4.61E-06 | -2.143 | -0.859 | |
| | **dorsal body surface area (in SD)** | - | - | - | - | - | 0.317 | 0.314 | 3.13E-01 | -0.299 | 0.934 | 0.207 | 0.366 | 5.72E-01 | -0.511 | 0.925 | |
| | **triglyceride levels (in SD)** | - | - | - | - | - | - | - | - | - | - | -0.043 | 0.232 | 8.52E-01 | -0.499 | 0.412 | |
| | **glucose levels (in SD)** | - | - | - | - | - | - | - | - | - | - | 0.193 | 0.241 | 4.23E-01 | -0.280 | 0.667 | |
| | **batch 1** | -1.376 | 1.000 | 1.69E-01 | -3.335 | 0.584 | -1.183 | 0.820 | 1.49E-01 | -2.790 | 0.424 | -0.936 | 0.866 | 2.80E-01 | -2.632 | 0.761 | |
| | **batch 2** | -2.393 | 0.517 | 3.60E-06 | -3.406 | -1.381 | -0.732 | 0.654 | 2.63E-01 | -2.015 | 0.551 | -0.237 | 0.674 | 7.25E-01 | -1.558 | 1.084 | |
| | **batch 3** | -1.164 | 0.563 | 3.87E-02 | -2.267 | -0.061 | -0.051 | 0.588 | 9.32E-01 | -1.204 | 1.103 | 0.349 | 0.796 | 6.61E-01 | -1.211 | 1.909 | |
| | **batch 4** | -0.811 | 0.479 | 9.01E-02 | -1.750 | 0.127 | 0.545 | 0.732 | 4.56E-01 | -0.889 | 1.979 | 0.841 | 0.767 | 2.73E-01 | -0.662 | 2.344 | |
| | **intercept** | 1.997 | 0.526 | 1.45E-04 | 0.967 | 3.027 | 1.290 | 0.612 | 3.52E-02 | 0.090 | 2.491 | 0.737 | 0.676 | 2.76E-01 | -0.588 | 2.062 | |

| | | **Vascular co-localization of macrophages with neutrophils** | | | | | | | | | | | | | | | |
|---|---|---|---|---|---|---|---|---|---|---|---|---|---|---|---|---|---|
| | | **Model 1 (n=394)** | | | | | **Model 2 (n=394)** | | | | | **Model 3 (n=349)** | | | | | |
| | | **Effect** | **SE** | ***P*** | **lci** | **uci** | **Effect** | **SE** | ***P*** | **lci** | **uci** | **Effect** | **SE** | ***P*** | **lci** | **uci** | |
| **fixed factors** | **atorvastatin and ezetimibe** | 0.139 | 0.092 | 1.29E-01 | -0.041 | 0.319 | 0.136 | 0.092 | 1.39E-01 | -0.044 | 0.316 | 0.159 | 0.100 | 1.10E-01 | -0.036 | 0.355 | *no observations* |
| | **time of day (in hours since 9AM)** | 0.033 | 0.023 | 1.45E-01 | -0.011 | 0.078 | 0.034 | 0.024 | 1.49E-01 | -0.012 | 0.080 | 0.026 | 0.025 | 2.97E-01 | -0.023 | 0.076 | |
| | **body length (in SD)** | - | - | - | - | - | 0.036 | 0.060 | 5.48E-01 | -0.082 | 0.154 | 0.062 | 0.063 | 3.28E-01 | -0.062 | 0.186 | |
| | **dorsal body surface area (in SD)** | - | - | - | - | - | -0.016 | 0.063 | 8.04E-01 | -0.140 | 0.109 | -0.092 | 0.070 | 1.89E-01 | -0.230 | 0.045 | |
| | **triglyceride levels (in SD)** | - | - | - | - | - | - | - | - | - | - | 0.077 | 0.057 | 1.80E-01 | -0.035 | 0.188 | |
| | **glucose levels (in SD)** | - | - | - | - | - | - | - | - | - | - | 0.085 | 0.044 | 5.55E-02 | -0.002 | 0.172 | |
| | **intercept** | -0.178 | 0.149 | 2.32E-01 | -0.471 | 0.114 | -0.182 | 0.149 | 2.23E-01 | -0.474 | 0.111 | -0.190 | 0.166 | 2.51E-01 | -0.515 | 0.135 | |
| **random factors** | ***variation by batch*** | *0.200* | *0.080* | - | *0.092* | *0.438* | *0.194* | *0.079* | - | *0.087* | *0.432* | *0.234* | *0.094* | - | *0.107* | *0.512* | |
| | ***residual*** | *0.906* | *0.032* | - | *0.845* | *0.972* | *0.906* | *0.032* | - | *0.844* | *0.972* | *0.894* | *0.034* | - | *0.830* | *0.964* | |

| | | **Endothelial thickness** | | | | | | | | | | | | | | | | | | | |
|---|---|---|---|---|---|---|---|---|---|---|---|---|---|---|---|---|---|---|---|---|---|
| | | **Model 1 (n=185)** | | | | | **Model 2 (n=185)** | | | | | **Model 3 (n=184)** | | | | | **Model 4 (n=162)** | | | | |
| | | **Effect** | **SE** | ***P*** | **lci** | **uci** | **Effect** | **SE** | ***P*** | **lci** | **uci** | **Effect** | **SE** | ***P*** | **lci** | **uci** | **Effect** | **SE** | ***P*** | **lci** | **uci** |
| **fixed factors** | **atorvastatin and ezetimibe** | -0.085 | 0.163 | 6.00E-01 | -0.404 | 0.234 | -0.083 | 0.164 | 6.14E-01 | -0.405 | 0.239 | -0.036 | 0.160 | 8.22E-01 | -0.350 | 0.277 | -0.036 | 0.174 | 8.36E-01 | -0.377 | 0.305 |
| | **time of day (in hours since 9AM)** | 0.031 | 0.037 | 4.05E-01 | -0.042 | 0.103 | 0.030 | 0.037 | 4.15E-01 | -0.042 | 0.102 | 0.014 | 0.036 | 6.87E-01 | -0.056 | 0.085 | -0.002 | 0.035 | 9.48E-01 | -0.071 | 0.066 |
| | **body length (in SD)** | - | - | - | - | - | -0.029 | 0.094 | 7.59E-01 | -0.214 | 0.156 | -0.113 | 0.094 | 2.30E-01 | -0.297 | 0.071 | -0.113 | 0.094 | 2.32E-01 | -0.297 | 0.072 |
| | **dorsal body surface area (in SD)** | - | - | - | - | - | 0.075 | 0.093 | 4.19E-01 | -0.107 | 0.257 | 0.020 | 0.092 | 8.31E-01 | -0.161 | 0.200 | 0.007 | 0.098 | 9.39E-01 | -0.185 | 0.200 |
| | **LDL cholesterol levels (in SD)** | - | - | - | - | - | - | - | - | - | - | - | - | - | - | - | 0.001 | 0.092 | 9.92E-01 | -0.180 | 0.182 |
| | **HDL cholesterol levels (in SD)** | - | - | - | - | - | - | - | - | - | - | - | - | - | - | - | -0.057 | 0.105 | 5.84E-01 | -0.263 | 0.148 |
| | **triglyceride levels (in SD)** | - | - | - | - | - | - | - | - | - | - | 0.390 | 0.107 | 2.71E-04 | 0.180 | 0.600 | 0.428 | 0.113 | 1.47E-04 | 0.207 | 0.648 |
| | **glucose levels (in SD)** | - | - | - | - | - | - | - | - | - | - | -0.341 | 0.180 | 5.76E-02 | -0.693 | 0.011 | -0.241 | 0.179 | 1.79E-01 | -0.593 | 0.111 |
| | **intercept** | 0.073 | 0.190 | 7.03E-01 | -0.301 | 0.446 | 0.015 | 0.197 | 9.39E-01 | -0.371 | 0.401 | -0.122 | 0.190 | 5.20E-01 | -0.495 | 0.250 | -0.127 | 0.190 | 5.04E-01 | -0.499 | 0.245 |
| **random factors** | ***variation by batch*** | *0.181* | *0.106* | - | *0.058* | *0.572* | *0.167* | *0.105* | - | *0.048* | *0.576* | *0.114* | *0.105* | - | *0.019* | *0.692* | *0.000* | *0.000* | - | *0.000* | *1650* |
| | ***residual*** | *0.976* | *0.051* | - | *0.881* | *1.081* | *0.974* | *0.051* | - | *0.879* | *1.080* | *0.938* | *0.049* | - | *0.846* | *1.040* | *0.890* | *0.049* | - | *0.798* | *0.992* |

Endothelial thickness is defined as surface area of the endothelium normalized for surface area of the circulating lipids. Associations were examined using negative binomial regresion for outcomes that showed a negative binomial distribution; and using hierarchical linear models on inverse normally transformed outcomes for outcomes that were (borderline) normally distributed (i.e. vascular accumulation of oxLDL; vascular infiltration by macrophages and neutrophils; endothelial thickness). Model 1: adjusted for time of day, transgenic background and batch; Model 2: additionally adjusted for body length and dorsal body surface area; Model 3: additionally adjusted for whole-body LDL cholesterol, HDL cholesterol, triglyceride and glucose levels. Dorsal body surface area was normalized for body length using residuals; whole-body LDL cholesterol, HDL cholesterol, triglyceride and glucose levels were normalized for protein level using residuals. Effects shown for atorvastatin and ezetimibe treatment are compared with untreated controls. Lci and uci are lower and upper boundaries of the 95% confidence interval.

**Supplementary Table 10 - The effect of treatment with atorvastatin and ezetimibe on suboptimal image or image quantification quality**

| | Vasculature not properly detected | | | | | | | | | | | | | | | | | | | |
|---|---|---|---|---|---|---|---|---|---|---|---|---|---|---|---|---|---|---|---|---|
| | Model 1 (n=927) | | | | | Model 2 (n=927) | | | | | Model 3 (n=876) | | | | | Model 4 (n=454) | | | | |
| | OR | SE | *P* | lci | uci | OR | SE | *P* | lci | uci | OR | SE | *P* | lci | uci | OR | SE | *P* | lci | uci |
| **atorvastatin and ezetimibe** | 0.910 | 0.200 | 6.85E-01 | 0.590 | 1.410 | 0.890 | 0.200 | 6.04E-01 | 0.580 | 1.380 | 0.910 | 0.210 | 6.95E-01 | 0.580 | 1.440 | 0.390 | 0.140 | 6.59E-03 | 0.200 | 0.770 |
| **time of day (in hours since 9AM)** | 1.040 | 0.050 | 4.72E-01 | 0.940 | 1.150 | 1.030 | 0.050 | 5.35E-01 | 0.930 | 1.140 | 1.050 | 0.050 | 3.38E-01 | 0.950 | 1.160 | 1.100 | 0.080 | 1.69E-01 | 0.960 | 1.270 |
| **body length (in SD)** | - | - | - | - | - | 0.880 | 0.100 | 2.85E-01 | 0.700 | 1.110 | 0.870 | 0.110 | 2.43E-01 | 0.680 | 1.100 | 0.840 | 0.170 | 3.89E-01 | 0.570 | 1.250 |
| **dorsal body surface area (in SD)** | - | - | - | - | - | 1.180 | 0.130 | 1.48E-01 | 0.940 | 1.460 | 1.420 | 0.180 | 4.62E-03 | 1.110 | 1.820 | 1.570 | 0.310 | 2.20E-02 | 1.070 | 2.320 |
| **LDL cholesterol levels (in SD)** | - | - | - | - | - | - | - | - | - | - | - | - | - | - | - | 0.880 | 0.150 | 4.53E-01 | 0.630 | 1.230 |
| **HDL cholesterol levels (in SD)** | - | - | - | - | - | - | - | - | - | - | - | - | - | - | - | 0.780 | 0.140 | 1.54E-01 | 0.550 | 1.100 |
| **triglyceride levels (in SD)** | - | - | - | - | - | - | - | - | - | - | 0.670 | 0.080 | 1.26E-03 | 0.520 | 0.850 | 0.600 | 0.150 | 3.77E-02 | 0.380 | 0.970 |
| **glucose levels (in SD)** | - | - | - | - | - | - | - | - | - | - | 0.850 | 0.100 | 1.70E-01 | 0.680 | 1.070 | 0.930 | 0.240 | 7.86E-01 | 0.560 | 1.550 |
| **intercept** | 0.100 | 0.030 | 8.39E-18 | 0.060 | 0.170 | 0.100 | 0.030 | 3.43E-17 | 0.060 | 0.170 | 0.090 | 0.030 | 9.41E-18 | 0.050 | 0.160 | 0.110 | 0.040 | 4.05E-08 | 0.050 | 0.250 |
| | **Many false positive lipid deposits** | | | | | | | | | | | | | | | | | | | |
| | Model 1 (n=853) | | | | | Model 2 (n=853) | | | | | Model 3 (n=804) | | | | | Model 4 (n=421) | | | | |
| | OR | SE | *P* | lci | uci | OR | SE | *P* | lci | uci | OR | SE | *P* | lci | uci | OR | SE | *P* | lci | uci |
| **atorvastatin and ezetimibe** | 0.440 | 0.230 | 1.23E-01 | 0.150 | 1.250 | 0.460 | 0.250 | 1.48E-01 | 0.160 | 1.320 | 0.780 | 0.440 | 6.62E-01 | 0.260 | 2.370 | 0.520 | 0.320 | 2.87E-01 | 0.150 | 1.740 |
| **time of day (in hours since 9AM)** | 1.010 | 0.110 | 9.45E-01 | 0.820 | 1.250 | 1.010 | 0.110 | 9.39E-01 | 0.820 | 1.250 | 0.960 | 0.100 | 7.20E-01 | 0.780 | 1.190 | 1.030 | 0.120 | 7.80E-01 | 0.830 | 1.280 |
| **body length (in SD)** | - | - | - | - | - | 1.430 | 0.380 | 1.69E-01 | 0.860 | 2.400 | 1.010 | 0.280 | 9.58E-01 | 0.590 | 1.750 | 1.000 | 0.310 | 9.93E-01 | 0.540 | 1.840 |
| **dorsal body surface area (in SD)** | - | - | - | - | - | 1.170 | 0.280 | 5.23E-01 | 0.730 | 1.870 | 0.950 | 0.270 | 8.44E-01 | 0.550 | 1.640 | 1.350 | 0.470 | 3.91E-01 | 0.680 | 2.680 |
| **LDL cholesterol levels (in SD)** | - | - | - | - | - | - | - | - | - | - | - | - | - | - | - | 0.650 | 0.180 | 1.09E-01 | 0.380 | 1.100 |
| **HDL cholesterol levels (in SD)** | - | - | - | - | - | - | - | - | - | - | - | - | - | - | - | 0.950 | 0.250 | 8.48E-01 | 0.570 | 1.590 |
| **triglyceride levels (in SD)** | - | - | - | - | - | - | - | - | - | - | 2.500 | 0.820 | 5.08E-03 | 1.320 | 4.750 | 2.640 | 1.050 | 1.49E-02 | 1.210 | 5.770 |
| **glucose levels (in SD)** | - | - | - | - | - | - | - | - | - | - | 0.370 | 0.110 | 8.64E-04 | 0.210 | 0.670 | 0.400 | 0.150 | 1.70E-02 | 0.190 | 0.850 |
| **intercept** | 0.030 | 0.020 | 3.80E-11 | 0.010 | 0.080 | 0.030 | 0.010 | 2.63E-11 | 0.010 | 0.070 | 0.010 | 0.010 | 2.61E-12 | 0.000 | 0.050 | 0.020 | 0.010 | 4.01E-09 | 0.010 | 0.080 |
| | **Larva moved during imaging** | | | | | | | | | | | | | | | | | | | |
| | Model 1 (n=873) | | | | | Model 2 (n=873) | | | | | Model 3 (n=824) | | | | | Model 4 (n=440) | | | | |
| | OR | SE | *P* | lci | uci | OR | SE | *P* | lci | uci | OR | SE | *P* | lci | uci | OR | SE | *P* | lci | uci |
| **atorvastatin and ezetimibe** | 3.510 | 1.370 | 1.23E-03 | 1.640 | 7.530 | 3.410 | 1.370 | 2.17E-03 | 1.560 | 7.480 | 3.380 | 1.360 | 2.43E-03 | 1.540 | 7.430 | 2.060 | 0.950 | 1.19E-01 | 0.830 | 5.080 |
| **time of day (in hours since 9AM)** | 1.200 | 0.100 | 3.36E-02 | 1.010 | 1.410 | 1.210 | 0.110 | 4.02E-02 | 1.010 | 1.450 | 1.220 | 0.110 | 3.11E-02 | 1.020 | 1.470 | 1.250 | 0.130 | 3.24E-02 | 1.020 | 1.540 |
| **body length (in SD)** | - | - | - | - | - | 0.390 | 0.070 | 5.54E-07 | 0.270 | 0.560 | 0.410 | 0.080 | 3.24E-06 | 0.280 | 0.590 | 0.700 | 0.200 | 2.04E-01 | 0.400 | 1.220 |
| **dorsal body surface area (in SD)** | - | - | - | - | - | 1.430 | 0.240 | 3.66E-02 | 1.020 | 2.000 | 1.460 | 0.270 | 4.07E-02 | 1.020 | 2.100 | 1.580 | 0.390 | 6.33E-02 | 0.970 | 2.560 |
| **LDL cholesterol levels (in SD)** | - | - | - | - | - | - | - | - | - | - | - | - | - | - | - | 0.820 | 0.200 | 4.10E-01 | 0.500 | 1.320 |
| **HDL cholesterol levels (in SD)** | - | - | - | - | - | - | - | - | - | - | - | - | - | - | - | 0.600 | 0.160 | 5.92E-02 | 0.350 | 1.020 |
| **triglyceride levels (in SD)** | - | - | - | - | - | - | - | - | - | - | 0.920 | 0.200 | 7.07E-01 | 0.610 | 1.400 | 0.340 | 0.130 | 3.91E-03 | 0.160 | 0.700 |
| **glucose levels (in SD)** | - | - | - | - | - | - | - | - | - | - | 1.030 | 0.230 | 8.80E-01 | 0.670 | 1.590 | 8.660 | 4.850 | 1.16E-04 | 2.890 | 25.960 |
| **intercept** | 0.010 | 0.000 | 5.15E-18 | 0.000 | 0.030 | 0.010 | 0.000 | 2.00E-17 | 0.000 | 0.020 | 0.010 | 0.000 | 6.94E-17 | 0.000 | 0.020 | 0.010 | 0.010 | 1.60E-09 | 0.000 | 0.060 |
| | **Many false positive oxLDL deposits** | | | | | | | | | | | | | | | | | | | |
| | Model 1 (n=236) | | | | | Model 2 (n=236) | | | | | Model 3 (n=233) | | | | | Model 4 (n=229) | | | | |
| | OR | SE | *P* | lci | uci | OR | SE | *P* | lci | uci | OR | SE | *P* | lci | uci | OR | SE | *P* | lci | uci |
| **atorvastatin and ezetimibe** | 0.480 | 0.130 | 7.66E-03 | 0.280 | 0.820 | 0.520 | 0.150 | 1.96E-02 | 0.300 | 0.900 | 0.510 | 0.150 | 2.09E-02 | 0.290 | 0.900 | 0.520 | 0.160 | 3.14E-02 | 0.290 | 0.940 |
| **time of day (in hours since 9AM)** | 0.970 | 0.060 | 5.51E-01 | 0.860 | 1.080 | 0.960 | 0.060 | 5.12E-01 | 0.850 | 1.080 | 0.960 | 0.060 | 5.09E-01 | 0.850 | 1.080 | 0.960 | 0.060 | 5.32E-01 | 0.850 | 1.090 |
| **body length (in SD)** | - | - | - | - | - | 1.570 | 0.250 | 4.91E-03 | 1.150 | 2.150 | 1.730 | 0.360 | 7.91E-03 | 1.160 | 2.600 | 1.820 | 0.390 | 4.50E-03 | 1.200 | 2.760 |
| **dorsal body surface area (in SD)** | - | - | - | - | - | 1.370 | 0.230 | 5.87E-02 | 0.990 | 1.890 | 1.540 | 0.330 | 4.19E-02 | 1.020 | 2.340 | 1.640 | 0.350 | 2.27E-02 | 1.070 | 2.500 |
| **LDL cholesterol levels (in SD)** | - | - | - | - | - | - | - | - | - | - | - | - | - | - | - | 1.070 | 0.170 | 6.76E-01 | 0.780 | 1.470 |
| **HDL cholesterol levels (in SD)** | - | - | - | - | - | - | - | - | - | - | - | - | - | - | - | 1.440 | 0.260 | 3.96E-02 | 1.020 | 2.040 |
| **triglyceride levels (in SD)** | - | - | - | - | - | - | - | - | - | - | 0.770 | 0.170 | 2.31E-01 | 0.490 | 1.190 | 0.810 | 0.190 | 3.67E-01 | 0.500 | 1.290 |
| **glucose levels (in SD)** | - | - | - | - | - | - | - | - | - | - | 1.050 | 0.270 | 8.43E-01 | 0.640 | 1.730 | 0.890 | 0.260 | 6.92E-01 | 0.510 | 1.570 |
| **intercept** | 2.130 | 0.680 | 1.88E-02 | 1.130 | 4.000 | 2.250 | 0.830 | 2.74E-02 | 1.090 | 4.620 | 2.320 | 1.070 | 6.90E-02 | 0.940 | 5.730 | 1.750 | 0.880 | 2.60E-01 | 0.660 | 4.670 |

Associations are shown for criteria that resulted in the exclusion of at least 10 larvae. Vasculature not properly detected typically resulted from weak staining, possibly due to low levels of circulating lipids; Many false positives: >20% of true negative objects were falsely detected by the qualtification pipeline; Many false negatives: <20% of true positive objects were detected by the qualtification pipeline. Associations were examined using logistic regression models. Model 1: adjusted for time of day; Model 2: additionally adjusted for body length and dorsal body surface area; Model 3: additionally adjusted for whole-body triglyceride and glucose levels; Model 4: additionally adjusted for whole-body LDL and HDL cholesterol levels. Dorsal body surface area was normalized for body length; whole-body LDL cholesterol, HDL cholesterol, triglyceride and glucose levels were normalized for protein level. Adjusting for transgenic background and batch would have excluded approximately half the larvae. Effects shown for atorvastatin and ezetimibe treatment are compared with untreated controls. Lci and uci are lower and upper boundaries of the 95% confidence interval.

**Supplementary Table 11 - Orthologues of proof-of-concept genes for dyslipidemia, atherosclerosis and coronary artery disease**

| Human gene | ENSG | Zebrafish orthologue | ENSDARG | Target %identity | Query %identity | Main human protein | Top hit BLAST | %identity (protein) | Conserved genes in locus |
|---|---|---|---|---|---|---|---|---|---|
| *APOE* | ENSG00000130203 | *apoea* | ENSDARG00000102004 | 25.65 | 21.77 | ENSP00000252486 | ENSDARP00000137865 | 27.78 | *TOMM40* |
| | | *apoeb* | ENSDARG00000040295 | 28.11 | 24.92 | | ENSDARP00000119141 | 32.04 | *BCAM, NECTIN2* |
| *APOB* | ENSG00000084674 | *apoba* | ENSDARG00000042780 | 33.70 | 32.63 | ENSP00000233242 | ENSDARP00000062792 | 34.51 | *C2orf43, GDF7, ITSN2, PFN4* |
| | | *apobb.1* | ENSDARG00000022767 | 29.68 | 24.26 | | ENSDARP00000119179 | 30.33 | NA |
| | | *apobb.2* | ENSDARG00000075016 | 29.64 | 16.46 | | ENSDARP00000144532 | 37.37 | NA |
| *LDLR* | ENSG00000130164 | *ldlra* | ENSDARG00000029476 | 52.80 | 55.93 | ENSP00000252444 | ENSDARP00000115492 | 58.69 | *SMARCA4, KRI1, SPC24* |
| | | *ldlrb* | ENSDARG00000026759 | 54.06 | 49.53 | | ENSDARP00000141207 | 58.57 | *SMARCA4, AP1M2, CDKN2D* |

Target %identity: percentage of the orthologous sequence matching the human sequence; Query %identity: percentage of the human sequence matching the sequence of the orthologue; Main human protein: Ensembl protein ID for the main transcript; %identity protein: percentage of the aligned query (input sequence, i.e. main human protein) which is identical to the subject (hit) sequence; conserved genes in locus: neighbouring genes conserved across danio rerio and homo sapiens locus according to Genomicus.

**Supplementary Table 12 - Identification of moderate-to-highly active CRISPR-Cas9 guide RNAs for proof-of-concept genes**

| Human gene | Zebrafish orthologue | CRISPR gRNA target sequence | Genomic location (danRer11/GRCz11) | | Exon | Strand | GC (%) | Self-complementarity | Off-targets | | | | Predicted efficiency | CRISPRscan score | Canonical (yes/no) | Target activity (NA[a], no[b], low[c], moderate[d], high[e] or very high[f]) | Forward primer | Reverse primer | Product size |
|---|---|---|---|---|---|---|---|---|---|---|---|---|---|---|---|---|---|---|---|
| | | | Chr | Pos | | | | | 0 | 1 | 2 | 3 | | | | | | | |
| ***APOE*** | ***apoea*** | **GGCTCTCTCCTGCGCGTAAG** | **19** | **10,856,063** | **3 of 4** | **-** | **65** | **0** | **0** | **0** | **0** | **0** | **0.42** | **54** | **yes** | **high** | **AGCACACTGATCTCTGACAGC** | **GATCCTTCGCCTCCTCCATG** | **160** |
| | | GGATGAGCCAAGAAGCCGCT | 19 | 10,855,768 | 2 of 4 | + | 60 | 2 | 0 | 0 | 0 | 0 | 0.52 | 37 | yes | moderate | TCCGTTTTGACTTTGCAGGC | GAGCTGAGTGGCCTTGATGT | 155 |
| | ***apoeb*** | GgGGGGCATCAGCCTGGAAC | 16 | 23,961,545 | 2 of 4 | - | 65 | 1 | 0 | 0 | 0 | 1 | 0.61 | 59 | no | very high | TGCCTGACTTGCTAATTGTGAT | CCGTCAGTTTGTGTGTTGAGTT | 178 |
| | | GgTGACGTGAAGAACCGTGT | 16 | 23,962,014 | 3 of 4 | + | 50 | 0 | 0 | 0 | 0 | 2 | 0.62 | 68 | no | very high | AGTGAAAATCTCCAAACCCAGA | TGAAGGAGCATCCCAACTTACT | 269 |
| | | **GAGGGGCATCAGCCTGGAAC** | **16** | **23,961,545** | **2 of 4** | **-** | **65** | **1** | **0** | **0** | **0** | **1** | **0.61** | **59** | **no** | **high** | **CTGTAAATTGCCTGACTTGCTAA** | **GCCCTTGATGTTTTGCACCA** | **208** |
| | | GgTGACGTGAAGAACCGTGT | 16 | 23,962,014 | 3 of 4 | + | 50 | 0 | 0 | 0 | 0 | 2 | 0.62 | 68 | no | high | CTGCTGGTCAGCTCAGTAAAGA | TGAAGGAGCATCCCAACTTACT | 226 |
| | | GgGGATCTTCTGGGAGTAGG | 16 | 23,962,897 | 4 of 4 | - | 60 | 0 | 0 | 0 | 0 | 0 | 0.59 | 85 | no | moderate | AGGAGAAGCTGGAGGAGACAG | CTCTTAAGCCTGAGTGGGAAGA | 170 |
| | | GgCATACATGAGCCAGGCCC | 16 | 23,962,700 | 4 of 4 | + | 65 | 1 | 0 | 0 | 0 | 0 | 0.58 | 59 | no | moderate | GCAACCTACATGAGTGAGATGC | GTAGGTTCTCGGCTGTCTCCT | 220 |
| | | GAACTCAACACACAAACTGA | 16 | 23,961,615 | 2 of 4 | + | 40 | 0 | 0 | 0 | 0 | 8 | 0.70 | 25 | no | no | CTCCAAACCCAGATGACCCC | CAGTTTGCGTGTGTAGGTGC | 201 |
| ***APOB*** | ***apoba*** | **gGGGAGGGCTCTATCTTAGG** | **17** | **30,717,989** | **24 of 29** | **+** | **55** | **0** | **0** | **0** | **0** | **0** | **0.69** | **100** | **no** | **high** | **CGCACTTTGGAATTCTCCTTAC** | **TCAATTTTGTATGACAGGGGTG** | **185** |
| | | gGATGAGGCAGACAGAGAGG | 17 | 30,708,635 | 9 of 29 | + | 60 | 0 | 0 | 0 | 0 | 12 | 0.74 | 61 | no | high | CTCACATGGCAGACACTCTTTC | GGGCCATACTCAGCATATCTCT | 172 |
| | | GGACACATTCTGTGGTGCGG | 17 | 30,705,707 | 4 of 29 | - | 60 | 1 | 0 | 0 | 0 | 5 | 0.64 | 48 | yes | no | ATGTCACAACCTCTGCAGCTAA | ACTTCTCCATAGCTGCCTGAAA | 944 |
| | | GGAAGCACTGAGGTTGCTG | 17 | 30,704,770 | 2 of 29 | + | 55 | 1 | 0 | 0 | 0 | 7 | 0.60 | 35 | no | no | AGGCCGCCTATTAATTCAGTTT | ATGTCACAACCTCTGCAGCTAA | 173 |
| | | GgAGTATGGTATCTCCTCAG | 17 | 30,708,080 | 8 of 29 | + | 50 | 0 | 0 | 0 | 0 | 0 | 0.59 | 74 | no | no | ATTTTTGTAAAGGGTGGGAACA | AAAAAGCAACAACCCATTTCAT | 224 |
| | | GgTGCTTTTTGCACAGGCAG | 17 | 30,709,180 | 11 of 29 | - | 57 | 1 | 0 | 0 | 0 | 0 | 0.71 | 44 | no | no | TCATTGGTGTCATGGGAAAATA | CACTAGAATGCAGAAAATCCCC | 260 |
| | ***apobb.1*** | **GgAGCTGACAAAGTACCAAG** | **20** | **31,273,917** | **13 of 27** | **+** | **50** | **0** | **0** | **0** | **0** | **1** | **0.75** | **54** | **no** | **high** | **CCCTGATTGGTATTGATGGATT** | **GGCCTAGAGTGAGAGGAGAACA** | **228** |
| | | gGGAGTTGAGTTTGTGACGG | 20 | 31,274,651 | 16 of 27 | + | 50 | 0 | 0 | 0 | 0 | 0 | 0.61 | 81 | no | moderate | AACTTGCTCGTGACATGGTATG | TTGAGACCACTCTCGTGGTAGA | 227 |
| | | GAGCCAGTTCAGTGGGCTG | 20 | 31,273,334 | 11 of 27 | - | 60 | 2 | 0 | 0 | 0 | 22 | 0.47 | 38 | no | moderate | GCAGAGAGGTGCTAATGAAGGT | TGTATACTCACTCCCCGGTCTC | 222 |
| | | GgGTTCATCCAGATTTGCAG | 20 | 31,277,737 | 24 of 27 | - | 45 | 0 | 0 | 0 | 0 | 8 | 0.73 | 47 | no | moderate | TTGACACTTTGTTTGGAAATCG | CCATTGAATTTGTTCTGCAGTG | 217 |
| | | GAAATCAAGCAGCAAGATGG | 20 | 31,277,192 | 24 of 27 | + | 45 | 0 | 0 | 0 | 1 | 5 | 0.70 | - | no | moderate | TGTTTAGGATCAACCTTCCTGG | ATTCAAAGGACCAGCCTTGATA | 279 |
| | | GGCTCTATTTTCTCCATTTG | 20 | 31,272,363 | 8 of 27 | - | 40 | 0 | 0 | 0 | 0 | 8 | 0.32 | 21 | yes | no | CAGTCCATCCAATTCAAGACAA | TACAGAGGAACGGTCAAAGGTT | 275 |
| | ***apobb.2*** | **gGATTAGCTGAGCAAGAGGA** | **20** | **53,444,329** | **5 of 22** | **-** | **45** | **1** | **0** | **0** | **0** | **7** | **0.61** | **49** | **no** | **moderate** | **GTGGACCCAGCATAAGACATTT** | **AACAAAATAGCAGGGATGCACT** | **226** |
| | | gGGCAGCCTGTTGGACTGCA | 20 | 53,445,660 | 9 of 22 | - | 60 | 5 | 0 | 0 | 0 | 0 | 0.40 | 80 | no | moderate | AACATGGTGGCTGCACTAGG | AGCACTTCTCTTCCCTGTAGG | 227 |
| | | gGGAGTCACAAGTGAGATCC | 20 | 53,445,352 | 8 of 22 | + | 55 | 1 | 0 | 0 | 0 | 1 | 0.50 | 66 | no | low | TGCCAAGGTATTGGGTTAGATT | AAAAACATGTTCCTGGTCACCT | 216 |
| | | gGACAAGTTCAGACCCATCG | 20 | 53,443,193 | 4 of 22 | + | 50 | 1 | 0 | 0 | 0 | 2 | 0.59 | 38 | no | no | CCCAGCATCTATGGTCTGTGTA | TTTAATGGAATTGCACCAGTTG | 234 |
| | | GATCCCCCTCTTAATGTTG | 20 | 53,442,015 | 3 of 22 | - | 45 | 0 | 0 | 0 | 0 | 2 | 0.53 | 34 | no | no | AAATCCTCTGAAATTCACCGT | TGGTTTGAAGTGAAAGGACAAA | 238 |
| | | gGTGAAGGAAATGTTGGCGA | 20 | 53,445,098 | 7 of 22 | + | 45 | 0 | 0 | 0 | 0 | 4 | 0.77 | 62 | no | NA | GAGGTGGATGCTGCTGTATATG | AATCTAACCCAATACCTTGGCA | 201 |
| ***LDLR*** | ***ldlra*** | **GATTCACGGCAGTATCAGTG** | **3** | **19,304,761** | **2 of 18** | **+** | **50** | **1** | **0** | **0** | **0** | **25** | **0.50** | **50** | **no** | **high** | **TAGCGCATATATCACAACGGAC** | **CCATCACCACAGTCATCAGTTT** | **228** |
| | | GGAAGTGGGGAATGCATACA | 3 | 19,308,392 | 4 of 18 | + | 50 | 0 | 0 | 0 | 0 | 3 | 0.46 | 75 | yes | moderate | GACAATTCAGATGAGTTGCCTG | ATACCATCCAGTGATAATCGGC | 274 |
| | ***ldlrb*** | **GGAGCGGATTCTGCGAGCGG** | **6** | **102,541** | **1 of 5** | **-** | **70** | **0** | **0** | **0** | **0** | **0** | **0.79** | **66** | **yes** | **high** | **CCCTGGCCTCAACACTACAG** | **ACCCAGAAGAGCAGCAGAAC** | **215** |
| | | gGCGCTGAGGAGTTCCGCTG | 6 | 103,574 | 3 of 5 | + | 65 | 1 | 0 | 0 | 0 | 1 | 0.67 | 80 | no | high | CGAGCAGAACTGCGGTAAAT | TGCACTGGAATGCTGTGG | 249 |
| | | gGGGCACACACTCTCCGCTG | 6 | 103,852 | 3 of 5 | - | 70 | 2 | 0 | 0 | 0 | 2 | 0.68 | 78 | no | no | GTGTCCCACACACACACAC | gtACTCACTGCAGTTGTCCTCG | 271 |
| | | GgGTCACGTTAGAGCTCTGA | 6 | 102,589 | 1 of 5 | - | 55 | 0 | 0 | 0 | 0 | 0 | 0.52 | 61 | no | no | TTCTGCTCAGAGAGGGAGAATC | GATCAGTGAAACTCACCGGTTC | 182 |
| | | GgGGATTCTGCGAGCGGTGG | 6 | 102,538 | 1 of 5 | - | 70 | 0 | 0 | 0 | 1 | 0 | 0.56 | 94 | no | no | TAACATCACACCACTGCTGGAG | GATCAGTGAAACTCACCGGTTC | 270 |
| | | GGGAAGTCTGACTGTGAGAA | 6 | 103,441 | 2 of 5 | + | 50 | 2 | 0 | 0 | 0 | 5 | 0.51 | 63 | yes | no | TGCTAATGACTCCTCTGCGT | CACACACACAGCAGAGTCAC | 216 |
| | | GgGTCGGGTCAGTTCAGCTG | 6 | 103,375 | 2 of 5 | + | 60 | 0 | 0 | 0 | 0 | 2 | 0.66 | 70 | no | NA | AGACAGGGTGAAacacacacac | agcagagtcacacacCTGAGAG | 249 |
| | | GgGGAACTCATCCGGCCGGC | 6 | 104,079 | 4 of 5 | - | 70 | 1 | 0 | 0 | 0 | 0 | 0.49 | - | no | NA | AGTGAGCAAGCGGACACTAAAt | TGAGTGGGCTTGTGTGTTTAAG | 258 |
| | | GGCACTGCATCTGCTGCTCC | 6 | 103,831 | 3 of 5 | - | 65 | 1 | 0 | 0 | 0 | 8 | 0.47 | 47 | yes | NA | GGGTCAGACGAGTGGCTG | GTACTCACTGCAGTTGTCCTCG | 162 |
| | | GgGTTCCTCTGTGCAGACGG | 6 | 104,093 | 4 of 5 | + | 60 | 1 | 0 | 0 | 1 | 6 | 0.75 | - | no | NA | AGTGAGCAAGCGGACACTAAAt | TGAGTGGGCTTGTGTGTTTAAG | 258 |

CRISPR gRNA target sequences were preferably selected based on location (i.e. in an early exon that affects all transcripts), complementarity (i.e. no complementarity), and free from predicted off targets. Target activity was examined by micro-injections in eight fertilized eggs in multiplex, followed by fragment length PCR analysis at 3 days post-fertilization. Results from target efficiency testing are shown, where NA: Not available due to failed capilary electrophoresis while estimating the length of the targeted region of an exon; No: 8 of 8 larvae test-injected with the gRNA only showed wildtype sequences; Low: 8 of 8 larvae showed wildtype sequence and fewer than 4 of 8 also contained indel sequence; Moderate: 8 of 8 larvae showed wildtype sequence and >4 of 8 also contained indel sequence; High: 8 of 8 larvae showed wildtype as well as indel sequence; Very high: Fewer than 4 of 8 larvae showed wildtype sequence and all larvae showed indel sequence. Target sequences highlighted in bold were selected and used to generate multiplexed mutant zebrafish. Of note: after completion of the study a new version of the zebrafish genome was released. Data shown for gRNA target sequences are from the new version of the genome (built GRCz11).

**Supplementary Table 13 - Unique CRISPR-Cas9-induced mutations for orthologues of proof-of-concept genes**

| Zebrafish orthologue | Sequence | Annotation | Number of alleles | mean ± SD number of reads |
|---|---|---|---|---|
| *apoea* | **ATGATGGAGCTGCAAATGTACAGAGATGATCTGCACTCCAAACTGGCCCCTTACGCGCAGGAGAGAGCCCAGAAGTTCAACGAAGATCTGCAGTTGTTGGTCACCAAGCTCCGCACACA** | **119M** | **430** | **1079 ± 766** |
| | ATGATGGAGCTGCAAATGTACAGAGATGATCTGCACTCCAAACGCGCAGGAGAGAGCCCAGAAGTTCAACGAAGATCTGCAGTTGTTGGTCACCAAGCTCCGCACACA | 41M11D67M | 91 | 861 ± 515 |
| | ATGATGGAGCTGCAAATGTACAGAGATGATCTGCACTCCAAACTGGCCCCTGGAGAGAGCCCAGAAGTTCAACGAAGATCTGCAGTTGTTGGTCACCAAGCTCCGCACACA | 51M8D60M | 67 | 873 ± 498 |
| | ATGATGGAGCTGCAAATGTACAGAGATGATCTGCACTCCAAACTGGCCCGCGCAGGAGAGAGCCCAGAAGTTCAACGAAGATCTGCAGTTGTTGGTCACCAAGCTCCGCACACA | 48M5D66M | 50 | 726 ± 397 |
| | ATGATGGACAGCATGATGGAGCTGCAAATGTACAGAGATGATCTGCACTCCAAACGCGCAGGAGAGAGCCCAGAAGTTCAACGAAGATCTGCAGTTGTTGGTCACCAAGCTCCGCACACA | 2M4S47M11D67M | 30 | 735 ± 344 |
| | ATGATGGAGCTGCAAATGTACAGAGATGATCTGCACTCCAAACTGGCCCCTCCAAACGCGCAGGAGAGAGCCCAGAAGTTCAACGAAGATCTGCAGTTGTTGGTCACCAAGCTCCGCACACA | 51M1S3I67M | 29 | 698 ± 293 |
| | ATGATGGAGCTGCAAATGTACAGAGATGATCTGCACTCCAAACTGGCCCCAGAGGAGGCAGGAGAGAGCCCAGAAGTTCAACGAAGATCTGCAGTTGTTGGTCACCAAGCTCCGCACACA | 50M2S1M1S1M1S1I63M | 15 | 782 ± 366 |
| | ATGATGGAGCTGCAAATGTACAGAGATGATCTGCACTCCAAACTGGCCCCCCTCTCTCGCGCAGGAGAGAGCCCAGAAGTTCAACGAAGATCTGCAGTTGTTGGTCACCAAGCTCCGCACACA | 49M4I2M2S66M | 14 | 636 ± 204 |
| | ATGATGGAGCTGCAAATGTACAGAGATGATCTGCACTCCAAACTGGCCCAGGAGAGAGCCCAGAAGTTCAACGAAGATCTGCAGTTGTTGGTCACCAAGCTCCGCACACA | 48M9D62M | 12 | 577 ± 355 |
| | ATGATGGAGCTGCAAATGTACAGAGATGATCTGCACTCCAAACTGGCCCCTGGCCCGCAGGAGAGAGCCCAGAAGTTCAACGAAGATCTGCAGTTGTTGGTCACCAAGCTCCGCACACA | 51M2S1M1S64M | 7 | 824 ± 601 |
| | ATGATGGAGCTGCAAATGTACAGAGATGATCTGCACTCCAAACTGGCAGGAGAGAGCCCAGAAGTTCAACGAAGATCTGCAGTTGTTGGTCACCAAGCTCCGCACACA | 45M11D63M | 7 | 695 ± 234 |
| | ATGATGGAGCTGCAAATGTACAGAGATGATCTGCACTCCAAACTGGCCCCTTGGAGGAGAGAGCCCAGAAGTTCAACGAAGATCTGCAGTTGTTGGTCACCAAGCTCCGCACACA | 52M4D1M1S61M | 5 | 862 ± 430 |
| | ATGATGGAGCTGCAAATGTACAGAGATGATCTGCACAGGAGAGAGCCCAGAAGTTCAACGAAGATCTGCAGTTGTTGGTCACCAAGCTCCGCACACA | 35M22D62M | 2 | 828 ± 729 |
| | ATGATGGAGCTGCAAATGTACAGAGATGATCTGCACTCCAAACTGGCCCCTTGATACGCGCAGGAGAGAGCCCAGAAGTTCAACGAAGATCTGCAGTTGTTGGTCACCAAGCTCCGCACACA | 51M3I68M | 2 | 664 ± 109 |
| | ATGATGGAGCTGCAAATGTACAGAGATGATCTGCACTCCAAACTGGCCGCGCAGGAGAGAGCCCAGAAGTTCAACGAAGATCTGCAGTTGTTGGTCACCAAGCTCCGCACACA | 47M6D66M | 1 | 2374 |
| | ATGATGGAGCTGCAAATGTACAGAGATGATCTGCACTCCAAACTGGCCCCACGCGCAGGAGAGAGCCCAGAAGTTCAACGAAGATCTGCAGTTGTTGGTCACCAAGCTCCGCACACA | 50M2D67M | 1 | 552 |
| | ATGATGGAGCTGCAAATGTACAGAGATGATCTGCACTCCAAACTGGCGCAGGAGAGAGCCCAGAAGTTCAACGAAGATCTGCAGTTGTTGGTCACCAAGCTCCGCACACA | 45M9D65M | 1 | 21 |
| *apoeb* | TTGTGATTAACTTAGTTTGGTATAAAGTTCACAGTCTCTTCTCACCTCTTGTAGGCTGCCAGGCTCGTAGCCTGTTGCCCCTCGTATGCCCCTCAGCCCAGATGGGAGGAGATGGTGGACCGTTTCTGGCAGTATGTGTCTGAACTCAACACACAAACTGACGGCA | 31M1S44M1I2M3S1M2S81M | 157 | 664 ± 416 |
| | TTGTGATTAACTTAGTTTGGTATAAAGTTCACAGTCTCTTCTCACCTCTTGTAGGCTGCCAGGCTCGTAGCCTGTCAAGTGACAGGCTGATGCCCCTCAGCCCAGATGGGAGGAGATGGTGGACCGTTTCTGGCAGTATGTGTCTGAACTCAACACACAAACTGACGGCA | 31M1S43M1S5I1S88M | 116 | 683 ± 500 |
| | TTGTGATTAACTTAGTTTGGTATAAAGTTCACAGTCTCTTCTCACCTCTTGTAGGCTGCCAGGCTCGTAGCTGATGGCTGATGCCCCTCAGCCCAGATGGGAGGAGATGGTGGACCGTTTCTGGCAGTATGTGTCTGAACTCAACACACAAACTGACGGCA | 31M1S39M3S1M4D86M | 100 | 708 ± 413 |
| | TTGTGATTAACTTAGTTTGGTATAAAGTTCACAGTCTCTTCTCACCTCTTGTAGGCTGCCAGGCTCGTAGCCTGTTCCAGGCTGATGCCCCTCAGCCCAGATGGGAGGAGATGGTGGACCGTTTCTGGCAGTATGTGTCTGAACTCAACACACAAACTGACGGCA | 31M1S133M | 91 | 677 ± 465 |
| | TTGTGATTAACTTAGTTTGGTATAAAGTTCACAGTCTCTTCTCACCTCTTGTAGGCTGCCAGGCTCGTAGCCTGATGCCCCTCAGCCCAGATGGGAGGAGATGGTGGACCGTTTCTGGCAGTATGTGTCTGAACTCAACACACAAACTGACGGCA | 31M1S39M10D84M | 75 | 619 ± 399 |
| | TTGTGATTAACTTAGTTTGGTATAAAGTTCACAGTCTCTTCTCACCTCTTGTAGGCTGCCAGGCTCGTAGCCAGGCTGATGCCCCTCAGCCCAGATGGGAGGAGATGGTGGACCGTTTCTGGCAGTATGTGTCTGAACTCAACACACAAACTGACGGCA | 31M1S38M6D89M | 46 | 455 ± 273 |
| | TTGTGATTAACTTAGTTTGGTATAAAGTTCAGAGTCTCTTCTCACCTCTTGTAGGCTGCCAGGCTCGTAGCCTGTTATATATATATAACAGGCTGATGCCCCTCAGCCGAGATGGGAGGAGATGGTGAGATGGTGGACCGTTTCTGGCAGTATGTGTCTGAACTCAACACACAAACTGATGGCA | 76M9I1S20M1S9M8I53M1S4M | 31 | 47 ± 26 |
| | TTGTGATTAACTTAGTTTGGTATAAAGTTCACAGTCTCTTCTCACCTCTTGTAGGCTGCCAGGCTCGTAGCCTGTTATATATATATAACAGGCTGATGCCCCTCAGCCGAGATGGGAGGAGATGGTGAGATGGTGGACCGTTTCTGGCAGTATGTGTCTGAACTCAACACACAAACTGACGGCA | 31M1S44M9I1S20M1S9M8I58M | 29 | 32 ± 19 |
| | TTGTGATTAACTTAGTTTGGTATAAAGTTCACAGTCTCTTCTCACCTCTTGTAGGCTGCCAGGCTCGTAGCCTGTTGCCCCTCAGCCCAGATGGGAGGAGATGGTGGACCGTTTCTGGCAGTATGTGTCTGAACTCAACACACAAACTGACGGCA | 31M1S43M10D80M | 22 | 475 ± 380 |
| | TTGTGATTAACTTAGTTTGGTATAAAGTTCACAGTCTCTTCTCACCTCTTGTAGGCTGCCAGGCTCGTAGCCTAGGCTAATGCCCCTCAGCCAAGATGGGAGGAGATGGTGGACCGTTTCTGGCAGTATGTGTCTGAACTCAACACACAAACTGACGGCA | 31M1S41M5D5M1S13M1S67M | 16 | 357 ± 294 |
| | TTGTGATTAACTTAGTTTGGTATAAAGTTCACAGTCTCTTCTCACCTCTTGTAGGCTGCCAGGCTCGTAGCCTGCCCCTCAGCCCAGATGGGAGGAGATGGTGGACCGTTTCTGGCAGTATGTGTCTGAACTCAACACACAAACTGACGGCA | 31M1S40M13D80M | 11 | 596 ± 279 |
| | TTGTGATTAACTTAGTTTGGTATAAAGTTCAGAGTCTCTTCTCACCTCTTGTAGGCTGCCAGGCTCGTAGCCTGTTGATAGCCCCTCAGCCCAGATGGGAGGAGATGGTGGACCGTTTCTGGCAGTATGTGTCTGAACTCAACACACAAACTGACGGCA | 75M7D4M1I79M | 9 | 409 ± 291 |
| | TTGTGATTAACTTAGTTTGGTATAAAGTTCACAGTCTCTTCTCACCTCTTGTAGGCTGCCAGGCTCGTAGCCAGGCTCGTAGCCCCTCAGCCCAGATGGGAGGAGATGGTGGACCGTTTCTGGCAGTATGTGTCTGAACTCAACACACAAACTGACGGCA | 31M1S38M6D7M2S1M1I79M | 8 | 660 ± 346 |
| | TTGTGATTAACTTAGTTTGGTATAAAGTTCACAGTCTCTTCTCACCTCTTGTAGGCTGCCAGGCTGATGCCCCTCAGCCCAGATGGGAGGAGATGGTGGACCGTTTCTGGCAGTATGTGTCTGAACTCAACACACAAACTGACGGCA | 31M1S26M18D89M | 8 | 658 ± 158 |
| | TTGTGATTAACTTAGTTTGGTATAAAGTTCAGAGTCTCTTCTCACCTCTTGTAGGCTGCCAGGCTCGTAGCCCCTCAGCCCAGATGGGAGGAGATGGTGGACCGTTTCTGGCAGTATGTGTCTGAACTCAACACACAAACTGACGGCA | 69M17D79M | 7 | 576 ± 469 |
| | TTGTGATTAACTTAGTTTGGTATAAAGTTCACAGTCTCTTCTCACCTCTTGTAGGCTGCCAGGCTCGTAGCCATCTGGGCTGATGCCCCTCAGCCCAGATGGGAGGAGATGGTGGACCGTTTCTGGCAGTATGTGTCTGAACTCAACACACAAACTGACGGCA | 31M1S40M2D1S2M2S86M | 6 | 676 ± 235 |
| | TTGTGATTAACTTAGTTTGGTATAAAGTTCACAGTCTCTTCTCACCTCTTGTAGGCTGCCAGGCTCGTAGCCTGTTCCAGGCTGATGCCCCTCAGCCGAGATGGGAGGAGATGGTGGACCGTTTCTGGCAGTATGTGTCTGAACTCAACACACAAACTGATGGCA | 31M1S65M1S62M1S4M | 3 | 213 ± 81 |
| | TTGTGATTAACTTAGTTTGGTATAAAGTTCACAGTCTCTTCTCACCTCTTGTAGGCTGAGATGGGAGGAGATGGTGGACCGTTTCTGGCAGTATGTGTCTGAACTCAACACACAAACTGATGGCA | 31M1S26M40D62M1S4M | 3 | 205 ± 157 |
| | TTGTGATTAACTTAGTTTGGTATAAAGTTCAGAGTCTCTTCTCACCTCTTGTAGGCTGCCAGGCTCGTAGCCTGTTATATATATATAACAGGCTGATGCCCCTCAGCCGAGATGGGAGGAGATGGTGAGATGGTGGACCGTTTCTGGCAGTATGTGTCTGAACTCAACACACAAACTGACGGCA | 76M9I1S20M1S9M8I58M | 2 | 27 ± 10 |
| | TTGTGATTAACTTAGTTTGGTATAAAGTTCAGAGTCTCTTCTCACCTCTTGTAGGCTGCCAGGCTCGTAGCCCAGGCTGATGCCCCTCAGCCCAGATGGGAGGAGATGGTGGACCGTTTCTGGCAGTATGTGTCTGAACTCAACACACAAACTGACGGCA | 71M5D89M | 1 | 2044 |
| | TTGTGATTAACTTAGTTTGGTATAAAGTTCAGAGTCTCTTCTCACCTCTTGTAGGCTGCCAGGCTCGTAGCTGATGCCCCTCAGCCCAGATGGGAGGAGATGGTGGACCGTTTCTGGCAGTATGTGTCTGAACTCAACACACAAACTGACGGCA | 69M11D85M | 1 | 1032 |
| *apoba* | CCATGTTGGGTGTAGCAGAGCTGTCAGCTAAGATGAACAGCAACTTTTATGAATGGGAGGGCTCTATCTATCTATGAATGGGAGGGGCAACAACACTGTTGATGTTCCCAATTACATTGCTAAGTACAAAATAATGGCCAGCTGCCCCTTAA | 69M11I3S43M1S25M | 122 | 666 ± 434 |
| | CCATGTTGGGTGTAGCAGAGCTGTCAGCTAAGATGAACAGCAACTTTTATGAATGGGAGGGCAACAACACTGTTGATGTTCCCAATTACATTGCTAAGTACAAAATAATGGCCAGCTGCCCCTTAA | 58M15D42M1S25M | 119 | 782 ± 674 |
| | CCATGTTGGGTGTAGCAGAGCTGTCAGCTAAGATGAACAGCAACTTTTATGAATGGGAGGGCTCTAGGGGGCAACAACACTGTTGATGTTCCCAATTACATTGCTAAGTACAAAATAATGGCCAGCTGCCCCTTAA | 64M5D46M1S25M | 91 | 742 ± 433 |
| | CCATGTTGGGTGTAGCAGAGCTGTCAGCTAAGATGAACAGCAACTTTTATGAATGGGAGGGGCAACAACACTGTTGATGTTCCCAATTACATTGCTAAGTATAAAATAATGGCCAGCTGCCCCTTAA | 58M14D69M | 90 | 655 ± 474 |
| | CCATGTTGGGTGTAGCAGAGCTGTCAGCTAAGATGAACAGCAACTTTTATGAATGGGAGGGCTCTATCTGTGCAACAACACTGTTGATGTTCCCAATTACATTGCTAAGTACAAAATAATGGCCAGCTGCCCCTTAA | 69M4D1M1S40M1S25M | 86 | 752 ± 426 |
| | CCATGTTGGGTGTAGCAGAGCTGTCAGCTAAGATGAACAGCAACTTTTATGAATGGGAGGGCTCTATCTAGGGGGCAACAACACTGTTGATGTTCCCAATTACATTGCTAAGTACAAAATAATGGCCAGCTGCCCCTTAA | 68M1D46M1S25M | 76 | 758 ± 458 |
| | CCATGTTGGGTGTAGCAGAGCTGTCAGCTAAGATGAACAGCAACTTTTATGAATGGGAGGGCTCTAGGGGGCAACATCAACACTGTTGATGTTCCCAATTACATTGCTAAGTACAAAATAATGGCCAGCTGCCCCTTAA | 64M5D10M3I36M1S25M | 56 | 795 ± 540 |
| | CCATGTTGGGTGTAGCAGAGCTGTCAGCTAAGATGAACAGCAACTTTTATGAATGGGAGGGGCAACAACACTGTTGATGTTCCCAATTACATTGCTAAGTACAAAATAATGGCCAGCTGCCCCTTAA | 58M14D43M1S25M | 29 | 588 ± 284 |
| | CCATGTTGGGTGTAGCAGAGCTGTCAGCTAAGATGAACAGCAACTTTTATGAATGGGAGGGCTCTATCTCCCATTCATAACTGGGGCAACAACACTGTTGATGTTCCCAATTACATTGCTAAGTATAAAATAATGGCCAGCTGCCCCTTAA | 69M8I2M1S2I69M | 18 | 627 ± 372 |
| | CCATGTTGGGTGTAGCAGAGCTGTCAGCTAAGATGAACAGCAACTTTTATGAATGGGAGGGCTCTATCTAACACTGTTGATGTTCCCAATTACATTGCTAAGTACAAAATAATGGCCAGCTGCCCCTTAA | 69M11D35M1S25M | 16 | 805 ± 385 |
| | CCATGTTGGGTGTAGCAGAGCTGTCAGCTAAGATGAACAGCAACTTTTATGAATGGGAGGGCTCTATCTCTATCTGGAAGGGGGCAACAACACTGTTGATGTTCCCAATTACATTGCTAAGTACAAAATAATGGCCAGCTGCCCCTTAA | 69M1S8I45M1S25M | 12 | 612 ± 275 |
| | CCATGTTGGGTGTAGCAGAGCTGTCAGCTAAGATGAACAGCAACTTTTATGAATGGGAGGGCTCTATCTATAGGGGGCAACAACACTGTTGATGTTCCCAATTACATTGCTAAGTACAAAATAATGGCCAGCTGCCCCTTAA | 69M1I46M1S25M | 8 | 641 ± 302 |
| | CCATGTTGGGTGTAGCAGAGCTGTCAGCTAAGATGAACAGCAACTTTTATGAATGGGAGGGCAACAACACTGTTGATGTTCCCAATTACATTGCTAAGTATAAAATAATGGCCAGCTGCCCCTTAA | 58M15D68M | 7 | 380 ± 249 |
| | CCATGTTGGGTGTAGCAGAGCTGTCAGCTAAGATGAACAGCAACTTTTATGAATGGGAGGGCTCTATCTGTTGATGTTCCCAATTACATTGCTAAGTATAAAATAATGGCCAGCTGCCCCTTAA | 67M17D57M | 6 | 698 ± 295 |
| | CCATGTTGGGTGTAGCAGAGCTGTCAGCTAAGATGAACAGCAACTTTTATGAATGGGAGGGCTCTATCAGCAACTTTTATGAATGGGGGGCAACAACACTGTTGATGTTCCCAATTACATTGCTAAGTATAAAATAATGGCCAGCTGCCCCTTAA | 68M14I1S1M1S70M | 5 | 497 ± 382 |
| | CCATGTTGGGTGTAGCAGAGCTGTCAGCTAAGATGAACAGCAACTTTTATGAATGGGAGGGCTCTATCTTCACAGCTAGGGGGCAACAACACTGTTGATGTTCCCAATTACATTGCTAAGTACAAAATAATGGCCAGCTGCCCCTTAA | 69M7I46M1S25M | 4 | 666 ± 97 |
| | CCATGTTGGGTGTAGCAGAGCTGTCAGCTAAGATGAACAGCAACTTTTATGAATGGGAGGGCTCTATCTGGGGGGCAATGGGGGGCAACAACACTGTTGATGTTCCCAATTACATTGCTAAGTACAAAATAATGGCCAGCTGCCCCTTAA | 69M9I1M1S44M1S25M | 3 | 590 ± 32 |
| | CCATGTTGGGTGTAGCAGAGCTGTCAGCTAAGATGAACAGCAACTTTTATGAATGGGAGGGCTCTATCTACAACACTGTTGATGTTCCCAATTACATTGCTAAGTACAAAATAATGGCCAGCTGCCCCTTAA | 69M9D37M1S25M | 3 | 517 ± 460 |
| | CCATGTTGGGTGTAGCAGAGCTGTCAGCTAAGATGAACAGCAACTTTTATGAATGGGAGGGGGCAACAACACTGTTGATGTTCCCAATTACATTGCTAAGTACAAAATAATGGCCAGCTGCCCCTTAA | 57M13D45M1S25M | 2 | 816 ± 240 |
| | CCATGTTGGGTGTAGCAGAGCTGTCAGCTAAGATGAACAGCAACTTTTATGAATGGGAGGGCTCTACACTGTTGATGTTCCCAATTACATTGCTAAGTATAAAATAATGGCCAGCTGCCCCTTAA | 65M16D60M | 2 | 681 ± 57 |
| | CCATGTTGGGTGTAGCAGAGCTGTCAGCTAAGATGAACAGCAACTTTTATGAATGGGAGGGCTCTATCTATCTTTTATGAATGGGGGGGCAACAACACTGTTGATGTTCCCAATTACATTGCTAAGTATAAAATAATGGCCAGCTGCCCCTTAA | 68M13I1M2S70M | 1 | 1120 |

***continued*** **Supplementary Table 13**

| | | | | |
|---|---|---|---|---|
| ***apobb.1*** | CTTCCGTGACACCATGCAGAAGACCATTAACTATGCAGCTGACAAAGTACCAAGGGGCAATGACATTATGCAGAGCATGTTCCCAACCCTATGGAATAACATCAAAATGCAAAAGGTCTATAAAATAGAGTTTATTCTTTATTTCCACACTGATGGCATCTTTTTTTTGTTCGATTTCTCACTCA | 160M1I24M | 252 | 772 ± 562 |
| | **CTTCCGTGACACCATGCAGAAGACCATTAACTATGCAGCTGACAAAGTACCAAGGGGCAATGACATTATGCAGAGCATGTTCCCAACCCTATGGAATAACATCAAAATGCAAAAGGTCTATAAAATAGAGTTTATTCTTTATTTCCACACTGATGGCATCTTTTTTTGTTCGATTTCTCACTCA** | **184M** | **213** | **750 ± 617** |
| | CTTCCGTGACACCATGCAGAAGACCATTAACTATGCAGCTGACAAAGTACAATGACATTATGCAGAGCAATGACATTATGCAGAGCATGTTCCCAACCCTATGGAATAACATCAAAATGCAAAAGGTCTATAAAATAGAGTTTATTCTTTATTTCCACACTGATGGCATCTTTTTTTTGTTCGATTTCTCACTCA | 50M1S1M1S1M10I1M1S104M1I24M | 101 | 639 ± 427 |
| | CTTCCGTGACACCATGCAGAAGACCATTAACTATGCAGCTGACAAAGGGGCAATGACATTATGCAGAGCATGTTCCCAACCCTATGGAATAACATCAAAATGCAAAAGGTCTATAAAATAGAGTTTATTCTTTATTTCCACACTGATGGCATCTTTTTTTTGTTCGATTTCTCACTCA | 44M7D109M1I24M | 73 | 651 ± 346 |
| | CTTCCGTGACACCATGCAGAAGACCATTAACTATGCAGCTGACAAAGTAAAGGGGCAATGACATTATGCAGAGCATGTTCCCAACCCTATGGAATAACATCAAAATGCAAAAGGTCTATAAAATAGAGTTTATTCTTTATTTCCACACTGATGGCATCTTTTTTTTGTTCGATTTCTCACTCA | 49M2D109M1I24M | 24 | 444 ± 206 |
| | CTTCCGTGACACCATGCAGAAGACCATTAACTATGCAGCTGACAAAGGGGCAATGACATTATGCAGAGCATGTTCCCAACCCTATGGAATAACATCAAAATGCAAAAGGTCTATAAAATAGAGTTTATTCTTTATTTCCACACTGATGGCATCTTTTTTTGTTCGATTTCTCACTCA | 44M7D133M | 14 | 887 ± 372 |
| | CTTCCGTGACACCATGCAGAAGACCATTAACTATGCAGCTGACAAAGTACCTAAGGGGCAATGACATTATGCAGAGCATGTTCCCAACCCTATGGAATAACATCAAAATGCAAAAGGTCTATAAAATAGAGTTTATTCTTTATTTCCACACTGATGGCATCTTTTTTTTGTTCGATTTCTCACTCA | 51M1I109M1I24M | 11 | 678 ± 327 |
| | CTTCCGTGACACCATGCAGAAGACCATTAACTATGCAGCTGACAAAGTAATGTTAAGGGGCAATGACATTATGCAGAGCATGTTCCCAACCCTATGGAATAACATCAAAATGCAAAAGGTCTATAAAATAGAGTTTATTCTTTATTTCCACACTGATGGCATCTTTTTTTTGTTCGATTTCTCACTCA | 49M3I2S109M1I24M | 6 | 907 ± 409 |
| | CTTCCGTGACACCATGCAGAAGACCATTAACTATGCAGCTGACAAAGTACAATGGGGCAATGACATTATGCAGAGCATGTTCCCAACCCTATGGAATAACATCAAAATGCAAAAGGTCTATAAAATAGAGTTTATTCTTTATTTCCACACTGATGGCATCTTTTTTTTGTTCGATTTCTCACTCA | 50M1S1M1S107M1I24M | 2 | 616 ± 257 |
| | CTTCCGTGACACCATGCAGAAGACCATTAACTATGCAGCTGGGCAATGACATTATGCAGAGCATGTTCCCAACCCTATGGAATAACATCAAAATGCAAAAGGTCTATAAAATAGAGTTTATTCTTTATTTCCACACTGATGGCATCTTTTTTTTGTTCGATTTCTCACTCA | 40M14D106M1I24M | 2 | 577 ± 34 |
| | CTTCCGTGACACCATGCAGAAGACCATTAACTATGCAGCTGACAAAGTACAATGACAAAGGGGCAATGACATTATGCAGAGCATGTTCCCAACCCTATGGAATAACATCAAAATGCAAAAGGTCTATAAAATAGAGTTTATTCTTTATTTCCACACTGATGGCATCTTTTTTTTGTTCGATTTCTCACTCA | 48M6I2M1S109M1I24M | 1 | 627 |
| | CTTCCGTGACACCATGCAGAAGACCATTAACTATGCAGCTGACATTATGCAGAGCATGTTCCCAACCCTATGGAATAACATCAAAATGCAAAAGGTCTATAAAATAGAGTTTATTCTTTATTTCCACACTGATGGCATCTTTTTTTGTTCGATTTCTCACTCA | 39M21D124M | 1 | 444 |
| ***apobb.2*** | GTCTTTATTATTTCCAGAGCCATCTTGCTCAGCTAATCAGGAGCAACCAGACCTGCAACTACAAGTTTGACAAAGAGCAGGAGCACATGACCTCTGCTATTTGCACCGAAAAACATGTTCTTGTGCCCTTTTCACACAAGTAAGCATCAATGTTAGAAACTTTGTTTAGAGATCTGCCT | 22M3D157M | 519 | 912 ± 862 |
| | GTCTTTATTATTTCCAGAGCCATCCTTGCTCAGCTAATCAGGAGCAACCAGACCTGCAACTACAAGTTTGACAAAGAGCAGGAGCACATGACCTCTGCTATTTGCACCGAAAAACATGTTCTTGTGCCCTTTTCACACAAGTAAGCATCAATGTTAGAAACTTTGTTTAGAGATCTGCCT | 24M2D156M | 62 | 665 ± 475 |
| | GTCTTTATTATTTCCAGAGCCATCAGCTAATCAGGAGCAACCAGACCTGCAACTACAAGTTTGACAAAGAGCAGGAGCACATGACCTCTGCTATTTGCACCGAAAAACATGTTCTTGTGCCCTTTTCACACAAGTAAGCATCAATGTTAGAAACTTTGTTTAGAGATCTGCCT | 22M9D151M | 43 | 426 ± 240 |
| | **GTCTTTATTATTTCCAGAGCCATCCTCTTGCTCAGCTAATCAGGAGCAACCAGACCTGCAACTACAAGTTTGACAAAGAGCAGGAGCACATGACCTCTGCTATTTGCACCGAAAAACATGTTCTTGTGCCCTTTTCACACAAGTAAGCATCAATGTTAGAAACTTTGTTTAGAGATCTGCCT** | **182M** | **21** | **437 ± 240** |
| | GTCTTTATTATTTCCAGAGCCAATCAGGAGCAACCAGACCTGCAACTACAAGTTTGACAAAGAGCAGGAGCACATGACCTCTGCTATTTGCACCGAAAAACATGTTCTTGTGCCCTTTTCACACAAGTAAGCATCAATGTTAGAAACTTTGTTTAGAGATCTGCCT | 21M16D145M | 18 | 704 ± 578 |
| | GTCTTTATTATTTCCAGAGCCATCCTTGCTTAGCTTGCTCAGCTTGCTCAGCTAATCAGGAGCAACCAGACCTGCAACTACAAGTTTGACAAAGAGCAGGAGCACATGACCTCTGCTATTTGCACCGAAAAACATGTTCTTGTGCCCTTTTCACACAAGTAAGCATCAATGTTAGAAACTTTGTTTAGAGATCTGC | 26M16I156M | 13 | 508 ± 367 |
| | GTCTTTATTATTTCCAGAGCCATCCAGACAACCAGACAACCAGGAGCAACCAGACCTGCAACTACAAGTTTGACAAAGAGCAGGAGCACATGACCTCTGCTATTTGCACCGAAAAACATGTTCTTGTGCCCTTTTCACACAAGTAAGCATCAATGTTAGAAACTTTGTTTAGAGATCTGCCT | 25M7S3M2S2M1S142M | 9 | 467 ± 183 |
| | GTCTTTATTATTTCCAGAGCTCAGCTAATCAGGAGCAACCAGACCTGCAACTACAAGTTTGACAAAGAGCAGGAGCACATGACCTCTGCTATTTGCACCGAAAAACATGTTCTTGTGCCCTTTTCACACAAGTAAGCATCAATGTTAGAAACTTTGTTTAGAGATCTGCCT | 18M11D153M | 6 | 751 ± 461 |
| | GTCTTTATTATTTCCAGAGCCATCTTTCTTGCTCAGCTAATCAGGAGCAACCAGACCTGCAACTACAAGTTTGACAAAGAGCAGGAGCACATGACCTCTGCTATTTGCACCGAAAAACATGTTCTTGTGCCCTTTTCACACAAGTAAGCATCAATGTTAGAAACTTTGTTTAGAGATCTGCCT | 24M1I1S157M | 5 | 22 ± 35 |
| | GTCTTTATTATTTCCAGAGCCATCCTGGAAATAAGCTCAGCTAATCAGGAGCAACCAGACCTGCAACTACAAGTTTGACAAAGAGCAGGAGCACATGACCTCTGCTATTTGCACCGAAAAACATGTTCTTGTGCCCTTTTCACACAAGTAAGCATCAATGTTAGAAACTTTGTTTAGAGATCTGCCT | 26M5I3S153M | 4 | 170 ± 141 |
| | GTCTTTATTATTTCCAGAGCCATCAGAGCATCCAGAGCTTGCTCAGCTAATCAGGAGCAACCAGACCTGCAACTACAAGTTTGACAAAGAGCAGGAGCACATGACCTCTGCTATTTGCACCGAAAAACATGTTCTTGTGCCCTTTTCACACAAGTAAGCATCAATGTTAGAAACTTTGTTTAGAGATCTGCCT | 24M11I2S156M | 3 | 730 ± 640 |
| | GTCTTTATTATTTCCAGAGCCATCTTTCTTGCTCAGCTAATCAGGAGCAACCAGACCTGCAACTACAAGTTTGACAAAGAGCAGGAGCACATGACCTCTGCTATTTGCACCGAAAAACATGTTCTTGTGCCCTTTTCACACAAGTAAGCATCAATGTTAGAAACTTAGTTTAGAGATCTGCCT | 24M1I1S140M1S16M | 3 | 5 ± 3 |
| | GTCTTTATTATGCAGATATTATGCTCAGCTAATCAGGAGCAACCAGACCTGCAACTACAAGTTTGACAAAGAGCAGGAGCACATGACCTCTGCTATTTGCACCGAAAAACATGTTCTTGTGCCCTTTTCACACAAGTAAGCATCAATGTTAGAAACTTTGTTTAGAGATCTGCCT | 11M7D2M5S1M2S154M | 2 | 109 ± 13 |
| | GTCTTTATTATTTCCAGAGCCATCAGCTTGCTCAGCTAATCAGGAGCAACCAGACCTGCAACTACAAGTTTGACAAAGAGCAGGAGCACATGACCTCTGCTATTTGCACCGAAAAACATGTTCTTGTGCCCTTTTCACACAAGTAAGCATCAATGTTAGAAACTTTGTTTAGAGATCTGCCT | 24M2S156M | 1 | 1439 |
| | GTCTTTATTATTTCCAGAGCCATCCTCTTGCTCAGCTAATCAGGAGCAACCAGACCTGCAACTACAAGTTTGACAAAGAGCAGGAGCACATGACCTCTGCTATTTGCACCGAAAAACATGTTCTTGTGCCCTTTTCACACAAGTAAGCATCAATGTTAGAAACTTAGTTTAGAGATCTGCCT | 165M1S16M | 1 | 7 |
| ***ldlra*** | **GAGGCGTGTTTGTTTAGTTTTTTAAGTATCAGCATAGTTTGGAATAATTAGCGTACGGCACCCTTAGTCAAGGAGAATGTGTCACTTACAGTATTTCTCCTTCCAACAGGTGCCTTGACTTGTGATTCACGGCAGTATCAGTGTGGCAATGGAAAGTGCATCACGGCGAGATGGGTGTGTGATG** | **184M** | **241** | **1176 ± 838** |
| | GAGGCGTGTTTGTTTAGTTTTTTAAGTATCAGCATAGTTTGGAATAATTAGCGTACGGCACCCTTAGTCAAGGAGAATGTGTCACTTACAGTATTTCTCCTTCCAACAGGTGCCTTGACTTGTGATTCACGGCACTACAGTAGTGTGGCAATGGAAAGTGCATCACGGCGAGATGGGTGTGTGATG | 134M1S2M1S2I1S45M | 159 | 982 ± 595 |
| | GAGGCGTGTTTGTTTAGTTTTTTAAGTATCAGCATAGTTTGGAATAATTAGCGTACGGCACCCTTAGTCAAGGAGAATGTGTCACTTACAGTATTTCTCCTTCCAACAGGTGCCTTGACTTGTGATTCACGGCAGTATCATGTGGCAATGGAAAGTGCATCACGGCGAGATGGGTGTGTGATG | 140M1D43M | 119 | 1179 ± 722 |
| | GAGGCGTGTTTGTTTAGTTTTTTAAATATCAGCATAGTTTGGAATAATTAGCGTATGGCACCCTTAGTCAAGGAGAATGTGTCACTTACAGTATTTCTCCTTCCAACAGGTGCCTTGACTTGTGATTCACGGCAGTATCAGTGTGGCAATGGAAAGTGCATCACGGCGAGATGGGTGTGTGATG | 25M1S29M1S128M | 64 | 888 ± 609 |
| | GAGGCGTGTTTGTTTAGTTTTTTAAGTATCAGCATAGTTTGGAATAATTAGCGTACGGCACCCTTAGTCAAGGAGAATGTGTCACTTACAGTATTTCTCCTTCCAACAGGTGCCTTGACTTGTGATTCACGGCAGTATCAAGTGTGGCAATGGAAAGTGCATCACGGCGAGATGGGTGTGTGATG | 139M1I45M | 53 | 937 ± 655 |
| | GAGGCGTGTTTGTTTAGTTTTTTAAATATCAGCATAGTTTGGAATAATTAGCGTATGGCACCCTTAGTCAAGGAGAATGTGTCACTTACAGTATTTCTCCTTCCAACAGGTGCCTTGACTTGTGATTCACGGCAGTGGCAATGGAAAGTGCATCACGGCGAGATGGGTGTGTGATG | 25M1S29M1S78M8D42M | 47 | 891 ± 617 |
| | GAGGCGTGTTTGTTTAGTTTTTTAAGTATCAGCATAGTTTGGAATAATTAGCGTACGGCACCCTTAGTCAAGGAGAATGTGTCACTTACAGTATTTCTCCTTCCAACAGGTGCCTTGACTTGTGATTCACGGCAGTGGCAATGGAAAGTGCATCACGGCGAGATGGGTGTGTGATG | 134M8D42M | 42 | 832 ± 397 |
| | GAGGCGTGTTTGTTTAGTTTTTTAAGTATCAGCATAGTTTGGAATAATTAGCGTACGGCACCCTTAGTCAAGGAGAATGTGTCACTTACAGTATTTCTCCTTCCAACAGGTGCCTTGACTTGTGATTCACGGCAGTGTGGCAATGGAAAGTGCATCACGGCGAGATGGGTGTGTGATG | 132M6D46M | 21 | 741 ± 441 |
| | GAGGCGTGTTTGTTTAGTTTTTTAAATATCAGCATAGTTTGGAATAATTAGCGTATGGCACCCTTAGTCAAGGAGAATGTGTCACTTACAGTATTTCTCCTTCCAACAGGTGCCTTGACTTGTGATTCACGGCAGTGATTCACGGCAGTGTGGCAATGGAAAGTGCATCACGGCGAGATGGGTGTGTGATG | 25M1S29M1S80M7I2S46M | 7 | 528 ± 324 |
| | GAGGCGTGTTTGTTTAGTTTTTTAAATATCAGCATAGTTTGGAATAATTAGCGTATGGCACCCTTAGTCAAGGAGAATGTGTCACTTACAGTATTTCTCCTTCCAACAGGTGCCTTGACTTGTGATTCACGGCAGTATCAAGTGTGGCAATGGAAAGTGCATCACGGCGAGATGGGTGTGTGATG | 25M1S29M1S83M1I45M | 4 | 395 ± 245 |
| | GAGGCGTGTTTGTTTAGTTTTTTAAGTATCAGCATAGTTTGGAATAATTAGCGTACGGCACCCTTAGTCAAGGAGAATGTGTCACTTACAGTATTTCTCCTTCCAACAGGTGCCTTGACTTGTGATTCACGGCAGTATCAATGGAAAGTGGCAATGGAAAGTGCATCACGGCGAGATGGGTGTGTGATG | 140M1S1M5I42M | 2 | 1236 ± 827 |
| | GAGGCGTGTTTGTTTAGTTTTTTAAATATCAGCATAGTTTGGAATAATTAGCGTATGGCACCCTTAGTCAAGGAGAATGTGTCACTTACAGTATTTCTCCTTCCAACAGGTGCCTTGACTTGTGATTCACGGCAGTGTGGCAATGGAAAGTGCATCACGGCGAGATGGGTGTGTGATG | 25M1S29M1S76M6D46M | 2 | 655 ± 65 |
| | GAGGCGTGTTTGTTTAGTTTTTTAAATATCAGCATAGTTTGGAATAATTAGCGTATGGCACCCTTAGTCAAGGAGAATGTGTCACTTACAGTATTTCTCCTTCCAACAGGTGCCTTGACTTGTGATTCACGGCAGTATGGCAATGGAAAAGTGCATCACGGCGAGATGGGTGTGTGATG | 25M1S29M1S81M6D9M1I32M | 1 | 1152 |
| ***ldlrb*** | CCGGAGCCCCGCCCCCCGCGGCTTCATTCAAATGCTCGGCTTCTGCTCAGAGAGGGAGAATCTCCGCCACCGCAGAATCCGCTCCCGGTACAGAACAGCATCTCTATCCGCCCTCAGAGCTCTAACGTGACGCTGATGGAGCCCCGGGTCGGCGTTCGGCTCGTCTGCTCG | 70M4D101M | 208 | 484 ± 494 |
| | CCGGAGCCCCGCCCCCCGCGGCTTCATTCAAATGCTCGGCTTCTGCTCAGAGAGGGAGAATCTCCGCCACATTCTCGCAGAATCCGCTCCCGGTACAGAACAGCATCTCTATCCGCCCTCAGAGCTCTAACGTGACGCTGATGGAGCCCCGGGTCGGCGTTCGGCTCGTCTGCTCG | 70M1I2S103M | 178 | 276 ± 256 |
| | CCGGAGCCCCGCCCCCCGCGGCTTCATTCAAATGCTCGGCTTCTGCTCAGAGAGGGAGAATCTCCGCCACTCGCAGAATCCGCTCCCGGTACAGAACAGCATCTCTATCCGCCCTCAGAGCTCTAACGTGACGCTGATGGAGCCCCGGGTCGGCGTTCGGCTCGTCTGCTCG | 69M3D103M | 113 | 212 ± 180 |
| | CCGGAGCCCCGCCCCCCGCGGCTTCATTCAAATGCTCGGCTTCTGCTCAGAGAGGGAGAATCTCCGCCACCGCCATCGCAGAATCCGCTCCCGGTACAGAACAGCATCTCTATCCGCCCTCAGAGCTCTAACGTGACGCTGATGGAGCCCCGGGTCGGCGTTCGGCTCGTCTGCTCG | 73M2I102M | 56 | 238 ± 156 |
| | CCGGAGCCCCGCCCCCCGCGGCTTCATTCAAATGCTCGGCTTCTGCTCAGAGAGGGAGAATCTCCGCCACAGAATCCGCTCCCGGTACAGAACAGCATCTCTATCCGCCCTCAGAGCTCTAACGTGACGCTGATGGAGCCCCGGGTCGGCGTTCGGCTCGTCTGCTCG | 69M7D99M | 22 | 164 ± 239 |
| | CCGGAGCCCCGCCCCCCGCGGCTTCATTCAAATGCTCGGCTTCTGCTCAGAGAGGGAGAATCTCCGCCACCGAAGAATCGCAGAATCCGAGAATCGCAGAATCCGCTCCCGGTACAGAACAGCATCTCTATCCGCCCTCAGAGCTCTAACGTGACGCTGATGGAGCCCCGGGTCGGCGTTCGGCTCGTCTGCTCG | 72M20I1S102M | 16 | 297 ± 208 |
| | CCGGAGCCCCGCCCCCCGCGGCTTCATTCAAATGCTCGGCTTCTGCTCAGAGAGGGAGAATCTCCGCCACCGCAGGGAGAATCTCTCGCAGAATCCGCTCCCGGTACAGAACAGCATCTCTATCCGCCCTCAGAGCACTAACGTGACGCTGATGGAGCCCCGGGTCGGCGCTCGGCTCGTCTGCTCG | 72M14I52M1S33M1S16M | 15 | 303 ± 287 |
| | CCGGAGCCCCGCCCCCCGCGGCTTCATTCAAATGCTCGGCTTCTGCTCAGAGAGGGAGAATCTCCGCCTCTCTCGCAGAATCCGCTCCCGGTACAGAACAGCATCTCTATCCGCCCTCAGAGCTCTAACGTGACGCTGATGGAGCCCCGGGTCGGCGTTCGGCTCGTCTGCTCG | 68M1D1S1M1S103M | 12 | 143 ± 116 |
| | CCGGAGCCCCGCCCCCCGCGGCTTCATTCAAATGCTCGGCTTCTGCTCAGAGAGGGAGAATCTCTCGCAGAATCCGCTCCCGGTACAGAACAGCATCTCTATCCGCCCTCAGAGCACTAACGTGACGCTGATGGAGCCCCGGGTCGGCGCTCGGCTCGTCTGCTCG | 63M9D52M1S33M1S16M | 11 | 583 ± 305 |
| | CCGGAGCCCCGCCCCCCGCGGCTTCATTCAAATGCTCGGCTTCTGCTCAGAGAGGGAGAATCTCCGCCACCGCTCCCGGTACAGAACAGCATCTCTATCCGCCCTCAGAGCTCTAACGTGACGCTGATGGAGCCCCGGGTCGGCGTTCGGCTCGTCTGCTCG | 69M13D93M | 11 | 547 ± 354 |
| | CCGGAGCCCCGCCCCCCGCGGCTTCATTCAAATGCTCGGCTTCTGCTCAGAGAGGGAGAATCTCCGCCACCGATGGAGCCCCGGGTCGGCGTTCGGCTCGTCTGCTCG | 71M67D37M | 8 | 822 ± 500 |
| | CCGGAGCCCCGCCCCCCGCGGCTTCATTCAAATGCTCGGCTTCTGCTCAGAGAGGGAGAATCTCCGCCACCTCGCAGAATCCGCTCCCGGTACAGAACAGCATCTCTATCCGCCCTCAGAGCTCTAACGTGACGCTGATGGAGCCCCGGGTCGGCGTTCGGCTCGTCTGCTCG | 70M2D103M | 4 | 584 ± 872 |
| | CCGGAGCCCCGCCCCCCGCGGCTTCATTCAAATGCTCGGCTTCTGCTCAGAGAGGGAGAATCTCCGCCTCCGCATTCTGTTCGCAGAATCCGCTCCCGGTACAGAACAGCATCTCTATCCGCCCTCAGAGCTCTAACGTGACGCTGATGGAGCCCCGGGTCGGCGTTCGGCTCGTCTGCTCG | 68M1S4M7I102M | 2 | 1064 |
| | **CCGGAGCCCCGCCCCCCGCGGCTTCATTCAAATGCTCGGCTTCTGCTCAGAGAGGGAGAATCTCCGCCACCGCTCGCAGAATCCGCTCCCGGTACAGAACAGCATCTCTATCCGCCCTCAGAGCTCTAACGTGACGCTGATGGAGCCCCGGGTCGGCGTTCGGCTCGTCTGCTCG** | **175M** | **2** | **7** |

Annotation shows the sequential number of base pairs that - when compared with the reference genome from Ensembl - represent a match (M), deletion (D), insertion (I), or substitution (S). For each unique sequence, the number of alleles in which it was observed is shown, as well as the mean and standrard deviation for the number of reads that were observed for the sequence.

**Supplementary Table 14 - Unique CRISPR-Cas9-induced variants in the most prominently observed sequence(s) and their predicted functional consequences**

| Zebrafish orthologue | Chr | Start | End | Mutation | Nett base pair change | Annotation VEP | VEP impact | $n_{affected\ alleles}$ |
|---|---|---|---|---|---|---|---|---|
| | | 10,856,052 | 10,856,073 | CTCCAAACTGGCCCCTTACGCG/- | -22 | frameshift variant | high | 2 |
| | | 10,856,058 | 10,856,068 | ACTGGCCCCTT/- | -11 | frameshift variant | high | 116 |
| | | 10,856,062 | 10,856,070 | GCCCCTTAC/- | -9 | inframe deletion | moderate | 1 |
| | | 10,856,062 | 10,856,072 | GCCCCTTACGC/- | -11 | frameshift variant | high | 7 |
| | | 10,856,064 | 10,856,069 | CCCTTA/- | -6 | inframe deletion | moderate | 1 |
| | | 10,856,065 | 10,856,069 | CCTTA/- | -5 | frameshift variant | high | 49 |
| | | 10,856,065 | 10,856,073 | CCTTACGCG/- | -9 | inframe deletion | moderate | 12 |
| | | 10,856,066 | 10,856,065 | -/CCCT | 4 | frameshift variant | high | 14 |
| | | 10,856,067 | 10,856,068 | TT/AG | 0 | missense variant | moderate | 15 |
| | | 10,856,067 | 10,856,068 | TT/- | -2 | frameshift variant | high | 1 |
| *apoea* | 19 | 10,856,068 | 10,856,067 | -/TGA | 3 | stop gained,inframe insertion | high | 2 |
| | | 10,856,068 | 10,856,068 | T/C | 0 | missense variant | moderate | 29 |
| | | 10,856,068 | 10,856,069 | TA/GG | 0 | missense variant | moderate | 7 |
| | | 10,856,068 | 10,856,069 | TA/CT | 0 | missense variant | moderate | 14 |
| | | 10,856,068 | 10,856,075 | TACGCGCA/- | -8 | frameshift variant | high | 65 |
| | | 10,856,069 | 10,856,068 | -/CAA | 3 | protein altering variant | moderate | 29 |
| | | 10,856,069 | 10,856,072 | ACGC/- | -4 | frameshift variant | high | 5 |
| | | 10,856,070 | 10,856,070 | C/G | 0 | stop gained | high | 15 |
| | | 10,856,071 | 10,856,071 | G/C | 0 | missense variant | moderate | 7 |
| | | 10,856,072 | 10,856,072 | C/A | 0 | missense variant | moderate | 15 |
| | | 10,856,073 | 10,856,072 | -/G | 1 | frameshift variant | high | 15 |
| | | 10,856,074 | 10,856,074 | C/G | 0 | missense variant | moderate | 5 |
| | | 23,961,533 | 23,961,550 | CCAGGCTCGTAGCCTGTT/- | -18 | inframe deletion | moderate | 7 |
| | | 23,961,533 | 23,961,572 | CCAGGCTCGTAGCCTGTTCCAGGCTGATGCCCCTCAGCCC/- | -40 | frameshift variant | high | 3 |
| | | 23,961,544 | 23,961,554 | GCCTGTTCCAG/- | -11 | frameshift variant | high | 1 |
| | | 23,961,544 | 23,961,560 | GCCTGTTCCAGGCTGAT/- | -17 | frameshift variant | high | 4 |
| | | 23,961,545 | 23,961,550 | CCTGTT/- | -6 | inframe deletion | moderate | 47 |
| | | 23,961,546 | 23,961,548 | CTG/TGA | 0 | stop gained | high | 58 |
| | | 23,961,546 | 23,961,550 | CTGTT/- | -5 | frameshift variant | high | 1 |
| | | 23,961,546 | 23,961,555 | CTGTTCCAGG/- | -10 | frameshift variant | high | 45 |
| | | 23,961,547 | 23,961,548 | TG/- | -2 | frameshift variant | high | 5 |
| | | 23,961,547 | 23,961,559 | TGTTCCAGGCTGA/- | -13 | frameshift variant | high | 8 |
| | | 23,961,548 | 23,961,552 | GTTCC/- | -5 | frameshift variant | high | 11 |
| | | 23,961,549 | 23,961,549 | T/A | 0 | missense variant | moderate | 5 |
| | | 23,961,550 | 23,961,550 | T/C | 0 | missense variant | moderate | 68 |
| *apoeb* | 16 | 23,961,550 | 23,961,553 | TCCA/- | -4 | frameshift variant | high | 58 |
| | | 23,961,550 | 23,961,556 | TCCAGGC/- | -7 | frameshift variant | high | 8 |
| | | 23,961,550 | 23,961,559 | TCCAGGCTGA/- | -10 | frameshift variant | high | 17 |
| | | 23,961,551 | 23,961,550 | -/AAGTG | 5 | frameshift variant | high | 68 |
| | | 23,961,551 | 23,961,550 | -/ATATATATA | 9 | protein altering variant | moderate | 62 |
| | | 23,961,551 | 23,961,550 | -/G | 1 | frameshift variant | high | 97 |
| | | 23,961,551 | 23,961,551 | C/A | 0 | missense variant | moderate | 130 |
| | | 23,961,552 | 23,961,553 | CA/TG | 0 | missense variant | moderate | 5 |
| | | 23,961,553 | 23,961,555 | AGG/CCT | 0 | missense variant | moderate | 97 |
| | | 23,961,557 | 23,961,558 | TG/GT | 0 | missense variant | moderate | 97 |
| | | 23,961,558 | 23,961,558 | G/A | 0 | missense variant | moderate | 11 |
| | | 23,961,558 | 23,961,559 | GA/CG | 0 | missense variant | moderate | 7 |
| | | 23,961,561 | 23,961,560 | -/A | 1 | frameshift variant | high | 15 |
| | | 23,961,572 | 23,961,572 | C/A | 0 | synonymous variant | low | 11 |
| | | 30,717,993 | 30,718,005 | AGGGCTCTATCTT/- | -13 | frameshift variant | high | 2 |
| | | 30,717,994 | 30,718,007 | GGGCTCTATCTTAG/- | -14 | frameshift variant | high | 119 |
| | | 30,717,994 | 30,718,008 | GGGCTCTATCTTAGG/- | -15 | inframe deletion | moderate | 103 |
| | | 30,718,000 | 30,718,004 | TATCT/- | -5 | frameshift variant | high | 139 |
| | | 30,718,001 | 30,718,016 | ATCTTAGGGGGCAACA/- | -16 | frameshift variant | high | 2 |
| | | 30,718,003 | 30,718,019 | CTTAGGGGGCAACAACA/- | -17 | frameshift variant | high | 6 |
| | | 30,718,004 | 30,718,003 | -/AGCAACTTTTATGA | 14 | frameshift variant | high | 5 |
| | | 30,718,004 | 30,718,003 | -/TATCTTTTATGAA | 13 | stop gained,frameshift variant | high | 1 |
| | | 30,718,004 | 30,718,004 | T/- | -1 | frameshift variant | high | 68 |
| | | 30,718,004 | 30,718,004 | T/A | 0 | missense variant | moderate | 5 |
| | | 30,718,005 | 30,718,004 | -/ATCTATGAATG | 11 | stop gained,frameshift variant | high | 103 |
| | | 30,718,005 | 30,718,004 | -/TCACAGC | 7 | frameshift variant | high | 4 |
| | | 30,718,005 | 30,718,004 | -/GGGGGGCAA | 9 | protein altering variant | moderate | 3 |
| *apoba* | 17 | 30,718,005 | 30,718,004 | -/CCCATTCA | 8 | frameshift variant | high | 18 |
| | | 30,718,005 | 30,718,004 | -/A | 1 | frameshift variant | high | 8 |
| | | 30,718,005 | 30,718,005 | T/C | 0 | missense variant | moderate | 12 |
| | | 30,718,005 | 30,718,006 | TA/GG | 0 | missense variant | moderate | 1 |
| | | 30,718,005 | 30,718,007 | TAG/GGA | 0 | missense variant | moderate | 103 |
| | | 30,718,005 | 30,718,008 | TAGG/- | -4 | frameshift variant | high | 70 |
| | | 30,718,005 | 30,718,013 | TAGGGGGCA/- | -9 | inframe deletion | moderate | 3 |
| | | 30,718,005 | 30,718,015 | TAGGGGGCAAC/- | -11 | frameshift variant | high | 16 |
| | | 30,718,006 | 30,718,005 | -/TATCTGGA | 8 | frameshift variant | high | 12 |
| | | 30,718,006 | 30,718,006 | A/G | 0 | synonymous variant | low | 8 |
| | | 30,718,007 | 30,718,007 | G/A | 0 | missense variant | moderate | 18 |
| | | 30,718,008 | 30,718,007 | -/CT | 2 | frameshift variant | high | 18 |
| | | 30,718,010 | 30,718,010 | G/T | 0 | missense variant | moderate | 70 |
| | | 30,718,015 | 30,718,014 | -/CAT | 3 | inframe insertion | moderate | 48 |
| | | 31,273,922 | 31,273,942 | TGACAAAGTACCAAGGGGCAA/- | -21 | inframe deletion | moderate | 1 |
| | | 31,273,923 | 31,273,936 | GACAAAGTACCAAG/- | -14 | frameshift variant | high | 2 |
| *apobb.1* | 20 | 31,273,927 | 31,273,933 | AAGTACC/- | -7 | frameshift variant | high | 82 |
| | | 31,273,931 | 31,273,930 | -/ACAATG | 6 | stop gained,inframe insertion | high | 1 |
| | | 31,273,932 | 31,273,931 | -/ATG | 3 | inframe insertion | moderate | 6 |
| | | 31,273,932 | 31,273,933 | CC/TT | 0 | missense variant | moderate | 6 |

***continued*** **Supplementary Table 14**

| Zebrafish orthologue | Chr | Start | End | Mutation | Nett base pair change | Annotation VEP | VEP impact | $n_{affected\ alleles}$ |
|---|---|---|---|---|---|---|---|---|
| ***apobb.1*** | 20 | 31,273,932 | 31,273,933 | CC/- | -2 | frameshift variant | high | 24 |
| | | 31,273,933 | 31,273,933 | C/A | 0 | missense variant | moderate | 94 |
| | | 31,273,934 | 31,273,933 | -/T | 1 | frameshift variant | high | 11 |
| | | 31,273,935 | 31,273,935 | A/T | 0 | missense variant | moderate | 93 |
| | | 31,273,937 | 31,273,936 | -/ACATTATGCA | 10 | frameshift variant | high | 91 |
| | | 31,273,938 | 31,273,938 | G/A | 0 | missense variant | moderate | 91 |
| ***apobb.2*** | 20 | 53,444,321 | 53,444,327 | TTCCAGA/- | -7 | splice acceptor variant,coding sequence variant,intron variant | high | 2 |
| | | 53,444,328 | 53,444,338 | GCCATCCTCTT/- | -11 | frameshift variant,splice region variant | high | 3 |
| | | 53,444,330 | 53,444,334 | CATCC/AGATA | 0 | missense variant | moderate | 2 |
| | | 53,444,331 | 53,444,346 | ATCCTCTTGCTCAGCT/- | -16 | frameshift variant | high | 12 |
| | | 53,444,332 | 53,444,334 | TCC/- | -3 | inframe deletion | moderate | 307 |
| | | 53,444,332 | 53,444,340 | TCCTCTTGC/- | -9 | inframe deletion | moderate | 33 |
| | | 53,444,334 | 53,444,333 | -/AGAGCATCCAG | 11 | frameshift variant | high | 3 |
| | | 53,444,334 | 53,444,333 | -/T | 1 | frameshift variant | high | 8 |
| | | 53,444,334 | 53,444,334 | C/T | 0 | missense variant | moderate | 8 |
| | | 53,444,334 | 53,444,335 | CT/AG | 0 | missense variant | moderate | 4 |
| | | 53,444,334 | 53,444,335 | CT/- | -2 | frameshift variant | high | 51 |
| | | 53,444,335 | 53,444,341 | TCTTGCT/AGACAAC | 0 | missense variant | moderate | 9 |
| | | 53,444,336 | 53,444,335 | -/GGAAA | 5 | frameshift variant | high | 4 |
| | | 53,444,336 | 53,444,335 | -/TGCTTAGCTTGCTCAG | 16 | frameshift variant | high | 13 |
| | | 53,444,336 | 53,444,337 | CT/TA | 0 | missense variant | moderate | 2 |
| | | 53,444,336 | 53,444,338 | CTT/TAA | 0 | stop gained | high | 4 |
| | | 53,444,345 | 53,444,346 | CT/AC | 0 | missense variant | moderate | 9 |
| | | 53,444,349 | 53,444,349 | T/C | 0 | missense variant | moderate | 9 |
| ***ldlra*** | 3 | 19,304,770 | 19,304,775 | CAGTAT/- | -6 | inframe deletion | moderate | 22 |
| | | 19,304,772 | 19,304,772 | G/C | 0 | missense variant | moderate | 97 |
| | | 19,304,772 | 19,304,779 | GTATCAGT/- | -8 | frameshift variant | high | 63 |
| | | 19,304,774 | 19,304,773 | -/GATTCAC | 7 | stop gained,frameshift variant | high | 7 |
| | | 19,304,774 | 19,304,775 | AT/GG | 0 | missense variant | moderate | 7 |
| | | 19,304,775 | 19,304,775 | T/C | 0 | synonymous variant | low | 97 |
| | | 19,304,775 | 19,304,780 | TCAGTG/- | -6 | inframe deletion | moderate | 1 |
| | | 19,304,776 | 19,304,775 | -/AG | 2 | frameshift variant | high | 97 |
| | | 19,304,776 | 19,304,776 | C/T | 0 | stop gained | high | 97 |
| | | 19,304,777 | 19,304,776 | -/A | 1 | frameshift variant | high | 39 |
| | | 19,304,778 | 19,304,778 | G/- | -1 | frameshift variant | high | 73 |
| | | 19,304,778 | 19,304,778 | G/A | 0 | synonymous variant | low | 2 |
| | | 19,304,780 | 19,304,779 | -/GGAAA | 5 | frameshift variant | high | 2 |
| | | 19,304,790 | 19,304,789 | -/A | 1 | frameshift variant | high | 1 |
| ***ldlrb*** | 6 | 102,538 | 102,546 | CCGCCACCG/- | -9 | upstream gene variant | modifier | 7 |
| | | 102,543 | 102,543 | A/T | 0 | upstream gene variant | modifier | 1 |
| | | 102,543 | 102,543 | A/- | -1 | upstream gene variant | modifier | 7 |
| | | 102,544 | 102,544 | C/T | 0 | upstream gene variant | modifier | 7 |
| | | 102,544 | 102,546 | CCG/- | -3 | upstream gene variant | modifier | 75 |
| | | 102,544 | 102,550 | CCGCTCG/- | -7 | upstream gene variant | modifier | 15 |
| | | 102,544 | 102,556 | CCGCTCGCAGAAT/- | -13 | upstream gene variant | modifier | 7 |
| | | 102,545 | 102,544 | -/A | 1 | upstream gene variant | modifier | 120 |
| | | 102,545 | 102,546 | CG/- | -2 | upstream gene variant | modifier | 3 |
| | | 102,545 | 102,546 | CG/TT | 0 | upstream gene variant | modifier | 120 |
| | | 102,545 | 102,548 | CGCT/- | -4 | upstream gene variant | modifier | 140 |
| | | 102,546 | 102,546 | G/T | 0 | upstream gene variant | modifier | 7 |
| | | 102,546 | 102,612 | GCTCGCAGAATCCGCTCCCGGTA CAGAACAGCATCTCTATCCGCCC TCAGAGCTCTAACGTGACGCT/- | -67 | upstream gene variant | modifier | 5 |
| | | 102,547 | 102,546 | -/AAGAATCGCAGAATCCGAGA | 20 | upstream gene variant | modifier | 9 |
| | | 102,547 | 102,546 | -/CAGGGAGAATCTCT | 14 | upstream gene variant | modifier | 9 |
| | | 102,547 | 102,547 | C/A | 0 | upstream gene variant | modifier | 9 |
| | | 102,548 | 102,547 | -/CA | 2 | upstream gene variant | modifier | 40 |
| | | 102,548 | 102,547 | -/ATTCTGT | 7 | upstream gene variant | modifier | 1 |

VEP: Ensembl's variant effect predictor; $n_{affected\ alleles}$: the number of alleles across the 384 sequenced larvae in which the variant was observed (possible range 0 to 768)

**Supplementary Table 15 - Sequencing results expressed in number of mutated alleles for proof-of-concept genes**

| Zebrafish orthologue | Number of affected alleles | | | Missing genotypes | Total | Non-missing | Mutant allele freq | $P_{HWE\_LR}$ |
|---|---|---|---|---|---|---|---|---|
| | 0 | 1 | 2 | | | | | |
| ***apoea*** | 112 | 202 | 66 | 1 | 381 | 380 | 0.439 | 1.45E-01 |
| ***apoeb*** | 37 | 11 | 315 | 18 | 381 | 363 | 0.883 | 2.53E-35 |
| ***apoba*** | 0 | 0 | 377 | 4 | 381 | 377 | 1.000 | - |
| ***apobb.1*** | 149 | 166 | 34 | 32 | 381 | 349 | 0.335 | 1.89E-01 |
| ***apobb.2*** | 0 | 22 | 332 | 27 | 381 | 354 | 0.969 | - |
| ***ldlra*** | 120 | 63 | 197 | 1 | 381 | 380 | 0.601 | 8.60E-40 |
| ***ldlrb*** | 1 | 0 | 327 | 25 | 353 | 328 | 0.997 | - |

The number of affected alleles located in a ±30 base pair window around the CRISPR cut site, without taking into account the variants' probability of affecting protein function. $P_{HWE\_LR}$: *P*- value for a Hardy-Weinberg equilibrium (HWE) likelihood-ratio chi-squared statistic ($P$ <2.9E-3 is significant after Bonferroni correction). For both *apoeb* and *ldlra*, there were more larvae carrying two mutated alleles than expected under HWE.

**Supplementary Table 16 - The effect of a genetic burden score on body size**

| | | Body length (n=339) | | | | |
|---|---|---|---|---|---|---|
| | | **Effect** | **SE** | ***P*** | **lci** | **uci** |
| **fixed factors** | **genetic burden score** | -0.047 | 0.028 | 9.33E-02 | -0.103 | 0.008 |
| | ***apoba*** | 0.022 | 0.231 | 9.24E-01 | -0.431 | 0.475 |
| | ***apobb.2*** | 0.455 | 0.167 | 6.58E-03 | 0.127 | 0.783 |
| | ***ldlrb*** | 2.135 | 1.852 | 2.49E-01 | -1.495 | 5.766 |
| | **time of day (in hours since 9AM)** | -0.022 | 0.037 | 5.54E-01 | -0.095 | 0.051 |
| | **intercept** | -1.439 | 0.914 | 1.15E-01 | -3.230 | 0.351 |
| **random factors** | ***variance by batch*** | *0.469* | *0.129* | - | *0.274* | *0.804* |
| | ***residual*** | *0.724* | *0.028* | - | *0.671* | *0.781* |

| | | Dorsal body surface area (n=339) | | | | |
|---|---|---|---|---|---|---|
| | | **Effect** | **SE** | ***P*** | **lci** | **uci** |
| **fixed factors** | **genetic burden score** | 0.001 | 0.033 | 9.76E-01 | -0.063 | 0.065 |
| | ***apoba*** | 0.165 | 0.267 | 5.36E-01 | -0.358 | 0.689 |
| | ***apobb.2*** | -0.459 | 0.194 | 1.78E-02 | -0.838 | -0.079 |
| | ***ldlrb*** | -2.781 | 2.142 | 1.94E-01 | -6.979 | 1.417 |
| | **time of day (in hours since 9AM)** | 0.028 | 0.043 | 5.21E-01 | -0.057 | 0.112 |
| | **intercept** | 1.335 | 1.054 | 2.05E-01 | -0.731 | 3.400 |
| **random factors** | ***variance by batch*** | *0.503* | *0.140* | - | *0.291* | *0.868* |
| | ***residual*** | *0.837* | *0.033* | - | *0.776* | *0.904* |

| | | Lateral body surface area (n=335) | | | | |
|---|---|---|---|---|---|---|
| | | **Effect** | **SE** | ***P*** | **lci** | **uci** |
| **fixed factors** | **genetic burden score** | 0.005 | 0.034 | 8.79E-01 | -0.061 | 0.071 |
| | ***apoba*** | 0.153 | 0.276 | 5.79E-01 | -0.387 | 0.693 |
| | ***apobb.2*** | -0.221 | 0.198 | 2.63E-01 | -0.609 | 0.166 |
| | ***ldlrb*** | -2.306 | 2.189 | 2.92E-01 | -6.595 | 1.983 |
| | **time of day (in hours since 9AM)** | 0.020 | 0.044 | 6.52E-01 | -0.066 | 0.106 |
| | **intercept** | 0.814 | 1.078 | 4.50E-01 | -1.299 | 2.927 |
| **random factors** | ***variance by batch*** | *0.483* | *0.135* | - | *0.279* | *0.835* |
| | ***residual*** | *0.854* | *0.033* | - | *0.791* | *0.922* |

| | | Body volume (n=328) | | | | |
|---|---|---|---|---|---|---|
| | | **Effect** | **SE** | ***P*** | **lci** | **uci** |
| **fixed factors** | **genetic burden score** | 0.009 | 0.033 | 7.93E-01 | -0.056 | 0.074 |
| | ***apoba*** | 0.152 | 0.270 | 5.75E-01 | -0.378 | 0.682 |
| | ***apobb.2*** | -0.309 | 0.196 | 1.15E-01 | -0.694 | 0.075 |
| | ***ldlrb*** | -2.617 | 2.141 | 2.21E-01 | -6.814 | 1.579 |
| | **time of day (in hours since 9AM)** | 0.041 | 0.044 | 3.46E-01 | -0.045 | 0.128 |
| | **intercept** | 1.000 | 1.060 | 3.46E-01 | -1.078 | 3.078 |
| **random factors** | ***variance by batch*** | *0.494* | *0.137* | - | *0.287* | *0.851* |
| | ***residual*** | *0.834* | *0.033* | - | *0.772* | *0.902* |

A genetic burden score was calculated by summing the dosage scores for *apoea* , *apoeb* , *apobb.1* and *ldlra.* Dorsal and lateral body surface area and body volume were normalized for body length using residuals. All outcomes were inverse-normally transformed before the analysis. Associations were examined using hierarchical linear models and were adjusted for mutations in *apoba* , *apobb.2* and *ldlrb* , time of day and batch. Effects shown for the genetic burden score are for each additional mutated allele. Lci and uci are lower and upper boundaries of the 95% confidence interval.

**Supplementary Table 17 - The effect of a genetic burden score on whole-body lipid and glucose levels**

| | | LDL cholesterol levels (n=381) | | | | |
|---|---|---|---|---|---|---|
| | | **Effect** | **SE** | ***P*** | **lci** | **uci** |
| **fixed factors** | **genetic burden score** | 0.052 | 0.035 | 1.35E-01 | -0.016 | 0.120 |
| | ***apoba*** | 0.019 | 0.290 | 9.47E-01 | -0.549 | 0.588 |
| | ***apobb.2*** | 0.302 | 0.211 | 1.52E-01 | -0.111 | 0.715 |
| | ***ldlrb*** | -3.424 | 2.424 | 1.58E-01 | -8.175 | 1.327 |
| | **time of day (in hours since 9AM)** | 0.055 | 0.043 | 1.97E-01 | -0.029 | 0.139 |
| | **intercept** | 0.595 | 1.167 | 6.10E-01 | -1.693 | 2.883 |
| **random factors** | ***variance by batch*** | *0.355* | *0.123* | - | *0.181* | *0.699* |
| | ***residual*** | *0.951* | *0.035* | - | *0.885* | *1.022* |

| | | HDL cholesterol levels (n=381) | | | | |
|---|---|---|---|---|---|---|
| | | **Effect** | **SE** | ***P*** | **lci** | **uci** |
| **fixed factors** | **genetic burden score** | 0.067 | 0.032 | 3.40E-02 | 0.005 | 0.129 |
| | ***apoba*** | 0.050 | 0.263 | 8.49E-01 | -0.465 | 0.565 |
| | ***apobb.2*** | 0.312 | 0.190 | 1.01E-01 | -0.061 | 0.685 |
| | ***ldlrb*** | 0.841 | 2.192 | 7.01E-01 | -3.455 | 5.136 |
| | **time of day (in hours since 9AM)** | -0.028 | 0.040 | 4.77E-01 | -0.107 | 0.050 |
| | **intercept** | -1.071 | 1.070 | 3.17E-01 | -3.168 | 1.027 |
| **random factors** | ***variance by batch*** | *0.589* | *0.159* | - | *0.346* | *1.001* |
| | ***residual*** | *0.857* | *0.031* | - | *0.798* | *0.921* |

| | | Triglyceride levels (n=381) | | | | |
|---|---|---|---|---|---|---|
| | | **Effect** | **SE** | ***P*** | **lci** | **uci** |
| **fixed factors** | **genetic burden score** | 0.014 | 0.025 | 5.72E-01 | -0.035 | 0.064 |
| | ***apoba*** | -0.098 | 0.210 | 6.43E-01 | -0.510 | 0.315 |
| | ***apobb.2*** | -0.326 | 0.152 | 3.25E-02 | -0.625 | -0.027 |
| | ***ldlrb*** | -0.351 | 1.754 | 8.41E-01 | -3.790 | 3.088 |
| | **time of day (in hours since 9AM)** | -0.146 | 0.032 | 5.92E-06 | -0.210 | -0.083 |
| | **intercept** | 0.987 | 0.885 | 2.64E-01 | -0.747 | 2.721 |
| **random factors** | ***variance by batch*** | *0.781* | *0.200* | - | *0.473* | *1.290* |
| | ***residual*** | *0.686* | *0.025* | - | *0.638* | *0.737* |

| | | Total cholesterol levels (n=381) | | | | |
|---|---|---|---|---|---|---|
| | | **Effect** | **SE** | ***P*** | **lci** | **uci** |
| **fixed factors** | **genetic burden score** | -0.031 | 0.030 | 3.01E-01 | -0.091 | 0.028 |
| | ***apoba*** | 0.165 | 0.252 | 5.13E-01 | -0.329 | 0.659 |
| | ***apobb.2*** | -0.064 | 0.183 | 7.28E-01 | -0.422 | 0.294 |
| | ***ldlrb*** | -0.850 | 2.102 | 6.86E-01 | -4.970 | 3.269 |
| | **time of day (in hours since 9AM)** | 0.083 | 0.039 | 3.24E-02 | 0.007 | 0.158 |
| | **intercept** | 0.109 | 1.042 | 9.17E-01 | -1.933 | 2.150 |
| **random factors** | ***variance by batch*** | *0.754* | *0.199* | - | *0.449* | *1.267* |
| | ***residual*** | *0.822* | *0.030* | - | *0.765* | *0.883* |

***continued*** **Supplementary Table 17**

| | | Glucose levels (n=381) | | | | |
|---|---|---|---|---|---|---|
| | | **Effect** | **SE** | ***P*** | **lci** | **uci** |
| **fixed factors** | **genetic burden score** | -0.047 | 0.035 | 1.74E-01 | -0.116 | 0.021 |
| | ***apoba*** | -0.080 | 0.290 | 7.82E-01 | -0.649 | 0.489 |
| | ***apobb.2*** | -0.364 | 0.211 | 8.44E-02 | -0.777 | 0.049 |
| | ***ldlrb*** | -1.790 | 2.424 | 4.60E-01 | -6.542 | 2.961 |
| | **time of day (in hours since 9AM)** | 0.044 | 0.043 | 3.06E-01 | -0.040 | 0.129 |
| | **intercept** | 1.458 | 1.169 | 2.12E-01 | -0.833 | 3.748 |
| **random factors** | ***variance by batch*** | *0.380* | *0.120* | - | *0.204* | *0.706* |
| | ***residual*** | *0.951* | *0.035* | | *0.885* | *1.022* |

A genetic burden score was calculated by summing the dosage scores for *apoea* , *apoeb* , *apobb.1* and *ldlra.* Dorsal and lateral body surface area and body volume were normalized for body length using residuals. All outcomes were inverse-normally transformed before the analysis. Associations were examined using hierarchical linear models and were adjusted for mutations in *apoba* , *apobb.2* and *ldlrb* , time of day and batch. Effects shown for the genetic burden score are for each additional mutated allele. Lci and uci are lower and upper boundaries of the 95% confidence interval.

**Supplementary Table 18 - The effect of a genetic burden score on vascular atherogenic traits**

| | | Vascular lipid deposition | | | | | | | | | | | | | | |
|---|---|---|---|---|---|---|---|---|---|---|---|---|---|---|---|---|
| | | Model 1 (n=306) | | | | | Model 2 (n=272) | | | | | Model 3 (n=272) | | | | |
| | | Effect | SE | *P* | lci | uci | Effect | SE | *P* | lci | uci | Effect | SE | *P* | lci | uci |
| negative binomial terms | genetic burden score | 0.233 | 0.064 | 2.81E-04 | 0.107 | 0.359 | 0.202 | 0.073 | 5.81E-03 | 0.059 | 0.346 | 0.176 | 0.077 | 2.29E-02 | 0.024 | 0.327 |
| | ***apoba*** | -0.075 | 0.556 | 8.93E-01 | -1.165 | 1.016 | -0.147 | 0.572 | 7.98E-01 | -1.267 | 0.974 | 0.187 | 0.549 | 7.34E-01 | -0.890 | 1.263 |
| | ***apobb.2*** | -0.693 | 0.506 | 1.70E-01 | -1.684 | 0.297 | -0.666 | 0.471 | 1.57E-01 | -1.589 | 0.257 | -0.371 | 0.418 | 3.75E-01 | -1.191 | 0.448 |
| | ***ldlrb*** | 18.109 | 3.317 | 4.76E-08 | 11.608 | 24.609 | 19.711 | 3.363 | 4.61E-09 | 13.119 | 26.303 | 18.757 | 3.209 | 5.08E-09 | 12.466 | 25.047 |
| | **time of day (in hours since 9AM)** | -0.118 | 0.094 | 2.08E-01 | -0.302 | 0.066 | -0.198 | 0.107 | 6.37E-02 | -0.408 | 0.011 | -0.163 | 0.110 | 1.38E-01 | -0.379 | 0.052 |
| | **body length (in SD)** | - | - | - | - | - | -0.285 | 0.144 | 4.72E-02 | -0.567 | -0.004 | -0.207 | 0.153 | 1.77E-01 | -0.508 | 0.094 |
| | **dorsal body surface area (in SD)** | - | - | - | - | - | 0.150 | 0.129 | 2.43E-01 | -0.102 | 0.403 | 0.183 | 0.130 | 1.58E-01 | -0.071 | 0.438 |
| | **LDL cholesterol levels (in SD)** | - | - | - | - | - | - | - | - | - | - | -0.088 | 0.092 | 3.38E-01 | -0.270 | 0.093 |
| | **HDL cholesterol levels (in SD)** | - | - | - | - | - | - | - | - | - | - | -0.035 | 0.129 | 7.87E-01 | -0.287 | 0.218 |
| | **triglyceride levels (in SD)** | - | - | - | - | - | - | - | - | - | - | 0.318 | 0.182 | 8.06E-02 | -0.039 | 0.674 |
| | **glucose levels (in SD)** | - | - | - | - | - | - | - | - | - | - | -0.120 | 0.107 | 2.60E-01 | -0.330 | 0.089 |
| | **batch 1** | 2.442 | 0.485 | 4.84E-07 | 1.491 | 3.394 | 2.791 | 0.557 | 5.35E-07 | 1.700 | 3.882 | 2.808 | 0.611 | 4.29E-06 | 1.611 | 4.005 |
| | **batch 2** | 0.979 | 0.560 | 8.05E-02 | -0.119 | 2.076 | 1.060 | 0.596 | 7.52E-02 | -0.108 | 2.227 | 1.363 | 0.692 | 4.90E-02 | 0.006 | 2.719 |
| | **batch 3** | 1.114 | 0.510 | 2.89E-02 | 0.115 | 2.113 | 1.411 | 0.553 | 1.07E-02 | 0.327 | 2.495 | 1.363 | 0.568 | 1.64E-02 | 0.250 | 2.476 |
| | **batch 4** | 1.555 | 0.585 | 7.87E-03 | 0.408 | 2.702 | 1.548 | 0.583 | 7.97E-03 | 0.404 | 2.691 | 1.627 | 0.584 | 5.32E-03 | 0.483 | 2.771 |
| | **batch 5** | 2.749 | 0.595 | 3.76E-06 | 1.584 | 3.915 | 2.987 | 0.648 | 4.01E-06 | 1.718 | 4.257 | 2.541 | 0.703 | 3.01E-04 | 1.163 | 3.919 |
| | **batch 6** | 2.541 | 0.509 | 5.95E-07 | 1.543 | 3.538 | 3.058 | 0.609 | 5.12E-07 | 1.864 | 4.251 | 2.488 | 0.654 | 1.42E-04 | 1.206 | 3.769 |
| | **batch 7** | 1.695 | 0.564 | 2.64E-03 | 0.590 | 2.799 | 2.100 | 0.689 | 2.30E-03 | 0.750 | 3.450 | 1.791 | 0.697 | 1.02E-02 | 0.425 | 3.157 |
| | **intercept** | -3.897 | 1.815 | 3.18E-02 | -7.454 | -0.339 | -4.338 | 1.729 | 1.21E-02 | -7.728 | -0.949 | -4.795 | 1.685 | 4.44E-03 | -8.098 | -1.492 |

| | | Vascular infiltration by macrophages | | | | | | | | | | | | | | |
|---|---|---|---|---|---|---|---|---|---|---|---|---|---|---|---|---|
| | | Model 1 (n=368) | | | | | Model 2 (n=328) | | | | | Model 3 (n=328) | | | | |
| | | Effect | SE | *P* | lci | uci | Effect | SE | *P* | lci | uci | Effect | SE | *P* | lci | uci |
| fixed factors | **genetic burden score** | -0.076 | 0.032 | 1.76E-02 | -0.139 | -0.013 | -0.087 | 0.035 | 1.22E-02 | -0.155 | -0.019 | -0.099 | 0.035 | 5.04E-03 | -0.168 | -0.030 |
| | ***apoba*** | -0.115 | 0.267 | 6.68E-01 | -0.639 | 0.409 | -0.089 | 0.284 | 7.54E-01 | -0.645 | 0.467 | -0.101 | 0.283 | 7.20E-01 | -0.655 | 0.453 |
| | ***apobb.2*** | -0.264 | 0.194 | 1.73E-01 | -0.645 | 0.116 | -0.371 | 0.210 | 7.71E-02 | -0.782 | 0.040 | -0.427 | 0.212 | 4.38E-02 | -0.842 | -0.012 |
| | ***ldlrb*** | -0.253 | 2.213 | 9.09E-01 | -4.591 | 4.085 | -0.537 | 2.267 | 8.13E-01 | -4.980 | 3.906 | -0.522 | 2.266 | 8.18E-01 | -4.963 | 3.919 |
| | **time of day (in hours since 9AM)** | 0.036 | 0.041 | 3.82E-01 | -0.044 | 0.116 | 0.036 | 0.046 | 4.33E-01 | -0.054 | 0.126 | 0.038 | 0.046 | 4.16E-01 | -0.053 | 0.128 |
| | **body length (in SD)** | - | - | - | - | - | 0.072 | 0.071 | 3.09E-01 | -0.067 | 0.211 | 0.075 | 0.072 | 2.92E-01 | -0.065 | 0.216 |
| | **dorsal body surface area (in SD)** | - | - | - | - | - | -0.049 | 0.061 | 4.22E-01 | -0.167 | 0.070 | -0.046 | 0.061 | 4.47E-01 | -0.166 | 0.073 |
| | **LDL cholesterol levels (in SD)** | - | - | - | - | - | - | - | - | - | - | 0.055 | 0.055 | 3.11E-01 | -0.052 | 0.162 |
| | **HDL cholesterol levels (in SD)** | - | - | - | - | - | - | - | - | - | - | 0.086 | 0.061 | 1.60E-01 | -0.034 | 0.205 |
| | **triglyceride levels (in SD)** | - | - | - | - | - | - | - | - | - | - | 0.021 | 0.074 | 7.80E-01 | -0.124 | 0.165 |
| | **glucose levels (in SD)** | - | - | - | - | - | - | - | - | - | - | -0.040 | 0.054 | 4.62E-01 | -0.146 | 0.066 |
| | **intercept** | 0.932 | 1.078 | 3.87E-01 | -1.181 | 3.045 | 1.214 | 1.119 | 2.78E-01 | -0.980 | 3.408 | 1.358 | 1.119 | 2.25E-01 | -0.835 | 3.551 |
| random factors | ***variance by batch*** | *0.506* | *0.138* | - | *0.296* | *0.865* | *0.510* | *0.144* | - | *0.293* | *0.887* | *0.475* | *0.138* | - | *0.269* | *0.839* |
| | ***residual*** | *0.866* | *0.032* | - | *0.805* | *0.932* | *0.883* | *0.035* | - | *0.817* | *0.954* | *0.879* | *0.035* | - | *0.813* | *0.950* |

***continued*** **Supplementary Table 18**

| | | Vascular co-localization of lipids with macrophages | | | | | | | | | | | | | | |
|---|---|---|---|---|---|---|---|---|---|---|---|---|---|---|---|---|
| | | Model 1 (n=301) | | | | | Model 2 (n=269) | | | | | Model 3 (n=269) | | | | |
| | | Effect | SE | *P* | lci | uci | Effect | SE | *P* | lci | uci | Effect | SE | *P* | lci | uci |
| negative binomial terms | genetic burden score | 0.066 | 0.092 | 4.71E-01 | -0.114 | 0.247 | 0.029 | 0.095 | 7.59E-01 | -0.158 | 0.216 | 0.010 | 0.103 | 9.25E-01 | -0.192 | 0.211 |
| | *apoba* | 0.259 | 0.869 | 7.65E-01 | -1.443 | 1.962 | 0.072 | 0.832 | 9.31E-01 | -1.559 | 1.703 | 0.481 | 0.875 | 5.82E-01 | -1.233 | 2.196 |
| | *apobb.2* | -0.765 | 0.587 | 1.92E-01 | -1.917 | 0.386 | -0.754 | 0.557 | 1.76E-01 | -1.846 | 0.338 | -0.360 | 0.493 | 4.65E-01 | -1.327 | 0.607 |
| | *ldlrb* | 15.180 | 3.761 | 5.43E-05 | 7.809 | 22.551 | 17.068 | 3.698 | 3.93E-06 | 9.819 | 24.316 | 15.483 | 3.640 | 2.10E-05 | 8.350 | 22.616 |
| | time of day (in hours since 9AM) | 0.022 | 0.101 | 8.24E-01 | -0.175 | 0.219 | -0.041 | 0.113 | 7.16E-01 | -0.262 | 0.180 | 0.042 | 0.123 | 7.36E-01 | -0.200 | 0.283 |
| | body length (in SD) | - | - | - | - | - | -0.397 | 0.204 | 5.16E-02 | -0.796 | 0.003 | -0.338 | 0.230 | 1.42E-01 | -0.789 | 0.113 |
| | dorsal body surface area (in SD) | - | - | - | - | - | 0.067 | 0.148 | 6.52E-01 | -0.223 | 0.357 | 0.061 | 0.149 | 6.81E-01 | -0.231 | 0.354 |
| | LDL cholesterol levels (in SD) | - | - | - | - | - | - | - | - | - | - | -0.119 | 0.150 | 4.29E-01 | -0.414 | 0.176 |
| | HDL cholesterol levels (in SD) | - | - | - | - | - | - | - | - | - | - | 0.182 | 0.153 | 2.35E-01 | -0.118 | 0.482 |
| | triglyceride levels (in SD) | - | - | - | - | - | - | - | - | - | - | 0.482 | 0.241 | 4.59E-02 | 0.009 | 0.955 |
| | glucose levels (in SD) | - | - | - | - | - | - | - | - | - | - | -0.128 | 0.139 | 3.56E-01 | -0.400 | 0.144 |
| | batch 1 | 1.324 | 0.750 | 7.74E-02 | -0.146 | 2.793 | 1.525 | 0.835 | 6.76E-02 | -0.110 | 3.161 | 1.415 | 0.877 | 1.07E-01 | -0.303 | 3.133 |
| | batch 2 | 0.132 | 0.926 | 8.87E-01 | -1.683 | 1.946 | 0.012 | 0.969 | 9.90E-01 | -1.887 | 1.912 | 0.018 | 0.994 | 9.86E-01 | -1.931 | 1.967 |
| | batch 3 | 0.273 | 0.753 | 7.17E-01 | -1.203 | 1.748 | 0.252 | 0.781 | 7.46E-01 | -1.278 | 1.782 | 0.152 | 0.771 | 8.44E-01 | -1.360 | 1.663 |
| | batch 4 | 1.378 | 0.834 | 9.84E-02 | -0.256 | 3.011 | 0.986 | 0.849 | 2.46E-01 | -0.679 | 2.650 | 0.851 | 0.818 | 2.98E-01 | -0.752 | 2.454 |
| | batch 5 | 1.816 | 0.868 | 3.65E-02 | 0.114 | 3.517 | 1.766 | 0.917 | 5.43E-02 | -0.033 | 3.564 | 1.203 | 0.969 | 2.14E-01 | -0.696 | 3.102 |
| | batch 6 | 0.740 | 0.789 | 3.48E-01 | -0.806 | 2.286 | 1.264 | 0.933 | 1.76E-01 | -0.565 | 3.094 | 0.589 | 1.029 | 5.67E-01 | -1.427 | 2.606 |
| | batch 7 | -0.383 | 0.915 | 6.76E-01 | -2.176 | 1.411 | -0.144 | 1.066 | 8.92E-01 | -2.234 | 1.945 | -0.449 | 1.121 | 6.89E-01 | -2.645 | 1.748 |
| | intercept | -4.162 | 2.165 | 5.46E-02 | -8.406 | 0.082 | -4.308 | 2.087 | 3.90E-02 | -8.399 | -0.217 | -4.870 | 2.132 | 2.23E-02 | -9.049 | -0.692 |

| | | Vascular infiltration by neutrophils | | | | | | | | | | | | | | |
|---|---|---|---|---|---|---|---|---|---|---|---|---|---|---|---|---|
| | | Model 1 (n=371) | | | | | Model 2 (n=330) | | | | | Model 3 (n=330) | | | | |
| | | Effect | SE | *P* | lci | uci | Effect | SE | *P* | lci | uci | Effect | SE | *P* | lci | uci |
| fixed factors | genetic burden score | -0.018 | 0.033 | 5.78E-01 | -0.082 | 0.046 | -0.026 | 0.035 | 4.57E-01 | -0.094 | 0.042 | -0.029 | 0.035 | 4.10E-01 | -0.098 | 0.040 |
| | *apoba* | 0.096 | 0.272 | 7.24E-01 | -0.436 | 0.628 | 0.144 | 0.284 | 6.11E-01 | -0.412 | 0.701 | 0.140 | 0.283 | 6.20E-01 | -0.414 | 0.695 |
| | *apobb.2* | -0.084 | 0.198 | 6.71E-01 | -0.472 | 0.304 | 0.016 | 0.211 | 9.40E-01 | -0.397 | 0.429 | 0.003 | 0.213 | 9.90E-01 | -0.414 | 0.420 |
| | *ldlrb* | -2.023 | 2.259 | 3.71E-01 | -6.451 | 2.405 | -1.483 | 2.282 | 5.16E-01 | -5.955 | 2.990 | -1.459 | 2.286 | 5.23E-01 | -5.940 | 3.021 |
| | time of day (in hours since 9AM) | 0.012 | 0.041 | 7.65E-01 | -0.067 | 0.092 | -0.020 | 0.045 | 6.55E-01 | -0.107 | 0.067 | -0.022 | 0.045 | 6.20E-01 | -0.110 | 0.065 |
| | body length (in SD) | - | - | - | - | - | -0.122 | 0.070 | 7.94E-02 | -0.258 | 0.014 | -0.114 | 0.070 | 1.04E-01 | -0.252 | 0.023 |
| | dorsal body surface area (in SD) | - | - | - | - | - | 0.060 | 0.059 | 3.12E-01 | -0.056 | 0.176 | 0.075 | 0.060 | 2.11E-01 | -0.043 | 0.193 |
| | LDL cholesterol levels (in SD) | - | - | - | - | - | - | - | - | - | - | 0.002 | 0.053 | 9.65E-01 | -0.103 | 0.107 |
| | HDL cholesterol levels (in SD) | - | - | - | - | - | - | - | - | - | - | 0.081 | 0.060 | 1.79E-01 | -0.037 | 0.200 |
| | triglyceride levels (in SD) | - | - | - | - | - | - | - | - | - | - | -0.042 | 0.071 | 5.50E-01 | -0.181 | 0.097 |
| | glucose levels (in SD) | - | - | - | - | - | - | - | - | - | - | 0.060 | 0.054 | 2.71E-01 | -0.046 | 0.166 |
| | intercept | 0.845 | 1.091 | 4.38E-01 | -1.293 | 2.983 | 0.536 | 1.116 | 6.31E-01 | -1.651 | 2.723 | 0.570 | 1.117 | 6.10E-01 | -1.619 | 2.759 |
| random factors | *variance by batch* | *0.356* | *0.101* | - | *0.204* | *0.622* | *0.323* | *0.097* | - | *0.179* | *0.582* | *0.297* | *0.092* | - | *0.162* | *0.545* |
| | *residual* | *0.885* | *0.033* | - | *0.823* | *0.952* | *0.890* | *0.035* | - | *0.824* | *0.962* | *0.888* | *0.035* | - | *0.822* | *0.959* |

***continued*** **Supplementary Table 18**

| | | Vascular co-localization of lipids with neutrophils | | | | | | | | | | | | | | |
|---|---|---|---|---|---|---|---|---|---|---|---|---|---|---|---|---|
| | | Model 1 (n=304) | | | | | Model 2 (n=271) | | | | | Model 3 (n=271) | | | | |
| | | **Effect** | **SE** | ***P*** | **lci** | **uci** | **Effect** | **SE** | ***P*** | **lci** | **uci** | **Effect** | **SE** | ***P*** | **lci** | **uci** |
| **negative binomial terms** | **genetic burden score** | 0.454 | 0.100 | 5.63E-06 | 0.258 | 0.650 | 0.383 | 0.122 | 1.73E-03 | 0.144 | 0.623 | 0.276 | 0.143 | 5.37E-02 | -0.004 | 0.556 |
| | ***apoba*** | 0.301 | 1.127 | 7.90E-01 | -1.908 | 2.510 | 0.098 | 1.173 | 9.33E-01 | -2.201 | 2.397 | 1.046 | 1.072 | 3.29E-01 | -1.055 | 3.147 |
| | ***apobb.2*** | 0.231 | 0.553 | 6.76E-01 | -0.854 | 1.316 | 0.185 | 0.608 | 7.61E-01 | -1.006 | 1.376 | 0.909 | 0.674 | 1.78E-01 | -0.412 | 2.230 |
| | ***ldlrb*** | 9.320 | 3.741 | 1.27E-02 | 1.987 | 16.653 | 10.263 | 3.782 | 6.66E-03 | 2.850 | 17.677 | 11.151 | 3.863 | 3.89E-03 | 3.580 | 18.722 |
| | **time of day (in hours since 9AM)** | -0.074 | 0.165 | 6.54E-01 | -0.398 | 0.250 | -0.176 | 0.187 | 3.46E-01 | -0.542 | 0.190 | -0.198 | 0.169 | 2.43E-01 | -0.529 | 0.134 |
| | **body length (in SD)** | - | - | - | - | - | -0.513 | 0.274 | 6.16E-02 | -1.051 | 0.025 | -0.632 | 0.269 | 1.88E-02 | -1.159 | -0.105 |
| | **dorsal body surface area (in SD)** | - | - | - | - | - | -0.004 | 0.224 | 9.87E-01 | -0.443 | 0.436 | -0.074 | 0.208 | 7.23E-01 | -0.482 | 0.334 |
| | **LDL cholesterol levels (in SD)** | - | - | - | - | - | - | - | - | - | - | 0.540 | 0.255 | 3.42E-02 | 0.040 | 1.040 |
| | **HDL cholesterol levels (in SD)** | - | - | - | - | - | - | - | - | - | - | 0.627 | 0.304 | 3.93E-02 | 0.031 | 1.224 |
| | **triglyceride levels (in SD)** | - | - | - | - | - | - | - | - | - | - | 1.083 | 0.290 | 1.93E-04 | 0.513 | 1.652 |
| | **glucose levels (in SD)** | - | - | - | - | - | - | - | - | - | - | 0.095 | 0.204 | 6.40E-01 | -0.304 | 0.495 |
| | **batch 1** | 3.896 | 0.963 | 5.19E-05 | 2.009 | 5.782 | 4.110 | 1.000 | 3.93E-05 | 2.151 | 6.069 | 5.318 | 1.174 | 5.95E-06 | 3.016 | 7.620 |
| | **batch 2** | 1.723 | 1.095 | 1.16E-01 | -0.423 | 3.870 | 1.735 | 1.136 | 1.27E-01 | -0.492 | 3.961 | 4.159 | 1.448 | 4.08E-03 | 1.320 | 6.997 |
| | **batch 3** | 2.330 | 1.086 | 3.19E-02 | 0.201 | 4.459 | 2.510 | 1.103 | 2.29E-02 | 0.348 | 4.672 | 3.085 | 1.143 | 6.97E-03 | 0.844 | 5.327 |
| | **batch 4** | 3.533 | 1.056 | 8.17E-04 | 1.464 | 5.603 | 3.288 | 1.086 | 2.47E-03 | 1.159 | 5.416 | 4.296 | 1.067 | 5.67E-05 | 2.205 | 6.388 |
| | **batch 5** | 4.590 | 1.077 | 2.01E-05 | 2.480 | 6.700 | 4.720 | 1.103 | 1.86E-05 | 2.559 | 6.881 | 4.782 | 1.145 | 2.98E-05 | 2.537 | 7.027 |
| | **batch 6** | 3.473 | 1.027 | 7.17E-04 | 1.461 | 5.485 | 3.780 | 1.064 | 3.79E-04 | 1.695 | 5.865 | 4.115 | 1.142 | 3.15E-04 | 1.876 | 6.353 |
| | **batch 7** | 1.922 | 1.340 | 1.51E-01 | -0.704 | 4.549 | 2.988 | 1.521 | 4.95E-02 | 0.007 | 5.969 | 3.253 | 1.429 | 2.29E-02 | 0.451 | 6.054 |
| | **intercept** | -8.759 | 2.629 | 8.62E-04 | -13.911 | -3.607 | -8.322 | 2.696 | 2.03E-03 | -13.606 | -3.037 | -11.768 | 2.659 | 9.61E-06 | -16.980 | -6.557 |

| | | Vascular co-localization of macrophages with neutrophils | | | | | | | | | | | | | | |
|---|---|---|---|---|---|---|---|---|---|---|---|---|---|---|---|---|
| | | Model 1 (n=367) | | | | | Model 2 (n=327) | | | | | Model 3 (n=327) | | | | |
| | | **Effect** | **SE** | ***P*** | **lci** | **uci** | **Effect** | **SE** | ***P*** | **lci** | **uci** | **Effect** | **SE** | ***P*** | **lci** | **uci** |
| **negative binomial terms** | **genetic burden score** | -0.034 | 0.051 | 5.09E-01 | -0.135 | 0.067 | -0.038 | 0.054 | 4.75E-01 | -0.143 | 0.067 | -0.031 | 0.054 | 5.64E-01 | -0.137 | 0.075 |
| | ***apoba*** | 0.113 | 0.404 | 7.80E-01 | -0.679 | 0.904 | 0.332 | 0.399 | 4.05E-01 | -0.450 | 1.115 | 0.379 | 0.398 | 3.41E-01 | -0.401 | 1.159 |
| | ***apobb.2*** | -0.264 | 0.319 | 4.07E-01 | -0.888 | 0.360 | -0.414 | 0.358 | 2.47E-01 | -1.116 | 0.287 | -0.319 | 0.341 | 3.50E-01 | -0.988 | 0.350 |
| | ***ldlrb*** | 0.251 | 0.678 | 7.11E-01 | -1.077 | 1.580 | 0.266 | 0.795 | 7.38E-01 | -1.293 | 1.824 | 0.856 | 0.918 | 3.51E-01 | -0.944 | 2.656 |
| | **time of day (in hours since 9AM)** | 0.069 | 0.067 | 2.98E-01 | -0.061 | 0.200 | 0.085 | 0.071 | 2.29E-01 | -0.053 | 0.223 | 0.069 | 0.074 | 3.50E-01 | -0.076 | 0.214 |
| | **body length (in SD)** | - | - | - | - | - | 0.304 | 0.112 | 6.88E-03 | 0.083 | 0.524 | 0.252 | 0.121 | 3.66E-02 | 0.016 | 0.489 |
| | **dorsal body surface area (in SD)** | - | - | - | - | - | 0.261 | 0.110 | 1.83E-02 | 0.044 | 0.477 | 0.253 | 0.116 | 2.90E-02 | 0.026 | 0.479 |
| | **LDL cholesterol levels (in SD)** | - | - | - | - | - | - | - | - | - | - | 0.086 | 0.094 | 3.62E-01 | -0.098 | 0.270 |
| | **HDL cholesterol levels (in SD)** | - | - | - | - | - | - | - | - | - | - | -0.035 | 0.102 | 7.33E-01 | -0.235 | 0.165 |
| | **triglyceride levels (in SD)** | - | - | - | - | - | - | - | - | - | - | -0.037 | 0.127 | 7.72E-01 | -0.285 | 0.211 |
| | **glucose levels (in SD)** | - | - | - | - | - | - | - | - | - | - | 0.106 | 0.083 | 2.04E-01 | -0.058 | 0.270 |
| | **batch 1** | 0.408 | 0.310 | 1.87E-01 | -0.199 | 1.015 | 0.202 | 0.333 | 5.44E-01 | -0.450 | 0.854 | 0.379 | 0.391 | 3.33E-01 | -0.388 | 1.147 |
| | **batch 2** | 0.285 | 0.518 | 5.82E-01 | -0.730 | 1.301 | 0.309 | 0.482 | 5.21E-01 | -0.636 | 1.254 | 0.488 | 0.498 | 3.27E-01 | -0.488 | 1.464 |
| | **batch 3** | -0.610 | 0.376 | 1.04E-01 | -1.346 | 0.126 | -0.727 | 0.373 | 5.13E-02 | -1.459 | 0.004 | -0.553 | 0.403 | 1.69E-01 | -1.343 | 0.236 |
| | **batch 4** | 0.938 | 0.390 | 1.60E-02 | 0.175 | 1.702 | 1.045 | 0.410 | 1.07E-02 | 0.242 | 1.849 | 1.091 | 0.443 | 1.39E-02 | 0.222 | 1.960 |
| | **batch 5** | 0.216 | 0.351 | 5.38E-01 | -0.472 | 0.903 | -0.098 | 0.362 | 7.88E-01 | -0.807 | 0.612 | 0.113 | 0.419 | 7.88E-01 | -0.709 | 0.935 |
| | **batch 6** | -1.562 | 0.358 | 1.27E-05 | -2.263 | -0.861 | -2.156 | 0.413 | 1.80E-07 | -2.966 | -1.346 | -2.009 | 0.434 | 3.73E-06 | -2.860 | -1.158 |
| | **batch 7** | -0.454 | 0.460 | 3.24E-01 | -1.355 | 0.448 | -1.079 | 0.517 | 3.67E-02 | -2.092 | -0.067 | -0.986 | 0.599 | 9.96E-02 | -2.160 | 0.188 |
| | **intercept** | 3.911 | 0.862 | 5.66E-06 | 2.222 | 5.600 | 3.929 | 0.927 | 2.24E-05 | 2.113 | 5.746 | 3.325 | 0.975 | 6.47E-04 | 1.415 | 5.236 |

A genetic burden score was calculated by summing the dosage scores for *apoea* , *apoeb* , *apobb.1* and *ldlra* . Dorsal and lateral body surface area and body volume were normalized for body length using residuals. All outcomes were inverse-normally transformed before the analysis. Associations were examined using hierarchical linear models and were adjusted for mutations in *apoba* , *apobb.2* and *ldlrb* , time of day and batch. Effects shown for the genetic burden score are for each additional mutated allele. Lci and uci are lower and upper boundaries of the 95% confidence interval.

**Supplementary Table 19 - The effect of two vs. zero mutated alleles in apoea, apoeb, apobb.1 or ldlra on body size**

| | | **Body length** | | | | | | | | | | | | | | | | | | | |
|---|---|---|---|---|---|---|---|---|---|---|---|---|---|---|---|---|---|---|---|---|---|
| | | ***apoea*** | | | | | ***apoeb*** | | | | | ***apobb.1*** | | | | | ***ldlra*** | | | | |
| | | **44 vs. 96 larvae with 2 vs 0 mutated alleles** | | | | | **212 vs. 34 larvae with 2 vs 0 mutated alleles** | | | | | **30 vs. 130 larvae with 2 vs 0 mutated alleles** | | | | | **165 vs. 105 larvae with 2 vs 0 mutated** | | | | |
| | | **Effect** | **SE** | ***P*** | **lci** | **uci** | **Effect** | **SE** | ***P*** | **lci** | **uci** | **Effect** | **SE** | ***P*** | **lci** | **uci** | **Effect** | **SE** | ***P*** | **lci** | **uci** |
| | **2 vs. 0 mutated alleles** | 0.057 | 0.146 | 6.95E-01 | -0.229 | 0.343 | -0.086 | 0.157 | 5.84E-01 | -0.394 | 0.222 | -1.071 | 0.149 | 7.19E-13 | -1.364 | -0.779 | 0.155 | 0.114 | 1.73E-01 | -0.068 | 0.379 |
| | ***apoea*** | - | - | - | - | - | 0.036 | 0.074 | 6.27E-01 | -0.109 | 0.181 | -0.013 | 0.089 | 8.80E-01 | -0.187 | 0.160 | -0.002 | 0.071 | 9.77E-01 | -0.141 | 0.137 |
| | ***apoeb*** | -0.035 | 0.107 | 7.40E-01 | -0.244 | 0.174 | - | - | - | - | - | 0.012 | 0.084 | 8.84E-01 | -0.152 | 0.177 | 0.014 | 0.075 | 8.49E-01 | -0.133 | 0.162 |
| | ***apobb.1*** | -0.460 | 0.096 | 1.67E-06 | -0.648 | -0.272 | -0.221 | 0.081 | 6.38E-03 | -0.380 | -0.062 | - | - | - | - | - | -0.397 | 0.075 | 1.19E-07 | -0.544 | -0.250 |
| **fixed** | ***ldlra*** | 0.044 | 0.078 | 5.69E-01 | -0.108 | 0.197 | 0.022 | 0.065 | 7.36E-01 | -0.106 | 0.150 | 0.081 | 0.080 | 3.08E-01 | -0.075 | 0.238 | - | - | - | - | - |
| **factors** | **time of day (in hours since 9AM)** | 0.006 | 0.054 | 9.15E-01 | -0.101 | 0.112 | -0.026 | 0.044 | 5.56E-01 | -0.111 | 0.060 | 0.077 | 0.047 | 1.02E-01 | -0.015 | 0.168 | -0.057 | 0.040 | 1.58E-01 | -0.136 | 0.022 |
| | **intercept** | -4.855 | 59.467 | 9.35E-01 | -121.409 | 111.699 | 15.036 | 51.182 | 7.69E-01 | -85.278 | 115.351 | 104.827 | 63.695 | 9.98E-02 | -20.013 | 229.667 | -1.214 | 0.932 | 1.93E-01 | -3.041 | 0.614 |
| | ***apoba*** | 0.041 | 0.370 | 9.13E-01 | -0.685 | 0.766 | 0.266 | 0.291 | 3.60E-01 | -0.304 | 0.835 | -0.408 | 0.320 | 2.02E-01 | -1.035 | 0.219 | -0.046 | 0.278 | 8.69E-01 | -0.590 | 0.499 |
| | ***apobb.2*** | 0.404 | 0.247 | 1.02E-01 | -0.081 | 0.889 | 0.423 | 0.183 | 2.12E-02 | 0.063 | 0.782 | 0.526 | 0.196 | 7.37E-03 | 0.141 | 0.911 | 0.359 | 0.178 | 4.39E-02 | 0.010 | 0.708 |
| | ***ldlrb*** | 10.731 | 148.645 | 9.42E-01 | -280.607 | 302.069 | -40.149 | 128.002 | 7.54E-01 | -291.028 | 210.729 | -263.194 | 159.306 | 9.85E-02 | -575.428 | 49.041 | 2.336 | 1.790 | 1.92E-01 | -1.172 | 5.844 |
| **random** | ***variance by batch*** | *0.445* | *0.136* | - | *0.245* | *0.810* | *0.508* | *0.142* | - | *0.294* | *0.879* | *0.454* | *0.136* | - | *0.252* | *0.815* | *0.472* | *0.131* | - | *0.274* | *0.812* |
| **factors** | ***residual*** | *0.666* | *0.041* | - | *0.590* | *0.752* | *0.695* | *0.032* | - | *0.635* | *0.760* | *0.633* | *0.036* | - | *0.566* | *0.709* | *0.694* | *0.030* | - | *0.637* | *0.756* |

| | | **Dorsal body surface area** | | | | | | | | | | | | | | | | | | | |
|---|---|---|---|---|---|---|---|---|---|---|---|---|---|---|---|---|---|---|---|---|---|
| | | ***apoea*** | | | | | ***apoeb*** | | | | | ***apobb.1*** | | | | | ***ldlra*** | | | | |
| | | **44 vs. 96 larvae with 2 vs 0 mutated alleles** | | | | | **212 vs. 34 larvae with 2 vs 0 mutated alleles** | | | | | **30 vs. 130 larvae with 2 vs 0 mutated alleles** | | | | | **165 vs. 105 larvae with 2 vs 0 mutated** | | | | |
| | | **Effect** | **SE** | ***P*** | **lci** | **uci** | **Effect** | **SE** | ***P*** | **lci** | **uci** | **Effect** | **SE** | ***P*** | **lci** | **uci** | **Effect** | **SE** | ***P*** | **lci** | **uci** |
| | **2 vs. 0 mutated alleles** | -0.138 | 0.162 | 3.93E-01 | -0.456 | 0.179 | 0.261 | 0.181 | 1.50E-01 | -0.094 | 0.615 | 0.215 | 0.190 | 2.58E-01 | -0.158 | 0.588 | -0.166 | 0.139 | 2.32E-01 | -0.439 | 0.106 |
| | ***apoea*** | - | - | - | - | - | -0.126 | 0.085 | 1.39E-01 | -0.293 | 0.041 | -0.257 | 0.112 | 2.20E-02 | -0.478 | -0.037 | -0.117 | 0.087 | 1.77E-01 | -0.287 | 0.053 |
| | ***apoeb*** | -0.015 | 0.119 | 8.98E-01 | -0.248 | 0.218 | - | - | - | - | - | -0.006 | 0.107 | 9.56E-01 | -0.216 | 0.204 | 0.064 | 0.092 | 4.87E-01 | -0.116 | 0.244 |
| | ***apobb.1*** | 0.287 | 0.107 | 7.43E-03 | 0.077 | 0.498 | 0.037 | 0.093 | 6.96E-01 | -0.147 | 0.220 | - | - | - | - | - | 0.137 | 0.092 | 1.34E-01 | -0.042 | 0.317 |
| **fixed** | ***ldlra*** | -0.078 | 0.087 | 3.72E-01 | -0.249 | 0.093 | -0.081 | 0.075 | 2.80E-01 | -0.229 | 0.066 | 0.048 | 0.101 | 6.36E-01 | -0.151 | 0.247 | - | - | - | - | - |
| **factors** | **time of day (in hours since 9AM)** | 0.106 | 0.059 | 7.53E-02 | -0.011 | 0.222 | 0.044 | 0.050 | 3.79E-01 | -0.053 | 0.141 | 0.000 | 0.058 | 9.95E-01 | -0.114 | 0.115 | 0.066 | 0.049 | 1.76E-01 | -0.029 | 0.161 |
| | **intercept** | 13.639 | 66.393 | 8.37E-01 | -116.489 | 143.766 | 73.698 | 58.862 | 2.11E-01 | -41.669 | 189.065 | -102.710 | 81.214 | 2.06E-01 | -261.885 | 56.466 | 0.913 | 1.132 | 4.20E-01 | -1.306 | 3.131 |
| | ***apoba*** | 0.509 | 0.414 | 2.19E-01 | -0.302 | 1.321 | -0.089 | 0.334 | 7.90E-01 | -0.745 | 0.567 | -0.083 | 0.406 | 8.38E-01 | -0.880 | 0.714 | 0.273 | 0.339 | 4.21E-01 | -0.392 | 0.938 |
| | ***apobb.2*** | -0.305 | 0.277 | 2.70E-01 | -0.847 | 0.237 | -0.372 | 0.211 | 7.87E-02 | -0.786 | 0.043 | -0.542 | 0.250 | 3.04E-02 | -1.032 | -0.051 | -0.400 | 0.218 | 6.65E-02 | -0.826 | 0.027 |
| | ***ldlrb*** | -36.318 | 165.957 | 8.27E-01 | -361.589 | 288.952 | -183.074 | 147.212 | 2.14E-01 | -471.605 | 105.458 | 259.291 | 203.127 | 2.02E-01 | -138.832 | 657.413 | -2.739 | 2.185 | 2.10E-01 | -7.021 | 1.544 |
| **random** | ***variance by batch*** | *0.403* | *0.146* | - | *0.199* | *0.818* | *0.451* | *0.135* | - | *0.251* | *0.812* | *0.425* | *0.133* | - | *0.230* | *0.783* | *0.453* | *0.131* | - | *0.256* | *0.800* |
| **factors** | ***residual*** | *0.745* | *0.046* | - | *0.660* | *0.842* | *0.801* | *0.037* | - | *0.732* | *0.877* | *0.808* | *0.046* | - | *0.722* | *0.905* | *0.849* | *0.037* | - | *0.779* | *0.925* |

| | | **Lateral body surface area** | | | | | | | | | | | | | | | | | | | |
|---|---|---|---|---|---|---|---|---|---|---|---|---|---|---|---|---|---|---|---|---|---|
| | | ***apoea*** | | | | | ***apoeb*** | | | | | ***apobb.1*** | | | | | ***ldlra*** | | | | |
| | | **43 vs. 95 larvae with 2 vs 0 mutated alleles** | | | | | **211 vs. 33 larvae with 2 vs 0 mutated alleles** | | | | | **30 vs. 128 larvae with 2 vs 0 mutated alleles** | | | | | **164 vs. 103 larvae with 2 vs 0 mutated** | | | | |
| | | **Effect** | **SE** | ***P*** | **lci** | **uci** | **Effect** | **SE** | ***P*** | **lci** | **uci** | **Effect** | **SE** | ***P*** | **lci** | **uci** | **Effect** | **SE** | ***P*** | **lci** | **uci** |
| | **2 vs. 0 mutated alleles** | -0.260 | 0.167 | 1.20E-01 | -0.587 | 0.068 | 0.337 | 0.193 | 8.04E-02 | -0.041 | 0.714 | 0.041 | 0.195 | 8.35E-01 | -0.341 | 0.422 | -0.027 | 0.141 | 8.46E-01 | -0.305 | 0.250 |
| | ***apoea*** | - | - | - | - | - | -0.150 | 0.089 | 9.23E-02 | -0.324 | 0.025 | -0.343 | 0.116 | 3.02E-03 | -0.570 | -0.116 | -0.143 | 0.088 | 1.02E-01 | -0.315 | 0.029 |
| | ***apoeb*** | 0.091 | 0.123 | 4.63E-01 | -0.151 | 0.332 | - | - | - | - | - | 0.037 | 0.110 | 7.35E-01 | -0.178 | 0.252 | 0.128 | 0.094 | 1.72E-01 | -0.056 | 0.313 |
| | ***apobb.1*** | 0.259 | 0.112 | 2.05E-02 | 0.040 | 0.479 | 0.000 | 0.098 | 9.99E-01 | -0.191 | 0.192 | - | - | - | - | - | 0.041 | 0.093 | 6.58E-01 | -0.141 | 0.223 |
| **fixed** | ***ldlra*** | -0.020 | 0.091 | 8.25E-01 | -0.199 | 0.159 | -0.058 | 0.080 | 4.64E-01 | -0.214 | 0.098 | 0.054 | 0.105 | 6.08E-01 | -0.152 | 0.259 | - | - | - | - | - |
| **factors** | **time of day (in hours since 9AM)** | 0.072 | 0.061 | 2.40E-01 | -0.048 | 0.191 | 0.067 | 0.052 | 1.95E-01 | -0.034 | 0.168 | -0.017 | 0.060 | 7.73E-01 | -0.135 | 0.100 | 0.034 | 0.049 | 4.92E-01 | -0.062 | 0.130 |
| | **intercept** | 2.580 | 68.557 | 9.70E-01 | -131.789 | 136.950 | 66.633 | 61.642 | 2.80E-01 | -54.183 | 187.448 | -95.130 | 82.967 | 2.52E-01 | -257.743 | 67.483 | 0.881 | 1.146 | 4.42E-01 | -1.364 | 3.126 |
| | ***apoba*** | 0.228 | 0.430 | 5.97E-01 | -0.616 | 1.071 | -0.071 | 0.355 | 8.42E-01 | -0.766 | 0.624 | -0.171 | 0.418 | 6.82E-01 | -0.989 | 0.647 | -0.040 | 0.345 | 9.08E-01 | -0.717 | 0.637 |
| | ***apobb.2*** | 0.181 | 0.286 | 5.27E-01 | -0.380 | 0.742 | -0.066 | 0.221 | 7.64E-01 | -0.498 | 0.366 | -0.345 | 0.256 | 1.77E-01 | -0.847 | 0.156 | -0.198 | 0.220 | 3.67E-01 | -0.629 | 0.232 |
| | ***ldlrb*** | -9.404 | 171.364 | 9.56E-01 | -345.272 | 326.464 | -166.932 | 154.152 | 2.79E-01 | -469.064 | 135.199 | 240.229 | 207.510 | 2.47E-01 | -166.484 | 646.941 | -1.960 | 2.207 | 3.74E-01 | -6.287 | 2.366 |
| **random** | ***variance by batch*** | *0.382* | *0.141* | - | *0.185* | *0.787* | *0.455* | *0.135* | - | *0.254* | *0.812* | *0.432* | *0.137* | - | *0.231* | *0.805* | *0.442* | *0.127* | - | *0.252* | *0.776* |
| **factors** | ***residual*** | *0.770* | *0.048* | - | *0.681* | *0.870* | *0.836* | *0.039* | - | *0.764* | *0.915* | *0.826* | *0.048* | - | *0.737* | *0.925* | *0.855* | *0.038* | - | *0.784* | *0.932* |

***continued*** **Supplementary Table 19**

| | | **Body volume** | | | | | | | | | | | | | | | | | | | |
|---|---|---|---|---|---|---|---|---|---|---|---|---|---|---|---|---|---|---|---|---|---|
| | | ***apoea*** | | | | | ***apoeb*** | | | | | ***apobb.1*** | | | | | ***ldlra*** | | | | |
| | | **43 vs. 90 larvae with 2 vs 0 mutated alleles** | | | | | **206 vs. 32 larvae with 2 vs 0 mutated alleles** | | | | | **29 vs. 125 larvae with 2 vs 0 mutated alleles** | | | | | **161 vs. 100 larvae with 2 vs 0 mutated** | | | | |
| | | **Effect** | **SE** | ***P*** | **lci** | **uci** | **Effect** | **SE** | ***P*** | **lci** | **uci** | **Effect** | **SE** | ***P*** | **lci** | **uci** | **Effect** | **SE** | ***P*** | **lci** | **uci** |
| **fixed factors** | **2 vs. 0 mutated alleles** | -0.209 | 0.161 | 1.94E-01 | -0.525 | 0.107 | 0.352 | 0.191 | 6.49E-02 | -0.022 | 0.726 | 0.074 | 0.193 | 7.02E-01 | -0.304 | 0.452 | -0.086 | 0.142 | 5.44E-01 | -0.365 | 0.193 |
| | ***apoea*** | - | - | - | - | - | -0.126 | 0.089 | 1.56E-01 | -0.299 | 0.048 | -0.288 | 0.115 | 1.24E-02 | -0.514 | -0.062 | -0.108 | 0.089 | 2.24E-01 | -0.281 | 0.066 |
| | ***apoeb*** | 0.058 | 0.121 | 6.29E-01 | -0.178 | 0.295 | - | - | - | - | - | 0.042 | 0.107 | 6.93E-01 | -0.168 | 0.253 | 0.111 | 0.095 | 2.41E-01 | -0.075 | 0.298 |
| | ***apobb.1*** | 0.282 | 0.110 | 1.02E-02 | 0.067 | 0.497 | 0.009 | 0.097 | 9.23E-01 | -0.182 | 0.200 | - | - | - | - | - | 0.063 | 0.094 | 5.07E-01 | -0.122 | 0.247 |
| | ***ldlra*** | -0.014 | 0.088 | 8.72E-01 | -0.188 | 0.159 | -0.059 | 0.078 | 4.47E-01 | -0.213 | 0.094 | 0.086 | 0.104 | 4.10E-01 | -0.118 | 0.289 | - | - | - | - | - |
| | **time of day (in hours since 9AM)** | 0.123 | 0.060 | 4.13E-02 | 0.005 | 0.242 | 0.078 | 0.052 | 1.30E-01 | -0.023 | 0.180 | 0.006 | 0.060 | 9.25E-01 | -0.112 | 0.124 | 0.062 | 0.050 | 2.16E-01 | -0.036 | 0.159 |
| | **intercept** | 13.835 | 65.406 | 8.32E-01 | -114.357 | 142.028 | 56.403 | 61.175 | 3.57E-01 | -63.497 | 176.303 | -92.716 | 81.151 | 2.53E-01 | -251.770 | 66.338 | 0.730 | 1.147 | 5.25E-01 | -1.518 | 2.978 |
| | ***apoba*** | 0.332 | 0.411 | 4.19E-01 | -0.474 | 1.138 | -0.167 | 0.346 | 6.30E-01 | -0.845 | 0.512 | -0.234 | 0.409 | 5.68E-01 | -1.034 | 0.567 | 0.167 | 0.346 | 6.30E-01 | -0.511 | 0.844 |
| | ***apobb.2*** | 0.022 | 0.285 | 9.39E-01 | -0.536 | 0.579 | -0.182 | 0.219 | 4.06E-01 | -0.611 | 0.247 | -0.424 | 0.253 | 9.33E-02 | -0.920 | 0.071 | -0.269 | 0.223 | 2.27E-01 | -0.707 | 0.168 |
| | ***ldlrb*** | -37.762 | 163.498 | 8.17E-01 | -358.212 | 282.688 | -140.614 | 152.985 | 3.58E-01 | -440.459 | 159.231 | 234.401 | 202.978 | 2.48E-01 | -163.428 | 632.231 | -2.413 | 2.199 | 2.73E-01 | -6.723 | 1.898 |
| **random factors** | ***variance by batch*** | *0.368* | *0.134* | - | *0.181* | *0.750* | *0.449* | *0.134* | - | *0.251* | *0.805* | *0.434* | *0.134* | - | *0.237* | *0.796* | *0.468* | *0.133* | - | *0.269* | *0.817* |
| | ***residual*** | *0.732* | *0.047* | - | *0.646* | *0.829* | *0.814* | *0.038* | - | *0.742* | *0.892* | *0.804* | *0.047* | - | *0.717* | *0.902* | *0.850* | *0.038* | - | *0.779* | *0.927* |

Dorsal and lateral body surface area and body volume were normalized for body length using residuals. All outcomes were inverse-normally transformed before the analysis and examined using hierarchical linear models. Effects shown are for larvae with two mutated alleles that are highly likely to affect protein function as predicted by Ensembl's Varient Effect Predictor (VEP) compared with larvae with zero CRISPR-mutated alleles. Associations were adjusted for the number of mutated alleles in the other six orthologues, weighted by their predicted effect on protein function, as well as for time of day and batch. Lci and uci are lower and upper boundaries of the 95% confidence interval.

**Supplementary Table 20 - The effect of two vs. zero mutated alleles in *apoea* , *apoeb* , *apobb.1* or *ldlra* on whole-body lipid and glucose levels**

| | | LDL cholesterol levels | | | | | | | | | | | | | | | | | | | |
|---|---|---|---|---|---|---|---|---|---|---|---|---|---|---|---|---|---|---|---|---|---|
| | | ***apoea*** | | | | | ***apoeb*** | | | | | ***apobb.1*** | | | | | ***ldlra*** | | | | |
| | | 44 vs. 96 larvae with 2 vs 0 mutated alleles | | | | | 212 vs. 34 larvae with 2 vs 0 mutated alleles | | | | | 30 vs. 130 larvae with 2 vs 0 mutated alleles | | | | | 165 vs. 105 larvae with 2 vs 0 mutated alleles | | | | |
| | | Effect | SE | *P* | lci | uci | Effect | SE | *P* | lci | uci | Effect | SE | *P* | lci | uci | Effect | SE | *P* | lci | uci |
| fixed factors | **2 vs. 0 mutated alleles** | 0.494 | 0.194 | 1.08E-02 | 0.114 | 0.873 | 0.164 | 0.206 | 4.25E-01 | -0.239 | 0.567 | -0.205 | 0.218 | 3.48E-01 | -0.632 | 0.223 | -0.080 | 0.147 | 5.87E-01 | -0.367 | 0.208 |
| | ***apoea*** | - | - | - | - | - | 0.242 | 0.097 | 1.21E-02 | 0.053 | 0.431 | 0.163 | 0.123 | 1.85E-01 | -0.078 | 0.404 | 0.133 | 0.094 | 1.57E-01 | -0.051 | 0.317 |
| | ***apoeb*** | 0.030 | 0.141 | 8.31E-01 | -0.247 | 0.307 | - | - | - | - | - | 0.040 | 0.118 | 7.32E-01 | -0.191 | 0.272 | 0.075 | 0.099 | 4.47E-01 | -0.118 | 0.268 |
| | ***apobb.1*** | -0.026 | 0.134 | 8.46E-01 | -0.288 | 0.236 | 0.014 | 0.108 | 8.95E-01 | -0.197 | 0.225 | - | - | - | - | - | 0.003 | 0.101 | 9.74E-01 | -0.195 | 0.201 |
| | ***ldlra*** | -0.087 | 0.109 | 4.22E-01 | -0.301 | 0.126 | -0.015 | 0.083 | 8.62E-01 | -0.178 | 0.149 | -0.022 | 0.111 | 8.44E-01 | -0.240 | 0.196 | - | - | - | - | - |
| | **time of day (in hours since 9AM)** | -0.051 | 0.063 | 4.20E-01 | -0.175 | 0.073 | 0.068 | 0.051 | 1.83E-01 | -0.032 | 0.168 | 0.046 | 0.060 | 4.42E-01 | -0.072 | 0.164 | 0.047 | 0.050 | 3.45E-01 | -0.050 | 0.144 |
| | **intercept** | 27.315 | 78.833 | 7.29E-01 | -127.196 | 181.825 | 42.957 | 64.470 | 5.05E-01 | -83.401 | 169.315 | 96.277 | 89.628 | 2.83E-01 | -79.391 | 271.946 | 0.132 | 1.258 | 9.16E-01 | -2.332 | 2.597 |
| | ***apoba*** | 0.466 | 0.525 | 3.75E-01 | -0.563 | 1.496 | 0.042 | 0.378 | 9.11E-01 | -0.698 | 0.782 | 0.044 | 0.459 | 9.24E-01 | -0.855 | 0.942 | 0.364 | 0.368 | 3.22E-01 | -0.356 | 1.085 |
| | ***apobb.2*** | 0.557 | 0.350 | 1.11E-01 | -0.128 | 1.242 | 0.412 | 0.243 | 9.00E-02 | -0.064 | 0.888 | 0.326 | 0.281 | 2.46E-01 | -0.224 | 0.876 | 0.265 | 0.238 | 2.66E-01 | -0.202 | 0.731 |
| | ***ldlrb*** | -72.128 | 197.119 | 7.14E-01 | -458.474 | 314.217 | -110.280 | 161.325 | 4.94E-01 | -426.471 | 205.911 | -243.081 | 224.383 | 2.79E-01 | -682.863 | 196.702 | -3.696 | 2.490 | 1.38E-01 | -8.576 | 1.184 |
| random factors | ***variance by batch*** | *0.202* | *0.140* | - | *0.052* | *0.787* | *0.315* | *0.126* | - | *0.144* | *0.689* | *0.203* | *0.383* | - | *0.005* | *8.145* | *0.356* | *0.136* | | *0.168* | *0.753* |
| | ***residual*** | *0.977* | *0.057* | - | *0.871* | *1.094* | *0.967* | *0.042* | - | *0.889* | *1.052* | *0.958* | *0.063* | - | *0.842* | *1.090* | *0.972* | *0.040* | | *0.896* | *1.054* |

| | | HDL cholesterol levels | | | | | | | | | | | | | | | | | | | |
|---|---|---|---|---|---|---|---|---|---|---|---|---|---|---|---|---|---|---|---|---|---|
| | | ***apoea*** | | | | | ***apoeb*** | | | | | ***apobb.1*** | | | | | ***ldlra*** | | | | |
| | | Effect | SE | *P* | lci | uci | Effect | SE | *P* | lci | uci | Effect | SE | *P* | lci | uci | Effect | SE | *P* | lci | uci |
| fixed factors | **2 vs. 0 mutated alleles** | 0.123 | 0.190 | 5.15E-01 | -0.248 | 0.495 | 0.223 | 0.180 | 2.16E-01 | -0.130 | 0.577 | 0.253 | 0.189 | 1.79E-01 | -0.116 | 0.623 | 0.072 | 0.132 | 5.85E-01 | -0.187 | 0.331 |
| | ***apoea*** | - | - | - | - | - | 0.035 | 0.086 | 6.80E-01 | -0.132 | 0.203 | -0.040 | 0.109 | 7.12E-01 | -0.254 | 0.174 | -0.010 | 0.085 | 9.02E-01 | -0.177 | 0.156 |
| | ***apoeb*** | -0.058 | 0.132 | 6.59E-01 | -0.317 | 0.200 | - | - | - | - | - | 0.148 | 0.102 | 1.47E-01 | -0.052 | 0.347 | 0.073 | 0.088 | 4.08E-01 | -0.100 | 0.246 |
| | ***apobb.1*** | 0.011 | 0.124 | 9.29E-01 | -0.233 | 0.255 | 0.037 | 0.095 | 6.93E-01 | -0.148 | 0.223 | - | - | - | - | - | 0.118 | 0.090 | 1.90E-01 | -0.059 | 0.296 |
| | ***ldlra*** | 0.130 | 0.101 | 1.99E-01 | -0.068 | 0.328 | 0.104 | 0.073 | 1.56E-01 | -0.040 | 0.248 | 0.090 | 0.098 | 3.60E-01 | -0.103 | 0.283 | - | - | - | - | - |
| | **time of day (in hours since 9AM)** | -0.077 | 0.066 | 2.42E-01 | -0.206 | 0.052 | -0.065 | 0.047 | 1.66E-01 | -0.157 | 0.027 | -0.109 | 0.056 | 5.18E-02 | -0.219 | 0.001 | -0.014 | 0.046 | 7.59E-01 | -0.105 | 0.076 |
| | **intercept** | 79.225 | 73.332 | 2.80E-01 | -64.504 | 222.954 | -2.809 | 57.043 | 9.61E-01 | -114.611 | 108.993 | -22.711 | 77.230 | 7.69E-01 | -174.080 | 128.657 | -0.979 | 1.141 | 3.91E-01 | -3.216 | 1.257 |
| | ***apoba*** | 0.053 | 0.489 | 9.13E-01 | -0.905 | 1.012 | -0.412 | 0.333 | 2.16E-01 | -1.065 | 0.240 | 0.014 | 0.404 | 9.73E-01 | -0.777 | 0.805 | 0.088 | 0.330 | 7.89E-01 | -0.558 | 0.735 |
| | ***apobb.2*** | 0.444 | 0.324 | 1.70E-01 | -0.191 | 1.078 | 0.196 | 0.214 | 3.60E-01 | -0.223 | 0.614 | 0.210 | 0.243 | 3.88E-01 | -0.266 | 0.686 | 0.229 | 0.213 | 2.84E-01 | -0.189 | 0.646 |
| | ***ldlrb*** | -199.333 | 183.370 | 2.77E-01 | -558.731 | 160.065 | 7.909 | 142.733 | 9.56E-01 | -271.843 | 287.661 | 55.951 | 193.319 | 7.72E-01 | -322.947 | 434.849 | 0.810 | 2.237 | 7.17E-01 | -3.574 | 5.194 |
| random factors | ***variance by batch*** | *0.570* | *0.172* | - | *0.316* | *1.029* | *0.579* | *0.160* | - | *0.336* | *0.997* | *0.528* | *0.161* | - | *0.290* | *0.958* | *0.584* | *0.162* | - | *0.340* | *1.005* |
| | ***residual*** | *0.901* | *0.052* | - | *0.804* | *1.008* | *0.847* | *0.036* | - | *0.779* | *0.921* | *0.822* | *0.045* | - | *0.739* | *0.914* | *0.869* | *0.036* | - | *0.801* | *0.942* |

| | | Triglyceride levels | | | | | | | | | | | | | | | | | | | |
|---|---|---|---|---|---|---|---|---|---|---|---|---|---|---|---|---|---|---|---|---|---|
| | | ***apoea*** | | | | | ***apoeb*** | | | | | ***apobb.1*** | | | | | ***ldlra*** | | | | |
| | | Effect | SE | *P* | lci | uci | Effect | SE | *P* | lci | uci | Effect | SE | *P* | lci | uci | Effect | SE | *P* | lci | uci |
| fixed factors | **2 vs. 0 mutated alleles** | -0.265 | 0.141 | 6.00E-02 | -0.541 | 0.011 | -0.052 | 0.145 | 7.21E-01 | -0.336 | 0.233 | 0.251 | 0.139 | 7.06E-02 | -0.021 | 0.522 | 0.034 | 0.104 | 7.41E-01 | -0.169 | 0.238 |
| | ***apoea*** | - | - | - | - | - | -0.075 | 0.069 | 2.77E-01 | -0.210 | 0.060 | -0.093 | 0.081 | 2.51E-01 | -0.251 | 0.066 | -0.037 | 0.067 | 5.80E-01 | -0.168 | 0.094 |
| | ***apoeb*** | -0.021 | 0.097 | 8.26E-01 | -0.210 | 0.168 | - | - | - | - | - | -0.057 | 0.075 | 4.44E-01 | -0.204 | 0.089 | -0.037 | 0.069 | 5.93E-01 | -0.173 | 0.099 |
| | ***apobb.1*** | 0.012 | 0.091 | 8.93E-01 | -0.166 | 0.190 | 0.115 | 0.076 | 1.30E-01 | -0.034 | 0.265 | - | - | - | - | - | 0.078 | 0.071 | 2.75E-01 | -0.062 | 0.217 |
| | ***ldlra*** | 0.034 | 0.074 | 6.46E-01 | -0.111 | 0.179 | 0.022 | 0.059 | 7.15E-01 | -0.094 | 0.138 | 0.045 | 0.073 | 5.33E-01 | -0.097 | 0.188 | - | - | - | - | - |
| | **time of day (in hours since 9AM)** | -0.146 | 0.050 | 3.27E-03 | -0.244 | -0.049 | -0.126 | 0.038 | 9.97E-04 | -0.201 | -0.051 | -0.145 | 0.042 | 5.90E-04 | -0.228 | -0.062 | -0.181 | 0.037 | 8.37E-07 | -0.253 | -0.109 |
| | **intercept** | 43.986 | 53.653 | 4.12E-01 | -61.173 | 149.145 | 21.252 | 46.003 | 6.44E-01 | -68.911 | 111.415 | 36.478 | 56.734 | 5.20E-01 | -74.719 | 147.675 | 0.872 | 0.924 | 3.46E-01 | -0.940 | 2.684 |
| | ***apoba*** | 0.079 | 0.358 | 8.25E-01 | -0.622 | 0.780 | -0.091 | 0.268 | 7.35E-01 | -0.616 | 0.435 | 0.025 | 0.298 | 9.32E-01 | -0.558 | 0.609 | -0.012 | 0.259 | 9.64E-01 | -0.520 | 0.497 |
| | ***apobb.2*** | -0.401 | 0.236 | 8.97E-02 | -0.865 | 0.062 | -0.402 | 0.172 | 1.96E-02 | -0.739 | -0.064 | -0.538 | 0.179 | 2.59E-03 | -0.889 | -0.188 | -0.246 | 0.167 | 1.42E-01 | -0.574 | 0.082 |
| | ***ldlrb*** | -108.252 | 134.160 | 4.20E-01 | -371.202 | 154.697 | -50.755 | 115.105 | 6.59E-01 | -276.357 | 174.846 | -88.498 | 142.009 | 5.33E-01 | -366.830 | 189.835 | -0.338 | 1.759 | 8.48E-01 | -3.786 | 3.110 |
| random factors | ***variance by batch*** | *0.789* | *0.209* | - | *0.470* | *1.326* | *0.810* | *0.209* | - | *0.489* | *1.342* | 0.720 | 0.188 | - | 0.431 | 1.203 | *0.784* | *0.202* | - | *0.473* | *1.299* |
| | ***residual*** | *0.657* | *0.038* | - | *0.587* | *0.736* | *0.681* | *0.029* | - | *0.626* | *0.741* | 0.603 | 0.033 | - | 0.543 | 0.671 | *0.682* | *0.028* | - | *0.629* | *0.740* |

*continued* **Supplementary Table 20**

| | | **Total cholesterol levels** | | | | | | | | | | | | | | | | | | | |
|---|---|---|---|---|---|---|---|---|---|---|---|---|---|---|---|---|---|---|---|---|---|
| | | ***apoea*** | | | | | ***apoeb*** | | | | | ***apobb.1*** | | | | | ***ldlra*** | | | | |
| | | **Effect** | **SE** | ***P*** | **lci** | **uci** | **Effect** | **SE** | ***P*** | **lci** | **uci** | **Effect** | **SE** | ***P*** | **lci** | **uci** | **Effect** | **SE** | ***P*** | **lci** | **uci** |
| **fixed factors** | **2 vs. 0 mutated alleles** | 0.204 | 0.168 | 2.25E-01 | -0.126 | 0.534 | 0.049 | 0.178 | 7.82E-01 | -0.300 | 0.399 | -0.881 | 0.184 | 1.63E-06 | -1.241 | -0.521 | -0.084 | 0.116 | 4.70E-01 | -0.311 | 0.144 |
| | ***apoea*** | - | - | - | - | - | 0.135 | 0.085 | 1.11E-01 | -0.031 | 0.301 | 0.074 | 0.107 | 4.88E-01 | -0.135 | 0.283 | 0.066 | 0.075 | 3.80E-01 | -0.081 | 0.212 |
| | ***apoeb*** | 0.076 | 0.116 | 5.09E-01 | -0.150 | 0.303 | - | - | - | - | - | 0.007 | 0.099 | 9.47E-01 | -0.188 | 0.201 | 0.031 | 0.077 | 6.89E-01 | -0.121 | 0.183 |
| | ***apobb.1*** | -0.351 | 0.109 | 1.29E-03 | -0.565 | -0.137 | -0.221 | 0.094 | 1.84E-02 | -0.404 | -0.037 | - | - | - | - | - | -0.171 | 0.079 | 3.09E-02 | -0.327 | -0.016 |
| | ***ldlra*** | -0.018 | 0.089 | 8.43E-01 | -0.191 | 0.156 | -0.007 | 0.073 | 9.21E-01 | -0.150 | 0.135 | 0.070 | 0.096 | 4.64E-01 | -0.118 | 0.259 | - | - | - | - | - |
| | **time of day (in hours since 9AM)** | 0.133 | 0.059 | 2.44E-02 | 0.017 | 0.249 | 0.094 | 0.047 | 4.32E-02 | 0.003 | 0.186 | 0.097 | 0.055 | 8.01E-02 | -0.012 | 0.205 | 0.088 | 0.041 | 3.17E-02 | 0.008 | 0.168 |
| | **intercept** | 28.011 | 64.334 | 6.63E-01 | -98.081 | 154.104 | 17.441 | 56.457 | 7.57E-01 | -93.213 | 128.096 | 108.667 | 75.188 | 1.48E-01 | -38.698 | 256.031 | 0.097 | 1.023 | 9.24E-01 | -1.908 | 2.102 |
| | ***apoba*** | 0.657 | 0.429 | 1.26E-01 | -0.184 | 1.498 | 0.071 | 0.329 | 8.30E-01 | -0.575 | 0.716 | 0.042 | 0.394 | 9.16E-01 | -0.730 | 0.814 | 0.196 | 0.290 | 4.98E-01 | -0.371 | 0.764 |
| | ***apobb.2*** | -0.009 | 0.284 | 9.74E-01 | -0.565 | 0.547 | -0.176 | 0.211 | 4.04E-01 | -0.591 | 0.238 | -0.248 | 0.237 | 2.95E-01 | -0.712 | 0.216 | -0.145 | 0.187 | 4.40E-01 | -0.511 | 0.222 |
| | ***ldlrb*** | -73.671 | 160.868 | 6.47E-01 | -388.966 | 241.625 | -43.732 | 141.267 | 7.57E-01 | -320.610 | 233.145 | -271.674 | 188.202 | 1.49E-01 | -640.542 | 97.195 | -0.969 | 1.965 | 6.22E-01 | -4.821 | 2.882 |
| **random factors** | ***variance by batch*** | *0.761* | *0.209* | - | *0.444* | *1.304* | *0.721* | *0.196* | - | *0.424* | *1.228* | *0.695* | *0.206* | - | *0.388* | *1.243* | *0.780* | *0.206* | - | *0.464* | *1.310* |
| | ***residual*** | *0.789* | *0.045* | - | *0.704* | *0.883* | *0.837* | *0.036* | - | *0.770* | *0.910* | *0.800* | *0.043* | - | *0.719* | *0.890* | *0.762* | *0.032* | - | *0.703* | *0.826* |

| | | **Glucose levels** | | | | | | | | | | | | | | | | | | | |
|---|---|---|---|---|---|---|---|---|---|---|---|---|---|---|---|---|---|---|---|---|---|
| | | ***apoea*** | | | | | ***apoeb*** | | | | | ***apobb.1*** | | | | | ***ldlra*** | | | | |
| | | **Effect** | **SE** | ***P*** | **lci** | **uci** | **Effect** | **SE** | ***P*** | **lci** | **uci** | **Effect** | **SE** | ***P*** | **lci** | **uci** | **Effect** | **SE** | ***P*** | **lci** | **uci** |
| **fixed factors** | **2 vs. 0 mutated alleles** | -0.270 | 0.199 | 1.74E-01 | -0.659 | 0.120 | 0.202 | 0.205 | 3.24E-01 | -0.199 | 0.603 | -0.485 | 0.210 | 2.10E-02 | -0.897 | -0.073 | -0.153 | 0.140 | 2.73E-01 | -0.427 | 0.121 |
| | ***apoea*** | - | - | - | - | - | -0.045 | 0.096 | 6.42E-01 | -0.234 | 0.144 | -0.009 | 0.120 | 9.37E-01 | -0.245 | 0.226 | -0.055 | 0.089 | 5.41E-01 | -0.230 | 0.121 |
| | ***apoeb*** | 0.065 | 0.140 | 6.42E-01 | -0.210 | 0.340 | - | - | - | - | - | -0.001 | 0.114 | 9.90E-01 | -0.224 | 0.221 | 0.112 | 0.094 | 2.33E-01 | -0.072 | 0.295 |
| | ***apobb.1*** | -0.226 | 0.132 | 8.73E-02 | -0.486 | 0.033 | -0.137 | 0.107 | 2.00E-01 | -0.347 | 0.073 | - | - | - | - | - | -0.138 | 0.096 | 1.50E-01 | -0.326 | 0.050 |
| | ***ldlra*** | -0.130 | 0.108 | 2.26E-01 | -0.341 | 0.081 | -0.007 | 0.083 | 9.35E-01 | -0.170 | 0.156 | -0.091 | 0.108 | 4.00E-01 | -0.304 | 0.121 | - | - | - | - | - |
| | **time of day (in hours since 9AM)** | -0.018 | 0.068 | 7.89E-01 | -0.151 | 0.115 | 0.046 | 0.052 | 3.72E-01 | -0.055 | 0.147 | 0.046 | 0.060 | 4.41E-01 | -0.071 | 0.163 | 0.028 | 0.047 | 5.51E-01 | -0.065 | 0.121 |
| | **intercept** | -13.418 | 78.005 | 8.63E-01 | -166.306 | 139.469 | 35.006 | 64.294 | 5.86E-01 | -91.008 | 161.019 | 7.275 | 86.223 | 9.33E-01 | -161.719 | 176.269 | 1.685 | 1.197 | 1.59E-01 | -0.661 | 4.031 |
| | ***apoba*** | 0.100 | 0.520 | 8.47E-01 | -0.920 | 1.120 | -0.202 | 0.376 | 5.92E-01 | -0.939 | 0.536 | 0.130 | 0.446 | 7.70E-01 | -0.743 | 1.004 | -0.207 | 0.350 | 5.54E-01 | -0.892 | 0.478 |
| | ***apobb.2*** | -0.790 | 0.345 | 2.19E-02 | -1.466 | -0.114 | -0.376 | 0.242 | 1.20E-01 | -0.850 | 0.098 | -0.523 | 0.271 | 5.36E-02 | -1.053 | 0.008 | -0.357 | 0.226 | 1.15E-01 | -0.800 | 0.087 |
| | ***ldlrb*** | 36.552 | 195.054 | 8.51E-01 | -345.747 | 418.850 | -85.721 | 160.882 | 5.94E-01 | -401.044 | 229.602 | -16.791 | 215.844 | 9.38E-01 | -439.838 | 406.255 | -2.133 | 2.369 | 3.68E-01 | -6.776 | 2.510 |
| **random factors** | ***variance by batch*** | *0.414* | *0.152* | - | *0.201* | *0.851* | *0.379* | *0.129* | - | *0.194* | *0.740* | *0.302* | *0.159* | - | *0.107* | *0.849* | *0.360* | *0.120* | - | *0.187* | *0.693* |
| | ***residual*** | *0.960* | *0.056* | - | *0.857* | *1.076* | *0.961* | *0.041* | - | *0.884* | *1.046* | *0.920* | *0.051* | - | *0.826* | *1.025* | *0.924* | *0.038* | - | *0.852* | *1.002* |

Dorsal and lateral body surface area and body volume were normalized for body length using residuals. All outcomes were inverse-normally transformed before the analysis and examined using hierarchical linear models. Effects shown are for larvae with two mutated alleles that are highly likely to affect protein function as predicted by Ensembl's Varient Effect Predictor (VEP) compared with larvae with zero CRISPR-mutated alleles. Associations were adjusted for the number of mutated alleles in the other six orthologues, weighted by their predicted effect on protein function, as well as for time of day and batch. Lci and uci are lower and upper boundaries of the 95% confidence interval.

**Supplementary Table 21 - The effect of two vs. zero mutated alleles in *apoea* , *apoeb* , *apobb.1* or *ldlra* on vascular atherogenic traits**

| | | | | | | | | | | | | | | | | |
|---|---|---|---|---|---|---|---|---|---|---|---|---|---|---|---|---|
| | | ***apoea*** | | | | | | | | | | | | | | |
| | | **Vascular lipid deposition** | | | | | | | | | | | | | | |
| | | **Model 1** | | | | | **Model 2** | | | | | **Model 3** | | | | |
| | | **31 vs. 83 larvae with 2 vs. 0 mutated alleles (n=114)** | | | | | **30 vs. 67 larvae with 2 vs. 0 mutated alleles (n=97)** | | | | | **30 vs. 67 larvae with 2 vs. 0 mutated alleles (n=97)** | | | | |
| | | **Effect** | **SE** | ***P*** | **lci** | **uci** | **Effect** | **SE** | ***P*** | **lci** | **uci** | **Effect** | **SE** | ***P*** | **lci** | **uci** |
| **negative binomial terms** | **2 vs. 0 mutated alleles** | -0.773 | 0.419 | 6.55E-02 | -1.595 | 0.050 | -0.802 | 0.471 | 8.88E-02 | -1.725 | 0.122 | -0.491 | 0.655 | 4.54E-01 | -1.776 | 0.793 |
| | ***apoeb*** | 0.365 | 0.195 | 6.08E-02 | -0.017 | 0.747 | 0.500 | 0.269 | 6.28E-02 | -0.027 | 1.027 | 0.466 | 0.282 | 9.80E-02 | -0.086 | 1.018 |
| | ***apobb.1*** | 1.007 | 0.233 | 1.59E-05 | 0.550 | 1.464 | 0.930 | 0.340 | 6.26E-03 | 0.263 | 1.597 | 0.875 | 0.349 | 1.22E-02 | 0.191 | 1.560 |
| | ***ldlra*** | 0.244 | 0.174 | 1.61E-01 | -0.098 | 0.586 | 0.229 | 0.201 | 2.54E-01 | -0.165 | 0.622 | 0.187 | 0.227 | 4.10E-01 | -0.258 | 0.632 |
| | **body length (in SD)** | - | - | - | - | - | 0.101 | 0.275 | 7.12E-01 | -0.437 | 0.640 | 0.205 | 0.280 | 4.64E-01 | -0.343 | 0.753 |
| | **dorsal body surface area (in SD)** | - | - | - | - | - | 0.470 | 0.274 | 8.59E-02 | -0.066 | 1.007 | 0.529 | 0.322 | 9.99E-02 | -0.101 | 1.160 |
| | **LDL cholesterol levels (in SD)** | - | - | - | - | - | - | - | - | - | - | -0.156 | 0.176 | 3.76E-01 | -0.501 | 0.189 |
| | **HDL cholesterol levels (in SD)** | - | - | - | - | - | - | - | - | - | - | 0.051 | 0.251 | 8.39E-01 | -0.442 | 0.544 |
| | **triglyceride levels (in SD)** | - | - | - | - | - | - | - | - | - | - | 0.423 | 0.172 | 1.39E-02 | 0.086 | 0.761 |
| | **glucose levels (in SD)** | - | - | - | - | - | - | - | - | - | - | 0.133 | 0.224 | 5.52E-01 | -0.306 | 0.573 |
| | **time of day (in hours since 9AM)** | 0.148 | 0.116 | 1.99E-01 | -0.078 | 0.375 | 0.029 | 0.179 | 8.72E-01 | -0.323 | 0.381 | 0.046 | 0.193 | 8.11E-01 | -0.331 | 0.424 |
| | **batch 2** | 0.070 | 0.527 | 8.94E-01 | -0.963 | 1.104 | 0.485 | 0.670 | 4.69E-01 | -0.829 | 1.799 | 0.913 | 0.828 | 2.70E-01 | -0.709 | 2.535 |
| | **batch 3** | -0.101 | 0.715 | 8.88E-01 | -1.502 | 1.301 | 0.264 | 0.737 | 7.20E-01 | -1.182 | 1.709 | 0.069 | 1.177 | 9.54E-01 | -2.239 | 2.376 |
| | **batch 4** | -0.760 | 0.525 | 1.48E-01 | -1.789 | 0.270 | -0.268 | 0.679 | 6.93E-01 | -1.599 | 1.063 | -0.372 | 0.785 | 6.35E-01 | -1.910 | 1.166 |
| | **batch 5** | 0.164 | 0.449 | 7.15E-01 | -0.716 | 1.045 | 0.462 | 0.602 | 4.42E-01 | -0.717 | 1.642 | -0.118 | 0.910 | 8.97E-01 | -1.903 | 1.666 |
| | **batch 6** | 0.086 | 0.415 | 8.36E-01 | -0.727 | 0.899 | -0.073 | 0.568 | 8.97E-01 | -1.187 | 1.040 | -0.803 | 0.874 | 3.59E-01 | -2.516 | 0.911 |
| | **batch 7** | -0.961 | 0.462 | 3.74E-02 | -1.866 | -0.056 | -1.047 | 0.624 | 9.33E-02 | -2.270 | 0.176 | -1.468 | 1.048 | 1.61E-01 | -3.522 | 0.586 |
| | **intercept** | -91.057 | 172.280 | 5.97E-01 | -428.720 | 246.607 | -109.553 | 223.730 | 6.24E-01 | -548.056 | 328.950 | -88.167 | 263.319 | 7.38E-01 | -604.262 | 427.928 |
| | ***apoba*** | -1.053 | 1.031 | 3.07E-01 | -3.074 | 0.968 | -1.141 | 1.295 | 3.78E-01 | -3.680 | 1.397 | -1.129 | 1.312 | 3.89E-01 | -3.699 | 1.442 |
| | ***apobb.2*** | -0.319 | 0.517 | 5.37E-01 | -1.333 | 0.694 | -0.360 | 0.594 | 5.44E-01 | -1.525 | 0.804 | 0.074 | 0.699 | 9.16E-01 | -1.296 | 1.443 |
| | ***ldlrb*** | 241.106 | 431.619 | 5.76E-01 | -604.850 | 1087.063 | 287.842 | 559.627 | 6.07E-01 | -809.006 | 1384.690 | 233.629 | 658.235 | 7.23E-01 | -1100.000 | 1523.746 |

| | | | | | | | | | | | | | | | | |
|---|---|---|---|---|---|---|---|---|---|---|---|---|---|---|---|---|
| | | **Vascular infiltration by macrophages** | | | | | | | | | | | | | | |
| | | **47 vs. 104 larvae with 2 vs. 0 mutated alleles (n=151)** | | | | | **44 vs. 89 larvae with 2 vs. 0 mutated alleles (n=133)** | | | | | **44 vs. 89 larvae with 2 vs. 0 mutated alleles (n=133)** | | | | |
| | | **Effect** | **SE** | ***P*** | **lci** | **uci** | **Effect** | **SE** | ***P*** | **lci** | **uci** | **Effect** | **SE** | ***P*** | **lci** | **uci** |
| **fixed factors** | **2 vs. 0 mutated alleles** | -0.063 | 0.177 | 7.23E-01 | -0.410 | 0.284 | -0.162 | 0.184 | 3.80E-01 | -0.524 | 0.200 | -0.187 | 0.187 | 3.18E-01 | -0.552 | 0.179 |
| | ***apoeb*** | -0.013 | 0.126 | 9.18E-01 | -0.260 | 0.234 | -0.006 | 0.139 | 9.67E-01 | -0.277 | 0.266 | 0.027 | 0.138 | 8.47E-01 | -0.244 | 0.297 |
| | ***apobb.1*** | -0.240 | 0.123 | 5.12E-02 | -0.481 | 0.001 | -0.294 | 0.137 | 3.17E-02 | -0.563 | -0.026 | -0.315 | 0.137 | 2.13E-02 | -0.583 | -0.047 |
| | ***ldlra*** | 0.088 | 0.097 | 3.64E-01 | -0.102 | 0.278 | 0.073 | 0.103 | 4.76E-01 | -0.128 | 0.274 | 0.048 | 0.102 | 6.39E-01 | -0.152 | 0.249 |
| | **body length (in SD)** | - | - | - | - | - | -0.193 | 0.110 | 7.85E-02 | -0.409 | 0.022 | -0.187 | 0.111 | 9.15E-02 | -0.403 | 0.030 |
| | **dorsal body surface area (in SD)** | - | - | - | - | - | -0.118 | 0.100 | 2.39E-01 | -0.313 | 0.078 | -0.109 | 0.101 | 2.83E-01 | -0.307 | 0.090 |
| | **LDL cholesterol levels (in SD)** | - | - | - | - | - | - | - | - | - | - | 0.057 | 0.083 | 4.92E-01 | -0.105 | 0.219 |
| | **HDL cholesterol levels (in SD)** | - | - | - | - | - | - | - | - | - | - | 0.108 | 0.089 | 2.27E-01 | -0.067 | 0.282 |
| | **triglyceride levels (in SD)** | - | - | - | - | - | - | - | - | - | - | 0.105 | 0.112 | 3.47E-01 | -0.114 | 0.324 |
| | **glucose levels (in SD)** | - | - | - | - | - | - | - | - | - | - | -0.123 | 0.084 | 1.44E-01 | -0.289 | 0.042 |
| | **time of day (in hours since 9AM)** | 0.049 | 0.062 | 4.29E-01 | -0.073 | 0.171 | 0.064 | 0.069 | 3.53E-01 | -0.071 | 0.200 | 0.075 | 0.068 | 2.71E-01 | -0.059 | 0.209 |
| | **intercept_random** | -9.339 | 69.740 | 8.93E-01 | -146.027 | 127.350 | 14.432 | 77.040 | 8.51E-01 | -136.563 | 165.427 | 8.768 | 76.105 | 9.08E-01 | -140.396 | 157.931 |
| | ***apoba*** | 0.070 | 0.463 | 8.80E-01 | -0.837 | 0.977 | 0.225 | 0.478 | 6.39E-01 | -0.713 | 1.162 | 0.150 | 0.473 | 7.51E-01 | -0.778 | 1.078 |
| | ***apobb.2*** | -0.383 | 0.303 | 2.06E-01 | -0.977 | 0.211 | -0.392 | 0.319 | 2.19E-01 | -1.017 | 0.233 | -0.547 | 0.327 | 9.47E-02 | -1.189 | 0.095 |
| | ***ldlrb*** | 24.231 | 174.423 | 8.90E-01 | -317.631 | 366.094 | -35.876 | 192.602 | 8.52E-01 | -413.370 | 341.618 | -20.891 | 190.282 | 9.13E-01 | -393.836 | 352.054 |
| **random factors** | ***variance by batch*** | *0.463* | *0.149* | *-* | *0.247* | *0.869* | *0.407* | *0.141* | *-* | *0.206* | *0.803* | *0.373* | *0.140* | *-* | *0.178* | *0.779* |
| | ***residual*** | *0.840* | *0.050* | *-* | *0.747* | *0.944* | *0.844* | *0.054* | *-* | *0.745* | *0.957* | *0.833* | *0.053* | *-* | *0.735* | *0.944* |

***continued*** **Supplementary Table 21**

| | | Vascular co-localization of lipids with macrophages | | | | | | | | | | | | | | |
|---|---|---|---|---|---|---|---|---|---|---|---|---|---|---|---|---|
| | | 31 vs. 80 larvae with 2 vs. 0 mutated alleles (n=111) | | | | | 30 vs. 65 larvae with 2 vs. 0 mutated alleles (n=95) | | | | | 30 vs. 65 larvae with 2 vs. 0 mutated alleles (n=95) | | | | |
| | | Effect | SE | *P* | lci | uci | Effect | SE | *P* | lci | uci | Effect | SE | *P* | lci | uci |
| negative binomial terms | **2 vs. 0 mutated alleles** | -0.387 | 0.699 | 5.80E-01 | -1.758 | 0.984 | -0.748 | 0.875 | 3.93E-01 | -2.463 | 0.967 | -0.531 | 0.976 | 5.86E-01 | -2.444 | 1.381 |
| | ***apoeb*** | 0.021 | 0.295 | 9.43E-01 | -0.558 | 0.600 | 0.251 | 0.314 | 4.24E-01 | -0.365 | 0.868 | 0.337 | 0.345 | 3.29E-01 | -0.339 | 1.014 |
| | ***apobb.1*** | 1.362 | 0.326 | 2.93E-05 | 0.723 | 2.001 | 1.142 | 0.513 | 2.61E-02 | 0.136 | 2.148 | 0.694 | 0.859 | 4.19E-01 | -0.990 | 2.378 |
| | ***ldlra*** | 0.166 | 0.351 | 6.35E-01 | -0.521 | 0.854 | 0.195 | 0.454 | 6.68E-01 | -0.696 | 1.086 | -0.096 | 0.468 | 8.37E-01 | -1.014 | 0.821 |
| | **body length (in SD)** | - | - | - | - | - | 0.078 | 0.479 | 8.70E-01 | -0.860 | 1.016 | 0.023 | 0.611 | 9.71E-01 | -1.174 | 1.219 |
| | **dorsal body surface area (in SD)** | - | - | - | - | - | 0.481 | 0.297 | 1.05E-01 | -0.101 | 1.062 | 0.426 | 0.645 | 5.09E-01 | -0.838 | 1.689 |
| | **LDL cholesterol levels (in SD)** | - | - | - | - | - | - | - | - | - | - | -0.081 | 0.327 | 8.03E-01 | -0.722 | 0.559 |
| | **HDL cholesterol levels (in SD)** | - | - | - | - | - | - | - | - | - | - | -0.016 | 0.498 | 9.74E-01 | -0.993 | 0.960 |
| | **triglyceride levels (in SD)** | - | - | - | - | - | - | - | - | - | - | 1.035 | 0.501 | 3.89E-02 | 0.053 | 2.018 |
| | **glucose levels (in SD)** | - | - | - | - | - | - | - | - | - | - | -0.228 | 0.522 | 6.62E-01 | -1.250 | 0.794 |
| | **time of day (in hours since 9AM)** | 0.215 | 0.140 | 1.25E-01 | -0.060 | 0.489 | 0.183 | 0.237 | 4.39E-01 | -0.281 | 0.648 | 0.327 | 0.361 | 3.65E-01 | -0.380 | 1.034 |
| | **batch 2** | 1.037 | 1.056 | 3.26E-01 | -1.033 | 3.107 | 0.982 | 1.145 | 3.91E-01 | -1.263 | 3.227 | 0.438 | 1.648 | 7.90E-01 | -2.792 | 3.668 |
| | **batch 3** | -1.382 | 0.712 | 5.22E-02 | -2.777 | 0.013 | -1.267 | 0.862 | 1.42E-01 | -2.956 | 0.423 | -1.449 | 1.537 | 3.46E-01 | -4.460 | 1.563 |
| | **batch 4** | -0.418 | 0.699 | 5.50E-01 | -1.789 | 0.952 | -0.393 | 0.911 | 6.66E-01 | -2.179 | 1.393 | -0.693 | 1.405 | 6.22E-01 | -3.447 | 2.061 |
| | **batch 5** | 0.438 | 0.619 | 4.80E-01 | -0.776 | 1.651 | 0.017 | 0.789 | 9.83E-01 | -1.529 | 1.563 | -1.515 | 1.427 | 2.89E-01 | -4.312 | 1.283 |
| | **batch 6** | -0.877 | 0.644 | 1.73E-01 | -2.139 | 0.386 | -1.652 | 0.760 | 2.98E-02 | -3.142 | -0.162 | -3.576 | 1.802 | 4.72E-02 | -7.107 | -0.044 |
| | **batch 7** | -2.995 | 0.769 | 9.76E-05 | -4.501 | -1.488 | -3.887 | 0.920 | 2.40E-05 | -5.691 | -2.084 | -4.793 | 2.075 | 2.09E-02 | -8.859 | -0.726 |
| | **intercept** | -233.734 | 227.970 | 3.05E-01 | -680.548 | 213.080 | -196.479 | 316.933 | 5.35E-01 | -817.657 | 424.699 | -194.514 | 380.442 | 6.09E-01 | -940.167 | 551.139 |
| | ***apoba*** | -1.350 | 1.667 | 4.18E-01 | -4.617 | 1.916 | -1.213 | 1.698 | 4.75E-01 | -4.540 | 2.115 | -0.244 | 2.329 | 9.16E-01 | -4.809 | 4.320 |
| | ***apobb.2*** | -0.885 | 0.700 | 2.06E-01 | -2.256 | 0.486 | -0.837 | 0.713 | 2.41E-01 | -2.235 | 0.561 | -0.087 | 1.485 | 9.53E-01 | -2.997 | 2.824 |
| | ***ldlrb*** | 596.038 | 572.764 | 2.98E-01 | -526.558 | 1718.634 | 502.398 | 792.141 | 5.26E-01 | -1100.000 | 2054.967 | 492.382 | 952.213 | 6.05E-01 | -1400.000 | 2358.685 |

| | | Vascular infiltration by neutrophils | | | | | | | | | | | | | | |
|---|---|---|---|---|---|---|---|---|---|---|---|---|---|---|---|---|
| | | 47 vs. 106 larvae with 2 vs. 0 mutated alleles (n=153) | | | | | 44 vs. 91 larvae with 2 vs. 0 mutated alleles (n=135) | | | | | 44 vs. 91 larvae with 2 vs. 0 mutated alleles (n=135) | | | | |
| | | Effect | SE | *P* | lci | uci | Effect | SE | *P* | lci | uci | Effect | SE | *P* | lci | uci |
| fixed factors | **2 vs. 0 mutated alleles** | -0.064 | 0.179 | 7.20E-01 | -0.416 | 0.287 | -0.080 | 0.176 | 6.48E-01 | -0.426 | 0.265 | -0.138 | 0.179 | 4.40E-01 | -0.489 | 0.213 |
| | ***apoeb*** | -0.108 | 0.128 | 3.98E-01 | -0.359 | 0.142 | -0.086 | 0.131 | 5.12E-01 | -0.344 | 0.171 | -0.083 | 0.131 | 5.30E-01 | -0.340 | 0.175 |
| | ***apobb.1*** | 0.140 | 0.125 | 2.65E-01 | -0.106 | 0.386 | -0.113 | 0.131 | 3.89E-01 | -0.369 | 0.144 | -0.112 | 0.132 | 3.95E-01 | -0.370 | 0.146 |
| | ***ldlra*** | -0.008 | 0.099 | 9.33E-01 | -0.202 | 0.185 | 0.069 | 0.098 | 4.84E-01 | -0.124 | 0.261 | 0.063 | 0.098 | 5.19E-01 | -0.129 | 0.256 |
| | **body length (in SD)** | - | - | - | - | - | -0.315 | 0.103 | 2.28E-03 | -0.517 | -0.113 | -0.325 | 0.103 | 1.59E-03 | -0.527 | -0.123 |
| | **dorsal body surface area (in SD)** | - | - | - | - | - | 0.191 | 0.094 | 4.25E-02 | 0.006 | 0.376 | 0.216 | 0.096 | 2.41E-02 | 0.028 | 0.404 |
| | **LDL cholesterol levels (in SD)** | - | - | - | - | - | - | - | - | - | - | 0.130 | 0.077 | 9.08E-02 | -0.021 | 0.282 |
| | **HDL cholesterol levels (in SD)** | - | - | - | - | - | - | - | - | - | - | 0.091 | 0.085 | 2.85E-01 | -0.076 | 0.257 |
| | **triglyceride levels (in SD)** | - | - | - | - | - | - | - | - | - | - | 0.047 | 0.103 | 6.48E-01 | -0.155 | 0.249 |
| | **glucose levels (in SD)** | - | - | - | - | - | - | - | - | - | - | 0.008 | 0.080 | 9.18E-01 | -0.149 | 0.165 |
| | **time of day (in hours since 9AM)** | 0.006 | 0.062 | 9.22E-01 | -0.115 | 0.128 | -0.030 | 0.065 | 6.45E-01 | -0.158 | 0.098 | -0.017 | 0.064 | 7.90E-01 | -0.142 | 0.108 |
| | **intercept_random** | -120.289 | 71.401 | 9.20E-02 | -260.231 | 19.654 | -125.307 | 73.902 | 9.00E-02 | -270.152 | 19.539 | -134.596 | 73.421 | 6.68E-02 | -278.498 | 9.306 |
| | ***apoba*** | 0.345 | 0.471 | 4.63E-01 | -0.577 | 1.268 | 0.182 | 0.456 | 6.89E-01 | -0.711 | 1.076 | 0.087 | 0.455 | 8.49E-01 | -0.805 | 0.978 |
| | ***apobb.2*** | -0.028 | 0.310 | 9.29E-01 | -0.636 | 0.580 | 0.174 | 0.306 | 5.71E-01 | -0.426 | 0.773 | 0.103 | 0.315 | 7.45E-01 | -0.515 | 0.721 |
| | ***ldlrb*** | 299.596 | 178.573 | 9.34E-02 | -50.401 | 649.593 | 312.556 | 184.759 | 9.07E-02 | -49.565 | 674.677 | 336.427 | 183.574 | 6.69E-02 | -23.371 | 696.226 |
| random factors | ***variance by batch*** | *0.375* | *0.121* | - | *0.199* | *0.706* | *0.334* | *0.121* | - | *0.164* | *0.681* | *0.271* | *0.117* | - | *0.116* | *0.632* |
| | ***residual*** | *0.861* | *0.051* | - | *0.768* | *0.967* | *0.812* | *0.051* | - | *0.717* | *0.919* | *0.807* | *0.051* | - | *0.713* | *0.913* |

***continued*** **Supplementary Table 21**

| | | Vascular co-localization of lipids with neutrophils | | | | | | | | | | | | | | |
|---|---|---|---|---|---|---|---|---|---|---|---|---|---|---|---|---|
| | | 31 vs. 75 larvae with 2 vs. 0 mutated alleles (n=106) | | | | | 30 vs. 60 larvae with 2 vs. 0 mutated alleles (n=90) | | | | | 30 vs. 60 larvae with 2 vs. 0 mutated alleles (n=90) | | | | |
| | | Effect | SE | *P* | lci | uci | Effect | SE | *P* | lci | uci | Effect | SE | *P* | lci | uci |
| **negative binomial terms** | **2 vs. 0 mutated alleles** | -0.823 | 0.679 | 2.25E-01 | -2.154 | 0.507 | -0.243 | 0.568 | 6.69E-01 | -1.355 | 0.870 | -0.523 | 0.644 | 4.17E-01 | -1.786 | 0.740 |
| | ***apoeb*** | 0.219 | 0.437 | 6.17E-01 | -0.638 | 1.075 | 0.780 | 0.294 | 7.90E-03 | 0.204 | 1.355 | 0.770 | 0.322 | 1.67E-02 | 0.140 | 1.401 |
| | ***apobb.1*** | 1.968 | 0.327 | 1.80E-09 | 1.327 | 2.610 | 2.615 | 0.502 | 1.90E-07 | 1.631 | 3.599 | 2.243 | 0.549 | 4.37E-05 | 1.167 | 3.319 |
| | ***ldlra*** | 0.702 | 0.327 | 3.20E-02 | 0.061 | 1.343 | 0.724 | 0.372 | 5.17E-02 | -0.005 | 1.452 | 0.562 | 0.354 | 1.13E-01 | -0.132 | 1.256 |
| | **body length (in SD)** | - | - | - | - | - | 0.073 | 0.477 | 8.79E-01 | -0.862 | 1.008 | 0.181 | 0.533 | 7.34E-01 | -0.863 | 1.225 |
| | **dorsal body surface area (in SD)** | - | - | - | - | - | -0.512 | 0.503 | 3.08E-01 | -1.497 | 0.473 | -0.284 | 0.538 | 5.98E-01 | -1.338 | 0.770 |
| | **LDL cholesterol levels (in SD)** | - | - | - | - | - | - | - | - | - | - | 0.033 | 0.332 | 9.22E-01 | -0.617 | 0.683 |
| | **HDL cholesterol levels (in SD)** | - | - | - | - | - | - | - | - | - | - | 0.646 | 0.379 | 8.87E-02 | -0.098 | 1.389 |
| | **triglyceride levels (in SD)** | - | - | - | - | - | - | - | - | - | - | 0.655 | 0.340 | 5.43E-02 | -0.012 | 1.322 |
| | **glucose levels (in SD)** | - | - | - | - | - | - | - | - | - | - | 0.106 | 0.262 | 6.85E-01 | -0.407 | 0.620 |
| | **time of day (in hours since 9AM)** | 0.469 | 0.179 | 9.00E-03 | 0.117 | 0.821 | 0.784 | 0.279 | 4.96E-03 | 0.237 | 1.331 | 0.642 | 0.305 | 3.50E-02 | 0.045 | 1.239 |
| | **batch 2** | 0.551 | 1.161 | 6.35E-01 | -1.725 | 2.827 | 1.792 | 1.096 | 1.02E-01 | -0.356 | 3.940 | 2.003 | 1.165 | 8.55E-02 | -0.280 | 4.286 |
| | **batch 3** | -1.322 | 1.155 | 2.52E-01 | -3.586 | 0.941 | -0.854 | 1.017 | 4.01E-01 | -2.847 | 1.139 | -0.100 | 1.270 | 9.38E-01 | -2.589 | 2.390 |
| | **batch 4** | -0.266 | 0.953 | 7.80E-01 | -2.134 | 1.603 | 0.286 | 0.971 | 7.68E-01 | -1.616 | 2.189 | 0.975 | 1.147 | 3.95E-01 | -1.273 | 3.223 |
| | **batch 5** | -0.247 | 0.830 | 7.66E-01 | -1.874 | 1.380 | 0.722 | 0.745 | 3.32E-01 | -0.738 | 2.182 | 0.661 | 1.019 | 5.16E-01 | -1.336 | 2.659 |
| | **batch 6** | -2.620 | 0.959 | 6.28E-03 | -4.499 | -0.741 | -1.984 | 0.865 | 2.19E-02 | -3.680 | -0.288 | -2.230 | 1.129 | 4.83E-02 | -4.443 | -0.017 |
| | **intercept** | -791.606 | 250.244 | 1.56E-03 | -1300.000 | -301.137 | -918.003 | 298.459 | 2.10E-03 | -1500.000 | -333.034 | -853.180 | 303.486 | 4.93E-03 | -1400.000 | -258.358 |
| | ***apoba*** | -3.117 | 1.464 | 3.32E-02 | -5.987 | -0.248 | -2.355 | 1.211 | 5.17E-02 | -4.728 | 0.017 | -2.021 | 1.458 | 1.66E-01 | -4.879 | 0.837 |
| | ***apobb.2*** | 1.520 | 0.907 | 9.38E-02 | -0.258 | 3.298 | -0.202 | 0.809 | 8.02E-01 | -1.787 | 1.383 | 0.105 | 0.976 | 9.14E-01 | -1.808 | 2.018 |
| | ***ldlrb*** | 1982.551 | 626.570 | 1.56E-03 | 754.497 | 3210.604 | 2291.772 | 746.022 | 2.13E-03 | 829.596 | 3753.949 | 2128.907 | 756.916 | 4.91E-03 | 645.379 | 3612.435 |

| | | Vascular co-localization of macrophages with neutrophils | | | | | | | | | | | | | | |
|---|---|---|---|---|---|---|---|---|---|---|---|---|---|---|---|---|
| | | 47 vs. 104 larvae with 2 vs. 0 mutated alleles (n=151) | | | | | 44 vs. 89 larvae with 2 vs. 0 mutated alleles (n=133) | | | | | 44 vs. 89 larvae with 2 vs. 0 mutated alleles (n=133) | | | | |
| | | Effect | SE | *P* | lci | uci | Effect | SE | *P* | lci | uci | Effect | SE | *P* | lci | uci |
| **negative binomial terms** | **2 vs. 0 mutated alleles** | -0.335 | 0.255 | 1.88E-01 | -0.834 | 0.164 | -0.252 | 0.247 | 3.07E-01 | -0.735 | 0.231 | -0.218 | 0.245 | 3.75E-01 | -0.699 | 0.263 |
| | ***apoeb*** | -0.434 | 0.195 | 2.61E-02 | -0.816 | -0.052 | -0.336 | 0.200 | 9.26E-02 | -0.728 | 0.056 | -0.367 | 0.197 | 6.30E-02 | -0.754 | 0.020 |
| | ***apobb.1*** | -0.105 | 0.162 | 5.18E-01 | -0.422 | 0.213 | -0.416 | 0.205 | 4.26E-02 | -0.818 | -0.014 | -0.427 | 0.201 | 3.36E-02 | -0.821 | -0.033 |
| | ***ldlra*** | 0.198 | 0.138 | 1.50E-01 | -0.071 | 0.468 | 0.203 | 0.143 | 1.57E-01 | -0.078 | 0.483 | 0.177 | 0.145 | 2.22E-01 | -0.107 | 0.462 |
| | **body length (in SD)** | - | - | - | - | - | 0.009 | 0.192 | 9.61E-01 | -0.366 | 0.385 | -0.019 | 0.190 | 9.18E-01 | -0.391 | 0.352 |
| | **dorsal body surface area (in SD)** | - | - | - | - | - | 0.741 | 0.289 | 1.04E-02 | 0.174 | 1.308 | 0.701 | 0.287 | 1.47E-02 | 0.138 | 1.264 |
| | **LDL cholesterol levels (in SD)** | - | - | - | - | - | - | - | - | - | - | 0.229 | 0.114 | 4.45E-02 | 0.006 | 0.451 |
| | **HDL cholesterol levels (in SD)** | - | - | - | - | - | - | - | - | - | - | 0.176 | 0.146 | 2.29E-01 | -0.110 | 0.461 |
| | **triglyceride levels (in SD)** | - | - | - | - | - | - | - | - | - | - | 0.293 | 0.146 | 4.49E-02 | 0.007 | 0.580 |
| | **glucose levels (in SD)** | - | - | - | - | - | - | - | - | - | - | 0.173 | 0.141 | 2.20E-01 | -0.104 | 0.449 |
| | **time of day (in hours since 9AM)** | 0.062 | 0.109 | 5.65E-01 | -0.150 | 0.275 | -0.112 | 0.142 | 4.27E-01 | -0.390 | 0.165 | -0.104 | 0.135 | 4.43E-01 | -0.368 | 0.161 |
| | **batch 1** | -0.433 | 0.383 | 2.58E-01 | -1.183 | 0.317 | -0.120 | 0.550 | 8.28E-01 | -1.199 | 0.959 | 0.085 | 0.667 | 8.99E-01 | -1.223 | 1.393 |
| | **batch 2** | -0.964 | 0.392 | 1.38E-02 | -1.732 | -0.197 | -0.997 | 0.445 | 2.52E-02 | -1.870 | -0.124 | -0.329 | 0.542 | 5.43E-01 | -1.391 | 0.732 |
| | **batch 3** | -1.324 | 0.485 | 6.36E-03 | -2.276 | -0.373 | -1.012 | 0.547 | 6.45E-02 | -2.085 | 0.061 | -0.777 | 0.552 | 1.59E-01 | -1.858 | 0.304 |
| | **batch 4** | -0.112 | 0.466 | 8.09E-01 | -1.025 | 0.800 | 0.237 | 0.546 | 6.63E-01 | -0.832 | 1.307 | 0.430 | 0.558 | 4.40E-01 | -0.663 | 1.524 |
| | **batch 5** | 0.030 | 0.469 | 9.48E-01 | -0.890 | 0.951 | 0.003 | 0.489 | 9.95E-01 | -0.956 | 0.962 | 0.060 | 0.535 | 9.10E-01 | -0.989 | 1.109 |
| | **batch 6** | -2.377 | 0.454 | 1.66E-07 | -3.267 | -1.487 | -3.075 | 0.624 | 8.32E-07 | -4.298 | -1.852 | -3.125 | 0.630 | 7.01E-07 | -4.359 | -1.890 |
| | **batch 7** | -1.110 | 0.731 | 1.29E-01 | -2.542 | 0.323 | -1.873 | 0.731 | 1.04E-02 | -3.306 | -0.440 | -1.714 | 0.733 | 1.93E-02 | -3.151 | -0.278 |
| | **intercept** | -55.609 | 120.106 | 6.43E-01 | -291.013 | 179.796 | -196.032 | 143.724 | 1.73E-01 | -477.726 | 85.662 | -287.996 | 137.895 | 3.68E-02 | -558.265 | -17.727 |
| | ***apoba*** | 1.159 | 0.625 | 6.37E-02 | -0.066 | 2.384 | 1.815 | 0.750 | 1.56E-02 | 0.344 | 3.286 | 1.848 | 0.744 | 1.30E-02 | 0.389 | 3.307 |
| | ***apobb.2*** | -0.200 | 0.324 | 5.38E-01 | -0.835 | 0.436 | -0.270 | 0.373 | 4.69E-01 | -1.002 | 0.461 | -0.283 | 0.404 | 4.84E-01 | -1.075 | 0.510 |
| | ***ldlrb*** | 146.298 | 300.491 | 6.26E-01 | -442.654 | 735.250 | 495.866 | 359.236 | 1.67E-01 | -208.223 | 1199.955 | 725.424 | 344.591 | 3.53E-02 | 50.039 | 1400.809 |

***continued*** **Supplementary Table 21**

| | | ***apoeb*** | | | | | | | | | | | | | | |
|---|---|---|---|---|---|---|---|---|---|---|---|---|---|---|---|---|
| | | **Vascular lipid deposition** | | | | | | | | | | | | | | |
| | | **Model 1** | | | | | **Model 2** | | | | | **Model 3** | | | | |
| | | **179 vs. 29 larvae with 2 vs. 0 mutated alleles (n=208)** | | | | | **153 vs. 26 larvae with 2 vs. 0 mutated alleles (n=179)** | | | | | **153 vs. 26 larvae with 2 vs. 0 mutated alleles (n=179)** | | | | |
| | | **Effect** | **SE** | ***P*** | **lci** | **uci** | **Effect** | **SE** | ***P*** | **lci** | **uci** | **Effect** | **SE** | ***P*** | **lci** | **uci** |
| **negative binomial terms** | **2 vs. 0 mutated alleles** | 0.479 | 0.305 | 1.16E-01 | -0.118 | 1.076 | 0.592 | 0.347 | 8.76E-02 | -0.087 | 1.272 | 0.564 | 0.370 | 1.28E-01 | -0.162 | 1.290 |
| | ***apoea*** | -0.238 | 0.188 | 2.06E-01 | -0.606 | 0.131 | -0.331 | 0.210 | 1.15E-01 | -0.742 | 0.081 | -0.263 | 0.207 | 2.04E-01 | -0.667 | 0.142 |
| | ***apobb.1*** | 0.730 | 0.174 | 2.64E-05 | 0.390 | 1.071 | 0.843 | 0.199 | 2.33E-05 | 0.452 | 1.234 | 0.671 | 0.201 | 8.13E-04 | 0.278 | 1.064 |
| | ***ldlra*** | 0.250 | 0.130 | 5.34E-02 | -0.004 | 0.504 | 0.267 | 0.149 | 7.25E-02 | -0.024 | 0.558 | 0.276 | 0.143 | 5.38E-02 | -0.004 | 0.556 |
| | **body length (in SD)** | - | - | - | - | - | 0.238 | 0.159 | 1.34E-01 | -0.074 | 0.550 | 0.336 | 0.158 | 3.36E-02 | 0.026 | 0.646 |
| | **dorsal body surface area (in SD)** | - | - | - | - | - | 0.217 | 0.148 | 1.43E-01 | -0.073 | 0.507 | 0.282 | 0.145 | 5.25E-02 | -0.003 | 0.567 |
| | **LDL cholesterol levels (in SD)** | - | - | - | - | - | - | - | - | - | - | -0.136 | 0.104 | 1.94E-01 | -0.340 | 0.069 |
| | **HDL cholesterol levels (in SD)** | - | - | - | - | - | - | - | - | - | - | -0.055 | 0.168 | 7.43E-01 | -0.384 | 0.274 |
| | **triglyceride levels (in SD)** | - | - | - | - | - | - | - | - | - | - | 0.435 | 0.167 | 9.32E-03 | 0.107 | 0.764 |
| | **glucose levels (in SD)** | - | - | - | - | - | - | - | - | - | - | -0.202 | 0.102 | 4.80E-02 | -0.403 | -0.002 |
| | **time of day (in hours since 9AM)** | -0.044 | 0.086 | 6.12E-01 | -0.213 | 0.125 | -0.103 | 0.105 | 3.28E-01 | -0.310 | 0.104 | -0.018 | 0.110 | 8.68E-01 | -0.233 | 0.197 |
| | **batch 2** | -1.404 | 0.457 | 2.14E-03 | -2.300 | -0.508 | -1.319 | 0.527 | 1.24E-02 | -2.352 | -0.285 | -0.923 | 0.528 | 8.03E-02 | -1.957 | 0.111 |
| | **batch 3** | -1.085 | 0.389 | 5.31E-03 | -1.847 | -0.322 | -1.113 | 0.409 | 6.48E-03 | -1.914 | -0.312 | -1.198 | 0.414 | 3.85E-03 | -2.010 | -0.386 |
| | **batch 4** | -1.068 | 0.442 | 1.56E-02 | -1.934 | -0.203 | -0.943 | 0.476 | 4.73E-02 | -1.876 | -0.011 | -0.826 | 0.435 | 5.74E-02 | -1.677 | 0.026 |
| | **batch 5** | -0.155 | 0.283 | 5.84E-01 | -0.709 | 0.399 | -0.238 | 0.325 | 4.64E-01 | -0.875 | 0.399 | -0.903 | 0.466 | 5.29E-02 | -1.817 | 0.011 |
| | **batch 6** | -0.217 | 0.295 | 4.63E-01 | -0.796 | 0.362 | -0.543 | 0.372 | 1.44E-01 | -1.273 | 0.186 | -1.323 | 0.550 | 1.62E-02 | -2.402 | -0.245 |
| | **batch 7** | -0.867 | 0.388 | 2.55E-02 | -1.628 | -0.106 | -1.179 | 0.401 | 3.24E-03 | -1.965 | -0.394 | -1.622 | 0.595 | 6.37E-03 | -2.787 | -0.457 |
| | **intercept** | 58.235 | 136.197 | 6.69E-01 | -208.706 | 325.175 | 41.113 | 152.198 | 7.87E-01 | -257.191 | 339.416 | 31.011 | 152.088 | 8.38E-01 | -267.075 | 329.097 |
| | ***apoba*** | -0.199 | 0.608 | 7.43E-01 | -1.391 | 0.992 | -0.357 | 0.650 | 5.83E-01 | -1.632 | 0.918 | -0.180 | 0.646 | 7.81E-01 | -1.447 | 1.087 |
| | ***apobb.2*** | 0.001 | 0.463 | 9.98E-01 | -0.907 | 0.909 | -0.029 | 0.491 | 9.52E-01 | -0.992 | 0.933 | 0.336 | 0.478 | 4.82E-01 | -0.601 | 1.274 |
| | ***ldlrb*** | -133.537 | 341.561 | 6.96E-01 | -802.985 | 535.911 | -89.418 | 381.314 | 8.15E-01 | -836.779 | 657.943 | -66.125 | 381.349 | 8.62E-01 | -813.556 | 681.305 |

| | | **Vascular infiltration by macrophages** | | | | | | | | | | | | | | |
|---|---|---|---|---|---|---|---|---|---|---|---|---|---|---|---|---|
| | | **237 vs. 37 larvae with 2 vs. 0 mutated alleles (n=274)** | | | | | **206 vs. 34 larvae with 2 vs. 0 mutated alleles (n=240)** | | | | | **206 vs. 34 larvae with 2 vs. 0 mutated alleles (n=240)** | | | | |
| | | **Effect** | **SE** | ***P*** | **lci** | **uci** | **Effect** | **SE** | ***P*** | **lci** | **uci** | **Effect** | **SE** | ***P*** | **lci** | **uci** |
| **fixed factors** | **2 vs. 0 mutated alleles** | -0.257 | 0.184 | 1.62E-01 | -0.617 | 0.103 | -0.218 | 0.203 | 2.83E-01 | -0.616 | 0.180 | -0.245 | 0.204 | 2.30E-01 | -0.646 | 0.155 |
| | ***apoea*** | -0.117 | 0.088 | 1.82E-01 | -0.289 | 0.055 | -0.148 | 0.096 | 1.24E-01 | -0.336 | 0.041 | -0.160 | 0.097 | 9.97E-02 | -0.350 | 0.030 |
| | ***apobb.1*** | -0.168 | 0.098 | 8.57E-02 | -0.361 | 0.024 | -0.173 | 0.107 | 1.05E-01 | -0.383 | 0.036 | -0.172 | 0.108 | 1.13E-01 | -0.385 | 0.040 |
| | ***ldlra*** | 0.013 | 0.075 | 8.67E-01 | -0.135 | 0.160 | 0.018 | 0.085 | 8.35E-01 | -0.149 | 0.184 | 0.002 | 0.085 | 9.81E-01 | -0.165 | 0.169 |
| | **body length (in SD)** | - | - | - | - | - | 0.023 | 0.086 | 7.91E-01 | -0.146 | 0.192 | 0.024 | 0.088 | 7.85E-01 | -0.148 | 0.195 |
| | **dorsal body surface area (in SD)** | - | - | - | - | - | -0.037 | 0.075 | 6.27E-01 | -0.184 | 0.111 | -0.039 | 0.076 | 6.07E-01 | -0.187 | 0.109 |
| | **LDL cholesterol levels (in SD)** | - | - | - | - | - | - | - | - | - | - | 0.029 | 0.063 | 6.45E-01 | -0.095 | 0.153 |
| | **HDL cholesterol levels (in SD)** | - | - | - | - | - | - | - | - | - | - | 0.112 | 0.074 | 1.30E-01 | -0.033 | 0.256 |
| | **triglyceride levels (in SD)** | - | - | - | - | - | - | - | - | - | - | 0.000 | 0.086 | 1.00E+00 | -0.169 | 0.169 |
| | **glucose levels (in SD)** | - | - | - | - | - | - | - | - | - | - | 0.031 | 0.064 | 6.34E-01 | -0.095 | 0.156 |
| | **time of day (in hours since 9AM)** | 0.020 | 0.048 | 6.72E-01 | -0.074 | 0.114 | 0.018 | 0.056 | 7.43E-01 | -0.091 | 0.128 | 0.023 | 0.056 | 6.79E-01 | -0.086 | 0.133 |
| | **intercept_random** | -49.906 | 57.788 | 3.88E-01 | -163.168 | 63.355 | -42.117 | 65.580 | 5.21E-01 | -170.650 | 86.417 | -34.466 | 65.536 | 5.99E-01 | -162.914 | 93.982 |
| | ***apoba*** | -0.187 | 0.340 | 5.82E-01 | -0.853 | 0.479 | -0.318 | 0.374 | 3.96E-01 | -1.051 | 0.416 | -0.283 | 0.374 | 4.49E-01 | -1.017 | 0.450 |
| | ***apobb.2*** | -0.359 | 0.217 | 9.75E-02 | -0.783 | 0.066 | -0.391 | 0.238 | 9.97E-02 | -0.857 | 0.075 | -0.416 | 0.242 | 8.58E-02 | -0.891 | 0.059 |
| | ***ldlrb*** | 127.830 | 144.603 | 3.77E-01 | -155.586 | 411.246 | 109.097 | 164.010 | 5.06E-01 | -212.356 | 430.551 | 89.972 | 163.907 | 5.83E-01 | -231.281 | 411.224 |
| **random factors** | ***variance by batch*** | *0.510* | *0.143* | - | *0.295* | *0.883* | *0.492* | *0.143* | - | *0.278* | *0.870* | *0.452* | *0.136* | - | *0.250* | *0.816* |
| | ***residual*** | *0.857* | *0.037* | - | *0.787* | *0.933* | *0.886* | *0.041* | - | *0.809* | *0.971* | *0.884* | *0.041* | - | *0.806* | *0.968* |

*continued* **Supplementary Table 21**

| | | Vascular co-localization of lipids with macrophages | | | | | | | | | | | | | | |
|---|---|---|---|---|---|---|---|---|---|---|---|---|---|---|---|---|
| | | **174 vs. 29 larvae with 2 vs. 0 mutated alleles (n=203)** | | | | | **150 vs. 26 larvae with 2 vs. 0 mutated alleles (n=176)** | | | | | **150 vs. 26 larvae with 2 vs. 0 mutated alleles (n=176)** | | | | |
| | | **Effect** | **SE** | ***P*** | **lci** | **uci** | **Effect** | **SE** | ***P*** | **lci** | **uci** | **Effect** | **SE** | ***P*** | **lci** | **uci** |
| **negative binomial terms** | **2 vs. 0 mutated alleles** | -0.468 | 0.389 | 2.29E-01 | -1.231 | 0.295 | -0.477 | 0.443 | 2.81E-01 | -1.345 | 0.391 | -0.283 | 0.464 | 5.41E-01 | -1.192 | 0.625 |
| | ***apoea*** | -0.247 | 0.253 | 3.28E-01 | -0.742 | 0.248 | -0.429 | 0.292 | 1.42E-01 | -1.000 | 0.143 | -0.402 | 0.343 | 2.41E-01 | -1.074 | 0.271 |
| | ***apobb.1*** | 0.934 | 0.220 | 2.27E-05 | 0.502 | 1.366 | 1.015 | 0.287 | 3.99E-04 | 0.453 | 1.576 | 0.692 | 0.352 | 4.91E-02 | 0.003 | 1.382 |
| | ***ldlra*** | 0.242 | 0.185 | 1.91E-01 | -0.121 | 0.606 | 0.306 | 0.221 | 1.65E-01 | -0.126 | 0.739 | 0.368 | 0.220 | 9.40E-02 | -0.063 | 0.799 |
| | **body length (in SD)** | - | - | - | - | - | 0.242 | 0.260 | 3.53E-01 | -0.268 | 0.751 | 0.234 | 0.279 | 4.02E-01 | -0.313 | 0.780 |
| | **dorsal body surface area (in SD)** | - | - | - | - | - | 0.471 | 0.184 | 1.07E-02 | 0.109 | 0.832 | 0.446 | 0.182 | 1.46E-02 | 0.088 | 0.804 |
| | **LDL cholesterol levels (in SD)** | - | - | - | - | - | - | - | - | - | - | -0.307 | 0.232 | 1.87E-01 | -0.762 | 0.149 |
| | **HDL cholesterol levels (in SD)** | - | - | - | - | - | - | - | - | - | - | 0.051 | 0.279 | 8.56E-01 | -0.496 | 0.597 |
| | **triglyceride levels (in SD)** | - | - | - | - | - | - | - | - | - | - | 0.467 | 0.265 | 7.77E-02 | -0.052 | 0.986 |
| | **glucose levels (in SD)** | - | - | - | - | - | - | - | - | - | - | -0.139 | 0.163 | 3.92E-01 | -0.459 | 0.180 |
| | **time of day (in hours since 9AM)** | 0.097 | 0.098 | 3.24E-01 | -0.095 | 0.289 | 0.200 | 0.124 | 1.05E-01 | -0.042 | 0.443 | 0.338 | 0.138 | 1.47E-02 | 0.067 | 0.609 |
| | **batch 2** | -0.438 | 0.824 | 5.95E-01 | -2.053 | 1.177 | -0.016 | 0.967 | 9.87E-01 | -1.911 | 1.880 | -0.594 | 0.914 | 5.16E-01 | -2.384 | 1.197 |
| | **batch 3** | -1.015 | 0.444 | 2.21E-02 | -1.885 | -0.146 | -0.976 | 0.490 | 4.65E-02 | -1.936 | -0.015 | -1.036 | 0.648 | 1.10E-01 | -2.305 | 0.234 |
| | **batch 4** | -0.450 | 0.471 | 3.40E-01 | -1.373 | 0.474 | -0.505 | 0.568 | 3.74E-01 | -1.619 | 0.608 | -0.674 | 0.600 | 2.62E-01 | -1.851 | 0.503 |
| | **batch 5** | 0.033 | 0.377 | 9.30E-01 | -0.706 | 0.772 | -0.486 | 0.464 | 2.95E-01 | -1.396 | 0.424 | -1.300 | 0.662 | 4.97E-02 | -2.597 | -0.002 |
| | **batch 6** | -0.548 | 0.370 | 1.38E-01 | -1.273 | 0.176 | -0.968 | 0.532 | 6.91E-02 | -2.011 | 0.076 | -1.528 | 0.962 | 1.12E-01 | -3.413 | 0.357 |
| | **batch 7** | -1.275 | 0.742 | 8.58E-02 | -2.729 | 0.180 | -2.061 | 0.725 | 4.46E-03 | -3.482 | -0.640 | -2.677 | 1.017 | 8.47E-03 | -4.671 | -0.684 |
| | **intercept** | 20.330 | 157.888 | 8.98E-01 | -289.125 | 329.784 | 52.867 | 187.217 | 7.78E-01 | -314.072 | 419.806 | 162.185 | 229.589 | 4.80E-01 | -287.800 | 612.170 |
| | ***apoba*** | -0.470 | 0.923 | 6.11E-01 | -2.278 | 1.339 | -0.523 | 1.012 | 6.06E-01 | -2.507 | 1.461 | -0.572 | 1.353 | 6.73E-01 | -3.223 | 2.080 |
| | ***apobb.2*** | -0.205 | 0.494 | 6.78E-01 | -1.173 | 0.762 | -0.180 | 0.536 | 7.38E-01 | -1.229 | 0.870 | 0.175 | 0.536 | 7.45E-01 | -0.876 | 1.225 |
| | ***ldlrb*** | -42.152 | 396.237 | 9.15E-01 | -818.763 | 734.458 | -123.631 | 468.918 | 7.92E-01 | -1000.000 | 795.432 | -398.212 | 576.562 | 4.90E-01 | -1500.000 | 731.829 |

| | | Vascular infiltration by neutrophils | | | | | | | | | | | | | | |
|---|---|---|---|---|---|---|---|---|---|---|---|---|---|---|---|---|
| | | **240 vs. 37 larvae with 2 vs. 0 mutated alleles (n=277)** | | | | | **208 vs. 34 larvae with 2 vs. 0 mutated alleles (n=242)** | | | | | **208 vs. 34 larvae with 2 vs. 0 mutated alleles (n=242)** | | | | |
| | | **Effect** | **SE** | ***P*** | **lci** | **uci** | **Effect** | **SE** | ***P*** | **lci** | **uci** | **Effect** | **SE** | ***P*** | **lci** | **uci** |
| **fixed factors** | **2 vs. 0 mutated alleles** | 0.100 | 0.179 | 5.77E-01 | -0.251 | 0.451 | 0.163 | 0.192 | 3.96E-01 | -0.214 | 0.540 | 0.128 | 0.192 | 5.06E-01 | -0.249 | 0.505 |
| | ***apoea*** | -0.032 | 0.085 | 7.08E-01 | -0.199 | 0.135 | -0.049 | 0.091 | 5.90E-01 | -0.227 | 0.129 | -0.073 | 0.091 | 4.22E-01 | -0.253 | 0.106 |
| | ***apobb.1*** | 0.125 | 0.095 | 1.89E-01 | -0.061 | 0.311 | 0.104 | 0.101 | 3.02E-01 | -0.093 | 0.301 | 0.106 | 0.102 | 2.96E-01 | -0.093 | 0.305 |
| | ***ldlra*** | -0.083 | 0.073 | 2.58E-01 | -0.227 | 0.061 | -0.078 | 0.080 | 3.32E-01 | -0.236 | 0.079 | -0.099 | 0.080 | 2.18E-01 | -0.256 | 0.058 |
| | **body length (in SD)** | - | - | - | - | - | -0.027 | 0.081 | 7.42E-01 | -0.186 | 0.132 | -0.018 | 0.081 | 8.26E-01 | -0.177 | 0.141 |
| | **dorsal body surface area (in SD)** | - | - | - | - | - | 0.070 | 0.070 | 3.17E-01 | -0.068 | 0.209 | 0.076 | 0.070 | 2.80E-01 | -0.062 | 0.213 |
| | **LDL cholesterol levels (in SD)** | - | - | - | - | - | - | - | - | - | - | 0.071 | 0.058 | 2.22E-01 | -0.043 | 0.185 |
| | **HDL cholesterol levels (in SD)** | - | - | - | - | - | - | - | - | - | - | 0.138 | 0.069 | 4.47E-02 | 0.003 | 0.272 |
| | **triglyceride levels (in SD)** | - | - | - | - | - | - | - | - | - | - | -0.012 | 0.079 | 8.77E-01 | -0.167 | 0.143 |
| | **glucose levels (in SD)** | - | - | - | - | - | - | - | - | - | - | 0.019 | 0.060 | 7.48E-01 | -0.098 | 0.137 |
| | **time of day (in hours since 9AM)** | 0.041 | 0.046 | 3.69E-01 | -0.049 | 0.132 | 0.002 | 0.052 | 9.72E-01 | -0.100 | 0.104 | 0.006 | 0.052 | 9.02E-01 | -0.095 | 0.108 |
| | **intercept_random** | 53.743 | 56.805 | 3.44E-01 | -57.594 | 165.079 | 26.564 | 62.657 | 6.72E-01 | -96.241 | 149.369 | 35.841 | 62.150 | 5.64E-01 | -85.971 | 157.654 |
| | ***apoba*** | 0.077 | 0.330 | 8.15E-01 | -0.570 | 0.725 | 0.102 | 0.353 | 7.73E-01 | -0.590 | 0.794 | 0.131 | 0.351 | 7.09E-01 | -0.557 | 0.820 |
| | ***apobb.2*** | 0.042 | 0.211 | 8.44E-01 | -0.373 | 0.456 | 0.070 | 0.225 | 7.55E-01 | -0.371 | 0.512 | 0.012 | 0.228 | 9.59E-01 | -0.435 | 0.459 |
| | ***ldlrb*** | -135.183 | 142.138 | 3.42E-01 | -413.770 | 143.403 | -67.148 | 156.701 | 6.68E-01 | -374.276 | 239.980 | -90.157 | 155.444 | 5.62E-01 | -394.822 | 214.507 |
| **random factors** | ***variance by batch*** | *0.407* | *0.115* | - | *0.234* | *0.708* | *0.395* | *0.118* | - | *0.220* | *0.709* | *0.343* | *0.108* | - | *0.186* | *0.634* |
| | ***residual*** | *0.838* | *0.036* | - | *0.770* | *0.912* | *0.842* | *0.039* | - | *0.768* | *0.922* | *0.834* | *0.039* | - | *0.762* | *0.914* |

***continued*** **Supplementary Table 21**

| | | Vascular co-localization of lipids with neutrophils | | | | | | | | | | | | | | |
|---|---|---|---|---|---|---|---|---|---|---|---|---|---|---|---|---|
| | | 168 vs. 27 larvae with 2 vs. 0 mutated alleles (n=195) | | | | | 143 vs. 24 larvae with 2 vs. 0 mutated alleles (n=167) | | | | | 143 vs. 24 larvae with 2 vs. 0 mutated alleles (n=167) | | | | |
| | | Effect | SE | *P* | lci | uci | Effect | SE | *P* | lci | uci | Effect | SE | *P* | lci | uci |
| negative binomial terms | **2 vs. 0 mutated alleles** | -0.389 | 0.565 | 4.91E-01 | -1.495 | 0.717 | 0.227 | 0.643 | 7.24E-01 | -1.033 | 1.488 | -0.109 | 0.648 | 8.67E-01 | -1.379 | 1.162 |
| | ***apoea*** | -0.241 | 0.339 | 4.77E-01 | -0.906 | 0.423 | 0.048 | 0.373 | 8.99E-01 | -0.684 | 0.779 | -0.974 | 0.391 | 1.27E-02 | -1.740 | -0.208 |
| | ***apobb.1*** | 1.748 | 0.280 | 4.46E-10 | 1.198 | 2.297 | 1.812 | 0.380 | 1.90E-06 | 1.066 | 2.557 | 1.613 | 0.377 | 1.88E-05 | 0.874 | 2.351 |
| | ***ldlra*** | 0.240 | 0.315 | 4.46E-01 | -0.377 | 0.858 | 0.092 | 0.322 | 7.76E-01 | -0.540 | 0.723 | 0.074 | 0.272 | 7.85E-01 | -0.459 | 0.607 |
| | **body length (in SD)** | - | - | - | - | - | 0.071 | 0.504 | 8.88E-01 | -0.917 | 1.060 | -0.058 | 0.414 | 8.88E-01 | -0.870 | 0.754 |
| | **dorsal body surface area (in SD)** | - | - | - | - | - | -0.177 | 0.246 | 4.71E-01 | -0.659 | 0.305 | -0.160 | 0.245 | 5.14E-01 | -0.640 | 0.320 |
| | **LDL cholesterol levels (in SD)** | - | - | - | - | - | - | - | - | - | - | 0.261 | 0.286 | 3.61E-01 | -0.299 | 0.821 |
| | **HDL cholesterol levels (in SD)** | - | - | - | - | - | - | - | - | - | - | 0.751 | 0.406 | 6.40E-02 | -0.044 | 1.546 |
| | **triglyceride levels (in SD)** | - | - | - | - | - | - | - | - | - | - | 1.409 | 0.367 | 1.24E-04 | 0.689 | 2.129 |
| | **glucose levels (in SD)** | - | - | - | - | - | - | - | - | - | - | -0.008 | 0.215 | 9.71E-01 | -0.428 | 0.413 |
| | **time of day (in hours since 9AM)** | 0.062 | 0.169 | 7.16E-01 | -0.270 | 0.393 | 0.188 | 0.195 | 3.36E-01 | -0.195 | 0.570 | 0.500 | 0.228 | 2.84E-02 | 0.053 | 0.947 |
| | **batch 2** | -1.617 | 0.813 | 4.68E-02 | -3.211 | -0.023 | -1.605 | 1.080 | 1.37E-01 | -3.722 | 0.511 | -1.421 | 0.967 | 1.42E-01 | -3.316 | 0.474 |
| | **batch 3** | -0.752 | 0.834 | 3.67E-01 | -2.387 | 0.882 | -0.720 | 0.791 | 3.63E-01 | -2.269 | 0.830 | -1.398 | 0.714 | 5.01E-02 | -2.797 | 0.000 |
| | **batch 4** | -0.887 | 0.621 | 1.53E-01 | -2.104 | 0.330 | -0.638 | 0.817 | 4.35E-01 | -2.240 | 0.964 | -1.172 | 0.794 | 1.40E-01 | -2.728 | 0.384 |
| | **batch 5** | -0.277 | 0.510 | 5.87E-01 | -1.277 | 0.722 | -0.327 | 0.550 | 5.52E-01 | -1.405 | 0.751 | -2.983 | 0.935 | 1.42E-03 | -4.816 | -1.150 |
| | **batch 6** | -2.640 | 0.635 | 3.23E-05 | -3.885 | -1.395 | -2.827 | 0.837 | 7.28E-04 | -4.467 | -1.187 | -4.597 | 1.110 | 3.43E-05 | -6.771 | -2.422 |
| | **intercept** | -101.955 | 224.820 | 6.50E-01 | -542.594 | 338.684 | 7.020 | 339.258 | 9.83E-01 | -657.913 | 671.953 | 444.862 | 301.233 | 1.40E-01 | -145.544 | 1035.268 |
| | ***apoba*** | 0.531 | 1.282 | 6.79E-01 | -1.982 | 3.045 | 1.061 | 1.155 | 3.58E-01 | -1.203 | 3.324 | 2.821 | 1.272 | 2.66E-02 | 0.327 | 5.315 |
| | ***apobb.2*** | 0.600 | 0.670 | 3.71E-01 | -0.714 | 1.913 | -0.122 | 0.678 | 8.57E-01 | -1.451 | 1.206 | 1.634 | 0.789 | 3.84E-02 | 0.087 | 3.180 |
| | ***ldlrb*** | 251.587 | 564.907 | 6.56E-01 | -855.610 | 1358.785 | -23.731 | 850.679 | 9.78E-01 | -1700.000 | 1643.569 | -1100.000 | 755.388 | 1.35E-01 | -2600.000 | 350.537 |

| | | Vascular co-localization of macrophages with neutrophils | | | | | | | | | | | | | | |
|---|---|---|---|---|---|---|---|---|---|---|---|---|---|---|---|---|
| | | 236 vs. 37 larvae with 2 vs. 0 mutated alleles (n=273) | | | | | 205 vs. 34 larvae with 2 vs. 0 mutated alleles (n=239) | | | | | 205 vs. 34 larvae with 2 vs. 0 mutated alleles (n=239) | | | | |
| | | Effect | SE | *P* | lci | uci | Effect | SE | *P* | lci | uci | Effect | SE | *P* | lci | uci |
| negative binomial terms | **2 vs. 0 mutated alleles** | 0.504 | 0.278 | 7.02E-02 | -0.042 | 1.050 | 0.498 | 0.280 | 7.53E-02 | -0.051 | 1.047 | 0.444 | 0.280 | 1.13E-01 | -0.105 | 0.993 |
| | ***apoea*** | -0.111 | 0.148 | 4.54E-01 | -0.400 | 0.179 | -0.100 | 0.157 | 5.26E-01 | -0.407 | 0.208 | -0.073 | 0.160 | 6.50E-01 | -0.386 | 0.241 |
| | ***apobb.1*** | 0.111 | 0.164 | 5.00E-01 | -0.211 | 0.432 | 0.142 | 0.163 | 3.84E-01 | -0.178 | 0.462 | 0.223 | 0.162 | 1.69E-01 | -0.095 | 0.542 |
| | ***ldlra*** | -0.181 | 0.126 | 1.50E-01 | -0.428 | 0.066 | -0.178 | 0.121 | 1.41E-01 | -0.415 | 0.059 | -0.201 | 0.119 | 9.11E-02 | -0.434 | 0.032 |
| | **body length (in SD)** | - | - | - | - | - | 0.471 | 0.145 | 1.16E-03 | 0.187 | 0.755 | 0.445 | 0.145 | 2.21E-03 | 0.160 | 0.729 |
| | **dorsal body surface area (in SD)** | - | - | - | - | - | 0.289 | 0.133 | 3.01E-02 | 0.028 | 0.550 | 0.250 | 0.130 | 5.49E-02 | -0.005 | 0.506 |
| | **LDL cholesterol levels (in SD)** | - | - | - | - | - | - | - | - | - | - | 0.073 | 0.102 | 4.76E-01 | -0.127 | 0.273 |
| | **HDL cholesterol levels (in SD)** | - | - | - | - | - | - | - | - | - | - | 0.082 | 0.119 | 4.89E-01 | -0.151 | 0.315 |
| | **triglyceride levels (in SD)** | - | - | - | - | - | - | - | - | - | - | -0.078 | 0.149 | 6.01E-01 | -0.370 | 0.215 |
| | **glucose levels (in SD)** | - | - | - | - | - | - | - | - | - | - | 0.256 | 0.089 | 3.79E-03 | 0.083 | 0.430 |
| | **time of day (in hours since 9AM)** | 0.112 | 0.077 | 1.43E-01 | -0.038 | 0.263 | 0.117 | 0.090 | 1.93E-01 | -0.059 | 0.293 | 0.120 | 0.092 | 1.94E-01 | -0.061 | 0.301 |
| | **batch 1** | 0.658 | 0.409 | 1.07E-01 | -0.143 | 1.459 | 0.246 | 0.463 | 5.95E-01 | -0.661 | 1.154 | 0.254 | 0.535 | 6.35E-01 | -0.794 | 1.302 |
| | **batch 2** | 0.750 | 0.544 | 1.68E-01 | -0.316 | 1.817 | 0.756 | 0.498 | 1.29E-01 | -0.221 | 1.732 | 0.924 | 0.538 | 8.61E-02 | -0.131 | 1.979 |
| | **batch 3** | -0.204 | 0.517 | 6.93E-01 | -1.216 | 0.809 | -0.543 | 0.525 | 3.01E-01 | -1.573 | 0.486 | -0.473 | 0.551 | 3.91E-01 | -1.552 | 0.606 |
| | **batch 4** | 1.501 | 0.490 | 2.20E-03 | 0.540 | 2.462 | 1.571 | 0.519 | 2.46E-03 | 0.554 | 2.587 | 1.458 | 0.561 | 9.42E-03 | 0.357 | 2.558 |
| | **batch 5** | 0.456 | 0.449 | 3.10E-01 | -0.424 | 1.335 | -0.114 | 0.493 | 8.17E-01 | -1.080 | 0.852 | 0.084 | 0.546 | 8.77E-01 | -0.985 | 1.154 |
| | **batch 6** | -1.210 | 0.466 | 9.40E-03 | -2.123 | -0.297 | -2.083 | 0.588 | 3.99E-04 | -3.236 | -0.930 | -1.925 | 0.620 | 1.90E-03 | -3.139 | -0.710 |
| | **batch 7** | -0.190 | 0.604 | 7.53E-01 | -1.374 | 0.994 | -1.063 | 0.767 | 1.66E-01 | -2.567 | 0.441 | -1.109 | 0.819 | 1.75E-01 | -2.714 | 0.495 |
| | **intercept** | 69.161 | 106.517 | 5.16E-01 | -139.609 | 277.931 | -25.456 | 109.673 | 8.16E-01 | -240.411 | 189.499 | -63.428 | 102.485 | 5.36E-01 | -264.296 | 137.440 |
| | ***apoba*** | -0.102 | 0.538 | 8.50E-01 | -1.157 | 0.953 | -0.007 | 0.546 | 9.90E-01 | -1.076 | 1.063 | 0.098 | 0.569 | 8.63E-01 | -1.017 | 1.214 |
| | ***apobb.2*** | 0.017 | 0.337 | 9.61E-01 | -0.644 | 0.678 | -0.223 | 0.379 | 5.57E-01 | -0.965 | 0.520 | -0.055 | 0.374 | 8.83E-01 | -0.788 | 0.677 |
| | ***ldlrb*** | -164.973 | 266.850 | 5.36E-01 | -687.989 | 358.042 | 73.008 | 274.916 | 7.91E-01 | -465.817 | 611.833 | 166.585 | 256.848 | 5.17E-01 | -336.827 | 669.997 |

***continued*** **Supplementary Table 21**

| | | ***apobb.1*** | | | | | | | | | | | | | | |
|---|---|---|---|---|---|---|---|---|---|---|---|---|---|---|---|---|
| | | **Vascular lipid deposition** | | | | | | | | | | | | | | |
| | | **Model 1** | | | | | **Model 2** | | | | | **Model 3** | | | | |
| | | **20 vs. 113 larvae with 2 vs. 0 mutated alleles (n=133)** | | | | | **20 vs. 95 larvae with 2 vs. 0 mutated alleles (n=115)** | | | | | **20 vs. 95 larvae with 2 vs. 0 mutated alleles (n=115)** | | | | |
| | | **Effect** | **SE** | ***P*** | **lci** | **uci** | **Effect** | **SE** | ***P*** | **lci** | **uci** | **Effect** | **SE** | ***P*** | **lci** | **uci** |
| **negative binomial terms** | **2 vs. 0 mutated alleles** | 1.765 | 0.360 | 9.32E-07 | 1.060 | 2.470 | 2.914 | 0.809 | 3.17E-04 | 1.328 | 4.499 | 2.482 | 0.850 | 3.50E-03 | 0.816 | 4.147 |
| | ***apoea*** | -0.406 | 0.264 | 1.24E-01 | -0.923 | 0.111 | -0.282 | 0.331 | 3.95E-01 | -0.930 | 0.367 | -0.299 | 0.339 | 3.78E-01 | -0.964 | 0.366 |
| | ***apoeb*** | 0.191 | 0.190 | 3.14E-01 | -0.181 | 0.563 | 0.271 | 0.230 | 2.40E-01 | -0.181 | 0.722 | 0.311 | 0.247 | 2.07E-01 | -0.173 | 0.795 |
| | ***ldlra*** | 0.200 | 0.174 | 2.51E-01 | -0.142 | 0.542 | 0.105 | 0.193 | 5.88E-01 | -0.274 | 0.483 | 0.088 | 0.199 | 6.59E-01 | -0.303 | 0.479 |
| | **body length (in SD)** | - | - | - | - | - | 0.704 | 0.296 | 1.74E-02 | 0.124 | 1.284 | 0.549 | 0.337 | 1.03E-01 | -0.111 | 1.210 |
| | **dorsal body surface area (in SD)** | - | - | - | - | - | 0.510 | 0.185 | 5.93E-03 | 0.147 | 0.874 | 0.425 | 0.201 | 3.44E-02 | 0.031 | 0.819 |
| | **LDL cholesterol levels (in SD)** | - | - | - | - | - | - | - | - | - | - | -0.051 | 0.135 | 7.09E-01 | -0.316 | 0.215 |
| | **HDL cholesterol levels (in SD)** | - | - | - | - | - | - | - | - | - | - | -0.147 | 0.227 | 5.16E-01 | -0.593 | 0.298 |
| | **triglyceride levels (in SD)** | - | - | - | - | - | - | - | - | - | - | 0.317 | 0.203 | 1.19E-01 | -0.081 | 0.715 |
| | **glucose levels (in SD)** | - | - | - | - | - | - | - | - | - | - | -0.060 | 0.140 | 6.69E-01 | -0.333 | 0.214 |
| | **time of day (in hours since 9AM)** | -0.075 | 0.114 | 5.09E-01 | -0.299 | 0.148 | -0.155 | 0.148 | 2.96E-01 | -0.445 | 0.136 | -0.066 | 0.171 | 6.98E-01 | -0.402 | 0.270 |
| | **batch 2** | -1.216 | 0.674 | 7.13E-02 | -2.538 | 0.106 | -0.149 | 0.900 | 8.68E-01 | -1.913 | 1.615 | -0.215 | 0.960 | 8.23E-01 | -2.096 | 1.666 |
| | **batch 3** | -1.276 | 0.499 | 1.06E-02 | -2.255 | -0.297 | -0.998 | 0.595 | 9.38E-02 | -2.164 | 0.169 | -1.209 | 0.651 | 6.32E-02 | -2.484 | 0.066 |
| | **batch 4** | -0.845 | 0.467 | 7.07E-02 | -1.761 | 0.071 | -0.047 | 0.624 | 9.40E-01 | -1.269 | 1.175 | -0.399 | 0.722 | 5.80E-01 | -1.815 | 1.016 |
| | **batch 5** | -0.211 | 0.387 | 5.85E-01 | -0.970 | 0.547 | -0.011 | 0.561 | 9.85E-01 | -1.111 | 1.090 | -0.654 | 0.712 | 3.59E-01 | -2.050 | 0.743 |
| | **batch 6** | -0.162 | 0.381 | 6.70E-01 | -0.910 | 0.585 | -0.524 | 0.419 | 2.11E-01 | -1.345 | 0.297 | -1.131 | 0.595 | 5.72E-02 | -2.297 | 0.034 |
| | **batch 7** | -1.058 | 0.462 | 2.19E-02 | -1.963 | -0.153 | -1.057 | 0.544 | 5.19E-02 | -2.122 | 0.009 | -1.729 | 0.830 | 3.72E-02 | -3.355 | -0.103 |
| | **intercept** | 183.351 | 148.640 | 2.17E-01 | -107.977 | 474.679 | 174.210 | 171.135 | 3.09E-01 | -161.209 | 509.629 | 131.729 | 171.727 | 4.43E-01 | -204.849 | 468.307 |
| | ***apoba*** | 0.954 | 0.808 | 2.37E-01 | -0.629 | 2.537 | 0.769 | 0.923 | 4.05E-01 | -1.040 | 2.577 | 0.695 | 0.964 | 4.71E-01 | -1.195 | 2.584 |
| | ***apobb.2*** | -0.031 | 0.473 | 9.48E-01 | -0.958 | 0.895 | -0.008 | 0.509 | 9.88E-01 | -1.006 | 0.990 | 0.307 | 0.554 | 5.80E-01 | -0.779 | 1.394 |
| | ***ldlrb*** | -450.568 | 372.285 | 2.26E-01 | -1200.000 | 279.096 | -427.256 | 428.905 | 3.19E-01 | -1300.000 | 413.383 | -321.569 | 430.563 | 4.55E-01 | -1200.000 | 522.320 |

| | | **Vascular infiltration by macrophages** | | | | | | | | | | | | | | |
|---|---|---|---|---|---|---|---|---|---|---|---|---|---|---|---|---|
| | | **26 vs. 143 larvae with 2 vs. 0 mutated alleles (n=169)** | | | | | **26 vs. 126 larvae with 2 vs. 0 mutated alleles (n=152)** | | | | | **26 vs. 126 larvae with 2 vs. 0 mutated alleles (n=152)** | | | | |
| | | **Effect** | **SE** | ***P*** | **lci** | **uci** | **Effect** | **SE** | ***P*** | **lci** | **uci** | **Effect** | **SE** | ***P*** | **lci** | **uci** |
| **fixed factors** | **2 vs. 0 mutated alleles** | -0.270 | 0.214 | 2.08E-01 | -0.690 | 0.150 | -0.300 | 0.255 | 2.39E-01 | -0.799 | 0.199 | -0.092 | 0.264 | 7.28E-01 | -0.609 | 0.426 |
| | ***apoea*** | -0.110 | 0.122 | 3.66E-01 | -0.350 | 0.129 | -0.178 | 0.135 | 1.89E-01 | -0.443 | 0.087 | -0.229 | 0.134 | 8.83E-02 | -0.492 | 0.034 |
| | ***apoeb*** | -0.089 | 0.113 | 4.30E-01 | -0.311 | 0.132 | -0.098 | 0.125 | 4.35E-01 | -0.343 | 0.148 | -0.143 | 0.125 | 2.54E-01 | -0.387 | 0.102 |
| | ***ldlra*** | 0.067 | 0.112 | 5.47E-01 | -0.152 | 0.286 | 0.086 | 0.122 | 4.79E-01 | -0.153 | 0.326 | 0.104 | 0.120 | 3.87E-01 | -0.132 | 0.340 |
| | **body length (in SD)** | - | - | - | - | - | -0.046 | 0.121 | 7.06E-01 | -0.284 | 0.192 | 0.010 | 0.123 | 9.36E-01 | -0.231 | 0.251 |
| | **dorsal body surface area (in SD)** | - | - | - | - | - | -0.194 | 0.098 | 4.73E-02 | -0.386 | -0.002 | -0.157 | 0.100 | 1.16E-01 | -0.352 | 0.039 |
| | **LDL cholesterol levels (in SD)** | - | - | - | - | - | - | - | - | - | - | 0.120 | 0.085 | 1.57E-01 | -0.046 | 0.286 |
| | **HDL cholesterol levels (in SD)** | - | - | - | - | - | - | - | - | - | - | 0.055 | 0.101 | 5.87E-01 | -0.144 | 0.254 |
| | **triglyceride levels (in SD)** | - | - | - | - | - | - | - | - | - | - | -0.258 | 0.117 | 2.79E-02 | -0.488 | -0.028 |
| | **glucose levels (in SD)** | - | - | - | - | - | - | - | - | - | - | 0.063 | 0.088 | 4.76E-01 | -0.110 | 0.236 |
| | **time of day (in hours since 9AM)** | 0.103 | 0.063 | 1.01E-01 | -0.020 | 0.225 | 0.117 | 0.069 | 9.19E-02 | -0.019 | 0.253 | 0.068 | 0.069 | 3.23E-01 | -0.067 | 0.203 |
| | **intercept_random** | 11.258 | 88.626 | 8.99E-01 | -162.446 | 184.962 | -2.079 | 99.580 | 9.83E-01 | -197.253 | 193.094 | 1.267 | 98.915 | 9.90E-01 | -192.603 | 195.136 |
| | ***apoba*** | -0.485 | 0.450 | 2.81E-01 | -1.367 | 0.397 | -0.542 | 0.478 | 2.58E-01 | -1.479 | 0.396 | -0.539 | 0.470 | 2.51E-01 | -1.460 | 0.382 |
| | ***apobb.2*** | -0.189 | 0.269 | 4.82E-01 | -0.715 | 0.337 | -0.389 | 0.303 | 1.99E-01 | -0.983 | 0.204 | -0.606 | 0.314 | 5.36E-02 | -1.221 | 0.010 |
| | ***ldlrb*** | -24.951 | 221.876 | 9.10E-01 | -459.821 | 409.919 | 9.325 | 249.131 | 9.70E-01 | -478.963 | 497.613 | 2.196 | 247.481 | 9.93E-01 | -482.857 | 487.250 |
| **random factors** | ***variance by batch*** | *0.574* | *0.172* | - | *0.319* | *1.031* | *0.490* | *0.173* | - | *0.245* | *0.980* | *0.366* | *0.143* | - | *0.170* | *0.787* |
| | ***residual*** | *0.899* | *0.050* | - | *0.805* | *1.003* | *0.931* | *0.055* | - | *0.828* | *1.046* | *0.918* | *0.055* | - | *0.817* | *1.031* |

*continued* **Supplementary Table 21**

| | | Vascular co-localization of lipids with macrophages | | | | | | | | | | | | | | |
|---|---|---|---|---|---|---|---|---|---|---|---|---|---|---|---|---|
| | | **20 vs. 108 larvae with 2 vs. 0 mutated alleles (n=128)** | | | | | **20 vs. 92 larvae with 2 vs. 0 mutated alleles (n=112)** | | | | | **20 vs. 92 larvae with 2 vs. 0 mutated alleles (n=112)** | | | | |
| | | **Effect** | **SE** | ***P*** | **lci** | **uci** | **Effect** | **SE** | ***P*** | **lci** | **uci** | **Effect** | **SE** | ***P*** | **lci** | **uci** |
| **negative binomial terms** | **2 vs. 0 mutated alleles** | 2.295 | 0.547 | 2.72E-05 | 1.223 | 3.367 | 4.048 | 1.068 | 1.50E-04 | 1.956 | 6.141 | 3.351 | 1.013 | 9.44E-04 | 1.365 | 5.337 |
| | ***apoea*** | -0.269 | 0.419 | 5.21E-01 | -1.089 | 0.552 | -0.374 | 0.514 | 4.66E-01 | -1.381 | 0.632 | -0.199 | 0.538 | 7.12E-01 | -1.253 | 0.856 |
| | ***apoeb*** | -0.236 | 0.256 | 3.57E-01 | -0.738 | 0.266 | -0.311 | 0.283 | 2.72E-01 | -0.867 | 0.244 | -0.408 | 0.276 | 1.40E-01 | -0.950 | 0.133 |
| | ***ldlra*** | 0.360 | 0.208 | 8.28E-02 | -0.047 | 0.767 | 0.256 | 0.220 | 2.45E-01 | -0.175 | 0.686 | 0.408 | 0.233 | 7.91E-02 | -0.047 | 0.864 |
| | **body length (in SD)** | - | - | - | - | - | 1.231 | 0.470 | 8.75E-03 | 0.311 | 2.151 | 1.121 | 0.485 | 2.08E-02 | 0.170 | 2.071 |
| | **dorsal body surface area (in SD)** | - | - | - | - | - | 0.847 | 0.259 | 1.07E-03 | 0.339 | 1.354 | 0.792 | 0.300 | 8.21E-03 | 0.205 | 1.380 |
| | **LDL cholesterol levels (in SD)** | - | - | - | - | - | - | - | - | - | - | -0.363 | 0.280 | 1.95E-01 | -0.911 | 0.186 |
| | **HDL cholesterol levels (in SD)** | - | - | - | - | - | - | - | - | - | - | -0.033 | 0.288 | 9.10E-01 | -0.597 | 0.532 |
| | **triglyceride levels (in SD)** | - | - | - | - | - | - | - | - | - | - | 0.096 | 0.427 | 8.22E-01 | -0.741 | 0.932 |
| | **glucose levels (in SD)** | - | - | - | - | - | - | - | - | - | - | -0.232 | 0.248 | 3.50E-01 | -0.718 | 0.254 |
| | **time of day (in hours since 9AM)** | 0.215 | 0.148 | 1.48E-01 | -0.076 | 0.505 | 0.313 | 0.164 | 5.67E-02 | -0.009 | 0.634 | 0.437 | 0.176 | 1.29E-02 | 0.092 | 0.782 |
| | **batch 2** | 0.570 | 1.135 | 6.16E-01 | -1.655 | 2.795 | 1.855 | 1.345 | 1.68E-01 | -0.782 | 4.491 | 1.208 | 1.298 | 3.52E-01 | -1.337 | 3.753 |
| | **batch 3** | -1.739 | 0.589 | 3.14E-03 | -2.893 | -0.585 | -1.631 | 0.721 | 2.37E-02 | -3.045 | -0.218 | -1.931 | 0.799 | 1.57E-02 | -3.498 | -0.364 |
| | **batch 4** | -0.373 | 0.601 | 5.35E-01 | -1.550 | 0.805 | 0.295 | 0.854 | 7.30E-01 | -1.379 | 1.968 | 0.269 | 0.934 | 7.73E-01 | -1.561 | 2.099 |
| | **batch 5** | -0.139 | 0.595 | 8.16E-01 | -1.306 | 1.028 | -0.859 | 0.762 | 2.60E-01 | -2.353 | 0.635 | -0.795 | 0.879 | 3.66E-01 | -2.518 | 0.928 |
| | **batch 6** | -0.716 | 0.577 | 2.14E-01 | -1.846 | 0.414 | -2.288 | 0.702 | 1.12E-03 | -3.664 | -0.912 | -2.405 | 1.136 | 3.43E-02 | -4.631 | -0.178 |
| | **batch 7** | -0.845 | 0.788 | 2.84E-01 | -2.390 | 0.701 | -2.164 | 0.826 | 8.77E-03 | -3.782 | -0.546 | -2.372 | 1.171 | 4.27E-02 | -4.667 | -0.078 |
| | **intercept** | 108.949 | 133.134 | 4.13E-01 | -151.988 | 369.886 | 105.728 | 213.116 | 6.20E-01 | -311.971 | 523.427 | 251.845 | 259.958 | 3.33E-01 | -257.664 | 761.353 |
| | ***apoba*** | -0.919 | 1.185 | 4.38E-01 | -3.242 | 1.405 | -0.768 | 1.116 | 4.92E-01 | -2.956 | 1.420 | -0.610 | 1.095 | 5.77E-01 | -2.757 | 1.536 |
| | ***apobb.2*** | 0.031 | 0.591 | 9.59E-01 | -1.127 | 1.188 | 0.132 | 0.639 | 8.36E-01 | -1.121 | 1.385 | 0.210 | 0.725 | 7.73E-01 | -1.212 | 1.631 |
| | ***ldlrb*** | -263.818 | 334.248 | 4.30E-01 | -918.933 | 391.296 | -256.522 | 532.363 | 6.30E-01 | -1300.000 | 786.891 | -623.857 | 649.781 | 3.37E-01 | -1900.000 | 649.691 |

| | | Vascular infiltration by neutrophils | | | | | | | | | | | | | | |
|---|---|---|---|---|---|---|---|---|---|---|---|---|---|---|---|---|
| | | **26 vs. 147 larvae with 2 vs. 0 mutated alleles (n=173)** | | | | | **26 vs. 129 larvae with 2 vs. 0 mutated alleles (n=155)** | | | | | **26 vs. 129 larvae with 2 vs. 0 mutated alleles (n=155)** | | | | |
| | | **Effect** | **SE** | ***P*** | **lci** | **uci** | **Effect** | **SE** | ***P*** | **lci** | **uci** | **Effect** | **SE** | ***P*** | **lci** | **uci** |
| **fixed factors** | **2 vs. 0 mutated alleles** | 0.428 | 0.206 | 3.77E-02 | 0.024 | 0.832 | 0.281 | 0.235 | 2.31E-01 | -0.179 | 0.742 | 0.466 | 0.245 | 5.72E-02 | -0.014 | 0.947 |
| | ***apoea*** | -0.152 | 0.116 | 1.92E-01 | -0.380 | 0.076 | -0.217 | 0.124 | 8.05E-02 | -0.460 | 0.026 | -0.236 | 0.124 | 5.78E-02 | -0.479 | 0.008 |
| | ***apoeb*** | 0.008 | 0.108 | 9.40E-01 | -0.204 | 0.220 | 0.021 | 0.115 | 8.58E-01 | -0.205 | 0.247 | -0.014 | 0.116 | 9.05E-01 | -0.241 | 0.213 |
| | ***ldlra*** | -0.088 | 0.107 | 4.10E-01 | -0.297 | 0.121 | -0.042 | 0.112 | 7.08E-01 | -0.261 | 0.177 | -0.019 | 0.111 | 8.62E-01 | -0.237 | 0.198 |
| | **body length (in SD)** | - | - | - | - | - | -0.112 | 0.110 | 3.08E-01 | -0.328 | 0.103 | -0.061 | 0.112 | 5.89E-01 | -0.280 | 0.159 |
| | **dorsal body surface area (in SD)** | - | - | - | - | - | 0.040 | 0.089 | 6.53E-01 | -0.135 | 0.215 | 0.074 | 0.092 | 4.19E-01 | -0.105 | 0.253 |
| | **LDL cholesterol levels (in SD)** | - | - | - | - | - | - | - | - | - | - | 0.054 | 0.076 | 4.76E-01 | -0.095 | 0.204 |
| | **HDL cholesterol levels (in SD)** | - | - | - | - | - | - | - | - | - | - | 0.049 | 0.094 | 6.02E-01 | -0.135 | 0.232 |
| | **triglyceride levels (in SD)** | - | - | - | - | - | - | - | - | - | - | -0.186 | 0.107 | 8.18E-02 | -0.395 | 0.023 |
| | **glucose levels (in SD)** | - | - | - | - | - | - | - | - | - | - | 0.122 | 0.082 | 1.34E-01 | -0.038 | 0.282 |
| | **time of day (in hours since 9AM)** | 0.029 | 0.059 | 6.29E-01 | -0.088 | 0.145 | 0.017 | 0.064 | 7.95E-01 | -0.108 | 0.141 | -0.015 | 0.064 | 8.14E-01 | -0.140 | 0.110 |
| | **intercept_random** | -182.126 | 85.408 | 3.30E-02 | -349.522 | -14.729 | -203.375 | 92.181 | 2.74E-02 | -384.047 | -22.702 | -198.131 | 92.230 | 3.17E-02 | -378.899 | -17.363 |
| | ***apoba*** | 0.233 | 0.431 | 5.89E-01 | -0.612 | 1.078 | 0.169 | 0.441 | 7.02E-01 | -0.695 | 1.033 | 0.163 | 0.436 | 7.09E-01 | -0.693 | 1.018 |
| | ***apobb.2*** | 0.004 | 0.259 | 9.87E-01 | -0.503 | 0.511 | 0.142 | 0.280 | 6.12E-01 | -0.407 | 0.692 | 0.045 | 0.292 | 8.77E-01 | -0.527 | 0.617 |
| | ***ldlrb*** | 454.749 | 213.818 | 3.34E-02 | 35.674 | 873.823 | 507.788 | 230.619 | 2.77E-02 | 55.782 | 959.793 | 495.311 | 230.758 | 3.18E-02 | 43.035 | 947.588 |
| **random factors** | ***variance by batch*** | *0.468* | *0.142* | - | *0.258* | *0.850* | *0.407* | *0.140* | - | *0.207* | *0.798* | *0.308* | *0.126* | - | *0.138* | *0.686* |
| | ***residual*** | *0.867* | *0.048* | - | *0.778* | *0.966* | *0.862* | *0.051* | - | *0.769* | *0.967* | *0.856* | *0.050* | - | *0.763* | *0.961* |

*continued* **Supplementary Table 21**

**Vascular co-localization of lipids with neutrophils**

| | | 19 vs. 106 larvae with 2 vs. 0 mutated alleles (n=125) | | | | | 19 vs. 89 larvae with 2 vs. 0 mutated alleles (n=108) | | | | | 19 vs. 89 larvae with 2 vs. 0 mutated alleles (n=108) | | | | |
|---|---|---|---|---|---|---|---|---|---|---|---|---|---|---|---|---|
| | | **Effect** | **SE** | ***P*** | **lci** | **uci** | **Effect** | **SE** | ***P*** | **lci** | **uci** | **Effect** | **SE** | ***P*** | **lci** | **uci** |
| **negative binomial terms** | **2 vs. 0 mutated alleles** | 4.082 | 0.541 | 4.44E-14 | 3.022 | 5.142 | 6.315 | 1.281 | 8.31E-07 | 3.803 | 8.826 | 7.962 | 1.975 | 5.54E-05 | 4.091 | 11.833 |
| | ***apoea*** | -1.717 | 0.466 | 2.28E-04 | -2.630 | -0.804 | -1.992 | 0.694 | 4.13E-03 | -3.353 | -0.631 | -1.881 | 0.797 | 1.82E-02 | -3.443 | -0.320 |
| | ***apoeb*** | -0.611 | 0.382 | 1.09E-01 | -1.359 | 0.137 | -0.545 | 0.503 | 2.79E-01 | -1.531 | 0.441 | 0.150 | 0.569 | 7.92E-01 | -0.964 | 1.264 |
| | ***ldlra*** | 0.031 | 0.290 | 9.15E-01 | -0.538 | 0.599 | 0.112 | 0.288 | 6.98E-01 | -0.453 | 0.677 | -0.199 | 0.375 | 5.96E-01 | -0.933 | 0.536 |
| | **body length (in SD)** | - | - | - | - | - | 1.422 | 0.684 | 3.75E-02 | 0.082 | 2.763 | 1.521 | 0.820 | 6.37E-02 | -0.087 | 3.128 |
| | **dorsal body surface area (in SD)** | - | - | - | - | - | -0.259 | 0.340 | 4.46E-01 | -0.925 | 0.407 | -0.435 | 0.374 | 2.45E-01 | -1.169 | 0.298 |
| | **LDL cholesterol levels (in SD)** | - | - | - | - | - | - | - | - | - | - | 0.411 | 0.337 | 2.22E-01 | -0.249 | 1.071 |
| | **HDL cholesterol levels (in SD)** | - | - | - | - | - | - | - | - | - | - | -0.177 | 0.558 | 7.51E-01 | -1.272 | 0.917 |
| | **triglyceride levels (in SD)** | - | - | - | - | - | - | - | - | - | - | -0.144 | 0.616 | 8.15E-01 | -1.352 | 1.063 |
| | **glucose levels (in SD)** | - | - | - | - | - | - | - | - | - | - | 1.340 | 0.467 | 4.13E-03 | 0.424 | 2.256 |
| | **time of day (in hours since 9AM)** | -0.506 | 0.180 | 5.00E-03 | -0.860 | -0.153 | -0.598 | 0.315 | 5.76E-02 | -1.215 | 0.019 | -0.547 | 0.350 | 1.19E-01 | -1.233 | 0.140 |
| | **batch 2** | -3.868 | 1.152 | 7.90E-04 | -6.126 | -1.609 | -3.200 | 1.213 | 8.33E-03 | -5.577 | -0.823 | -0.841 | 1.754 | 6.32E-01 | -4.278 | 2.596 |
| | **batch 3** | -1.358 | 0.854 | 1.12E-01 | -3.033 | 0.316 | -1.330 | 0.923 | 1.49E-01 | -3.138 | 0.478 | -1.396 | 1.122 | 2.14E-01 | -3.596 | 0.804 |
| | **batch 4** | -1.785 | 0.808 | 2.72E-02 | -3.369 | -0.201 | -1.256 | 0.998 | 2.08E-01 | -3.212 | 0.700 | -1.802 | 1.075 | 9.37E-02 | -3.909 | 0.305 |
| | **batch 5** | -1.321 | 0.655 | 4.36E-02 | -2.604 | -0.038 | -1.559 | 0.883 | 7.76E-02 | -3.290 | 0.173 | -1.916 | 1.500 | 2.02E-01 | -4.855 | 1.024 |
| | **batch 6** | -3.031 | 0.662 | 4.74E-06 | -4.329 | -1.733 | -2.775 | 0.935 | 2.99E-03 | -4.607 | -0.943 | -2.871 | 1.403 | 4.08E-02 | -5.621 | -0.121 |
| | **intercept** | -214.663 | 420.786 | 6.10E-01 | -1000.000 | 610.062 | -472.077 | 522.037 | 3.66E-01 | -1500.000 | 551.098 | -942.198 | 584.076 | 1.07E-01 | -2100.000 | 202.571 |
| | ***apoba*** | 3.615 | 1.392 | 9.41E-03 | 0.886 | 6.343 | 3.721 | 1.556 | 1.68E-02 | 0.671 | 6.771 | 4.649 | 2.299 | 4.31E-02 | 0.144 | 9.154 |
| | ***apobb.2*** | 0.569 | 0.805 | 4.79E-01 | -1.008 | 2.146 | 0.248 | 0.958 | 7.96E-01 | -1.629 | 2.125 | 1.111 | 1.180 | 3.46E-01 | -1.202 | 3.425 |
| | ***ldlrb*** | 530.344 | 1051.605 | 6.14E-01 | -1500.000 | 2591.452 | 1174.217 | 1307.275 | 3.69E-01 | -1400.000 | 3736.429 | 2338.624 | 1464.591 | 1.10E-01 | -531.921 | 5209.169 |

**Vascular co-localization of macrophages with neutrophils**

| | | 26 vs. 143 larvae with 2 vs. 0 mutated alleles (n=169) | | | | | 26 vs. 126 larvae with 2 vs. 0 mutated alleles (n=152) | | | | | 26 vs. 126 larvae with 2 vs. 0 mutated alleles (n=152) | | | | |
|---|---|---|---|---|---|---|---|---|---|---|---|---|---|---|---|---|
| | | **Effect** | **SE** | ***P*** | **lci** | **uci** | **Effect** | **SE** | ***P*** | **lci** | **uci** | **Effect** | **SE** | ***P*** | **lci** | **uci** |
| **negative binomial terms** | **2 vs. 0 mutated alleles** | 0.080 | 0.279 | 7.73E-01 | -0.467 | 0.628 | 0.002 | 0.312 | 9.96E-01 | -0.609 | 0.613 | 0.101 | 0.328 | 7.58E-01 | -0.542 | 0.743 |
| | ***apoea*** | -0.230 | 0.158 | 1.45E-01 | -0.540 | 0.080 | -0.287 | 0.172 | 9.53E-02 | -0.624 | 0.050 | -0.435 | 0.181 | 1.63E-02 | -0.790 | -0.080 |
| | ***apoeb*** | -0.011 | 0.149 | 9.41E-01 | -0.302 | 0.280 | -0.040 | 0.162 | 8.03E-01 | -0.359 | 0.278 | -0.066 | 0.156 | 6.72E-01 | -0.371 | 0.239 |
| | ***ldlra*** | -0.041 | 0.137 | 7.66E-01 | -0.310 | 0.228 | -0.042 | 0.132 | 7.52E-01 | -0.301 | 0.218 | 0.047 | 0.132 | 7.21E-01 | -0.212 | 0.307 |
| | **body length (in SD)** | - | - | - | - | - | -0.026 | 0.165 | 8.73E-01 | -0.350 | 0.297 | -0.019 | 0.201 | 9.23E-01 | -0.414 | 0.375 |
| | **dorsal body surface area (in SD)** | - | - | - | - | - | 0.123 | 0.142 | 3.86E-01 | -0.155 | 0.401 | 0.103 | 0.150 | 4.92E-01 | -0.191 | 0.397 |
| | **LDL cholesterol levels (in SD)** | - | - | - | - | - | - | - | - | - | - | 0.012 | 0.143 | 9.35E-01 | -0.268 | 0.292 |
| | **HDL cholesterol levels (in SD)** | - | - | - | - | - | - | - | - | - | - | -0.166 | 0.147 | 2.60E-01 | -0.454 | 0.122 |
| | **triglyceride levels (in SD)** | - | - | - | - | - | - | - | - | - | - | -0.284 | 0.219 | 1.95E-01 | -0.714 | 0.145 |
| | **glucose levels (in SD)** | - | - | - | - | - | - | - | - | - | - | 0.304 | 0.112 | 6.68E-03 | 0.084 | 0.524 |
| | **time of day (in hours since 9AM)** | 0.253 | 0.079 | 1.25E-03 | 0.099 | 0.407 | 0.235 | 0.086 | 6.31E-03 | 0.066 | 0.404 | 0.185 | 0.093 | 4.62E-02 | 0.003 | 0.367 |
| | **batch 1** | -0.542 | 0.505 | 2.83E-01 | -1.533 | 0.448 | -0.413 | 0.576 | 4.74E-01 | -1.542 | 0.717 | -0.276 | 0.782 | 7.24E-01 | -1.809 | 1.257 |
| | **batch 2** | -0.959 | 0.494 | 5.22E-02 | -1.927 | 0.009 | -1.027 | 0.525 | 5.05E-02 | -2.057 | 0.002 | -0.697 | 0.619 | 2.60E-01 | -1.910 | 0.515 |
| | **batch 3** | -1.403 | 0.500 | 4.99E-03 | -2.383 | -0.424 | -1.360 | 0.508 | 7.39E-03 | -2.354 | -0.365 | -1.238 | 0.639 | 5.27E-02 | -2.489 | 0.014 |
| | **batch 4** | 0.258 | 0.523 | 6.23E-01 | -0.768 | 1.284 | 0.275 | 0.520 | 5.96E-01 | -0.743 | 1.294 | 0.232 | 0.632 | 7.13E-01 | -1.007 | 1.471 |
| | **batch 5** | -0.934 | 0.444 | 3.53E-02 | -1.804 | -0.064 | -1.058 | 0.501 | 3.46E-02 | -2.040 | -0.076 | -0.619 | 0.603 | 3.04E-01 | -1.800 | 0.562 |
| | **batch 6** | -2.488 | 0.504 | 7.97E-07 | -3.476 | -1.500 | -2.571 | 0.611 | 2.54E-05 | -3.767 | -1.374 | -2.227 | 0.664 | 8.01E-04 | -3.529 | -0.925 |
| | **batch 7** | -2.134 | 0.793 | 7.15E-03 | -3.689 | -0.579 | -2.135 | 0.849 | 1.19E-02 | -3.799 | -0.471 | -2.588 | 1.027 | 1.17E-02 | -4.600 | -0.575 |
| | **intercept** | -185.934 | 104.046 | 7.39E-02 | -389.861 | 17.993 | -142.067 | 123.760 | 2.51E-01 | -384.633 | 100.498 | -147.810 | 126.894 | 2.44E-01 | -396.517 | 100.898 |
| | ***apoba*** | -0.090 | 0.607 | 8.82E-01 | -1.281 | 1.100 | 0.110 | 0.598 | 8.54E-01 | -1.063 | 1.282 | 0.303 | 0.577 | 5.99E-01 | -0.827 | 1.434 |
| | ***apobb.2*** | -0.339 | 0.366 | 3.55E-01 | -1.056 | 0.379 | -0.520 | 0.407 | 2.02E-01 | -1.318 | 0.279 | -0.419 | 0.388 | 2.79E-01 | -1.179 | 0.340 |
| | ***ldlrb*** | 477.129 | 261.076 | 6.76E-02 | -34.570 | 988.829 | 367.600 | 310.089 | 2.36E-01 | -240.163 | 975.364 | 380.686 | 317.971 | 2.31E-01 | -242.525 | 1003.897 |

***continued*** **Supplementary Table 21**

| | | ***ldlra*** | | | | | | | | | | | | | | |
|---|---|---|---|---|---|---|---|---|---|---|---|---|---|---|---|---|
| | | **Vascular lipid deposition** | | | | | | | | | | | | | | |
| | | **Model 1** | | | | | **Model 2** | | | | | **Model 3** | | | | |
| | | **130 vs. 90 larvae with 2 vs. 0 mutated alleles (n=220)** | | | | | **117 vs. 80 larvae with 2 vs. 0 mutated alleles (n=197)** | | | | | **117 vs. 80 larvae with 2 vs. 0 mutated alleles (n=197)** | | | | |
| | | **Effect** | **SE** | ***P*** | **lci** | **uci** | **Effect** | **SE** | ***P*** | **lci** | **uci** | **Effect** | **SE** | ***P*** | **lci** | **uci** |
| **negative binomial terms** | **2 vs. 0 mutated alleles** | 0.111 | 0.251 | 6.58E-01 | -0.381 | 0.603 | 0.104 | 0.266 | 6.97E-01 | -0.418 | 0.626 | 0.073 | 0.275 | 7.91E-01 | -0.466 | 0.612 |
| | ***apoea*** | -0.189 | 0.195 | 3.32E-01 | -0.571 | 0.193 | -0.183 | 0.212 | 3.89E-01 | -0.598 | 0.233 | -0.172 | 0.209 | 4.10E-01 | -0.582 | 0.238 |
| | ***apoeb*** | 0.308 | 0.164 | 6.09E-02 | -0.014 | 0.629 | 0.394 | 0.186 | 3.46E-02 | 0.029 | 0.759 | 0.392 | 0.191 | 4.03E-02 | 0.017 | 0.768 |
| | ***apobb.1*** | 0.996 | 0.161 | 5.70E-10 | 0.681 | 1.310 | 0.937 | 0.198 | 2.13E-06 | 0.549 | 1.324 | 0.898 | 0.210 | 1.91E-05 | 0.486 | 1.310 |
| | **body length (in SD)** | - | - | - | - | - | -0.071 | 0.167 | 6.70E-01 | -0.398 | 0.256 | -0.041 | 0.175 | 8.16E-01 | -0.384 | 0.302 |
| | **dorsal body surface area (in SD)** | - | - | - | - | - | 0.197 | 0.141 | 1.63E-01 | -0.080 | 0.474 | 0.213 | 0.145 | 1.43E-01 | -0.072 | 0.498 |
| | **LDL cholesterol levels (in SD)** | - | - | - | - | - | - | - | - | - | - | -0.015 | 0.107 | 8.85E-01 | -0.225 | 0.194 |
| | **HDL cholesterol levels (in SD)** | - | - | - | - | - | - | - | - | - | - | -0.045 | 0.144 | 7.53E-01 | -0.327 | 0.237 |
| | **triglyceride levels (in SD)** | - | - | - | - | - | - | - | - | - | - | 0.140 | 0.178 | 4.33E-01 | -0.209 | 0.489 |
| | **glucose levels (in SD)** | - | - | - | - | - | - | - | - | - | - | -0.042 | 0.121 | 7.30E-01 | -0.279 | 0.195 |
| | **time of day (in hours since 9AM)** | -0.062 | 0.094 | 5.07E-01 | -0.246 | 0.122 | -0.107 | 0.117 | 3.60E-01 | -0.335 | 0.122 | -0.086 | 0.118 | 4.66E-01 | -0.318 | 0.146 |
| | **batch 2** | -1.659 | 0.483 | 5.94E-04 | -2.606 | -0.712 | -1.871 | 0.575 | 1.13E-03 | -2.998 | -0.745 | -1.705 | 0.605 | 4.79E-03 | -2.890 | -0.520 |
| | **batch 3** | -0.467 | 0.382 | 2.21E-01 | -1.216 | 0.282 | -0.569 | 0.399 | 1.54E-01 | -1.351 | 0.213 | -0.644 | 0.442 | 1.45E-01 | -1.509 | 0.222 |
| | **batch 4** | -0.714 | 0.416 | 8.63E-02 | -1.529 | 0.102 | -0.820 | 0.449 | 6.77E-02 | -1.699 | 0.060 | -0.818 | 0.467 | 8.02E-02 | -1.734 | 0.098 |
| | **batch 5** | 0.372 | 0.321 | 2.46E-01 | -0.257 | 1.001 | 0.208 | 0.367 | 5.72E-01 | -0.512 | 0.928 | -0.019 | 0.503 | 9.70E-01 | -1.005 | 0.967 |
| | **batch 6** | 0.156 | 0.281 | 5.80E-01 | -0.395 | 0.707 | 0.092 | 0.318 | 7.72E-01 | -0.531 | 0.716 | -0.232 | 0.535 | 6.64E-01 | -1.281 | 0.816 |
| | **batch 7** | -0.554 | 0.363 | 1.27E-01 | -1.266 | 0.157 | -0.730 | 0.377 | 5.27E-02 | -1.468 | 0.008 | -0.959 | 0.554 | 8.37E-02 | -2.046 | 0.128 |
| | **intercept** | 280.829 | 147.422 | 5.68E-02 | -8.113 | 569.772 | 320.998 | 153.007 | 3.59E-02 | 21.110 | 620.886 | 319.158 | 154.393 | 3.87E-02 | 16.554 | 621.762 |
| | ***apoba*** | -0.166 | 0.630 | 7.92E-01 | -1.401 | 1.069 | -0.357 | 0.644 | 5.79E-01 | -1.619 | 0.905 | -0.211 | 0.668 | 7.52E-01 | -1.520 | 1.098 |
| | ***apobb.2*** | -0.046 | 0.404 | 9.10E-01 | -0.837 | 0.746 | -0.138 | 0.409 | 7.36E-01 | -0.940 | 0.665 | -0.060 | 0.419 | 8.86E-01 | -0.881 | 0.761 |
| | ***ldlrb*** | -690.958 | 368.791 | 6.10E-02 | -1400.000 | 31.859 | -789.795 | 382.814 | 3.91E-02 | -1500.000 | -39.493 | -785.887 | 386.208 | 4.19E-02 | -1500.000 | -28.933 |

| | | **Vascular infiltration by macrophages** | | | | | | | | | | | | | | |
|---|---|---|---|---|---|---|---|---|---|---|---|---|---|---|---|---|
| | | **174 vs. 117 larvae with 2 vs. 0 mutated alleles (n=291)** | | | | | **158 vs. 102 larvae with 2 vs. 0 mutated alleles (n=260)** | | | | | **158 vs. 102 larvae with 2 vs. 0 mutated alleles (n=260)** | | | | |
| | | **Effect** | **SE** | ***P*** | **lci** | **uci** | **Effect** | **SE** | ***P*** | **lci** | **uci** | **Effect** | **SE** | ***P*** | **lci** | **uci** |
| **fixed factors** | **2 vs. 0 mutated alleles** | 0.077 | 0.136 | 5.70E-01 | -0.189 | 0.343 | 0.029 | 0.152 | 8.50E-01 | -0.269 | 0.327 | 0.000 | 0.152 | 1.00E+00 | -0.297 | 0.297 |
| | ***apoea*** | -0.126 | 0.087 | 1.48E-01 | -0.297 | 0.045 | -0.184 | 0.094 | 5.13E-02 | -0.369 | 0.001 | -0.197 | 0.094 | 3.66E-02 | -0.382 | -0.012 |
| | ***apoeb*** | -0.181 | 0.090 | 4.53E-02 | -0.357 | -0.004 | -0.170 | 0.099 | 8.64E-02 | -0.365 | 0.024 | -0.167 | 0.099 | 9.21E-02 | -0.362 | 0.027 |
| | ***apobb.1*** | -0.190 | 0.096 | 4.64E-02 | -0.378 | -0.003 | -0.186 | 0.106 | 7.95E-02 | -0.393 | 0.022 | -0.207 | 0.107 | 5.29E-02 | -0.417 | 0.003 |
| | **body length (in SD)** | - | - | - | - | - | -0.004 | 0.083 | 9.63E-01 | -0.167 | 0.159 | -0.009 | 0.084 | 9.12E-01 | -0.175 | 0.156 |
| | **dorsal body surface area (in SD)** | - | - | - | - | - | -0.069 | 0.068 | 3.11E-01 | -0.202 | 0.064 | -0.071 | 0.069 | 3.00E-01 | -0.206 | 0.063 |
| | **LDL cholesterol levels (in SD)** | - | - | - | - | - | - | - | - | - | - | 0.086 | 0.061 | 1.57E-01 | -0.033 | 0.204 |
| | **HDL cholesterol levels (in SD)** | - | - | - | - | - | - | - | - | - | - | 0.073 | 0.068 | 2.86E-01 | -0.061 | 0.207 |
| | **triglyceride levels (in SD)** | - | - | - | - | - | - | - | - | - | - | 0.060 | 0.085 | 4.82E-01 | -0.106 | 0.226 |
| | **glucose levels (in SD)** | - | - | - | - | - | - | - | - | - | - | -0.074 | 0.064 | 2.47E-01 | -0.199 | 0.051 |
| | **time of day (in hours since 9AM)** | 0.008 | 0.047 | 8.71E-01 | -0.084 | 0.099 | 0.012 | 0.052 | 8.18E-01 | -0.091 | 0.115 | 0.017 | 0.053 | 7.53E-01 | -0.087 | 0.120 |
| | **intercept_random** | 0.946 | 1.148 | 4.10E-01 | -1.304 | 3.196 | 1.163 | 1.212 | 3.37E-01 | -1.212 | 3.538 | 1.342 | 1.209 | 2.67E-01 | -1.027 | 3.711 |
| | ***apoba*** | -0.123 | 0.336 | 7.14E-01 | -0.782 | 0.535 | -0.127 | 0.365 | 7.28E-01 | -0.841 | 0.588 | -0.186 | 0.363 | 6.08E-01 | -0.897 | 0.525 |
| | ***apobb.2*** | -0.173 | 0.218 | 4.27E-01 | -0.601 | 0.254 | -0.228 | 0.238 | 3.38E-01 | -0.695 | 0.239 | -0.278 | 0.239 | 2.45E-01 | -0.747 | 0.191 |
| | ***ldlrb*** | -0.088 | 2.256 | 9.69E-01 | -4.511 | 4.334 | -0.312 | 2.335 | 8.94E-01 | -4.889 | 4.266 | -0.245 | 2.333 | 9.16E-01 | -4.817 | 4.327 |
| **random factors** | ***variance by batch*** | *0.442* | *0.129* | - | *0.249* | *0.784* | *0.421* | *0.130* | - | *0.230* | *0.773* | *0.396* | *0.128* | - | *0.211* | *0.744* |
| | ***residual*** | *0.878* | *0.037* | - | *0.808* | *0.954* | *0.903* | *0.040* | - | *0.828* | *0.986* | *0.896* | *0.040* | - | *0.821* | *0.979* |

*continued* **Supplementary Table 21**

| | | Vascular co-localization of lipids with macrophages | | | | | | | | | | | | | | |
|---|---|---|---|---|---|---|---|---|---|---|---|---|---|---|---|---|
| | | 128 vs. 89 larvae with 2 vs. 0 mutated alleles (n=217) | | | | | 115 vs. 79 larvae with 2 vs. 0 mutated alleles (n=194) | | | | | 115 vs. 79 larvae with 2 vs. 0 mutated alleles (n=194) | | | | |
| | | **Effect** | **SE** | ***P*** | **lci** | **uci** | **Effect** | **SE** | ***P*** | **lci** | **uci** | **Effect** | **SE** | ***P*** | **lci** | **uci** |
| **negative binomial terms** | **2 vs. 0 mutated alleles** | 0.184 | 0.338 | 5.87E-01 | -0.479 | 0.846 | 0.244 | 0.377 | 5.19E-01 | -0.496 | 0.983 | 0.208 | 0.380 | 5.84E-01 | -0.536 | 0.952 |
| | ***apoea*** | -0.008 | 0.252 | 9.76E-01 | -0.502 | 0.487 | -0.147 | 0.270 | 5.87E-01 | -0.676 | 0.382 | -0.168 | 0.269 | 5.31E-01 | -0.696 | 0.359 |
| | ***apoeb*** | 0.023 | 0.231 | 9.22E-01 | -0.431 | 0.476 | 0.141 | 0.254 | 5.80E-01 | -0.358 | 0.639 | 0.136 | 0.260 | 6.01E-01 | -0.373 | 0.645 |
| | ***apobb.1*** | 1.145 | 0.230 | 6.01E-07 | 0.696 | 1.595 | 0.895 | 0.296 | 2.49E-03 | 0.315 | 1.474 | 0.799 | 0.363 | 2.75E-02 | 0.088 | 1.510 |
| | **body length (in SD)** | - | - | - | - | - | -0.164 | 0.238 | 4.90E-01 | -0.631 | 0.302 | -0.215 | 0.281 | 4.45E-01 | -0.766 | 0.336 |
| | **dorsal body surface area (in SD)** | - | - | - | - | - | 0.319 | 0.150 | 3.34E-02 | 0.025 | 0.613 | 0.308 | 0.149 | 3.88E-02 | 0.016 | 0.601 |
| | **LDL cholesterol levels (in SD)** | - | - | - | - | - | - | - | - | - | - | 0.094 | 0.171 | 5.83E-01 | -0.241 | 0.429 |
| | **HDL cholesterol levels (in SD)** | - | - | - | - | - | - | - | - | - | - | 0.118 | 0.224 | 5.98E-01 | -0.321 | 0.558 |
| | **triglyceride levels (in SD)** | - | - | - | - | - | - | - | - | - | - | 0.178 | 0.275 | 5.18E-01 | -0.362 | 0.717 |
| | **glucose levels (in SD)** | - | - | - | - | - | - | - | - | - | - | -0.088 | 0.162 | 5.86E-01 | -0.406 | 0.229 |
| | **time of day (in hours since 9AM)** | 0.091 | 0.112 | 4.16E-01 | -0.128 | 0.310 | 0.088 | 0.132 | 5.04E-01 | -0.170 | 0.347 | 0.117 | 0.141 | 4.07E-01 | -0.160 | 0.394 |
| | **batch 2** | -2.643 | 0.693 | 1.38E-04 | -4.002 | -1.284 | -3.005 | 0.793 | 1.50E-04 | -4.559 | -1.452 | -3.007 | 0.802 | 1.77E-04 | -4.579 | -1.435 |
| | **batch 3** | -1.141 | 0.418 | 6.35E-03 | -1.961 | -0.322 | -1.133 | 0.497 | 2.27E-02 | -2.107 | -0.158 | -1.092 | 0.549 | 4.67E-02 | -2.169 | -0.016 |
| | **batch 4** | -0.119 | 0.500 | 8.12E-01 | -1.099 | 0.861 | -0.367 | 0.567 | 5.18E-01 | -1.477 | 0.744 | -0.283 | 0.635 | 6.56E-01 | -1.528 | 0.961 |
| | **batch 5** | 0.440 | 0.405 | 2.76E-01 | -0.353 | 1.233 | 0.151 | 0.506 | 7.65E-01 | -0.840 | 1.143 | 0.003 | 0.757 | 9.97E-01 | -1.482 | 1.487 |
| | **batch 6** | -0.124 | 0.392 | 7.52E-01 | -0.891 | 0.644 | -0.142 | 0.452 | 7.53E-01 | -1.028 | 0.744 | -0.318 | 0.969 | 7.42E-01 | -2.217 | 1.580 |
| | **batch 7** | -0.761 | 0.790 | 3.35E-01 | -2.309 | 0.787 | -1.260 | 0.764 | 9.92E-02 | -2.758 | 0.238 | -1.208 | 0.973 | 2.14E-01 | -3.115 | 0.699 |
| | **intercept** | 563.870 | 199.665 | 4.74E-03 | 172.534 | 955.207 | 660.449 | 204.076 | 1.21E-03 | 260.468 | 1060.430 | 696.194 | 205.187 | 6.91E-04 | 294.035 | 1098.352 |
| | ***apoba*** | -0.860 | 1.155 | 4.57E-01 | -3.122 | 1.403 | -1.185 | 1.185 | 3.18E-01 | -3.508 | 1.138 | -0.979 | 1.236 | 4.28E-01 | -3.401 | 1.443 |
| | ***apobb.2*** | -0.139 | 0.519 | 7.89E-01 | -1.156 | 0.878 | -0.355 | 0.519 | 4.94E-01 | -1.372 | 0.663 | -0.287 | 0.561 | 6.09E-01 | -1.386 | 0.813 |
| | ***ldlrb*** | -1400.000 | 498.871 | 4.96E-03 | -2400.000 | -423.915 | -1600.000 | 509.990 | 1.30E-03 | -2600.000 | -640.783 | -1700.000 | 512.929 | 7.40E-04 | -2700.000 | -725.512 |

| | | Vascular infiltration by neutrophils | | | | | | | | | | | | | | |
|---|---|---|---|---|---|---|---|---|---|---|---|---|---|---|---|---|
| | | 175 vs. 118 larvae with 2 vs. 0 mutated alleles (n=293) | | | | | 159 vs. 103 larvae with 2 vs. 0 mutated alleles (n=262) | | | | | 159 vs. 103 larvae with 2 vs. 0 mutated alleles (n=262) | | | | |
| | | **Effect** | **SE** | ***P*** | **lci** | **uci** | **Effect** | **SE** | ***P*** | **lci** | **uci** | **Effect** | **SE** | ***P*** | **lci** | **uci** |
| **fixed factors** | **2 vs. 0 mutated alleles** | -0.224 | 0.131 | 8.60E-02 | -0.480 | 0.032 | -0.222 | 0.141 | 1.14E-01 | -0.498 | 0.053 | -0.218 | 0.140 | 1.21E-01 | -0.493 | 0.057 |
| | ***apoea*** | -0.033 | 0.083 | 6.88E-01 | -0.197 | 0.130 | -0.071 | 0.087 | 4.18E-01 | -0.242 | 0.100 | -0.067 | 0.087 | 4.42E-01 | -0.238 | 0.104 |
| | ***apoeb*** | -0.053 | 0.087 | 5.38E-01 | -0.223 | 0.116 | -0.014 | 0.092 | 8.81E-01 | -0.194 | 0.166 | -0.030 | 0.092 | 7.45E-01 | -0.210 | 0.150 |
| | ***apobb.1*** | 0.126 | 0.092 | 1.71E-01 | -0.054 | 0.306 | 0.034 | 0.098 | 7.28E-01 | -0.158 | 0.225 | 0.057 | 0.099 | 5.62E-01 | -0.137 | 0.251 |
| | **body length (in SD)** | - | - | - | - | - | -0.161 | 0.076 | 3.44E-02 | -0.310 | -0.012 | -0.138 | 0.078 | 7.50E-02 | -0.290 | 0.014 |
| | **dorsal body surface area (in SD)** | - | - | - | - | - | 0.007 | 0.062 | 9.06E-01 | -0.114 | 0.129 | 0.030 | 0.063 | 6.39E-01 | -0.094 | 0.153 |
| | **LDL cholesterol levels (in SD)** | - | - | - | - | - | - | - | - | - | - | -0.033 | 0.055 | 5.50E-01 | -0.141 | 0.075 |
| | **HDL cholesterol levels (in SD)** | - | - | - | - | - | - | - | - | - | - | 0.053 | 0.063 | 4.00E-01 | -0.070 | 0.177 |
| | **triglyceride levels (in SD)** | - | - | - | - | - | - | - | - | - | - | -0.096 | 0.077 | 2.10E-01 | -0.247 | 0.054 |
| | **glucose levels (in SD)** | - | - | - | - | - | - | - | - | - | - | 0.065 | 0.059 | 2.67E-01 | -0.050 | 0.181 |
| | **time of day (in hours since 9AM)** | -0.014 | 0.044 | 7.59E-01 | -0.100 | 0.073 | -0.036 | 0.048 | 4.48E-01 | -0.129 | 0.057 | -0.043 | 0.048 | 3.69E-01 | -0.137 | 0.051 |
| | **intercept_random** | 0.244 | 1.101 | 8.25E-01 | -1.914 | 2.402 | -0.103 | 1.119 | 9.27E-01 | -2.297 | 2.091 | -0.103 | 1.119 | 9.26E-01 | -2.296 | 2.089 |
| | ***apoba*** | 0.488 | 0.323 | 1.30E-01 | -0.144 | 1.120 | 0.557 | 0.337 | 9.81E-02 | -0.103 | 1.218 | 0.584 | 0.336 | 8.20E-02 | -0.074 | 1.243 |
| | ***apobb.2*** | -0.169 | 0.210 | 4.21E-01 | -0.581 | 0.243 | -0.093 | 0.221 | 6.73E-01 | -0.526 | 0.340 | -0.094 | 0.222 | 6.72E-01 | -0.529 | 0.341 |
| | ***ldlrb*** | -1.665 | 2.175 | 4.44E-01 | -5.927 | 2.598 | -1.151 | 2.169 | 5.96E-01 | -5.403 | 3.100 | -1.232 | 2.173 | 5.71E-01 | -5.492 | 3.027 |
| **random factors** | ***variance by batch*** | *0.343* | *0.102* | - | *0.192* | *0.615* | *0.314* | *0.098* | - | *0.171* | *0.578* | *0.303* | *0.096* | - | *0.163* | *0.563* |
| | ***residual*** | *0.847* | *0.036* | - | *0.780* | *0.919* | *0.839* | *0.037* | - | *0.769* | *0.915* | *0.834* | *0.037* | - | *0.765* | *0.910* |

***continued*** **Supplementary Table 21**

| | | Vascular co-localization of lipids with neutrophils | | | | | | | | | | | | | | |
|---|---|---|---|---|---|---|---|---|---|---|---|---|---|---|---|---|
| | | **123 vs. 85 larvae with 2 vs. 0 mutated alleles (n=208)** | | | | | **110 vs. 75 larvae with 2 vs. 0 mutated alleles (n=185)** | | | | | **110 vs. 75 larvae with 2 vs. 0 mutated alleles (n=185)** | | | | |
| | | **Effect** | **SE** | ***P*** | **lci** | **uci** | **Effect** | **SE** | ***P*** | **lci** | **uci** | **Effect** | **SE** | ***P*** | **lci** | **uci** |
| **negative binomial terms** | **2 vs. 0 mutated alleles** | 0.043 | 0.496 | 9.31E-01 | -0.929 | 1.015 | 0.299 | 0.483 | 5.36E-01 | -0.647 | 1.245 | -0.089 | 0.481 | 8.53E-01 | -1.031 | 0.853 |
| | ***apoea*** | -0.944 | 0.371 | 1.09E-02 | -1.672 | -0.217 | -1.037 | 0.412 | 1.18E-02 | -1.844 | -0.229 | -1.329 | 0.471 | 4.77E-03 | -2.252 | -0.406 |
| | ***apoeb*** | -0.379 | 0.342 | 2.69E-01 | -1.050 | 0.292 | -0.270 | 0.324 | 4.05E-01 | -0.905 | 0.365 | -0.234 | 0.315 | 4.58E-01 | -0.851 | 0.384 |
| | ***apobb.1*** | 2.008 | 0.316 | 2.06E-10 | 1.389 | 2.626 | 1.756 | 0.391 | 6.97E-06 | 0.990 | 2.521 | 2.266 | 0.444 | 3.34E-07 | 1.396 | 3.136 |
| | **body length (in SD)** | - | - | - | - | - | -0.593 | 0.294 | 4.38E-02 | -1.169 | -0.016 | -0.817 | 0.276 | 3.06E-03 | -1.358 | -0.276 |
| | **dorsal body surface area (in SD)** | - | - | - | - | - | -0.419 | 0.210 | 4.56E-02 | -0.830 | -0.008 | -0.645 | 0.235 | 5.97E-03 | -1.105 | -0.185 |
| | **LDL cholesterol levels (in SD)** | - | - | - | - | - | - | - | - | - | - | 0.859 | 0.339 | 1.14E-02 | 0.194 | 1.524 |
| | **HDL cholesterol levels (in SD)** | - | - | - | - | - | - | - | - | - | - | 0.974 | 0.369 | 8.26E-03 | 0.251 | 1.696 |
| | **triglyceride levels (in SD)** | - | - | - | - | - | - | - | - | - | - | 0.482 | 0.387 | 2.12E-01 | -0.276 | 1.240 |
| | **glucose levels (in SD)** | - | - | - | - | - | - | - | - | - | - | 0.588 | 0.284 | 3.80E-02 | 0.032 | 1.144 |
| | **time of day (in hours since 9AM)** | -0.130 | 0.173 | 4.50E-01 | -0.469 | 0.208 | 0.204 | 0.227 | 3.68E-01 | -0.240 | 0.648 | 0.345 | 0.236 | 1.45E-01 | -0.119 | 0.808 |
| | **batch 2** | -3.290 | 0.866 | 1.44E-04 | -4.987 | -1.594 | -3.777 | 0.835 | 6.12E-06 | -5.414 | -2.140 | -3.831 | 0.924 | 3.38E-05 | -5.642 | -2.020 |
| | **batch 3** | 0.046 | 0.833 | 9.56E-01 | -1.587 | 1.680 | -0.065 | 0.817 | 9.36E-01 | -1.667 | 1.536 | -0.456 | 0.766 | 5.51E-01 | -1.958 | 1.045 |
| | **batch 4** | -0.858 | 0.698 | 2.19E-01 | -2.225 | 0.509 | -1.467 | 0.741 | 4.77E-02 | -2.919 | -0.015 | -2.263 | 0.873 | 9.53E-03 | -3.974 | -0.552 |
| | **batch 5** | 0.197 | 0.640 | 7.59E-01 | -1.058 | 1.451 | -0.536 | 0.667 | 4.22E-01 | -1.842 | 0.771 | -1.466 | 0.902 | 1.04E-01 | -3.235 | 0.302 |
| | **batch 6** | -1.714 | 0.692 | 1.32E-02 | -3.070 | -0.358 | -1.552 | 0.687 | 2.39E-02 | -2.899 | -0.205 | -1.854 | 1.026 | 7.07E-02 | -3.865 | 0.156 |
| | **intercept** | 469.980 | 217.000 | 3.03E-02 | 44.667 | 895.292 | 696.320 | 210.590 | 9.45E-04 | 283.570 | 1109.069 | 820.268 | 241.156 | 6.70E-04 | 347.611 | 1292.926 |
| | ***apoba*** | 2.535 | 1.153 | 2.79E-02 | 0.275 | 4.795 | 2.308 | 1.180 | 5.05E-02 | -0.005 | 4.620 | 4.189 | 1.468 | 4.33E-03 | 1.311 | 7.067 |
| | ***apobb.2*** | 1.439 | 0.734 | 5.01E-02 | 0.000 | 2.878 | 1.334 | 0.721 | 6.42E-02 | -0.078 | 2.747 | 2.106 | 0.792 | 7.84E-03 | 0.554 | 3.658 |
| | ***ldlrb*** | -1200.000 | 543.813 | 2.89E-02 | -2300.000 | -122.607 | -1800.000 | 528.345 | 8.90E-04 | -2800.000 | -720.288 | -2100.000 | 605.508 | 5.99E-04 | -3300.000 | -891.491 |

| | | Vascular co-localization of macrophages with neutrophils | | | | | | | | | | | | | | |
|---|---|---|---|---|---|---|---|---|---|---|---|---|---|---|---|---|
| | | **173 vs. 117 larvae with 2 vs. 0 mutated alleles (n=290)** | | | | | **157 vs. 102 larvae with 2 vs. 0 mutated alleles (n=259)** | | | | | **157 vs. 102 larvae with 2 vs. 0 mutated alleles (n=259)** | | | | |
| | | **Effect** | **SE** | ***P*** | **lci** | **uci** | **Effect** | **SE** | ***P*** | **lci** | **uci** | **Effect** | **SE** | ***P*** | **lci** | **uci** |
| **negative binomial terms** | **2 vs. 0 mutated alleles** | -0.299 | 0.223 | 1.80E-01 | -0.736 | 0.138 | -0.455 | 0.232 | 4.99E-02 | -0.910 | 0.000 | -0.465 | 0.233 | 4.57E-02 | -0.922 | -0.009 |
| | ***apoea*** | -0.150 | 0.128 | 2.39E-01 | -0.401 | 0.100 | -0.158 | 0.134 | 2.38E-01 | -0.420 | 0.104 | -0.156 | 0.136 | 2.50E-01 | -0.422 | 0.110 |
| | ***apoeb*** | 0.072 | 0.131 | 5.83E-01 | -0.185 | 0.329 | 0.118 | 0.140 | 4.01E-01 | -0.157 | 0.393 | 0.095 | 0.138 | 4.92E-01 | -0.176 | 0.365 |
| | ***apobb.1*** | -0.072 | 0.153 | 6.36E-01 | -0.372 | 0.227 | -0.085 | 0.147 | 5.64E-01 | -0.373 | 0.203 | -0.039 | 0.147 | 7.91E-01 | -0.328 | 0.250 |
| | **body length (in SD)** | - | - | - | - | - | 0.236 | 0.122 | 5.37E-02 | -0.004 | 0.475 | 0.228 | 0.131 | 8.27E-02 | -0.030 | 0.486 |
| | **dorsal body surface area (in SD)** | - | - | - | - | - | 0.350 | 0.116 | 2.46E-03 | 0.123 | 0.576 | 0.357 | 0.121 | 3.14E-03 | 0.120 | 0.595 |
| | **LDL cholesterol levels (in SD)** | - | - | - | - | - | - | - | - | - | - | 0.031 | 0.106 | 7.70E-01 | -0.177 | 0.238 |
| | **HDL cholesterol levels (in SD)** | - | - | - | - | - | - | - | - | - | - | 0.024 | 0.112 | 8.31E-01 | -0.196 | 0.244 |
| | **triglyceride levels (in SD)** | - | - | - | - | - | - | - | - | - | - | -0.104 | 0.150 | 4.86E-01 | -0.398 | 0.189 |
| | **glucose levels (in SD)** | - | - | - | - | - | - | - | - | - | - | 0.125 | 0.095 | 1.88E-01 | -0.061 | 0.310 |
| | **time of day (in hours since 9AM)** | 0.025 | 0.080 | 7.50E-01 | -0.131 | 0.181 | 0.011 | 0.083 | 8.94E-01 | -0.151 | 0.173 | -0.006 | 0.086 | 9.45E-01 | -0.174 | 0.162 |
| | **batch 1** | 0.300 | 0.321 | 3.51E-01 | -0.330 | 0.930 | 0.107 | 0.354 | 7.63E-01 | -0.588 | 0.802 | 0.093 | 0.430 | 8.29E-01 | -0.750 | 0.935 |
| | **batch 2** | 0.189 | 0.490 | 7.00E-01 | -0.771 | 1.149 | 0.246 | 0.462 | 5.94E-01 | -0.659 | 1.152 | 0.208 | 0.501 | 6.78E-01 | -0.773 | 1.189 |
| | **batch 3** | -0.660 | 0.441 | 1.34E-01 | -1.525 | 0.204 | -0.643 | 0.433 | 1.38E-01 | -1.492 | 0.206 | -0.589 | 0.479 | 2.19E-01 | -1.528 | 0.349 |
| | **batch 4** | 0.442 | 0.381 | 2.46E-01 | -0.304 | 1.187 | 0.573 | 0.399 | 1.51E-01 | -0.209 | 1.355 | 0.502 | 0.447 | 2.61E-01 | -0.374 | 1.378 |
| | **batch 5** | 0.067 | 0.387 | 8.63E-01 | -0.692 | 0.826 | -0.183 | 0.405 | 6.51E-01 | -0.976 | 0.610 | -0.028 | 0.457 | 9.52E-01 | -0.923 | 0.868 |
| | **batch 6** | -1.557 | 0.403 | 1.11E-04 | -2.346 | -0.768 | -2.058 | 0.465 | 9.81E-06 | -2.970 | -1.145 | -1.927 | 0.489 | 8.25E-05 | -2.886 | -0.968 |
| | **batch 7** | -0.537 | 0.550 | 3.28E-01 | -1.614 | 0.540 | -1.034 | 0.592 | 8.08E-02 | -2.195 | 0.127 | -1.022 | 0.734 | 1.64E-01 | -2.461 | 0.416 |
| | **intercept** | 3.443 | 0.942 | 2.58E-04 | 1.596 | 5.291 | 3.341 | 0.936 | 3.57E-04 | 1.507 | 5.176 | 2.978 | 0.976 | 2.29E-03 | 1.064 | 4.892 |
| | ***apoba*** | 0.309 | 0.508 | 5.43E-01 | -0.686 | 1.304 | 0.480 | 0.492 | 3.30E-01 | -0.485 | 1.444 | 0.565 | 0.483 | 2.42E-01 | -0.382 | 1.513 |
| | ***apobb.2*** | -0.009 | 0.273 | 9.73E-01 | -0.544 | 0.526 | -0.039 | 0.276 | 8.89E-01 | -0.579 | 0.502 | 0.004 | 0.276 | 9.88E-01 | -0.538 | 0.546 |
| | ***ldlrb*** | 0.307 | 0.742 | 6.79E-01 | -1.146 | 1.760 | 0.522 | 0.879 | 5.52E-01 | -1.200 | 2.244 | 0.893 | 0.997 | 3.71E-01 | -1.062 | 2.848 |

Dorsal and lateral body surface area and body volume were normalized for body length using residuals. All outcomes were inverse-normally transformed before the analysis and examined using hierarchical linear models. Effects shown are for larvae with two mutated alleles that are highly likely to affect protein function as predicted by Ensembl's Varient Effect Predictor (VEP) compared with larvae with zero CRISPR-mutated alleles. Associations were adjusted for the number of mutated alleles in the other six orthologues, weighted by their predicted effect on protein function, as well as for time of day and batch. Lci and uci are lower and upper boundaries of the 95% confidence interval.

**Supplementary Table 22 - The additive effect of mutated alleles in *apoea*, *apoeb*, *apobb.1* and *ldlra* on body size**

| | | Body length (n=339) | | | | |
|---|---|---|---|---|---|---|
| | | **Effect** | **SE** | ***P*** | **lci** | **uci** |
| **fixed factors** | ***apoea*** | 0.008 | 0.063 | 8.96E-01 | -0.115 | 0.131 |
| | ***apoeb*** | -0.010 | 0.070 | 8.89E-01 | -0.146 | 0.127 |
| | ***apoba*** | -0.039 | 0.233 | 8.67E-01 | -0.495 | 0.417 |
| | ***apobb.1*** | -0.336 | 0.065 | 2.57E-07 | -0.464 | -0.208 |
| | ***apobb.2*** | 0.393 | 0.162 | 1.52E-02 | 0.076 | 0.711 |
| | ***ldlra*** | 0.058 | 0.054 | 2.85E-01 | -0.048 | 0.164 |
| | **time of day (in hours since 9AM)** | -0.039 | 0.036 | 2.76E-01 | -0.110 | 0.032 |
| | **intercept** | -1.203 | 0.891 | 1.77E-01 | -2.950 | 0.544 |
| | ***ldlrb*** | 2.088 | 1.789 | 2.43E-01 | -1.418 | 5.595 |
| **random factors** | ***variance by batch*** | *0.497* | *0.135* | - | *0.292* | *0.846* |
| | ***residual*** | *0.698* | *0.027* | - | *0.647* | *0.753* |

| | | Dorsal body surface area (n=339) | | | | |
|---|---|---|---|---|---|---|
| | | **Effect** | **SE** | ***P*** | **lci** | **uci** |
| **fixed factors** | ***apoea*** | -0.120 | 0.074 | 1.08E-01 | -0.266 | 0.026 |
| | ***apoeb*** | 0.122 | 0.083 | 1.39E-01 | -0.040 | 0.284 |
| | ***apoba*** | 0.072 | 0.276 | 7.94E-01 | -0.469 | 0.613 |
| | ***apobb.1*** | 0.114 | 0.077 | 1.39E-01 | -0.037 | 0.266 |
| | ***apobb.2*** | -0.426 | 0.192 | 2.67E-02 | -0.802 | -0.049 |
| | ***ldlra*** | -0.069 | 0.064 | 2.80E-01 | -0.195 | 0.056 |
| | **time of day (in hours since 9AM)** | 0.029 | 0.043 | 4.99E-01 | -0.055 | 0.112 |
| | **intercept** | 1.396 | 1.050 | 1.83E-01 | -0.661 | 3.454 |
| | ***ldlrb*** | -2.842 | 2.122 | 1.80E-01 | -7.001 | 1.316 |
| **random factors** | ***variance by batch*** | *0.475* | *0.134* | - | *0.273* | *0.827* |
| | ***residual*** | *0.828* | *0.032* | - | *0.768* | *0.894* |

| | | Lateral body surface area (n=335) | | | | |
|---|---|---|---|---|---|---|
| | | **Effect** | **SE** | ***P*** | **lci** | **uci** |
| **fixed factors** | ***apoea*** | -0.131 | 0.076 | 8.49E-02 | -0.281 | 0.018 |
| | ***apoeb*** | 0.169 | 0.086 | 4.83E-02 | 0.001 | 0.337 |
| | ***apoba*** | -0.013 | 0.286 | 9.63E-01 | -0.573 | 0.547 |
| | ***apobb.1*** | 0.020 | 0.079 | 8.04E-01 | -0.136 | 0.175 |
| | ***apobb.2*** | -0.207 | 0.196 | 2.92E-01 | -0.592 | 0.178 |
| | ***ldlra*** | -0.026 | 0.066 | 6.99E-01 | -0.155 | 0.104 |
| | **time of day (in hours since 9AM)** | 0.014 | 0.044 | 7.51E-01 | -0.072 | 0.099 |
| | **intercept** | 1.049 | 1.076 | 3.30E-01 | -1.061 | 3.158 |
| | ***ldlrb*** | -2.429 | 2.170 | 2.63E-01 | -6.682 | 1.823 |
| **random factors** | ***variance by batch*** | *0.469* | *0.132* | - | *0.270* | *0.814* |
| | ***residual*** | *0.845* | *0.033* | - | *0.783* | *0.913* |

| | | Body volume (n=328) | | | | |
|---|---|---|---|---|---|---|
| | | **Effect** | **SE** | ***P*** | **lci** | **uci** |
| **fixed factors** | ***apoea*** | -0.111 | 0.076 | 1.43E-01 | -0.259 | 0.037 |
| | ***apoeb*** | 0.170 | 0.085 | 4.48E-02 | 0.004 | 0.336 |
| | ***apoba*** | 0.015 | 0.281 | 9.57E-01 | -0.535 | 0.566 |
| | ***apobb.1*** | 0.047 | 0.079 | 5.52E-01 | -0.107 | 0.201 |
| | ***apobb.2*** | -0.294 | 0.195 | 1.32E-01 | -0.676 | 0.089 |
| | ***ldlra*** | -0.043 | 0.065 | 5.11E-01 | -0.171 | 0.085 |
| | **time of day (in hours since 9AM)** | 0.036 | 0.044 | 4.09E-01 | -0.050 | 0.122 |
| | **intercept** | 1.177 | 1.058 | 2.66E-01 | -0.897 | 3.250 |
| | ***ldlrb*** | -2.747 | 2.123 | 1.96E-01 | -6.909 | 1.415 |
| **random factors** | ***variance by batch*** | *0.478* | *0.134* | - | *0.276* | *0.827* |
| | ***residual*** | *0.826* | *0.033* | - | *0.765* | *0.893* |

All outcomes were normalized for length using residuals, and inverse-normally transformed before the analysis. Associations were examined using hierarchical linear models. Effects shown are for each additional mutated allele in *apoea*, *apoeb*, *apoba*, *apobb.1*, *apobb.2*, *ldlra* and *ldlrb*, weighted by the allele's predicted effect on protein function (i.e. additive model, mutually adjusted). Associations were adjusted for time of day and batch. Lci and uci are lower and upper boundaries of the 95% confidence interval.

**Supplementary Table 23 - The additive effect of mutated alleles in *apoea* , *apoeb* , *apobb.1* and *ldlra* on**

| | | LDL cholesterol levels (n=381) | | | | |
|---|---|---|---|---|---|---|
| | | **Effect** | **SE** | ***P*** | **lci** | **uci** |
| **fixed factors** | ***apoea*** | 0.183 | 0.080 | 2.31E-02 | 0.025 | 0.341 |
| | ***apoeb*** | 0.068 | 0.089 | 4.43E-01 | -0.106 | 0.243 |
| | ***apoba*** | 0.164 | 0.300 | 5.86E-01 | -0.425 | 0.753 |
| | ***apobb.1*** | 0.043 | 0.085 | 6.17E-01 | -0.124 | 0.209 |
| | ***apobb.2*** | 0.293 | 0.211 | 1.65E-01 | -0.120 | 0.706 |
| | ***ldlra*** | -0.027 | 0.068 | 6.94E-01 | -0.160 | 0.107 |
| | **body length (in SD)** | - | - | - | - | - |
| | **dorsal body surface area (in SD)** | - | - | - | - | - |
| | **time of day (in hours since 9AM)** | 0.055 | 0.043 | 2.00E-01 | -0.029 | 0.138 |
| | **intercept** | 0.392 | 1.169 | 7.38E-01 | -1.900 | 2.684 |
| | ***ldlrb*** | -3.665 | 2.418 | 1.30E-01 | -8.405 | 1.074 |
| **random factors** | ***variance by batch*** | *0.331* | *0.119* | - | *0.164* | *0.668* |
| | ***residual*** | *0.948* | *0.035* | - | *0.882* | *1.018* |

| | | HDL cholesterol levels (n=381) | | | | |
|---|---|---|---|---|---|---|
| | | **Effect** | **SE** | ***P*** | **lci** | **uci** |
| **fixed factors** | ***apoea*** | 0.031 | 0.073 | 6.76E-01 | -0.113 | 0.174 |
| | ***apoeb*** | 0.092 | 0.081 | 2.54E-01 | -0.066 | 0.250 |
| | ***apoba*** | 0.013 | 0.273 | 9.63E-01 | -0.522 | 0.547 |
| | ***apobb.1*** | 0.079 | 0.077 | 3.06E-01 | -0.072 | 0.230 |
| | ***apobb.2*** | 0.318 | 0.191 | 9.60E-02 | -0.056 | 0.693 |
| | ***ldlra*** | 0.066 | 0.062 | 2.83E-01 | -0.055 | 0.187 |
| | **body length (in SD)** | - | - | - | - | - |
| | **dorsal body surface area (in SD)** | - | - | - | - | - |
| | **time of day (in hours since 9AM)** | -0.028 | 0.040 | 4.83E-01 | -0.107 | 0.050 |
| | **intercept** | -1.030 | 1.076 | 3.39E-01 | -3.138 | 1.079 |
| | ***ldlrb*** | 0.847 | 2.194 | 6.99E-01 | -3.453 | 5.147 |
| **random factors** | ***variance by batch*** | *0.592* | *0.160* | - | *0.348* | *1.006* |
| | ***residual*** | *0.857* | *0.031* | - | *0.797* | *0.921* |

| | | Triglyceride levels (n=381) | | | | |
|---|---|---|---|---|---|---|
| | | **Effect** | **SE** | ***P*** | **lci** | **uci** |
| **fixed factors** | ***apoea*** | -0.083 | 0.058 | 1.53E-01 | -0.198 | 0.031 |
| | ***apoeb*** | 0.008 | 0.064 | 9.06E-01 | -0.118 | 0.133 |
| | ***apoba*** | -0.178 | 0.217 | 4.11E-01 | -0.604 | 0.247 |
| | ***apobb.1*** | 0.108 | 0.061 | 7.86E-02 | -0.012 | 0.228 |
| | ***apobb.2*** | -0.298 | 0.152 | 4.99E-02 | -0.596 | 0.000 |
| | ***ldlra*** | 0.025 | 0.049 | 6.05E-01 | -0.071 | 0.122 |
| | **body length (in SD)** | - | - | - | - | - |
| | **dorsal body surface area (in SD)** | - | - | - | - | - |
| | **time of day (in hours since 9AM)** | -0.143 | 0.032 | 9.33E-06 | -0.206 | -0.080 |
| | **intercept** | 1.064 | 0.883 | 2.28E-01 | -0.667 | 2.795 |
| | ***ldlrb*** | -0.236 | 1.746 | 8.93E-01 | -3.658 | 3.187 |
| **random factors** | ***variance by batch*** | *0.771* | *0.198* | - | *0.467* | *1.275* |
| | ***residual*** | *0.681* | *0.025* | - | *0.634* | *0.732* |

***continued* Supplementary Table 23**

| | | Total cholesterol levels (n=381) | | | | |
|---|---|---|---|---|---|---|
| | | **Effect** | **SE** | ***P*** | **lci** | **uci** |
| **fixed factors** | ***apoea*** | 0.046 | 0.070 | 5.08E-01 | -0.090 | 0.182 |
| | ***apoeb*** | 0.049 | 0.076 | 5.20E-01 | -0.101 | 0.199 |
| | ***apoba*** | 0.192 | 0.259 | 4.57E-01 | -0.315 | 0.700 |
| | ***apobb.1*** | -0.219 | 0.073 | 2.69E-03 | -0.363 | -0.076 |
| | ***apobb.2*** | -0.109 | 0.181 | 5.47E-01 | -0.465 | 0.246 |
| | ***ldlra*** | -0.027 | 0.059 | 6.46E-01 | -0.142 | 0.088 |
| | **body length (in SD)** | - | - | - | - | - |
| | **dorsal body surface area (in SD)** | - | - | - | - | - |
| | **time of day (in hours since 9AM)** | 0.075 | 0.038 | 5.04E-02 | 0.000 | 0.150 |
| | **intercept** | 0.130 | 1.036 | 9.00E-01 | -1.899 | 2.160 |
| | ***ldlrb*** | -1.012 | 2.082 | 6.27E-01 | -5.092 | 3.068 |
| **random factors** | ***variance by batch*** | *0.748* | *0.198* | - | *0.445* | *1.255* |
| | ***residual*** | *0.812* | *0.030* | - | *0.756* | *0.873* |

| | | Glucose levels (n=381) | | | | |
|---|---|---|---|---|---|---|
| | | **Effect** | **SE** | ***P*** | **lci** | **uci** |
| **fixed factors** | ***apoea*** | -0.062 | 0.080 | 4.36E-01 | -0.220 | 0.095 |
| | ***apoeb*** | 0.121 | 0.089 | 1.71E-01 | -0.053 | 0.295 |
| | ***apoba*** | -0.141 | 0.300 | 6.38E-01 | -0.728 | 0.446 |
| | ***apobb.1*** | -0.172 | 0.085 | 4.27E-02 | -0.338 | -0.006 |
| | ***apobb.2*** | -0.385 | 0.210 | 6.69E-02 | -0.797 | 0.027 |
| | ***ldlra*** | -0.076 | 0.068 | 2.63E-01 | -0.209 | 0.057 |
| | **body length (in SD)** | - | - | - | - | - |
| | **dorsal body surface area (in SD)** | - | - | - | - | - |
| | **time of day (in hours since 9AM)** | 0.038 | 0.043 | 3.72E-01 | -0.046 | 0.122 |
| | **intercept** | 1.544 | 1.167 | 1.86E-01 | -0.744 | 3.832 |
| | ***ldlrb*** | -1.971 | 2.410 | 4.13E-01 | -6.694 | 2.752 |
| **random factors** | ***variance by batch*** | *0.384* | *0.121* | - | *0.207* | *0.710* |
| | ***residual*** | *0.943* | *0.035* | - | *0.878* | *1.014* |

All outcomes were normalized for length using residuals, and inverse-normally transformed before the analysis. Associations were examined using hierarchical linear models. Effects shown are for each additional mutated allele in *apoea* , *apoeb* , *apoba* , *apobb.1,* *apobb.2* , *ldlra* and *ldlrb,* weighted by the allele's predicted effect on protein function (i.e. additive model, mutually adjusted). Associations were adjusted for time of day and batch. Lci and uci are lower and upper boundaries of the 95% confidence interval.

**Supplementary Table 24 - The additive effect of mutated alleles in *apoea* , *apoeb* , *apobb.1* and *ldlra* on image-based vascular atherogenic traits**

| | | Vascular lipid deposition | | | | | | | | | | | | | | |
|---|---|---|---|---|---|---|---|---|---|---|---|---|---|---|---|---|
| | | Model 1 (n=306) | | | | | Model 2 (n=272) | | | | | Model 3 (n=272) | | | | |
| | | Effect | SE | *P* | lci | uci | Effect | SE | *P* | lci | uci | Effect | SE | *P* | lci | uci |
| negative binomial terms | ***apoea*** | -0.120 | 0.158 | 4.47E-01 | -0.431 | 0.190 | -0.169 | 0.173 | 3.28E-01 | -0.509 | 0.170 | -0.124 | 0.172 | 4.71E-01 | -0.462 | 0.214 |
| | ***apoeb*** | 0.189 | 0.146 | 1.96E-01 | -0.098 | 0.476 | 0.231 | 0.160 | 1.49E-01 | -0.083 | 0.546 | 0.227 | 0.162 | 1.61E-01 | -0.090 | 0.545 |
| | ***apoba*** | 0.003 | 0.508 | 9.95E-01 | -0.993 | 0.999 | -0.093 | 0.521 | 8.58E-01 | -1.114 | 0.928 | 0.113 | 0.515 | 8.26E-01 | -0.896 | 1.122 |
| | ***apobb.1*** | 0.972 | 0.144 | 1.52E-11 | 0.690 | 1.255 | 0.972 | 0.183 | 1.02E-07 | 0.614 | 1.329 | 0.947 | 0.196 | 1.30E-06 | 0.564 | 1.331 |
| | ***apobb.2*** | -0.330 | 0.410 | 4.20E-01 | -1.133 | 0.473 | -0.341 | 0.412 | 4.09E-01 | -1.148 | 0.467 | -0.152 | 0.405 | 7.07E-01 | -0.946 | 0.641 |
| | ***ldlra*** | 0.007 | 0.137 | 9.62E-01 | -0.261 | 0.274 | 0.018 | 0.142 | 8.98E-01 | -0.260 | 0.296 | 0.001 | 0.144 | 9.95E-01 | -0.280 | 0.282 |
| | **body length (in SD)** | - | - | - | - | - | -0.030 | 0.158 | 8.47E-01 | -0.340 | 0.279 | 0.017 | 0.161 | 9.16E-01 | -0.299 | 0.333 |
| | **dorsal body surface area (in SD)** | - | - | - | - | - | 0.176 | 0.126 | 1.61E-01 | -0.070 | 0.423 | 0.196 | 0.128 | 1.25E-01 | -0.055 | 0.447 |
| | **LDL cholesterol levels (in SD)** | - | - | - | - | - | - | - | - | - | - | -0.060 | 0.092 | 5.18E-01 | -0.241 | 0.121 |
| | **HDL cholesterol levels (in SD)** | - | - | - | - | - | - | - | - | - | - | -0.118 | 0.126 | 3.49E-01 | -0.364 | 0.128 |
| | **triglyceride levels (in SD)** | - | - | - | - | - | - | - | - | - | - | 0.193 | 0.170 | 2.56E-01 | -0.140 | 0.527 |
| | **glucose levels (in SD)** | - | - | - | - | - | - | - | - | - | - | -0.049 | 0.104 | 6.40E-01 | -0.254 | 0.156 |
| | **time of day (in hours since 9AM)** | -0.107 | 0.080 | 1.81E-01 | -0.263 | 0.050 | -0.177 | 0.099 | 7.39E-02 | -0.372 | 0.017 | -0.148 | 0.104 | 1.56E-01 | -0.352 | 0.056 |
| | **batch 1** | 2.477 | 0.423 | 4.67E-09 | 1.648 | 3.305 | 2.692 | 0.492 | 4.40E-08 | 1.728 | 3.656 | 2.861 | 0.549 | 1.90E-07 | 1.784 | 3.937 |
| | **batch 2** | 1.303 | 0.517 | 1.17E-02 | 0.290 | 2.315 | 1.415 | 0.531 | 7.69E-03 | 0.374 | 2.455 | 1.764 | 0.613 | 3.99E-03 | 0.563 | 2.965 |
| | **batch 3** | 1.771 | 0.500 | 3.96E-04 | 0.791 | 2.752 | 1.970 | 0.524 | 1.69E-04 | 0.943 | 2.996 | 1.986 | 0.545 | 2.69E-04 | 0.918 | 3.055 |
| | **batch 4** | 1.559 | 0.513 | 2.38E-03 | 0.553 | 2.565 | 1.753 | 0.528 | 8.92E-04 | 0.719 | 2.788 | 1.914 | 0.541 | 4.06E-04 | 0.853 | 2.975 |
| | **batch 5** | 2.972 | 0.518 | 9.40E-09 | 1.957 | 3.986 | 3.193 | 0.592 | 6.96E-08 | 2.032 | 4.353 | 2.949 | 0.656 | 6.99E-06 | 1.663 | 4.235 |
| | **batch 6** | 2.623 | 0.443 | 3.33E-09 | 1.754 | 3.492 | 2.794 | 0.553 | 4.44E-07 | 1.709 | 3.878 | 2.462 | 0.596 | 3.55E-05 | 1.295 | 3.629 |
| | **batch 7** | 1.974 | 0.515 | 1.25E-04 | 0.965 | 2.982 | 2.088 | 0.611 | 6.26E-04 | 0.892 | 3.285 | 1.827 | 0.631 | 3.76E-03 | 0.591 | 3.063 |
| | **intercept** | -4.563 | 1.643 | 5.47E-03 | -7.783 | -1.344 | -4.567 | 1.623 | 4.90E-03 | -7.748 | -1.385 | -5.112 | 1.615 | 1.55E-03 | -8.276 | -1.947 |
| | ***ldlrb*** | 17.601 | 3.301 | 9.70E-08 | 11.131 | 24.070 | 18.155 | 3.236 | 2.01E-08 | 11.813 | 24.496 | 17.829 | 3.207 | 2.72E-08 | 11.543 | 24.115 |

| | | Vascular infiltration by macrophages | | | | | | | | | | | | | | |
|---|---|---|---|---|---|---|---|---|---|---|---|---|---|---|---|---|
| | | Model 1 (n=368) | | | | | Model 2 (n=328) | | | | | Model 3 (n=328) | | | | |
| | | Effect | SE | *P* | lci | uci | Effect | SE | *P* | lci | uci | Effect | SE | *P* | lci | uci |
| fixed factors | ***apoea*** | -0.067 | 0.075 | 3.70E-01 | -0.213 | 0.079 | -0.111 | 0.081 | 1.67E-01 | -0.269 | 0.047 | -0.126 | 0.081 | 1.20E-01 | -0.285 | 0.033 |
| | ***apoeb*** | -0.145 | 0.082 | 7.49E-02 | -0.305 | 0.015 | -0.125 | 0.089 | 1.59E-01 | -0.299 | 0.049 | -0.129 | 0.089 | 1.49E-01 | -0.303 | 0.046 |
| | ***apoba*** | -0.144 | 0.275 | 6.02E-01 | -0.683 | 0.396 | -0.165 | 0.295 | 5.76E-01 | -0.743 | 0.413 | -0.178 | 0.294 | 5.44E-01 | -0.754 | 0.398 |
| | ***apobb.1*** | -0.222 | 0.080 | 5.54E-03 | -0.379 | -0.065 | -0.208 | 0.087 | 1.72E-02 | -0.379 | -0.037 | -0.222 | 0.089 | 1.23E-02 | -0.396 | -0.048 |
| | ***apobb.2*** | -0.300 | 0.193 | 1.21E-01 | -0.678 | 0.079 | -0.373 | 0.209 | 7.41E-02 | -0.782 | 0.036 | -0.425 | 0.211 | 4.37E-02 | -0.838 | -0.012 |
| | ***ldlra*** | 0.034 | 0.063 | 5.91E-01 | -0.090 | 0.158 | 0.013 | 0.070 | 8.50E-01 | -0.124 | 0.150 | -0.002 | 0.070 | 9.74E-01 | -0.139 | 0.135 |
| | **body length (in SD)** | - | - | - | - | - | 0.045 | 0.072 | 5.36E-01 | -0.097 | 0.186 | 0.046 | 0.073 | 5.25E-01 | -0.097 | 0.190 |
| | **dorsal body surface area (in SD)** | - | - | - | - | - | -0.044 | 0.061 | 4.67E-01 | -0.163 | 0.075 | -0.045 | 0.061 | 4.67E-01 | -0.165 | 0.076 |
| | **LDL cholesterol levels (in SD)** | - | - | - | - | - | - | - | - | - | - | 0.058 | 0.055 | 2.87E-01 | -0.049 | 0.165 |
| | **HDL cholesterol levels (in SD)** | - | - | - | - | - | - | - | - | - | - | 0.079 | 0.061 | 1.91E-01 | -0.040 | 0.199 |
| | **triglyceride levels (in SD)** | - | - | - | - | - | - | - | - | - | - | 0.034 | 0.074 | 6.50E-01 | -0.112 | 0.179 |
| | **glucose levels (in SD)** | - | - | - | - | - | - | - | - | - | - | -0.043 | 0.054 | 4.32E-01 | -0.149 | 0.064 |
| | **time of day (in hours since 9AM)** | 0.031 | 0.041 | 4.41E-01 | -0.048 | 0.111 | 0.030 | 0.046 | 5.16E-01 | -0.060 | 0.120 | 0.032 | 0.046 | 4.87E-01 | -0.058 | 0.122 |
| | **intercept** | 1.063 | 1.075 | 3.23E-01 | -1.044 | 3.170 | 1.346 | 1.122 | 2.30E-01 | -0.853 | 3.544 | 1.478 | 1.121 | 1.87E-01 | -0.719 | 3.675 |
| | ***ldlrb*** | -0.104 | 2.197 | 9.62E-01 | -4.411 | 4.202 | -0.347 | 2.258 | 8.78E-01 | -4.773 | 4.079 | -0.329 | 2.258 | 8.84E-01 | -4.755 | 4.097 |
| random factors | ***variation by batch*** | *0.514* | *0.140* | - | *0.301* | *0.877* | *0.509* | *0.143* | - | *0.293* | *0.884* | *0.479* | *0.138* | - | *0.272* | *0.844* |
| | ***residual*** | *0.858* | *0.032* | - | *0.798* | *0.923* | *0.878* | *0.035* | - | *0.812* | *0.949* | *0.874* | *0.035* | - | *0.809* | *0.944* |

***continued*** **Supplementary Table 24**

| | | Vascular co-localization of lipids with macrophages | | | | | | | | | | | | | | |
|---|---|---|---|---|---|---|---|---|---|---|---|---|---|---|---|---|
| | | Model 1 (n=301) | | | | | Model 2 (n=269) | | | | | Model 3 (n=269) | | | | |
| | | Effect | SE | *P* | lci | uci | Effect | SE | *P* | lci | uci | Effect | SE | *P* | lci | uci |
| negative binomial terms | ***apoea*** | -0.299 | 0.234 | 2.02E-01 | -0.758 | 0.160 | -0.410 | 0.246 | 9.58E-02 | -0.893 | 0.073 | -0.351 | 0.244 | 1.51E-01 | -0.830 | 0.128 |
| | ***apoeb*** | -0.142 | 0.180 | 4.31E-01 | -0.496 | 0.212 | -0.123 | 0.198 | 5.34E-01 | -0.512 | 0.265 | -0.113 | 0.198 | 5.67E-01 | -0.501 | 0.275 |
| | ***apoba*** | 0.123 | 0.760 | 8.71E-01 | -1.367 | 1.613 | 0.108 | 0.745 | 8.85E-01 | -1.353 | 1.569 | 0.456 | 0.805 | 5.71E-01 | -1.122 | 2.035 |
| | ***apobb.1*** | 1.049 | 0.204 | 2.71E-07 | 0.649 | 1.449 | 0.963 | 0.248 | 1.05E-04 | 0.476 | 1.449 | 0.881 | 0.273 | 1.27E-03 | 0.345 | 1.416 |
| | ***apobb.2*** | -0.154 | 0.475 | 7.46E-01 | -1.086 | 0.778 | -0.233 | 0.499 | 6.41E-01 | -1.211 | 0.746 | 0.002 | 0.496 | 9.96E-01 | -0.970 | 0.975 |
| | ***ldlra*** | -0.096 | 0.193 | 6.22E-01 | -0.475 | 0.284 | -0.079 | 0.191 | 6.79E-01 | -0.454 | 0.296 | -0.102 | 0.191 | 5.95E-01 | -0.476 | 0.273 |
| | **body length (in SD)** | - | - | - | - | - | -0.093 | 0.245 | 7.05E-01 | -0.574 | 0.388 | -0.060 | 0.256 | 8.14E-01 | -0.563 | 0.442 |
| | **dorsal body surface area (in SD)** | - | - | - | - | - | 0.096 | 0.152 | 5.30E-01 | -0.203 | 0.394 | 0.068 | 0.154 | 6.58E-01 | -0.233 | 0.369 |
| | **LDL cholesterol levels (in SD)** | - | - | - | - | - | - | - | - | - | - | -0.086 | 0.151 | 5.69E-01 | -0.382 | 0.210 |
| | **HDL cholesterol levels (in SD)** | - | - | - | - | - | - | - | - | - | - | 0.009 | 0.170 | 9.58E-01 | -0.325 | 0.343 |
| | **triglyceride levels (in SD)** | - | - | - | - | - | - | - | - | - | - | 0.316 | 0.235 | 1.78E-01 | -0.144 | 0.776 |
| | **glucose levels (in SD)** | - | - | - | - | - | - | - | - | - | - | -0.116 | 0.137 | 3.95E-01 | -0.384 | 0.151 |
| | **time of day (in hours since 9AM)** | 0.031 | 0.090 | 7.32E-01 | -0.146 | 0.208 | 0.026 | 0.115 | 8.23E-01 | -0.199 | 0.250 | 0.092 | 0.122 | 4.48E-01 | -0.146 | 0.331 |
| | **batch 1** | 1.673 | 0.598 | 5.14E-03 | 0.501 | 2.845 | 1.703 | 0.705 | 1.57E-02 | 0.321 | 3.086 | 1.674 | 0.786 | 3.32E-02 | 0.134 | 3.214 |
| | **batch 2** | 1.054 | 0.863 | 2.22E-01 | -0.637 | 2.745 | 0.884 | 0.862 | 3.05E-01 | -0.805 | 2.574 | 0.758 | 0.928 | 4.14E-01 | -1.060 | 2.577 |
| | **batch 3** | 0.934 | 0.618 | 1.30E-01 | -0.277 | 2.145 | 0.910 | 0.646 | 1.59E-01 | -0.356 | 2.176 | 0.785 | 0.664 | 2.37E-01 | -0.517 | 2.086 |
| | **batch 4** | 1.449 | 0.652 | 2.62E-02 | 0.171 | 2.727 | 1.271 | 0.677 | 6.02E-02 | -0.055 | 2.598 | 1.260 | 0.720 | 7.98E-02 | -0.150 | 2.671 |
| | **batch 5** | 2.334 | 0.705 | 9.37E-04 | 0.951 | 3.717 | 2.246 | 0.827 | 6.64E-03 | 0.624 | 3.867 | 1.773 | 0.883 | 4.46E-02 | 0.043 | 3.504 |
| | **batch 6** | 1.190 | 0.648 | 6.62E-02 | -0.080 | 2.460 | 1.252 | 0.865 | 1.48E-01 | -0.443 | 2.947 | 0.663 | 0.932 | 4.77E-01 | -1.164 | 2.489 |
| | **batch 7** | 0.737 | 0.971 | 4.48E-01 | -1.166 | 2.640 | 0.545 | 1.090 | 6.17E-01 | -1.592 | 2.682 | 0.141 | 1.122 | 9.00E-01 | -2.057 | 2.340 |
| | **intercept** | -5.120 | 1.899 | 7.02E-03 | -8.842 | -1.398 | -4.899 | 1.863 | 8.54E-03 | -8.550 | -1.248 | -5.421 | 1.971 | 5.95E-03 | -9.284 | -1.558 |
| | ***ldlrb*** | 14.867 | 3.709 | 6.11E-05 | 7.598 | 22.136 | 15.184 | 3.380 | 7.03E-06 | 8.560 | 21.808 | 14.218 | 3.406 | 2.98E-05 | 7.543 | 20.893 |

| | | Vascular infiltration by neutrophils | | | | | | | | | | | | | | |
|---|---|---|---|---|---|---|---|---|---|---|---|---|---|---|---|---|
| | | Model 1 (n=371) | | | | | Model 2 (n=330) | | | | | Model 3 (n=330) | | | | |
| | | Effect | SE | *P* | lci | uci | Effect | SE | *P* | lci | uci | Effect | SE | *P* | lci | uci |
| fixed factors | ***apoea*** | -0.047 | 0.076 | 5.36E-01 | -0.196 | 0.102 | -0.076 | 0.081 | 3.52E-01 | -0.234 | 0.083 | -0.083 | 0.082 | 3.11E-01 | -0.244 | 0.078 |
| | ***apoeb*** | 0.001 | 0.084 | 9.87E-01 | -0.162 | 0.165 | 0.028 | 0.090 | 7.54E-01 | -0.147 | 0.204 | 0.010 | 0.090 | 9.09E-01 | -0.166 | 0.186 |
| | ***apoba*** | 0.099 | 0.281 | 7.25E-01 | -0.452 | 0.650 | 0.111 | 0.296 | 7.07E-01 | -0.469 | 0.692 | 0.110 | 0.295 | 7.10E-01 | -0.469 | 0.689 |
| | ***apobb.1*** | 0.103 | 0.082 | 2.07E-01 | -0.057 | 0.264 | 0.042 | 0.088 | 6.32E-01 | -0.130 | 0.214 | 0.061 | 0.089 | 4.96E-01 | -0.114 | 0.235 |
| | ***apobb.2*** | -0.056 | 0.198 | 7.79E-01 | -0.443 | 0.332 | 0.021 | 0.210 | 9.19E-01 | -0.391 | 0.434 | 0.006 | 0.212 | 9.76E-01 | -0.410 | 0.423 |
| | ***ldlra*** | -0.074 | 0.064 | 2.51E-01 | -0.200 | 0.052 | -0.066 | 0.070 | 3.47E-01 | -0.204 | 0.072 | -0.069 | 0.070 | 3.29E-01 | -0.207 | 0.069 |
| | **body length (in SD)** | - | - | - | - | - | -0.111 | 0.071 | 1.19E-01 | -0.250 | 0.028 | -0.098 | 0.072 | 1.71E-01 | -0.239 | 0.043 |
| | **dorsal body surface area (in SD)** | - | - | - | - | - | 0.050 | 0.060 | 4.00E-01 | -0.067 | 0.168 | 0.067 | 0.060 | 2.67E-01 | -0.051 | 0.186 |
| | **LDL cholesterol levels (in SD)** | - | - | - | - | - | - | - | - | - | - | 0.006 | 0.054 | 9.15E-01 | -0.099 | 0.111 |
| | **HDL cholesterol levels (in SD)** | - | - | - | - | - | - | - | - | - | - | 0.083 | 0.060 | 1.72E-01 | -0.036 | 0.201 |
| | **triglyceride levels (in SD)** | - | - | - | - | - | - | - | - | - | - | -0.054 | 0.072 | 4.50E-01 | -0.195 | 0.086 |
| | **glucose levels (in SD)** | - | - | - | - | - | - | - | - | - | - | 0.062 | 0.054 | 2.55E-01 | -0.045 | 0.169 |
| | **time of day (in hours since 9AM)** | 0.016 | 0.041 | 6.95E-01 | -0.064 | 0.095 | -0.020 | 0.045 | 6.63E-01 | -0.107 | 0.068 | -0.022 | 0.045 | 6.28E-01 | -0.109 | 0.066 |
| | **intercept** | 0.777 | 1.093 | 4.77E-01 | -1.364 | 2.919 | 0.584 | 1.122 | 6.03E-01 | -1.615 | 2.782 | 0.617 | 1.122 | 5.82E-01 | -1.582 | 2.815 |
| | ***ldlrb*** | -2.049 | 2.254 | 3.63E-01 | -6.467 | 2.369 | -1.571 | 2.281 | 4.91E-01 | -6.041 | 2.900 | -1.525 | 2.285 | 5.05E-01 | -6.005 | 2.954 |
| random factors | ***variation by batch*** | *0.352* | *0.100* | - | *0.201* | *0.615* | *0.323* | *0.097* | - | *0.179* | *0.583* | *0.296* | *0.092* | - | *0.161* | *0.543* |
| | ***residual*** | *0.882* | *0.033* | - | *0.820* | *0.948* | *0.888* | *0.035* | - | *0.822* | *0.959* | *0.885* | *0.035* | - | *0.819* | *0.956* |

***continued*** **Supplementary Table 24**

| | | Vascular co-localization of lipids with neutrophils | | | | | | | | | | | | | | |
|---|---|---|---|---|---|---|---|---|---|---|---|---|---|---|---|---|
| | | Model 1 (n=282) | | | | | Model 2 (n=250) | | | | | Model 3 (n=250) | | | | |
| | | Effect | SE | *P* | lci | uci | Effect | SE | *P* | lci | uci | Effect | SE | *P* | lci | uci |
| negative binomial terms | ***apoea*** | -0.154 | 0.268 | 5.65E-01 | -0.679 | 0.371 | -0.015 | 0.284 | 9.58E-01 | -0.572 | 0.542 | -0.361 | 0.318 | 2.57E-01 | -0.985 | 0.263 |
| | ***apoeb*** | 0.079 | 0.270 | 7.71E-01 | -0.451 | 0.608 | 0.177 | 0.278 | 5.24E-01 | -0.369 | 0.723 | 0.181 | 0.277 | 5.14E-01 | -0.361 | 0.723 |
| | ***apoba*** | 0.520 | 0.995 | 6.01E-01 | -1.431 | 2.471 | 0.985 | 0.931 | 2.90E-01 | -0.840 | 2.811 | 1.882 | 1.019 | 6.48E-02 | -0.116 | 3.880 |
| | ***apobb.1*** | 1.722 | 0.263 | 5.80E-11 | 1.206 | 2.237 | 1.619 | 0.309 | 1.54E-07 | 1.015 | 2.224 | 1.547 | 0.357 | 1.43E-05 | 0.848 | 2.246 |
| | ***apobb.2*** | 0.441 | 0.640 | 4.91E-01 | -0.814 | 1.696 | 0.187 | 0.654 | 7.75E-01 | -1.095 | 1.468 | 1.103 | 0.719 | 1.25E-01 | -0.308 | 2.513 |
| | ***ldlra*** | 0.033 | 0.247 | 8.93E-01 | -0.451 | 0.517 | 0.040 | 0.254 | 8.76E-01 | -0.458 | 0.537 | -0.071 | 0.245 | 7.71E-01 | -0.551 | 0.408 |
| | **body length (in SD)** | - | - | - | - | - | -0.035 | 0.259 | 8.94E-01 | -0.542 | 0.473 | -0.272 | 0.275 | 3.22E-01 | -0.811 | 0.267 |
| | **dorsal body surface area (in SD)** | - | - | - | - | - | 0.122 | 0.231 | 5.96E-01 | -0.330 | 0.575 | -0.014 | 0.233 | 9.52E-01 | -0.471 | 0.443 |
| | **LDL cholesterol levels (in SD)** | - | - | - | - | - | - | - | - | - | - | 0.620 | 0.267 | 2.04E-02 | 0.096 | 1.143 |
| | **HDL cholesterol levels (in SD)** | - | - | - | - | - | - | - | - | - | - | 0.513 | 0.331 | 1.22E-01 | -0.137 | 1.162 |
| | **triglyceride levels (in SD)** | - | - | - | - | - | - | - | - | - | - | 1.085 | 0.342 | 1.49E-03 | 0.415 | 1.754 |
| | **glucose levels (in SD)** | - | - | - | - | - | - | - | - | - | - | 0.339 | 0.202 | 9.27E-02 | -0.056 | 0.735 |
| | **time of day (in hours since 9AM)** | -0.184 | 0.167 | 2.69E-01 | -0.511 | 0.142 | -0.151 | 0.207 | 4.66E-01 | -0.557 | 0.255 | -0.085 | 0.207 | 6.82E-01 | -0.491 | 0.321 |
| | **batch 2** | -1.524 | 0.842 | 7.03E-02 | -3.174 | 0.126 | -1.617 | 0.913 | 7.67E-02 | -3.407 | 0.173 | -0.683 | 1.032 | 5.08E-01 | -2.706 | 1.340 |
| | **batch 3** | -0.627 | 0.721 | 3.84E-01 | -2.039 | 0.785 | -0.692 | 0.679 | 3.08E-01 | -2.023 | 0.638 | -1.417 | 0.659 | 3.15E-02 | -2.709 | -0.125 |
| | **batch 4** | -0.239 | 0.580 | 6.80E-01 | -1.376 | 0.897 | -0.154 | 0.654 | 8.14E-01 | -1.437 | 1.128 | -0.859 | 0.667 | 1.98E-01 | -2.167 | 0.449 |
| | **batch 5** | 0.980 | 0.549 | 7.45E-02 | -0.097 | 2.056 | 0.862 | 0.629 | 1.71E-01 | -0.371 | 2.094 | -1.191 | 0.923 | 1.97E-01 | -3.001 | 0.619 |
| | **batch 6** | -1.064 | 0.550 | 5.31E-02 | -2.143 | 0.014 | -0.926 | 0.600 | 1.23E-01 | -2.102 | 0.250 | -2.069 | 0.970 | 3.30E-02 | -3.971 | -0.168 |
| | **batch 7** | -1.512 | 0.951 | 1.12E-01 | -3.377 | 0.353 | -1.677 | 1.009 | 9.65E-02 | -3.654 | 0.300 | -3.366 | 1.259 | 7.52E-03 | -5.834 | -0.898 |
| | **intercept** | -28.073 | 216.656 | 8.97E-01 | -452.711 | 396.565 | 61.871 | 235.125 | 7.92E-01 | -398.965 | 522.707 | 201.886 | 254.650 | 4.28E-01 | -297.219 | 700.991 |
| | ***ldlrb*** | 66.921 | 543.258 | 9.02E-01 | -997.846 | 1131.688 | -160.054 | 589.423 | 7.86E-01 | -1300.000 | 995.195 | -515.165 | 638.098 | 4.19E-01 | -1800.000 | 735.485 |

| | | Vascular co-localization of macrophages with neutrophils | | | | | | | | | | | | | | |
|---|---|---|---|---|---|---|---|---|---|---|---|---|---|---|---|---|
| | | Model 1 (n=367) | | | | | Model 2 (n=327) | | | | | Model 3 (n=327) | | | | |
| | | Effect | SE | *P* | lci | uci | Effect | SE | *P* | lci | uci | Effect | SE | *P* | lci | uci |
| negative binomial terms | ***apoea*** | -0.102 | 0.116 | 3.77E-01 | -0.329 | 0.125 | -0.094 | 0.123 | 4.45E-01 | -0.334 | 0.147 | -0.092 | 0.124 | 4.58E-01 | -0.335 | 0.151 |
| | ***apoeb*** | 0.133 | 0.118 | 2.60E-01 | -0.099 | 0.366 | 0.136 | 0.126 | 2.81E-01 | -0.111 | 0.383 | 0.121 | 0.124 | 3.30E-01 | -0.122 | 0.363 |
| | ***apoba*** | 0.055 | 0.417 | 8.96E-01 | -0.762 | 0.871 | 0.302 | 0.411 | 4.63E-01 | -0.504 | 1.108 | 0.350 | 0.410 | 3.93E-01 | -0.453 | 1.153 |
| | ***apobb.1*** | 0.036 | 0.140 | 7.96E-01 | -0.237 | 0.310 | 0.079 | 0.141 | 5.77E-01 | -0.198 | 0.356 | 0.109 | 0.143 | 4.45E-01 | -0.172 | 0.391 |
| | ***apobb.2*** | -0.252 | 0.300 | 4.01E-01 | -0.840 | 0.336 | -0.387 | 0.335 | 2.48E-01 | -1.044 | 0.269 | -0.297 | 0.320 | 3.53E-01 | -0.924 | 0.330 |
| | ***ldlra*** | -0.141 | 0.108 | 1.90E-01 | -0.352 | 0.070 | -0.189 | 0.108 | 7.87E-02 | -0.401 | 0.022 | -0.178 | 0.109 | 1.02E-01 | -0.392 | 0.035 |
| | **body length (in SD)** | - | - | - | - | - | 0.308 | 0.112 | 5.73E-03 | 0.090 | 0.527 | 0.269 | 0.120 | 2.51E-02 | 0.034 | 0.505 |
| | **dorsal body surface area (in SD)** | - | - | - | - | - | 0.271 | 0.110 | 1.35E-02 | 0.056 | 0.487 | 0.267 | 0.116 | 2.16E-02 | 0.039 | 0.495 |
| | **LDL cholesterol levels (in SD)** | - | - | - | - | - | - | - | - | - | - | 0.066 | 0.092 | 4.72E-01 | -0.114 | 0.247 |
| | **HDL cholesterol levels (in SD)** | - | - | - | - | - | - | - | - | - | - | -0.016 | 0.100 | 8.73E-01 | -0.213 | 0.181 |
| | **triglyceride levels (in SD)** | - | - | - | - | - | - | - | - | - | - | -0.049 | 0.123 | 6.93E-01 | -0.290 | 0.193 |
| | **glucose levels (in SD)** | - | - | - | - | - | - | - | - | - | - | 0.119 | 0.081 | 1.44E-01 | -0.041 | 0.278 |
| | **time of day (in hours since 9AM)** | 0.074 | 0.068 | 2.75E-01 | -0.059 | 0.206 | 0.086 | 0.073 | 2.38E-01 | -0.057 | 0.228 | 0.073 | 0.075 | 3.33E-01 | -0.075 | 0.220 |
| | **batch 1** | 0.378 | 0.325 | 2.45E-01 | -0.259 | 1.014 | 0.123 | 0.361 | 7.34E-01 | -0.585 | 0.831 | 0.236 | 0.418 | 5.72E-01 | -0.583 | 1.056 |
| | **batch 2** | 0.213 | 0.485 | 6.61E-01 | -0.738 | 1.165 | 0.222 | 0.454 | 6.26E-01 | -0.669 | 1.112 | 0.353 | 0.477 | 4.60E-01 | -0.583 | 1.289 |
| | **batch 3** | -0.552 | 0.397 | 1.64E-01 | -1.331 | 0.226 | -0.656 | 0.403 | 1.04E-01 | -1.445 | 0.134 | -0.534 | 0.429 | 2.13E-01 | -1.376 | 0.307 |
| | **batch 4** | 0.911 | 0.392 | 2.01E-02 | 0.143 | 1.679 | 0.999 | 0.417 | 1.65E-02 | 0.182 | 1.816 | 0.992 | 0.445 | 2.57E-02 | 0.120 | 1.863 |
| | **batch 5** | 0.171 | 0.373 | 6.46E-01 | -0.560 | 0.903 | -0.182 | 0.394 | 6.44E-01 | -0.955 | 0.590 | -0.012 | 0.441 | 9.78E-01 | -0.877 | 0.852 |
| | **batch 6** | -1.591 | 0.383 | 3.32E-05 | -2.342 | -0.840 | -2.219 | 0.456 | 1.13E-06 | -3.113 | -1.326 | -2.105 | 0.475 | 9.32E-06 | -3.036 | -1.174 |
| | **batch 7** | -0.432 | 0.491 | 3.79E-01 | -1.395 | 0.531 | -1.048 | 0.569 | 6.57E-02 | -2.163 | 0.068 | -1.007 | 0.646 | 1.19E-01 | -2.273 | 0.258 |
| | **intercept** | 3.948 | 0.846 | 3.06E-06 | 2.290 | 5.605 | 3.908 | 0.912 | 1.82E-05 | 2.121 | 5.695 | 3.376 | 0.937 | 3.16E-04 | 1.539 | 5.213 |
| | ***ldlrb*** | 0.044 | 0.776 | 9.54E-01 | -1.477 | 1.566 | 0.090 | 0.894 | 9.20E-01 | -1.662 | 1.843 | 0.663 | 1.014 | 5.14E-01 | -1.326 | 2.651 |

All outcomes were normalized for length using residuals, and inverse-normally transformed before the analysis. Associations were examined using hierarchical linear models. Effects shown are for each additional mutated allele in *apoea* , *apoeb* , *apoba* , *apobb.1,* *apobb.2* , *ldlra* and *ldlrb,* weighted by the allele's predicted effect on protein function (i.e. additive model, mutually adjusted). Associations were adjusted for time of day and batch. Lci and uci are lower and upper boundaries of the 95% confidence interval.

**Supplementary Table 25 - The additive effect of mutated alleles in *apoea*, *apoeb*, *apoba*, *apobb.1*, *apobb.2*, *ldlra* and *ldlrb* on image and image quantification quality**

| | Debris included in the segmentation for body size (n=374) | | | | |
|---|---|---|---|---|---|
| | OR | SE | *P* | lci | uci |
| ***apoea*** | 0.580 | 0.179 | 7.70E-02 | 0.317 | 1.061 |
| ***apoeb*** | 1.511 | 0.525 | 2.34E-01 | 0.765 | 2.986 |
| ***apoba*** | 2.543 | 3.029 | 4.33E-01 | 0.246 | 26.260 |
| ***apobb.1*** | 0.569 | 0.189 | 8.91E-02 | 0.297 | 1.090 |
| ***apobb.2*** | 2.123 | 1.781 | 3.69E-01 | 0.410 | 10.993 |
| ***ldlra*** | 0.791 | 0.187 | 3.22E-01 | 0.497 | 1.258 |
| ***ldlrb*** | 355 | 13000 | 8.74E-01 | 0.000 | - |
| **time of day (in hours since 9AM)** | 1.088 | 0.148 | 5.33E-01 | 0.834 | 1.420 |
| **intercept** | 0.001 | 0.009 | 6.20E-01 | 0.000 | - |

| | Many false positive lipid deposits | | | | | | | | | | | | | | | | | | | |
|---|---|---|---|---|---|---|---|---|---|---|---|---|---|---|---|---|---|---|---|---|
| | Model 1 (n=373) | | | | | Model 2 (n=245) | | | | | Model 3 (n=245) | | | | | Model 4 (n=245) | | | | |
| | OR | SE | *P* | lci | uci | OR | SE | *P* | lci | uci | OR | SE | *P* | lci | uci | OR | SE | *P* | lci | uci |
| ***apoea*** | 0.837 | 0.187 | 4.26E-01 | 0.539 | 1.298 | 0.979 | 0.295 | 9.44E-01 | 0.542 | 1.767 | 1.081 | 0.343 | 8.07E-01 | 0.580 | 2.014 | 1.284 | 0.429 | 4.54E-01 | 0.667 | 2.472 |
| ***apoeb*** | 0.918 | 0.234 | 7.38E-01 | 0.557 | 1.514 | 0.899 | 0.311 | 7.57E-01 | 0.456 | 1.771 | 0.919 | 0.322 | 8.09E-01 | 0.462 | 1.827 | 0.907 | 0.324 | 7.85E-01 | 0.450 | 1.828 |
| ***apoba*** | 0.845 | 0.698 | 8.38E-01 | 0.167 | 4.264 | 0.189 | 0.230 | 1.72E-01 | 0.017 | 2.065 | 0.189 | 0.240 | 1.89E-01 | 0.016 | 2.265 | 0.213 | 0.279 | 2.37E-01 | 0.017 | 2.755 |
| ***apobb.1*** | 0.842 | 0.201 | 4.70E-01 | 0.527 | 1.344 | 0.658 | 0.212 | 1.94E-01 | 0.349 | 1.238 | 0.640 | 0.206 | 1.66E-01 | 0.340 | 1.203 | 0.660 | 0.217 | 2.06E-01 | 0.347 | 1.256 |
| ***apobb.2*** | 0.514 | 0.300 | 2.54E-01 | 0.164 | 1.611 | 0.202 | 0.244 | 1.85E-01 | 0.019 | 2.145 | 0.296 | 0.371 | 3.31E-01 | 0.025 | 3.455 | 0.343 | 0.453 | 4.18E-01 | 0.026 | 4.571 |
| ***ldlra*** | 1.175 | 0.222 | 3.93E-01 | 0.812 | 1.701 | 1.507 | 0.385 | 1.09E-01 | 0.913 | 2.487 | 1.544 | 0.402 | 9.54E-02 | 0.927 | 2.573 | 1.623 | 0.442 | 7.50E-02 | 0.952 | 2.766 |
| ***ldlrb*** | 1927 | 58000 | 8.03E-01 | 0.000 | - | - | - | - | - | - | - | - | - | - | - | - | - | - | - | - |
| **time of day (in hours since 9AM)** | 1.243 | 0.129 | 3.62E-02 | 1.014 | 1.524 | 0.998 | 0.143 | 9.90E-01 | 0.754 | 1.321 | 0.989 | 0.142 | 9.37E-01 | 0.747 | 1.309 | 0.977 | 0.143 | 8.77E-01 | 0.733 | 1.303 |
| **body length (in SD)** | - | - | - | - | - | 1.838 | 0.423 | 8.20E-03 | 1.170 | 2.885 | 1.613 | 0.385 | 4.52E-02 | 1.010 | 2.576 | 1.667 | 0.427 | 4.58E-02 | 1.010 | 2.754 |
| **dorsal body surface area (in SD)** | - | - | - | - | - | 2.156 | 0.448 | 2.17E-04 | 1.435 | 3.240 | 1.959 | 0.427 | 2.02E-03 | 1.278 | 3.002 | 1.901 | 0.418 | 3.44E-03 | 1.236 | 2.924 |
| **LDL cholesterol levels (in SD)** | - | - | - | - | - | - | - | - | - | - | - | - | - | - | - | 0.580 | 0.113 | 5.30E-03 | 0.395 | 0.851 |
| **HDL cholesterol levels (in SD)** | - | - | - | - | - | - | - | - | - | - | - | - | - | - | - | 0.915 | 0.194 | 6.75E-01 | 0.605 | 1.385 |
| **triglyceride levels (in SD)** | - | - | - | - | - | - | - | - | - | - | 1.678 | 0.345 | 1.18E-02 | 1.122 | 2.509 | 1.718 | 0.364 | 1.06E-02 | 1.134 | 2.602 |
| **glucose levels (in SD)** | - | - | - | - | - | - | - | - | - | - | 1.035 | 0.176 | 8.40E-01 | 0.742 | 1.444 | 1.077 | 0.189 | 6.74E-01 | 0.763 | 1.519 |
| **intercept** | 0.022 | 0.270 | 7.56E-01 | 0.000 | | 38.951 | 132.322 | 2.81E-01 | 0.050 | 30000 | 19.083 | 67.600 | 4.05E-01 | 0.018 | 20000 | 9.277 | 34.040 | 5.44E-01 | 0.007 | 12000 |

Associations are shown for criteria that resulted in the exclusion of at least 10 larvae. Many false positives: >20% of true negative objects were falsely detected by the quantification pipeline. Associations were examined using logistic regression models. Model 1: adjusted for time of day; Model 2: additionally adjusted for body length and dorsal surface area; Model 3: additionally adjusted for whole-body triglyceride and glucose levels; Model 4: additionally adjusted for whole-body LDL and HDL cholesterol levels. Dorsal body surface area was normalized for body length using residuals; whole-body LDL cholesterol, HDL cholesterol, triglyceride and glucose levels were normalized for protein level using residuals. Effects shown for *apoea*, *apoeb*, *apoba*, *apobb.1*, *apobb.2* and *ldlra* are for each additional mutated allele. Adjusting for batch would have resulted in the exclusion of larvae. Lci and uci are lower and upper boundaries of the 95% confidence

**Supplementary Table 26 - The effect of gene x gene interactions on body size**

| | | Body length (n=339) | | | | | Dorsal body surface area (n=339) | | | | | Lateral body surface area (n=335) | | | | | Body volume (n=328) | | | | |
|---|---|---|---|---|---|---|---|---|---|---|---|---|---|---|---|---|---|---|---|---|---|
| | | **Effect** | **SE** | ***P*** | **lci** | **uci** | **Effect** | **SE** | ***P*** | **lci** | **uci** | **Effect** | **SE** | ***P*** | **lci** | **uci** | **Effect** | **SE** | ***P*** | **lci** | **uci** |
| **fixed factors** | ***apoea*** | -0.223 | 0.170 | 1.90E-01 | -0.557 | 0.110 | -0.155 | 0.204 | 4.49E-01 | -0.555 | 0.246 | -0.078 | 0.207 | 7.08E-01 | -0.484 | 0.328 | -0.055 | 0.206 | 7.88E-01 | -0.460 | 0.349 |
| | ***apoeb*** | -0.090 | 0.121 | 4.56E-01 | -0.327 | 0.147 | 0.069 | 0.145 | 6.36E-01 | -0.216 | 0.353 | 0.205 | 0.148 | 1.66E-01 | -0.085 | 0.495 | 0.167 | 0.149 | 2.61E-01 | -0.124 | 0.458 |
| | ***apobb.1*** | -0.119 | 0.215 | 5.78E-01 | -0.540 | 0.301 | -0.018 | 0.257 | 9.43E-01 | -0.523 | 0.486 | -0.180 | 0.266 | 4.99E-01 | -0.702 | 0.342 | -0.118 | 0.262 | 6.52E-01 | -0.633 | 0.396 |
| | ***ldlra*** | 0.179 | 0.208 | 3.89E-01 | -0.228 | 0.586 | -0.165 | 0.249 | 5.08E-01 | -0.653 | 0.324 | -0.223 | 0.254 | 3.80E-01 | -0.721 | 0.275 | -0.217 | 0.250 | 3.87E-01 | -0.707 | 0.274 |
| | ***apoea* x *apoeb*** | 0.087 | 0.107 | 4.19E-01 | -0.124 | 0.297 | 0.040 | 0.129 | 7.56E-01 | -0.212 | 0.293 | -0.071 | 0.131 | 5.88E-01 | -0.328 | 0.186 | -0.025 | 0.131 | 8.46E-01 | -0.281 | 0.231 |
| | ***apoea* x *apobb.1*** | 0.081 | 0.101 | 4.22E-01 | -0.117 | 0.278 | -0.059 | 0.121 | 6.26E-01 | -0.296 | 0.178 | 0.010 | 0.124 | 9.33E-01 | -0.232 | 0.253 | -0.024 | 0.124 | 8.46E-01 | -0.267 | 0.219 |
| | ***apoea* x *ldlra*** | 0.034 | 0.077 | 6.60E-01 | -0.116 | 0.184 | 0.002 | 0.092 | 9.85E-01 | -0.178 | 0.182 | 0.040 | 0.094 | 6.73E-01 | -0.144 | 0.223 | -0.007 | 0.093 | 9.36E-01 | -0.190 | 0.175 |
| | ***apoeb* x *apobb.1*** | -0.075 | 0.122 | 5.39E-01 | -0.315 | 0.165 | 0.083 | 0.147 | 5.71E-01 | -0.204 | 0.371 | 0.052 | 0.151 | 7.33E-01 | -0.245 | 0.348 | 0.057 | 0.149 | 7.02E-01 | -0.235 | 0.349 |
| | ***apoeb* x *ldlra*** | -0.036 | 0.103 | 7.30E-01 | -0.237 | 0.166 | 0.046 | 0.123 | 7.11E-01 | -0.196 | 0.288 | 0.060 | 0.126 | 6.34E-01 | -0.187 | 0.308 | 0.074 | 0.124 | 5.50E-01 | -0.170 | 0.318 |
| | ***apobb.1* x *ldlra*** | -0.156 | 0.081 | 5.56E-02 | -0.316 | 0.004 | 0.058 | 0.098 | 5.53E-01 | -0.133 | 0.250 | 0.106 | 0.099 | 2.88E-01 | -0.089 | 0.301 | 0.098 | 0.099 | 3.23E-01 | -0.096 | 0.291 |
| | **time of day (in hours since 9AM)** | -0.054 | 0.036 | 1.36E-01 | -0.125 | 0.017 | 0.039 | 0.043 | 3.60E-01 | -0.045 | 0.124 | 0.023 | 0.044 | 5.99E-01 | -0.063 | 0.109 | 0.047 | 0.044 | 2.82E-01 | -0.039 | 0.133 |
| | **intercept** | 0.284 | 0.288 | 3.25E-01 | -0.281 | 0.849 | -0.173 | 0.316 | 5.84E-01 | -0.793 | 0.447 | -0.231 | 0.319 | 4.68E-01 | -0.856 | 0.394 | -0.311 | 0.319 | 3.29E-01 | -0.936 | 0.314 |
| **random factors** | ***variance by batch*** | *0.510* | *0.138* | - | *0.300* | *0.866* | *0.469* | *0.133* | - | *0.269* | *0.818* | *0.467* | *0.131* | - | *0.269* | *0.809* | *0.469* | *0.131* | - | *0.271* | *0.813* |
| | ***residual*** | *0.696* | *0.027* | - | *0.645* | *0.751* | *0.835* | *0.033* | - | *0.774* | *0.902* | *0.845* | *0.033* | - | *0.783* | *0.912* | *0.829* | *0.033* | - | *0.767* | *0.895* |

Dorsal and lateral body surface area and body volume were normalized for body length using residuals. All outcomes were inverse-normally transformed before the analysis. Associations and interactions were examined using hierarchical linear models in larvae carrying two mutated alleles in apoba, apobb.2 and ldlrb. Associations and interactions were examined for each additional mutated allele, weighted by its predicted effect on protein function, i.e. using an additive model. Associations were adjusted for time of day and batch. Lci and uci are lower and upper boundaries of the 95% confidence interval.

**Supplementary Table 27 - The effect of gene x gene interactions on whole-body lipid and glucose levels**

| | | LDL cholesterol levels (n=381) | | | | |
|---|---|---|---|---|---|---|
| | | **Effect** | **SE** | ***P*** | **lci** | **uci** |
| **fixed factors** | ***apoea*** | -0.183 | 0.216 | 3.97E-01 | -0.606 | 0.240 |
| | ***apoeb*** | -0.014 | 0.147 | 9.22E-01 | -0.303 | 0.274 |
| | ***apobb.1*** | 0.103 | 0.284 | 7.16E-01 | -0.453 | 0.660 |
| | ***ldlra*** | 0.026 | 0.273 | 9.25E-01 | -0.510 | 0.561 |
| | ***apoea* x *apoeb*** | 0.103 | 0.136 | 4.48E-01 | -0.163 | 0.369 |
| | ***apoea* x *apobb.1*** | 0.070 | 0.132 | 5.97E-01 | -0.188 | 0.328 |
| | ***apoea* x *ldlra*** | 0.122 | 0.098 | 2.12E-01 | -0.070 | 0.314 |
| | ***apoeb* x *apobb.1*** | -0.023 | 0.161 | 8.86E-01 | -0.339 | 0.293 |
| | ***apoeb* x *ldlra*** | -0.049 | 0.135 | 7.17E-01 | -0.313 | 0.215 |
| | ***apobb.1* x *ldlra*** | -0.078 | 0.105 | 4.57E-01 | -0.285 | 0.128 |
| | **time of day (in hours since 9AM)** | 0.044 | 0.043 | 3.01E-01 | -0.039 | 0.128 |
| | **intercept** | -0.091 | 0.299 | 7.60E-01 | -0.678 | 0.495 |
| **random factors** | ***variance by batch*** | *0.350* | *0.120* | - | *0.179* | *0.685* |
| | ***residual*** | *0.945* | *0.035* | - | *0.880* | *1.016* |

| | | HDL cholesterol levels (n=381) | | | | |
|---|---|---|---|---|---|---|
| | | **Effect** | **SE** | ***P*** | **lci** | **uci** |
| **fixed factors** | ***apoea*** | 0.001 | 0.196 | 9.94E-01 | -0.383 | 0.386 |
| | ***apoeb*** | 0.036 | 0.134 | 7.89E-01 | -0.226 | 0.298 |
| | ***apobb.1*** | 0.188 | 0.258 | 4.66E-01 | -0.317 | 0.693 |
| | ***ldlra*** | 0.102 | 0.249 | 6.83E-01 | -0.386 | 0.589 |
| | ***apoea* x *apoeb*** | 0.113 | 0.123 | 3.62E-01 | -0.129 | 0.355 |
| | ***apoea* x *apobb.1*** | -0.077 | 0.120 | 5.21E-01 | -0.312 | 0.158 |
| | ***apoea* x *ldlra*** | -0.082 | 0.089 | 3.54E-01 | -0.257 | 0.092 |
| | ***apoeb* x *apobb.1*** | -0.074 | 0.147 | 6.14E-01 | -0.361 | 0.213 |
| | ***apoeb* x *ldlra*** | -0.004 | 0.123 | 9.72E-01 | -0.244 | 0.236 |
| | ***apobb.1* x *ldlra*** | 0.033 | 0.096 | 7.31E-01 | -0.155 | 0.220 |
| | **time of day (in hours since 9AM)** | -0.035 | 0.040 | 3.82E-01 | -0.114 | 0.044 |
| | **intercept** | -0.158 | 0.325 | 6.28E-01 | -0.795 | 0.480 |
| **random factors** | ***variance by batch*** | *0.592* | *0.160* | - | *0.348* | *1.006* |
| | ***residual*** | *0.858* | *0.031* | - | *0.799* | *0.922* |

| | | Triglyceride levels (n=381) | | | | |
|---|---|---|---|---|---|---|
| | | **Effect** | **SE** | ***P*** | **lci** | **uci** |
| **fixed factors** | ***apoea*** | -0.164 | 0.156 | 2.93E-01 | -0.469 | 0.141 |
| | ***apoeb*** | -0.123 | 0.106 | 2.47E-01 | -0.331 | 0.085 |
| | ***apobb.1*** | -0.163 | 0.205 | 4.26E-01 | -0.563 | 0.238 |
| | ***ldlra*** | -0.102 | 0.197 | 6.05E-01 | -0.489 | 0.285 |
| | ***apoea* x *apoeb*** | 0.054 | 0.098 | 5.84E-01 | -0.138 | 0.246 |
| | ***apoea* x *apobb.1*** | 0.063 | 0.095 | 5.09E-01 | -0.123 | 0.249 |
| | ***apoea* x *ldlra*** | -0.047 | 0.071 | 5.06E-01 | -0.185 | 0.091 |
| | ***apoeb* x *apobb.1*** | 0.138 | 0.116 | 2.34E-01 | -0.089 | 0.366 |
| | ***apoeb* x *ldlra*** | 0.088 | 0.097 | 3.68E-01 | -0.103 | 0.278 |
| | ***apobb.1* x *ldlra*** | 0.009 | 0.076 | 9.03E-01 | -0.139 | 0.158 |
| | **time of day (in hours since 9AM)** | -0.131 | 0.032 | 4.36E-05 | -0.194 | -0.068 |
| | **intercept** | 0.379 | 0.335 | 2.58E-01 | -0.277 | 1.035 |
| **random factors** | ***variance by batch*** | *0.763* | *0.196* | - | *0.462* | *1.261* |
| | ***residual*** | *0.680* | *0.025* | - | *0.633* | *0.731* |

| | | Total cholesterol levels (n=381) | | | | |
|---|---|---|---|---|---|---|
| | | **Effect** | **SE** | ***P*** | **lci** | **uci** |
| **fixed factors** | ***apoea*** | -0.041 | 0.185 | 8.26E-01 | -0.404 | 0.322 |
| | ***apoeb*** | 0.001 | 0.126 | 9.94E-01 | -0.246 | 0.248 |
| | ***apobb.1*** | -0.127 | 0.243 | 6.02E-01 | -0.604 | 0.350 |
| | ***ldlra*** | -0.088 | 0.235 | 7.07E-01 | -0.548 | 0.372 |
| | ***apoea* x *apoeb*** | 0.013 | 0.116 | 9.08E-01 | -0.215 | 0.242 |
| | ***apoea* x *apobb.1*** | 0.140 | 0.113 | 2.16E-01 | -0.082 | 0.361 |
| | ***apoea* x *ldlra*** | -0.030 | 0.084 | 7.16E-01 | -0.195 | 0.134 |
| | ***apoeb* x *apobb.1*** | -0.026 | 0.138 | 8.49E-01 | -0.297 | 0.245 |
| | ***apoeb* x *ldlra*** | 0.100 | 0.116 | 3.85E-01 | -0.126 | 0.327 |
| | ***apobb.1* x *ldlra*** | -0.104 | 0.090 | 2.50E-01 | -0.281 | 0.073 |
| | **time of day (in hours since 9AM)** | 0.079 | 0.038 | 3.92E-02 | 0.004 | 0.153 |
| | **intercept** | -0.074 | 0.353 | 8.35E-01 | -0.766 | 0.619 |
| **random factors** | ***variance by batch*** | *0.746* | *0.197* | - | *0.445* | *1.250* |
| | ***residual*** | *0.810* | *0.030* | - | *0.753* | *0.870* |

| | | Glucose levels (n=381) | | | | |
|---|---|---|---|---|---|---|
| | | **Effect** | **SE** | ***P*** | **lci** | **uci** |
| **fixed factors** | ***apoea*** | -0.068 | 0.216 | 7.51E-01 | -0.491 | 0.354 |
| | ***apoeb*** | -0.004 | 0.147 | 9.77E-01 | -0.292 | 0.284 |
| | ***apobb.1*** | -0.477 | 0.283 | 9.22E-02 | -1.032 | 0.078 |
| | ***ldlra*** | -0.101 | 0.273 | 7.12E-01 | -0.635 | 0.434 |
| | ***apoea* x *apoeb*** | 0.101 | 0.136 | 4.55E-01 | -0.165 | 0.367 |
| | ***apoea* x *apobb.1*** | -0.090 | 0.131 | 4.95E-01 | -0.347 | 0.168 |
| | ***apoea* x *ldlra*** | -0.087 | 0.098 | 3.75E-01 | -0.278 | 0.105 |
| | ***apoeb* x *apobb.1*** | 0.171 | 0.161 | 2.89E-01 | -0.145 | 0.486 |
| | ***apoeb* x *ldlra*** | 0.021 | 0.134 | 8.74E-01 | -0.242 | 0.285 |
| | ***apobb.1* x *ldlra*** | 0.087 | 0.105 | 4.08E-01 | -0.119 | 0.293 |
| | **time of day (in hours since 9AM)** | 0.052 | 0.043 | 2.20E-01 | -0.031 | 0.136 |
| | **intercept** | 0.046 | 0.303 | 8.78E-01 | -0.547 | 0.640 |
| **random factors** | ***variance by batch*** | *0.375* | *0.119* | - | *0.201* | *0.699* |
| | ***residual*** | *0.943* | *0.035* | - | *0.878* | *1.013* |

Dorsal and lateral body surface area and body volume were normalized for body length using residuals. All outcomes were inverse-normally transformed before the analysis. Associations and interactions were examined using hierarchical linear models in larvae carrying two mutated alleles in apoba, apobb.2 and ldlrb. Associations and interactions were examined for each additional mutated allele, weighted by its predicted effect on protein function, i.e. using an additive model. Associations were adjusted for time of day and batch. Lci and uci are lower and upper boundaries of the 95% confidence interval.

**Supplementary Table 28 - The effect of gene x gene interactions on image-based vascular atherogenic traits**

| | | Vascular lipid deposition | | | | | | | | | |
|---|---|---|---|---|---|---|---|---|---|---|---|
| | | 2 vs. 0 mutated alleles (n=62) | | | | | Additive model (n=272) | | | | |
| | | Effect | SE | *P* | lci | uci | Effect | SE | *P* | lci | uci |
| negative binomial terms | ***apoea*** | -3.380 | 0.895 | 1.59E-04 | -5.134 | -1.626 | -1.173 | 0.618 | 5.77E-02 | -2.385 | 0.038 |
| | ***apoeb*** | 0.078 | 0.312 | 8.01E-01 | -0.533 | 0.690 | 0.148 | 0.374 | 6.92E-01 | -0.586 | 0.882 |
| | ***apobb.1*** | 1.104 | 0.294 | 1.75E-04 | 0.527 | 1.680 | 0.675 | 0.563 | 2.30E-01 | -0.428 | 1.778 |
| | ***ldlra*** | -0.726 | 0.367 | 4.75E-02 | -1.445 | -0.008 | -0.481 | 0.520 | 3.55E-01 | -1.500 | 0.538 |
| | ***apoea* x *apoeb*** | - | - | - | - | - | 0.129 | 0.354 | 7.16E-01 | -0.564 | 0.821 |
| | ***apoea* x *apobb.1*** | - | - | - | - | - | 0.134 | 0.278 | 6.30E-01 | -0.411 | 0.678 |
| | ***apoea* x *ldlra*** | 4.368 | 1.256 | 5.07E-04 | 1.906 | 6.830 | 0.518 | 0.200 | 9.51E-03 | 0.127 | 0.910 |
| | ***apoeb* x *apobb.1*** | - | - | - | - | - | 0.008 | 0.328 | 9.80E-01 | -0.634 | 0.650 |
| | ***apoeb* x *ldlra*** | - | - | - | - | - | -0.024 | 0.260 | 9.28E-01 | -0.533 | 0.486 |
| | ***apobb.1* x *ldlra*** | - | - | - | - | - | 0.188 | 0.192 | 3.28E-01 | -0.189 | 0.565 |
| | **time of day (in hours since 9AM)** | 0.144 | 0.177 | 4.16E-01 | -0.204 | 0.492 | -0.168 | 0.108 | 1.19E-01 | -0.379 | 0.043 |
| | **body length (in SD)** | -0.129 | 0.270 | 6.32E-01 | -0.659 | 0.401 | -0.044 | 0.170 | 7.97E-01 | -0.377 | 0.289 |
| | **dorsal body surface area (in SD)** | -0.190 | 0.319 | 5.50E-01 | -0.815 | 0.434 | 0.156 | 0.125 | 2.13E-01 | -0.090 | 0.401 |
| | **batch 1** | - | - | - | - | - | 3.176 | 0.477 | 2.89E-11 | 2.241 | 4.112 |
| | **batch 2** | -0.500 | 1.066 | 6.39E-01 | -2.589 | 1.589 | 1.667 | 0.523 | 1.42E-03 | 0.643 | 2.691 |
| | **batch 3** | -0.479 | 0.871 | 5.82E-01 | -2.186 | 1.228 | 2.417 | 0.505 | 1.73E-06 | 1.426 | 3.407 |
| | **batch 4** | -1.364 | 0.832 | 1.01E-01 | -2.996 | 0.267 | 2.344 | 0.516 | 5.59E-06 | 1.333 | 3.356 |
| | **batch 5** | -0.177 | 0.716 | 8.05E-01 | -1.580 | 1.226 | 3.737 | 0.618 | 1.49E-09 | 2.526 | 4.949 |
| | **batch 6** | 1.265 | 0.763 | 9.76E-02 | -0.231 | 2.761 | 3.373 | 0.582 | 6.84E-09 | 2.232 | 4.514 |
| | **batch 7** | 0.398 | 0.843 | 6.37E-01 | -1.255 | 2.051 | 2.776 | 0.669 | 3.32E-05 | 1.465 | 4.088 |
| | **intercept** | 3.518 | 0.862 | 4.53E-05 | 1.827 | 5.208 | 2.330 | 0.646 | 3.09E-04 | 1.064 | 3.596 |

| | | Vascular co-localization of lipids with macrophages | | | | | | | | | |
|---|---|---|---|---|---|---|---|---|---|---|---|
| | | 2 vs. 0 mutated alleles (n=63) | | | | | Additive model (n=269) | | | | |
| | | Effect | SE | *P* | lci | uci | Effect | SE | *P* | lci | uci |
| negative binomial terms | ***apoea*** | -17.245 | 1.068 | 1.15E-58 | -19.338 | -15.152 | -0.324 | 0.757 | 6.68E-01 | -1.808 | 1.159 |
| | ***apoeb*** | -0.189 | 0.297 | 5.25E-01 | -0.771 | 0.393 | 0.264 | 0.477 | 5.80E-01 | -0.670 | 1.199 |
| | ***apobb.1*** | 1.360 | 0.463 | 3.32E-03 | 0.452 | 2.268 | 1.309 | 0.889 | 1.41E-01 | -0.433 | 3.050 |
| | ***ldlra*** | -1.833 | 0.671 | 6.34E-03 | -3.149 | -0.517 | -0.974 | 0.628 | 1.21E-01 | -2.205 | 0.257 |
| | ***apoea* x *apoeb*** | - | - | - | - | - | -0.419 | 0.419 | 3.18E-01 | -1.241 | 0.404 |
| | ***apoea* x *apobb.1*** | - | - | - | - | - | -0.491 | 0.463 | 2.89E-01 | -1.398 | 0.417 |
| | ***apoea* x *ldlra*** | 19.351 | 1.238 | 4.67E-55 | 16.924 | 21.778 | 0.843 | 0.327 | 1.00E-02 | 0.202 | 1.485 |
| | ***apoeb* x *apobb.1*** | - | - | - | - | - | 0.067 | 0.437 | 8.78E-01 | -0.789 | 0.923 |
| | ***apoeb* x *ldlra*** | - | - | - | - | - | 0.040 | 0.272 | 8.84E-01 | -0.493 | 0.572 |
| | ***apobb.1* x *ldlra*** | - | - | - | - | - | 0.066 | 0.283 | 8.16E-01 | -0.490 | 0.621 |
| | **time of day (in hours since 9AM)** | 0.429 | 0.204 | 3.55E-02 | 0.029 | 0.829 | -0.022 | 0.128 | 8.62E-01 | -0.272 | 0.228 |
| | **body length (in SD)** | -1.065 | 0.446 | 1.71E-02 | -1.940 | -0.190 | -0.123 | 0.255 | 6.29E-01 | -0.622 | 0.376 |
| | **dorsal body surface area (in SD)** | -0.253 | 0.382 | 5.08E-01 | -1.003 | 0.496 | 0.077 | 0.159 | 6.28E-01 | -0.234 | 0.388 |
| | **batch 1** | 3.022 | 1.075 | 4.93E-03 | 0.915 | 5.129 | 2.389 | 0.665 | 3.24E-04 | 1.087 | 3.692 |
| | **batch 2** | 0.013 | 1.734 | 9.94E-01 | -3.385 | 3.411 | 1.704 | 0.900 | 5.84E-02 | -0.061 | 3.468 |
| | **batch 3** | 0.181 | 1.070 | 8.66E-01 | -1.916 | 2.278 | 1.347 | 0.621 | 3.02E-02 | 0.129 | 2.564 |
| | **batch 4** | 0.071 | 1.197 | 9.53E-01 | -2.276 | 2.417 | 2.070 | 0.688 | 2.61E-03 | 0.723 | 3.418 |
| | **batch 5** | 1.672 | 1.212 | 1.68E-01 | -0.704 | 4.047 | 3.066 | 0.810 | 1.54E-04 | 1.478 | 4.654 |
| | **batch 6** | 2.193 | 1.367 | 1.09E-01 | -0.486 | 4.872 | 2.085 | 0.839 | 1.30E-02 | 0.440 | 3.729 |
| | **batch 7** | 0.385 | 1.696 | 8.20E-01 | -2.939 | 3.709 | 1.495 | 1.121 | 1.82E-01 | -0.702 | 3.692 |
| | **intercept** | -1.275 | 1.320 | 3.34E-01 | -3.862 | 1.312 | 0.298 | 0.848 | 7.25E-01 | -1.364 | 1.960 |

Dorsal and lateral body surface area and body volume were normalized for body length using residuals. All outcomes were inverse-normally transformed before the analysis. Associations and interactions were examined using hierarchical linear models in larvae carrying two mutated alleles in apoba, apobb.2 and ldlrb. Associations and interactions were examined for each additional mutated allele, weighted by its predicted effect on protein function, i.e. using an additive model. Associations were adjusted for time of day and batch. Lci and uci are lower and upper boundaries of the 95% confidence interval.

**Supplementary Table 29 - The association of image-based vascular atherogenic traits with whole-body lipid and glucose levels**

| | | Vascular lipid deposition (n=1,118) | | | | |
|---|---|---|---|---|---|---|
| | | Effect | SE | *P* | lci | uci |
| negative binomial terms | LDL cholesterol levels (in SD) | -0.010 | 0.080 | 8.92E-01 | -0.160 | 0.140 |
| | HDL cholesterol levels (in SD) | -0.060 | 0.080 | 4.65E-01 | -0.220 | 0.100 |
| | triglyceride levels (in SD) | 0.600 | 0.110 | 2.00E-07 | 0.370 | 0.820 |
| | glucose levels (in SD) | -0.370 | 0.100 | 3.57E-04 | -0.570 | -0.170 |
| | body length (in SD) | 0.110 | 0.110 | 3.07E-01 | -0.100 | 0.310 |
| | dorsal body surface area (in SD) | 0.070 | 0.100 | 4.95E-01 | -0.130 | 0.260 |
| | Tg(hsp70:IK17:EGFP; mpeg1:mCherry) carriers vs. Tg(mpo:EGFP) & Tg(mpeg1:mCherry) carriers | -3.040 | 0.640 | 2.18E-06 | -4.290 | -1.780 |
| | Tg(flk:EGFP) carriers vs. Tg(mpo:EGFP; mpeg1:mCherry) carriers | -1.710 | 0.410 | 3.72E-05 | -2.520 | -0.900 |
| | batch 15 | 0.790 | 0.580 | 1.71E-01 | -0.340 | 1.920 |
| | batch 16 | 0.360 | 0.590 | 5.42E-01 | -0.790 | 1.510 |
| | batch 17 | 0.980 | 0.600 | 1.03E-01 | -0.200 | 2.160 |
| | batch 18 | 0.290 | 0.590 | 6.26E-01 | -0.870 | 1.450 |
| | batch 19 | 0.000 | 0.650 | 1.00E+00 | -1.270 | 1.270 |
| | batch 33 | -0.620 | 0.410 | 1.31E-01 | -1.430 | 0.190 |
| | batch 34 | -0.280 | 0.460 | 5.42E-01 | -1.190 | 0.630 |
| | batch 36 | 0.380 | 0.660 | 5.61E-01 | -0.910 | 1.680 |
| | batch 37 | -19.770 | 0.560 | 1.35E-269 | -20.870 | -18.660 |
| | batch 38 | 1.660 | 0.550 | 2.63E-03 | 0.580 | 2.740 |
| | batch 39 | -1.180 | 0.570 | 3.73E-02 | -2.300 | -0.070 |
| | batch 40 | 1.320 | 0.420 | 1.61E-03 | 0.500 | 2.140 |
| | batch 41 | 0.260 | 0.620 | 6.77E-01 | -0.960 | 1.470 |
| | batch 42 | 0.060 | 0.410 | 8.82E-01 | -0.750 | 0.870 |
| | batch 43 | 0.620 | 0.500 | 2.19E-01 | -0.370 | 1.600 |
| | batch 44 | 0.840 | 0.500 | 8.90E-02 | -0.130 | 1.820 |
| | batch 45 | 0.510 | 0.340 | 1.35E-01 | -0.160 | 1.180 |
| | intercept | 4.450 | 0.300 | 5.28E-51 | 3.870 | 5.030 |

| | | Vascular infiltration by oxidized LDL (n=677) | | | | |
|---|---|---|---|---|---|---|
| | | Effect | SE | *P* | lci | uci |
| negative binomial terms | LDL cholesterol levels (in SD) | -0.040 | 0.040 | 2.26E-01 | -0.110 | 0.030 |
| | HDL cholesterol levels (in SD) | 0.080 | 0.040 | 1.85E-02 | 0.010 | 0.160 |
| | triglyceride levels (in SD) | 0.090 | 0.060 | 1.31E-01 | -0.030 | 0.210 |
| | glucose levels (in SD) | 0.080 | 0.060 | 2.43E-01 | -0.050 | 0.200 |
| | body length (in SD) | 0.130 | 0.050 | 1.08E-02 | 0.030 | 0.240 |
| | dorsal body surface area (in SD) | 0.100 | 0.050 | 4.49E-02 | 0.000 | 0.190 |
| | batch 15 | 1.280 | 0.170 | 1.21E-14 | 0.960 | 1.610 |
| | batch 16 | 0.530 | 0.180 | 2.62E-03 | 0.190 | 0.880 |
| | batch 17 | 0.520 | 0.160 | 9.04E-04 | 0.210 | 0.830 |
| | batch 18 | 1.190 | 0.170 | 1.86E-12 | 0.860 | 1.520 |
| | batch 19 | 1.240 | 0.170 | 1.80E-13 | 0.910 | 1.570 |
| | batch 36 | 1.590 | 0.160 | 1.73E-22 | 1.270 | 1.910 |
| | batch 37 | 0.870 | 0.170 | 3.02E-07 | 0.540 | 1.210 |
| | batch 38 | 1.420 | 0.150 | 4.02E-21 | 1.130 | 1.720 |
| | intercept | 5.410 | 0.140 | 0.00E+00 | 5.140 | 5.680 |

| | | Vascular co-localization of lipids and macrophages (n=908) | | | | |
|---|---|---|---|---|---|---|
| | | Effect | SE | *P* | lci | uci |
| negative binomial terms | LDL cholesterol levels (in SD) | 0.140 | 0.130 | 2.89E-01 | -0.120 | 0.400 |
| | HDL cholesterol levels (in SD) | 0.030 | 0.130 | 8.38E-01 | -0.240 | 0.290 |
| | triglyceride levels (in SD) | 0.820 | 0.180 | 7.91E-06 | 0.460 | 1.190 |
| | glucose levels (in SD) | -0.340 | 0.160 | 3.14E-02 | -0.650 | -0.030 |
| | body length (in SD) | -0.280 | 0.190 | 1.33E-01 | -0.650 | 0.090 |
| | dorsal body surface area (in SD) | 0.200 | 0.170 | 2.32E-01 | -0.130 | 0.520 |
| | batch 39 | 0.380 | 0.920 | 6.82E-01 | -1.430 | 2.180 |
| | batch 40 | 2.480 | 0.680 | 2.75E-04 | 1.150 | 3.820 |
| | batch 41 | 1.380 | 0.900 | 1.23E-01 | -0.370 | 3.140 |
| | batch 42 | 0.910 | 0.620 | 1.44E-01 | -0.310 | 2.130 |
| | batch 43 | 2.030 | 0.710 | 4.22E-03 | 0.640 | 3.420 |
| | batch 44 | 1.970 | 0.700 | 5.04E-03 | 0.590 | 3.350 |
| | batch 45 | 0.780 | 0.590 | 1.83E-01 | -0.370 | 1.930 |
| | intercept | 1.180 | 0.530 | 2.46E-02 | 0.150 | 2.220 |

***continued*** **Supplementary Table 29**

| | | Vascular co-localization of macrophages and oxidized LDL (n=619) | | | | |
|---|---|---|---|---|---|---|
| | | **Effect** | **SE** | ***P*** | **lci** | **uci** |
| **negative binomial terms** | **LDL cholesterol levels (in SD)** | -0.080 | 0.050 | 1.43E-01 | -0.190 | 0.030 |
| | **HDL cholesterol levels (in SD)** | 0.020 | 0.050 | 7.23E-01 | -0.070 | 0.100 |
| | **triglyceride levels (in SD)** | 0.320 | 0.100 | 1.67E-03 | 0.120 | 0.530 |
| | **glucose levels (in SD)** | -0.190 | 0.090 | 2.77E-02 | -0.360 | -0.020 |
| | **body length (in SD)** | -0.050 | 0.080 | 5.52E-01 | -0.200 | 0.110 |
| | **dorsal body surface area (in SD)** | 0.010 | 0.070 | 9.07E-01 | -0.130 | 0.150 |
| | **batch 15** | 0.560 | 0.270 | 3.84E-02 | 0.030 | 1.080 |
| | **batch 16** | 0.230 | 0.290 | 4.35E-01 | -0.340 | 0.800 |
| | **batch 17** | -0.650 | 0.300 | 2.77E-02 | -1.240 | -0.070 |
| | **batch 18** | 0.430 | 0.270 | 1.18E-01 | -0.110 | 0.960 |
| | **batch 19** | 0.730 | 0.250 | 3.48E-03 | 0.240 | 1.210 |
| | **batch 36** | 0.320 | 0.250 | 2.11E-01 | -0.180 | 0.810 |
| | **batch 37** | 0.320 | 0.290 | 2.58E-01 | -0.240 | 0.880 |
| | **batch 38** | 1.250 | 0.240 | 1.80E-07 | 0.780 | 1.710 |
| | **intercept** | 2.540 | 0.230 | 3.49E-29 | 2.090 | 2.980 |

| | | Vascular co-localization of lipids and neutrophils (n=271) | | | | |
|---|---|---|---|---|---|---|
| | | **Effect** | **SE** | ***P*** | **lci** | **uci** |
| **negative binomial terms** | **LDL cholesterol levels (in SD)** | 0.540 | 0.230 | 1.83E-02 | 0.090 | 1.000 |
| | **HDL cholesterol levels (in SD)** | 0.730 | 0.310 | 1.81E-02 | 0.120 | 1.330 |
| | **triglyceride levels (in SD)** | 1.100 | 0.260 | 3.53E-05 | 0.580 | 1.610 |
| | **glucose levels (in SD)** | 0.060 | 0.190 | 7.65E-01 | -0.320 | 0.440 |
| | **body length (in SD)** | -0.830 | 0.220 | 2.05E-04 | -1.270 | -0.390 |
| | **dorsal body surface area (in SD)** | -0.200 | 0.200 | 3.13E-01 | -0.600 | 0.190 |
| | **batch 40** | 4.600 | 1.190 | 1.03E-04 | 2.280 | 6.930 |
| | **batch 41** | 3.710 | 1.520 | 1.44E-02 | 0.740 | 6.680 |
| | **batch 42** | 2.740 | 1.130 | 1.50E-02 | 0.530 | 4.950 |
| | **batch 43** | 3.020 | 1.030 | 3.24E-03 | 1.010 | 5.030 |
| | **batch 44** | 3.770 | 1.010 | 1.97E-04 | 1.790 | 5.760 |
| | **batch 45** | 3.770 | 1.110 | 6.87E-04 | 1.590 | 5.940 |
| | **batch 46** | 3.040 | 1.410 | 3.14E-02 | 0.270 | 5.800 |
| | **intercept** | -2.790 | 0.980 | 4.32E-03 | -4.710 | -0.870 |

| | | Vascular co-localization of macrophages and neutrophils (n=327) | | | | |
|---|---|---|---|---|---|---|
| | | **Effect** | **SE** | ***P*** | **lci** | **uci** |
| **negative binomial terms** | **LDL cholesterol levels (in SD)** | 0.110 | 0.090 | 2.36E-01 | -0.070 | 0.280 |
| | **HDL cholesterol levels (in SD)** | -0.060 | 0.100 | 5.65E-01 | -0.260 | 0.140 |
| | **triglyceride levels (in SD)** | -0.040 | 0.130 | 7.29E-01 | -0.290 | 0.210 |
| | **glucose levels (in SD)** | 0.110 | 0.080 | 1.69E-01 | -0.050 | 0.270 |
| | **body length (in SD)** | 0.200 | 0.130 | 1.21E-01 | -0.050 | 0.450 |
| | **dorsal body surface area (in SD)** | 0.230 | 0.120 | 4.87E-02 | 0.000 | 0.450 |
| | **batch 40** | 0.550 | 0.380 | 1.51E-01 | -0.200 | 1.300 |
| | **batch 41** | 0.660 | 0.500 | 1.89E-01 | -0.330 | 1.650 |
| | **batch 42** | -0.330 | 0.360 | 3.48E-01 | -1.030 | 0.360 |
| | **batch 43** | 1.400 | 0.500 | 4.56E-03 | 0.430 | 2.370 |
| | **batch 44** | 0.480 | 0.350 | 1.63E-01 | -0.200 | 1.160 |
| | **batch 45** | -1.630 | 0.390 | 2.39E-05 | -2.390 | -0.870 |
| | **batch 46** | -0.620 | 0.510 | 2.23E-01 | -1.620 | 0.380 |
| | **intercept** | 3.740 | 0.290 | 6.31E-38 | 3.170 | 4.310 |

Associations were examined using negative binomial regresion using data from the dietary, drug treatment and genetic proof-of-concept interventions combined. Dorsal body surface area was normalized for body length using residuals; whole-body LDL cholesterol, HDL cholesterol, triglyceride and glucose levels were normalized for protein level using residuals. Lci and uci are lower and upper boundaries of the 95% confidence interval and have been calculated using robust standard errors.

**Supplementary Table 30 - Orthologues of candidate genes in the triglyceride, LDLc and total cholesterol-associated locus on chr 19p13.11**

| Human gene | ENSG | Zebrafish orthologue | ENSDARG | Chr | Target %identity | Query %identity | Main human protein | Top hit BLAST | %identity (protein) | Conserved genes in locus |
|---|---|---|---|---|---|---|---|---|---|---|
| *LPAR2* | ENSG00000064547 | *lpar2a* | ENSDARG00000042338 | 3 | 49.16 | 49.86 | ENSP0000384665 | ENSDARP00000154087 | 54.95 | *PBX4, ATP13A1, GMIP* |
| | | *lpar2b* | ENSDARG00000042561 | 1 | 51.76 | 54.42 | | ENSDARP00000062425 | 60.00 | none |
| *GMIP* | ENSG00000089639 | *gmip* | ENSDARG00000077249 | 3 | 36.91 | 34.02 | ENSP00000203556 | ENSDARP00000131373 | 43.92 | *ATP13A1, PBX4, LPAR2* |
| *GATAD2A* | ENSG00000167491 | *gatad2ab* | ENSDARG00000006192 | 22 | 53.16 | 53.00 | ENSP00000351552 | ENSDARP00000115930 | 53.8 | *GMIP, CILP2, YJEFN3, TSSK6, TM6SF2, HAPLN4, NCAN, NR2C2AP, RFXANK, BORCS8, MEF2B, TMEM161A, SLC25A42, ARMC6, HOMER3* |
| *TM6SF2* | ENSG00000213996 | *tm6sf2* | ENSDARG00000029057 | 22 | 43.32 | 42.97 | ENSP00000374014 | ENSDARP00000118571 | 45.86 | *GMIP, HAPLN4, NCAN, NR2C2AP, RFXANK, BORCS8, MEF2B, TMEM161A, GATAD2A, TSSK6, YJEFN3,* |
| | | *zgc:85843* | ENSDARG00000105208 | 2 | 43.86 | 39.79 | | ENSDARP00000130339 | 45.67 | *HOMER3, GMIP, SCL25A42, ARMC6* |

Target %identity: percentage of the orthologous sequence matching the human sequence; Query %identity: percentage of the human sequence matching the sequence of the orthologue; Main human protein: Ensembl protein ID for the main transcript; %identity protein: percentage of the aligned query (input sequence, i.e. main human protein) which is identical to the subject (hit) sequence; conserved genes in locus: neighbouring genes conserved across danio rerio and homo sapiens locus according to Genomicus.

**Supplementary Table 31 - Identification of moderate-to-highly active CRISPR-Cas9 guide RNAs for 19p13.11 candidate genes**

| Human gene | Zebrafish orthologue | CRISPR gRNA target sequen | Genomic location (dabRer11/GRCz11) Chr | Pos | Exon | Strand | GC (%) | Self-complementarity | Off-targets 0 | 1 | 2 | 3 | Predicted efficiency | CRISPRscan score | Target activity (NA[a], no[b], low[c], moderate[d], high[e] or very high[f]) | Forward primer | Reverse primer | product size |
|---|---|---|---|---|---|---|---|---|---|---|---|---|---|---|---|---|---|---|
| *LPAR2* | *lpar2a* | GATGCCAGCGAAGAGGTC | 3 | 53,091,511 | 2 of 3 | + | 60 | 0 | 0 | 0 | 0 | 6 | 0.65 | - | NA | ATTGTCAACCGAAAGTTCCACT | TGAGGCTCATGTTAATCAATGC | 175 |
| | | **GgGCACTCATGACTTTGT** | **3** | **53,091,654** | **2 of 3** | **-** | **55** | **1** | **0** | **0** | **0** | **1** | **0.63** | **53** | **high** | **TTAACAGCTGTGAGGCTTTTGA** | **AGTGGAACTTTCGGTTGACAAT** | **186** |
| | *lpar2b* | **GGGGACTAAAGCGAAGG** | **1** | **59,206,929** | **2 of 7** | **-** | **65** | **1** | **0** | **0** | **0** | **0** | **0.76** | **76** | **very high** | **TTTTCTTCTACAGCATCAGCCA** | **CCAGCAGGTAATAAATGGGGTA** | **174** |
| | | GGTCAAACACAAGGAGAC | 1 | 59,204,504 | 3 of 7 | - | 55 | 0 | 0 | 0 | 1 | 1 | 0.70 | 49 | high, but only inframe variants | CTACACGCGCATCTTCATTTAC | CGCTAAAATGGAAGCTGAATCT | 187 |
| *GMIP* | *gmip* | **GgTCCACGCCATCCTCGC** | **3** | **52,763,869** | **5 of 20** | **-** | **70** | **1** | **0** | **0** | **0** | **0** | **0.68** | **-** | **moderate** | **GTTGATGCTGATGTGTTCGTTT** | **TCCAGCCTAATACAATGTGTCG** | **243** |
| | | gGTGAAGATCTGCCGTGAA | 3 | 52,753,647 | 4 of 20 | + | 50 | 0 | 0 | 0 | 0 | 1 | 0.62 | - | moderate | TGCCACTGGGAATATTAAAAGC | ACGAGTTCTCATCCCAAATCTC | 205 |
| *GATAD2A* | *gatad2ab* | GAGAAGATGTCAGAAGAC | 22 | 18,353,216 | 2 of 11 | + | 50 | 0 | 0 | 0 | 0 | 3 | 0.70 | 51 | NA | GGTCAGACCACTTGTATGTCCA | TCTACAGCTCCAGGTTCTCCTC | 207 |
| | | **GGTGAAGGCCACCATCA** | **22** | **18,353,404** | **2 of 11** | **+** | **60** | **1** | **0** | **0** | **1** | **1** | **0.63** | **51** | **moderate/high** | **GAGGAGAACCTGGAGCTGTAGA** | **ATGATCGACCCTAGTTAGCAGC** | **194** |
| *TM6SF2* | *tm6sf2* | GGTTATTCTTGAAATTGGC | 22 | 17,790,559 | 3 of 11 | + | 40 | 0 | 0 | 0 | 0 | 7 | 0.60 | 10 | no | ACAACCTCAGCATCCTATTTCC | GGATCCTTAGGAGGATTGAAGC | 239 |
| | | GAGGTCAATGACACACGT | 22 | 17,792,919 | 4 of 11 | - | 50 | 1 | 0 | 0 | 0 | 0 | 0.63 | 62 | NA | ACACCACAACACTGAAAGCAAT | ACGCTAACCGTTTTCTGGTAGA | 186 |
| | | **GgGTAGCGAGGTAGACCA** | **22** | **17,790,601** | **3 of 11** | **-** | **65** | **1** | **0** | **0** | **0** | **0** | **0.67** | **69** | **very high** | **ttctatcccacagcaagctaaa** | **aggatgaaatgcagaatgtgg** | **229** |
| | | GAGGATTGAAGCGCGTAG | 22 | 17,790,613 | 3 of 11 | - | 60 | 1 | 0 | 0 | 0 | 0 | 0.63 | 58 | very high | ttctatcccacagcaagctaaa | aggatgaaatgcagaatgtgg | 229 |
| | *zgc:85843* | GATGCTGACAACAGACGT | 2 | 56,897,287 | 3 of 10 | - | 50 | 0 | 0 | 0 | 4 | 2 | 0.63 | 54 | moderate | GATGGGGAGGAGAGAGGGAG* | AAAGTCCATGAAGCCTTTGATG* | 231 |
| | | gGGTCTTACCGTAGAACAG | 2 | 56,896,546 | 2 of 10 | - | 45 | 1 | 0 | 0 | 0 | 1 | 0.57 | 46 | NA | TGGATGTGTTGTGTGAAATGAT | GCCTAATTATCCTAACCTGCCCA | 241 |
| | | GgGATGATGAGCTACTGGG | 2 | 56,904,057 | 4 of 10 | + | 50 | 0 | 0 | 0 | 0 | 6 | 0.68 | 70 | very high | CTTGTGCTGTTGTTGTAGGCAG | cacaggagggcgaatagtttag | 244 |
| | | **gGACACTGAGGTTTGCCA** | **2** | **56,896,504** | **2 of 10** | **-** | **55** | **0** | **0** | **0** | **0** | **4** | **0.66** | **40** | **very high** | **TGAATGGATGTGTTGTGTGAAA** | **ttgtctccagaacaaacacctg** | **280** |

CRISPR gRNA target sequences were preferably selected based on location (i.e. in an early exon that affects all transcripts), complementarity (i.e. no complementarity), and free from predicted off targets. Target activity was examined by micro-injections in eight fertilized eggs in multiplex, followed by fragment length PCR analysis at 3 days post-fertilization. Results from target efficiency testing are shown, where NA: Not available due to failed capilary electrophoresis while estimating the length of the targeted region of an exon; No: 8 of 8 larvae test-injected with the gRNA only showed wildtype sequences; Low: 8 of 8 larvae showed wildtype sequence and fewer than 4 of 8 also contained indel sequence; Moderate: 8 of 8 larvae showed wildtype sequence and >4 of 8 also contained indel sequence; High: 8 of 8 larvae showed wildtype as well as indel sequence; Very high: Fewer than 4 of 8 larvae showed wildtype sequence and all larvae showed indel sequence. Target sequences highlighted in bold were selected and used to generate multiplexed mutant zebrafish.

**Supplementary Table 32 - Unique CRISPR-Cas9-induced mutations for 19p13.11 candidate genes**

| Zebrafish orthologue | Sequence | Annotation | Number of alleles | mean ± SD number of reads |
|---|---|---|---|---|
| *lpar2a* | **TATAGCCGCTATGACCAACATATTGGCCAGGATGATGAAAACACAAATGGGGAGCCCCAGACCGACCACCACAAAGTCATGAGTGCGCCAGTTTGGACTAATAGGCTTGCCAGTCCTGTTGTAGAAGTAAGACACTGGATTG** | **142M (wildtype reference)** | **1083** | **742 ± 537** |
| | TATAGCCGCTATGACCAACATATTGGCCAGGATGATGAAAACACAAATGGGGAGCCCCAGACCGACCACCAAAGTCATGAGTGCGCCAGTTTGGACTAATAGGCTTGCCAGTCCTGTTGTAGAAGTAAGACACTGGATTG | 69M2D71M | 8 | 512 ± 569 |
| | TATAGCCGCTATGACCAACATATTGGCCAGGATGATGAAAACACAAATGGGGAGCCCCAGACCGACCACCATGAGTGCGCCAGTTTGGACTAATAGGCTTGCCAGTCCTGTTGTAGAAGTAAGACACTGGATTG | 69M8D65M | 2 | 327 ± 65 |
| | TATAGCCGCTATGACCAACATATTGGCCAGGATGATGAAAACACAAATGGGGAGCCCCAGACCGACCAAAGTCATGAGTGCGCCAGTTTGGACTAATAGGCTTGCCAGTCCTGTTGTAGAAGTAAGACACTGGATTG | 66M5D71M | 1 | 673 |
| *lpar2b* | **GTGAAAGCGGCGGTTCATGAAGATGGCCGCCATCACCAGGATGTTGGTGAAGATGACAAAAACACTGACCAGCAGCCCCATACCCACCACCGCCTTCGCTTTAGTCCCCCACGTATCGCTGATGTTCTTA** | **130M (wildtype reference)** | **1086** | **310 ± 174** |
| | GTGAAAGCGGCGGTTCATGAAGATGGCCGCCATCACCAGGATGTTGGTGAAGATGACAAAAACACTGACCAGCAGCCCCATACCCACCACCTTCGCTTTAGTCCCCCACGTATCGCTGATGTTCTTA | 89M3D38M | 6 | 278 ± 91 |
| *gatad2ab* | **TGGACTCAAGACCAAAAGCGAACAGGCCAACAAGGTGGCGAACATCCTGCGGGCCGGAGAGGTGAAGGCCACCATCAAGGTGGAGGTGCAGACCAGCGACGAGCCCGTGGACATGAGCACATCCAAGAGGTTGGTCATGAAATAAGTCCA** | **150M (wildtype reference)** | **1062** | **1091 ± 567** |
| | TGGACTCAAGACCAAAAGCGAACAGGCCAACAAGGTGGCGAACATCCTGCGGGCCGGAGAGGTGAAGGTGGAGGTGCAGACCAGCGACGAGCCCGTGGACATGAGCACATCCAAGAGGTTGGTCATGAAATAAGTCCA | 64M12D74M | 27 | 584 ± 349 |
| | TGGACTCAAGACCAAAAGCGAACAGGCCAACAAGGTGGCGAACATCCTGCGGGCCGGAGAGGTGAAGGCCACCATCAGGTGGTGCAGACCAGCGGTGGTGCAGACCAGCGACGAGCCCGTGGACATGAGCACATCCAAGAGGTTGGTCATGAAATAAGTC[illegible] | 77M1S1M12I1M1S2M1S66M | 2 | 424 ± 98 |
| | TGAACTCAAGACCAAAAGCGAACAGGCCAACAAGGTGGCGAACATCCTGCGGGCCGGAGAGGTGAAGGCCAAGGTGGAGGTGCAGACCAGCGACGAGCCCGTGGACATGAGCACATCCAAGAGGTTGGTCATGAAATAAGTCCA | 2M1S66M6D75M | 2 | 323 ± 48 |
| | TGGACTCAAGACCAAAAGCGAACAGGCCAACAAGGTGGCGAACATCCTGCGGGCCGGAGAGGTGAAGGCCGGTGGAGGTGCAGACCAGCGACGAGCCCGTGGACATGAGCACATCCAAGAGGTTGGTCATGAAATAAGTCCA | 70M8D72M | 1 | 436 |

Annotation shows the sequential number of base pairs that - when compared with the reference genome from Ensembl - represent a match (M), deletion (D), insertion (I), or substitution (S). For each unique sequence, the number of alleles in which it was observed is shown, as well as the mean and standrard deviation for the number of reads that were observed for the sequence.

**Supplementary Table 33 - Unique CRISPR-Cas9-induced mutations in 19p13.11 candidate genes and their predicted functional consequences**

| Zebrafish orthologue | Chr | Start | End | Mutation | Nett base pair change | Annotation VEP | VEP impact | $n_{affected\ alleles}$ |
|---|---|---|---|---|---|---|---|---|
| ***lpar2a*** | 3 | 53,091,655 | 53,091,659 | CACCA/- | -5 | frameshift variant | high | 1 |
| | | 53,091,658 | 53,091,659 | CA/- | -2 | frameshift variant | high | 8 |
| | | 53,091,658 | 53,091,665 | CACAAAGT/- | -8 | frameshift variant | high | 2 |
| ***lpar2b*** | 1 | 59,206,932 | 59,206,934 | CCG/- | -3 | inframe deletion | moderate | 6 |
| ***gatad2ab*** | 22 | 18,353,408 | 18,353,419 | AAGGCCACCATC/- | -12 | inframe deletion | moderate | 27 |
| | | 18,353,413 | 18,353,418 | CACCAT/- | -6 | inframe deletion | moderate | 2 |
| | | 18,353,414 | 18,353,421 | ACCATCAA/- | -8 | frameshift variant | high | 1 |
| | | 18,353,421 | 18,353,421 | A/G | 0 | missense variant | moderate | 2 |
| | | 18,353,423 | 18,353,422 | -/TGGTGCAGACCA | 12 | inframe insertion | moderate | 2 |
| | | 18,353,424 | 18,353,424 | T/C | 0 | missense variant | moderate | 2 |
| | | 18,353,427 | 18,353,427 | A/T | 0 | missense variant | moderate | 2 |

VEP: Ensembl's variant effect predictor; $n_{affected\ alleles}$: the number of alleles across the sequenced larvae in which the variant was observed.

**Supplementary Table 34 - Sequencing results expressed in number of mutated alleles for 19p13.11 candidate genes**

| Zebrafish orthologue | Number of mutated alleles | | | Missing genotypes | Total | Non-missing | Mutant allele freq | $P_{HWE\_LR}$ |
|---|---|---|---|---|---|---|---|---|
| | **0** | **1** | **2** | | | | | |
| ***lpar2a*** | 536 | 11 | 0 | 5 | 552 | 547 | 0.010 | 0.99 |
| ***lpar2b*** | 540 | 6 | 0 | 6 | 552 | 546 | 0.005 | 0.02 |
| ***gatad2ab*** | 515 | 32 | 0 | 5 | 552 | 547 | 0.029 | 0.47 |

The number of mutated alleles does not take into account the mutation's probability of affecting protein function. Dosage scores summed across both alleles were used in the association analyses. Four larvae were wildtype controls used to exclude variants that are inherently present from influencing the results, and five more larvae were excluded for having more than two missing calls across the seven successfully sequenced orthologues. Only genes for which some larvae were successfully mutated are shown. $P_{HWE\_LR}$: *P*- value for a Hardy-Weinberg equilibrium likelihood-ratio chi-squared statistic, considering a ±30 base pair window around the CRISPR cut site as one locus.

**Supplementary Table 35 - The effect of a mutated allele in *lpar2a* , *lpar2b* and *gatad2ab* on body size**

| | | Body length (n=505) | | | | |
|---|---|---|---|---|---|---|
| | | **Effect** | **SE** | ***P*** | **lci** | **uci** |
| **fixed factors** | ***lpar2a*** | -0.016 | 0.243 | 9.47E-01 | -0.493 | 0.460 |
| | ***lpar2b*** | 1.075 | 0.490 | 2.80E-02 | 0.116 | 2.035 |
| | ***gatad2ab*** | -0.161 | 0.224 | 4.71E-01 | -0.600 | 0.277 |
| | **age (11dpf vs. 10dpf)** | 0.301 | 0.101 | 2.94E-03 | 0.103 | 0.500 |
| | **time of day (in hours since 9AM)** | 0.000 | 0.019 | 9.79E-01 | -0.036 | 0.037 |
| | **intercept** | 0.114 | 0.290 | 6.95E-01 | -0.454 | 0.682 |
| **random factors** | ***variance by batch*** | *0.779* | *0.025* | - | *0.732* | *0.829* |
| | ***residual*** | *0.673* | *0.200* | - | *0.376* | *1.204* |

| | | Dorsal body surface area (n=505) | | | | |
|---|---|---|---|---|---|---|
| | | **Effect** | **SE** | ***P*** | **lci** | **uci** |
| **fixed factors** | ***lpar2a*** | 0.016 | 0.263 | 9.53E-01 | -0.500 | 0.531 |
| | ***lpar2b*** | -0.831 | 0.530 | 1.17E-01 | -1.869 | 0.207 |
| | ***gatad2ab*** | 0.825 | 0.242 | 6.52E-04 | 0.351 | 1.299 |
| | **age (11dpf vs. 10dpf)** | 0.214 | 0.109 | 4.99E-02 | 0.000 | 0.429 |
| | **time of day (in hours since 9AM)** | 0.007 | 0.020 | 7.45E-01 | -0.033 | 0.046 |
| | **intercept** | 0.012 | 0.237 | 9.58E-01 | -0.452 | 0.477 |
| **random factors** | ***variance by batch*** | *0.843* | *0.027* | - | *0.792* | *0.897* |
| | ***residual*** | *0.527* | *0.158* | - | *0.292* | *0.949* |

| | | Lateral body surface area (n=502) | | | | |
|---|---|---|---|---|---|---|
| | | **Effect** | **SE** | ***P*** | **lci** | **uci** |
| **fixed factors** | ***lpar2a*** | 0.281 | 0.279 | 3.14E-01 | -0.265 | 0.827 |
| | ***lpar2b*** | 0.070 | 0.561 | 9.01E-01 | -1.030 | 1.169 |
| | ***gatad2ab*** | 0.710 | 0.256 | 5.59E-03 | 0.208 | 1.212 |
| | **age (11dpf vs. 10dpf)** | 0.263 | 0.115 | 2.27E-02 | 0.037 | 0.489 |
| | **time of day (in hours since 9AM)** | -0.040 | 0.021 | 6.13E-02 | -0.082 | 0.002 |
| | **intercept** | 0.170 | 0.213 | 4.26E-01 | -0.248 | 0.588 |
| **random factors** | ***variance by batch*** | *0.892* | *0.028* | - | *0.839* | *0.950* |
| | ***residual*** | *0.454* | *0.140* | - | *0.248* | *0.831* |

| | | Body volume (n=495) | | | | |
|---|---|---|---|---|---|---|
| | | **Effect** | **SE** | ***P*** | **lci** | **uci** |
| **fixed factors** | ***lpar2a*** | 0.200 | 0.276 | 4.69E-01 | -0.341 | 0.740 |
| | ***lpar2b*** | 0.152 | 0.606 | 8.03E-01 | -1.037 | 1.340 |
| | ***gatad2ab*** | 0.924 | 0.254 | 2.72E-04 | 0.426 | 1.421 |
| | **age (11dpf vs. 10dpf)** | 0.286 | 0.115 | 1.31E-02 | 0.060 | 0.512 |
| | **time of day (in hours since 9AM)** | -0.005 | 0.021 | 8.04E-01 | -0.047 | 0.037 |
| | **intercept** | 0.023 | 0.220 | 9.16E-01 | -0.408 | 0.454 |
| **random factors** | ***variance by batch*** | *0.883* | *0.028* | - | *0.830* | *0.940* |
| | ***residual*** | *0.474* | *0.146* | - | *0.259* | *0.866* |

All outcomes were normalized for length using residuals, and inverse-normally transformed before the analysis. Associations were examined using hierarchical linear models. Effects shown are for the effect of carrying a mutated allele in *lpar2a, lpar2b and gatad2ab* , weighted by the allele's predicted effect on protein function (i.e. additive model, mutually adjusted). All associations were adjusted for age (i.e. 11 vs. 10 days post fertilization (dpf), time of day and batch. Lci and uci are lower and upper boundaries of the 95% confidence interval.

**Supplementary Table 36 - The effect of a mutated allele in *lpar2a, lpar2b* and *gatad2ab* on whole-body lipid**

| | | LDL cholesterol levels (n=513) | | | | |
|---|---|---|---|---|---|---|
| | | **Effect** | **SE** | ***P*** | **lci** | **uci** |
| **fixed factors** | ***lpar2a*** | -0.607 | 0.310 | 5.00E-02 | -1.215 | 0.000 |
| | ***lpar2b*** | -0.539 | 0.724 | 4.57E-01 | -1.959 | 0.881 |
| | ***gatad2ab*** | -0.159 | 0.262 | 5.45E-01 | -0.673 | 0.356 |
| | **age (11dpf vs. 10dpf)** | -0.263 | 0.121 | 2.98E-02 | -0.500 | -0.026 |
| | **time of day (in hours since 9AM)** | 0.010 | 0.024 | 6.86E-01 | -0.037 | 0.056 |
| | **intercept** | 0.108 | 0.160 | 5.01E-01 | -0.206 | 0.422 |
| **random factors** | ***variance by batch*** | *0.946* | *0.030* | - | *0.889* | *1.006* |
| | ***residual*** | *0.284* | *0.101* | - | *0.141* | *0.572* |

| | | HDL cholesterol levels (n=513) | | | | |
|---|---|---|---|---|---|---|
| | | **Effect** | **SE** | ***P*** | **lci** | **uci** |
| **fixed factors** | ***lpar2a*** | -0.346 | 0.318 | 2.77E-01 | -0.970 | 0.278 |
| | ***lpar2b*** | -1.264 | 0.744 | 8.96E-02 | -2.723 | 0.195 |
| | ***gatad2ab*** | -0.486 | 0.269 | 7.11E-02 | -1.014 | 0.042 |
| | **age (11dpf vs. 10dpf)** | -0.122 | 0.122 | 3.17E-01 | -0.362 | 0.117 |
| | **time of day (in hours since 9AM)** | 0.020 | 0.024 | 4.03E-01 | -0.027 | 0.067 |
| | **intercept** | -0.059 | 0.144 | 6.81E-01 | -0.341 | 0.223 |
| **random factors** | ***variance by batch*** | 0.972 | 0.031 | - | 0.914 | 1.034 |
| | ***residual*** | 0.218 | 0.084 | - | 0.103 | 0.462 |

| | | Triglyceride levels (n=513) | | | | |
|---|---|---|---|---|---|---|
| | | **Effect** | **SE** | ***P*** | **lci** | **uci** |
| **fixed factors** | ***lpar2a*** | -0.652 | 0.278 | 1.92E-02 | -1.197 | -0.106 |
| | ***lpar2b*** | -0.001 | 0.650 | 9.99E-01 | -1.276 | 1.274 |
| | ***gatad2ab*** | 0.394 | 0.236 | 9.45E-02 | -0.068 | 0.857 |
| | **age (11dpf vs. 10dpf)** | 0.007 | 0.111 | 9.52E-01 | -0.210 | 0.224 |
| | **time of day (in hours since 9AM)** | 0.085 | 0.022 | 7.12E-05 | 0.043 | 0.128 |
| | **intercept** | -0.328 | 0.232 | 1.57E-01 | -0.783 | 0.126 |
| **random factors** | ***variance by batch*** | *0.849* | *0.027* | - | *0.798* | *0.903* |
| | ***residual*** | *0.512* | *0.156* | - | *0.282* | *0.929* |

| | | Total cholesterol levels (n=513) | | | | |
|---|---|---|---|---|---|---|
| | | **Effect** | **SE** | ***P*** | **lci** | **uci** |
| **fixed factors** | ***lpar2a*** | -0.739 | 0.306 | 1.58E-02 | -1.340 | -0.139 |
| | ***lpar2b*** | 0.369 | 0.716 | 6.07E-01 | -1.035 | 1.773 |
| | ***gatad2ab*** | 0.118 | 0.260 | 6.48E-01 | -0.390 | 0.627 |
| | **age (11dpf vs. 10dpf)** | -0.487 | 0.120 | 4.89E-05 | -0.722 | -0.252 |
| | **time of day (in hours since 9AM)** | 0.018 | 0.023 | 4.39E-01 | -0.028 | 0.064 |
| | **intercept** | 0.034 | 0.166 | 8.37E-01 | -0.291 | 0.360 |
| **random factors** | ***variance by batch*** | *0.935* | *0.029* | - | *0.879* | *0.994* |
| | ***residual*** | *0.306* | *0.105* | - | *0.156* | *0.598* |

***continued* Supplementary Table 36**

| | | Glucose levels (n=513) | | | | |
|---|---|---|---|---|---|---|
| | | **Effect** | **SE** | ***P*** | **lci** | **uci** |
| **fixed factors** | ***lpar2a*** | -0.146 | 0.308 | 6.36E-01 | -0.748 | 0.457 |
| | ***lpar2b*** | -0.001 | 0.719 | 9.99E-01 | -1.410 | 1.408 |
| | ***gatad2ab*** | -0.371 | 0.260 | 1.55E-01 | -0.881 | 0.140 |
| | **age (11dpf vs. 10dpf)** | 0.147 | 0.120 | 2.23E-01 | -0.089 | 0.383 |
| | **time of day (in hours since 9AM)** | -0.080 | 0.024 | 6.71E-04 | -0.126 | -0.034 |
| | **intercept** | 0.240 | 0.168 | 1.53E-01 | -0.089 | 0.569 |
| **random factors** | ***variance by batch*** | *0.938* | *0.029* | - | *0.882* | *0.998* |
| | ***residual*** | *0.311* | *0.103* | - | *0.162* | *0.594* |

All outcomes were normalized for protein level using residuals, and inverse-normally transformed before the analysis. Associations were examined using hierarchical linear models. Effects shown are for the effect of carrying a mutated allele in *lpar2a* , *lpar2b* and *gatad2ab* , weighted by the allele's predicted effect on protein function (i.e. additive model, mutually adjusted). Associations were adjusted for age (i.e. 11 vs. 10 days post fertilization (dpf), time of day and batch. Lci and uci are lower and upper boundaries of the 95% confidence interval.

**Supplementary Table 37 - The effect of a mutated allele in *lpar2a, lpar2b* and *gatad2ab* on vascular atherogenic traits**

| | | Vascular lipid deposition | | | | | | | | | | | | | | |
|---|---|---|---|---|---|---|---|---|---|---|---|---|---|---|---|---|
| | | Model 1 (n=280) | | | | | | Model 2 (n=258) | | | | | Model 3 (n=233) | | | |
| | | Effect | SE | *P* | lci | uci | | Effect | SE | *P* | lci | uci | Effect | SE | *P* | lci | uci |
| negative binomial terms | ***lpar2a*** | 1.870 | 1.003 | 6.23E-02 | -0.096 | 3.836 | | 1.622 | 1.001 | 1.05E-01 | -0.340 | 3.584 | 1.795 | 0.995 | 7.13E-02 | -0.156 | 3.746 |
| | ***lpar2b*** | 0.994 | 1.191 | 4.04E-01 | -1.340 | 3.328 | | 0.861 | 1.139 | 4.50E-01 | -1.371 | 3.093 | 0.912 | 1.351 | 5.00E-01 | -1.736 | 3.560 |
| | ***gatad2ab*** | -0.006 | 0.334 | 9.87E-01 | -0.659 | 0.648 | | -0.058 | 0.316 | 8.55E-01 | -0.676 | 0.561 | -0.277 | 0.313 | 3.75E-01 | -0.890 | 0.336 |
| | **body length (in SD)** | - | - | - | - | - | | 0.152 | 0.116 | 1.92E-01 | -0.076 | 0.379 | 0.194 | 0.126 | 1.24E-01 | -0.054 | 0.442 |
| | **dorsal body surface area (in SD)** | - | - | - | - | - | | 0.153 | 0.131 | 2.44E-01 | -0.104 | 0.410 | 0.202 | 0.139 | 1.46E-01 | -0.070 | 0.474 |
| | **age (11dpf vs. 10dpf)** | 0.263 | 0.503 | 6.01E-01 | -0.722 | 1.249 | | 0.328 | 0.497 | 5.09E-01 | -0.646 | 1.302 | 0.229 | 0.462 | 6.20E-01 | -0.677 | 1.136 |
| | **time of day (in hours since 9AM)** | -0.079 | 0.038 | 4.00E-02 | -0.154 | -0.004 | | -0.075 | 0.041 | 7.06E-02 | -0.156 | 0.006 | -0.045 | 0.052 | 3.93E-01 | -0.147 | 0.058 |
| | **batch 1** | 3.239 | 0.635 | 3.42E-07 | 1.994 | 4.484 | | 3.299 | 0.621 | 1.09E-07 | 2.082 | 4.517 | 3.364 | 0.580 | 6.46E-09 | 2.228 | 4.500 |
| | **batch 2** | 4.939 | 0.746 | 3.50E-11 | 3.477 | 6.400 | | 4.771 | 0.782 | 1.03E-09 | 3.239 | 6.304 | 4.686 | 0.744 | 3.00E-10 | 3.228 | 6.144 |
| | **batch 3** | 4.581 | 0.725 | 2.66E-10 | 3.160 | 6.003 | | 4.361 | 0.749 | 5.83E-09 | 2.893 | 5.829 | 4.486 | 0.683 | 4.99E-11 | 3.148 | 5.824 |
| | **batch 4** | 3.656 | 0.738 | 7.39E-07 | 2.209 | 5.103 | | 3.855 | 0.726 | 1.11E-07 | 2.431 | 5.279 | 3.953 | 0.653 | 1.46E-09 | 2.672 | 5.234 |
| | **batch 5** | 3.903 | 0.732 | 9.59E-08 | 2.469 | 5.337 | | 3.705 | 0.751 | 8.02E-07 | 2.233 | 5.176 | 3.635 | 0.689 | 1.31E-07 | 2.285 | 4.985 |
| | **intercept** | 0.802 | 0.720 | 2.65E-01 | -0.609 | 2.212 | | 0.766 | 0.702 | 2.75E-01 | -0.610 | 2.143 | 0.597 | 0.639 | 3.50E-01 | -0.656 | 1.850 |
| | **HDL cholesterol levels (in SD)** | - | - | - | - | - | | - | - | - | - | - | -0.189 | 0.096 | 4.94E-02 | -0.377 | 0.000 |
| | **LDL cholesterol levels (in SD)** | - | - | - | - | - | | - | - | - | - | - | 0.084 | 0.075 | 2.65E-01 | -0.063 | 0.231 |
| | **glucose levels (in SD)** | - | - | - | - | - | | - | - | - | - | - | 0.070 | 0.105 | 5.09E-01 | -0.137 | 0.276 |
| | **triglyceride levels (in SD)** | - | - | - | - | - | | - | - | - | - | - | -0.016 | 0.108 | 8.82E-01 | -0.227 | 0.195 |

| | | Vascular infiltration by macrophages | | | | | | | | | | | | | | |
|---|---|---|---|---|---|---|---|---|---|---|---|---|---|---|---|---|
| | | Model 1 (n=363) | | | | | | Model 2 (n=331) | | | | | Model 3 (n=305) | | | |
| | | Effect | SE | *P* | lci | uci | | Effect | SE | *P* | lci | uci | Effect | SE | *P* | lci | uci |
| fixed factors | ***lpar2a*** | 0.307 | 0.374 | 4.12E-01 | -0.426 | 1.041 | 363 | 0.282 | 0.368 | 4.44E-01 | -0.440 | 1.003 | 0.238 | 0.420 | 5.70E-01 | -0.584 | 1.061 |
| | ***lpar2b*** | 0.528 | 0.520 | 3.09E-01 | -0.490 | 1.546 | | 0.597 | 0.512 | 2.43E-01 | -0.406 | 1.601 | 0.649 | 0.636 | 3.08E-01 | -0.598 | 1.896 |
| | ***gatad2ab*** | 0.404 | 0.254 | 1.11E-01 | -0.093 | 0.902 | | 0.472 | 0.261 | 7.08E-02 | -0.040 | 0.984 | 0.458 | 0.267 | 8.68E-02 | -0.066 | 0.982 |
| | **body length (in SD)** | - | - | - | - | - | | 0.020 | 0.059 | 7.38E-01 | -0.096 | 0.136 | 0.029 | 0.062 | 6.36E-01 | -0.092 | 0.150 |
| | **dorsal body surface area (in SD)** | - | - | - | - | - | | 0.077 | 0.055 | 1.63E-01 | -0.031 | 0.186 | 0.056 | 0.057 | 3.28E-01 | -0.056 | 0.168 |
| | **age (11dpf vs. 10dpf)** | -0.149 | 0.151 | 3.24E-01 | -0.445 | 0.147 | | -0.146 | 0.148 | 3.25E-01 | -0.436 | 0.144 | -0.124 | 0.147 | 3.97E-01 | -0.411 | 0.163 |
| | **time of day (in hours since 9AM)** | 0.042 | 0.023 | 6.53E-02 | -0.003 | 0.086 | | 0.043 | 0.023 | 6.58E-02 | -0.003 | 0.088 | 0.010 | 0.027 | 7.21E-01 | -0.043 | 0.063 |
| | **intercept** | -0.101 | 0.172 | 5.56E-01 | -0.438 | 0.235 | | -0.139 | 0.169 | 4.12E-01 | -0.470 | 0.193 | -0.029 | 0.157 | 8.55E-01 | -0.336 | 0.278 |
| | **HDL cholesterol levels (in SD)** | - | - | - | - | - | | - | - | - | - | - | -0.001 | 0.048 | 9.81E-01 | -0.095 | 0.092 |
| | **LDL cholesterol levels (in SD)** | - | - | - | - | - | | - | - | - | - | - | -0.012 | 0.046 | 7.88E-01 | -0.103 | 0.078 |
| | **glucose levels (in SD)** | - | - | - | - | - | | - | - | - | - | - | -0.021 | 0.052 | 6.84E-01 | -0.123 | 0.080 |
| | **triglyceride levels (in SD)** | - | - | - | - | - | | - | - | - | - | - | 0.094 | 0.057 | 9.68E-02 | -0.017 | 0.205 |
| random factors | ***intercept*** | *0.822* | *0.031* | - | *0.764* | *0.885* | | *0.805* | *0.032* | | *0.746* | *0.870* | *0.812* | *0.033* | | *0.749* | *0.879* |
| | ***variation by batch*** | *0.311* | *0.103* | - | *0.162* | *0.595* | | *0.294* | *0.099* | | *0.151* | *0.569* | *0.218* | *0.086* | | *0.100* | *0.474* |

***continued*** **Supplementary Table 37**

| | | Vascular co-localization of lipids with macrophages | | | | | | | | | | | | | | |
|---|---|---|---|---|---|---|---|---|---|---|---|---|---|---|---|---|
| | | **Model 1 (n=263)** | | | | | **Model 2 (n=241)** | | | | | **Model 3 (n=217)** | | | | |
| | | **Effect** | **SE** | ***P*** | **lci** | **uci** | **Effect** | **SE** | ***P*** | **lci** | **uci** | **Effect** | **SE** | ***P*** | **lci** | **uci** |
| **negative binomial terms** | ***lpar2a*** | 3.463 | 1.204 | 4.02E-03 | 1.103 | 5.823 | 3.511 | 1.207 | 3.63E-03 | 1.145 | 5.877 | 3.408 | 1.167 | 3.50E-03 | 1.120 | 5.696 |
| | ***lpar2b*** | 0.851 | 1.083 | 4.32E-01 | -1.270 | 2.973 | 0.747 | 1.018 | 4.63E-01 | -1.248 | 2.743 | 0.284 | 1.368 | 8.35E-01 | -2.396 | 2.965 |
| | ***gatad2ab*** | 0.248 | 0.491 | 6.13E-01 | -0.714 | 1.210 | 0.495 | 0.518 | 3.39E-01 | -0.520 | 1.509 | 0.404 | 0.582 | 4.87E-01 | -0.737 | 1.545 |
| | **body length (in SD)** | - | - | - | - | - | 0.101 | 0.190 | 5.94E-01 | -0.271 | 0.473 | 0.223 | 0.197 | 2.57E-01 | -0.162 | 0.609 |
| | **dorsal body surface area (in SD)** | - | - | - | - | - | -0.063 | 0.179 | 7.25E-01 | -0.414 | 0.288 | 0.059 | 0.200 | 7.70E-01 | -0.334 | 0.452 |
| | **age (11dpf vs. 10dpf)** | 0.248 | 0.642 | 6.99E-01 | -1.010 | 1.506 | 0.261 | 0.634 | 6.81E-01 | -0.981 | 1.502 | -0.196 | 0.678 | 7.73E-01 | -1.524 | 1.133 |
| | **time of day (in hours since 9AM)** | -0.129 | 0.054 | 1.65E-02 | -0.235 | -0.024 | -0.150 | 0.060 | 1.17E-02 | -0.267 | -0.033 | -0.124 | 0.087 | 1.55E-01 | -0.294 | 0.047 |
| | **batch 1** | 4.433 | 0.824 | 7.38E-08 | 2.819 | 6.048 | 4.418 | 0.831 | 1.06E-07 | 2.789 | 6.048 | 4.521 | 0.825 | 4.31E-08 | 2.904 | 6.139 |
| | **batch 2** | 5.985 | 0.974 | 7.89E-10 | 4.076 | 7.893 | 5.875 | 0.994 | 3.37E-09 | 3.928 | 7.823 | 5.222 | 1.122 | 3.25E-06 | 3.023 | 7.422 |
| | **batch 3** | 5.635 | 0.928 | 1.28E-09 | 3.815 | 7.454 | 5.649 | 0.948 | 2.51E-09 | 3.792 | 7.507 | 5.258 | 1.068 | 8.46E-07 | 3.165 | 7.351 |
| | **batch 4** | 4.252 | 1.026 | 3.40E-05 | 2.241 | 6.262 | 4.352 | 1.082 | 5.73E-05 | 2.232 | 6.472 | 4.120 | 1.109 | 2.02E-04 | 1.947 | 6.293 |
| | **batch 5** | 4.694 | 0.939 | 5.84E-07 | 2.853 | 6.535 | 4.494 | 0.969 | 3.55E-06 | 2.594 | 6.394 | 4.039 | 1.033 | 9.29E-05 | 2.014 | 6.065 |
| | **intercept** | -1.384 | 0.956 | 1.48E-01 | -3.258 | 0.490 | -1.303 | 0.965 | 1.77E-01 | -3.195 | 0.589 | -1.141 | 1.126 | 3.11E-01 | -3.348 | 1.066 |
| | **HDL cholesterol levels (in SD)** | - | - | - | - | - | - | - | - | - | - | -0.098 | 0.139 | 4.82E-01 | -0.371 | 0.175 |
| | **LDL cholesterol levels (in SD)** | - | - | - | - | - | - | - | - | - | - | 0.105 | 0.107 | 3.23E-01 | -0.104 | 0.314 |
| | **glucose levels (in SD)** | - | - | - | - | - | - | - | - | - | - | -0.274 | 0.167 | 1.01E-01 | -0.601 | 0.053 |
| | **triglyceride levels (in SD)** | - | - | - | - | - | - | - | - | - | - | -0.170 | 0.139 | 2.22E-01 | -0.442 | 0.103 |

| | | Vascular infiltration by neutrophils | | | | | | | | | | | | | | |
|---|---|---|---|---|---|---|---|---|---|---|---|---|---|---|---|---|
| | | **Model 1 (n=363)** | | | | | **Model 2 (n=334)** | | | | | **Model 3 (n=307)** | | | | |
| | | **Effect** | **SE** | ***P*** | **lci** | **uci** | **Effect** | **SE** | ***P*** | **lci** | **uci** | **Effect** | **SE** | ***P*** | **lci** | **uci** |
| **fixed factors** | ***lpar2a*** | 0.743 | 0.387 | 5.46E-02 | -0.015 | 1.501 | 0.678 | 0.386 | 7.89E-02 | -0.078 | 1.435 | 0.566 | 0.423 | 1.80E-01 | -0.262 | 1.395 |
| | ***lpar2b*** | 0.130 | 0.537 | 8.09E-01 | -0.922 | 1.182 | 0.035 | 0.538 | 9.48E-01 | -1.019 | 1.090 | 0.045 | 0.642 | 9.44E-01 | -1.213 | 1.303 |
| | ***gatad2ab*** | -0.315 | 0.267 | 2.38E-01 | -0.837 | 0.208 | -0.452 | 0.278 | 1.04E-01 | -0.996 | 0.093 | -0.353 | 0.274 | 1.97E-01 | -0.889 | 0.183 |
| | **body length (in SD)** | - | - | - | - | - | 0.145 | 0.061 | 1.76E-02 | 0.025 | 0.264 | 0.197 | 0.063 | 1.72E-03 | 0.074 | 0.320 |
| | **dorsal body surface area (in SD)** | - | - | - | - | - | 0.053 | 0.059 | 3.68E-01 | -0.062 | 0.168 | 0.016 | 0.060 | 7.88E-01 | -0.101 | 0.134 |
| | **age (11dpf vs. 10dpf)** | -0.120 | 0.152 | 4.30E-01 | -0.418 | 0.178 | -0.157 | 0.150 | 2.97E-01 | -0.451 | 0.138 | -0.159 | 0.150 | 2.89E-01 | -0.453 | 0.135 |
| | **time of day (in hours since 9AM)** | 0.025 | 0.023 | 2.83E-01 | -0.021 | 0.070 | 0.025 | 0.024 | 2.92E-01 | -0.022 | 0.072 | 0.019 | 0.027 | 4.92E-01 | -0.034 | 0.072 |
| | **intercept** | 0.033 | 0.195 | 8.65E-01 | -0.348 | 0.415 | 0.023 | 0.179 | 9.00E-01 | -0.329 | 0.374 | 0.038 | 0.186 | 8.36E-01 | -0.325 | 0.402 |
| | **HDL cholesterol levels (in SD)** | - | - | - | - | - | - | - | - | - | - | 0.029 | 0.048 | 5.54E-01 | -0.066 | 0.123 |
| | **LDL cholesterol levels (in SD)** | - | - | - | - | - | - | - | - | - | - | -0.135 | 0.046 | 3.67E-03 | -0.226 | -0.044 |
| | **glucose levels (in SD)** | - | - | - | - | - | - | - | - | - | - | 0.033 | 0.052 | 5.35E-01 | -0.070 | 0.135 |
| | **triglyceride levels (in SD)** | - | - | - | - | - | - | - | - | - | - | -0.049 | 0.057 | 3.88E-01 | -0.160 | 0.062 |
| **random factors** | ***intercept*** | *0.849* | *0.032* | - | *0.789* | *0.914* | *0.846* | *0.033* | | *0.783* | *0.913* | *0.818* | *0.033* | | *0.755* | *0.886* |
| | ***variation by batch*** | *0.379* | *0.121* | - | *0.202* | *0.710* | *0.320* | *0.111* | | *0.162* | *0.630* | *0.329* | *0.114* | | *0.167* | *0.649* |

*continued* **Supplementary Table 37**

| | | **Vascular co-localization of lipids with neutrophils** | | | | | | | | | | | | | | |
|---|---|---|---|---|---|---|---|---|---|---|---|---|---|---|---|---|
| | | **Model 1 (n=260)** | | | | | **Model 2 (n=242)** | | | | | **Model 3 (n=217)** | | | | |
| | | **Effect** | **SE** | ***P*** | **lci** | **uci** | **Effect** | **SE** | ***P*** | **lci** | **uci** | **Effect** | **SE** | ***P*** | **lci** | **uci** |
| **negative binomial terms** | ***lpar2a*** | 2.718 | 1.321 | 3.96E-02 | 0.129 | 5.307 | 2.614 | 1.314 | 4.67E-02 | 0.038 | 5.189 | 3.548 | 1.292 | 6.03E-03 | 1.016 | 6.081 |
| | ***lpar2b*** | -0.034 | 1.109 | 9.75E-01 | -2.207 | 2.139 | -0.260 | 1.167 | 8.24E-01 | -2.547 | 2.027 | -2.627 | 1.687 | 1.20E-01 | -5.934 | 0.681 |
| | ***gatad2ab*** | -1.383 | 1.031 | 1.80E-01 | -3.403 | 0.638 | -1.613 | 1.014 | 1.12E-01 | -3.601 | 0.375 | -2.236 | 1.025 | 2.91E-02 | -4.245 | -0.227 |
| | **body length (in SD)** | - | - | - | - | - | 0.260 | 0.182 | 1.53E-01 | -0.097 | 0.617 | 0.178 | 0.185 | 3.37E-01 | -0.185 | 0.542 |
| | **dorsal body surface area (in SD)** | - | - | - | - | - | 0.077 | 0.182 | 6.72E-01 | -0.280 | 0.434 | 0.000 | 0.187 | 9.98E-01 | -0.367 | 0.368 |
| | **age (11dpf vs. 10dpf)** | 0.838 | 0.610 | 1.69E-01 | -0.357 | 2.033 | 0.853 | 0.608 | 1.60E-01 | -0.338 | 2.044 | 0.390 | 0.632 | 5.38E-01 | -0.849 | 1.628 |
| | **time of day (in hours since 9AM)** | -0.126 | 0.069 | 6.76E-02 | -0.262 | 0.009 | -0.134 | 0.068 | 4.74E-02 | -0.267 | -0.002 | -0.219 | 0.078 | 5.16E-03 | -0.372 | -0.065 |
| | **batch 1** | 3.057 | 1.022 | 2.78E-03 | 1.054 | 5.060 | 3.136 | 0.981 | 1.39E-03 | 1.213 | 5.059 | 3.287 | 0.840 | 9.06E-05 | 1.641 | 4.933 |
| | **batch 2** | 4.126 | 1.202 | 5.98E-04 | 1.770 | 6.482 | 3.865 | 1.213 | 1.44E-03 | 1.488 | 6.243 | 4.036 | 1.086 | 2.01E-04 | 1.908 | 6.164 |
| | **batch 3** | 3.450 | 1.162 | 2.99E-03 | 1.173 | 5.727 | 3.265 | 1.137 | 4.09E-03 | 1.036 | 5.494 | 3.350 | 1.010 | 9.10E-04 | 1.371 | 5.330 |
| | **batch 4** | 3.911 | 1.181 | 9.27E-04 | 1.596 | 6.225 | 4.210 | 1.145 | 2.36E-04 | 1.966 | 6.453 | 4.467 | 0.971 | 4.21E-06 | 2.564 | 6.370 |
| | **batch 5** | 2.912 | 1.171 | 1.29E-02 | 0.617 | 5.208 | 3.073 | 1.122 | 6.16E-03 | 0.874 | 5.272 | 3.193 | 0.965 | 9.33E-04 | 1.303 | 5.084 |
| | **intercept** | -1.356 | 1.157 | 2.41E-01 | -3.624 | 0.911 | -1.388 | 1.105 | 2.09E-01 | -3.554 | 0.778 | -1.071 | 0.999 | 2.83E-01 | -3.029 | 0.886 |
| | **HDL cholesterol levels (in SD)** | - | - | - | - | - | - | - | - | - | - | -0.312 | 0.165 | 5.79E-02 | -0.635 | 0.010 |
| | **LDL cholesterol levels (in SD)** | - | - | - | - | - | - | - | - | - | - | 0.088 | 0.132 | 5.05E-01 | -0.171 | 0.347 |
| | **glucose levels (in SD)** | - | - | - | - | - | - | - | - | - | - | 0.134 | 0.145 | 3.53E-01 | -0.149 | 0.417 |
| | **triglyceride levels (in SD)** | - | - | - | - | - | - | - | - | - | - | 0.135 | 0.171 | 4.30E-01 | -0.200 | 0.471 |

| | | **Vascular co-localization of macrophages with neutrophils** | | | | | | | | | | | | | | |
|---|---|---|---|---|---|---|---|---|---|---|---|---|---|---|---|---|
| | | **Model 1 (n=345)** | | | | | **Model 2 (n=317)** | | | | | **Model 3 (n=291)** | | | | |
| | | **Effect** | **SE** | ***P*** | **lci** | **uci** | **Effect** | **SE** | ***P*** | **lci** | **uci** | **Effect** | **SE** | ***P*** | **lci** | **uci** |
| **negative binomial terms** | ***lpar2a*** | 0.219 | 0.301 | 4.67E-01 | -0.371 | 0.808 | 0.185 | 0.304 | 5.44E-01 | -0.411 | 0.781 | 0.023 | 0.315 | 9.41E-01 | -0.594 | 0.641 |
| | ***lpar2b*** | -0.073 | 0.473 | 8.77E-01 | -1.000 | 0.853 | 0.022 | 0.452 | 9.62E-01 | -0.864 | 0.907 | 0.225 | 0.513 | 6.61E-01 | -0.780 | 1.229 |
| | ***gatad2ab*** | -0.409 | 0.357 | 2.52E-01 | -1.108 | 0.290 | -0.514 | 0.417 | 2.18E-01 | -1.330 | 0.303 | -0.676 | 0.384 | 7.84E-02 | -1.429 | 0.077 |
| | **body length (in SD)** | - | - | - | - | - | -0.084 | 0.073 | 2.49E-01 | -0.226 | 0.059 | -0.027 | 0.077 | 7.29E-01 | -0.177 | 0.124 |
| | **dorsal body surface area (in SD)** | - | - | - | - | - | 0.100 | 0.066 | 1.26E-01 | -0.028 | 0.229 | 0.096 | 0.068 | 1.60E-01 | -0.038 | 0.230 |
| | **age (11dpf vs. 10dpf)** | -0.209 | 0.165 | 2.05E-01 | -0.532 | 0.114 | -0.186 | 0.165 | 2.59E-01 | -0.510 | 0.137 | -0.199 | 0.174 | 2.51E-01 | -0.540 | 0.141 |
| | **time of day (in hours since 9AM)** | 0.019 | 0.028 | 4.97E-01 | -0.036 | 0.075 | 0.021 | 0.029 | 4.65E-01 | -0.035 | 0.077 | 0.030 | 0.032 | 3.60E-01 | -0.034 | 0.093 |
| | **batch 1** | 0.026 | 0.176 | 8.83E-01 | -0.318 | 0.370 | 0.084 | 0.178 | 6.39E-01 | -0.266 | 0.433 | 0.038 | 0.205 | 8.54E-01 | -0.365 | 0.440 |
| | **batch 2** | 0.491 | 0.255 | 5.39E-02 | -0.008 | 0.991 | 0.528 | 0.311 | 8.94E-02 | -0.081 | 1.137 | 0.512 | 0.349 | 1.43E-01 | -0.173 | 1.196 |
| | **batch 3** | 1.144 | 0.236 | 1.26E-06 | 0.681 | 1.606 | 1.137 | 0.267 | 2.00E-05 | 0.615 | 1.660 | 1.223 | 0.314 | 9.78E-05 | 0.608 | 1.838 |
| | **batch 4** | -0.467 | 0.259 | 7.09E-02 | -0.975 | 0.040 | -0.514 | 0.258 | 4.65E-02 | -1.021 | -0.008 | -0.560 | 0.255 | 2.78E-02 | -1.059 | -0.061 |
| | **batch 5** | -0.311 | 0.228 | 1.73E-01 | -0.758 | 0.136 | -0.313 | 0.238 | 1.88E-01 | -0.779 | 0.153 | -0.299 | 0.264 | 2.58E-01 | -0.816 | 0.219 |
| | **intercept** | 4.047 | 0.208 | 1.95E-84 | 3.639 | 4.454 | 4.021 | 0.211 | 8.39E-81 | 3.607 | 4.435 | 3.978 | 0.253 | 7.40E-56 | 3.483 | 4.473 |
| | **HDL cholesterol levels (in SD)** | - | - | - | - | - | - | - | - | - | - | -0.035 | 0.054 | 5.11E-01 | -0.141 | 0.070 |
| | **LDL cholesterol levels (in SD)** | - | - | - | - | - | - | - | - | - | - | -0.105 | 0.060 | 7.96E-02 | -0.222 | 0.012 |
| | **glucose levels (in SD)** | - | - | - | - | - | - | - | - | - | - | -0.005 | 0.062 | 9.36E-01 | -0.126 | 0.116 |
| | **triglyceride levels (in SD)** | - | - | - | - | - | - | - | - | - | - | -0.024 | 0.072 | 7.35E-01 | -0.166 | 0.117 |

Associations were examined using negative binomial regression for outcomes that showed a negative binomial distribution; and using hierarchical linear models on inverse-normally transformed outcomes for outcomes that were (borderline) normally distributed (i.e. vascular accumulation of macrophages and neutrophils). Effects shown are for the effect of carrying a mutated allele in *lpar2a* , *lpar2b* and *gatad2ab* , weighted by the allele's predicted effect on protein function (i.e. additive model, mutually adjusted). Model 1: adjusted for time of day and batch; Model 2: additionally adjusted for body length and dorsal body surface area normalized for length; Model 3: additionally adjusted for whole-body LDL cholesterol, HDL cholesterol, triglyceride and glucose levels normalized for protein level. Lci and uci are lower and upper boundaries of the 95% confidence interval.

**Supplementary Table 38 - The effect of a mutated allele in *lpar2a*, *lpar2b* and *gatad2ab* on image and image quantification quality**

| | Region of interest not detected | | | | | | | | | | | | | | |
|---|---|---|---|---|---|---|---|---|---|---|---|---|---|---|---|
| | Model 1 (n=297) | | | | | Model 2 (n=276) | | | | | Model 3 (n=253) | | | | |
| | Effect | SE | *P* | lci | uci | Effect | SE | *P* | lci | uci | Effect | SE | *P* | lci | uci |
| ***lpar2a*** | 1.021 | 1.184 | 9.85E-01 | 0.105 | 9.899 | 3.326 | 4.200 | 3.41E-01 | 0.280 | 39.519 | 2.918 | 3.880 | 4.21E-01 | 0.215 | 39.528 |
| ***lpar2b*** | all 39 larvae with an undetected vessel are wildtype for mutations in *lpar2b* ; 0.74 mutant larvae were expected. | | | | | | | | | | | | | | |
| ***gatad2ab*** | all 39 larvae with an undetected vessel are wildtype for mutations in *gatad2ab* ; 1.84 mutant larvae were expected. | | | | | | | | | | | | | | |
| **age (11dpf vs. 10dpf)** | 3.904 | 1.404 | 1.52E-04 | 1.929 | 7.898 | 2.172 | 0.848 | 4.71E-02 | 1.010 | 4.670 | 2.011 | 0.802 | 7.99E-02 | 0.920 | 4.395 |
| **time of day (in hours since 9AM)** | 0.923 | 0.080 | 3.56E-01 | 0.780 | 1.093 | 0.999 | 0.099 | 9.92E-01 | 0.823 | 1.212 | 1.060 | 0.109 | 5.68E-01 | 0.867 | 1.297 |
| **body length (in SD)** | - | - | - | - | - | 0.334 | 0.083 | 9.35E-06 | 0.206 | 0.543 | 0.344 | 0.091 | 5.17E-05 | 0.205 | 0.576 |
| **dorsal body surface area (in SD)** | - | - | - | - | - | 0.345 | 0.089 | 3.82E-05 | 0.208 | 0.573 | 0.356 | 0.097 | 1.59E-04 | 0.208 | 0.609 |
| **LDLc levels (in SD)** | - | - | - | - | - | - | - | - | - | - | 1.076 | 0.262 | 7.63E-01 | 0.668 | 1.734 |
| **HDLc levels (in SD)** | - | - | - | - | - | - | - | - | - | - | 1.161 | 0.247 | 4.81E-01 | 0.766 | 1.761 |
| **triglyceride levels (in SD)** | - | - | - | - | - | - | - | - | - | - | 0.634 | 0.157 | 6.58E-02 | 0.390 | 1.030 |
| **glucose levels** | - | - | - | - | - | - | - | - | - | - | 1.054 | 0.239 | 8.16E-01 | 0.676 | 1.643 |
| **intercept** | 0.139 | 0.063 | 1.19E-05 | 0.057 | 0.336 | 0.096 | 0.050 | 6.94E-06 | 0.035 | 0.267 | 0.069 | 0.038 | 1.68E-06 | 0.023 | 0.205 |

| | Many false positive lipid deposits | | | | | | | | | | | | | | |
|---|---|---|---|---|---|---|---|---|---|---|---|---|---|---|---|
| | Model 1 (n=364) | | | | | Model 2 (n=331) | | | | | Model 3 (n=305) | | | | |
| | Effect | SE | *P* | lci | uci | Effect | SE | *P* | lci | uci | Effect | SE | *P* | lci | uci |
| ***lpar2a*** | all 90 larvae with many false positive lipid deposits were wildtype for mutations in *lpar2a* ; 1.22 mutant larvae were expected. | | | | | | | | | | | | | | |
| ***lpar2b*** | all 90 larvae with many false positive lipid deposits were wildtype or had a missed call for mutations in *lpar2b* ; 1.46 mutant larvae were expected. | | | | | | | | | | | | | | |
| ***gatad2ab*** | 3.137 | 1.951 | 6.59E-02 | 0.927 | 10.613 | 4.301 | 2.900 | 3.05E-02 | 1.147 | 16.125 | 3.799 | 2.631 | 5.40E-02 | 0.977 | 14.766 |
| **age (11dpf vs. 10dpf)** | 0.772 | 0.265 | 4.50E-01 | 0.394 | 1.512 | 0.639 | 0.228 | 2.09E-01 | 0.317 | 1.286 | 0.642 | 0.232 | 2.20E-01 | 0.316 | 1.304 |
| **time of day (in hours since 9AM)** | 0.968 | 0.059 | 5.92E-01 | 0.859 | 1.090 | 0.981 | 0.066 | 7.76E-01 | 0.859 | 1.120 | 0.985 | 0.074 | 8.41E-01 | 0.850 | 1.141 |
| **body length (in SD)** | - | - | - | - | - | 0.753 | 0.113 | 5.92E-02 | 0.561 | 1.011 | 0.764 | 0.126 | 1.04E-01 | 0.552 | 1.057 |
| **dorsal body surface area (in SD)** | - | - | - | - | - | 0.697 | 0.099 | 1.13E-02 | 0.527 | 0.921 | 0.708 | 0.109 | 2.45E-02 | 0.523 | 0.957 |
| **LDLc levels (in SD)** | - | - | - | - | - | - | - | - | - | - | 0.867 | 0.116 | 2.86E-01 | 0.667 | 1.127 |
| **HDLc levels (in SD)** | - | - | - | - | - | - | - | - | - | - | 1.105 | 0.152 | 4.69E-01 | 0.843 | 1.448 |
| **triglyceride levels (in SD)** | - | - | - | - | - | - | - | - | - | - | 1.113 | 0.178 | 5.03E-01 | 0.813 | 1.524 |
| **glucose levels** | - | - | - | - | - | - | - | - | - | - | 0.843 | 0.122 | 2.39E-01 | 0.635 | 1.120 |
| **intercept** | 0.382 | 0.122 | 2.67E-03 | 0.204 | 0.716 | 0.386 | 0.134 | 5.99E-03 | 0.196 | 0.761 | 0.384 | 0.140 | 8.74E-03 | 0.188 | 0.785 |

*continued* **Supplementary Table 38**

| | Many false negative lipid deposits | | | | | | | | | | | | | | |
|---|---|---|---|---|---|---|---|---|---|---|---|---|---|---|---|
| | Model 1 (n=287) | | | | | Model 2 (n=265) | | | | | Model 3 (n=243) | | | | |
| | **Effect** | **SE** | ***P*** | **lci** | **uci** | **Effect** | **SE** | ***P*** | **lci** | **uci** | **Effect** | **SE** | ***P*** | **lci** | **uci** |
| ***lpar2a*** | all 19 larvae with many false negative lipid deposits were wildtype for mutations in *lpar2a* ; 0.32 mutant larvae were expected. | | | | | | | | | | | | | | |
| ***lpar2b*** | all 19 larvae with many false negative lipid deposits were wildtype for mutations in *lpar2b* ; 0.38 mutant larvae were expected. | | | | | | | | | | | | | | |
| ***gatad2ab*** | 0.934 | 1.481 | 9.66E-01 | 0.042 | 20.905 | 1.128 | 1.885 | 9.42E-01 | 0.043 | 29.838 | 2.649 | 4.643 | 5.78E-01 | 0.085 | 82.204 |
| **age (11dpf vs. 10dpf)** | 2.011 | 1.051 | 1.81E-01 | 0.722 | 5.601 | 1.209 | 0.651 | 7.25E-01 | 0.421 | 3.476 | 1.270 | 0.702 | 6.66E-01 | 0.430 | 3.752 |
| **time of day (in hours since 9AM)** | 0.872 | 0.103 | 2.47E-01 | 0.691 | 1.100 | 0.917 | 0.117 | 4.97E-01 | 0.715 | 1.177 | 1.000 | 0.133 | 1.00E+00 | 0.770 | 1.299 |
| **body length (in SD)** | - | - | - | - | - | 0.542 | 0.158 | 3.51E-02 | 0.307 | 0.958 | 0.564 | 0.179 | 7.11E-02 | 0.303 | 1.050 |
| **dorsal body surface area (in SD)** | - | - | - | - | - | 0.449 | 0.139 | 9.81E-03 | 0.244 | 0.824 | 0.479 | 0.157 | 2.49E-02 | 0.252 | 0.911 |
| **LDLc levels (in SD)** | - | - | - | - | - | - | - | - | - | - | 1.064 | 0.293 | 8.22E-01 | 0.620 | 1.826 |
| **HDLc levels (in SD)** | - | - | - | - | - | - | - | - | - | - | 1.437 | 0.375 | 1.65E-01 | 0.861 | 2.397 |
| **triglyceride levels (in SD)** | - | - | - | - | - | - | - | - | - | - | 0.749 | 0.231 | 3.49E-01 | 0.409 | 1.371 |
| **glucose levels** | - | - | - | - | - | - | - | - | - | - | 1.389 | 0.367 | 2.13E-01 | 0.828 | 2.332 |
| **intercept** | 0.110 | 0.064 | 1.38E-04 | 0.035 | 0.342 | 0.106 | 0.065 | 2.63E-04 | 0.032 | 0.355 | 0.063 | 0.044 | 5.84E-05 | 0.017 | 0.243 |

| | Transparant larvae | | | | | | | | | | | | | | |
|---|---|---|---|---|---|---|---|---|---|---|---|---|---|---|---|
| | Model 1 (n=280) | | | | | Model 2 (n=258) | | | | | Model 3 (n=236) | | | | |
| | **Effect** | **SE** | ***P*** | **lci** | **uci** | **Effect** | **SE** | ***P*** | **lci** | **uci** | **Effect** | **SE** | ***P*** | **lci** | **uci** |
| ***lpar2a*** | all 12 larvae that appeared particularly transparant were wildtype for mutations in *lpar2a* ; 0.21 mutant larvae were expected. | | | | | | | | | | | | | | |
| ***lpar2b*** | all 12 larvae that appeared particularly transparant were wildtype for mutations in *lpar2b* ; 0.25 mutant larvae were expected. | | | | | | | | | | | | | | |
| ***gatad2ab*** | 1.849 | 2.961 | 7.01E-01 | 0.080 | 42.635 | 1.376 | 2.245 | 8.45E-01 | 0.056 | 33.657 | 0.753 | 1.386 | 8.77E-01 | 0.020 | 27.745 |
| **age (11dpf vs. 10dpf)** | 2.166 | 1.382 | 2.26E-01 | 0.620 | 7.565 | 3.442 | 2.524 | 9.18E-02 | 0.818 | 14.488 | 2.309 | 1.942 | 3.20E-01 | 0.444 | 12.004 |
| **time of day (in hours since 9AM)** | 0.900 | 0.135 | 4.83E-01 | 0.671 | 1.207 | 0.871 | 0.131 | 3.60E-01 | 0.648 | 1.170 | 0.902 | 0.185 | 6.17E-01 | 0.603 | 1.350 |
| **body length (in SD)** | - | - | - | - | - | 1.344 | 0.452 | 3.80E-01 | 0.695 | 2.598 | 0.762 | 0.295 | 4.83E-01 | 0.357 | 1.626 |
| **dorsal body surface area (in SD)** | - | - | - | - | - | 1.923 | 0.681 | 6.49E-02 | 0.960 | 3.851 | 1.344 | 0.553 | 4.73E-01 | 0.599 | 3.012 |
| **LDLc levels (in SD)** | - | - | - | - | - | - | - | - | - | - | 1.160 | 0.335 | 6.07E-01 | 0.659 | 2.045 |
| **HDLc levels (in SD)** | - | - | - | - | - | - | - | - | - | - | 0.915 | 0.293 | 7.81E-01 | 0.489 | 1.714 |
| **triglyceride levels (in SD)** | - | - | - | - | - | - | - | - | - | - | 4.396 | 1.989 | 1.06E-03 | 1.811 | 10.669 |
| **glucose levels** | - | - | - | - | - | - | - | - | - | - | 0.756 | 0.268 | 4.30E-01 | 0.377 | 1.516 |
| **intercept** | 0.057 | 0.043 | 1.51E-04 | 0.013 | 0.251 | 0.043 | 0.035 | 1.12E-04 | 0.009 | 0.211 | 0.019 | 0.023 | 8.75E-04 | 0.002 | 0.197 |

***continued*** **Supplementary Table 38**

| | | | | | | | | | | | | | | | | |
|---|---|---|---|---|---|---|---|---|---|---|---|---|---|---|---|---|
| | **Bad quality images of macrophages** | | | | | | | | | | | | | | | |
| | **Model 1 (n=323)** | | | | | **Model 2 (n=291)** | | | | | | **Model 3 (n=269)** | | | | |
| | **Effect** | **SE** | ***P*** | **lci** | **uci** | **Effect** | **SE** | ***P*** | **lci** | **uci** | | **Effect** | **SE** | ***P*** | **lci** | **uci** |
| ***lpar2a*** | all 12 larvae that had bad quality images for macrophages were wildtype for mutations in *lpar2a* ; 0.27 mutant larvae were expected. | | | | | | | | | | | | | | | |
| ***lpar2b*** | all 12 larvae that had bad quality images for macrophages were wildtype for mutations in *lpar2b* ; 0.13 mutant larvae were expected. | | | | | | | | | | | | | | | |
| ***gatad2ab*** | all 12 larvae that had bad quality images for macrophages were wildtype for mutations in *gatad2ab* ; 0.83 mutant larvae were expected. | | | | | | | | | | | | | | | |
| **time of day (in hours since 9AM)** | 1.004 | 0.144 | 9.79E-01 | 0.758 | 1.329 | 1.097 | 0.201 | 6.12E-01 | 0.767 | 1.570 | 291 | 1.256 | 0.277 | 3.01E-01 | 0.815 | 1.935 |
| **body length (in SD)** | - | - | - | - | - | 0.301 | 0.110 | 1.00E-03 | 0.147 | 0.615 | 291 | 0.309 | 0.120 | 2.39E-03 | 0.145 | 0.660 |
| **dorsal body surface area (in SD)** | - | - | - | - | - | 1.353 | 0.508 | 4.20E-01 | 0.649 | 2.825 | 291 | 1.734 | 0.737 | 1.95E-01 | 0.754 | 3.988 |
| **LDLc levels (in SD)** | - | - | - | - | - | - | - | - | - | - | 291 | 0.942 | 0.366 | 8.77E-01 | 0.440 | 2.016 |
| **HDLc levels (in SD)** | - | - | - | - | - | - | - | - | - | - | 291 | 1.071 | 0.358 | 8.38E-01 | 0.556 | 2.063 |
| **triglyceride levels (in SD)** | - | - | - | - | - | - | - | - | - | - | 291 | 0.668 | 0.284 | 3.42E-01 | 0.290 | 1.536 |
| **glucose levels** | - | - | - | - | - | - | - | - | - | - | 291 | 0.709 | 0.250 | 3.30E-01 | 0.355 | 1.416 |
| **intercept** | 0.038 | 0.027 | 5.64E-06 | 0.009 | 0.156 | 0.015 | 0.015 | 1.26E-05 | 0.002 | 0.100 | 291 | 0.008 | 0.009 | 3.87E-05 | 0.001 | 0.078 |

| | | | | | | | | | | | | | | | |
|---|---|---|---|---|---|---|---|---|---|---|---|---|---|---|---|
| | **Many false positve macrophages** | | | | | | | | | | | | | | |
| | **Model 1 (n=401)** | | | | | **Model 2 (n=371)** | | | | | **Model 3 (n=348)** | | | | |
| | **Effect** | **SE** | ***P*** | **lci** | **uci** | **Effect** | **SE** | ***P*** | **lci** | **uci** | **Effect** | **SE** | ***P*** | **lci** | **uci** |
| ***lpar2a*** | all 10 larvae that had many false positive macrophages were wildtype for mutations in *lpar2a* ; 0.22 mutant larvae were expected. | | | | | | | | | | | | | | |
| ***lpar2b*** | all 10 larvae that had many false positive macrophages were wildtype for mutations in *lpar2b* ; 0.11 mutant larvae were expected. | | | | | | | | | | | | | | |
| ***gatad2ab*** | all 10 larvae that had many false positive macrophages were wildtype for mutations in *gatad2ab* ; 0.69 mutant larvae were expected. | | | | | | | | | | | | | | |
| **age (11dpf vs. 10dpf)** | 1.645 | 1.155 | 4.78E-01 | 0.416 | 6.513 | 1.454 | 1.060 | 6.07E-01 | 0.349 | 6.069 | 2.040 | 1.563 | 3.52E-01 | 0.455 | 9.154 |
| **time of day (in hours since 9AM)** | 0.907 | 0.143 | 5.35E-01 | 0.666 | 1.235 | 0.909 | 0.143 | 5.42E-01 | 0.668 | 1.236 | 0.733 | 0.149 | 1.27E-01 | 0.492 | 1.092 |
| **body length (in SD)** | - | - | - | - | - | 0.718 | 0.246 | 3.34E-01 | 0.367 | 1.406 | 0.646 | 0.255 | 2.68E-01 | 0.298 | 1.400 |
| **dorsal body surface area (in SD)** | - | - | - | - | - | 1.489 | 0.513 | 2.48E-01 | 0.758 | 2.924 | 1.005 | 0.378 | 9.90E-01 | 0.481 | 2.101 |
| **LDLc levels (in SD)** | - | - | - | - | - | - | - | - | - | - | 2.512 | 1.011 | 2.21E-02 | 1.141 | 5.528 |
| **HDLc levels (in SD)** | - | - | - | - | - | - | - | - | - | - | 0.728 | 0.297 | 4.35E-01 | 0.327 | 1.618 |
| **triglyceride levels (in SD)** | - | - | - | - | - | - | - | - | - | - | 1.196 | 0.509 | 6.73E-01 | 0.520 | 2.752 |
| **glucose levels** | - | - | - | - | - | - | - | - | - | - | 0.843 | 0.317 | 6.51E-01 | 0.404 | 1.762 |
| **intercept** | 0.035 | 0.026 | 9.99E-06 | 0.008 | 0.154 | 0.036 | 0.026 | 7.34E-06 | 0.008 | 0.153 | 0.050 | 0.042 | 3.21E-04 | 0.010 | 0.257 |

*continued* **Supplementary Table 38**

| | Clumped macrophages | | | | | | | | | | | | | | |
|---|---|---|---|---|---|---|---|---|---|---|---|---|---|---|---|
| | Model 1 (n=408) | | | | | Model 2 (n=378) | | | | | Model 3 (n=353) | | | | |
| | **Effect** | **SE** | ***P*** | **lci** | **uci** | **Effect** | **SE** | ***P*** | **lci** | **uci** | **Effect** | **SE** | ***P*** | **lci** | **uci** |
| ***lpar2a*** | all 11 larvae with clumped macrophages were wildtype for mutations in *lpar2a* ; 0.25 mutant larvae were expected. | | | | | | | | | | | | | | |
| ***lpar2b*** | 28.540 | 53.945 | 7.62E-02 | 0.702 | 1159.755 | 29.048 | 56.199 | 8.16E-02 | 0.655 | 1287.961 | 82.944 | 202.163 | 6.99E-02 | 0.698 | 9850.414 |
| ***gatad2ab*** | all 11 larvae with clumped macrophages were wildtype for mutations in *gatad2ab* ; 0.76 mutant larvae were expected. | | | | | | | | | | | | | | |
| **age (11dpf vs. 10dpf)** | 0.380 | 0.404 | 3.63E-01 | 0.047 | 3.054 | 0.368 | 0.395 | 3.51E-01 | 0.045 | 3.014 | 0.333 | 0.368 | 3.19E-01 | 0.038 | 2.904 |
| **time of day (in hours since 9AM)** | 0.615 | 0.117 | 1.07E-02 | 0.424 | 0.893 | 0.628 | 0.120 | 1.49E-02 | 0.432 | 0.913 | 0.557 | 0.136 | 1.67E-02 | 0.345 | 0.899 |
| **body length (in SD)** | - | - | - | - | - | 0.871 | 0.304 | 6.93E-01 | 0.439 | 1.727 | 0.605 | 0.251 | 2.26E-01 | 0.268 | 1.364 |
| **dorsal body surface area (in SD)** | - | - | - | - | - | 1.251 | 0.414 | 4.99E-01 | 0.654 | 2.394 | 1.297 | 0.480 | 4.83E-01 | 0.628 | 2.678 |
| **LDLc levels (in SD)** | - | - | - | - | - | - | - | - | - | - | 1.771 | 0.732 | 1.66E-01 | 0.788 | 3.981 |
| **HDLc levels (in SD)** | - | - | - | - | - | - | - | - | - | - | 0.618 | 0.221 | 1.79E-01 | 0.306 | 1.247 |
| **triglyceride levels (in SD)** | - | - | - | - | - | - | - | - | - | - | 0.681 | 0.281 | 3.53E-01 | 0.303 | 1.530 |
| **glucose levels** | - | - | - | - | - | - | - | - | - | - | 1.245 | 0.449 | 5.44E-01 | 0.614 | 2.524 |
| **intercept** | 0.171 | 0.111 | 6.52E-03 | 0.048 | 0.611 | 0.171 | 0.112 | 7.11E-03 | 0.047 | 0.619 | 0.170 | 0.134 | 2.40E-02 | 0.037 | 0.792 |

| | Macrophages outside the region of interest | | | | | | | | | | | | | | |
|---|---|---|---|---|---|---|---|---|---|---|---|---|---|---|---|
| | Model 1 (n=406) | | | | | Model 2 (n=376) | | | | | Model 3 (n=354) | | | | |
| | **Effect** | **SE** | ***P*** | **lci** | **uci** | **Effect** | **SE** | ***P*** | **lci** | **uci** | **Effect** | **SE** | ***P*** | **lci** | **uci** |
| ***lpar2a*** | all 15 larvae with many macropahages outside the region of interest were wildtype for mutations in *lpar2a* ; 0.33 mutant larvae were expected. | | | | | | | | | | | | | | |
| ***lpar2b*** | all 15 larvae with many macropahages outside the region of interest were wildtype for mutations in *lpar2b* ; 0.17 mutant larvae were expected. | | | | | | | | | | | | | | |
| ***gatad2ab*** | all 15 larvae with many macropahages outside the region of interest were wildtype for mutations in *gatad2ab* ; 1.02 mutant larvae were expected. | | | | | | | | | | | | | | |
| **age (11dpf vs. 10dpf)** | 1.361 | 0.819 | 6.08E-01 | 0.418 | 4.429 | 1.031 | 0.636 | 9.61E-01 | 0.308 | 3.453 | 1.141 | 0.731 | 8.37E-01 | 0.325 | 4.003 |
| **time of day (in hours since 9AM)** | 0.742 | 0.107 | 3.76E-02 | 0.560 | 0.983 | 0.848 | 0.130 | 2.82E-01 | 0.628 | 1.145 | 0.818 | 0.133 | 2.18E-01 | 0.594 | 1.126 |
| **body length (in SD)** | - | - | - | - | - | 0.310 | 0.121 | 2.80E-03 | 0.144 | 0.668 | 0.245 | 0.103 | 7.82E-04 | 0.108 | 0.557 |
| **dorsal body surface area (in SD)** | - | - | - | - | - | 0.573 | 0.208 | 1.25E-01 | 0.282 | 1.167 | 0.541 | 0.198 | 9.35E-02 | 0.264 | 1.109 |
| **LDLc levels (in SD)** | - | - | - | - | - | - | - | - | - | - | 2.382 | 0.801 | 9.83E-03 | 1.233 | 4.604 |
| **HDLc levels (in SD)** | - | - | - | - | - | - | - | - | - | - | 1.436 | 0.512 | 3.11E-01 | 0.713 | 2.890 |
| **triglyceride levels (in SD)** | - | - | - | - | - | - | - | - | - | - | 0.965 | 0.359 | 9.23E-01 | 0.465 | 1.999 |
| **glucose levels** | - | - | - | - | - | - | - | - | - | - | 1.103 | 0.355 | 7.60E-01 | 0.587 | 2.071 |
| **intercept** | 0.116 | 0.068 | 2.27E-04 | 0.037 | 0.364 | 0.045 | 0.034 | 3.76E-05 | 0.010 | 0.196 | 0.037 | 0.030 | 6.52E-05 | 0.007 | 0.186 |

***continued*** **Supplementary Table 38**

| | Clumped neutrophils | | | | | | | | | | | | | | |
|---|---|---|---|---|---|---|---|---|---|---|---|---|---|---|---|
| | Model 1 (n=456) | | | | | Model 2 (n=427) | | | | | Model 3 (n=402) | | | | |
| | **Effect** | **SE** | ***P*** | **lci** | **uci** | **Effect** | **SE** | ***P*** | **lci** | **uci** | **Effect** | **SE** | ***P*** | **lci** | **uci** |
| ***lpar2a*** | 3.869 | 3.250 | 1.07E-01 | 0.746 | 20.074 | 3.695 | 3.109 | 1.20E-01 | 0.710 | 19.221 | 3.609 | 3.246 | 1.54E-01 | 0.619 | 21.040 |
| ***lpar2b*** | all 34 larvae with clumped neutrophils were wildtype for mutations in *lpar2b* ; 0.44 mutant larvae were expected. | | | | | | | | | | | | | | |
| ***gatad2ab*** | 0.892 | 1.010 | 9.20E-01 | 0.097 | 8.215 | 0.835 | 0.955 | 8.75E-01 | 0.089 | 7.863 | 1.235 | 1.439 | 8.56E-01 | 0.126 | 12.110 |
| **age (11dpf vs. 10dpf)** | 0.891 | 0.404 | 8.00E-01 | 0.367 | 2.166 | 0.790 | 0.367 | 6.12E-01 | 0.318 | 1.965 | 0.767 | 0.371 | 5.83E-01 | 0.298 | 1.978 |
| **time of day (in hours since 9AM)** | 0.980 | 0.086 | 8.17E-01 | 0.825 | 1.164 | 0.997 | 0.092 | 9.75E-01 | 0.832 | 1.194 | 0.961 | 0.103 | 7.10E-01 | 0.779 | 1.185 |
| **body length (in SD)** | - | - | - | - | - | 0.859 | 0.166 | 4.33E-01 | 0.588 | 1.255 | 1.023 | 0.232 | 9.20E-01 | 0.657 | 1.594 |
| **dorsal body surface area (in SD)** | - | - | - | - | - | 1.073 | 0.207 | 7.16E-01 | 0.735 | 1.566 | 0.916 | 0.197 | 6.85E-01 | 0.601 | 1.397 |
| **LDLc levels (in SD)** | - | - | - | - | - | - | - | - | - | - | 0.876 | 0.182 | 5.25E-01 | 0.584 | 1.316 |
| **HDLc levels (in SD)** | - | - | - | - | - | - | - | - | - | - | 0.959 | 0.184 | 8.28E-01 | 0.659 | 1.396 |
| **triglyceride levels (in SD)** | - | - | - | - | - | - | - | - | - | - | 0.838 | 0.186 | 4.28E-01 | 0.542 | 1.296 |
| **glucose levels** | - | - | - | - | - | - | - | - | - | - | 1.151 | 0.223 | 4.69E-01 | 0.787 | 1.684 |
| **intercept** | 0.087 | 0.038 | 2.33E-08 | 0.037 | 0.204 | 0.089 | 0.040 | 5.86E-08 | 0.037 | 0.213 | 0.091 | 0.044 | 6.33E-07 | 0.036 | 0.234 |

| | Partly imaged larvae (n=386) | | | | |
|---|---|---|---|---|---|
| | **Effect** | **SE** | ***P*** | **lci** | **uci** |
| ***lpar2a*** | all 29 larvae that were only partly imaged were wildtype for mutations in *lpar2a* ; 0.58 mutant larvae were expected. | | | | |
| ***lpar2b*** | all 29 larvae that were only partly imaged were wildtype for mutations in *lpar2b* ; 0.35 mutant larvae were expected. | | | | |
| ***gatad2ab*** | 0.889 | 1.012 | 9.18E-01 | 0.096 | 8.270 |
| **time of day (in hours since 9AM)** | 1.170 | 0.112 | 1.02E-01 | 0.969 | 1.411 |
| **intercept** | 0.039 | 0.020 | 5.32E-10 | 0.014 | 0.108 |

Associations are shown for criteria that resulted in the exclusion of at least 10 larvae. Many false positives: >20% of true negative objects were falsely detected by the quantification pipeline. Many false negatives: >20% of true positives objects were falsely excluded by the quantification pipeline.Associations were examined using logistic regression models. Model 1: adjusted for time of day; Model 2: additionally adjusted for body length and dorsal surface area; Model 3: additionally adjusted for whole-body triglyceride and glucose levels. Dorsal body surface area was normalized for body length using residuals; whole-body LDL cholesterol, HDL cholesterol, triglyceride and glucose levels were normalized for protein level using residuals. Effects shown for *lpar2a*, *lpar2b* and *gatad2ab* are for each additional mutated allele. Adjusting for batch would have resulted in the exclusion of larvae. Lci and uci are lower and upper boundaries of the 95% confidence interval.
